# WWP2 osteoarthritis risk allele rs1052429-A confers risk by affecting cartilage matrix deposition via hypoxia associated genes

**DOI:** 10.1101/2022.03.31.486523

**Authors:** Margo Tuerlings, George M.C. Janssen, Ilja Boone, Marcella van Hoolwerff, Alejandro Rodriguez Ruiz, Evelyn Houtman, Eka H.E.D. Suchiman, Robert J.P. van der Wal, Rob G.H.H. Nelissen, Rodrigo Coutinho de Almeida, Peter A. van Veelen, Yolande F.M. Ramos, Ingrid Meulenbelt

## Abstract

**Objective:** To explore the co-expression network of the osteoarthritis (OA) risk gene *WWP2* in articular cartilage and study cartilage characteristics when mimicking the effect of OA risk allele rs1052429-A on *WWP2* expression in a human 3D *in vitro* model of cartilage.

**Methods:** Co-expression behavior of *WWP2* with genes expressed in lesioned OA articular cartilage (N=35 samples) was explored. By applying lentiviral particle mediated *WWP2* upregulation in 3D *in vitro* pellet cultures of human primary chondrocytes (N=8 donors) the effects of upregulation on cartilage matrix deposition was evaluated. Finally, we transfected primary chondrocytes with miR-140 mimics to evaluate whether miR-140 and WWP2 are involved in similar pathways.

**Results:** Upon performing Spearman correlations in lesioned OA cartilage, 98 highly correlating genes (|ρ|>0.7) were identified. Among these genes, we identified *GJA1, GDF10, STC2, WDR1*, and *WNK4*. Subsequent upregulation of WWP2 on 3D chondrocyte pellet cultures resulted in a decreased expression of *COL2A1* and *ACAN* and an increase in *EPAS1* expression. Additionally, we observed a decreased expression of *GDF10, STC2*, and *GJA1*. Proteomics analysis identified 42 proteins being differentially expressed with WWP2 upregulation, which were enriched for ubiquitin conjugating enzyme activity. Finally, upregulation of miR-140 in 2D chondrocytes resulted in significant upregulation of *WWP2* and *WDR1*.

**Conclusions:** Mimicking the effect of OA risk allele rs1052429-A on *WWP2* expression initiates detrimental processes in the cartilage shown by a response in hypoxia associated genes *EPAS1,GDF10*, and *GJA1* and a decrease in anabolic markers, *COL2A1* and *ACAN*.

## INTRODUCTION

Globally, osteoarthritis (OA) is a highly prevalent and disabling joint disease which confers high social and economic burden to society. Risk factors for OA include sex, abnormal joint loading, obesity, metabolic diseases, and genetic factors [1]. To discover genes and underlying disease pathways, large genome wide association meta-analyses have been performed and multiple robust single nucleotide polymorphisms (SNPs) were identified significantly conferring risk to initiation and progression of OA [2-4]. Similar to other complex traits, these risk alleles have subsequently been found to affect expression of positional genes *in cis* in disease relevant tissues, also known as allelic imbalance (AI)[5, 6]. Founded by this assumption, we previously used RNA sequencing data of OA articular cartilage to report on genome-wide AI expression of SNPs in cartilage specific genes, as such, providing an AI expression database to *in silico* check functional aspects of identified and/or future OA risk SNPs [7]. One of the top findings was SNP rs1052429 located in the 3’UTR of the *WWP2* gene showing highly significant AI, with risk allele rs1052429-A marking higher expression of *WWP2* relative to rs1052429-G. Among the OA risk SNPs identified in a large genome-wide meta-analysis of Icelandic and UK knee OA patients was rs34195470, located in *WWP2* gene and a proxy of our AI SNP rs1052429 (r^2^=0.6) [2]. Recently, rs34195470 was confirmed being OA risk SNP in the largest genome-wide meta-analysis so far, including individuals from 9 populations [4]. Together these data demonstrate that *WWP2* with risk alleles rs34195470-G and rs1052429-A confer robust risk to human OA and are associated to higher expression of *WWP2*. We also previously identified transcription of *WWP2* in cartilage being epigenetically regulated [8], as well as being responsive in the OA pathophysiological process [9]. Moreover, *WWP2* was previously shown to be a marker for hypertrophic chondrocytes in OA knee joints [10].

WWP2 is a member of the Nedd4 superfamily, a small group within the E3 ubiquitin ligase enzymes and is involved in post-translational modifications. The WWP2 protein contains four double tryptophan (WW) domains, which allow specific protein-protein interactions and it is expressed in multiple organs throughout the body [11]. More specifically to cartilage, Nakamura et al. [12] showed that WWP2 interacts with SOX9 to form a complex that facilitates nuclear translocation of SOX9, as such enabling SOX9 transcriptional activity. Despite the association between the risk allele and higher expression levels of *WWP2* in human cartilage, the effect of *WWP2* knockout (KO) in mice with age-related and surgically induced models of OA showed that lack of *WWP2* expression resulted in increased expression of catabolic cartilage markers *RUNX2* and *ADAMTS5* [13]. In a different context, *WWP2* was found to be a host gene for microRNA-140 (miR-140), a miRNA highly expressed in cartilage and shown to be differentially expressed between preserved and lesioned OA cartilage [9]. As such, it was suggested that expression of miR-140 and the C-terminal transcript of *WWP2* (*WWP2-C*, also called *WWP2* isoform 2) are co-regulated [14, 15].

In the current study, we set out to explore a *WWP2* co-expression network in our previous whole-transcriptome OA cartilage dataset [9]. Moreover, to study the effect of the genetic risk allele, we functionally assessed the effect of lentiviral-mediated upregulation of *WWP2* in a 3D *in vitro* model using primary human chondrocytes. Apart from conventional anabolic and catabolic cartilage markers, genes identified in the *WWP2* co-expression network were used as a read-out to evaluate the effect of *WWP2* upregulation. Since WWP2 is involved in post-translational modifications, we explored the effect of *WWP2* upregulation on protein level by performing proteomic analysis. Finally, we explored the effects of upregulation of miR-140 in primary chondrocytes by transfection with miR-140 mimics.

## METHODS

### Sample description

RNA-sequencing data was included of macroscopically lesioned OA cartilage (N=35 samples, RAAK-study). Classification of macroscopically preserved and lesioned OA cartilage was done as described previously [16]. Chondrocytes isolated from the preserved areas of N=7 participants of the RAAK-study were included to perform lentiviral transduction. Finally, chondrocytes of N=7 participants of the RAAK-study were included to perform miR-140 transfection. For all sample characteristics see **Table S1**. The RAAK-study is approved by the medical ethics committee of the Leiden University Medical Center (P08.239/P19.013).

### RNA-sequencing

RNA was isolated from the articular cartilage using Qiagen RNeasy Mini Kit (Qiagen, GmbH, Hilden, Germany). Paired-end 2×100 bp RNA-sequencing (Illumina TruSeq RNA Library Prep Kit, Illumina HiSeq2000 and Illumina HiSeq4000) was performed. Strand specific RNA-seq libraries were generated which yielded a mean of 20 million reads per sample. Data from both Illumina platforms were integrated and analyzed with the same in-house pipeline. RNA-seq reads were aligned using GSNAP [17] against GRCh38 using default parameters. Read abundances per sample was estimated using HTSeq count v0.11.1 [18]. Only uniquely mapping reads were used for estimating expression. The quality of the raw reads for RNA-sequencing was checked using MultiQC v1.7. [19]. To identify outliers, principal component analysis (PCA) was applied. The DESeq2 package [20] was used to normalize the RNA-seq data, as a variance-stabilizing transformation was performed.

### Creating a co-expression network

We explored co-expression behavior of WWP2 with progression of OA by correlating (Spearman correlation) *WWP2* expression levels in our RNA sequencing dataset with expression levels of all genes expressed in OA articular cartilage (N=20048 genes)[9]. To correct for multiple testing, the Benjamini-Hochberg method was used, as indicated by the false discovery rate (FDR) with a significance cutoff value of 0.05. To include the most informative genes a threshold of |ρ|>0.7 was selected, corresponding to approximately the top 1% of the total significant correlations.

### Lentiviral transduction

The full length WWP2 plasmid (NM_001270454.1) was ordered in pcDNA3.1 with XhoI/XbaI cloning sites (Genscript Biotech). The plasmid was digested and inserted into the XhoI/XbaI sites of the pLV-CMV-IRES-eGFP lentiviral backbone (kindly provided by Prof. Dr. Hoeben, Dept. of Molecular Cell Biology, Leiden University Medical Center). Lentiviral production was performed in HEK 293T cells, using Lenti-vpak Lentiviral Packaging Kit (Origene Technologies, Inc.). The HEK 293T cells were expanded in DMEM (high glucose; Gibco) supplemented with 10% fetal calf serum (FBS, Gibco) and 100 U/ml penicillin and 100 ug/ml streptomycin (Gibco). The pLV-CMV-IRES-eGFP lentiviral backbone without the WWP2 insert was used as a control. Primary chondrocytes were isolated from the articular cartilage of human joints and expanded in DMEM (high glucose; Gibco) supplemented with 10% FBS (Gibco), 100 U/ml penicillin and 100 ug/ml streptomycin (Gibco), and 0.5 ng/ml FGF-2 (PeproTech), as described previously [21]. After expansion, the chondrocytes were seeded in a density of 3.5×105 cells per 10 cm culture dish (passage 2) and left overnight. Then, the lentivirus was added in a MOI of 1 in addition of 15 ug/ml Polybrene (Sigma-Aldrich). After incubation of approximately 16 hours, the lentiviral solution was replaced by normal culture medium. The chondrocytes were passaged and expanded afterwards.

### *In vitro* 3D pellet cultures

3D pellet cultures were formed by adding 2.5×105 cells in their expansion medium to a 15 ml Falcon tube and subsequently expose them to centrifugal forces (1200 rpm, 4 minutes). After 24 hours, the expansion medium was replaced by chondrogenic differentiation medium (DMEM (high glucose; Gibco), supplemented with Ascorbic acid (50 μg/ml; Sigma-Aldrich), L-Proline (40 μg/ml; Sigma-Aldrich), Sodium Puryvate (100 μg/ml; Sigma-Aldrich), Dexamethasone (0.1 μM; Sigma-Aldrich), ITS+, 100 U/ml penicillin and 100 ug/ml streptomycin (Gibco) and TGF-β1 (10 ng/ml; PeproTech)), as described previously [21]. Medium was refreshed every 3-4 days and the caps of the Falcon tubes were open for the first 7 days to allow oxygen entering the tubes. The pellets were harvested at different timepoints; after 24 hours (day 0) and after 7 days. The harvested materials were lysed using RNABee (Bio-connect) and stored at -80°C until further processing.

### RT-qPCR

RNA was isolated from the samples using the RNeasy Mini Kit (Qiagen). cDNA synthesis was performed using the First Strand cDNA Synthesis Kit (Roche Applied Science). Subsequently, RT-qPCR was performed with the Biomark™ 96.96 Dynamic Arrays (Fluidigm) according to the manufacturer’s protocol. Additional RT-qPCR was performed using SYBR Green without the ROX reference dye (Roche Applied Science) and the QuantStudio 6 Real-Time PCR system (Applied Biosystems). In both methods, GAPDH and SDHA were used as housekeeping genes. The measured gene expression levels were corrected for the housekeeping genes GAPDH and SDHA, and the fold changes were calculated using the 2^-ΔΔCT^ method. All values were calculated relative to the control groups. The paired sample t-test was used to calculate significance (P<0.05).

### Quantitative Proteomics Using TMT Labeling

Lysis, digestion, TMT labeling and mass spectrometry analysis was essentially performed as described [22]. In short, 3D pellet cultures were washed with PBS, extracted with 100 ul 5% SDS, 100 mM Tris HCl pH 7.6 each, sonicated twice for 10 min and incubated for 20 min at 95°C. Insoluble material was removed by centrifugation for 2 minutes at 15,000 rpm in an Eppendorf centrifuge. Proteins were reduced, alkylated, subjected to chloroform methanol precipitation and digested with trypsin as described earlier [22]. Peptide concentration was determined by BCA Gold protein assay (Pierce) and 10ug of the peptide was labeled using TMT10plex reagent (Thermo). Three separate TMT10 sets were prepared with a common reference sample consisting of a mixture of all peptide samples. TMT-labeled peptides were dissolved in 0.1% formic acid and subsequently analyzed by online C18 nano-HPLC MS/MS with a system consisting of an Easy nLC 1200 gradient HPLC system (Thermo, Bremen, Germany), and an Orbitrap Fusion LUMOS mass spectrometer in synchronous precursor selection (Thermo). For peptide identification, MS/MS spectra were searched against the human database (20596 entries) using Mascot Version 2.2.07 (Matrix Science) with the following settings: 10 ppm and 0.6 Da deviation for precursor and fragment masses, respectively. Trypsin was set as enzyme and two missed cleavages were allowed. Carbamidomethyl on cysteines and TMT6plex on Lys and N-term were set as fixed modifications. Variable modifications were oxidation (on Met and Pro) and acetylation on the protein N-terminus. All searches and subsequent data analysis, including Percolator and abundance ratio calculation, were performed using Proteome Discoverer 2.4 (Thermo Scientific). Peptide-spectrum matches were adjusted to a 1% FDR. Proteins were filtered on a minimal unique peptide count of 2. Since the sample size was rather small and we did observe significant increased expression of cartilage markers already at day three of pellet culture, we pooled the data of day 3 and day 7 for further analysis.

### Histochemistry

The 3D chondrocyte pellet cultures were fixed with 4% formaldehyde and subsequently embedded in paraffin. The sections were stained for glycosaminoglycan (GAG) deposition using the Alcian Blue staining. The staining was quantified by loading the images in Fiji and splitting the color channels. Subsequently, the grey values were measured and corrected for the grey value of the background and for the number of cells present. The paired sample t-test was again used to calculate significance.

### Transfection with miR-140 mimics

Primary chondrocytes were passaged in a concentration of 4.0×104 cells per well. After 24 hours, the cells were transfected with hsa-miR-3p mimic (Invitrogen) or a control mimic at 5 nM final concentration using Opti-MEM (Gibco) and Lipofectamine RNAiMax Transfection reagent (Invitrogen) according to manufacturer’s protocol. Approximately 16 hours after transfection, the cells were lysed using TRizol (Thermo Fisher Scientific) for RNA isolation and stored at -80°C until further processing. An overview of the applied strategy can be seen in **Fig. 1**. The RNA sequencing data of the articular cartilage is deposited at ArrayExpress (E-MTAB-7313). Further data generated and used in this study is not openly available due to reasons of sensitivity and are available from the corresponding author upon reasonable request.

**Fig. 1.**
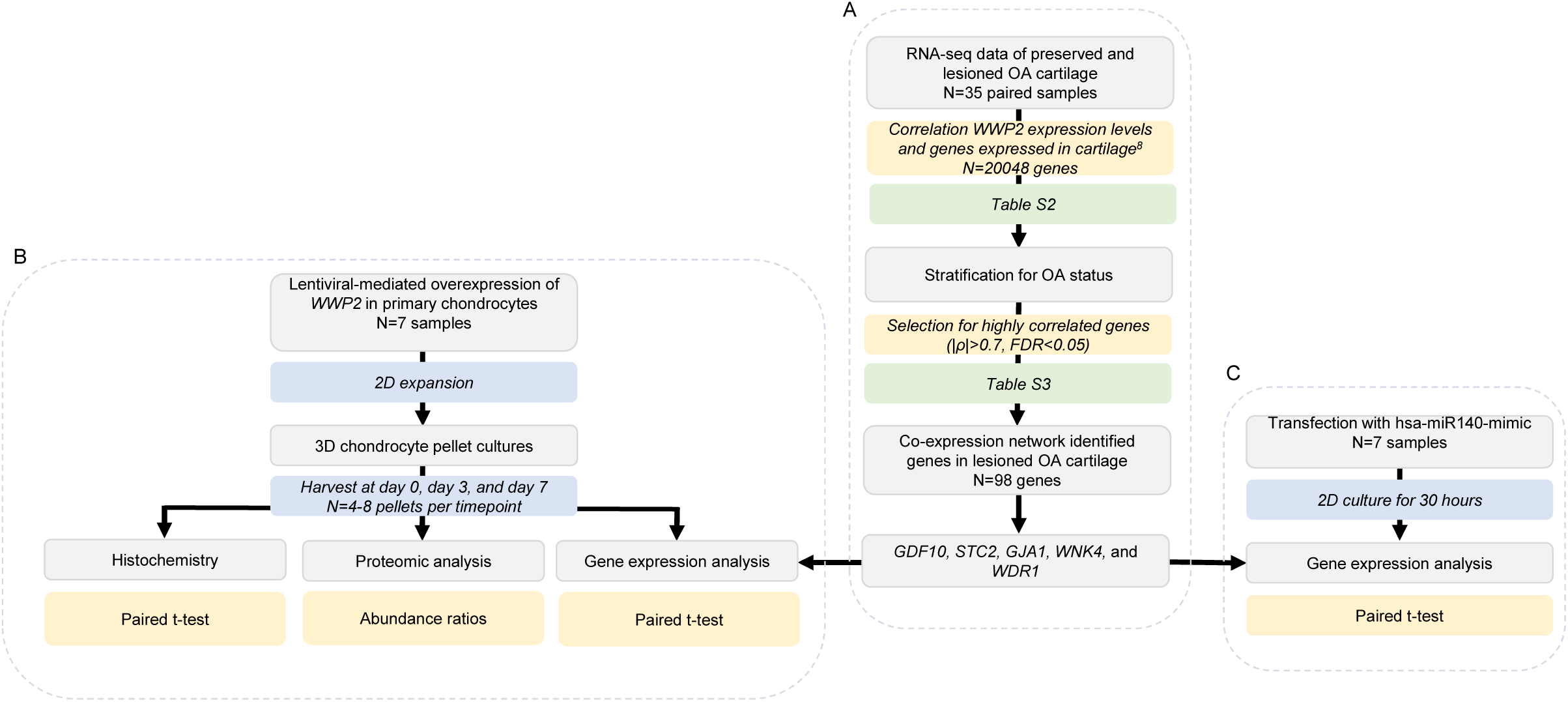
Schematic overview of applied strategy

## RESULTS

### Co-expression network of *WWP2*

To identify genes that are regulated by, or co-expressed with, *WWP2* in OA cartilage, we used RNA sequencing data of lesioned OA cartilage (N=35 samples, **Table S1A**) to perform Spearman correlation between expression levels of *WWP2* and genes expressed in cartilage (N=20048 genes, **Table S2, Fig. 1A**). We identified 98 genes highly correlating (|ρ|>0.7) to *WWP2*. These 98 genes were significantly enriched for, amongst others, GO-terms Extracellular exosome (GO:0070062, 36 genes), characterized by expression of *GJA1* (encoding gap junction alpha 1), *SMO* (encoding smoothened frizzled class receptor), and *WDR1* (encoding WD repeat domain 1), and Myelin sheath (GO:0043209, 10 genes), characterized by expression of *WDR1, RALA* (encoding RAS like proto-oncogene A), and *CCT5* (encoding chaperonin containing TCP1 subunit 5) (**Table S3**). As shown in **Fig. S1**, genes highly correlating to *WWP2* (N=98) formed a highly interconnected network, i.e. genes that are all highly correlating with each other. In this network we identified direct and indirect relations with *WWP2*, including *GJA1* (ρ=-0.81, 70 connections, i.e. highly correlating to 70 genes in the network), *WNK4* (encoding WNK lysine deficient protein kinase 4, ρ=0.81, 37 connections), *ACAN* (encoding aggrecan, ρ=0.78, 16 connections), and *STC2* (encoding stanniocalcin 2, ρ=0.77, 17 connections).

### Lentiviral particle-mediated upregulation of *WWP2*

Although lower expression of *WWP2* was observed in lesioned compared to preserved cartilage in our previous study [9], genetic evidence suggests that higher expression of *WWP2* predisposes to development of OA, indicating that downregulation in OA pathophysiology is merely a beneficial attempt of chondrocytes to reverse the OA state [7]. Therefore, effect of upregulation of *WWP2* was studied on cartilaginous matrix deposition in *in vitro* 3D chondrocyte pellet cultures, by creating a lentiviral particle mediated upregulation of *WWP2*. 3D pellet cultures were harvested after three or seven days of culturing and gene (N=16 pellet cultures of N=8 donors) and protein (N=16 pellet cultures of N=4 donors) expression levels were measured (**Table S1B**). First, we confirmed whether *WWP2* upregulation was successful by measuring both gene and protein expression levels at day zero of the 3D chondrocyte pellet culture, and we observed a significant increase in *WWP2* gene expression levels (P=1.0×10^−5^, **Fig. S2A-B**), which was confirmed on protein level (P=3.2×10^−6^, **Fig. S2C**).

### Effect of *WWP2* upregulation on cartilage matrix deposition

Next, we evaluated effect of *WWP2* upregulation on expression levels of conventional cartilage genes during 3D pellet culture of seven days (**Fig. 1B**). As shown in **Fig. 2A**, we found significant reduced gene expression of *ACAN* (FC=0.80, *P*=0.04) and *COL2A1* (encoding collagen type 2 alpha chain 1, FC=0.77, *P*=0.01), in *WWP2* upregulated pellets compared to their controls at day seven. Moreover, we showed significant increased gene expression of degeneration markers *EPAS1* (encoding endothelial PAS domain protein 1, FC=1.56, *P*=0.004) (**Fig. 2B**). Notably, *SOX9, ADAMTS5,* and *RUNX2*, which were previously linked to WWP2 function, were not consistently changed upon WWP2 upregulation. Moreover, we stained 3D pellet cultures for presence of glycosaminoglycans (GAGs) using Alcian Blue staining, and observed a trend towards decreased Alcian Blue intensity when comparing *WWP2* upregulated pellets with controls (**Fig. 2C**).

**Fig. 2.**
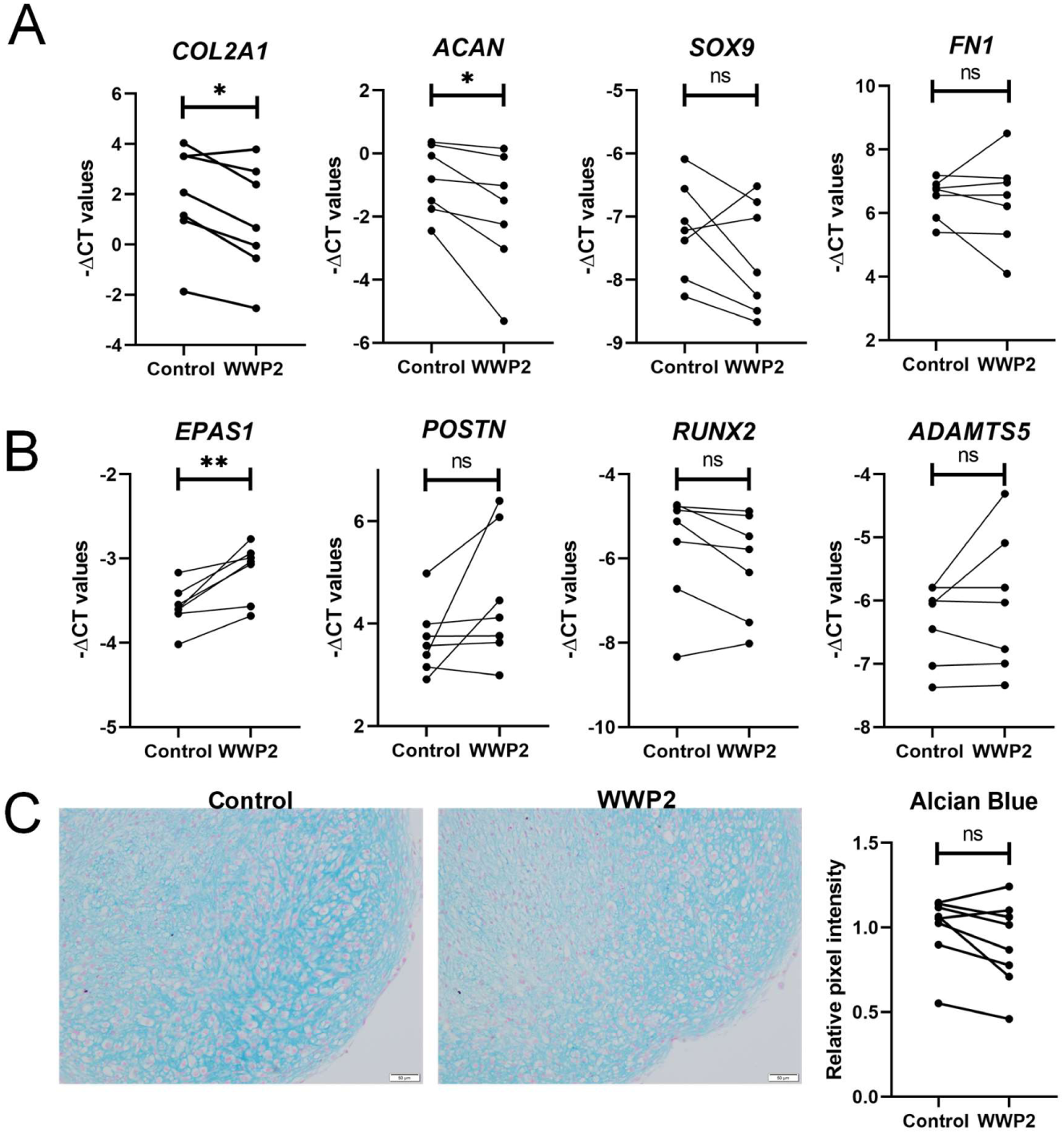
mRNA expression levels of cartilage matrix markers (A) and cartilage degeneration markers (B) for pellets with WWP2 upregulation and their controls at day 7 (N=11-12 pellet cultures, N=7 donors). (C) Alcian Blue staining visualizing GAGs deposition in WWP2 overexpressed pellets and their controls after 7 days of culturing (N= 26 pellet cultures, N=8 donors). Scale bar indicates 50 um. Ns: not significant, *P<0.05, ** P<0.005 upon performing a Paired sample t-test.

### Effect of *WWP2* upregulation on genes correlated to *WWP2*

To investigate functional relationships between *WWP2* and identified correlating and highly interconnected genes, we selected *GDF10* (encoding growth differentiation factor 10, ρ=0.72, 18 connections), *STC2* (ρ=0.77, 17 connections), *GJA1* (ρ=-0.81, 70 connections), *WDR1* (encoding WD repeat-containing protein 1, ρ=-0.70, 5 connections), and *WNK4 (*ρ=0.81, 37 connections) from the network (**Fig. 1**) to use as read-out of WWP2 upregulation in 3D chondrocyte pellet cultures (**Fig. 1B**). As shown in **Fig. 3**, we observed significant decreased gene expression of *GDF10* (FC=0.62, *P*=0.002) and *STC2* (FC=0.73, *P*=0.04) with upregulation of *WWP2*. Albeit not significant, gene expression of *GJA1* (FC=0.76, *P*=0.08) was also consistently lower in *WWP2* upregulated pellets. Together, these data suggest that *GDF10, STC2*, and *GJA1* are downstream of *WWP2* either by direct or indirect activity. In contrast, *WNK4* and *WDR1* did not show consistent changes in expression with upregulation of *WWP2*, suggesting *WNK4* and *WDR1* are rather upstream in the pathway of *WWP2*.

**Fig. 3.**
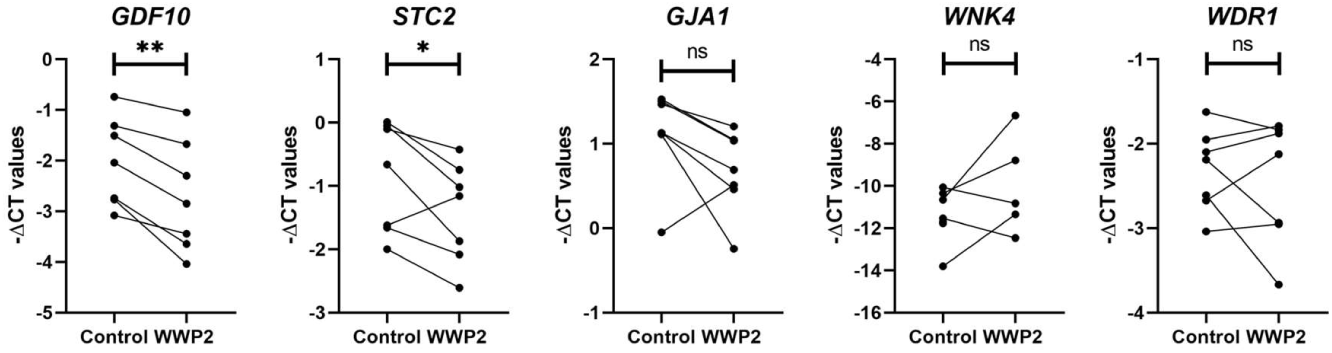
mRNA expression levels of genes correlating with *WWP2* in WWP2 upregulated 3D chondrocyte pellet cultures compared to their controls after 7 days of culturing (N=6-12 pellet cultures, N=7 donors). Ns: not significant, * P<0.05, ** P<0.05 upon performing a Paired sample t-test.

### Proteomics

To study the extent to which gene expression levels translate to protein levels, we performed proteomics analysis. Prior to differential expression analysis of pellet cultures with and without WWP2 upregulation, we explored protein expression levels of cartilage markers in our control pellet cultures at day three and day seven of 3D pellet culture. Upon comparing day three and day seven with day zero of control pellets, we observed increased protein expression of cartilage markers COL2A1 (FC=2.68 and FC=32.94, respectively), ACAN (FC=5.28 and FC=13.75, respectively), COMP (cartilage oligomeric matrix protein, FC=5.31 and FC=20.25, respectively), and FN1 (fibronectin, FC=2.12 and FC=3.51, respectively) (**Table S4, Fig. S3**). Moreover, mesenchymal markers CD44 (FC=0.83 and FC=0.46, respectively) and CD166 (FC=0.79 and FC=0.62, respectively) and IGFBP3 (insulin growth factor binding protein 3, FC=0.10 and FC=0.10, respectively) were downregulated on both days.

Together, this indicates that cartilage-like matrix is produced by chondrocytes already at day three, but is increasing towards day seven. Notably, SOX9 was not detected in the proteomics analysis.

Next, we evaluated the effect of WWP2 upregulation on cartilage matrix deposition on protein level (**Fig. 1B**). Since we observed increased protein expression of cartilage markers in control pellet cultures already on day three (**Fig. S3A**), we pooled day three and day seven for further analysis to increase power. Upon comparing pellet cultures with and without WWP2 upregulation, we found WWP2 still being significantly upregulated after three and seven days of culturing. Furthermore, we found 42 proteins significantly differentially expressed (**Fig. 4A, Table S5**), of which GJA1 (FC=0.54) was most significantly downregulated in WWP2 upregulated pellet cultures, confirming the downregulation observed on gene expression level (FC=0.76). Oppositely, the observed changes in *ACAN, COL2A1, FN1*, and *POSTN* gene expression levels were not confirmed on protein level. Proteins encoded by *SOX9, EPAS1, RUNX2*, and *ADAMTS5* were either not identified or did not show unique peptides. Upon performing enrichment analysis on the 42 differentially expressed proteins, we found significant enrichment for ubiquitin conjugating enzyme activity (5 proteins, FDR=0.002) and ubiquitin-protein transferase activity (6 proteins, FDR=0.03), both terms characterized by expression of, amongst others, UBE2D4, UBE2L3, and UBE2D1. Furthermore, these 42 proteins showed significant protein-protein interactions (P=0.02, **Fig. 4B**), also representing ubiquitin conjugating enzyme activity.

**Fig. 4.**
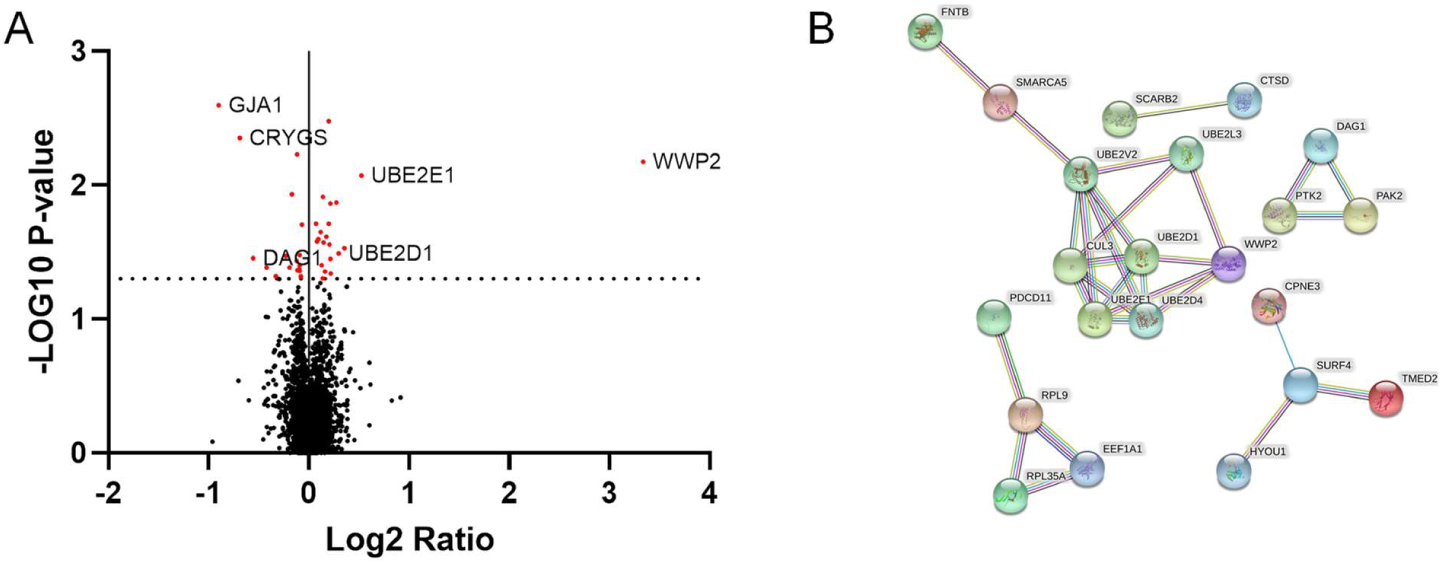
(A) Volcano plot of proteins differentially expressed between *WWP2* transduced 3D pellet cultures and their controls at day three (N=8 pellets, N=4 donors) and day seven (N=8 pellets, N=4 donors) together (N=16 pellets, N=4 donors). The red dots indicate the significantly differentially expressed proteins. (B) Protein-protein interaction network in STRING.

### miR-140-3p and WWP2

Since it has been suggested that *WWP2* and miR-140 are co-expressed [14, 15], we transfected primary chondrocytes with miR-140-3p mimics, to assess whether this miRNA regulates similar genes as involved in the *WWP2* co-expression network (N=7, **Table S1B, Fig. 1C**). As shown in **Fig. 5A**, we observed significant increased expression levels of *WWP2* (FC=1.22, *P*=0.02) with upregulation of miR-140-3p. Moreover, we observed consistent increased expression of *WWP2* isoform 6 (FC=1.29, *P*=0.06), while significant decreased expression of isoform 4 (FC=0.63, *P*=0.02). Notably, we did not see effect on expression levels of *WWP2* isoform 2, also called *WWP2-C*. With respect to highly correlated genes, we observed increased expression of *WDR1* (FC=1.79, *P*=1.00×10^−3^), one of the genes that was not consistently changed with *WWP2* upregulation (**Fig. 5B**). Also contradictory to the effects of *WWP2* upregulation, we observed an increased expression of *STC2* (FC=1.55, *P*=0.08) and we did not observe consistent effects on *GJA1* expression level.

**Fig. 5.**
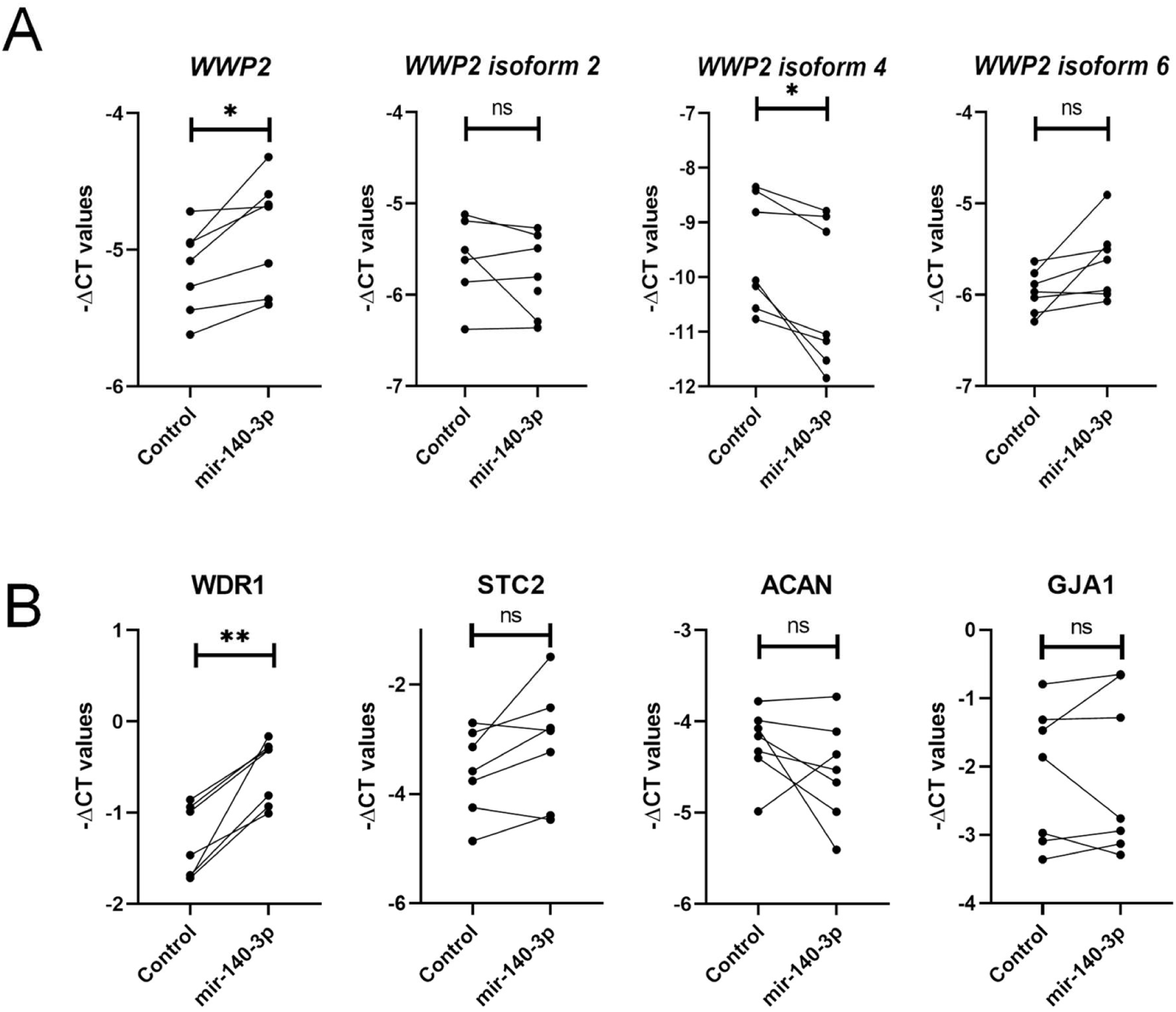
mRNA expression levels upon transfection with miR-140-3p. (A) expression levels of WWP2 and its isoforms. (B) Expression levels of genes correlated to *WWP2* (N=8 wells, N=4 donors). Ns: not significant, * p<0.05, ** p<0.005 upon performing a Paired sample t-test.

## DISCUSSION

By combining a genome-wide screen for cartilage specific allelic imbalance [7] and large scale GWAS [2, 4], we hypothesized that upregulated expression of *WWP2* confers robust risk to OA. Here, we set out to functionally investigate the role of *WWP2* in cartilage by exploring the *WWP2* co-expression network in a previously assessed RNA sequencing dataset [9]. Moreover, lentiviral-mediated upregulation of *WWP2* was shown to have detrimental effects on cartilage matrix deposition, as shown by downregulation of *COL2A1* and *ACAN* and upregulation of *EPAS1*. Apart from conventional anabolic and catabolic cartilage markers, genes identified in the *WWP2* co-expression network were used as read-out, showing *GDF10, STC2*, and *GJA1* being responsive to *WWP2* upregulation. Furthermore, to explore effects of miR-140-3p, that was suggested to be co-regulated with *WWP2*, we transfected primary chondrocytes with miR-140-3p mimics.

Based on AI expression of the OA risk SNP rs1052429, we hypothesized that *WWP2* confers risk to OA onset by upregulated expression, whereas *WWP2* exhibited FDR significantly lower expression in lesioned compared to preserved cartilage (FC=0.78, FDR=5.3×10^−3^), together suggesting that lower expression levels of *WWP2* in lesioned OA cartilage are rather an attempt of chondrocytes to reverse the OA state than a cause to the OA process [24, 25]. Concomitantly, co-expression network analyses showed 98 highly and significantly (|ρ|>0.7, FDR<0.05) correlating genes to *WWP2*, including *GJA1* (ρ=-0.81), *WNK4* (ρ=0.81), *ACAN* (ρ=0.78), and *STC2* (ρ=0.77) (**Table S2**). Previously, it was shown that WWP2 interacts with SOX9 and that it regulates SOX9 transcriptional activity [12]. Although *SOX9* is highly expressed in cartilage, *SOX9* was not among the high and significant correlations (ρ=0.5). On the other hand, *SOX9* was previously shown to regulate the expression of, amongst others *ACAN* [26, 27], which was here shown to be highly correlated to *WWP2* (ρ=0.78).

Upon studying the effect of upregulation of *WWP2*, we found *EPAS1* and *GDF10* being genes that had most consistent and significant changed levels of gene expression. *EPAS1*, encoding hypoxia-inducible factor 2 alpha, is known for its role in endochondral ossification and is a known cartilage degradation marker in OA [28, 29]. *GDF10*, also known as bone morphogenic protein 3, is involved in osteogenesis as it inhibits osteoblast differentiation via *SMAD2* and *SMAD3 [25, 30]*. The latter also being previously identified as OA susceptibility gene [2]. Furthermore, lower expression level of *GDF10* was associated with OA severity in both bone and cartilage [31]. Interestingly, *GDF10* was shown to be a hypoxia inducible gene, like *EPAS1*, which is regulated by SOX9 and was identified as marker for differentiated chondrocytes as it inhibits adipogenesis and osteogenesis [32]. Both *EPAS1* and *GDF10* were not in the proteomics analysis, either because they were not measured (EPAS1) or they did not show unique peptides (GDF10). Additionally, we observed a decrease in *STC2* and *GJA1* expression. Downregulation of *GJA1* did not reach statistical significance on gene expression level, while on protein level GJA1 was the most significantly downregulated protein. STC2 is a glycoprotein and upregulation of *STC2* in mice has been shown to delay endochondral ossification [33, 34]. Moreover, it was shown that *STC2* was higher expressed in healthy cartilage compared to osteophytic cartilage [35], suggesting its potential role in initiation and progression of OA in presence of higher *WWP2* expression. GJA1, also known as connexin 43, is a major protein of functional gap junctions which allows for cell-cell communication. More specific to cartilage, connexin 43 is essential in mechanotransduction [36]. Alterations in connexin 43 expression and localization affects this cell-cell communication, by which homeostasis to maintain cartilage tissue gets disturbed [37]. Notably, like EPAS1 and GDF10, the function of connexin 43 is regulated by oxygen levels [38]. Upregulation of *EPAS1* and downregulation of *GDF10, STC2* and *GJA1* suggests that increased level of WWP2 has detrimental effects on cartilage matrix deposition, which acts via hypoxia associated chondrocyte dedifferentiation. This is in line with decreased gene expression levels of *COL2A1* and *ACAN*, two major cartilage markers.

To evaluate the effects of WWP2 upregulation on cartilage matrix deposition on protein level, we performed proteomics analysis. We did confirm upregulation of WWP2 on day zero, which was still present at day three and seven (**Fig. 4, Fig. S2**,). Moreover, we found 42 significantly differentially expressed proteins upon comparing pellet cultures with and without WWP2 upregulation, of which GJA1 was most significantly differentially expressed and showing the highest fold change (FC=0.49). We were not able to confirm differences we observed in gene expression levels of *COL2A1,ACAN,FN1*, and *POSTN* on protein level (**Table S5**), which might be due to the relatively low sample size (N=4 donors) or due to suboptimal timepoint chosen to evaluate the effect of WWP2 on either gene or protein expression level. Since WWP2 is a E3 ubiquitin ligase and the differentially expressed proteins were significantly enriched for ubiquitin conjugating enzyme activity and ubiquitin-protein transferase activity, upregulation of WWP2 could also affect proteins cellular location, activity, and protein-protein interactions without changing expression levels itself, which is not captured by our read-outs. Moreover, it should be noted that the here observed fold differences were relatively low and additional validation and replication are necessary.

Since it has been suggested that miR-140 is co-expressed with *WWP2-C* [14], *WWP2* isoform 2, we generated upregulation of miR-140 in 2D primary chondrocytes to explore whether *WWP2* and miR-140 are involved in similar pathways. In the miR-140 upregulated cells, we observed significant increased expression levels of *WWP2*, indicating miR-140 indeed targets *WWP2*. Nonetheless, given the predicted *WWP2* target site of miR-140 (3’UTR of *WWP2* isoform 6 [23]) and the absence of SNPs in this region in linkage disequilibrium with the OA risk SNP rs1052429, the genetic *WWP2* risk nor the allelic imbalance is brought about via an aberrant miR-140 binding to *WWP2*. The fact that we did not observe consistent changes in expression levels of *WWP2* isoform 2, suggests that *WWP2* isoform 2 and miR-140 indeed share the intron 10 (*WWP2* full length) promotor as hypothesized previously by Rice at al. [15].

To our surprise, Styrkarsdottir et al. [2] reported on *WWP2* expression in adipose tissue as function of SNP rs4985453-G, a proxy of their identified OA risk allele rs34195470-G (R^2^=0.79) and our AI SNP rs1052429-A (R^2^=0.77), highlighting the OA risk allele being associated with lower expression levels of *WWP2*. Upon investigating the GTEx eQTL data of *WWP2* with the highlighted SNPs [39], we found only data showing consistently higher expression of *WWP2* as function of OA risk alleles of the respective SNPs across multiple tissues (**Fig. S4**), underscoring the aberrant effects observed here with *WWP2* upregulation. Although Mokuda et al. [13] showed that lack of *WWP2* expression in mice resulted in increased expression of *RUNX2* and *ADAMTS5*, we here did not observe consistent changes in expression of *RUNX2* or *ADAMTS5* upon upregulation of *WWP2* in our human chondrocyte pellet cultures and culturing for seven days. This difference could be due to translational limitations from mice to humans. Alternatively, we here create neocartilage, and the effect on *RUNX2* and *ADAMTS5* may be a temporal or time-dependent effect, which we do not observe at day seven of culturing.

In conclusion, our data provide support to our hypothesis that high levels of *WWP2* have detrimental effects on cartilage homeostasis. We identified *EPAS1, GJA1, GDF10*, and *STC2*, all genes involved in chondrocyte dedifferentiation, to be involved in the *WWP2* pathway. Moreover, we showed that miR-140 is likely involved in similar pathways as *WWP2* and miR-140 might play a role in regulating *WWP2* expression. Together these data contribute to a better understanding of how *WWP2* confers risk to OA and is a step towards translation from bench to bedside.

## ACKNOWLEDGEMENTS

We thank all the participants of the RAAK study. The LUMC has and is supporting the RAAK study. We thank all the members of our group for valuable discussion and feedback. We also thank Enrike van der Linden, Demiën Broekhuis, Peter van Schie, Shaho Hasan, Maartje Meijer, Daisy Latijnhouwers, Anika Rabelink-Hoogenstraaten, and Geert Spierenburg for collecting the RAAK material. We thank the Sequence Analysis Support Core (SASC) of the Leiden University Medical Center for their support. The study was funded by the Dutch Scientific Research council NWO/ZonMW VICI scheme (nr 91816631/528), Dutch Arthritis Society (DAA_10_1-402), BBMRI-NL complementation project (CP2013-83), European Commission Seventh Framework programme (TreatOA,200800), and Ana fonds (O2015-27). Data is generated within the scope of the Medical Delta programs Regenerative Medicine 4D: Generating complex tissues with stem cells and printing technology and Improving Mobility with Technology.

## STATEMENTS AND DECLARATIONS

### Data availability

The RNA sequencing data of the articular cartilage is deposited at ArrayExpress (E-MTAB-7313). Further data generated and used in this study is not openly available due to reasons of sensitivity and are available from the corresponding author upon reasonable request.

### Ethics approval

Samples used in this study are collected as part of the Research Arthritis and Articular Cartilage (RAAK study). The RAAK-study is approved by the medical ethics committee of the Leiden University Medical Center (P08.239/P19.013).

## SUPPORTING INFORMATION

**Fig. S1** - Interconnected network of genes highly correlating (|ρ|>0.7, FDR<0.05) to WWP2 in lesioned OA articular cartilage, with the positive correlations in green and the negative correlations in orange. The size of the nodes in the network indicate the number of connections.

**Fig. S2** – Overexpression of WWP2. (A) Monolayer of chondrocytes 3 days after transduction. (B) -ΔCT values of WWP2 overexpressed pellets and their controls at day 0 of the 3D pellet culture. (C) Volcano plot of proteomics analysis at day 0 showing WWP2 being upregulated

**Fig. S3** – Volcano plot of proteins differentially expressed between day three and day zero (A) and day seven and day zero (B) of pellet culture in the control pellets (N=16 pellet cultures, N=4 donors)

**Fig. S4** – GTEx violin plots of WWP2 expression as function of the three OA susceptibility alleles. In all three cases, the OA risk allele is associated to higher expression levels of WWP2 across multiple tissues.

**Table S1** – Baseline characteristics of material included in the current study

**Table S1A** – Sample characteristics of RNA-seq data for correlation

**Table S1B** – Sample characteristics of functional experiments

**Table S2 -** Spearman correlations between expression levels of WWP2 and genes expressed in articular cartilage (N=20048 genes) in lesioned OA cartilage samples.

**Table S3 -** Significant gene enrichment of 98 genes correlating to WWP2

**Table S4 -** Differential protein expression on day three and day seven relative to day zero of pellet culture in our 3D chondrocyte pellet cultures.

**Table S5 -** Differential protein expression between chondrocyte pellet cultures with and without upregulation of WWP2

