## SUPPORTING INFORMATION for "WWP2 osteoarthritis risk allele rs1052429-A confers risk by affecting cartilage matrix deposition via hypoxia associated genes"

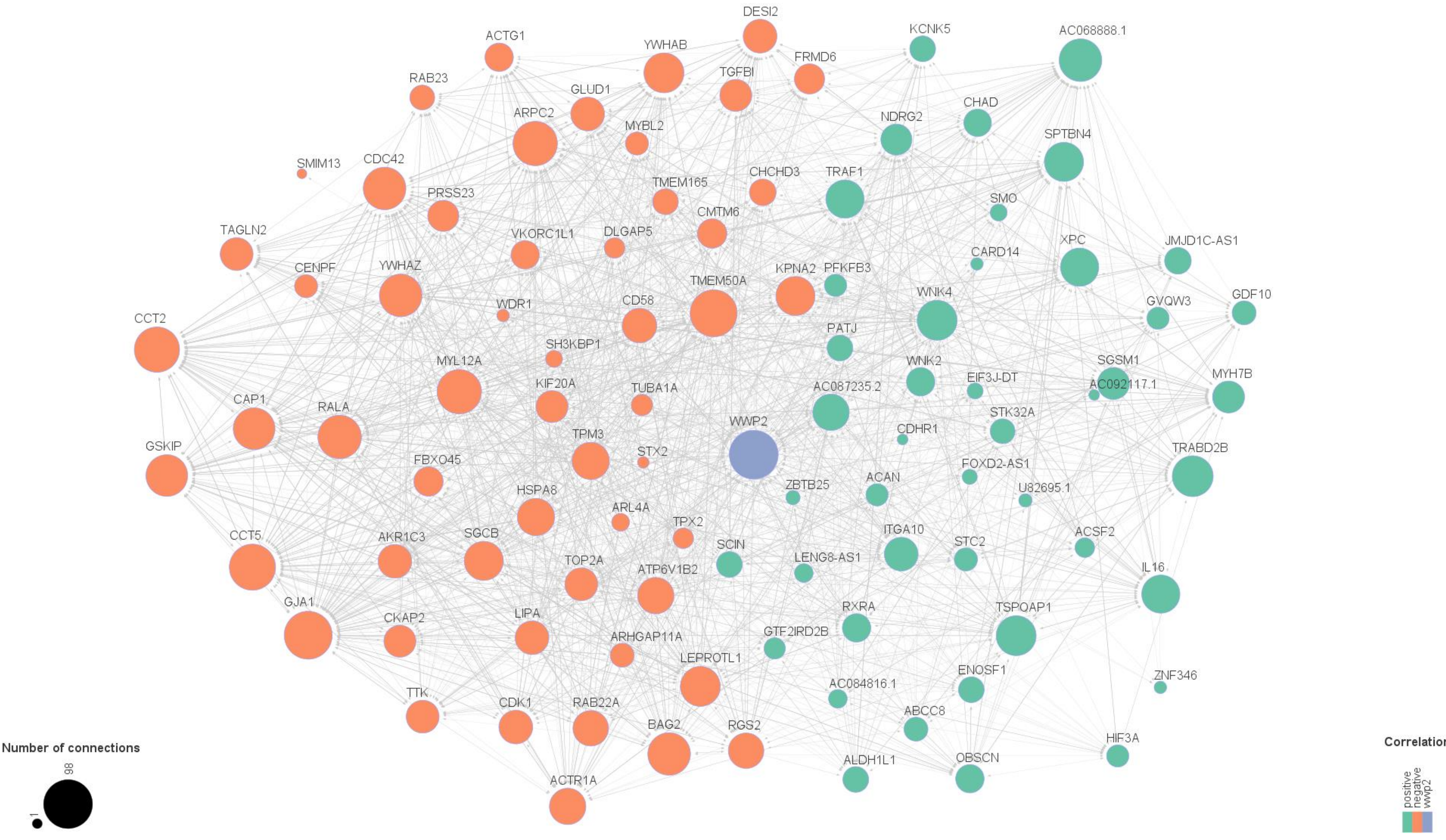


**Figure S1** – Interconnected network of genes highly correlating (|ρ|>0.7, FDR<0.05) to *WWP2* in lesioned OA articular cartilage, with the positive correlations in green and the negative correlations in orange. The size of the nodes in the network indicate the number of connections.

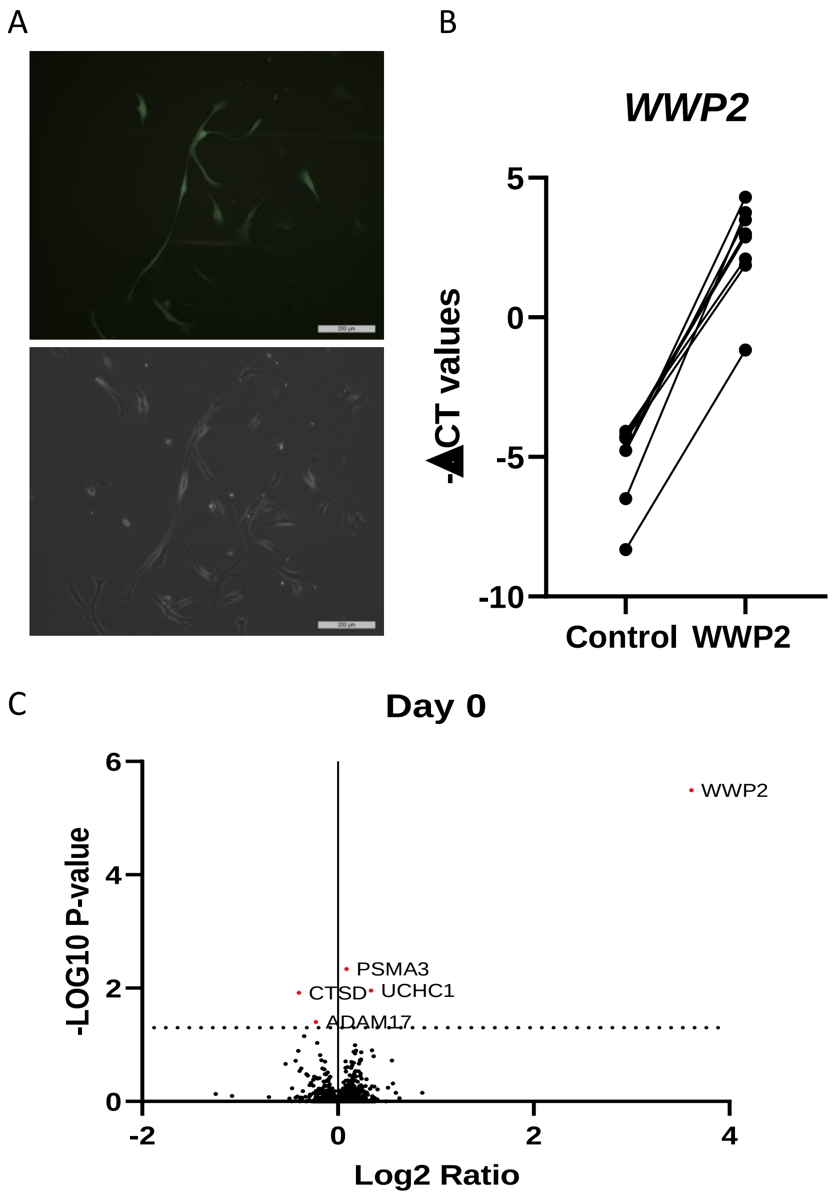


**Figure S2** – Overexpression of *WWP2*. (A) Monolayer of chondrocytes 3 days after transduction. (B) -ΔCT values of *WWP2* overexpressed pellets and their controls at day 0 of the 3D pellet culture. (C) Volcano plot of proteomics analysis at day 0 showing WWP2 being upregulated

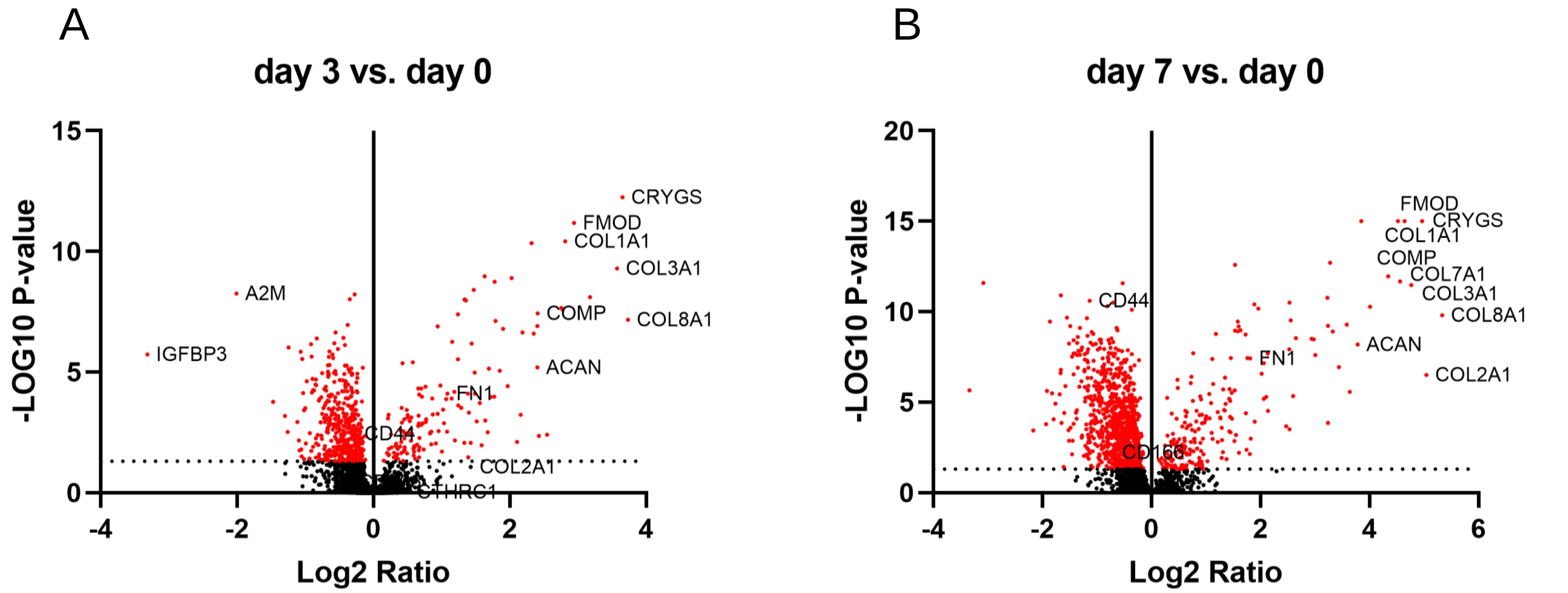


**Figure S3** – Volcano plot of proteins differentially expressed between day three and day zero (A) and day seven and day zero (B) of pellet culture in the control pellets (N=16 pellet cultures, N=4 donors)

Risk allele: Expression level WWP2:

rs34195470-G

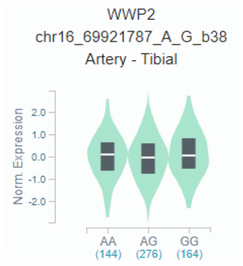


rs4985453-G

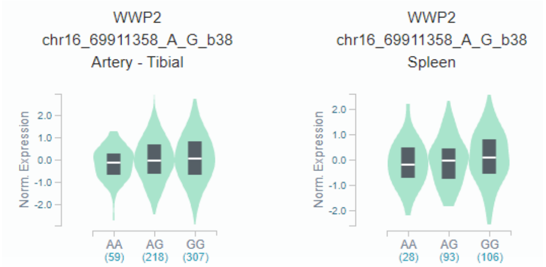


rs1052429-A

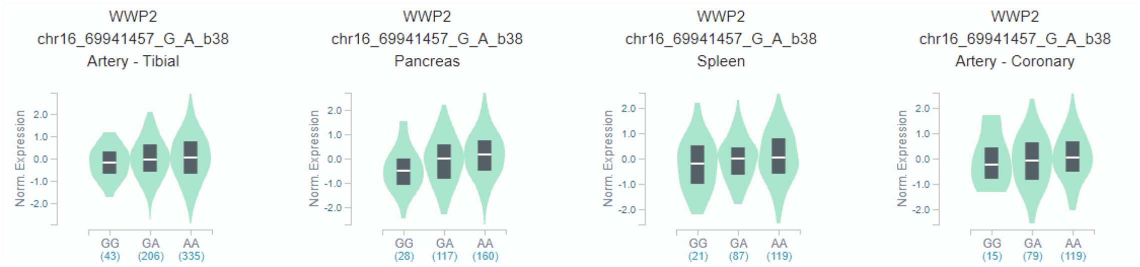


**Figure S4** – GTEx violin plots of WWP2 expression as function of the three OA susceptibility alleles. In all three cases, the OA risk allele is associated to higher expression levels of WWP2 across multiple tissues.

**Table S1**: Baseline characteristics of material included in the current study

**Table S1A**: Sample characteristics of RNA-seq data for correlation

| | **RNA-seq data (N=35)** |
|---|---|
| **Participants** | 35 |
| **Age (SD)** | 68,6 (9,0) |
| **Knees (Hips)** | 28 (7) |
| **Females (Males)** | 27 (7) |

**Table S1B**: Sample characteristics of functional experiments

| | **Lentiviral particle mediated overexpression of WWP2 (Gene expression, N=8)** | **Lentiviral particle mediated overexpression of WWP2 (proteomics, N=4)** | **Transfection miR-140 (N=7)** |
|---|---|---|---|
| **Age (SD)** | 71,1 (9,2) | 75,5 (11,6) | 71,6 (9,8) |
| **Knees (Hips)** | 8 (0) | 4 (0) | 7 (0) |
| **Females (Males)** | 5(3) | 2 (2) | 5(2) |

**Table S2** - Top 25 % Spearman correlations (|ρ|) between expression level of WWP2 and genes expressed in lesioned OA articular cartilage (N=20048 genes)

| Ensembl ID | Gene Name | ρ | Pval | Padj |
|---|---|---|---|---|
| ENSG00000256321 | AC087235.2 | 0.84 | 3.26E-10 | 6.54E-06 |
| ENSG00000152661 | GJA1 | -0.81 | 2.93E-09 | 2.93E-05 |
| ENSG00000126562 | WNK4 | 0.81 | 4.56E-09 | 3.04E-05 |
| ENSG00000196715 | VKORC1L1 | -0.80 | 6.71E-09 | 3.36E-05 |
| ENSG00000100744 | GSKIP | -0.79 | 1.59E-08 | 5.64E-05 |
| ENSG00000138107 | ACTR1A | -0.79 | 1.75E-08 | 5.64E-05 |
| ENSG00000269113 | TRABD2B | 0.79 | 1.97E-08 | 5.64E-05 |
| ENSG00000112208 | BAG2 | -0.79 | 2.30E-08 | 5.76E-05 |
| ENSG00000128602 | SMO | 0.78 | 2.89E-08 | 6.44E-05 |
| ENSG00000112742 | TTK | -0.78 | 4.36E-08 | 8.58E-05 |
| ENSG00000182481 | KPNA2 | -0.77 | 4.95E-08 | 8.58E-05 |
| ENSG00000089775 | ZBTB25 | 0.77 | 5.13E-08 | 8.58E-05 |
| ENSG00000113739 | STC2 | 0.77 | 6.96E-08 | 1.01E-04 |
| ENSG00000154767 | XPC | 0.77 | 7.08E-08 | 1.01E-04 |
| ENSG00000150753 | CCT5 | -0.76 | 9.05E-08 | 1.19E-04 |
| ENSG00000157766 | ACAN | 0.76 | 9.53E-08 | 1.19E-04 |
| ENSG00000257337 | AC068888.1 | 0.76 | 1.08E-07 | 1.22E-04 |
| ENSG00000174013 | FBXO45 | -0.76 | 1.09E-07 | 1.22E-04 |
| ENSG00000131747 | TOP2A | -0.76 | 1.21E-07 | 1.24E-04 |
| ENSG00000164626 | KCNK5 | 0.76 | 1.30E-07 | 1.24E-04 |
| ENSG00000256995 | AC084816.1 | 0.76 | 1.30E-07 | 1.24E-04 |
| ENSG00000141527 | CARD14 | 0.76 | 1.36E-07 | 1.24E-04 |
| ENSG00000111450 | STX2 | -0.75 | 1.66E-07 | 1.45E-04 |
| ENSG00000131236 | CAP1 | -0.75 | 1.90E-07 | 1.52E-04 |
| ENSG00000167037 | SGSM1 | 0.75 | 1.90E-07 | 1.52E-04 |
| ENSG00000147010 | SH3KBP1 | -0.75 | 2.13E-07 | 1.59E-04 |
| ENSG00000158710 | TAGLN2 | -0.75 | 2.27E-07 | 1.59E-04 |
| ENSG00000226696 | LENG8-AS1 | 0.75 | 2.27E-07 | 1.59E-04 |
| ENSG00000056558 | TRAF1 | 0.75 | 2.31E-07 | 1.59E-04 |
| ENSG00000166226 | CCT2 | -0.75 | 2.58E-07 | 1.70E-04 |
| ENSG00000136457 | CHAD | 0.75 | 2.75E-07 | 1.70E-04 |
| ENSG00000170312 | CDK1 | -0.75 | 2.75E-07 | 1.70E-04 |
| ENSG00000183726 | TMEM50A | -0.74 | 2.88E-07 | 1.70E-04 |
| ENSG00000091317 | CMTM6 | -0.74 | 2.93E-07 | 1.70E-04 |
| ENSG00000124209 | RAB22A | -0.74 | 2.97E-07 | 1.70E-04 |
| ENSG00000136108 | CKAP2 | -0.74 | 3.22E-07 | 1.72E-04 |
| ENSG00000172349 | IL16 | 0.74 | 3.22E-07 | 1.72E-04 |
| ENSG00000150687 | PRSS23 | -0.74 | 3.27E-07 | 1.72E-04 |
| ENSG00000148672 | GLUD1 | -0.74 | 3.59E-07 | 1.85E-04 |
| ENSG00000184009 | ACTG1 | -0.74 | 3.76E-07 | 1.88E-04 |
| ENSG00000186350 | RXRA | 0.74 | 3.88E-07 | 1.90E-04 |
| ENSG00000132849 | PATJ | 0.74 | 4.06E-07 | 1.94E-04 |

| Ensembl ID | Gene Name | ρ | Pval | Padj |
|---|---|---|---|---|
| ENSG00000276791 | AC092117.1 | 0.74 | 4.45E-07 | 2.01E-04 |
| ENSG00000104660 | LEPROTL1 | -0.74 | 4.52E-07 | 2.01E-04 |
| ENSG00000116741 | RGS2 | -0.74 | 4.52E-07 | 2.01E-04 |
| ENSG00000224963 | U82695.1 | 0.74 | 4.66E-07 | 2.03E-04 |
| ENSG00000144908 | ALDH1L1 | 0.74 | 4.95E-07 | 2.07E-04 |
| ENSG00000167552 | TUBA1A | -0.74 | 4.95E-07 | 2.07E-04 |
| ENSG00000006451 | RALA | -0.73 | 5.50E-07 | 2.25E-04 |
| ENSG00000163466 | ARPC2 | -0.73 | 6.48E-07 | 2.60E-04 |
| ENSG00000196139 | AKR1C3 | -0.73 | 6.88E-07 | 2.70E-04 |
| ENSG00000169302 | STK32A | 0.73 | 7.29E-07 | 2.81E-04 |
| ENSG00000101608 | MYL12A | -0.73 | 7.62E-07 | 2.88E-04 |
| ENSG00000088325 | TPX2 | -0.73 | 7.84E-07 | 2.90E-04 |
| ENSG00000107798 | LIPA | -0.73 | 7.96E-07 | 2.90E-04 |
| ENSG00000120708 | TGFBI | -0.73 | 8.19E-07 | 2.93E-04 |
| ENSG00000112984 | KIF20A | -0.72 | 8.53E-07 | 3.00E-04 |
| ENSG00000147416 | ATP6V1B2 | -0.72 | 8.94E-07 | 3.08E-04 |
| ENSG00000266524 | GDF10 | 0.72 | 9.07E-07 | 3.08E-04 |
| ENSG00000167107 | ACSF2 | 0.72 | 9.33E-07 | 3.12E-04 |
| ENSG00000237424 | FOXD2-AS1 | 0.72 | 1.02E-06 | 3.34E-04 |
| ENSG00000165238 | WNK2 | 0.72 | 1.05E-06 | 3.37E-04 |
| ENSG00000224531 | SMIM13 | -0.72 | 1.06E-06 | 3.37E-04 |
| ENSG00000134851 | TMEM165 | -0.72 | 1.08E-06 | 3.37E-04 |
| ENSG00000166913 | YWHAB | -0.72 | 1.15E-06 | 3.50E-04 |
| ENSG00000006071 | ABCC8 | 0.72 | 1.17E-06 | 3.50E-04 |
| ENSG00000170525 | PFKFB3 | 0.72 | 1.17E-06 | 3.50E-04 |
| ENSG00000143549 | TPM3 | -0.72 | 1.22E-06 | 3.60E-04 |
| ENSG00000272767 | JMJD1C-AS1 | 0.72 | 1.24E-06 | 3.60E-04 |
| ENSG00000174428 | GTF2IRD2B | 0.72 | 1.26E-06 | 3.60E-04 |
| ENSG00000078814 | MYH7B | 0.72 | 1.29E-06 | 3.60E-04 |
| ENSG00000148600 | CDHR1 | 0.72 | 1.29E-06 | 3.60E-04 |
| ENSG00000126787 | DLGAP5 | -0.72 | 1.34E-06 | 3.68E-04 |
| ENSG00000112210 | RAB23 | -0.71 | 1.40E-06 | 3.75E-04 |
| ENSG00000165795 | NDRG2 | 0.71 | 1.40E-06 | 3.75E-04 |
| ENSG00000179240 | GVQW3 | 0.71 | 1.46E-06 | 3.86E-04 |
| ENSG00000113761 | ZNF346 | 0.71 | 1.53E-06 | 3.97E-04 |
| ENSG00000139926 | FRMD6 | -0.71 | 1.55E-06 | 3.98E-04 |
| ENSG00000179523 | EIF3J-DT | 0.71 | 1.57E-06 | 3.98E-04 |
| ENSG00000101057 | MYBL2 | -0.71 | 1.70E-06 | 4.27E-04 |
| ENSG00000116815 | CD58 | -0.71 | 1.80E-06 | 4.45E-04 |
| ENSG00000121644 | DESI2 | -0.71 | 1.82E-06 | 4.46E-04 |
| ENSG00000164924 | YWHAZ | -0.71 | 1.87E-06 | 4.52E-04 |
| ENSG00000198826 | ARHGAP11A | -0.71 | 2.00E-06 | 4.78E-04 |
| ENSG00000109971 | HSPA8 | -0.71 | 2.11E-06 | 4.99E-04 |
| ENSG00000005379 | TSPOAP1 | 0.71 | 2.26E-06 | 5.21E-04 |
| ENSG00000122644 | ARL4A | -0.71 | 2.26E-06 | 5.21E-04 |

| Ensembl ID | Gene Name | ρ | Pval | Padj |
|---|---|---|---|---|
| ENSG00000106554 | CHCHD3 | -0.70 | 2.35E-06 | 5.25E-04 |
| ENSG00000160460 | SPTBN4 | 0.70 | 2.35E-06 | 5.25E-04 |
| ENSG00000070831 | CDC42 | -0.70 | 2.38E-06 | 5.25E-04 |
| ENSG00000117724 | CENPF | -0.70 | 2.38E-06 | 5.25E-04 |
| ENSG00000124440 | HIF3A | 0.70 | 2.45E-06 | 5.27E-04 |
| ENSG00000143127 | ITGA10 | 0.70 | 2.45E-06 | 5.27E-04 |
| ENSG00000071127 | WDR1 | -0.70 | 2.48E-06 | 5.29E-04 |
| ENSG00000132199 | ENOSF1 | 0.70 | 2.58E-06 | 5.44E-04 |
| ENSG00000163069 | SGCB | -0.70 | 2.65E-06 | 5.53E-04 |
| ENSG00000154358 | OBSCN | 0.70 | 2.72E-06 | 5.62E-04 |
| ENSG00000006747 | SCIN | 0.70 | 2.75E-06 | 5.63E-04 |
| ENSG00000104490 | NCALD | -0.70 | 2.94E-06 | 5.95E-04 |
| ENSG00000048544 | MRPS10 | -0.70 | 2.98E-06 | 5.97E-04 |
| ENSG00000088832 | FKBP1A | -0.70 | 3.02E-06 | 5.99E-04 |
| ENSG00000198556 | ZNF789 | 0.70 | 3.22E-06 | 6.33E-04 |
| ENSG00000077684 | JADE1 | 0.70 | 3.35E-06 | 6.51E-04 |
| ENSG00000235601 | BARX1-DT | 0.70 | 3.43E-06 | 6.62E-04 |
| ENSG00000162704 | ARPC5 | -0.70 | 3.48E-06 | 6.64E-04 |
| ENSG00000081760 | AACS | 0.69 | 3.76E-06 | 7.10E-04 |
| ENSG00000182718 | ANXA2 | -0.69 | 4.10E-06 | 7.69E-04 |
| ENSG00000162734 | PEA15 | -0.69 | 4.26E-06 | 7.91E-04 |
| ENSG00000088367 | EPB41L1 | 0.69 | 4.37E-06 | 8.04E-04 |
| ENSG00000006007 | GDE1 | -0.69 | 4.54E-06 | 8.15E-04 |
| ENSG00000109906 | ZBTB16 | 0.69 | 4.54E-06 | 8.15E-04 |
| ENSG00000016391 | CHDH | 0.69 | 4.60E-06 | 8.15E-04 |
| ENSG00000100307 | CBX7 | 0.69 | 4.60E-06 | 8.15E-04 |
| ENSG00000171488 | LRRC8C | -0.69 | 4.95E-06 | 8.67E-04 |
| ENSG00000106105 | GARS | -0.69 | 5.02E-06 | 8.67E-04 |
| ENSG00000165244 | ZNF367 | -0.69 | 5.02E-06 | 8.67E-04 |
| ENSG00000122756 | CNTFR | 0.69 | 5.08E-06 | 8.70E-04 |
| ENSG00000107438 | PDLIM1 | -0.69 | 5.21E-06 | 8.70E-04 |
| ENSG00000100568 | VTI1B | -0.69 | 5.27E-06 | 8.70E-04 |
| ENSG00000168615 | ADAM9 | -0.69 | 5.27E-06 | 8.70E-04 |
| ENSG00000116237 | ICMT | -0.69 | 5.34E-06 | 8.70E-04 |
| ENSG00000163884 | KLF15 | 0.69 | 5.34E-06 | 8.70E-04 |
| ENSG00000164050 | PLXNB1 | 0.69 | 5.34E-06 | 8.70E-04 |
| ENSG00000100522 | GNPNAT1 | -0.69 | 5.40E-06 | 8.73E-04 |
| ENSG00000175203 | DCTN2 | -0.69 | 5.54E-06 | 8.88E-04 |
| ENSG00000113161 | HMGCR | -0.69 | 5.61E-06 | 8.92E-04 |
| ENSG00000168078 | PBK | -0.68 | 5.70E-06 | 8.99E-04 |
| ENSG00000239779 | WBP1 | 0.68 | 5.75E-06 | 9.00E-04 |
| ENSG00000166446 | CDYL2 | -0.68 | 5.82E-06 | 9.04E-04 |
| ENSG00000231160 | KLF3-AS1 | 0.68 | 5.89E-06 | 9.08E-04 |
| ENSG00000077549 | CAPZB | -0.68 | 5.96E-06 | 9.12E-04 |
| ENSG00000078124 | ACER3 | -0.68 | 6.11E-06 | 9.21E-04 |

| Ensembl ID | Gene Name | ρ | Pval | Padj |
|---|---|---|---|---|
| ENSG00000105974 | CAV1 | -0.68 | 6.11E-06 | 9.21E-04 |
| ENSG00000185100 | ADSSL1 | 0.68 | 6.18E-06 | 9.25E-04 |
| ENSG00000108960 | MMD | -0.68 | 6.26E-06 | 9.27E-04 |
| ENSG00000081087 | OSTM1 | -0.68 | 6.34E-06 | 9.27E-04 |
| ENSG00000160111 | CPAMD8 | 0.68 | 6.34E-06 | 9.27E-04 |
| ENSG00000073670 | ADAM11 | 0.68 | 6.56E-06 | 9.41E-04 |
| ENSG00000115993 | TRAK2 | -0.68 | 6.57E-06 | 9.41E-04 |
| ENSG00000142731 | PLK4 | -0.68 | 6.57E-06 | 9.41E-04 |
| ENSG00000114346 | ECT2 | -0.68 | 6.73E-06 | 9.57E-04 |
| ENSG00000164294 | GPX8 | -0.68 | 6.81E-06 | 9.62E-04 |
| ENSG00000129195 | PIMREG | -0.68 | 7.27E-06 | 1.00E-03 |
| ENSG00000158560 | DYNC1I1 | 0.68 | 7.33E-06 | 1.00E-03 |
| ENSG00000261094 | AC007066.2 | 0.68 | 7.35E-06 | 1.00E-03 |
| ENSG00000105971 | CAV2 | -0.68 | 7.41E-06 | 1.00E-03 |
| ENSG00000149547 | EI24 | -0.68 | 7.41E-06 | 1.00E-03 |
| ENSG00000171017 | LRRC8E | -0.68 | 7.41E-06 | 1.00E-03 |
| ENSG00000170759 | KIF5B | -0.68 | 7.59E-06 | 1.02E-03 |
| ENSG00000183291 | SELENOF | -0.68 | 7.78E-06 | 1.04E-03 |
| ENSG00000066279 | ASPM | -0.68 | 7.87E-06 | 1.04E-03 |
| ENSG00000075223 | SEMA3C | -0.68 | 8.16E-06 | 1.07E-03 |
| ENSG00000108474 | PIGL | 0.68 | 8.16E-06 | 1.07E-03 |
| ENSG00000123416 | TUBA1B | -0.68 | 8.45E-06 | 1.10E-03 |
| ENSG00000054392 | HHAT | 0.68 | 8.66E-06 | 1.11E-03 |
| ENSG00000165275 | TRMT10B | 0.68 | 8.66E-06 | 1.11E-03 |
| ENSG00000123999 | INHA | 0.67 | 8.72E-06 | 1.11E-03 |
| ENSG00000274292 | AC084018.2 | 0.67 | 8.86E-06 | 1.12E-03 |
| ENSG00000260641 | AC114811.2 | 0.67 | 8.88E-06 | 1.12E-03 |
| ENSG00000100596 | SPTLC2 | -0.67 | 8.97E-06 | 1.12E-03 |
| ENSG00000116157 | GPX7 | -0.67 | 9.08E-06 | 1.13E-03 |
| ENSG00000117399 | CDC20 | -0.67 | 9.18E-06 | 1.13E-03 |
| ENSG00000260261 | AC124944.3 | 0.67 | 9.18E-06 | 1.13E-03 |
| ENSG00000146386 | ABRACL | -0.67 | 9.29E-06 | 1.14E-03 |
| ENSG00000198805 | PNP | -0.67 | 9.51E-06 | 1.16E-03 |
| ENSG00000244242 | IFITM10 | 0.67 | 9.62E-06 | 1.16E-03 |
| ENSG00000179406 | LINC00174 | 0.67 | 9.97E-06 | 1.20E-03 |
| ENSG00000196199 | MPHOSPH8 | 0.67 | 1.01E-05 | 1.20E-03 |
| ENSG00000123130 | ACOT9 | -0.67 | 1.02E-05 | 1.21E-03 |
| ENSG00000256982 | AC135782.1 | 0.67 | 1.03E-05 | 1.21E-03 |
| ENSG00000197006 | METTL9 | -0.67 | 1.04E-05 | 1.22E-03 |
| ENSG00000077721 | UBE2A | -0.67 | 1.09E-05 | 1.27E-03 |
| ENSG00000123485 | HJURP | -0.67 | 1.10E-05 | 1.27E-03 |
| ENSG00000188770 | OPTC | 0.67 | 1.11E-05 | 1.28E-03 |
| ENSG00000103187 | COTL1 | -0.67 | 1.12E-05 | 1.28E-03 |
| ENSG00000185168 | LINC00482 | 0.67 | 1.13E-05 | 1.29E-03 |
| ENSG00000101191 | DIDO1 | 0.67 | 1.16E-05 | 1.31E-03 |

| Ensembl ID | Gene Name | ρ | Pval | Padj |
|---|---|---|---|---|
| ENSG00000231920 | NEBL-AS1 | 0.67 | 1.16E-05 | 1.31E-03 |
| ENSG00000101166 | PRELID3B | -0.67 | 1.19E-05 | 1.33E-03 |
| ENSG00000169508 | GPR183 | -0.67 | 1.20E-05 | 1.33E-03 |
| ENSG00000075035 | WSCD2 | 0.67 | 1.21E-05 | 1.34E-03 |
| ENSG00000196954 | CASP4 | -0.67 | 1.23E-05 | 1.35E-03 |
| ENSG00000129244 | ATP1B2 | 0.67 | 1.24E-05 | 1.35E-03 |
| ENSG00000197747 | S100A10 | -0.67 | 1.24E-05 | 1.35E-03 |
| ENSG00000077585 | GPR137B | -0.67 | 1.26E-05 | 1.36E-03 |
| ENSG00000171954 | CYP4F22 | 0.67 | 1.30E-05 | 1.40E-03 |
| ENSG00000141429 | GALNT1 | -0.66 | 1.35E-05 | 1.44E-03 |
| ENSG00000223475 | AC099681.1 | 0.66 | 1.37E-05 | 1.45E-03 |
| ENSG00000140157 | NIPA2 | -0.66 | 1.38E-05 | 1.45E-03 |
| ENSG00000117632 | STMN1 | -0.66 | 1.39E-05 | 1.45E-03 |
| ENSG00000181045 | SLC26A11 | 0.66 | 1.39E-05 | 1.45E-03 |
| ENSG00000272645 | GTF2IP20 | 0.66 | 1.39E-05 | 1.45E-03 |
| ENSG00000113916 | BCL6 | 0.66 | 1.41E-05 | 1.46E-03 |
| ENSG00000187792 | ZNF70 | 0.66 | 1.44E-05 | 1.49E-03 |
| ENSG00000121964 | GTDC1 | -0.66 | 1.47E-05 | 1.51E-03 |
| ENSG00000112406 | HECA | -0.66 | 1.49E-05 | 1.51E-03 |
| ENSG00000113575 | PPP2CA | -0.66 | 1.49E-05 | 1.51E-03 |
| ENSG00000275120 | AC048382.5 | 0.66 | 1.49E-05 | 1.51E-03 |
| ENSG00000121152 | NCAPH | -0.66 | 1.53E-05 | 1.54E-03 |
| ENSG00000183688 | RFLNB | -0.66 | 1.56E-05 | 1.56E-03 |
| ENSG00000086062 | B4GALT1 | -0.66 | 1.63E-05 | 1.61E-03 |
| ENSG00000151690 | MFSD6 | -0.66 | 1.63E-05 | 1.61E-03 |
| ENSG00000081665 | ZNF506 | 0.66 | 1.65E-05 | 1.61E-03 |
| ENSG00000119771 | KLHL29 | 0.66 | 1.65E-05 | 1.61E-03 |
| ENSG00000152642 | GPD1L | -0.66 | 1.65E-05 | 1.61E-03 |
| ENSG00000100226 | GTPBP1 | 0.66 | 1.68E-05 | 1.64E-03 |
| ENSG00000123505 | AMD1 | -0.66 | 1.78E-05 | 1.69E-03 |
| ENSG00000144857 | BOC | 0.66 | 1.78E-05 | 1.69E-03 |
| ENSG00000173457 | PPP1R14B | -0.66 | 1.78E-05 | 1.69E-03 |
| ENSG00000004139 | SARM1 | 0.66 | 1.80E-05 | 1.69E-03 |
| ENSG00000011426 | ANLN | -0.66 | 1.80E-05 | 1.69E-03 |
| ENSG00000065135 | GNAI3 | -0.66 | 1.80E-05 | 1.69E-03 |
| ENSG00000106462 | EZH2 | -0.66 | 1.80E-05 | 1.69E-03 |
| ENSG00000137364 | TPMT | -0.66 | 1.82E-05 | 1.71E-03 |
| ENSG00000136449 | MYCBPAP | 0.66 | 1.88E-05 | 1.75E-03 |
| ENSG00000162913 | OBSCN-AS1 | 0.66 | 1.90E-05 | 1.75E-03 |
| ENSG00000166780 | C16orf45 | -0.66 | 1.90E-05 | 1.75E-03 |
| ENSG00000100554 | ATP6V1D | -0.66 | 1.92E-05 | 1.75E-03 |
| ENSG00000163191 | S100A11 | -0.66 | 1.92E-05 | 1.75E-03 |
| ENSG00000277449 | CEBPB-AS1 | 0.66 | 1.92E-05 | 1.75E-03 |
| ENSG00000099365 | STX1B | 0.66 | 1.95E-05 | 1.76E-03 |
| ENSG00000144711 | IQSEC1 | 0.65 | 1.97E-05 | 1.77E-03 |

| Ensembl ID | Gene Name | ρ | Pval | Padj |
|---|---|---|---|---|
| ENSG00000175106 | TVP23C | 0.65 | 1.97E-05 | 1.77E-03 |
| ENSG00000124006 | OBSL1 | 0.65 | 1.99E-05 | 1.77E-03 |
| ENSG00000188825 | LINC00910 | 0.65 | 1.99E-05 | 1.77E-03 |
| ENSG00000172667 | ZMAT3 | -0.65 | 2.01E-05 | 1.78E-03 |
| ENSG00000259146 | AC005476.2 | 0.65 | 2.01E-05 | 1.78E-03 |
| ENSG00000160712 | IL6R | 0.65 | 2.03E-05 | 1.79E-03 |
| ENSG00000012822 | CALCOCO1 | 0.65 | 2.08E-05 | 1.81E-03 |
| ENSG00000274925 | ZKSCAN2-DT | 0.65 | 2.08E-05 | 1.81E-03 |
| ENSG00000235954 | TTC28-AS1 | 0.65 | 2.12E-05 | 1.84E-03 |
| ENSG00000159720 | ATP6V0D1 | -0.65 | 2.15E-05 | 1.85E-03 |
| ENSG00000198951 | NAGA | -0.65 | 2.15E-05 | 1.85E-03 |
| ENSG00000122861 | PLAU | -0.65 | 2.18E-05 | 1.86E-03 |
| ENSG00000080986 | NDC80 | -0.65 | 2.19E-05 | 1.86E-03 |
| ENSG00000184988 | TMEM106A | 0.65 | 2.19E-05 | 1.86E-03 |
| ENSG00000167191 | GPRC5B | 0.65 | 2.24E-05 | 1.90E-03 |
| ENSG00000213189 | BTF3L4P2 | -0.65 | 2.27E-05 | 1.91E-03 |
| ENSG00000131773 | KHDRBS3 | -0.65 | 2.29E-05 | 1.92E-03 |
| ENSG00000182220 | ATP6AP2 | -0.65 | 2.32E-05 | 1.93E-03 |
| ENSG00000226891 | LINC01359 | 0.65 | 2.43E-05 | 2.02E-03 |
| ENSG00000229589 | ACVR2B-AS1 | 0.65 | 2.47E-05 | 2.05E-03 |
| ENSG00000226029 | LINC01772 | 0.65 | 2.50E-05 | 2.06E-03 |
| ENSG00000038295 | TLL1 | -0.65 | 2.52E-05 | 2.06E-03 |
| ENSG00000110455 | ACCS | 0.65 | 2.52E-05 | 2.06E-03 |
| ENSG00000281392 | AC107204.1 | 0.65 | 2.52E-05 | 2.06E-03 |
| ENSG00000132718 | SYT11 | -0.65 | 2.61E-05 | 2.11E-03 |
| ENSG00000198833 | UBE2J1 | -0.65 | 2.61E-05 | 2.11E-03 |
| ENSG00000184743 | ATL3 | -0.65 | 2.63E-05 | 2.11E-03 |
| ENSG00000250644 | AC068580.4 | 0.65 | 2.63E-05 | 2.11E-03 |
| ENSG00000187595 | ZNF385C | 0.65 | 2.66E-05 | 2.11E-03 |
| ENSG00000198356 | ASNA1 | -0.65 | 2.66E-05 | 2.11E-03 |
| ENSG00000236603 | RANP1 | -0.65 | 2.66E-05 | 2.11E-03 |
| ENSG00000155465 | SLC7A7 | -0.65 | 2.69E-05 | 2.12E-03 |
| ENSG00000229132 | EIF4A1P10 | -0.65 | 2.69E-05 | 2.12E-03 |
| ENSG00000246223 | LINC01550 | 0.65 | 2.70E-05 | 2.12E-03 |
| ENSG00000148773 | MKI67 | -0.65 | 2.72E-05 | 2.12E-03 |
| ENSG00000147576 | ADHFE1 | 0.65 | 2.75E-05 | 2.13E-03 |
| ENSG00000233251 | AC007743.1 | 0.65 | 2.75E-05 | 2.13E-03 |
| ENSG00000132646 | PCNA | -0.65 | 2.87E-05 | 2.20E-03 |
| ENSG00000136867 | SLC31A2 | -0.65 | 2.87E-05 | 2.20E-03 |
| ENSG00000215452 | ZNF663P | 0.65 | 2.87E-05 | 2.20E-03 |
| ENSG00000117360 | PRPF3 | 0.64 | 2.90E-05 | 2.20E-03 |
| ENSG00000119403 | PHF19 | 0.64 | 2.90E-05 | 2.20E-03 |
| ENSG00000167553 | TUBA1C | -0.64 | 2.93E-05 | 2.22E-03 |
| ENSG00000114405 | C3orf14 | -0.64 | 2.99E-05 | 2.26E-03 |
| ENSG00000166816 | LDHD | 0.64 | 3.02E-05 | 2.27E-03 |

| Ensembl ID | Gene Name | ρ | Pval | Padj |
|---|---|---|---|---|
| ENSG00000106004 | HOXA5 | 0.64 | 3.06E-05 | 2.29E-03 |
| ENSG00000143847 | PPFIA4 | 0.64 | 3.09E-05 | 2.29E-03 |
| ENSG00000205413 | SAMD9 | -0.64 | 3.09E-05 | 2.29E-03 |
| ENSG00000148426 | PROSER2 | 0.64 | 3.12E-05 | 2.31E-03 |
| ENSG00000135046 | ANXA1 | -0.64 | 3.15E-05 | 2.33E-03 |
| ENSG00000087586 | AURKA | -0.64 | 3.19E-05 | 2.33E-03 |
| ENSG00000187653 | TMSB4XP8 | -0.64 | 3.19E-05 | 2.33E-03 |
| ENSG00000121075 | TBX4 | 0.64 | 3.22E-05 | 2.35E-03 |
| ENSG00000143622 | RIT1 | -0.64 | 3.26E-05 | 2.36E-03 |
| ENSG00000162073 | PAQR4 | -0.64 | 3.32E-05 | 2.41E-03 |
| ENSG00000109814 | UGDH | -0.64 | 3.36E-05 | 2.42E-03 |
| ENSG00000080815 | PSEN1 | -0.64 | 3.39E-05 | 2.44E-03 |
| ENSG00000043143 | JADE2 | 0.64 | 3.43E-05 | 2.46E-03 |
| ENSG00000089847 | ANKRD24 | 0.64 | 3.47E-05 | 2.47E-03 |
| ENSG00000186153 | WWOX | 0.64 | 3.50E-05 | 2.49E-03 |
| ENSG00000157557 | ETS2 | 0.64 | 3.54E-05 | 2.51E-03 |
| ENSG00000115762 | PLEKHB2 | -0.64 | 3.58E-05 | 2.52E-03 |
| ENSG00000165304 | MELK | -0.64 | 3.59E-05 | 2.52E-03 |
| ENSG00000135052 | GOLM1 | -0.64 | 3.61E-05 | 2.53E-03 |
| ENSG00000275549 | STPG3-AS1 | 0.64 | 3.65E-05 | 2.55E-03 |
| ENSG00000227354 | RBM26-AS1 | 0.64 | 3.69E-05 | 2.57E-03 |
| ENSG00000110218 | PANX1 | -0.64 | 3.73E-05 | 2.59E-03 |
| ENSG00000049759 | NEDD4L | -0.64 | 3.77E-05 | 2.59E-03 |
| ENSG00000067560 | RHOA | -0.64 | 3.77E-05 | 2.59E-03 |
| ENSG00000137812 | KNL1 | -0.64 | 3.77E-05 | 2.59E-03 |
| ENSG00000100321 | SYNGR1 | 0.64 | 3.84E-05 | 2.63E-03 |
| ENSG00000122574 | WIPF3 | 0.64 | 3.86E-05 | 2.63E-03 |
| ENSG00000131462 | TUBG1 | -0.64 | 3.88E-05 | 2.64E-03 |
| ENSG00000077152 | UBE2T | -0.64 | 3.94E-05 | 2.66E-03 |
| ENSG00000105767 | CADM4 | -0.64 | 3.96E-05 | 2.66E-03 |
| ENSG00000170027 | YWHAG | -0.64 | 3.96E-05 | 2.66E-03 |
| ENSG00000258704 | SRP54-AS1 | 0.64 | 3.96E-05 | 2.66E-03 |
| ENSG00000253141 | AC008632.1 | 0.64 | 3.98E-05 | 2.66E-03 |
| ENSG00000178741 | COX5A | -0.64 | 4.05E-05 | 2.70E-03 |
| ENSG00000179335 | CLK3 | 0.64 | 4.13E-05 | 2.72E-03 |
| ENSG00000226824 | AC006001.2 | 0.64 | 4.13E-05 | 2.72E-03 |
| ENSG00000269837 | IPO5P1 | 0.64 | 4.13E-05 | 2.72E-03 |
| ENSG00000215241 | LINC02449 | 0.63 | 4.22E-05 | 2.77E-03 |
| ENSG00000135930 | EIF4E2 | -0.63 | 4.26E-05 | 2.79E-03 |
| ENSG00000134853 | PDGFRA | -0.63 | 4.30E-05 | 2.81E-03 |
| ENSG00000188626 | GOLGA8M | 0.63 | 4.35E-05 | 2.83E-03 |
| ENSG00000283683 | MYOCOS | 0.63 | 4.40E-05 | 2.85E-03 |
| ENSG00000182628 | SKA2 | -0.63 | 4.44E-05 | 2.87E-03 |
| ENSG00000259658 | AC027559.1 | 0.63 | 4.45E-05 | 2.87E-03 |
| ENSG00000130653 | PNPLA7 | 0.63 | 4.62E-05 | 2.95E-03 |

| Ensembl ID | Gene Name | ρ | Pval | Padj |
|---|---|---|---|---|
| ENSG00000136238 | RAC1 | -0.63 | 4.62E-05 | 2.95E-03 |
| ENSG00000235888 | AF064858.1 | 0.63 | 4.64E-05 | 2.95E-03 |
| ENSG00000166685 | COG1 | 0.63 | 4.67E-05 | 2.95E-03 |
| ENSG00000169908 | TM4SF1 | -0.63 | 4.67E-05 | 2.95E-03 |
| ENSG00000267221 | C17orf113 | 0.63 | 4.67E-05 | 2.95E-03 |
| ENSG00000255650 | FAM222A-AS1 | 0.63 | 4.68E-05 | 2.95E-03 |
| ENSG00000185352 | HS6ST3 | 0.63 | 4.70E-05 | 2.95E-03 |
| ENSG00000159307 | SCUBE1 | 0.63 | 4.71E-05 | 2.95E-03 |
| ENSG00000092020 | PPP2R3C | -0.63 | 4.76E-05 | 2.96E-03 |
| ENSG00000103769 | RAB11A | -0.63 | 4.76E-05 | 2.96E-03 |
| ENSG00000131797 | CLUHP3 | 0.63 | 4.76E-05 | 2.96E-03 |
| ENSG00000101160 | CTSZ | -0.63 | 4.81E-05 | 2.97E-03 |
| ENSG00000120509 | PDZD11 | -0.63 | 4.81E-05 | 2.97E-03 |
| ENSG00000100526 | CDKN3 | -0.63 | 5.10E-05 | 3.14E-03 |
| ENSG00000228463 | AP006222.1 | 0.63 | 5.12E-05 | 3.14E-03 |
| ENSG00000101935 | AMMECR1 | -0.63 | 5.16E-05 | 3.14E-03 |
| ENSG00000114503 | NCBP2 | -0.63 | 5.16E-05 | 3.14E-03 |
| ENSG00000167460 | TPM4 | -0.63 | 5.16E-05 | 3.14E-03 |
| ENSG00000154813 | DPH3 | -0.63 | 5.21E-05 | 3.15E-03 |
| ENSG00000156026 | MCU | -0.63 | 5.21E-05 | 3.15E-03 |
| ENSG00000137804 | NUSAP1 | -0.63 | 5.27E-05 | 3.16E-03 |
| ENSG00000198522 | GPN1 | -0.63 | 5.27E-05 | 3.16E-03 |
| ENSG00000198690 | FAN1 | 0.63 | 5.32E-05 | 3.17E-03 |
| ENSG00000198901 | PRC1 | -0.63 | 5.32E-05 | 3.17E-03 |
| ENSG00000086061 | DNAJA1 | -0.63 | 5.37E-05 | 3.20E-03 |
| ENSG00000215912 | TTC34 | 0.63 | 5.43E-05 | 3.22E-03 |
| ENSG00000143147 | GPR161 | -0.63 | 5.53E-05 | 3.26E-03 |
| ENSG00000204054 | LINC00963 | 0.63 | 5.53E-05 | 3.26E-03 |
| ENSG00000184702 | SEPT5 | 0.63 | 5.59E-05 | 3.29E-03 |
| ENSG00000172172 | MRPL13 | -0.63 | 5.65E-05 | 3.30E-03 |
| ENSG00000179387 | ELMOD2 | -0.63 | 5.65E-05 | 3.30E-03 |
| ENSG00000164418 | GRIK2 | 0.63 | 5.66E-05 | 3.30E-03 |
| ENSG00000176593 | AC008969.1 | 0.63 | 5.76E-05 | 3.34E-03 |
| ENSG00000183798 | EMILIN3 | 0.63 | 5.76E-05 | 3.34E-03 |
| ENSG00000029153 | ARNTL2 | -0.62 | 5.99E-05 | 3.45E-03 |
| ENSG00000162627 | SNX7 | -0.62 | 5.99E-05 | 3.45E-03 |
| ENSG00000077063 | CTTNBP2 | 0.62 | 6.11E-05 | 3.50E-03 |
| ENSG00000155115 | GTF3C6 | -0.62 | 6.11E-05 | 3.50E-03 |
| ENSG00000259363 | AC090825.1 | 0.62 | 6.17E-05 | 3.51E-03 |
| ENSG00000273749 | CYFIP1 | -0.62 | 6.17E-05 | 3.51E-03 |
| ENSG00000064726 | BTBD1 | -0.62 | 6.23E-05 | 3.52E-03 |
| ENSG00000137731 | FXYD2 | 0.62 | 6.23E-05 | 3.52E-03 |
| ENSG00000138160 | KIF11 | -0.62 | 6.23E-05 | 3.52E-03 |
| ENSG00000080822 | CLDND1 | -0.62 | 6.36E-05 | 3.57E-03 |
| ENSG00000261324 | AC010168.2 | 0.62 | 6.36E-05 | 3.57E-03 |

| Ensembl ID | Gene Name | ρ | Pval | Padj |
|---|---|---|---|---|
| ENSG00000029993 | HMGB3 | -0.62 | 6.42E-05 | 3.59E-03 |
| ENSG00000183520 | UTP11 | -0.62 | 6.55E-05 | 3.64E-03 |
| ENSG00000204248 | COL11A2 | 0.62 | 6.55E-05 | 3.64E-03 |
| ENSG00000112079 | STK38 | -0.62 | 6.61E-05 | 3.66E-03 |
| ENSG00000162998 | FRZB | 0.62 | 6.61E-05 | 3.66E-03 |
| ENSG00000135632 | SMYD5 | 0.62 | 6.67E-05 | 3.68E-03 |
| ENSG00000188643 | S100A16 | -0.62 | 6.67E-05 | 3.68E-03 |
| ENSG00000173692 | PSMD1 | -0.62 | 6.87E-05 | 3.77E-03 |
| ENSG00000076351 | SLC46A1 | 0.62 | 6.94E-05 | 3.79E-03 |
| ENSG00000251359 | WWC2-AS2 | 0.62 | 6.94E-05 | 3.79E-03 |
| ENSG00000101367 | MAPRE1 | -0.62 | 7.01E-05 | 3.81E-03 |
| ENSG00000187068 | C3orf70 | 0.62 | 7.01E-05 | 3.81E-03 |
| ENSG00000172757 | CFL1 | -0.62 | 7.08E-05 | 3.83E-03 |
| ENSG00000140598 | EFL1 | -0.62 | 7.15E-05 | 3.86E-03 |
| ENSG00000280339 | AP001528.3 | 0.62 | 7.24E-05 | 3.90E-03 |
| ENSG00000087302 | RTRAF | -0.62 | 7.29E-05 | 3.91E-03 |
| ENSG00000132341 | RAN | -0.62 | 7.29E-05 | 3.91E-03 |
| ENSG00000123329 | ARHGAP9 | -0.62 | 7.43E-05 | 3.96E-03 |
| ENSG00000124766 | SOX4 | -0.62 | 7.43E-05 | 3.96E-03 |
| ENSG00000182397 | DNM1P46 | 0.62 | 7.48E-05 | 3.98E-03 |
| ENSG00000101928 | MOSPD1 | -0.62 | 7.65E-05 | 4.03E-03 |
| ENSG00000108829 | LRRC59 | -0.62 | 7.65E-05 | 4.03E-03 |
| ENSG00000154734 | ADAMTS1 | -0.62 | 7.65E-05 | 4.03E-03 |
| ENSG00000115091 | ACTR3 | -0.62 | 7.72E-05 | 4.05E-03 |
| ENSG00000126001 | CEP250 | 0.62 | 7.72E-05 | 4.05E-03 |
| ENSG00000026508 | CD44 | -0.62 | 7.79E-05 | 4.07E-03 |
| ENSG00000128245 | YWHAH | -0.62 | 7.79E-05 | 4.07E-03 |
| ENSG00000224728 | IMPDH1P8 | 0.62 | 7.98E-05 | 4.15E-03 |
| ENSG00000105058 | FAM32A | -0.62 | 8.10E-05 | 4.18E-03 |
| ENSG00000205542 | TMSB4X | -0.62 | 8.10E-05 | 4.18E-03 |
| ENSG00000263412 | AC004477.1 | 0.62 | 8.10E-05 | 4.18E-03 |
| ENSG00000102226 | USP11 | 0.62 | 8.18E-05 | 4.20E-03 |
| ENSG00000176490 | DIRAS1 | -0.62 | 8.18E-05 | 4.20E-03 |
| ENSG00000124731 | TREM1 | -0.62 | 8.34E-05 | 4.27E-03 |
| ENSG00000071539 | TRIP13 | -0.61 | 8.48E-05 | 4.34E-03 |
| ENSG00000034510 | TMSB10 | -0.61 | 8.74E-05 | 4.46E-03 |
| ENSG00000175063 | UBE2C | -0.61 | 8.81E-05 | 4.47E-03 |
| ENSG00000235363 | SNRPGP10 | -0.61 | 8.81E-05 | 4.47E-03 |
| ENSG00000261959 | AC015909.3 | 0.61 | 8.98E-05 | 4.52E-03 |
| ENSG00000118894 | EEF2KMT | -0.61 | 9.00E-05 | 4.52E-03 |
| ENSG00000154096 | THY1 | -0.61 | 9.00E-05 | 4.52E-03 |
| ENSG00000188042 | ARL4C | -0.61 | 9.00E-05 | 4.52E-03 |
| ENSG00000109919 | MTCH2 | -0.61 | 9.08E-05 | 4.52E-03 |
| ENSG00000213569 | AL390039.1 | -0.61 | 9.08E-05 | 4.52E-03 |
| ENSG00000237649 | KIFC1 | -0.61 | 9.08E-05 | 4.52E-03 |

| Ensembl ID | Gene Name | ρ | Pval | Padj |
|---|---|---|---|---|
| ENSG00000101447 | FAM83D | -0.61 | 9.14E-05 | 4.52E-03 |
| ENSG00000066322 | ELOVL1 | -0.61 | 9.17E-05 | 4.52E-03 |
| ENSG00000140688 | C16orf58 | 0.61 | 9.17E-05 | 4.52E-03 |
| ENSG00000150938 | CRIM1 | 0.61 | 9.17E-05 | 4.52E-03 |
| ENSG00000236609 | ZNF853 | 0.61 | 9.17E-05 | 4.52E-03 |
| ENSG00000165480 | SKA3 | -0.61 | 9.29E-05 | 4.56E-03 |
| ENSG00000134057 | CCNB1 | -0.61 | 9.34E-05 | 4.56E-03 |
| ENSG00000163359 | COL6A3 | -0.61 | 9.34E-05 | 4.56E-03 |
| ENSG00000176974 | SHMT1 | 0.61 | 9.34E-05 | 4.56E-03 |
| ENSG00000146731 | CCT6A | -0.61 | 9.43E-05 | 4.58E-03 |
| ENSG00000186866 | POFUT2 | 0.61 | 9.43E-05 | 4.58E-03 |
| ENSG00000137845 | ADAM10 | -0.61 | 9.52E-05 | 4.61E-03 |
| ENSG00000126785 | RHOJ | 0.61 | 9.61E-05 | 4.62E-03 |
| ENSG00000177889 | UBE2N | -0.61 | 9.61E-05 | 4.62E-03 |
| ENSG00000215414 | PSMA6P1 | -0.61 | 9.61E-05 | 4.62E-03 |
| ENSG00000109079 | TNFAIP1 | -0.61 | 9.70E-05 | 4.62E-03 |
| ENSG00000115468 | EFHD1 | 0.61 | 9.70E-05 | 4.62E-03 |
| ENSG00000156042 | CFAP70 | 0.61 | 9.70E-05 | 4.62E-03 |
| ENSG00000184752 | NDUFA12 | -0.61 | 9.70E-05 | 4.62E-03 |
| ENSG00000171208 | NETO2 | -0.61 | 9.74E-05 | 4.63E-03 |
| ENSG00000041357 | PSMA4 | -0.61 | 9.79E-05 | 4.64E-03 |
| ENSG00000121671 | CRY2 | 0.61 | 9.89E-05 | 4.66E-03 |
| ENSG00000253320 | AZIN1-AS1 | 0.61 | 9.89E-05 | 4.66E-03 |
| ENSG00000060656 | PTPRU | 0.61 | 9.98E-05 | 4.67E-03 |
| ENSG00000108518 | PFN1 | -0.61 | 9.98E-05 | 4.67E-03 |
| ENSG00000116489 | CAPZA1 | -0.61 | 9.98E-05 | 4.67E-03 |
| ENSG00000132359 | RAP1GAP2 | 0.61 | 1.01E-04 | 4.67E-03 |
| ENSG00000137807 | KIF23 | -0.61 | 1.01E-04 | 4.67E-03 |
| ENSG00000169184 | MN1 | 0.61 | 1.01E-04 | 4.67E-03 |
| ENSG00000228109 | MELTF-AS1 | 0.61 | 1.01E-04 | 4.67E-03 |
| ENSG00000257696 | AC010203.2 | 0.61 | 1.01E-04 | 4.68E-03 |
| ENSG00000145241 | CENPC | 0.61 | 1.03E-04 | 4.73E-03 |
| ENSG00000236548 | RNF217-AS1 | 0.61 | 1.03E-04 | 4.73E-03 |
| ENSG00000074621 | SLC24A1 | 0.61 | 1.04E-04 | 4.75E-03 |
| ENSG00000143365 | RORC | 0.61 | 1.04E-04 | 4.75E-03 |
| ENSG00000005513 | SOX8 | 0.61 | 1.06E-04 | 4.80E-03 |
| ENSG00000021574 | SPAST | -0.61 | 1.06E-04 | 4.80E-03 |
| ENSG00000116701 | NCF2 | -0.61 | 1.06E-04 | 4.80E-03 |
| ENSG00000123609 | NMI | -0.61 | 1.06E-04 | 4.80E-03 |
| ENSG00000154589 | LY96 | -0.61 | 1.06E-04 | 4.81E-03 |
| ENSG00000198547 | C20orf203 | 0.61 | 1.07E-04 | 4.82E-03 |
| ENSG00000005022 | SLC25A5 | -0.61 | 1.08E-04 | 4.84E-03 |
| ENSG00000168282 | MGAT2 | -0.61 | 1.08E-04 | 4.84E-03 |
| ENSG00000105088 | OLFM2 | -0.61 | 1.08E-04 | 4.86E-03 |
| ENSG00000267640 | AC016582.3 | 0.61 | 1.09E-04 | 4.87E-03 |

| Ensembl ID | Gene Name | ρ | Pval | Padj |
|---|---|---|---|---|
| ENSG00000133226 | SRRM1 | 0.61 | 1.10E-04 | 4.90E-03 |
| ENSG00000171476 | HOPX | -0.61 | 1.10E-04 | 4.92E-03 |
| ENSG00000111319 | SCNN1A | 0.61 | 1.11E-04 | 4.92E-03 |
| ENSG00000134531 | EMP1 | -0.61 | 1.11E-04 | 4.92E-03 |
| ENSG00000114738 | MAPKAPK3 | -0.61 | 1.12E-04 | 4.95E-03 |
| ENSG00000135835 | KIAA1614 | 0.61 | 1.13E-04 | 4.99E-03 |
| ENSG00000103091 | WDR59 | 0.61 | 1.14E-04 | 5.01E-03 |
| ENSG00000131711 | MAP1B | -0.61 | 1.14E-04 | 5.01E-03 |
| ENSG00000104549 | SQLE | -0.61 | 1.15E-04 | 5.02E-03 |
| ENSG00000176700 | SCAND2P | 0.61 | 1.15E-04 | 5.02E-03 |
| ENSG00000260563 | AC132872.1 | 0.61 | 1.15E-04 | 5.02E-03 |
| ENSG00000175352 | NRIP3 | -0.61 | 1.16E-04 | 5.05E-03 |
| ENSG00000233937 | AC008443.1 | 0.61 | 1.16E-04 | 5.05E-03 |
| ENSG00000114573 | ATP6V1A | -0.61 | 1.17E-04 | 5.08E-03 |
| ENSG00000003756 | RBM5 | 0.61 | 1.18E-04 | 5.12E-03 |
| ENSG00000142657 | PGD | -0.60 | 1.19E-04 | 5.16E-03 |
| ENSG00000138166 | DUSP5 | -0.60 | 1.20E-04 | 5.17E-03 |
| ENSG00000184208 | C22orf46 | 0.60 | 1.20E-04 | 5.17E-03 |
| ENSG00000253626 | EIF5AL1 | -0.60 | 1.20E-04 | 5.17E-03 |
| ENSG00000129465 | RIPK3 | -0.60 | 1.22E-04 | 5.24E-03 |
| ENSG00000263072 | ZNF213-AS1 | 0.60 | 1.22E-04 | 5.24E-03 |
| ENSG00000197768 | STPG3 | 0.60 | 1.25E-04 | 5.35E-03 |
| ENSG00000023909 | GCLM | -0.60 | 1.26E-04 | 5.36E-03 |
| ENSG00000175287 | PHYHD1 | 0.60 | 1.26E-04 | 5.36E-03 |
| ENSG00000038945 | MSR1 | -0.60 | 1.26E-04 | 5.36E-03 |
| ENSG00000169016 | E2F6 | -0.60 | 1.27E-04 | 5.38E-03 |
| ENSG00000226578 | AL132657.1 | 0.60 | 1.28E-04 | 5.42E-03 |
| ENSG00000279900 | AP001767.4 | 0.60 | 1.29E-04 | 5.45E-03 |
| ENSG00000175782 | SLC35E3 | 0.60 | 1.31E-04 | 5.50E-03 |
| ENSG00000253848 | AC010834.2 | 0.60 | 1.31E-04 | 5.50E-03 |
| ENSG00000156471 | PTDSS1 | -0.60 | 1.32E-04 | 5.52E-03 |
| ENSG00000136810 | TXN | -0.60 | 1.33E-04 | 5.54E-03 |
| ENSG00000161800 | RACGAP1 | -0.60 | 1.33E-04 | 5.54E-03 |
| ENSG00000189056 | RELN | 0.60 | 1.33E-04 | 5.54E-03 |
| ENSG00000113407 | TARS | -0.60 | 1.34E-04 | 5.56E-03 |
| ENSG00000151239 | TWF1 | -0.60 | 1.34E-04 | 5.56E-03 |
| ENSG00000204623 | ZNRD1ASP | 0.60 | 1.34E-04 | 5.56E-03 |
| ENSG00000162241 | SLC25A45 | 0.60 | 1.35E-04 | 5.59E-03 |
| ENSG00000120915 | EPHX2 | 0.60 | 1.37E-04 | 5.60E-03 |
| ENSG00000153291 | SLC25A27 | 0.60 | 1.37E-04 | 5.60E-03 |
| ENSG00000197555 | SIPA1L1 | 0.60 | 1.37E-04 | 5.60E-03 |
| ENSG00000245694 | CRNDE | -0.60 | 1.37E-04 | 5.60E-03 |
| ENSG00000205744 | DENND1C | -0.60 | 1.39E-04 | 5.68E-03 |
| ENSG00000110955 | ATP5F1B | -0.60 | 1.40E-04 | 5.71E-03 |
| ENSG00000122591 | FAM126A | -0.60 | 1.40E-04 | 5.71E-03 |

| Ensembl ID | Gene Name | ρ | Pval | Padj |
|---|---|---|---|---|
| ENSG00000171241 | SHCBP1 | -0.60 | 1.40E-04 | 5.71E-03 |
| ENSG00000111711 | GOLT1B | -0.60 | 1.42E-04 | 5.74E-03 |
| ENSG00000244617 | ASPRV1 | 0.60 | 1.42E-04 | 5.74E-03 |
| ENSG00000005961 | ITGA2B | 0.60 | 1.43E-04 | 5.74E-03 |
| ENSG00000143198 | MGST3 | -0.60 | 1.43E-04 | 5.74E-03 |
| ENSG00000143845 | ETNK2 | 0.60 | 1.43E-04 | 5.74E-03 |
| ENSG00000163636 | PSMD6 | -0.60 | 1.43E-04 | 5.74E-03 |
| ENSG00000137497 | NUMA1 | 0.60 | 1.44E-04 | 5.78E-03 |
| ENSG00000227053 | AC105446.1 | -0.60 | 1.45E-04 | 5.80E-03 |
| ENSG00000007402 | CACNA2D2 | 0.60 | 1.46E-04 | 5.81E-03 |
| ENSG00000275367 | AC092111.1 | 0.60 | 1.47E-04 | 5.85E-03 |
| ENSG00000138670 | RASGEF1B | -0.60 | 1.48E-04 | 5.88E-03 |
| ENSG00000181638 | ZFP41 | 0.60 | 1.48E-04 | 5.88E-03 |
| ENSG00000196230 | TUBB | -0.60 | 1.50E-04 | 5.91E-03 |
| ENSG00000214194 | SMIM30 | -0.60 | 1.50E-04 | 5.91E-03 |
| ENSG00000272692 | AC010997.4 | 0.60 | 1.50E-04 | 5.92E-03 |
| ENSG00000099869 | IGF2-AS | 0.60 | 1.52E-04 | 5.94E-03 |
| ENSG00000012660 | ELOVL5 | -0.60 | 1.52E-04 | 5.94E-03 |
| ENSG00000137103 | TMEM8B | 0.60 | 1.52E-04 | 5.94E-03 |
| ENSG00000139734 | DIAPH3 | -0.60 | 1.52E-04 | 5.94E-03 |
| ENSG00000173918 | C1QTNF1 | -0.60 | 1.52E-04 | 5.94E-03 |
| ENSG00000232445 | AC006329.1 | 0.60 | 1.52E-04 | 5.94E-03 |
| ENSG00000006377 | DLX6 | 0.60 | 1.54E-04 | 5.97E-03 |
| ENSG00000162851 | TFB2M | -0.60 | 1.54E-04 | 5.97E-03 |
| ENSG00000181751 | C5orf30 | -0.60 | 1.55E-04 | 6.01E-03 |
| ENSG00000111348 | ARHGDIB | -0.60 | 1.56E-04 | 6.04E-03 |
| ENSG00000163902 | RPN1 | -0.60 | 1.56E-04 | 6.04E-03 |
| ENSG00000160180 | TFF3 | -0.60 | 1.57E-04 | 6.04E-03 |
| ENSG00000136827 | TOR1A | -0.60 | 1.58E-04 | 6.04E-03 |
| ENSG00000178053 | MLF1 | -0.60 | 1.58E-04 | 6.04E-03 |
| ENSG00000272501 | AL662844.4 | 0.60 | 1.58E-04 | 6.04E-03 |
| ENSG00000089685 | BIRC5 | -0.60 | 1.59E-04 | 6.04E-03 |
| ENSG00000168036 | CTNNB1 | -0.60 | 1.59E-04 | 6.04E-03 |
| ENSG00000170779 | CDCA4 | -0.60 | 1.59E-04 | 6.04E-03 |
| ENSG00000078098 | FAP | -0.60 | 1.61E-04 | 6.04E-03 |
| ENSG00000119729 | RHOQ | -0.60 | 1.61E-04 | 6.04E-03 |
| ENSG00000146670 | CDCA5 | -0.60 | 1.61E-04 | 6.04E-03 |
| ENSG00000149218 | ENDOD1 | -0.60 | 1.61E-04 | 6.04E-03 |
| ENSG00000170919 | TPT1-AS1 | 0.60 | 1.61E-04 | 6.04E-03 |
| ENSG00000206527 | HACD2 | -0.60 | 1.61E-04 | 6.04E-03 |
| ENSG00000241288 | LINC02614 | 0.60 | 1.61E-04 | 6.04E-03 |
| ENSG00000107954 | NEURL1 | 0.60 | 1.62E-04 | 6.07E-03 |
| ENSG00000244879 | GABPB1-AS1 | 0.60 | 1.62E-04 | 6.07E-03 |
| ENSG00000166451 | CENPN | -0.59 | 1.64E-04 | 6.12E-03 |
| ENSG00000185070 | FLRT2 | -0.59 | 1.65E-04 | 6.16E-03 |

| Ensembl ID | Gene Name | ρ | Pval | Padj |
|---|---|---|---|---|
| ENSG00000159899 | NPR2 | 0.59 | 1.66E-04 | 6.20E-03 |
| ENSG00000005893 | LAMP2 | -0.59 | 1.68E-04 | 6.25E-03 |
| ENSG00000115963 | RND3 | -0.59 | 1.69E-04 | 6.29E-03 |
| ENSG00000112305 | SMAP1 | -0.59 | 1.71E-04 | 6.30E-03 |
| ENSG00000114126 | TFDP2 | 0.59 | 1.71E-04 | 6.30E-03 |
| ENSG00000175130 | MARCKSL1 | -0.59 | 1.71E-04 | 6.30E-03 |
| ENSG00000198932 | GPRASP1 | 0.59 | 1.71E-04 | 6.30E-03 |
| ENSG00000005059 | MCUB | -0.59 | 1.74E-04 | 6.38E-03 |
| ENSG00000068078 | FGFR3 | 0.59 | 1.74E-04 | 6.38E-03 |
| ENSG00000204071 | TCEAL6 | 0.59 | 1.74E-04 | 6.38E-03 |
| ENSG00000157456 | CCNB2 | -0.59 | 1.75E-04 | 6.39E-03 |
| ENSG00000142621 | FHAD1 | -0.59 | 1.76E-04 | 6.42E-03 |
| ENSG00000179399 | GPC5 | 0.59 | 1.77E-04 | 6.46E-03 |
| ENSG00000081818 | PCDHB4 | 0.59 | 1.79E-04 | 6.46E-03 |
| ENSG00000102218 | RP2 | -0.59 | 1.79E-04 | 6.46E-03 |
| ENSG00000116586 | LAMTOR2 | -0.59 | 1.79E-04 | 6.46E-03 |
| ENSG00000128536 | CDHR3 | 0.59 | 1.79E-04 | 6.46E-03 |
| ENSG00000140931 | CMTM3 | -0.59 | 1.79E-04 | 6.46E-03 |
| ENSG00000134376 | CRB1 | 0.59 | 1.79E-04 | 6.47E-03 |
| ENSG00000261713 | SSTR5-AS1 | 0.59 | 1.80E-04 | 6.49E-03 |
| ENSG00000166411 | IDH3A | -0.59 | 1.82E-04 | 6.54E-03 |
| ENSG00000156970 | BUB1B | -0.59 | 1.82E-04 | 6.54E-03 |
| ENSG00000253608 | AC022915.2 | 0.59 | 1.83E-04 | 6.54E-03 |
| ENSG00000256745 | AP002784.2 | -0.59 | 1.84E-04 | 6.56E-03 |
| ENSG00000021762 | OSBPL5 | 0.59 | 1.85E-04 | 6.59E-03 |
| ENSG00000073910 | FRY | 0.59 | 1.85E-04 | 6.59E-03 |
| ENSG00000120992 | LYPLA1 | -0.59 | 1.88E-04 | 6.70E-03 |
| ENSG00000188760 | TMEM198 | 0.59 | 1.90E-04 | 6.75E-03 |
| ENSG00000113361 | CDH6 | -0.59 | 1.91E-04 | 6.75E-03 |
| ENSG00000136240 | KDELR2 | -0.59 | 1.92E-04 | 6.75E-03 |
| ENSG00000163444 | TMEM183A | -0.59 | 1.92E-04 | 6.75E-03 |
| ENSG00000168374 | ARF4 | -0.59 | 1.92E-04 | 6.75E-03 |
| ENSG00000277715 | AC079174.2 | 0.59 | 1.92E-04 | 6.75E-03 |
| ENSG00000084733 | RAB10 | -0.59 | 1.93E-04 | 6.76E-03 |
| ENSG00000164611 | PTTG1 | -0.59 | 1.93E-04 | 6.76E-03 |
| ENSG00000204386 | NEU1 | -0.59 | 1.93E-04 | 6.76E-03 |
| ENSG00000271816 | AC022400.5 | 0.59 | 1.94E-04 | 6.76E-03 |
| ENSG00000171848 | RRM2 | -0.59 | 1.97E-04 | 6.86E-03 |
| ENSG00000104312 | RIPK2 | -0.59 | 1.99E-04 | 6.91E-03 |
| ENSG00000120705 | ETF1 | -0.59 | 2.00E-04 | 6.95E-03 |
| ENSG00000171862 | PTEN | -0.59 | 2.00E-04 | 6.95E-03 |
| ENSG00000103591 | AAGAB | -0.59 | 2.04E-04 | 7.06E-03 |
| ENSG00000138180 | CEP55 | -0.59 | 2.05E-04 | 7.06E-03 |
| ENSG00000092841 | MYL6 | -0.59 | 2.06E-04 | 7.06E-03 |
| ENSG00000125037 | EMC3 | -0.59 | 2.06E-04 | 7.06E-03 |

| Ensembl ID | Gene Name | ρ | Pval | Padj |
|---|---|---|---|---|
| ENSG00000130294 | KIF1A | 0.59 | 2.06E-04 | 7.06E-03 |
| ENSG00000163468 | CCT3 | -0.59 | 2.06E-04 | 7.06E-03 |
| ENSG00000143753 | DEGS1 | -0.59 | 2.07E-04 | 7.07E-03 |
| ENSG00000164332 | UBLCP1 | -0.59 | 2.07E-04 | 7.07E-03 |
| ENSG00000165323 | FAT3 | -0.59 | 2.07E-04 | 7.07E-03 |
| ENSG00000229180 | AC006001.3 | 0.59 | 2.07E-04 | 7.07E-03 |
| ENSG00000064763 | FAR2 | -0.59 | 2.09E-04 | 7.09E-03 |
| ENSG00000110013 | SIAE | -0.59 | 2.09E-04 | 7.09E-03 |
| ENSG00000142227 | EMP3 | -0.59 | 2.09E-04 | 7.09E-03 |
| ENSG00000158286 | RNF207 | 0.59 | 2.09E-04 | 7.09E-03 |
| ENSG00000142599 | RERE | 0.59 | 2.11E-04 | 7.14E-03 |
| ENSG00000159713 | TPPP3 | -0.59 | 2.13E-04 | 7.18E-03 |
| ENSG00000250343 | STK32A-AS1 | 0.59 | 2.13E-04 | 7.18E-03 |
| ENSG00000271646 | AC099343.3 | 0.59 | 2.15E-04 | 7.23E-03 |
| ENSG00000143621 | ILF2 | -0.59 | 2.17E-04 | 7.26E-03 |
| ENSG00000144843 | ADPRH | -0.59 | 2.17E-04 | 7.26E-03 |
| ENSG00000118680 | MYL12B | -0.59 | 2.19E-04 | 7.29E-03 |
| ENSG00000133619 | KRBA1 | 0.59 | 2.19E-04 | 7.29E-03 |
| ENSG00000142197 | DOP1B | -0.59 | 2.19E-04 | 7.29E-03 |
| ENSG00000133884 | DPF2 | 0.59 | 2.22E-04 | 7.37E-03 |
| ENSG00000151917 | BEND6 | -0.59 | 2.22E-04 | 7.37E-03 |
| ENSG00000176723 | ZNF843 | 0.59 | 2.22E-04 | 7.37E-03 |
| ENSG00000186298 | PPP1CC | -0.59 | 2.22E-04 | 7.37E-03 |
| ENSG00000276291 | FRG1HP | 0.58 | 2.24E-04 | 7.42E-03 |
| ENSG00000071054 | MAP4K4 | -0.58 | 2.26E-04 | 7.44E-03 |
| ENSG00000078295 | ADCY2 | 0.58 | 2.26E-04 | 7.44E-03 |
| ENSG00000151276 | MAGI1 | 0.58 | 2.26E-04 | 7.44E-03 |
| ENSG00000074410 | CA12 | -0.58 | 2.26E-04 | 7.44E-03 |
| ENSG00000255471 | AP001528.2 | 0.58 | 2.28E-04 | 7.46E-03 |
| ENSG00000137745 | MMP13 | -0.58 | 2.28E-04 | 7.46E-03 |
| ENSG00000183155 | RABIF | -0.58 | 2.28E-04 | 7.46E-03 |
| ENSG00000005007 | UPF1 | 0.58 | 2.30E-04 | 7.50E-03 |
| ENSG00000091483 | FH | -0.58 | 2.30E-04 | 7.50E-03 |
| ENSG00000234883 | MIR155HG | -0.58 | 2.31E-04 | 7.53E-03 |
| ENSG00000088808 | PPP1R13B | 0.58 | 2.32E-04 | 7.53E-03 |
| ENSG00000131844 | MCCC2 | 0.58 | 2.32E-04 | 7.53E-03 |
| ENSG00000106006 | HOXA6 | 0.58 | 2.36E-04 | 7.65E-03 |
| ENSG00000005302 | MSL3 | -0.58 | 2.38E-04 | 7.69E-03 |
| ENSG00000141956 | PRDM15 | 0.58 | 2.38E-04 | 7.69E-03 |
| ENSG00000142230 | SAE1 | -0.58 | 2.40E-04 | 7.73E-03 |
| ENSG00000260807 | AC009041.2 | 0.58 | 2.40E-04 | 7.73E-03 |
| ENSG00000132874 | SLC14A2 | 0.58 | 2.40E-04 | 7.73E-03 |
| ENSG00000187514 | PTMA | -0.58 | 2.42E-04 | 7.74E-03 |
| ENSG00000196639 | HRH1 | -0.58 | 2.42E-04 | 7.74E-03 |
| ENSG00000225205 | AC078883.1 | 0.58 | 2.42E-04 | 7.74E-03 |

| Ensembl ID | Gene Name | ρ | Pval | Padj |
|---|---|---|---|---|
| ENSG00000234062 | AL390879.1 | 0.58 | 2.42E-04 | 7.74E-03 |
| ENSG00000274536 | AL034397.3 | -0.58 | 2.43E-04 | 7.74E-03 |
| ENSG00000216657 | GLRX3P2 | -0.58 | 2.44E-04 | 7.78E-03 |
| ENSG00000024526 | DEPDC1 | -0.58 | 2.48E-04 | 7.87E-03 |
| ENSG00000114735 | HEMK1 | 0.58 | 2.49E-04 | 7.87E-03 |
| ENSG00000128342 | LIF | -0.58 | 2.49E-04 | 7.87E-03 |
| ENSG00000074181 | NOTCH3 | -0.58 | 2.53E-04 | 7.92E-03 |
| ENSG00000122641 | INHBA | -0.58 | 2.53E-04 | 7.92E-03 |
| ENSG00000140367 | UBE2Q2 | -0.58 | 2.53E-04 | 7.92E-03 |
| ENSG00000172992 | DCAKD | 0.58 | 2.53E-04 | 7.92E-03 |
| ENSG00000173085 | COQ2 | -0.58 | 2.53E-04 | 7.92E-03 |
| ENSG00000213366 | GSTM2 | 0.58 | 2.53E-04 | 7.92E-03 |
| ENSG00000275765 | AC091982.3 | 0.58 | 2.53E-04 | 7.92E-03 |
| ENSG00000237940 | LINC01238 | 0.58 | 2.54E-04 | 7.92E-03 |
| ENSG00000160867 | FGFR4 | 0.58 | 2.55E-04 | 7.92E-03 |
| ENSG00000162694 | EXTL2 | -0.58 | 2.55E-04 | 7.92E-03 |
| ENSG00000163617 | CCDC191 | 0.58 | 2.55E-04 | 7.92E-03 |
| ENSG00000170837 | GPR27 | 0.58 | 2.55E-04 | 7.92E-03 |
| ENSG00000136689 | IL1RN | -0.58 | 2.55E-04 | 7.92E-03 |
| ENSG00000186469 | GNG2 | -0.58 | 2.57E-04 | 7.97E-03 |
| ENSG00000196072 | BLOC1S2 | -0.58 | 2.59E-04 | 8.03E-03 |
| ENSG00000101265 | RASSF2 | -0.58 | 2.60E-04 | 8.04E-03 |
| ENSG00000111206 | FOXM1 | -0.58 | 2.62E-04 | 8.06E-03 |
| ENSG00000171552 | BCL2L1 | -0.58 | 2.62E-04 | 8.06E-03 |
| ENSG00000170579 | DLGAP1 | 0.58 | 2.62E-04 | 8.06E-03 |
| ENSG00000239704 | CDRT4 | 0.58 | 2.62E-04 | 8.06E-03 |
| ENSG00000092820 | EZR | -0.58 | 2.64E-04 | 8.07E-03 |
| ENSG00000119335 | SET | -0.58 | 2.64E-04 | 8.07E-03 |
| ENSG00000179930 | ZNF648 | 0.58 | 2.64E-04 | 8.07E-03 |
| ENSG00000102313 | ITIH6 | 0.58 | 2.66E-04 | 8.11E-03 |
| ENSG00000197162 | ZNF785 | 0.58 | 2.66E-04 | 8.11E-03 |
| ENSG00000105849 | TWISTNB | -0.58 | 2.68E-04 | 8.15E-03 |
| ENSG00000133318 | RTN3 | -0.58 | 2.68E-04 | 8.15E-03 |
| ENSG00000105479 | CCDC114 | 0.58 | 2.69E-04 | 8.17E-03 |
| ENSG00000163606 | CD200R1 | -0.58 | 2.70E-04 | 8.18E-03 |
| ENSG00000186056 | MATN1-AS1 | 0.58 | 2.71E-04 | 8.19E-03 |
| ENSG00000103966 | EHD4 | -0.58 | 2.73E-04 | 8.24E-03 |
| ENSG00000058799 | YIPF1 | -0.58 | 2.75E-04 | 8.29E-03 |
| ENSG00000164253 | WDR41 | -0.58 | 2.75E-04 | 8.29E-03 |
| ENSG00000116954 | RRAGC | -0.58 | 2.80E-04 | 8.40E-03 |
| ENSG00000228486 | C2orf92 | 0.58 | 2.80E-04 | 8.40E-03 |
| ENSG00000137404 | NRM | -0.58 | 2.82E-04 | 8.46E-03 |
| ENSG00000076650 | GPATCH1 | 0.58 | 2.85E-04 | 8.47E-03 |
| ENSG00000143369 | ECM1 | -0.58 | 2.85E-04 | 8.47E-03 |
| ENSG00000154845 | PPP4R1 | -0.58 | 2.85E-04 | 8.47E-03 |

| Ensembl ID | Gene Name | ρ | Pval | Padj |
|---|---|---|---|---|
| ENSG00000156273 | BACH1 | -0.58 | 2.85E-04 | 8.47E-03 |
| ENSG00000178093 | TSSK6 | 0.58 | 2.85E-04 | 8.47E-03 |
| ENSG00000250790 | AC127070.2 | 0.58 | 2.87E-04 | 8.52E-03 |
| ENSG00000157551 | KCNJ15 | -0.58 | 2.87E-04 | 8.52E-03 |
| ENSG00000133985 | TTC9 | -0.58 | 2.90E-04 | 8.56E-03 |
| ENSG00000141469 | SLC14A1 | 0.58 | 2.90E-04 | 8.56E-03 |
| ENSG00000253161 | LINC01605 | -0.58 | 2.91E-04 | 8.58E-03 |
| ENSG00000120457 | KCNJ5 | -0.58 | 2.93E-04 | 8.63E-03 |
| ENSG00000090266 | NDUFB2 | -0.58 | 2.94E-04 | 8.65E-03 |
| ENSG00000184900 | SUMO3 | -0.58 | 2.94E-04 | 8.65E-03 |
| ENSG00000266835 | GAPLINC | -0.58 | 2.95E-04 | 8.65E-03 |
| ENSG00000095585 | BLNK | -0.58 | 2.96E-04 | 8.67E-03 |
| ENSG00000114978 | MOB1A | -0.58 | 2.97E-04 | 8.67E-03 |
| ENSG00000169607 | CKAP2L | -0.58 | 2.97E-04 | 8.67E-03 |
| ENSG00000179051 | RCC2 | -0.58 | 2.97E-04 | 8.67E-03 |
| ENSG00000114251 | WNT5A | -0.58 | 2.99E-04 | 8.68E-03 |
| ENSG00000075624 | ACTB | -0.58 | 2.99E-04 | 8.68E-03 |
| ENSG00000139921 | TMX1 | -0.58 | 2.99E-04 | 8.68E-03 |
| ENSG00000146373 | RNF217 | 0.58 | 2.99E-04 | 8.68E-03 |
| ENSG00000149582 | TMEM25 | 0.58 | 2.99E-04 | 8.68E-03 |
| ENSG00000177082 | WDR73 | 0.58 | 3.02E-04 | 8.74E-03 |
| ENSG00000007968 | E2F2 | -0.57 | 3.04E-04 | 8.74E-03 |
| ENSG00000115556 | PLCD4 | 0.57 | 3.05E-04 | 8.74E-03 |
| ENSG00000132963 | POMP | -0.57 | 3.05E-04 | 8.74E-03 |
| ENSG00000184205 | TSPYL2 | 0.57 | 3.05E-04 | 8.74E-03 |
| ENSG00000264235 | AP005329.1 | -0.57 | 3.06E-04 | 8.74E-03 |
| ENSG00000128595 | CALU | -0.57 | 3.07E-04 | 8.74E-03 |
| ENSG00000147130 | ZMYM3 | 0.57 | 3.07E-04 | 8.74E-03 |
| ENSG00000169083 | AR | 0.57 | 3.07E-04 | 8.74E-03 |
| ENSG00000174021 | GNG5 | -0.57 | 3.07E-04 | 8.74E-03 |
| ENSG00000254860 | TMEM9B-AS1 | 0.57 | 3.07E-04 | 8.74E-03 |
| ENSG00000282057 | AC092807.3 | 0.57 | 3.07E-04 | 8.74E-03 |
| ENSG00000108788 | MLX | -0.57 | 3.10E-04 | 8.78E-03 |
| ENSG00000185267 | CDNF | 0.57 | 3.10E-04 | 8.78E-03 |
| ENSG00000235001 | EIF4A1P2 | -0.57 | 3.10E-04 | 8.78E-03 |
| ENSG00000065029 | ZNF76 | 0.57 | 3.12E-04 | 8.83E-03 |
| ENSG00000183718 | TRIM52 | 0.57 | 3.12E-04 | 8.83E-03 |
| ENSG00000279620 | AC099494.3 | 0.57 | 3.14E-04 | 8.84E-03 |
| ENSG00000132478 | UNK | 0.57 | 3.15E-04 | 8.84E-03 |
| ENSG00000163661 | PTX3 | -0.57 | 3.15E-04 | 8.84E-03 |
| ENSG00000168268 | NT5DC2 | -0.57 | 3.15E-04 | 8.84E-03 |
| ENSG00000197498 | RPF2 | -0.57 | 3.15E-04 | 8.84E-03 |
| ENSG00000224934 | AL391684.1 | 0.57 | 3.15E-04 | 8.84E-03 |
| ENSG00000117410 | ATP6V0B | -0.57 | 3.17E-04 | 8.86E-03 |
| ENSG00000162836 | ACP6 | 0.57 | 3.17E-04 | 8.86E-03 |

| Ensembl ID | Gene Name | ρ | Pval | Padj |
|---|---|---|---|---|
| ENSG00000181798 | LINC00471 | 0.57 | 3.17E-04 | 8.86E-03 |
| ENSG00000103855 | CD276 | -0.57 | 3.20E-04 | 8.89E-03 |
| ENSG00000132879 | FBXO44 | 0.57 | 3.20E-04 | 8.89E-03 |
| ENSG00000169554 | ZEB2 | -0.57 | 3.20E-04 | 8.89E-03 |
| ENSG00000173175 | ADCY5 | 0.57 | 3.20E-04 | 8.89E-03 |
| ENSG00000163013 | FBXO41 | 0.57 | 3.23E-04 | 8.93E-03 |
| ENSG00000165389 | SPTSSA | -0.57 | 3.23E-04 | 8.93E-03 |
| ENSG00000197147 | LRRC8B | -0.57 | 3.23E-04 | 8.93E-03 |
| ENSG00000261720 | AC009041.3 | 0.57 | 3.23E-04 | 8.93E-03 |
| ENSG00000088882 | CPXM1 | -0.57 | 3.24E-04 | 8.94E-03 |
| ENSG00000185432 | METTL7A | 0.57 | 3.25E-04 | 8.96E-03 |
| ENSG00000108953 | YWHAE | -0.57 | 3.31E-04 | 9.09E-03 |
| ENSG00000113387 | SUB1 | -0.57 | 3.31E-04 | 9.09E-03 |
| ENSG00000065911 | MTHFD2 | -0.57 | 3.34E-04 | 9.13E-03 |
| ENSG00000093000 | NUP50 | -0.57 | 3.34E-04 | 9.13E-03 |
| ENSG00000128294 | TPST2 | -0.57 | 3.34E-04 | 9.13E-03 |
| ENSG00000132005 | RFX1 | 0.57 | 3.36E-04 | 9.15E-03 |
| ENSG00000166949 | SMAD3 | 0.57 | 3.36E-04 | 9.15E-03 |
| ENSG00000177868 | SVBP | -0.57 | 3.36E-04 | 9.15E-03 |
| ENSG00000215045 | GRID2IP | 0.57 | 3.36E-04 | 9.15E-03 |
| ENSG00000158113 | LRRC43 | 0.57 | 3.42E-04 | 9.29E-03 |
| ENSG00000137819 | PAQR5 | -0.57 | 3.43E-04 | 9.30E-03 |
| ENSG00000167992 | VWCE | 0.57 | 3.45E-04 | 9.34E-03 |
| ENSG00000177951 | BET1L | 0.57 | 3.48E-04 | 9.41E-03 |
| ENSG00000069849 | ATP1B3 | -0.57 | 3.51E-04 | 9.43E-03 |
| ENSG00000124172 | ATP5F1E | -0.57 | 3.51E-04 | 9.43E-03 |
| ENSG00000157637 | SLC38A10 | 0.57 | 3.51E-04 | 9.43E-03 |
| ENSG00000179409 | GEMIN4 | 0.57 | 3.51E-04 | 9.43E-03 |
| ENSG00000163808 | KIF15 | -0.57 | 3.53E-04 | 9.44E-03 |
| ENSG00000115514 | TXNDC9 | -0.57 | 3.54E-04 | 9.44E-03 |
| ENSG00000137573 | SULF1 | -0.57 | 3.54E-04 | 9.44E-03 |
| ENSG00000166436 | TRIM66 | 0.57 | 3.54E-04 | 9.44E-03 |
| ENSG00000185345 | PRKN | 0.57 | 3.54E-04 | 9.44E-03 |
| ENSG00000197965 | MPZL1 | -0.57 | 3.54E-04 | 9.44E-03 |
| ENSG00000128944 | KNSTRN | -0.57 | 3.56E-04 | 9.48E-03 |
| ENSG00000169609 | C15orf40 | 0.57 | 3.56E-04 | 9.48E-03 |
| ENSG00000211448 | DIO2 | -0.57 | 3.56E-04 | 9.48E-03 |
| ENSG00000112799 | LY86 | -0.57 | 3.59E-04 | 9.53E-03 |
| ENSG00000066468 | FGFR2 | 0.57 | 3.59E-04 | 9.53E-03 |
| ENSG00000066697 | MSANTD3 | -0.57 | 3.62E-04 | 9.60E-03 |
| ENSG00000260992 | DOCK9-DT | 0.57 | 3.64E-04 | 9.62E-03 |
| ENSG00000018510 | AGPS | -0.57 | 3.65E-04 | 9.64E-03 |
| ENSG00000143933 | CALM2 | -0.57 | 3.65E-04 | 9.64E-03 |
| ENSG00000152270 | PDE3B | 0.57 | 3.68E-04 | 9.69E-03 |
| ENSG00000241014 | AC114490.1 | 0.57 | 3.68E-04 | 9.69E-03 |

| Ensembl ID | Gene Name | ρ | Pval | Padj |
|---|---|---|---|---|
| ENSG00000196611 | MMP1 | -0.57 | 3.71E-04 | 9.73E-03 |
| ENSG00000135631 | RAB11FIP5 | 0.57 | 3.71E-04 | 9.73E-03 |
| ENSG00000172315 | TP53RK | -0.57 | 3.71E-04 | 9.73E-03 |
| ENSG00000133703 | KRAS | -0.57 | 3.74E-04 | 9.79E-03 |
| ENSG00000163430 | FSTL1 | -0.57 | 3.74E-04 | 9.79E-03 |
| ENSG00000169490 | TM2D2 | -0.57 | 3.77E-04 | 9.85E-03 |
| ENSG00000152117 | AC073869.1 | 0.57 | 3.84E-04 | 9.98E-03 |
| ENSG00000185651 | UBE2L3 | -0.57 | 3.84E-04 | 9.98E-03 |
| ENSG00000235815 | AL136099.1 | 0.57 | 3.84E-04 | 9.98E-03 |
| ENSG00000057608 | GDI2 | -0.57 | 3.87E-04 | 1.00E-02 |
| ENSG00000146678 | IGFBP1 | -0.57 | 3.87E-04 | 1.00E-02 |
| ENSG00000145217 | SLC26A1 | 0.57 | 3.88E-04 | 1.00E-02 |
| ENSG00000130775 | THEMIS2 | -0.57 | 3.90E-04 | 1.01E-02 |
| ENSG00000161912 | ADCY10P1 | 0.57 | 3.90E-04 | 1.01E-02 |
| ENSG00000057657 | PRDM1 | -0.57 | 3.93E-04 | 1.01E-02 |
| ENSG00000104131 | EIF3J | -0.57 | 3.96E-04 | 1.02E-02 |
| ENSG00000205302 | SNX2 | -0.57 | 4.00E-04 | 1.03E-02 |
| ENSG00000111412 | C12orf49 | -0.57 | 4.03E-04 | 1.03E-02 |
| ENSG00000167720 | SRR | -0.57 | 4.03E-04 | 1.03E-02 |
| ENSG00000173418 | NAA20 | -0.57 | 4.03E-04 | 1.03E-02 |
| ENSG00000227496 | AC099066.2 | -0.57 | 4.05E-04 | 1.04E-02 |
| ENSG00000105879 | CBLL1 | -0.56 | 4.06E-04 | 1.04E-02 |
| ENSG00000244968 | LIFR-AS1 | 0.56 | 4.06E-04 | 1.04E-02 |
| ENSG00000153094 | BCL2L11 | -0.56 | 4.09E-04 | 1.04E-02 |
| ENSG00000239827 | SUGT1P3 | 0.56 | 4.09E-04 | 1.04E-02 |
| ENSG00000241684 | ADAMTS9-AS2 | 0.56 | 4.09E-04 | 1.04E-02 |
| ENSG00000044446 | PHKA2 | 0.56 | 4.13E-04 | 1.05E-02 |
| ENSG00000177103 | DSCAML1 | 0.56 | 4.13E-04 | 1.05E-02 |
| ENSG00000186812 | ZNF397 | 0.56 | 4.13E-04 | 1.05E-02 |
| ENSG00000072042 | RDH11 | -0.56 | 4.16E-04 | 1.05E-02 |
| ENSG00000119326 | CTNNAL1 | -0.56 | 4.19E-04 | 1.06E-02 |
| ENSG00000152953 | STK32B | 0.56 | 4.19E-04 | 1.06E-02 |
| ENSG00000169241 | SLC50A1 | -0.56 | 4.19E-04 | 1.06E-02 |
| ENSG00000197780 | TAF13 | -0.56 | 4.19E-04 | 1.06E-02 |
| ENSG00000145936 | KCNMB1 | -0.56 | 4.21E-04 | 1.06E-02 |
| ENSG00000213057 | C1orf220 | 0.56 | 4.22E-04 | 1.06E-02 |
| ENSG00000113810 | SMC4 | -0.56 | 4.23E-04 | 1.06E-02 |
| ENSG00000122729 | ACO1 | -0.56 | 4.23E-04 | 1.06E-02 |
| ENSG00000132793 | LPIN3 | 0.56 | 4.23E-04 | 1.06E-02 |
| ENSG00000140326 | CDAN1 | 0.56 | 4.23E-04 | 1.06E-02 |
| ENSG00000134690 | CDCA8 | -0.56 | 4.26E-04 | 1.06E-02 |
| ENSG00000146066 | HIGD2A | -0.56 | 4.26E-04 | 1.06E-02 |
| ENSG00000163249 | CCNYL1 | -0.56 | 4.26E-04 | 1.06E-02 |
| ENSG00000198856 | OSTC | -0.56 | 4.26E-04 | 1.06E-02 |
| ENSG00000228118 | MYL6P5 | -0.56 | 4.26E-04 | 1.06E-02 |

| Ensembl ID | Gene Name | ρ | Pval | Padj |
|---|---|---|---|---|
| ENSG00000116017 | ARID3A | -0.56 | 4.30E-04 | 1.07E-02 |
| ENSG00000269068 | AC009955.4 | 0.56 | 4.33E-04 | 1.07E-02 |
| ENSG00000205710 | C17orf107 | 0.56 | 4.33E-04 | 1.07E-02 |
| ENSG00000062485 | CS | -0.56 | 4.37E-04 | 1.08E-02 |
| ENSG00000116906 | GNPAT | -0.56 | 4.37E-04 | 1.08E-02 |
| ENSG00000178764 | ZHX2 | 0.56 | 4.37E-04 | 1.08E-02 |
| ENSG00000114544 | SLC41A3 | 0.56 | 4.40E-04 | 1.08E-02 |
| ENSG00000155959 | VBP1 | -0.56 | 4.40E-04 | 1.08E-02 |
| ENSG00000164484 | TMEM200A | -0.56 | 4.40E-04 | 1.08E-02 |
| ENSG00000168461 | RAB31 | -0.56 | 4.40E-04 | 1.08E-02 |
| ENSG00000064601 | CTSA | -0.56 | 4.44E-04 | 1.08E-02 |
| ENSG00000123684 | LPGAT1 | -0.56 | 4.44E-04 | 1.08E-02 |
| ENSG00000198039 | ZNF273 | 0.56 | 4.44E-04 | 1.08E-02 |
| ENSG00000054277 | OPN3 | -0.56 | 4.47E-04 | 1.09E-02 |
| ENSG00000180914 | OXTR | -0.56 | 4.49E-04 | 1.09E-02 |
| ENSG00000224769 | MUC20P1 | 0.56 | 4.50E-04 | 1.10E-02 |
| ENSG00000084444 | FAM234B | 0.56 | 4.51E-04 | 1.10E-02 |
| ENSG00000137714 | FDX1 | -0.56 | 4.51E-04 | 1.10E-02 |
| ENSG00000011600 | TYROBP | -0.56 | 4.54E-04 | 1.10E-02 |
| ENSG00000137200 | CMTR1 | 0.56 | 4.55E-04 | 1.10E-02 |
| ENSG00000176788 | BASP1 | -0.56 | 4.55E-04 | 1.10E-02 |
| ENSG00000186472 | PCLO | 0.56 | 4.55E-04 | 1.10E-02 |
| ENSG00000178055 | PRSS42 | 0.56 | 4.59E-04 | 1.11E-02 |
| ENSG00000068001 | HYAL2 | -0.56 | 4.62E-04 | 1.11E-02 |
| ENSG00000135451 | TROAP | -0.56 | 4.65E-04 | 1.12E-02 |
| ENSG00000100227 | POLDIP3 | 0.56 | 4.66E-04 | 1.12E-02 |
| ENSG00000143387 | CTSK | -0.56 | 4.66E-04 | 1.12E-02 |
| ENSG00000260027 | HOXB7 | -0.56 | 4.66E-04 | 1.12E-02 |
| ENSG00000198728 | LDB1 | 0.56 | 4.69E-04 | 1.12E-02 |
| ENSG00000250305 | TRMT9B | 0.56 | 4.69E-04 | 1.12E-02 |
| ENSG00000102802 | MEDAG | -0.56 | 4.73E-04 | 1.13E-02 |
| ENSG00000180113 | TDRD6 | 0.56 | 4.73E-04 | 1.13E-02 |
| ENSG00000184905 | TCEAL2 | 0.56 | 4.73E-04 | 1.13E-02 |
| ENSG00000213719 | CLIC1 | -0.56 | 4.73E-04 | 1.13E-02 |
| ENSG00000272462 | U91328.1 | 0.56 | 4.73E-04 | 1.13E-02 |
| ENSG00000138760 | SCARB2 | -0.56 | 4.77E-04 | 1.13E-02 |
| ENSG00000213865 | C8orf44 | 0.56 | 4.77E-04 | 1.13E-02 |
| ENSG00000065328 | MCM10 | -0.56 | 4.78E-04 | 1.13E-02 |
| ENSG00000157017 | GHRL | 0.56 | 4.80E-04 | 1.14E-02 |
| ENSG00000170145 | SIK2 | 0.56 | 4.81E-04 | 1.14E-02 |
| ENSG00000223501 | VPS52 | 0.56 | 4.81E-04 | 1.14E-02 |
| ENSG00000124181 | PLCG1 | 0.56 | 4.84E-04 | 1.14E-02 |
| ENSG00000235531 | MSC-AS1 | -0.56 | 4.85E-04 | 1.14E-02 |
| ENSG00000014914 | MTMR11 | 0.56 | 4.88E-04 | 1.15E-02 |
| ENSG00000086300 | SNX10 | -0.56 | 4.88E-04 | 1.15E-02 |

| Ensembl ID | Gene Name | ρ | Pval | Padj |
|---|---|---|---|---|
| ENSG00000100979 | PLTP | -0.56 | 4.88E-04 | 1.15E-02 |
| ENSG00000162512 | SDC3 | 0.56 | 4.88E-04 | 1.15E-02 |
| ENSG00000222032 | AC112721.2 | -0.56 | 4.88E-04 | 1.15E-02 |
| ENSG00000163083 | INHBB | -0.56 | 4.96E-04 | 1.16E-02 |
| ENSG00000163823 | CCR1 | -0.56 | 4.97E-04 | 1.16E-02 |
| ENSG00000254016 | ALG1L10P | 0.56 | 4.98E-04 | 1.16E-02 |
| ENSG00000140105 | WARS | -0.56 | 5.00E-04 | 1.17E-02 |
| ENSG00000249936 | RAC1P2 | -0.56 | 5.00E-04 | 1.17E-02 |
| ENSG00000129925 | TMEM8A | -0.56 | 5.04E-04 | 1.17E-02 |
| ENSG00000138834 | MAPK8IP3 | 0.56 | 5.04E-04 | 1.17E-02 |
| ENSG00000187715 | KBTBD12 | 0.56 | 5.04E-04 | 1.17E-02 |
| ENSG00000196275 | GTF2IRD2 | 0.56 | 5.04E-04 | 1.17E-02 |
| ENSG00000171860 | C3AR1 | -0.56 | 5.04E-04 | 1.17E-02 |
| ENSG00000008735 | MAPK8IP2 | 0.56 | 5.08E-04 | 1.17E-02 |
| ENSG00000134901 | KDELC1 | -0.56 | 5.08E-04 | 1.17E-02 |
| ENSG00000154518 | ATP5MC3 | -0.56 | 5.08E-04 | 1.17E-02 |
| ENSG00000247679 | AC139795.2 | 0.56 | 5.08E-04 | 1.17E-02 |
| ENSG00000143178 | TBX19 | 0.56 | 5.12E-04 | 1.17E-02 |
| ENSG00000145354 | CISD2 | -0.56 | 5.12E-04 | 1.17E-02 |
| ENSG00000163633 | C4orf36 | 0.56 | 5.12E-04 | 1.17E-02 |
| ENSG00000188015 | S100A3 | -0.56 | 5.12E-04 | 1.17E-02 |
| ENSG00000241644 | INMT | -0.56 | 5.12E-04 | 1.17E-02 |
| ENSG00000277443 | MARCKS | -0.56 | 5.12E-04 | 1.17E-02 |
| ENSG00000105880 | DLX5 | 0.56 | 5.16E-04 | 1.18E-02 |
| ENSG00000112972 | HMGCS1 | -0.56 | 5.16E-04 | 1.18E-02 |
| ENSG00000127863 | TNFRSF19 | -0.56 | 5.16E-04 | 1.18E-02 |
| ENSG00000166477 | LEO1 | -0.56 | 5.16E-04 | 1.18E-02 |
| ENSG00000169679 | BUB1 | -0.56 | 5.16E-04 | 1.18E-02 |
| ENSG00000233858 | LINC02599 | 0.56 | 5.16E-04 | 1.18E-02 |
| ENSG00000125835 | SNRPB | -0.56 | 5.20E-04 | 1.18E-02 |
| ENSG00000167613 | LAIR1 | -0.56 | 5.20E-04 | 1.18E-02 |
| ENSG00000260285 | AL133367.1 | 0.56 | 5.21E-04 | 1.18E-02 |
| ENSG00000230552 | AC092162.2 | 0.56 | 5.21E-04 | 1.18E-02 |
| ENSG00000129933 | MAU2 | 0.56 | 5.24E-04 | 1.18E-02 |
| ENSG00000144867 | SRPRB | -0.56 | 5.24E-04 | 1.18E-02 |
| ENSG00000248329 | APELA | -0.56 | 5.26E-04 | 1.19E-02 |
| ENSG00000027697 | IFNGR1 | -0.56 | 5.28E-04 | 1.19E-02 |
| ENSG00000118640 | VAMP8 | -0.56 | 5.28E-04 | 1.19E-02 |
| ENSG00000137547 | MRPL15 | -0.56 | 5.28E-04 | 1.19E-02 |
| ENSG00000006118 | TMEM132A | -0.56 | 5.30E-04 | 1.19E-02 |
| ENSG00000066382 | MPPED2 | 0.56 | 5.33E-04 | 1.19E-02 |
| ENSG00000243323 | PTPRVP | 0.56 | 5.33E-04 | 1.19E-02 |
| ENSG00000118689 | FOXO3 | 0.56 | 5.37E-04 | 1.20E-02 |
| ENSG00000138835 | RGS3 | -0.56 | 5.37E-04 | 1.20E-02 |
| ENSG00000166128 | RAB8B | -0.56 | 5.37E-04 | 1.20E-02 |

| Ensembl ID | Gene Name | ρ | Pval | Padj |
|---|---|---|---|---|
| ENSG00000196460 | RFX8 | -0.56 | 5.39E-04 | 1.20E-02 |
| ENSG00000144118 | RALB | -0.55 | 5.41E-04 | 1.21E-02 |
| ENSG00000105419 | MEIS3 | -0.55 | 5.45E-04 | 1.21E-02 |
| ENSG00000158859 | ADAMTS4 | -0.55 | 5.45E-04 | 1.21E-02 |
| ENSG00000144057 | ST6GAL2 | -0.55 | 5.49E-04 | 1.22E-02 |
| ENSG00000072401 | UBE2D1 | -0.55 | 5.50E-04 | 1.22E-02 |
| ENSG00000139318 | DUSP6 | -0.55 | 5.50E-04 | 1.22E-02 |
| ENSG00000183876 | ARSI | -0.55 | 5.50E-04 | 1.22E-02 |
| ENSG00000279662 | AC131649.2 | 0.55 | 5.50E-04 | 1.22E-02 |
| ENSG00000266538 | AC005838.2 | 0.55 | 5.52E-04 | 1.22E-02 |
| ENSG00000066248 | NGEF | 0.55 | 5.54E-04 | 1.22E-02 |
| ENSG00000119965 | C10orf88 | -0.55 | 5.54E-04 | 1.22E-02 |
| ENSG00000185222 | TCEAL9 | -0.55 | 5.54E-04 | 1.22E-02 |
| ENSG00000116254 | CHD5 | 0.55 | 5.58E-04 | 1.22E-02 |
| ENSG00000166924 | NYAP1 | 0.55 | 5.58E-04 | 1.22E-02 |
| ENSG00000167114 | SLC27A4 | -0.55 | 5.58E-04 | 1.22E-02 |
| ENSG00000182985 | CADM1 | -0.55 | 5.58E-04 | 1.22E-02 |
| ENSG00000092330 | TINF2 | -0.55 | 5.63E-04 | 1.23E-02 |
| ENSG00000099785 | MARCH2 | -0.55 | 5.63E-04 | 1.23E-02 |
| ENSG00000108010 | GLRX3 | -0.55 | 5.63E-04 | 1.23E-02 |
| ENSG00000168092 | PAFAH1B2 | -0.55 | 5.63E-04 | 1.23E-02 |
| ENSG00000178952 | TUFM | -0.55 | 5.63E-04 | 1.23E-02 |
| ENSG00000181061 | HIGD1A | -0.55 | 5.63E-04 | 1.23E-02 |
| ENSG00000270094 | AL670729.2 | -0.55 | 5.67E-04 | 1.23E-02 |
| ENSG00000136045 | PWP1 | -0.55 | 5.67E-04 | 1.23E-02 |
| ENSG00000268460 | AC006262.1 | 0.55 | 5.71E-04 | 1.24E-02 |
| ENSG00000168653 | NDUFS5 | -0.55 | 5.71E-04 | 1.24E-02 |
| ENSG00000068615 | REEP1 | -0.55 | 5.75E-04 | 1.25E-02 |
| ENSG00000230838 | LINC01614 | -0.55 | 5.76E-04 | 1.25E-02 |
| ENSG00000275055 | AC011468.5 | 0.55 | 5.82E-04 | 1.26E-02 |
| ENSG00000118785 | SPP1 | -0.55 | 5.85E-04 | 1.26E-02 |
| ENSG00000198938 | MT-CO3 | -0.55 | 5.90E-04 | 1.27E-02 |
| ENSG00000234880 | LINC00163 | 0.55 | 5.90E-04 | 1.27E-02 |
| ENSG00000261821 | AC090826.1 | 0.55 | 5.90E-04 | 1.27E-02 |
| ENSG00000140285 | FGF7 | -0.55 | 5.94E-04 | 1.27E-02 |
| ENSG00000078699 | CBFA2T2 | 0.55 | 5.94E-04 | 1.27E-02 |
| ENSG00000100221 | JOSD1 | -0.55 | 5.94E-04 | 1.27E-02 |
| ENSG00000136522 | MRPL47 | -0.55 | 5.94E-04 | 1.27E-02 |
| ENSG00000126368 | NR1D1 | 0.55 | 5.99E-04 | 1.28E-02 |
| ENSG00000138029 | HADHB | -0.55 | 5.99E-04 | 1.28E-02 |
| ENSG00000140416 | TPM1 | -0.55 | 5.99E-04 | 1.28E-02 |
| ENSG00000266412 | NCOA4 | -0.55 | 5.99E-04 | 1.28E-02 |
| ENSG00000092758 | COL9A3 | 0.55 | 6.03E-04 | 1.28E-02 |
| ENSG00000106479 | ZNF862 | 0.55 | 6.03E-04 | 1.28E-02 |
| ENSG00000143373 | ZNF687 | 0.55 | 6.03E-04 | 1.28E-02 |

| Ensembl ID | Gene Name | ρ | Pval | Padj |
|---|---|---|---|---|
| ENSG00000164309 | CMYA5 | 0.55 | 6.03E-04 | 1.28E-02 |
| ENSG00000183833 | MAATS1 | 0.55 | 6.03E-04 | 1.28E-02 |
| ENSG00000178999 | AURKB | -0.55 | 6.07E-04 | 1.28E-02 |
| ENSG00000033011 | ALG1 | -0.55 | 6.08E-04 | 1.28E-02 |
| ENSG00000056050 | HPF1 | -0.55 | 6.08E-04 | 1.28E-02 |
| ENSG00000165629 | ATP5F1C | -0.55 | 6.08E-04 | 1.28E-02 |
| ENSG00000172115 | CYCS | -0.55 | 6.08E-04 | 1.28E-02 |
| ENSG00000028839 | TBPL1 | -0.55 | 6.13E-04 | 1.29E-02 |
| ENSG00000085662 | AKR1B1 | -0.55 | 6.13E-04 | 1.29E-02 |
| ENSG00000197381 | ADARB1 | 0.55 | 6.13E-04 | 1.29E-02 |
| ENSG00000273142 | LINC02604 | 0.55 | 6.13E-04 | 1.29E-02 |
| ENSG00000099341 | PSMD8 | -0.55 | 6.18E-04 | 1.29E-02 |
| ENSG00000164647 | STEAP1 | -0.55 | 6.18E-04 | 1.29E-02 |
| ENSG00000167193 | CRK | -0.55 | 6.18E-04 | 1.29E-02 |
| ENSG00000172889 | EGFL7 | -0.55 | 6.18E-04 | 1.29E-02 |
| ENSG00000110060 | PUS3 | -0.55 | 6.22E-04 | 1.30E-02 |
| ENSG00000221944 | TIGD1 | 0.55 | 6.22E-04 | 1.30E-02 |
| ENSG00000065600 | TMEM206 | -0.55 | 6.27E-04 | 1.31E-02 |
| ENSG00000124126 | PREX1 | 0.55 | 6.27E-04 | 1.31E-02 |
| ENSG00000252835 | SCARNA21 | -0.55 | 6.29E-04 | 1.31E-02 |
| ENSG00000011198 | ABHD5 | -0.55 | 6.32E-04 | 1.31E-02 |
| ENSG00000151090 | THRB | 0.55 | 6.32E-04 | 1.31E-02 |
| ENSG00000008710 | PKD1 | 0.55 | 6.37E-04 | 1.32E-02 |
| ENSG00000212464 | SNORA12 | -0.55 | 6.37E-04 | 1.32E-02 |
| ENSG00000114744 | COMMD2 | -0.55 | 6.42E-04 | 1.33E-02 |
| ENSG00000163577 | EIF5A2 | -0.55 | 6.42E-04 | 1.33E-02 |
| ENSG00000055483 | USP36 | 0.55 | 6.47E-04 | 1.33E-02 |
| ENSG00000055813 | CCDC85A | 0.55 | 6.47E-04 | 1.33E-02 |
| ENSG00000154640 | BTG3 | -0.55 | 6.47E-04 | 1.33E-02 |
| ENSG00000168291 | PDHB | -0.55 | 6.47E-04 | 1.33E-02 |
| ENSG00000141030 | COPS3 | -0.55 | 6.52E-04 | 1.34E-02 |
| ENSG00000196154 | S100A4 | -0.55 | 6.52E-04 | 1.34E-02 |
| ENSG00000117650 | NEK2 | -0.55 | 6.57E-04 | 1.35E-02 |
| ENSG00000153902 | LGI4 | 0.55 | 6.57E-04 | 1.35E-02 |
| ENSG00000177106 | EPS8L2 | 0.55 | 6.57E-04 | 1.35E-02 |
| ENSG00000196591 | HDAC2 | -0.55 | 6.62E-04 | 1.36E-02 |
| ENSG00000231711 | LINC00899 | 0.55 | 6.62E-04 | 1.36E-02 |
| ENSG00000162520 | SYNC | -0.55 | 6.67E-04 | 1.36E-02 |
| ENSG00000119737 | GPR75 | 0.55 | 6.69E-04 | 1.37E-02 |
| ENSG00000271447 | MMP28 | -0.55 | 6.71E-04 | 1.37E-02 |
| ENSG00000101247 | NDUFAF5 | -0.55 | 6.72E-04 | 1.37E-02 |
| ENSG00000163170 | BOLA3 | -0.55 | 6.72E-04 | 1.37E-02 |
| ENSG00000188227 | ZNF793 | 0.55 | 6.72E-04 | 1.37E-02 |
| ENSG00000250132 | AC004803.1 | 0.55 | 6.72E-04 | 1.37E-02 |
| ENSG00000171700 | RGS19 | -0.55 | 6.77E-04 | 1.37E-02 |

| Ensembl ID | Gene Name | ρ | Pval | Padj |
|---|---|---|---|---|
| ENSG00000171824 | EXOSC10 | 0.55 | 6.77E-04 | 1.37E-02 |
| ENSG00000109674 | NEIL3 | -0.55 | 6.82E-04 | 1.38E-02 |
| ENSG00000225075 | AL603832.1 | 0.55 | 6.82E-04 | 1.38E-02 |
| ENSG00000159588 | CCDC17 | 0.55 | 6.82E-04 | 1.38E-02 |
| ENSG00000172965 | MIR4435-2HG | -0.55 | 6.82E-04 | 1.38E-02 |
| ENSG00000102753 | KPNA3 | -0.55 | 6.88E-04 | 1.38E-02 |
| ENSG00000113758 | DBN1 | -0.55 | 6.88E-04 | 1.38E-02 |
| ENSG00000164919 | COX6C | -0.55 | 6.88E-04 | 1.38E-02 |
| ENSG00000224861 | YBX1P1 | -0.55 | 6.88E-04 | 1.38E-02 |
| ENSG00000272556 | GTF2IP13 | 0.55 | 6.88E-04 | 1.38E-02 |
| ENSG00000278668 | AC005899.7 | 0.55 | 6.92E-04 | 1.39E-02 |
| ENSG00000070770 | CSNK2A2 | -0.55 | 6.93E-04 | 1.39E-02 |
| ENSG00000111186 | WNT5B | -0.55 | 6.93E-04 | 1.39E-02 |
| ENSG00000163297 | ANTXR2 | -0.55 | 6.93E-04 | 1.39E-02 |
| ENSG00000180389 | ATP5F1EP2 | -0.55 | 6.93E-04 | 1.39E-02 |
| ENSG00000184047 | DIABLO | 0.55 | 6.93E-04 | 1.39E-02 |
| ENSG00000088756 | ARHGAP28 | -0.55 | 6.98E-04 | 1.39E-02 |
| ENSG00000097096 | SYDE2 | 0.55 | 6.98E-04 | 1.39E-02 |
| ENSG00000128606 | LRRC17 | -0.55 | 6.98E-04 | 1.39E-02 |
| ENSG00000273247 | AC097376.2 | 0.55 | 6.98E-04 | 1.39E-02 |
| ENSG00000103035 | PSMD7 | -0.55 | 7.04E-04 | 1.40E-02 |
| ENSG00000137309 | HMGA1 | -0.55 | 7.04E-04 | 1.40E-02 |
| ENSG00000265354 | TIMM23 | -0.55 | 7.04E-04 | 1.40E-02 |
| ENSG00000084073 | ZMPSTE24 | -0.55 | 7.09E-04 | 1.40E-02 |
| ENSG00000113597 | TRAPPC13 | -0.55 | 7.09E-04 | 1.40E-02 |
| ENSG00000123095 | BHLHE41 | 0.55 | 7.09E-04 | 1.40E-02 |
| ENSG00000167850 | CD300C | -0.55 | 7.09E-04 | 1.40E-02 |
| ENSG00000169021 | UQCRFS1 | -0.55 | 7.09E-04 | 1.40E-02 |
| ENSG00000185305 | ARL15 | -0.55 | 7.09E-04 | 1.40E-02 |
| ENSG00000276067 | AC009542.2 | 0.55 | 7.10E-04 | 1.40E-02 |
| ENSG00000091140 | DLD | -0.54 | 7.14E-04 | 1.41E-02 |
| ENSG00000140199 | SLC12A6 | -0.54 | 7.20E-04 | 1.41E-02 |
| ENSG00000162542 | TMCO4 | 0.54 | 7.20E-04 | 1.41E-02 |
| ENSG00000166073 | GPR176 | -0.54 | 7.20E-04 | 1.41E-02 |
| ENSG00000242600 | MBL1P | 0.54 | 7.20E-04 | 1.41E-02 |
| ENSG00000274270 | AL137060.1 | 0.54 | 7.23E-04 | 1.41E-02 |
| ENSG00000084090 | STARD7 | -0.54 | 7.25E-04 | 1.41E-02 |
| ENSG00000132507 | EIF5A | -0.54 | 7.25E-04 | 1.41E-02 |
| ENSG00000172269 | DPAGT1 | -0.54 | 7.25E-04 | 1.41E-02 |
| ENSG00000179859 | RNF227 | 0.54 | 7.25E-04 | 1.41E-02 |
| ENSG00000185332 | TMEM105 | 0.54 | 7.25E-04 | 1.41E-02 |
| ENSG00000259327 | AC023906.3 | 0.54 | 7.30E-04 | 1.42E-02 |
| ENSG00000104695 | PPP2CB | -0.54 | 7.31E-04 | 1.42E-02 |
| ENSG00000143320 | CRABP2 | -0.54 | 7.31E-04 | 1.42E-02 |
| ENSG00000176407 | KCMF1 | -0.54 | 7.31E-04 | 1.42E-02 |

| Ensembl ID | Gene Name | ρ | Pval | Padj |
|---|---|---|---|---|
| ENSG00000101856 | PGRMC1 | -0.54 | 7.36E-04 | 1.43E-02 |
| ENSG00000153179 | RASSF3 | -0.54 | 7.36E-04 | 1.43E-02 |
| ENSG00000158163 | DZIP1L | 0.54 | 7.36E-04 | 1.43E-02 |
| ENSG00000271858 | Z84492.1 | 0.54 | 7.42E-04 | 1.44E-02 |
| ENSG00000235151 | AC131097.4 | 0.54 | 7.43E-04 | 1.44E-02 |
| ENSG00000143418 | CERS2 | -0.54 | 7.48E-04 | 1.44E-02 |
| ENSG00000139679 | LPAR6 | -0.54 | 7.53E-04 | 1.45E-02 |
| ENSG00000151893 | CACUL1 | -0.54 | 7.53E-04 | 1.45E-02 |
| ENSG00000172061 | LRRC15 | -0.54 | 7.53E-04 | 1.45E-02 |
| ENSG00000279943 | AC110771.1 | 0.54 | 7.56E-04 | 1.45E-02 |
| ENSG00000139644 | TMBIM6 | -0.54 | 7.59E-04 | 1.46E-02 |
| ENSG00000072571 | HMMR | -0.54 | 7.61E-04 | 1.46E-02 |
| ENSG00000134001 | EIF2S1 | -0.54 | 7.65E-04 | 1.47E-02 |
| ENSG00000159674 | SPON2 | -0.54 | 7.65E-04 | 1.47E-02 |
| ENSG00000090104 | RGS1 | -0.54 | 7.66E-04 | 1.47E-02 |
| ENSG00000107020 | PLGRKT | -0.54 | 7.71E-04 | 1.47E-02 |
| ENSG00000121281 | ADCY7 | -0.54 | 7.71E-04 | 1.47E-02 |
| ENSG00000232104 | RFX3-AS1 | 0.54 | 7.76E-04 | 1.48E-02 |
| ENSG00000013288 | MAN2B2 | 0.54 | 7.76E-04 | 1.48E-02 |
| ENSG00000140525 | FANCI | -0.54 | 7.82E-04 | 1.49E-02 |
| ENSG00000148248 | SURF4 | -0.54 | 7.82E-04 | 1.49E-02 |
| ENSG00000160360 | GPSM1 | 0.54 | 7.82E-04 | 1.49E-02 |
| ENSG00000179094 | AC129492.1 | 0.54 | 7.82E-04 | 1.49E-02 |
| ENSG00000171798 | KNDC1 | 0.54 | 7.88E-04 | 1.49E-02 |
| ENSG00000184471 | C1QTNF8 | 0.54 | 7.88E-04 | 1.49E-02 |
| ENSG00000188112 | C6orf132 | -0.54 | 7.88E-04 | 1.49E-02 |
| ENSG00000276649 | AL117335.1 | 0.54 | 7.88E-04 | 1.49E-02 |
| ENSG00000100097 | LGALS1 | -0.54 | 7.94E-04 | 1.50E-02 |
| ENSG00000135916 | ITM2C | 0.54 | 7.94E-04 | 1.50E-02 |
| ENSG00000198898 | CAPZA2 | -0.54 | 7.94E-04 | 1.50E-02 |
| ENSG00000100372 | SLC25A17 | -0.54 | 8.00E-04 | 1.50E-02 |
| ENSG00000111845 | PAK1IP1 | -0.54 | 8.00E-04 | 1.50E-02 |
| ENSG00000134108 | ARL8B | -0.54 | 8.00E-04 | 1.50E-02 |
| ENSG00000176834 | VSIG10 | 0.54 | 8.00E-04 | 1.50E-02 |
| ENSG00000126822 | PLEKHG3 | 0.54 | 8.06E-04 | 1.51E-02 |
| ENSG00000130558 | OLFM1 | 0.54 | 8.12E-04 | 1.52E-02 |
| ENSG00000166902 | MRPL16 | -0.54 | 8.12E-04 | 1.52E-02 |
| ENSG00000251432 | LINC02615 | 0.54 | 8.12E-04 | 1.52E-02 |
| ENSG00000101347 | SAMHD1 | -0.54 | 8.24E-04 | 1.54E-02 |
| ENSG00000104341 | LAPTM4B | -0.54 | 8.24E-04 | 1.54E-02 |
| ENSG00000256546 | AC156455.1 | 0.54 | 8.24E-04 | 1.54E-02 |
| ENSG00000129965 | INS-IGF2 | 0.54 | 8.27E-04 | 1.54E-02 |
| ENSG00000125266 | EFNB2 | -0.54 | 8.27E-04 | 1.54E-02 |
| ENSG00000113648 | H2AFY | -0.54 | 8.31E-04 | 1.54E-02 |
| ENSG00000118508 | RAB32 | -0.54 | 8.31E-04 | 1.54E-02 |

| Ensembl ID | Gene Name | ρ | Pval | Padj |
|---|---|---|---|---|
| ENSG00000147140 | NONO | -0.54 | 8.31E-04 | 1.54E-02 |
| ENSG00000236094 | LINC00545 | -0.54 | 8.31E-04 | 1.54E-02 |
| ENSG00000276805 | AL133216.2 | 0.54 | 8.31E-04 | 1.54E-02 |
| ENSG00000259295 | CSPG4P12 | 0.54 | 8.35E-04 | 1.54E-02 |
| ENSG00000141682 | PMAIP1 | -0.54 | 8.36E-04 | 1.54E-02 |
| ENSG00000067113 | PLPP1 | -0.54 | 8.37E-04 | 1.54E-02 |
| ENSG00000135441 | BLOC1S1 | -0.54 | 8.37E-04 | 1.54E-02 |
| ENSG00000171357 | LURAP1 | 0.54 | 8.37E-04 | 1.54E-02 |
| ENSG00000231290 | APCDD1L-DT | -0.54 | 8.37E-04 | 1.54E-02 |
| ENSG00000010270 | STARD3NL | -0.54 | 8.43E-04 | 1.55E-02 |
| ENSG00000049239 | H6PD | 0.54 | 8.43E-04 | 1.55E-02 |
| ENSG00000116977 | LGALS8 | -0.54 | 8.43E-04 | 1.55E-02 |
| ENSG00000151748 | SAV1 | 0.54 | 8.43E-04 | 1.55E-02 |
| ENSG00000197153 | HIST1H3J | -0.54 | 8.51E-04 | 1.56E-02 |
| ENSG00000266527 | AC005697.2 | 0.54 | 8.51E-04 | 1.56E-02 |
| ENSG00000223561 | AC005165.1 | 0.54 | 8.52E-04 | 1.56E-02 |
| ENSG00000011422 | PLAUR | -0.54 | 8.56E-04 | 1.56E-02 |
| ENSG00000117450 | PRDX1 | -0.54 | 8.56E-04 | 1.56E-02 |
| ENSG00000158747 | NBL1 | -0.54 | 8.56E-04 | 1.56E-02 |
| ENSG00000162702 | ZNF281 | -0.54 | 8.56E-04 | 1.56E-02 |
| ENSG00000163644 | PPM1K | 0.54 | 8.56E-04 | 1.56E-02 |
| ENSG00000163734 | CXCL3 | -0.54 | 8.58E-04 | 1.56E-02 |
| ENSG00000101079 | NDRG3 | -0.54 | 8.62E-04 | 1.57E-02 |
| ENSG00000105270 | CLIP3 | 0.54 | 8.62E-04 | 1.57E-02 |
| ENSG00000165704 | HPRT1 | -0.54 | 8.62E-04 | 1.57E-02 |
| ENSG00000175324 | LSM1 | -0.54 | 8.62E-04 | 1.57E-02 |
| ENSG00000140030 | GPR65 | -0.54 | 8.63E-04 | 1.57E-02 |
| ENSG00000105509 | HAS1 | -0.54 | 8.65E-04 | 1.57E-02 |
| ENSG00000048052 | HDAC9 | 0.54 | 8.69E-04 | 1.57E-02 |
| ENSG00000075651 | PLD1 | -0.54 | 8.69E-04 | 1.57E-02 |
| ENSG00000198042 | MAK16 | -0.54 | 8.69E-04 | 1.57E-02 |
| ENSG00000242294 | STAG3L5P | 0.54 | 8.69E-04 | 1.57E-02 |
| ENSG00000257557 | PPP1R12A-AS1 | 0.54 | 8.71E-04 | 1.57E-02 |
| ENSG00000153310 | FAM49B | -0.54 | 8.75E-04 | 1.58E-02 |
| ENSG00000167525 | PROCA1 | 0.54 | 8.75E-04 | 1.58E-02 |
| ENSG00000102898 | NUTF2 | -0.54 | 8.82E-04 | 1.58E-02 |
| ENSG00000179604 | CDC42EP4 | 0.54 | 8.82E-04 | 1.58E-02 |
| ENSG00000185561 | TLCD2 | 0.54 | 8.82E-04 | 1.58E-02 |
| ENSG00000248334 | WHAMMP2 | 0.54 | 8.82E-04 | 1.58E-02 |
| ENSG00000217835 | AL034397.1 | 0.54 | 8.83E-04 | 1.58E-02 |
| ENSG00000122034 | GTF3A | -0.54 | 8.88E-04 | 1.59E-02 |
| ENSG00000143624 | INTS3 | 0.54 | 8.88E-04 | 1.59E-02 |
| ENSG00000149600 | COMMD7 | -0.54 | 8.88E-04 | 1.59E-02 |
| ENSG00000230715 | AC018638.2 | 0.54 | 8.88E-04 | 1.59E-02 |
| ENSG00000251474 | RPL32P3 | 0.54 | 8.88E-04 | 1.59E-02 |

| Ensembl ID | Gene Name | ρ | Pval | Padj |
|---|---|---|---|---|
| ENSG00000279932 | AP006248.4 | 0.54 | 8.93E-04 | 1.59E-02 |
| ENSG00000125386 | FAM193A | 0.54 | 8.95E-04 | 1.59E-02 |
| ENSG00000239900 | ADSL | -0.54 | 8.95E-04 | 1.59E-02 |
| ENSG00000156463 | SH3RF2 | 0.54 | 9.01E-04 | 1.60E-02 |
| ENSG00000204060 | FOXO6 | 0.54 | 9.01E-04 | 1.60E-02 |
| ENSG00000105171 | POP4 | -0.54 | 9.08E-04 | 1.61E-02 |
| ENSG00000113643 | RARS | -0.54 | 9.08E-04 | 1.61E-02 |
| ENSG00000187446 | CHP1 | -0.54 | 9.08E-04 | 1.61E-02 |
| ENSG00000174721 | FGFBP3 | 0.54 | 9.15E-04 | 1.62E-02 |
| ENSG00000206344 | HCG27 | 0.54 | 9.15E-04 | 1.62E-02 |
| ENSG00000233514 | AL356653.1 | -0.54 | 9.16E-04 | 1.62E-02 |
| ENSG00000062716 | VMP1 | -0.54 | 9.22E-04 | 1.63E-02 |
| ENSG00000103335 | PIEZO1 | 0.54 | 9.28E-04 | 1.64E-02 |
| ENSG00000198355 | PIM3 | 0.54 | 9.28E-04 | 1.64E-02 |
| ENSG00000233006 | MIR3936HG | 0.54 | 9.28E-04 | 1.64E-02 |
| ENSG00000107719 | PALD1 | 0.53 | 9.35E-04 | 1.64E-02 |
| ENSG00000119812 | FAM98A | -0.53 | 9.35E-04 | 1.64E-02 |
| ENSG00000137285 | TUBB2B | 0.53 | 9.35E-04 | 1.64E-02 |
| ENSG00000149403 | GRIK4 | 0.53 | 9.35E-04 | 1.64E-02 |
| ENSG00000174282 | ZBTB4 | 0.53 | 9.35E-04 | 1.64E-02 |
| ENSG00000231991 | ANXA2P2 | -0.53 | 9.35E-04 | 1.64E-02 |
| ENSG00000017483 | SLC38A5 | -0.53 | 9.42E-04 | 1.65E-02 |
| ENSG00000100410 | PHF5A | -0.53 | 9.42E-04 | 1.65E-02 |
| ENSG00000170606 | HSPA4 | -0.53 | 9.42E-04 | 1.65E-02 |
| ENSG00000204396 | VWA7 | 0.53 | 9.42E-04 | 1.65E-02 |
| ENSG00000204959 | ARHGEF34P | 0.53 | 9.42E-04 | 1.65E-02 |
| ENSG00000086015 | MAST2 | 0.53 | 9.49E-04 | 1.65E-02 |
| ENSG00000150787 | PTS | -0.53 | 9.49E-04 | 1.65E-02 |
| ENSG00000275023 | MLLT6 | 0.53 | 9.49E-04 | 1.65E-02 |
| ENSG00000137507 | LRRC32 | -0.53 | 9.56E-04 | 1.66E-02 |
| ENSG00000156232 | WHAMM | 0.53 | 9.56E-04 | 1.66E-02 |
| ENSG00000037241 | RPL26L1 | -0.53 | 9.63E-04 | 1.67E-02 |
| ENSG00000136828 | RALGPS1 | 0.53 | 9.63E-04 | 1.67E-02 |
| ENSG00000196368 | NUDT11 | -0.53 | 9.63E-04 | 1.67E-02 |
| ENSG00000267287 | AC068473.3 | 0.53 | 9.63E-04 | 1.67E-02 |
| ENSG00000267546 | AC015802.4 | 0.53 | 9.65E-04 | 1.67E-02 |
| ENSG00000070476 | ZXDC | 0.53 | 9.70E-04 | 1.67E-02 |
| ENSG00000109436 | TBC1D9 | -0.53 | 9.70E-04 | 1.67E-02 |
| ENSG00000170558 | CDH2 | -0.53 | 9.70E-04 | 1.67E-02 |
| ENSG00000272693 | AC073107.1 | 0.53 | 9.70E-04 | 1.67E-02 |
| ENSG00000020256 | ZFP64 | 0.53 | 9.77E-04 | 1.68E-02 |
| ENSG00000148848 | ADAM12 | -0.53 | 9.77E-04 | 1.68E-02 |
| ENSG00000152234 | ATP5F1A | -0.53 | 9.77E-04 | 1.68E-02 |
| ENSG00000205105 | COX17P1 | -0.53 | 9.84E-04 | 1.69E-02 |
| ENSG00000089472 | HEPH | -0.53 | 9.85E-04 | 1.69E-02 |

| Ensembl ID | Gene Name | ρ | Pval | Padj |
|---|---|---|---|---|
| ENSG00000121579 | NAA50 | -0.53 | 9.85E-04 | 1.69E-02 |
| ENSG00000138771 | SHROOM3 | 0.53 | 9.85E-04 | 1.69E-02 |
| ENSG00000103978 | TMEM87A | -0.53 | 9.92E-04 | 1.69E-02 |
| ENSG00000141759 | TXNL4A | -0.53 | 9.92E-04 | 1.69E-02 |
| ENSG00000167283 | ATP5MG | -0.53 | 9.92E-04 | 1.69E-02 |
| ENSG00000185513 | L3MBTL1 | 0.53 | 9.92E-04 | 1.69E-02 |
| ENSG00000244560 | AC004890.2 | 0.53 | 9.92E-04 | 1.69E-02 |
| ENSG00000262580 | AC087741.1 | 0.53 | 9.94E-04 | 1.70E-02 |
| ENSG00000090520 | DNAJB11 | -0.53 | 9.99E-04 | 1.70E-02 |
| ENSG00000111199 | TRPV4 | 0.53 | 9.99E-04 | 1.70E-02 |
| ENSG00000213867 | AL049830.1 | -0.53 | 1.00E-03 | 1.70E-02 |
| ENSG00000149532 | CPSF7 | 0.53 | 1.01E-03 | 1.71E-02 |
| ENSG00000230989 | HSBP1 | -0.53 | 1.01E-03 | 1.71E-02 |
| ENSG00000143771 | CNIH4 | -0.53 | 1.01E-03 | 1.72E-02 |
| ENSG00000189058 | APOD | 0.53 | 1.01E-03 | 1.72E-02 |
| ENSG00000143851 | PTPN7 | -0.53 | 1.01E-03 | 1.72E-02 |
| ENSG00000255153 | TOLLIP-AS1 | 0.53 | 1.02E-03 | 1.73E-02 |
| ENSG00000136261 | BZW2 | -0.53 | 1.02E-03 | 1.73E-02 |
| ENSG00000154736 | ADAMTS5 | -0.53 | 1.02E-03 | 1.73E-02 |
| ENSG00000223478 | AL441992.1 | -0.53 | 1.03E-03 | 1.73E-02 |
| ENSG00000101134 | DOK5 | -0.53 | 1.03E-03 | 1.73E-02 |
| ENSG00000174348 | PODN | -0.53 | 1.03E-03 | 1.73E-02 |
| ENSG00000179933 | C14orf119 | -0.53 | 1.03E-03 | 1.73E-02 |
| ENSG00000197566 | ZNF624 | -0.53 | 1.03E-03 | 1.73E-02 |
| ENSG00000229127 | AC007038.1 | 0.53 | 1.03E-03 | 1.73E-02 |
| ENSG00000175183 | CSRP2 | -0.53 | 1.03E-03 | 1.73E-02 |
| ENSG00000104687 | GSR | -0.53 | 1.04E-03 | 1.74E-02 |
| ENSG00000171951 | SCG2 | -0.53 | 1.04E-03 | 1.74E-02 |
| ENSG00000060709 | RIMBP2 | 0.53 | 1.04E-03 | 1.74E-02 |
| ENSG00000126088 | UROD | -0.53 | 1.04E-03 | 1.74E-02 |
| ENSG00000131174 | COX7B | -0.53 | 1.04E-03 | 1.74E-02 |
| ENSG00000164733 | CTSB | -0.53 | 1.04E-03 | 1.74E-02 |
| ENSG00000169019 | COMMD8 | -0.53 | 1.04E-03 | 1.74E-02 |
| ENSG00000241935 | HOGA1 | 0.53 | 1.04E-03 | 1.74E-02 |
| ENSG00000277142 | LINC00235 | 0.53 | 1.05E-03 | 1.76E-02 |
| ENSG00000182873 | PRKCZ-AS1 | 0.53 | 1.06E-03 | 1.76E-02 |
| ENSG00000135473 | PAN2 | 0.53 | 1.06E-03 | 1.76E-02 |
| ENSG00000139651 | ZNF740 | 0.53 | 1.06E-03 | 1.76E-02 |
| ENSG00000174353 | STAG3L3 | 0.53 | 1.06E-03 | 1.76E-02 |
| ENSG00000234616 | JRK | 0.53 | 1.06E-03 | 1.76E-02 |
| ENSG00000261474 | AC026471.4 | 0.53 | 1.06E-03 | 1.76E-02 |
| ENSG00000234807 | LINC01135 | 0.53 | 1.06E-03 | 1.76E-02 |
| ENSG00000105227 | PRX | 0.53 | 1.07E-03 | 1.76E-02 |
| ENSG00000113732 | ATP6V0E1 | -0.53 | 1.07E-03 | 1.76E-02 |
| ENSG00000171497 | PPID | -0.53 | 1.07E-03 | 1.76E-02 |

| Ensembl ID | Gene Name | ρ | Pval | Padj |
|---|---|---|---|---|
| ENSG00000186063 | AIDA | -0.53 | 1.07E-03 | 1.76E-02 |
| ENSG00000188662 | HILS1 | 0.53 | 1.07E-03 | 1.76E-02 |
| ENSG00000268530 | AC008403.3 | -0.53 | 1.07E-03 | 1.76E-02 |
| ENSG00000087266 | SH3BP2 | 0.53 | 1.07E-03 | 1.77E-02 |
| ENSG00000131508 | UBE2D2 | -0.53 | 1.07E-03 | 1.77E-02 |
| ENSG00000150630 | VEGFC | -0.53 | 1.07E-03 | 1.77E-02 |
| ENSG00000220924 | OSTCP4 | -0.53 | 1.08E-03 | 1.77E-02 |
| ENSG00000259826 | AC072061.1 | 0.53 | 1.08E-03 | 1.78E-02 |
| ENSG00000281969 | AL592158.1 | 0.53 | 1.09E-03 | 1.79E-02 |
| ENSG00000153786 | ZDHHC7 | -0.53 | 1.10E-03 | 1.80E-02 |
| ENSG00000158864 | NDUFS2 | -0.53 | 1.10E-03 | 1.80E-02 |
| ENSG00000170340 | B3GNT2 | -0.53 | 1.10E-03 | 1.80E-02 |
| ENSG00000183230 | CTNNA3 | 0.53 | 1.10E-03 | 1.81E-02 |
| ENSG00000113389 | NPR3 | -0.53 | 1.11E-03 | 1.81E-02 |
| ENSG00000130988 | RGN | 0.53 | 1.11E-03 | 1.81E-02 |
| ENSG00000226457 | RPL22P3 | -0.53 | 1.11E-03 | 1.81E-02 |
| ENSG00000250012 | AC079848.1 | 0.53 | 1.11E-03 | 1.81E-02 |
| ENSG00000162377 | COA7 | -0.53 | 1.11E-03 | 1.81E-02 |
| ENSG00000197976 | AKAP17A | 0.53 | 1.11E-03 | 1.81E-02 |
| ENSG00000235437 | LINC01278 | 0.53 | 1.11E-03 | 1.81E-02 |
| ENSG00000125977 | EIF2S2 | -0.53 | 1.12E-03 | 1.82E-02 |
| ENSG00000189159 | JPT1 | -0.53 | 1.12E-03 | 1.82E-02 |
| ENSG00000044090 | CUL7 | 0.53 | 1.13E-03 | 1.83E-02 |
| ENSG00000129255 | MPDU1 | -0.53 | 1.13E-03 | 1.83E-02 |
| ENSG00000086848 | ALG9 | 0.53 | 1.14E-03 | 1.84E-02 |
| ENSG00000267100 | ILF3-DT | 0.53 | 1.14E-03 | 1.84E-02 |
| ENSG00000243970 | PPIEL | 0.53 | 1.15E-03 | 1.85E-02 |
| ENSG00000087303 | NID2 | -0.53 | 1.15E-03 | 1.85E-02 |
| ENSG00000134986 | NREP | -0.53 | 1.15E-03 | 1.86E-02 |
| ENSG00000227855 | DPY19L2P3 | 0.53 | 1.15E-03 | 1.86E-02 |
| ENSG00000233225 | AC004987.2 | 0.53 | 1.15E-03 | 1.86E-02 |
| ENSG00000143514 | TP53BP2 | -0.53 | 1.16E-03 | 1.87E-02 |
| ENSG00000105011 | ASF1B | -0.53 | 1.17E-03 | 1.88E-02 |
| ENSG00000070756 | PABPC1 | -0.53 | 1.17E-03 | 1.88E-02 |
| ENSG00000158716 | DUSP23 | -0.53 | 1.17E-03 | 1.88E-02 |
| ENSG00000171045 | TSNARE1 | 0.53 | 1.17E-03 | 1.88E-02 |
| ENSG00000224837 | GCSHP5 | -0.53 | 1.17E-03 | 1.88E-02 |
| ENSG00000050628 | PTGER3 | 0.53 | 1.18E-03 | 1.89E-02 |
| ENSG00000101911 | PRPS2 | -0.53 | 1.18E-03 | 1.89E-02 |
| ENSG00000132676 | DAP3 | -0.53 | 1.18E-03 | 1.89E-02 |
| ENSG00000138792 | ENPEP | -0.53 | 1.18E-03 | 1.89E-02 |
| ENSG00000180329 | CCDC43 | -0.53 | 1.18E-03 | 1.89E-02 |
| ENSG00000132622 | HSPA12B | -0.53 | 1.19E-03 | 1.91E-02 |
| ENSG00000103932 | RPAP1 | 0.53 | 1.20E-03 | 1.91E-02 |
| ENSG00000109111 | SUPT6H | 0.53 | 1.20E-03 | 1.91E-02 |

| Ensembl ID | Gene Name | ρ | Pval | Padj |
|---|---|---|---|---|
| ENSG00000172954 | LCLAT1 | -0.53 | 1.20E-03 | 1.91E-02 |
| ENSG00000213281 | NRAS | -0.53 | 1.20E-03 | 1.91E-02 |
| ENSG00000002919 | SNX11 | -0.52 | 1.21E-03 | 1.92E-02 |
| ENSG00000114686 | MRPL3 | -0.52 | 1.21E-03 | 1.92E-02 |
| ENSG00000133067 | LGR6 | 0.52 | 1.21E-03 | 1.92E-02 |
| ENSG00000136367 | ZFHX2 | 0.52 | 1.21E-03 | 1.92E-02 |
| ENSG00000124635 | HIST1H2BJ | -0.52 | 1.21E-03 | 1.92E-02 |
| ENSG00000072310 | SREBF1 | 0.52 | 1.21E-03 | 1.92E-02 |
| ENSG00000101871 | MID1 | -0.52 | 1.21E-03 | 1.92E-02 |
| ENSG00000116793 | PHTF1 | -0.52 | 1.21E-03 | 1.92E-02 |
| ENSG00000127804 | METTL16 | 0.52 | 1.21E-03 | 1.92E-02 |
| ENSG00000136052 | SLC41A2 | -0.52 | 1.21E-03 | 1.92E-02 |
| ENSG00000153815 | CMIP | -0.52 | 1.21E-03 | 1.92E-02 |
| ENSG00000181915 | ADO | -0.52 | 1.21E-03 | 1.92E-02 |
| ENSG00000147606 | SLC26A7 | 0.52 | 1.22E-03 | 1.92E-02 |
| ENSG00000137491 | SLCO2B1 | -0.52 | 1.22E-03 | 1.92E-02 |
| ENSG00000167461 | RAB8A | -0.52 | 1.22E-03 | 1.92E-02 |
| ENSG00000171793 | CTPS1 | -0.52 | 1.22E-03 | 1.92E-02 |
| ENSG00000147533 | GOLGA7 | -0.52 | 1.23E-03 | 1.93E-02 |
| ENSG00000188010 | MORN2 | -0.52 | 1.23E-03 | 1.93E-02 |
| ENSG00000034971 | MYOC | 0.52 | 1.24E-03 | 1.94E-02 |
| ENSG00000130254 | SAFB2 | 0.52 | 1.24E-03 | 1.94E-02 |
| ENSG00000150779 | TIMM8B | -0.52 | 1.24E-03 | 1.94E-02 |
| ENSG00000106952 | TNFSF8 | -0.52 | 1.25E-03 | 1.95E-02 |
| ENSG00000116459 | ATP5PB | -0.52 | 1.25E-03 | 1.95E-02 |
| ENSG00000213699 | SLC35F6 | -0.52 | 1.25E-03 | 1.95E-02 |
| ENSG00000278177 | AL354811.1 | 0.52 | 1.25E-03 | 1.95E-02 |
| ENSG00000242574 | HLA-DMB | -0.52 | 1.25E-03 | 1.95E-02 |
| ENSG00000087116 | ADAMTS2 | -0.52 | 1.26E-03 | 1.96E-02 |
| ENSG00000100823 | APEX1 | -0.52 | 1.26E-03 | 1.96E-02 |
| ENSG00000131470 | PSMC3IP | -0.52 | 1.26E-03 | 1.96E-02 |
| ENSG00000134575 | ACP2 | -0.52 | 1.26E-03 | 1.96E-02 |
| ENSG00000229886 | AC068533.2 | 0.52 | 1.26E-03 | 1.96E-02 |
| ENSG00000269416 | LINC01224 | 0.52 | 1.26E-03 | 1.96E-02 |
| ENSG00000196917 | HCAR1 | 0.52 | 1.27E-03 | 1.96E-02 |
| ENSG00000273148 | LINC00653 | 0.52 | 1.27E-03 | 1.96E-02 |
| ENSG00000124507 | PACSIN1 | 0.52 | 1.27E-03 | 1.97E-02 |
| ENSG00000063127 | SLC6A16 | 0.52 | 1.28E-03 | 1.97E-02 |
| ENSG00000135956 | TMEM127 | -0.52 | 1.28E-03 | 1.97E-02 |
| ENSG00000175581 | MRPL48 | -0.52 | 1.28E-03 | 1.97E-02 |
| ENSG00000219607 | PPP1R3G | 0.52 | 1.28E-03 | 1.97E-02 |
| ENSG00000247735 | AC120114.1 | 0.52 | 1.28E-03 | 1.98E-02 |
| ENSG00000070214 | SLC44A1 | -0.52 | 1.29E-03 | 1.98E-02 |
| ENSG00000108947 | EFNB3 | -0.52 | 1.29E-03 | 1.98E-02 |
| ENSG00000196352 | CD55 | -0.52 | 1.29E-03 | 1.98E-02 |

| Ensembl ID | Gene Name | ρ | Pval | Padj |
|---|---|---|---|---|
| ENSG00000259676 | AC103982.1 | 0.52 | 1.29E-03 | 1.98E-02 |
| ENSG00000130695 | CEP85 | 0.52 | 1.29E-03 | 1.99E-02 |
| ENSG00000067064 | IDI1 | -0.52 | 1.30E-03 | 2.00E-02 |
| ENSG00000130787 | HIP1R | 0.52 | 1.30E-03 | 2.00E-02 |
| ENSG00000156968 | MPV17L | 0.52 | 1.30E-03 | 2.00E-02 |
| ENSG00000198768 | APCDD1L | -0.52 | 1.30E-03 | 2.00E-02 |
| ENSG00000169813 | HNRNPF | -0.52 | 1.31E-03 | 2.01E-02 |
| ENSG00000228327 | AL669831.1 | 0.52 | 1.31E-03 | 2.01E-02 |
| ENSG00000004059 | ARF5 | -0.52 | 1.32E-03 | 2.02E-02 |
| ENSG00000004779 | NDUFAB1 | -0.52 | 1.32E-03 | 2.02E-02 |
| ENSG00000144749 | LRIG1 | 0.52 | 1.32E-03 | 2.02E-02 |
| ENSG00000162909 | CAPN2 | -0.52 | 1.32E-03 | 2.02E-02 |
| ENSG00000248019 | FAM13A-AS1 | 0.52 | 1.32E-03 | 2.02E-02 |
| ENSG00000275645 | AC068338.3 | 0.52 | 1.32E-03 | 2.02E-02 |
| ENSG00000135776 | ABCB10 | -0.52 | 1.33E-03 | 2.02E-02 |
| ENSG00000162980 | ARL5A | -0.52 | 1.33E-03 | 2.02E-02 |
| ENSG00000198625 | MDM4 | 0.52 | 1.33E-03 | 2.02E-02 |
| ENSG00000215386 | MIR99AHG | 0.52 | 1.33E-03 | 2.02E-02 |
| ENSG00000251003 | ZFPM2-AS1 | -0.52 | 1.33E-03 | 2.02E-02 |
| ENSG00000129538 | RNASE1 | -0.52 | 1.34E-03 | 2.03E-02 |
| ENSG00000133083 | DCLK1 | -0.52 | 1.34E-03 | 2.03E-02 |
| ENSG00000239467 | AC007405.3 | 0.52 | 1.34E-03 | 2.03E-02 |
| ENSG00000259721 | AC090877.2 | 0.52 | 1.34E-03 | 2.03E-02 |
| ENSG00000103510 | KAT8 | 0.52 | 1.35E-03 | 2.04E-02 |
| ENSG00000122778 | KIAA1549 | 0.52 | 1.35E-03 | 2.04E-02 |
| ENSG00000165934 | CPSF2 | -0.52 | 1.35E-03 | 2.04E-02 |
| ENSG00000171421 | MRPL36 | -0.52 | 1.35E-03 | 2.04E-02 |
| ENSG00000181523 | SGSH | 0.52 | 1.35E-03 | 2.04E-02 |
| ENSG00000272894 | AC004982.2 | 0.52 | 1.36E-03 | 2.04E-02 |
| ENSG00000178772 | CPN2 | -0.52 | 1.36E-03 | 2.04E-02 |
| ENSG00000234492 | RPL34-AS1 | 0.52 | 1.37E-03 | 2.06E-02 |
| ENSG00000198189 | HSD17B11 | -0.52 | 1.37E-03 | 2.06E-02 |
| ENSG00000116898 | MRPS15 | -0.52 | 1.37E-03 | 2.06E-02 |
| ENSG00000186193 | SAPCD2 | -0.52 | 1.37E-03 | 2.06E-02 |
| ENSG00000050555 | LAMC3 | 0.52 | 1.38E-03 | 2.06E-02 |
| ENSG00000129194 | SOX15 | 0.52 | 1.38E-03 | 2.06E-02 |
| ENSG00000145685 | LHFPL2 | -0.52 | 1.38E-03 | 2.06E-02 |
| ENSG00000203709 | MIR29B2CHG | 0.52 | 1.38E-03 | 2.06E-02 |
| ENSG00000234899 | SOX9-AS1 | 0.52 | 1.38E-03 | 2.06E-02 |
| ENSG00000149084 | HSD17B12 | -0.52 | 1.39E-03 | 2.08E-02 |
| ENSG00000140307 | GTF2A2 | -0.52 | 1.40E-03 | 2.09E-02 |
| ENSG00000155659 | VSIG4 | -0.52 | 1.40E-03 | 2.09E-02 |
| ENSG00000198915 | RASGEF1A | 0.52 | 1.40E-03 | 2.09E-02 |
| ENSG00000127564 | PKMYT1 | -0.52 | 1.41E-03 | 2.09E-02 |
| ENSG00000139291 | TMEM19 | -0.52 | 1.41E-03 | 2.09E-02 |

| Ensembl ID | Gene Name | ρ | Pval | Padj |
|---|---|---|---|---|
| ENSG00000162688 | AGL | -0.52 | 1.41E-03 | 2.09E-02 |
| ENSG00000164845 | FAM86FP | 0.52 | 1.41E-03 | 2.09E-02 |
| ENSG00000186564 | FOXD2 | 0.52 | 1.41E-03 | 2.09E-02 |
| ENSG00000134013 | LOXL2 | -0.52 | 1.42E-03 | 2.10E-02 |
| ENSG00000197320 | AC060834.1 | 0.52 | 1.42E-03 | 2.10E-02 |
| ENSG00000273230 | AC102953.2 | 0.52 | 1.42E-03 | 2.10E-02 |
| ENSG00000183496 | MEX3B | -0.52 | 1.43E-03 | 2.11E-02 |
| ENSG00000081377 | CDC14B | -0.52 | 1.43E-03 | 2.11E-02 |
| ENSG00000197093 | GAL3ST4 | -0.52 | 1.43E-03 | 2.11E-02 |
| ENSG00000224424 | PRKAR2A-AS1 | 0.52 | 1.43E-03 | 2.11E-02 |
| ENSG00000011347 | SYT7 | -0.52 | 1.43E-03 | 2.12E-02 |
| ENSG00000139410 | SDSL | -0.52 | 1.44E-03 | 2.12E-02 |
| ENSG00000121057 | AKAP1 | 0.52 | 1.44E-03 | 2.12E-02 |
| ENSG00000231394 | AC099681.2 | 0.52 | 1.45E-03 | 2.12E-02 |
| ENSG00000006625 | GGCT | -0.52 | 1.45E-03 | 2.12E-02 |
| ENSG00000141741 | MIEN1 | -0.52 | 1.45E-03 | 2.12E-02 |
| ENSG00000144152 | FBLN7 | 0.52 | 1.45E-03 | 2.12E-02 |
| ENSG00000174977 | AC026271.1 | -0.52 | 1.45E-03 | 2.12E-02 |
| ENSG00000180592 | SKIDA1 | 0.52 | 1.45E-03 | 2.12E-02 |
| ENSG00000186532 | SMYD4 | 0.52 | 1.45E-03 | 2.12E-02 |
| ENSG00000211445 | GPX3 | 0.52 | 1.45E-03 | 2.12E-02 |
| ENSG00000123892 | RAB38 | -0.52 | 1.46E-03 | 2.13E-02 |
| ENSG00000170113 | NIPA1 | -0.52 | 1.46E-03 | 2.13E-02 |
| ENSG00000171202 | TMEM126A | -0.52 | 1.46E-03 | 2.13E-02 |
| ENSG00000182117 | NOP10 | -0.52 | 1.46E-03 | 2.13E-02 |
| ENSG00000283050 | GTF2IP12 | 0.52 | 1.46E-03 | 2.13E-02 |
| ENSG00000106683 | LIMK1 | -0.52 | 1.47E-03 | 2.14E-02 |
| ENSG00000168256 | NKIRAS2 | -0.52 | 1.47E-03 | 2.14E-02 |
| ENSG00000250208 | FZD10-AS1 | -0.52 | 1.47E-03 | 2.14E-02 |
| ENSG00000058262 | SEC61A1 | -0.52 | 1.48E-03 | 2.15E-02 |
| ENSG00000108578 | BLMH | -0.52 | 1.49E-03 | 2.16E-02 |
| ENSG00000110092 | CCND1 | -0.52 | 1.49E-03 | 2.16E-02 |
| ENSG00000157227 | MMP14 | -0.52 | 1.49E-03 | 2.16E-02 |
| ENSG00000165458 | INPPL1 | 0.52 | 1.49E-03 | 2.16E-02 |
| ENSG00000170802 | FOXN2 | -0.52 | 1.49E-03 | 2.16E-02 |
| ENSG00000214357 | NEURL1B | -0.52 | 1.49E-03 | 2.16E-02 |
| ENSG00000154277 | UCHL1 | -0.52 | 1.50E-03 | 2.17E-02 |
| ENSG00000182963 | GJC1 | -0.52 | 1.50E-03 | 2.17E-02 |
| ENSG00000140941 | MAP1LC3B | -0.52 | 1.50E-03 | 2.17E-02 |
| ENSG00000154723 | ATP5PF | -0.52 | 1.50E-03 | 2.17E-02 |
| ENSG00000163541 | SUCLG1 | -0.52 | 1.50E-03 | 2.17E-02 |
| ENSG00000213478 | CFL1P2 | -0.52 | 1.51E-03 | 2.17E-02 |
| ENSG00000103342 | GSPT1 | -0.52 | 1.51E-03 | 2.17E-02 |
| ENSG00000109738 | GLRB | -0.52 | 1.51E-03 | 2.17E-02 |
| ENSG00000130066 | SAT1 | -0.52 | 1.51E-03 | 2.17E-02 |

| Ensembl ID | Gene Name | ρ | Pval | Padj |
|---|---|---|---|---|
| ENSG00000213341 | CHUK | -0.52 | 1.51E-03 | 2.17E-02 |
| ENSG00000101115 | SALL4 | 0.52 | 1.51E-03 | 2.18E-02 |
| ENSG00000001460 | STPG1 | -0.52 | 1.52E-03 | 2.18E-02 |
| ENSG00000070950 | RAD18 | -0.52 | 1.52E-03 | 2.18E-02 |
| ENSG00000083812 | ZNF324 | 0.52 | 1.52E-03 | 2.18E-02 |
| ENSG00000176720 | BOK | 0.52 | 1.52E-03 | 2.18E-02 |
| ENSG00000161664 | ASB16 | 0.52 | 1.52E-03 | 2.18E-02 |
| ENSG00000100105 | PATZ1 | 0.52 | 1.53E-03 | 2.19E-02 |
| ENSG00000146425 | DYNLT1 | -0.52 | 1.53E-03 | 2.19E-02 |
| ENSG00000184007 | PTP4A2 | -0.52 | 1.53E-03 | 2.19E-02 |
| ENSG00000204287 | HLA-DRA | -0.52 | 1.53E-03 | 2.19E-02 |
| ENSG00000266904 | LINC00663 | 0.52 | 1.53E-03 | 2.19E-02 |
| ENSG00000237950 | AL357079.1 | 0.52 | 1.53E-03 | 2.19E-02 |
| ENSG00000015479 | MATR3 | 0.52 | 1.54E-03 | 2.19E-02 |
| ENSG00000101335 | MYL9 | -0.52 | 1.54E-03 | 2.19E-02 |
| ENSG00000139343 | SNRPF | -0.52 | 1.54E-03 | 2.19E-02 |
| ENSG00000182512 | GLRX5 | -0.52 | 1.54E-03 | 2.19E-02 |
| ENSG00000115233 | PSMD14 | -0.51 | 1.55E-03 | 2.21E-02 |
| ENSG00000145088 | EAF2 | -0.51 | 1.55E-03 | 2.21E-02 |
| ENSG00000267702 | AP005131.6 | 0.51 | 1.55E-03 | 2.21E-02 |
| ENSG00000169413 | RNASE6 | -0.51 | 1.56E-03 | 2.22E-02 |
| ENSG00000099875 | MKNK2 | 0.51 | 1.56E-03 | 2.22E-02 |
| ENSG00000235194 | PPP1R3E | 0.51 | 1.56E-03 | 2.22E-02 |
| ENSG00000068489 | PRR11 | -0.51 | 1.57E-03 | 2.23E-02 |
| ENSG00000128849 | CGNL1 | 0.51 | 1.57E-03 | 2.23E-02 |
| ENSG00000163803 | PLB1 | 0.51 | 1.57E-03 | 2.23E-02 |
| ENSG00000054116 | TRAPPC3 | -0.51 | 1.59E-03 | 2.23E-02 |
| ENSG00000143158 | MPC2 | -0.51 | 1.59E-03 | 2.23E-02 |
| ENSG00000158769 | F11R | -0.51 | 1.59E-03 | 2.23E-02 |
| ENSG00000196754 | S100A2 | -0.51 | 1.59E-03 | 2.23E-02 |
| ENSG00000225138 | SLC9A3-AS1 | 0.51 | 1.59E-03 | 2.23E-02 |
| ENSG00000228544 | CCDC183-AS1 | 0.51 | 1.59E-03 | 2.23E-02 |
| ENSG00000257267 | ZNF271P | -0.51 | 1.59E-03 | 2.23E-02 |
| ENSG00000091592 | NLRP1 | 0.51 | 1.61E-03 | 2.26E-02 |
| ENSG00000116212 | LRRC42 | -0.51 | 1.62E-03 | 2.27E-02 |
| ENSG00000188352 | FOCAD | -0.51 | 1.62E-03 | 2.27E-02 |
| ENSG00000102996 | MMP15 | -0.51 | 1.63E-03 | 2.29E-02 |
| ENSG00000123342 | MMP19 | -0.51 | 1.63E-03 | 2.29E-02 |
| ENSG00000279619 | AC020907.5 | 0.51 | 1.64E-03 | 2.29E-02 |
| ENSG00000100330 | MTMR3 | 0.51 | 1.64E-03 | 2.29E-02 |
| ENSG00000101440 | ASIP | 0.51 | 1.64E-03 | 2.29E-02 |
| ENSG00000131148 | EMC8 | -0.51 | 1.64E-03 | 2.29E-02 |
| ENSG00000176014 | TUBB6 | -0.51 | 1.64E-03 | 2.29E-02 |
| ENSG00000198799 | LRIG2 | 0.51 | 1.64E-03 | 2.29E-02 |
| ENSG00000253671 | AC027117.1 | 0.51 | 1.64E-03 | 2.29E-02 |

| Ensembl ID | Gene Name | ρ | Pval | Padj |
|---|---|---|---|---|
| ENSG00000205221 | VIT | 0.51 | 1.65E-03 | 2.30E-02 |
| ENSG00000166257 | SCN3B | 0.51 | 1.66E-03 | 2.31E-02 |
| ENSG00000182158 | CREB3L2 | 0.51 | 1.66E-03 | 2.31E-02 |
| ENSG00000226711 | FAM66C | 0.51 | 1.66E-03 | 2.31E-02 |
| ENSG00000230454 | U73166.1 | 0.51 | 1.66E-03 | 2.31E-02 |
| ENSG00000110427 | KIAA1549L | -0.51 | 1.67E-03 | 2.32E-02 |
| ENSG00000103018 | CYB5B | -0.51 | 1.68E-03 | 2.32E-02 |
| ENSG00000123427 | EEF1AKMT3 | 0.51 | 1.68E-03 | 2.32E-02 |
| ENSG00000205581 | HMGN1 | -0.51 | 1.68E-03 | 2.32E-02 |
| ENSG00000132305 | IMMT | -0.51 | 1.69E-03 | 2.33E-02 |
| ENSG00000141452 | RMC1 | -0.51 | 1.69E-03 | 2.33E-02 |
| ENSG00000198715 | GLMP | -0.51 | 1.69E-03 | 2.33E-02 |
| ENSG00000245468 | LINC02447 | 0.51 | 1.69E-03 | 2.34E-02 |
| ENSG00000119638 | NEK9 | 0.51 | 1.70E-03 | 2.34E-02 |
| ENSG00000121310 | ECHDC2 | 0.51 | 1.70E-03 | 2.34E-02 |
| ENSG00000138944 | SHISAL1 | -0.51 | 1.70E-03 | 2.34E-02 |
| ENSG00000151623 | NR3C2 | 0.51 | 1.70E-03 | 2.34E-02 |
| ENSG00000197818 | SLC9A8 | 0.51 | 1.70E-03 | 2.34E-02 |
| ENSG00000215908 | CROCCP2 | 0.51 | 1.70E-03 | 2.34E-02 |
| ENSG00000244509 | APOBEC3C | -0.51 | 1.70E-03 | 2.34E-02 |
| ENSG00000156920 | ADGRG4 | 0.51 | 1.70E-03 | 2.34E-02 |
| ENSG00000133800 | LYVE1 | -0.51 | 1.71E-03 | 2.35E-02 |
| ENSG00000146674 | IGFBP3 | -0.51 | 1.71E-03 | 2.35E-02 |
| ENSG00000143476 | DTL | -0.51 | 1.72E-03 | 2.36E-02 |
| ENSG00000111229 | ARPC3 | -0.51 | 1.72E-03 | 2.36E-02 |
| ENSG00000116273 | PHF13 | 0.51 | 1.72E-03 | 2.36E-02 |
| ENSG00000125089 | SH3TC1 | 0.51 | 1.72E-03 | 2.36E-02 |
| ENSG00000182240 | BACE2 | -0.51 | 1.72E-03 | 2.36E-02 |
| ENSG00000213601 | KRT18P19 | 0.51 | 1.72E-03 | 2.36E-02 |
| ENSG00000272870 | AC097534.2 | 0.51 | 1.73E-03 | 2.37E-02 |
| ENSG00000108511 | HOXB6 | -0.51 | 1.74E-03 | 2.38E-02 |
| ENSG00000102393 | GLA | -0.51 | 1.75E-03 | 2.38E-02 |
| ENSG00000115641 | FHL2 | -0.51 | 1.75E-03 | 2.38E-02 |
| ENSG00000175505 | CLCF1 | -0.51 | 1.75E-03 | 2.38E-02 |
| ENSG00000182473 | EXOC7 | 0.51 | 1.75E-03 | 2.38E-02 |
| ENSG00000100027 | YPEL1 | -0.51 | 1.76E-03 | 2.39E-02 |
| ENSG00000146021 | KLHL3 | 0.51 | 1.76E-03 | 2.39E-02 |
| ENSG00000227008 | AL009174.1 | -0.51 | 1.76E-03 | 2.39E-02 |
| ENSG00000247095 | MIR210HG | 0.51 | 1.76E-03 | 2.39E-02 |
| ENSG00000107758 | PPP3CB | -0.51 | 1.77E-03 | 2.40E-02 |
| ENSG00000124313 | IQSEC2 | 0.51 | 1.77E-03 | 2.40E-02 |
| ENSG00000135698 | MPHOSPH6 | -0.51 | 1.77E-03 | 2.40E-02 |
| ENSG00000170209 | ANKK1 | 0.51 | 1.78E-03 | 2.41E-02 |
| ENSG00000039560 | RAI14 | -0.51 | 1.78E-03 | 2.41E-02 |
| ENSG00000110057 | UNC93B1 | -0.51 | 1.78E-03 | 2.41E-02 |

| Ensembl ID | Gene Name | ρ | Pval | Padj |
|---|---|---|---|---|
| ENSG00000177042 | TMEM80 | 0.51 | 1.78E-03 | 2.41E-02 |
| ENSG00000233695 | GAS6-AS1 | 0.51 | 1.78E-03 | 2.41E-02 |
| ENSG00000236535 | RC3H1-IT1 | 0.51 | 1.78E-03 | 2.41E-02 |
| ENSG00000279407 | AC007191.1 | 0.51 | 1.79E-03 | 2.42E-02 |
| ENSG00000140043 | PTGR2 | 0.51 | 1.79E-03 | 2.42E-02 |
| ENSG00000146411 | SLC2A12 | -0.51 | 1.79E-03 | 2.42E-02 |
| ENSG00000113971 | NPHP3 | 0.51 | 1.81E-03 | 2.43E-02 |
| ENSG00000142634 | EFHD2 | -0.51 | 1.81E-03 | 2.43E-02 |
| ENSG00000182141 | ZNF708 | 0.51 | 1.81E-03 | 2.43E-02 |
| ENSG00000115165 | CYTIP | -0.51 | 1.81E-03 | 2.44E-02 |
| ENSG00000146963 | LUC7L2 | 0.51 | 1.82E-03 | 2.44E-02 |
| ENSG00000197406 | DIO3 | -0.51 | 1.82E-03 | 2.44E-02 |
| ENSG00000259856 | RAB43P1 | 0.51 | 1.82E-03 | 2.44E-02 |
| ENSG00000076984 | MAP2K7 | 0.51 | 1.83E-03 | 2.44E-02 |
| ENSG00000101132 | PFDN4 | -0.51 | 1.83E-03 | 2.44E-02 |
| ENSG00000139737 | SLAIN1 | 0.51 | 1.83E-03 | 2.44E-02 |
| ENSG00000151617 | EDNRA | -0.51 | 1.83E-03 | 2.44E-02 |
| ENSG00000164619 | BMPER | 0.51 | 1.83E-03 | 2.44E-02 |
| ENSG00000173372 | C1QA | -0.51 | 1.83E-03 | 2.44E-02 |
| ENSG00000197265 | GTF2E2 | -0.51 | 1.83E-03 | 2.44E-02 |
| ENSG00000272293 | AC083964.1 | -0.51 | 1.84E-03 | 2.45E-02 |
| ENSG00000067208 | EVI5 | -0.51 | 1.84E-03 | 2.45E-02 |
| ENSG00000088766 | CRLS1 | -0.51 | 1.84E-03 | 2.45E-02 |
| ENSG00000114999 | TTL | -0.51 | 1.84E-03 | 2.45E-02 |
| ENSG00000162687 | KCNT2 | 0.51 | 1.84E-03 | 2.45E-02 |
| ENSG00000280693 | SH3PXD2A-AS1 | -0.51 | 1.84E-03 | 2.45E-02 |
| ENSG00000154839 | SKA1 | -0.51 | 1.85E-03 | 2.46E-02 |
| ENSG00000205740 | AL359878.1 | 0.51 | 1.85E-03 | 2.46E-02 |
| ENSG00000053371 | AKR7A2 | -0.51 | 1.86E-03 | 2.46E-02 |
| ENSG00000142507 | PSMB6 | -0.51 | 1.86E-03 | 2.46E-02 |
| ENSG00000144034 | TPRKB | -0.51 | 1.86E-03 | 2.46E-02 |
| ENSG00000171291 | ZNF439 | 0.51 | 1.86E-03 | 2.46E-02 |
| ENSG00000108963 | DPH1 | 0.51 | 1.87E-03 | 2.47E-02 |
| ENSG00000149177 | PTPRJ | -0.51 | 1.87E-03 | 2.47E-02 |
| ENSG00000186432 | KPNA4 | -0.51 | 1.87E-03 | 2.47E-02 |
| ENSG00000264578 | AC009630.3 | -0.51 | 1.88E-03 | 2.48E-02 |
| ENSG00000133657 | ATP13A3 | -0.51 | 1.88E-03 | 2.48E-02 |
| ENSG00000139173 | TMEM117 | -0.51 | 1.88E-03 | 2.48E-02 |
| ENSG00000189043 | NDUFA4 | -0.51 | 1.88E-03 | 2.48E-02 |
| ENSG00000246174 | KCTD21-AS1 | 0.51 | 1.88E-03 | 2.48E-02 |
| ENSG00000174611 | KY | 0.51 | 1.88E-03 | 2.48E-02 |
| ENSG00000232593 | KANTR | 0.51 | 1.89E-03 | 2.48E-02 |
| ENSG00000100739 | BDKRB1 | -0.51 | 1.89E-03 | 2.48E-02 |
| ENSG00000101198 | NKAIN4 | -0.51 | 1.89E-03 | 2.48E-02 |
| ENSG00000002746 | HECW1 | 0.51 | 1.89E-03 | 2.48E-02 |

| Ensembl ID | Gene Name | ρ | Pval | Padj |
|---|---|---|---|---|
| ENSG00000101605 | MYOM1 | 0.51 | 1.89E-03 | 2.48E-02 |
| ENSG00000168398 | BDKRB2 | -0.51 | 1.89E-03 | 2.48E-02 |
| ENSG00000173218 | VANGL1 | -0.51 | 1.89E-03 | 2.48E-02 |
| ENSG00000177119 | ANO6 | -0.51 | 1.89E-03 | 2.48E-02 |
| ENSG00000259660 | DNM1P47 | 0.51 | 1.90E-03 | 2.48E-02 |
| ENSG00000214391 | TUBAP2 | -0.51 | 1.91E-03 | 2.48E-02 |
| ENSG00000080608 | PUM3 | -0.51 | 1.91E-03 | 2.48E-02 |
| ENSG00000115363 | EVA1A | -0.51 | 1.91E-03 | 2.48E-02 |
| ENSG00000162636 | FAM102B | -0.51 | 1.91E-03 | 2.48E-02 |
| ENSG00000198131 | ZNF544 | -0.51 | 1.91E-03 | 2.48E-02 |
| ENSG00000249306 | LINC01411 | -0.51 | 1.91E-03 | 2.48E-02 |
| ENSG00000260267 | AC026471.1 | 0.51 | 1.91E-03 | 2.48E-02 |
| ENSG00000166763 | STRCP1 | 0.51 | 1.92E-03 | 2.50E-02 |
| ENSG00000066629 | EML1 | -0.51 | 1.92E-03 | 2.50E-02 |
| ENSG00000145494 | NDUFS6 | -0.51 | 1.92E-03 | 2.50E-02 |
| ENSG00000113211 | PCDHB6 | 0.51 | 1.93E-03 | 2.51E-02 |
| ENSG00000137460 | FHDC1 | 0.51 | 1.93E-03 | 2.51E-02 |
| ENSG00000165501 | LRR1 | -0.51 | 1.93E-03 | 2.51E-02 |
| ENSG00000170571 | EMB | -0.51 | 1.93E-03 | 2.51E-02 |
| ENSG00000174004 | NRROS | -0.51 | 1.93E-03 | 2.51E-02 |
| ENSG00000253854 | AC010834.3 | 0.51 | 1.94E-03 | 2.51E-02 |
| ENSG00000108557 | RAI1 | 0.51 | 1.95E-03 | 2.51E-02 |
| ENSG00000116151 | MORN1 | 0.51 | 1.95E-03 | 2.51E-02 |
| ENSG00000155926 | SLA | -0.51 | 1.95E-03 | 2.51E-02 |
| ENSG00000158122 | PRXL2C | -0.51 | 1.95E-03 | 2.51E-02 |
| ENSG00000183098 | GPC6 | 0.51 | 1.95E-03 | 2.51E-02 |
| ENSG00000267904 | AC024075.1 | 0.51 | 1.95E-03 | 2.51E-02 |
| ENSG00000262312 | AC004494.1 | 0.51 | 1.95E-03 | 2.51E-02 |
| ENSG00000117505 | DR1 | -0.51 | 1.96E-03 | 2.52E-02 |
| ENSG00000164086 | DUSP7 | 0.51 | 1.96E-03 | 2.52E-02 |
| ENSG00000171533 | MAP6 | 0.51 | 1.96E-03 | 2.52E-02 |
| ENSG00000236383 | CCDC200 | 0.51 | 1.97E-03 | 2.53E-02 |
| ENSG00000070882 | OSBPL3 | -0.51 | 1.97E-03 | 2.54E-02 |
| ENSG00000222047 | C10orf55 | -0.50 | 1.99E-03 | 2.55E-02 |
| ENSG00000100804 | PSMB5 | -0.50 | 1.99E-03 | 2.55E-02 |
| ENSG00000110583 | NAA40 | 0.50 | 1.99E-03 | 2.55E-02 |
| ENSG00000156052 | GNAQ | -0.50 | 2.00E-03 | 2.56E-02 |
| ENSG00000160633 | SAFB | 0.50 | 2.00E-03 | 2.56E-02 |
| ENSG00000181035 | SLC25A42 | 0.50 | 2.00E-03 | 2.56E-02 |
| ENSG00000196510 | ANAPC7 | -0.50 | 2.00E-03 | 2.56E-02 |
| ENSG00000233276 | GPX1 | -0.50 | 2.00E-03 | 2.56E-02 |
| ENSG00000091831 | ESR1 | 0.50 | 2.01E-03 | 2.57E-02 |
| ENSG00000233421 | LINC01783 | 0.50 | 2.01E-03 | 2.57E-02 |
| ENSG00000229893 | AC005091.1 | 0.50 | 2.02E-03 | 2.57E-02 |
| ENSG00000145002 | FAM86B2 | 0.50 | 2.03E-03 | 2.58E-02 |

| Ensembl ID | Gene Name | ρ | Pval | Padj |
|---|---|---|---|---|
| ENSG00000055070 | SZRD1 | -0.50 | 2.03E-03 | 2.58E-02 |
| ENSG00000115484 | CCT4 | -0.50 | 2.03E-03 | 2.58E-02 |
| ENSG00000123643 | SLC36A1 | -0.50 | 2.03E-03 | 2.58E-02 |
| ENSG00000197956 | S100A6 | -0.50 | 2.03E-03 | 2.58E-02 |
| ENSG00000279048 | AC080080.1 | 0.50 | 2.03E-03 | 2.58E-02 |
| ENSG00000007264 | MATK | -0.50 | 2.04E-03 | 2.60E-02 |
| ENSG00000047315 | POLR2B | -0.50 | 2.05E-03 | 2.60E-02 |
| ENSG00000072201 | LNX1 | -0.50 | 2.05E-03 | 2.60E-02 |
| ENSG00000164284 | GRPEL2 | -0.50 | 2.05E-03 | 2.60E-02 |
| ENSG00000182742 | HOXB4 | -0.50 | 2.06E-03 | 2.61E-02 |
| ENSG00000067606 | PRKCZ | 0.50 | 2.07E-03 | 2.61E-02 |
| ENSG00000149679 | CABLES2 | 0.50 | 2.07E-03 | 2.61E-02 |
| ENSG00000158258 | CLSTN2 | -0.50 | 2.07E-03 | 2.61E-02 |
| ENSG00000158470 | B4GALT5 | -0.50 | 2.07E-03 | 2.61E-02 |
| ENSG00000162337 | LRP5 | 0.50 | 2.07E-03 | 2.61E-02 |
| ENSG00000166130 | IKBIP | -0.50 | 2.07E-03 | 2.61E-02 |
| ENSG00000181619 | GPR135 | 0.50 | 2.07E-03 | 2.61E-02 |
| ENSG00000188747 | NOXA1 | 0.50 | 2.07E-03 | 2.61E-02 |
| ENSG00000224831 | TMEM183B | -0.50 | 2.07E-03 | 2.61E-02 |
| ENSG00000236782 | AL391650.1 | -0.50 | 2.08E-03 | 2.61E-02 |
| ENSG00000241570 | PAQR9-AS1 | 0.50 | 2.08E-03 | 2.61E-02 |
| ENSG00000120438 | TCP1 | -0.50 | 2.08E-03 | 2.61E-02 |
| ENSG00000134153 | EMC7 | -0.50 | 2.08E-03 | 2.61E-02 |
| ENSG00000245025 | AC107959.1 | 0.50 | 2.08E-03 | 2.61E-02 |
| ENSG00000079257 | LXN | -0.50 | 2.10E-03 | 2.63E-02 |
| ENSG00000160752 | FDPS | -0.50 | 2.10E-03 | 2.63E-02 |
| ENSG00000003137 | CYP26B1 | 0.50 | 2.11E-03 | 2.64E-02 |
| ENSG00000047621 | C12orf4 | -0.50 | 2.11E-03 | 2.64E-02 |
| ENSG00000167110 | GOLGA2 | 0.50 | 2.11E-03 | 2.64E-02 |
| ENSG00000280119 | AC093642.2 | 0.50 | 2.12E-03 | 2.65E-02 |
| ENSG00000109854 | HTATIP2 | -0.50 | 2.12E-03 | 2.65E-02 |
| ENSG00000117133 | RPF1 | -0.50 | 2.12E-03 | 2.65E-02 |
| ENSG00000162511 | LAPTM5 | -0.50 | 2.12E-03 | 2.65E-02 |
| ENSG00000197798 | FAM118B | -0.50 | 2.12E-03 | 2.65E-02 |
| ENSG00000108932 | SLC16A6 | -0.50 | 2.13E-03 | 2.66E-02 |
| ENSG00000048545 | GUCA1A | -0.50 | 2.13E-03 | 2.66E-02 |
| ENSG00000100196 | KDELR3 | -0.50 | 2.14E-03 | 2.66E-02 |
| ENSG00000108821 | COL1A1 | -0.50 | 2.14E-03 | 2.66E-02 |
| ENSG00000125398 | SOX9 | 0.50 | 2.14E-03 | 2.66E-02 |
| ENSG00000245556 | SCAMP1-AS1 | 0.50 | 2.14E-03 | 2.66E-02 |
| ENSG00000166323 | C11orf65 | 0.50 | 2.14E-03 | 2.66E-02 |
| ENSG00000079482 | OPHN1 | 0.50 | 2.17E-03 | 2.69E-02 |
| ENSG00000103472 | RRN3P2 | 0.50 | 2.17E-03 | 2.69E-02 |
| ENSG00000123297 | TSFM | -0.50 | 2.17E-03 | 2.69E-02 |
| ENSG00000185418 | TARSL2 | 0.50 | 2.17E-03 | 2.69E-02 |

| Ensembl ID | Gene Name | ρ | Pval | Padj |
|---|---|---|---|---|
| ENSG00000006327 | TNFRSF12A | -0.50 | 2.18E-03 | 2.69E-02 |
| ENSG00000091622 | PITPNM3 | -0.50 | 2.18E-03 | 2.69E-02 |
| ENSG00000115128 | SF3B6 | -0.50 | 2.18E-03 | 2.69E-02 |
| ENSG00000129235 | TXNDC17 | -0.50 | 2.18E-03 | 2.69E-02 |
| ENSG00000130830 | MPP1 | -0.50 | 2.18E-03 | 2.69E-02 |
| ENSG00000166484 | MAPK7 | 0.50 | 2.18E-03 | 2.69E-02 |
| ENSG00000168275 | COA6 | -0.50 | 2.18E-03 | 2.69E-02 |
| ENSG00000250490 | LINC02145 | 0.50 | 2.19E-03 | 2.70E-02 |
| ENSG00000089876 | DHX32 | -0.50 | 2.20E-03 | 2.70E-02 |
| ENSG00000132321 | IQCA1 | 0.50 | 2.20E-03 | 2.70E-02 |
| ENSG00000137193 | PIM1 | -0.50 | 2.20E-03 | 2.70E-02 |
| ENSG00000138646 | HERC5 | 0.50 | 2.20E-03 | 2.70E-02 |
| ENSG00000182195 | LDOC1 | 0.50 | 2.20E-03 | 2.70E-02 |
| ENSG00000182518 | FAM104B | -0.50 | 2.20E-03 | 2.70E-02 |
| ENSG00000279253 | AL121753.2 | 0.50 | 2.20E-03 | 2.70E-02 |
| ENSG00000168477 | TNXB | 0.50 | 2.21E-03 | 2.71E-02 |
| ENSG00000196236 | XPNPEP3 | 0.50 | 2.21E-03 | 2.71E-02 |
| ENSG00000267013 | LINC01929 | -0.50 | 2.22E-03 | 2.72E-02 |
| ENSG00000243498 | UBA52P5 | -0.50 | 2.22E-03 | 2.72E-02 |
| ENSG00000107130 | NCS1 | -0.50 | 2.22E-03 | 2.72E-02 |
| ENSG00000162415 | ZSWIM5 | 0.50 | 2.22E-03 | 2.72E-02 |
| ENSG00000198668 | CALM1 | -0.50 | 2.22E-03 | 2.72E-02 |
| ENSG00000260572 | AC069224.1 | 0.50 | 2.22E-03 | 2.72E-02 |
| ENSG00000204472 | AIF1 | -0.50 | 2.23E-03 | 2.73E-02 |
| ENSG00000058404 | CAMK2B | 0.50 | 2.24E-03 | 2.73E-02 |
| ENSG00000250218 | ALDH1L1-AS1 | 0.50 | 2.24E-03 | 2.73E-02 |
| ENSG00000237437 | ASS1P12 | 0.50 | 2.25E-03 | 2.74E-02 |
| ENSG00000110675 | ELMOD1 | -0.50 | 2.25E-03 | 2.74E-02 |
| ENSG00000111859 | NEDD9 | -0.50 | 2.25E-03 | 2.74E-02 |
| ENSG00000174851 | YIF1A | -0.50 | 2.25E-03 | 2.74E-02 |
| ENSG00000107566 | ERLIN1 | -0.50 | 2.27E-03 | 2.76E-02 |
| ENSG00000131459 | GFPT2 | -0.50 | 2.27E-03 | 2.76E-02 |
| ENSG00000273033 | LINC02035 | 0.50 | 2.27E-03 | 2.76E-02 |
| ENSG00000117616 | RSRP1 | 0.50 | 2.28E-03 | 2.77E-02 |
| ENSG00000157045 | NTAN1 | -0.50 | 2.28E-03 | 2.77E-02 |
| ENSG00000177575 | CD163 | -0.50 | 2.28E-03 | 2.77E-02 |
| ENSG00000255353 | AP003057.1 | 0.50 | 2.28E-03 | 2.77E-02 |
| ENSG00000109758 | HGFAC | 0.50 | 2.29E-03 | 2.78E-02 |
| ENSG00000137288 | UQCC2 | -0.50 | 2.31E-03 | 2.80E-02 |
| ENSG00000149577 | SIDT2 | 0.50 | 2.31E-03 | 2.80E-02 |
| ENSG00000164111 | ANXA5 | -0.50 | 2.31E-03 | 2.80E-02 |
| ENSG00000169057 | MECP2 | 0.50 | 2.31E-03 | 2.80E-02 |
| ENSG00000232303 | DFFBP1 | 0.50 | 2.33E-03 | 2.81E-02 |
| ENSG00000180992 | MRPL14 | -0.50 | 2.33E-03 | 2.81E-02 |
| ENSG00000197375 | SLC22A5 | 0.50 | 2.33E-03 | 2.81E-02 |

| Ensembl ID | Gene Name | ρ | Pval | Padj |
|---|---|---|---|---|
| ENSG00000158186 | MRAS | -0.50 | 2.35E-03 | 2.83E-02 |
| ENSG00000102384 | CENPI | -0.50 | 2.36E-03 | 2.84E-02 |
| ENSG00000065978 | YBX1 | -0.50 | 2.36E-03 | 2.84E-02 |
| ENSG00000104442 | ARMC1 | -0.50 | 2.36E-03 | 2.84E-02 |
| ENSG00000168874 | ATOH8 | 0.50 | 2.36E-03 | 2.84E-02 |
| ENSG00000272391 | POM121C | 0.50 | 2.36E-03 | 2.84E-02 |
| ENSG00000016864 | GLT8D1 | -0.50 | 2.38E-03 | 2.84E-02 |
| ENSG00000112312 | GMNN | -0.50 | 2.38E-03 | 2.84E-02 |
| ENSG00000116604 | MEF2D | 0.50 | 2.38E-03 | 2.84E-02 |
| ENSG00000134248 | LAMTOR5 | -0.50 | 2.38E-03 | 2.84E-02 |
| ENSG00000134717 | BTF3L4 | -0.50 | 2.38E-03 | 2.84E-02 |
| ENSG00000155097 | ATP6V1C1 | -0.50 | 2.38E-03 | 2.84E-02 |
| ENSG00000187189 | TSPYL4 | 0.50 | 2.38E-03 | 2.84E-02 |
| ENSG00000226674 | TEX41 | -0.50 | 2.38E-03 | 2.84E-02 |
| ENSG00000214114 | MYCBP | -0.50 | 2.39E-03 | 2.85E-02 |
| ENSG00000079332 | SAR1A | -0.50 | 2.39E-03 | 2.85E-02 |
| ENSG00000130592 | LSP1 | 0.50 | 2.39E-03 | 2.85E-02 |
| ENSG00000151322 | NPAS3 | 0.50 | 2.39E-03 | 2.85E-02 |
| ENSG00000163788 | SNRK | -0.50 | 2.39E-03 | 2.85E-02 |
| ENSG00000234892 | Z82214.1 | 0.50 | 2.40E-03 | 2.86E-02 |
| ENSG00000181026 | AEN | -0.50 | 2.41E-03 | 2.87E-02 |
| ENSG00000103415 | HMOX2 | -0.50 | 2.42E-03 | 2.89E-02 |
| ENSG00000187323 | DCC | 0.50 | 2.43E-03 | 2.89E-02 |
| ENSG00000258872 | FDPSP3 | 0.50 | 2.44E-03 | 2.89E-02 |
| ENSG00000081320 | STK17B | -0.50 | 2.44E-03 | 2.89E-02 |
| ENSG00000099256 | PRTFDC1 | -0.50 | 2.44E-03 | 2.89E-02 |
| ENSG00000108561 | C1QBP | -0.50 | 2.44E-03 | 2.89E-02 |
| ENSG00000138867 | GUCD1 | -0.50 | 2.44E-03 | 2.89E-02 |
| ENSG00000184574 | LPAR5 | -0.50 | 2.44E-03 | 2.89E-02 |
| ENSG00000197594 | ENPP1 | -0.50 | 2.44E-03 | 2.89E-02 |
| ENSG00000100055 | CYTH4 | -0.50 | 2.46E-03 | 2.90E-02 |
| ENSG00000128039 | SRD5A3 | -0.50 | 2.46E-03 | 2.90E-02 |
| ENSG00000155329 | ZCCHC10 | -0.50 | 2.46E-03 | 2.90E-02 |
| ENSG00000197629 | MPEG1 | -0.50 | 2.47E-03 | 2.92E-02 |
| ENSG00000272386 | AC015802.5 | 0.50 | 2.48E-03 | 2.93E-02 |
| ENSG00000136560 | TANK | -0.50 | 2.49E-03 | 2.93E-02 |
| ENSG00000137575 | SDCBP | -0.50 | 2.49E-03 | 2.93E-02 |
| ENSG00000167363 | FN3K | 0.50 | 2.49E-03 | 2.93E-02 |
| ENSG00000241549 | GUSBP2 | 0.50 | 2.49E-03 | 2.94E-02 |
| ENSG00000043462 | LCP2 | -0.50 | 2.50E-03 | 2.94E-02 |
| ENSG00000074047 | GLI2 | 0.49 | 2.50E-03 | 2.94E-02 |
| ENSG00000135404 | CD63 | -0.49 | 2.50E-03 | 2.94E-02 |
| ENSG00000167513 | CDT1 | -0.49 | 2.50E-03 | 2.94E-02 |
| ENSG00000182004 | SNRPE | -0.49 | 2.50E-03 | 2.94E-02 |
| ENSG00000245060 | LINC00847 | 0.49 | 2.50E-03 | 2.94E-02 |

| Ensembl ID | Gene Name | ρ | Pval | Padj |
|---|---|---|---|---|
| ENSG00000129084 | PSMA1 | -0.49 | 2.52E-03 | 2.96E-02 |
| ENSG00000135940 | COX5B | -0.49 | 2.52E-03 | 2.96E-02 |
| ENSG00000183287 | CCBE1 | -0.49 | 2.53E-03 | 2.97E-02 |
| ENSG00000105877 | DNAH11 | 0.49 | 2.54E-03 | 2.97E-02 |
| ENSG00000179532 | DNHD1 | 0.49 | 2.54E-03 | 2.97E-02 |
| ENSG00000272902 | TBC1D8-AS1 | 0.49 | 2.54E-03 | 2.97E-02 |
| ENSG00000145113 | MUC4 | 0.49 | 2.55E-03 | 2.98E-02 |
| ENSG00000100528 | CNIH1 | -0.49 | 2.55E-03 | 2.98E-02 |
| ENSG00000166794 | PPIB | -0.49 | 2.55E-03 | 2.98E-02 |
| ENSG00000178980 | SELENOW | -0.49 | 2.55E-03 | 2.98E-02 |
| ENSG00000268362 | AC092279.1 | 0.49 | 2.55E-03 | 2.98E-02 |
| ENSG00000270055 | AC127502.2 | 0.49 | 2.55E-03 | 2.98E-02 |
| ENSG00000125726 | CD70 | -0.49 | 2.57E-03 | 2.99E-02 |
| ENSG00000113312 | TTC1 | -0.49 | 2.57E-03 | 2.99E-02 |
| ENSG00000113734 | BNIP1 | -0.49 | 2.57E-03 | 2.99E-02 |
| ENSG00000133460 | SLC2A11 | 0.49 | 2.57E-03 | 2.99E-02 |
| ENSG00000182704 | TSKU | 0.49 | 2.57E-03 | 2.99E-02 |
| ENSG00000008282 | SYPL1 | -0.49 | 2.59E-03 | 2.99E-02 |
| ENSG00000076716 | GPC4 | -0.49 | 2.59E-03 | 2.99E-02 |
| ENSG00000100532 | CGRRF1 | -0.49 | 2.59E-03 | 2.99E-02 |
| ENSG00000108175 | ZMIZ1 | 0.49 | 2.59E-03 | 2.99E-02 |
| ENSG00000186001 | LRCH3 | 0.49 | 2.59E-03 | 2.99E-02 |
| ENSG00000198225 | FKBP1C | -0.49 | 2.59E-03 | 2.99E-02 |
| ENSG00000198931 | APRT | -0.49 | 2.59E-03 | 2.99E-02 |
| ENSG00000219438 | FAM19A5 | -0.49 | 2.59E-03 | 2.99E-02 |
| ENSG00000222041 | CYTOR | -0.49 | 2.59E-03 | 2.99E-02 |
| ENSG00000223745 | CCDC18-AS1 | 0.49 | 2.59E-03 | 2.99E-02 |
| ENSG00000279302 | AC013643.3 | 0.49 | 2.59E-03 | 2.99E-02 |
| ENSG00000184661 | CDCA2 | -0.49 | 2.60E-03 | 2.99E-02 |
| ENSG00000103365 | GGA2 | 0.49 | 2.60E-03 | 2.99E-02 |
| ENSG00000112695 | COX7A2 | -0.49 | 2.60E-03 | 2.99E-02 |
| ENSG00000153165 | RGPD3 | 0.49 | 2.60E-03 | 2.99E-02 |
| ENSG00000161249 | DMKN | 0.49 | 2.60E-03 | 2.99E-02 |
| ENSG00000169071 | ROR2 | 0.49 | 2.60E-03 | 2.99E-02 |
| ENSG00000182351 | CRIP1P4 | -0.49 | 2.60E-03 | 2.99E-02 |
| ENSG00000265158 | LRRC37A7P | 0.49 | 2.60E-03 | 2.99E-02 |
| ENSG00000269609 | RPARP-AS1 | 0.49 | 2.60E-03 | 2.99E-02 |
| ENSG00000272678 | AC112503.1 | 0.49 | 2.61E-03 | 2.99E-02 |
| ENSG00000171530 | TBCA | -0.49 | 2.62E-03 | 3.01E-02 |
| ENSG00000101182 | PSMA7 | -0.49 | 2.64E-03 | 3.02E-02 |
| ENSG00000101493 | ZNF516 | 0.49 | 2.64E-03 | 3.02E-02 |
| ENSG00000187840 | EIF4EBP1 | -0.49 | 2.64E-03 | 3.02E-02 |
| ENSG00000083457 | ITGAE | -0.49 | 2.66E-03 | 3.03E-02 |
| ENSG00000117122 | MFAP2 | -0.49 | 2.66E-03 | 3.03E-02 |
| ENSG00000148484 | RSU1 | -0.49 | 2.66E-03 | 3.03E-02 |

| Ensembl ID | Gene Name | ρ | Pval | Padj |
|---|---|---|---|---|
| ENSG00000240303 | ACAD11 | 0.49 | 2.66E-03 | 3.03E-02 |
| ENSG00000279805 | Z95114.3 | 0.49 | 2.66E-03 | 3.03E-02 |
| ENSG00000081041 | CXCL2 | -0.49 | 2.66E-03 | 3.03E-02 |
| ENSG00000091536 | MYO15A | 0.49 | 2.67E-03 | 3.04E-02 |
| ENSG00000109610 | SOD3 | 0.49 | 2.67E-03 | 3.04E-02 |
| ENSG00000137942 | FNBP1L | -0.49 | 2.67E-03 | 3.04E-02 |
| ENSG00000173369 | C1QB | -0.49 | 2.67E-03 | 3.04E-02 |
| ENSG00000176595 | KBTBD11 | 0.49 | 2.67E-03 | 3.04E-02 |
| ENSG00000253636 | AC022893.1 | 0.49 | 2.67E-03 | 3.04E-02 |
| ENSG00000164520 | RAET1E | -0.49 | 2.70E-03 | 3.07E-02 |
| ENSG00000151388 | ADAMTS12 | -0.49 | 2.70E-03 | 3.07E-02 |
| ENSG00000120697 | ALG5 | -0.49 | 2.71E-03 | 3.07E-02 |
| ENSG00000126746 | ZNF384 | 0.49 | 2.71E-03 | 3.07E-02 |
| ENSG00000127957 | PMS2P3 | 0.49 | 2.71E-03 | 3.07E-02 |
| ENSG00000198429 | ZNF69 | 0.49 | 2.71E-03 | 3.07E-02 |
| ENSG00000226471 | Z93930.2 | 0.49 | 2.71E-03 | 3.07E-02 |
| ENSG00000267185 | PTP4A2P1 | -0.49 | 2.71E-03 | 3.07E-02 |
| ENSG00000078668 | VDAC3 | -0.49 | 2.73E-03 | 3.08E-02 |
| ENSG00000127920 | GNG11 | -0.49 | 2.73E-03 | 3.08E-02 |
| ENSG00000180776 | ZDHHC20 | -0.49 | 2.73E-03 | 3.08E-02 |
| ENSG00000113273 | ARSB | -0.49 | 2.74E-03 | 3.10E-02 |
| ENSG00000171310 | CHST11 | -0.49 | 2.74E-03 | 3.10E-02 |
| ENSG00000176871 | WSB2 | -0.49 | 2.74E-03 | 3.10E-02 |
| ENSG00000182197 | EXT1 | -0.49 | 2.74E-03 | 3.10E-02 |
| ENSG00000134684 | YARS | -0.49 | 2.76E-03 | 3.11E-02 |
| ENSG00000231305 | AC112484.1 | 0.49 | 2.77E-03 | 3.12E-02 |
| ENSG00000110077 | MS4A6A | -0.49 | 2.78E-03 | 3.12E-02 |
| ENSG00000059804 | SLC2A3 | -0.49 | 2.78E-03 | 3.12E-02 |
| ENSG00000135002 | RFK | -0.49 | 2.78E-03 | 3.12E-02 |
| ENSG00000144744 | UBA3 | -0.49 | 2.78E-03 | 3.12E-02 |
| ENSG00000235387 | SPAAR | 0.49 | 2.78E-03 | 3.12E-02 |
| ENSG00000150625 | GPM6A | 0.49 | 2.79E-03 | 3.14E-02 |
| ENSG00000096060 | FKBP5 | 0.49 | 2.80E-03 | 3.14E-02 |
| ENSG00000154359 | LONRF1 | 0.49 | 2.80E-03 | 3.14E-02 |
| ENSG00000204084 | INPP5B | 0.49 | 2.80E-03 | 3.14E-02 |
| ENSG00000228801 | AC064807.1 | 0.49 | 2.80E-03 | 3.14E-02 |
| ENSG00000070010 | UFD1 | -0.49 | 2.81E-03 | 3.15E-02 |
| ENSG00000120526 | NUDCD1 | -0.49 | 2.81E-03 | 3.15E-02 |
| ENSG00000166164 | BRD7 | -0.49 | 2.81E-03 | 3.15E-02 |
| ENSG00000105341 | DMAC2 | -0.49 | 2.83E-03 | 3.17E-02 |
| ENSG00000109790 | KLHL5 | -0.49 | 2.83E-03 | 3.17E-02 |
| ENSG00000277072 | STAG3L2 | 0.49 | 2.83E-03 | 3.17E-02 |
| ENSG00000185862 | EVI2B | -0.49 | 2.85E-03 | 3.17E-02 |
| ENSG00000115461 | IGFBP5 | -0.49 | 2.85E-03 | 3.17E-02 |
| ENSG00000133110 | POSTN | -0.49 | 2.85E-03 | 3.17E-02 |

| Ensembl ID | Gene Name | ρ | Pval | Padj |
|---|---|---|---|---|
| ENSG00000137767 | SQOR | -0.49 | 2.85E-03 | 3.17E-02 |
| ENSG00000146376 | ARHGAP18 | -0.49 | 2.85E-03 | 3.17E-02 |
| ENSG00000173040 | EVC2 | 0.49 | 2.85E-03 | 3.17E-02 |
| ENSG00000198431 | TXNRD1 | -0.49 | 2.85E-03 | 3.17E-02 |
| ENSG00000253771 | TPTE2P1 | 0.49 | 2.85E-03 | 3.17E-02 |
| ENSG00000272970 | AC107294.2 | 0.49 | 2.85E-03 | 3.17E-02 |
| ENSG00000171067 | C11orf24 | -0.49 | 2.87E-03 | 3.19E-02 |
| ENSG00000205853 | RFPL3S | 0.49 | 2.87E-03 | 3.19E-02 |
| ENSG00000260532 | AL031598.1 | 0.49 | 2.88E-03 | 3.20E-02 |
| ENSG00000198354 | DCAF12L2 | 0.49 | 2.89E-03 | 3.20E-02 |
| ENSG00000110442 | COMMD9 | -0.49 | 2.89E-03 | 3.20E-02 |
| ENSG00000135476 | ESPL1 | -0.49 | 2.89E-03 | 3.20E-02 |
| ENSG00000176998 | HCG4 | 0.49 | 2.89E-03 | 3.20E-02 |
| ENSG00000215284 | AL512633.1 | -0.49 | 2.89E-03 | 3.20E-02 |
| ENSG00000081853 | PCDHGA2 | 0.49 | 2.91E-03 | 3.21E-02 |
| ENSG00000085063 | CD59 | -0.49 | 2.91E-03 | 3.21E-02 |
| ENSG00000181467 | RAP2B | -0.49 | 2.91E-03 | 3.21E-02 |
| ENSG00000234390 | USP27X-AS1 | 0.49 | 2.91E-03 | 3.21E-02 |
| ENSG00000241506 | PSMC1P1 | -0.49 | 2.91E-03 | 3.21E-02 |
| ENSG00000273382 | AL356488.3 | 0.49 | 2.91E-03 | 3.21E-02 |
| ENSG00000163739 | CXCL1 | -0.49 | 2.91E-03 | 3.21E-02 |
| ENSG00000134061 | CD180 | -0.49 | 2.91E-03 | 3.21E-02 |
| ENSG00000159423 | ALDH4A1 | -0.49 | 2.92E-03 | 3.21E-02 |
| ENSG00000183856 | IQGAP3 | -0.49 | 2.92E-03 | 3.21E-02 |
| ENSG00000223705 | NSUN5P1 | 0.49 | 2.92E-03 | 3.21E-02 |
| ENSG00000235316 | DUSP8P5 | 0.49 | 2.92E-03 | 3.21E-02 |
| ENSG00000261270 | AC012181.2 | 0.49 | 2.92E-03 | 3.21E-02 |
| ENSG00000171659 | GPR34 | -0.49 | 2.93E-03 | 3.22E-02 |
| ENSG00000008441 | NFIX | 0.49 | 2.94E-03 | 3.23E-02 |
| ENSG00000164087 | POC1A | -0.49 | 2.94E-03 | 3.23E-02 |
| ENSG00000175390 | EIF3F | -0.49 | 2.94E-03 | 3.23E-02 |
| ENSG00000176401 | EID2B | 0.49 | 2.94E-03 | 3.23E-02 |
| ENSG00000146648 | EGFR | 0.49 | 2.96E-03 | 3.24E-02 |
| ENSG00000184588 | PDE4B | -0.49 | 2.96E-03 | 3.24E-02 |
| ENSG00000198585 | NUDT16 | -0.49 | 2.96E-03 | 3.24E-02 |
| ENSG00000262877 | AC110285.2 | 0.49 | 2.96E-03 | 3.24E-02 |
| ENSG00000196735 | HLA-DQA1 | -0.49 | 2.97E-03 | 3.25E-02 |
| ENSG00000134809 | TIMM10 | -0.49 | 2.98E-03 | 3.25E-02 |
| ENSG00000156697 | UTP14A | -0.49 | 2.98E-03 | 3.25E-02 |
| ENSG00000175166 | PSMD2 | -0.49 | 2.98E-03 | 3.25E-02 |
| ENSG00000142945 | KIF2C | -0.49 | 3.00E-03 | 3.27E-02 |
| ENSG00000023902 | PLEKHO1 | -0.49 | 3.02E-03 | 3.29E-02 |
| ENSG00000010256 | UQCRC1 | -0.49 | 3.04E-03 | 3.30E-02 |
| ENSG00000065548 | ZC3H15 | -0.49 | 3.04E-03 | 3.30E-02 |
| ENSG00000107338 | SHB | -0.49 | 3.04E-03 | 3.30E-02 |

| Ensembl ID | Gene Name | ρ | Pval | Padj |
|---|---|---|---|---|
| ENSG00000119013 | NDUFB3 | -0.49 | 3.04E-03 | 3.30E-02 |
| ENSG00000129562 | DAD1 | -0.49 | 3.04E-03 | 3.30E-02 |
| ENSG00000163029 | SMC6 | -0.49 | 3.04E-03 | 3.30E-02 |
| ENSG00000165474 | GJB2 | -0.49 | 3.04E-03 | 3.30E-02 |
| ENSG00000196549 | MME | -0.49 | 3.04E-03 | 3.30E-02 |
| ENSG00000198804 | MT-CO1 | -0.49 | 3.04E-03 | 3.30E-02 |
| ENSG00000104951 | IL4I1 | -0.49 | 3.04E-03 | 3.30E-02 |
| ENSG00000279108 | AC008537.3 | 0.49 | 3.05E-03 | 3.30E-02 |
| ENSG00000139289 | PHLDA1 | -0.49 | 3.06E-03 | 3.31E-02 |
| ENSG00000143612 | C1orf43 | -0.49 | 3.06E-03 | 3.31E-02 |
| ENSG00000164405 | UQCRQ | -0.49 | 3.06E-03 | 3.31E-02 |
| ENSG00000181929 | PRKAG1 | -0.49 | 3.06E-03 | 3.31E-02 |
| ENSG00000183154 | AC138356.1 | 0.49 | 3.08E-03 | 3.32E-02 |
| ENSG00000106344 | RBM28 | 0.49 | 3.08E-03 | 3.32E-02 |
| ENSG00000103489 | XYLT1 | 0.49 | 3.10E-03 | 3.34E-02 |
| ENSG00000185947 | ZNF267 | -0.49 | 3.10E-03 | 3.34E-02 |
| ENSG00000260942 | CAPN10-DT | 0.49 | 3.10E-03 | 3.34E-02 |
| ENSG00000145864 | GABRB2 | -0.49 | 3.10E-03 | 3.34E-02 |
| ENSG00000030110 | BAK1 | -0.49 | 3.12E-03 | 3.35E-02 |
| ENSG00000043093 | DCUN1D1 | -0.49 | 3.12E-03 | 3.35E-02 |
| ENSG00000120049 | KCNIP2 | 0.49 | 3.12E-03 | 3.35E-02 |
| ENSG00000131015 | ULBP2 | -0.49 | 3.12E-03 | 3.35E-02 |
| ENSG00000156802 | ATAD2 | -0.49 | 3.12E-03 | 3.35E-02 |
| ENSG00000260805 | AC092803.2 | 0.49 | 3.13E-03 | 3.35E-02 |
| ENSG00000023734 | STRAP | -0.49 | 3.14E-03 | 3.35E-02 |
| ENSG00000026103 | FAS | -0.49 | 3.14E-03 | 3.35E-02 |
| ENSG00000125249 | RAP2A | -0.49 | 3.14E-03 | 3.35E-02 |
| ENSG00000139044 | B4GALNT3 | 0.49 | 3.14E-03 | 3.35E-02 |
| ENSG00000151665 | PIGF | -0.49 | 3.14E-03 | 3.35E-02 |
| ENSG00000162745 | OLFML2B | -0.49 | 3.14E-03 | 3.35E-02 |
| ENSG00000163354 | DCST2 | 0.49 | 3.14E-03 | 3.35E-02 |
| ENSG00000168913 | ENHO | 0.49 | 3.14E-03 | 3.35E-02 |
| ENSG00000261879 | AC087500.1 | 0.49 | 3.14E-03 | 3.35E-02 |
| ENSG00000019169 | MARCO | -0.48 | 3.17E-03 | 3.39E-02 |
| ENSG00000152804 | HHEX | -0.48 | 3.17E-03 | 3.39E-02 |
| ENSG00000106803 | SEC61B | -0.48 | 3.18E-03 | 3.39E-02 |
| ENSG00000169902 | TPST1 | -0.48 | 3.18E-03 | 3.39E-02 |
| ENSG00000197291 | RAMP2-AS1 | 0.48 | 3.18E-03 | 3.39E-02 |
| ENSG00000226648 | PLCG1-AS1 | 0.48 | 3.19E-03 | 3.40E-02 |
| ENSG00000053918 | KCNQ1 | -0.48 | 3.19E-03 | 3.40E-02 |
| ENSG00000134375 | TIMM17A | -0.48 | 3.20E-03 | 3.40E-02 |
| ENSG00000143228 | NUF2 | -0.48 | 3.20E-03 | 3.40E-02 |
| ENSG00000163479 | SSR2 | -0.48 | 3.20E-03 | 3.40E-02 |
| ENSG00000245849 | RAD51-AS1 | 0.48 | 3.20E-03 | 3.40E-02 |
| ENSG00000169429 | CXCL8 | -0.48 | 3.20E-03 | 3.40E-02 |

| Ensembl ID | Gene Name | ρ | Pval | Padj |
|---|---|---|---|---|
| ENSG00000091527 | CDV3 | -0.48 | 3.22E-03 | 3.41E-02 |
| ENSG00000117143 | UAP1 | -0.48 | 3.22E-03 | 3.41E-02 |
| ENSG00000123353 | ORMDL2 | -0.48 | 3.22E-03 | 3.41E-02 |
| ENSG00000255284 | AP006621.3 | 0.48 | 3.22E-03 | 3.41E-02 |
| ENSG00000060558 | GNA15 | -0.48 | 3.22E-03 | 3.41E-02 |
| ENSG00000215277 | RNF212B | 0.48 | 3.24E-03 | 3.42E-02 |
| ENSG00000049323 | LTBP1 | -0.48 | 3.24E-03 | 3.42E-02 |
| ENSG00000111775 | COX6A1 | -0.48 | 3.24E-03 | 3.42E-02 |
| ENSG00000156398 | SFXN2 | 0.48 | 3.24E-03 | 3.42E-02 |
| ENSG00000165495 | PKNOX2 | 0.48 | 3.24E-03 | 3.42E-02 |
| ENSG00000258227 | CLEC5A | -0.48 | 3.24E-03 | 3.42E-02 |
| ENSG00000225486 | AL663058.1 | -0.48 | 3.24E-03 | 3.42E-02 |
| ENSG00000105887 | MTPN | -0.48 | 3.26E-03 | 3.43E-02 |
| ENSG00000174807 | CD248 | -0.48 | 3.26E-03 | 3.43E-02 |
| ENSG00000176105 | YES1 | -0.48 | 3.26E-03 | 3.43E-02 |
| ENSG00000254893 | AC113404.3 | -0.48 | 3.26E-03 | 3.43E-02 |
| ENSG00000243742 | RPLP0P2 | -0.48 | 3.28E-03 | 3.44E-02 |
| ENSG00000100099 | HPS4 | 0.48 | 3.28E-03 | 3.44E-02 |
| ENSG00000101844 | ATG4A | -0.48 | 3.28E-03 | 3.44E-02 |
| ENSG00000105221 | AKT2 | 0.48 | 3.28E-03 | 3.44E-02 |
| ENSG00000140368 | PSTPIP1 | 0.48 | 3.28E-03 | 3.44E-02 |
| ENSG00000163162 | RNF149 | -0.48 | 3.28E-03 | 3.44E-02 |
| ENSG00000168496 | FEN1 | -0.48 | 3.28E-03 | 3.44E-02 |
| ENSG00000278600 | AC015871.3 | 0.48 | 3.28E-03 | 3.44E-02 |
| ENSG00000162444 | RBP7 | -0.48 | 3.28E-03 | 3.44E-02 |
| ENSG00000277632 | CCL3 | -0.48 | 3.28E-03 | 3.44E-02 |
| ENSG00000186407 | CD300E | -0.48 | 3.29E-03 | 3.45E-02 |
| ENSG00000133134 | BEX2 | 0.48 | 3.30E-03 | 3.45E-02 |
| ENSG00000115944 | COX7A2L | -0.48 | 3.30E-03 | 3.45E-02 |
| ENSG00000138698 | RAP1GDS1 | -0.48 | 3.30E-03 | 3.45E-02 |
| ENSG00000140795 | MYLK3 | 0.48 | 3.30E-03 | 3.45E-02 |
| ENSG00000198492 | YTHDF2 | -0.48 | 3.30E-03 | 3.45E-02 |
| ENSG00000260892 | AC105020.4 | 0.48 | 3.30E-03 | 3.45E-02 |
| ENSG00000087502 | ERGIC2 | -0.48 | 3.32E-03 | 3.46E-02 |
| ENSG00000090889 | KIF4A | -0.48 | 3.32E-03 | 3.46E-02 |
| ENSG00000173546 | CSPG4 | 0.48 | 3.32E-03 | 3.46E-02 |
| ENSG00000213585 | VDAC1 | -0.48 | 3.32E-03 | 3.46E-02 |
| ENSG00000246022 | ALDH1L1-AS2 | 0.48 | 3.34E-03 | 3.47E-02 |
| ENSG00000105576 | TNPO2 | 0.48 | 3.34E-03 | 3.47E-02 |
| ENSG00000138018 | SELENOI | -0.48 | 3.34E-03 | 3.47E-02 |
| ENSG00000175348 | TMEM9B | -0.48 | 3.34E-03 | 3.47E-02 |
| ENSG00000175745 | NR2F1 | -0.48 | 3.36E-03 | 3.49E-02 |
| ENSG00000127511 | SIN3B | 0.48 | 3.36E-03 | 3.49E-02 |
| ENSG00000258667 | HIF1A-AS2 | 0.48 | 3.36E-03 | 3.49E-02 |
| ENSG00000203650 | LINC01285 | 0.48 | 3.37E-03 | 3.49E-02 |

| Ensembl ID | Gene Name | ρ | Pval | Padj |
|---|---|---|---|---|
| ENSG00000056736 | IL17RB | 0.48 | 3.38E-03 | 3.50E-02 |
| ENSG00000128524 | ATP6V1F | -0.48 | 3.38E-03 | 3.50E-02 |
| ENSG00000147383 | NSDHL | -0.48 | 3.38E-03 | 3.50E-02 |
| ENSG00000148396 | SEC16A | 0.48 | 3.38E-03 | 3.50E-02 |
| ENSG00000198873 | GRK5 | 0.48 | 3.38E-03 | 3.50E-02 |
| ENSG00000254910 | AC136475.2 | 0.48 | 3.38E-03 | 3.50E-02 |
| ENSG00000261327 | AC134312.5 | -0.48 | 3.39E-03 | 3.50E-02 |
| ENSG00000189057 | FAM111B | -0.48 | 3.39E-03 | 3.50E-02 |
| ENSG00000082196 | C1QTNF3 | -0.48 | 3.41E-03 | 3.51E-02 |
| ENSG00000146147 | MLIP | -0.48 | 3.41E-03 | 3.51E-02 |
| ENSG00000154928 | EPHB1 | 0.48 | 3.41E-03 | 3.51E-02 |
| ENSG00000065989 | AC011529.1 | 0.48 | 3.43E-03 | 3.53E-02 |
| ENSG00000157992 | KRTCAP3 | 0.48 | 3.43E-03 | 3.53E-02 |
| ENSG00000197712 | FAM114A1 | -0.48 | 3.43E-03 | 3.53E-02 |
| ENSG00000221716 | SNORA11 | 0.48 | 3.43E-03 | 3.53E-02 |
| ENSG00000272631 | AC067750.1 | 0.48 | 3.43E-03 | 3.53E-02 |
| ENSG00000266019 | MIR3609 | -0.48 | 3.44E-03 | 3.54E-02 |
| ENSG00000091490 | SEL1L3 | -0.48 | 3.45E-03 | 3.54E-02 |
| ENSG00000140459 | CYP11A1 | 0.48 | 3.45E-03 | 3.54E-02 |
| ENSG00000248905 | FMN1 | 0.48 | 3.45E-03 | 3.54E-02 |
| ENSG00000275131 | AC241952.1 | 0.48 | 3.45E-03 | 3.54E-02 |
| ENSG00000159189 | C1QC | -0.48 | 3.46E-03 | 3.55E-02 |
| ENSG00000085982 | USP40 | 0.48 | 3.47E-03 | 3.55E-02 |
| ENSG00000109063 | MYH3 | 0.48 | 3.47E-03 | 3.55E-02 |
| ENSG00000141753 | IGFBP4 | -0.48 | 3.47E-03 | 3.55E-02 |
| ENSG00000163932 | PRKCD | 0.48 | 3.47E-03 | 3.55E-02 |
| ENSG00000093009 | CDC45 | -0.48 | 3.48E-03 | 3.56E-02 |
| ENSG00000110852 | CLEC2B | -0.48 | 3.48E-03 | 3.56E-02 |
| ENSG00000129422 | MTUS1 | 0.48 | 3.49E-03 | 3.56E-02 |
| ENSG00000133624 | ZNF767P | 0.48 | 3.49E-03 | 3.56E-02 |
| ENSG00000172531 | PPP1CA | -0.48 | 3.49E-03 | 3.56E-02 |
| ENSG00000183684 | ALYREF | -0.48 | 3.49E-03 | 3.56E-02 |
| ENSG00000272430 | LINC02637 | 0.48 | 3.50E-03 | 3.57E-02 |
| ENSG00000272791 | AC073389.3 | 0.48 | 3.50E-03 | 3.57E-02 |
| ENSG00000106392 | C1GALT1 | -0.48 | 3.51E-03 | 3.58E-02 |
| ENSG00000108219 | TSPAN14 | -0.48 | 3.51E-03 | 3.58E-02 |
| ENSG00000166716 | ZNF592 | 0.48 | 3.51E-03 | 3.58E-02 |
| ENSG00000052802 | MSMO1 | -0.48 | 3.54E-03 | 3.59E-02 |
| ENSG00000065526 | SPEN | 0.48 | 3.54E-03 | 3.59E-02 |
| ENSG00000188283 | ZNF383 | 0.48 | 3.54E-03 | 3.59E-02 |
| ENSG00000276997 | AL513314.2 | 0.48 | 3.54E-03 | 3.59E-02 |
| ENSG00000135446 | CDK4 | -0.48 | 3.56E-03 | 3.61E-02 |
| ENSG00000136205 | TNS3 | 0.48 | 3.56E-03 | 3.61E-02 |
| ENSG00000261360 | AC010491.1 | 0.48 | 3.57E-03 | 3.62E-02 |
| ENSG00000005020 | SKAP2 | -0.48 | 3.58E-03 | 3.63E-02 |

| Ensembl ID | Gene Name | ρ | Pval | Padj |
|---|---|---|---|---|
| ENSG00000150756 | ATPSCKMT | -0.48 | 3.58E-03 | 3.63E-02 |
| ENSG00000123146 | ADGRE5 | -0.48 | 3.60E-03 | 3.64E-02 |
| ENSG00000131067 | GGT7 | 0.48 | 3.60E-03 | 3.64E-02 |
| ENSG00000143155 | TIPRL | -0.48 | 3.60E-03 | 3.64E-02 |
| ENSG00000148143 | ZNF462 | 0.48 | 3.60E-03 | 3.64E-02 |
| ENSG00000144868 | TMEM108 | -0.48 | 3.62E-03 | 3.66E-02 |
| ENSG00000153879 | CEBPG | -0.48 | 3.62E-03 | 3.66E-02 |
| ENSG00000231154 | MORF4L2-AS1 | 0.48 | 3.63E-03 | 3.67E-02 |
| ENSG00000029534 | ANK1 | -0.48 | 3.64E-03 | 3.67E-02 |
| ENSG00000255198 | SNHG9 | -0.48 | 3.64E-03 | 3.67E-02 |
| ENSG00000120306 | CYSTM1 | -0.48 | 3.65E-03 | 3.67E-02 |
| ENSG00000136888 | ATP6V1G1 | -0.48 | 3.65E-03 | 3.67E-02 |
| ENSG00000138495 | COX17 | -0.48 | 3.65E-03 | 3.67E-02 |
| ENSG00000228305 | AC016734.1 | -0.48 | 3.65E-03 | 3.67E-02 |
| ENSG00000134463 | ECHDC3 | 0.48 | 3.67E-03 | 3.69E-02 |
| ENSG00000138794 | CASP6 | -0.48 | 3.67E-03 | 3.69E-02 |
| ENSG00000151694 | ADAM17 | -0.48 | 3.67E-03 | 3.69E-02 |
| ENSG00000232993 | AL445183.1 | -0.48 | 3.67E-03 | 3.69E-02 |
| ENSG00000143257 | NR1I3 | -0.48 | 3.69E-03 | 3.70E-02 |
| ENSG00000132424 | PNISR | 0.48 | 3.69E-03 | 3.70E-02 |
| ENSG00000168002 | POLR2G | -0.48 | 3.69E-03 | 3.70E-02 |
| ENSG00000196689 | TRPV1 | 0.48 | 3.69E-03 | 3.70E-02 |
| ENSG00000197734 | C14orf178 | 0.48 | 3.71E-03 | 3.71E-02 |
| ENSG00000106066 | CPVL | -0.48 | 3.71E-03 | 3.72E-02 |
| ENSG00000125657 | TNFSF9 | -0.48 | 3.72E-03 | 3.72E-02 |
| ENSG00000142082 | SIRT3 | 0.48 | 3.72E-03 | 3.72E-02 |
| ENSG00000166825 | ANPEP | -0.48 | 3.72E-03 | 3.72E-02 |
| ENSG00000092853 | CLSPN | -0.48 | 3.72E-03 | 3.72E-02 |
| ENSG00000197927 | C2orf27A | -0.48 | 3.74E-03 | 3.73E-02 |
| ENSG00000114054 | PCCB | -0.48 | 3.74E-03 | 3.73E-02 |
| ENSG00000137941 | TTLL7 | 0.48 | 3.74E-03 | 3.73E-02 |
| ENSG00000153898 | MCOLN2 | 0.48 | 3.74E-03 | 3.73E-02 |
| ENSG00000249592 | AC139887.2 | 0.48 | 3.74E-03 | 3.73E-02 |
| ENSG00000234500 | AC008267.3 | 0.48 | 3.75E-03 | 3.74E-02 |
| ENSG00000156030 | ELMSAN1 | 0.48 | 3.76E-03 | 3.74E-02 |
| ENSG00000166016 | ABTB2 | 0.48 | 3.76E-03 | 3.74E-02 |
| ENSG00000281912 | LINC01144 | 0.48 | 3.76E-03 | 3.74E-02 |
| ENSG00000176907 | TCIM | -0.48 | 3.78E-03 | 3.76E-02 |
| ENSG00000088986 | DYNLL1 | -0.48 | 3.79E-03 | 3.76E-02 |
| ENSG00000107789 | MINPP1 | -0.48 | 3.79E-03 | 3.76E-02 |
| ENSG00000128340 | RAC2 | -0.48 | 3.79E-03 | 3.76E-02 |
| ENSG00000128591 | FLNC | -0.48 | 3.79E-03 | 3.76E-02 |
| ENSG00000153250 | RBMS1 | -0.48 | 3.81E-03 | 3.78E-02 |
| ENSG00000198604 | BAZ1A | -0.48 | 3.81E-03 | 3.78E-02 |
| ENSG00000274904 | AC093512.1 | 0.48 | 3.82E-03 | 3.79E-02 |

| Ensembl ID | Gene Name | ρ | Pval | Padj |
|---|---|---|---|---|
| ENSG00000203805 | PLPP4 | -0.48 | 3.83E-03 | 3.79E-02 |
| ENSG00000181392 | SYNE4 | 0.48 | 3.83E-03 | 3.79E-02 |
| ENSG00000069535 | MAOB | 0.48 | 3.83E-03 | 3.79E-02 |
| ENSG00000086504 | MRPL28 | -0.48 | 3.83E-03 | 3.79E-02 |
| ENSG00000196126 | HLA-DRB1 | -0.48 | 3.83E-03 | 3.79E-02 |
| ENSG00000197043 | ANXA6 | -0.48 | 3.83E-03 | 3.79E-02 |
| ENSG00000130707 | ASS1 | 0.48 | 3.86E-03 | 3.81E-02 |
| ENSG00000221817 | PPP3CB-AS1 | 0.48 | 3.86E-03 | 3.81E-02 |
| ENSG00000230091 | TMEM254-AS1 | 0.48 | 3.86E-03 | 3.81E-02 |
| ENSG00000279353 | AP001324.3 | 0.48 | 3.87E-03 | 3.81E-02 |
| ENSG00000060688 | SNRNP40 | -0.48 | 3.88E-03 | 3.81E-02 |
| ENSG00000075142 | SRI | -0.48 | 3.88E-03 | 3.81E-02 |
| ENSG00000128463 | EMC4 | -0.48 | 3.88E-03 | 3.81E-02 |
| ENSG00000128709 | HOXD9 | 0.48 | 3.88E-03 | 3.81E-02 |
| ENSG00000130035 | GALNT8 | 0.48 | 3.88E-03 | 3.81E-02 |
| ENSG00000141570 | CBX8 | -0.48 | 3.88E-03 | 3.81E-02 |
| ENSG00000162714 | ZNF496 | 0.48 | 3.88E-03 | 3.81E-02 |
| ENSG00000176024 | ZNF613 | -0.48 | 3.88E-03 | 3.81E-02 |
| ENSG00000231764 | DLX6-AS1 | 0.48 | 3.88E-03 | 3.81E-02 |
| ENSG00000054938 | CHRDL2 | 0.48 | 3.89E-03 | 3.82E-02 |
| ENSG00000038210 | PI4K2B | -0.48 | 3.90E-03 | 3.82E-02 |
| ENSG00000100997 | ABHD12 | -0.48 | 3.90E-03 | 3.82E-02 |
| ENSG00000131966 | ACTR10 | -0.48 | 3.90E-03 | 3.82E-02 |
| ENSG00000172586 | CHCHD1 | -0.48 | 3.90E-03 | 3.82E-02 |
| ENSG00000175224 | ATG13 | -0.48 | 3.90E-03 | 3.82E-02 |
| ENSG00000176170 | SPHK1 | -0.48 | 3.90E-03 | 3.82E-02 |
| ENSG00000188763 | FZD9 | 0.48 | 3.90E-03 | 3.82E-02 |
| ENSG00000197479 | PCDHB11 | 0.48 | 3.90E-03 | 3.82E-02 |
| ENSG00000236281 | NDUFB9P2 | -0.48 | 3.92E-03 | 3.83E-02 |
| ENSG00000115307 | AUP1 | -0.48 | 3.93E-03 | 3.83E-02 |
| ENSG00000117528 | ABCD3 | -0.48 | 3.93E-03 | 3.83E-02 |
| ENSG00000126698 | DNAJC8 | -0.48 | 3.93E-03 | 3.83E-02 |
| ENSG00000162997 | PRORSD1P | 0.48 | 3.93E-03 | 3.83E-02 |
| ENSG00000170271 | FAXDC2 | 0.48 | 3.93E-03 | 3.83E-02 |
| ENSG00000185499 | MUC1 | 0.48 | 3.93E-03 | 3.83E-02 |
| ENSG00000277075 | HIST1H2AE | -0.48 | 3.93E-03 | 3.83E-02 |
| ENSG00000198223 | CSF2RA | -0.47 | 3.95E-03 | 3.84E-02 |
| ENSG00000168785 | TSPAN5 | -0.47 | 3.95E-03 | 3.84E-02 |
| ENSG00000182578 | CSF1R | -0.47 | 3.95E-03 | 3.84E-02 |
| ENSG00000196187 | TMEM63A | 0.47 | 3.95E-03 | 3.84E-02 |
| ENSG00000233806 | LINC01237 | 0.47 | 3.96E-03 | 3.85E-02 |
| ENSG00000049167 | ERCC8 | -0.47 | 3.98E-03 | 3.85E-02 |
| ENSG00000116030 | SUMO1 | -0.47 | 3.98E-03 | 3.85E-02 |
| ENSG00000126261 | UBA2 | -0.47 | 3.98E-03 | 3.85E-02 |
| ENSG00000168386 | FILIP1L | -0.47 | 3.98E-03 | 3.85E-02 |

| Ensembl ID | Gene Name | ρ | Pval | Padj |
|---|---|---|---|---|
| ENSG00000198252 | STYX | -0.47 | 3.98E-03 | 3.85E-02 |
| ENSG00000238105 | GOLGA2P5 | 0.47 | 3.98E-03 | 3.85E-02 |
| ENSG00000237840 | FAM21FP | 0.47 | 3.99E-03 | 3.86E-02 |
| ENSG00000101004 | NINL | 0.47 | 4.00E-03 | 3.86E-02 |
| ENSG00000106636 | YKT6 | -0.47 | 4.00E-03 | 3.86E-02 |
| ENSG00000128482 | RNF112 | 0.47 | 4.00E-03 | 3.86E-02 |
| ENSG00000128789 | PSMG2 | -0.47 | 4.00E-03 | 3.86E-02 |
| ENSG00000164713 | BRI3 | -0.47 | 4.00E-03 | 3.86E-02 |
| ENSG00000168214 | RBPJ | -0.47 | 4.00E-03 | 3.86E-02 |
| ENSG00000169252 | ADRB2 | 0.47 | 4.00E-03 | 3.86E-02 |
| ENSG00000175662 | TOM1L2 | 0.47 | 4.00E-03 | 3.86E-02 |
| ENSG00000123219 | CENPK | -0.47 | 4.03E-03 | 3.88E-02 |
| ENSG00000165617 | DACT1 | 0.47 | 4.03E-03 | 3.88E-02 |
| ENSG00000114790 | ARHGEF26 | 0.47 | 4.05E-03 | 3.90E-02 |
| ENSG00000132313 | MRPL35 | -0.47 | 4.05E-03 | 3.90E-02 |
| ENSG00000101916 | TLR8 | -0.47 | 4.06E-03 | 3.90E-02 |
| ENSG00000227888 | FAM66A | 0.47 | 4.06E-03 | 3.90E-02 |
| ENSG00000250608 | AC010210.1 | 0.47 | 4.07E-03 | 3.90E-02 |
| ENSG00000260121 | AC138028.4 | 0.47 | 4.07E-03 | 3.90E-02 |
| ENSG00000075945 | KIFAP3 | -0.47 | 4.08E-03 | 3.90E-02 |
| ENSG00000080573 | COL5A3 | -0.47 | 4.08E-03 | 3.90E-02 |
| ENSG00000140839 | CLEC18B | 0.47 | 4.08E-03 | 3.90E-02 |
| ENSG00000156261 | CCT8 | -0.47 | 4.08E-03 | 3.90E-02 |
| ENSG00000157259 | GATAD1 | 0.47 | 4.08E-03 | 3.90E-02 |
| ENSG00000166086 | JAM3 | -0.47 | 4.08E-03 | 3.90E-02 |
| ENSG00000213694 | S1PR3 | -0.47 | 4.08E-03 | 3.90E-02 |
| ENSG00000230487 | PSMG3-AS1 | 0.47 | 4.08E-03 | 3.90E-02 |
| ENSG00000114013 | CD86 | -0.47 | 4.08E-03 | 3.90E-02 |
| ENSG00000075884 | ARHGAP15 | -0.47 | 4.08E-03 | 3.90E-02 |
| ENSG00000105048 | TNNT1 | -0.47 | 4.08E-03 | 3.90E-02 |
| ENSG00000263050 | AC090617.5 | -0.47 | 4.09E-03 | 3.91E-02 |
| ENSG00000198502 | HLA-DRB5 | -0.47 | 4.10E-03 | 3.91E-02 |
| ENSG00000121297 | TSHZ3 | -0.47 | 4.10E-03 | 3.91E-02 |
| ENSG00000136378 | ADAMTS7 | -0.47 | 4.10E-03 | 3.91E-02 |
| ENSG00000075415 | SLC25A3 | -0.47 | 4.13E-03 | 3.93E-02 |
| ENSG00000081791 | DELE1 | 0.47 | 4.13E-03 | 3.93E-02 |
| ENSG00000128641 | MYO1B | -0.47 | 4.13E-03 | 3.93E-02 |
| ENSG00000160161 | CILP2 | 0.47 | 4.13E-03 | 3.93E-02 |
| ENSG00000223547 | ZNF844 | 0.47 | 4.13E-03 | 3.93E-02 |
| ENSG00000265972 | TXNIP | 0.47 | 4.13E-03 | 3.93E-02 |
| ENSG00000161970 | RPL26 | -0.47 | 4.15E-03 | 3.95E-02 |
| ENSG00000251143 | AP002490.1 | 0.47 | 4.15E-03 | 3.95E-02 |
| ENSG00000188033 | ZNF490 | 0.47 | 4.16E-03 | 3.95E-02 |
| ENSG00000260091 | AC093752.3 | 0.47 | 4.17E-03 | 3.96E-02 |
| ENSG00000142156 | COL6A1 | -0.47 | 4.18E-03 | 3.96E-02 |

| Ensembl ID | Gene Name | ρ | Pval | Padj |
|---|---|---|---|---|
| ENSG00000198794 | SCAMP5 | 0.47 | 4.18E-03 | 3.96E-02 |
| ENSG00000100167 | SEPT3 | -0.47 | 4.19E-03 | 3.97E-02 |
| ENSG00000123975 | CKS2 | -0.47 | 4.20E-03 | 3.97E-02 |
| ENSG00000178809 | TRIM73 | 0.47 | 4.20E-03 | 3.97E-02 |
| ENSG00000092445 | TYRO3 | -0.47 | 4.20E-03 | 3.97E-02 |
| ENSG00000136732 | GYPC | -0.47 | 4.20E-03 | 3.97E-02 |
| ENSG00000183808 | RBM12B | 0.47 | 4.20E-03 | 3.97E-02 |
| ENSG00000196850 | PPTC7 | -0.47 | 4.20E-03 | 3.97E-02 |
| ENSG00000247934 | AC022364.1 | 0.47 | 4.20E-03 | 3.97E-02 |
| ENSG00000257176 | AC009318.1 | 0.47 | 4.20E-03 | 3.97E-02 |
| ENSG00000137947 | GTF2B | -0.47 | 4.23E-03 | 3.99E-02 |
| ENSG00000153976 | HS3ST3A1 | 0.47 | 4.23E-03 | 3.99E-02 |
| ENSG00000271725 | AC103858.1 | 0.47 | 4.24E-03 | 4.00E-02 |
| ENSG00000085491 | SLC25A24 | -0.47 | 4.25E-03 | 4.00E-02 |
| ENSG00000157020 | SEC13 | -0.47 | 4.25E-03 | 4.00E-02 |
| ENSG00000173757 | STAT5B | 0.47 | 4.25E-03 | 4.00E-02 |
| ENSG00000242732 | RTL5 | 0.47 | 4.25E-03 | 4.00E-02 |
| ENSG00000260233 | SSSCA1-AS1 | 0.47 | 4.25E-03 | 4.00E-02 |
| ENSG00000280434 | AL031595.3 | 0.47 | 4.27E-03 | 4.01E-02 |
| ENSG00000097021 | ACOT7 | -0.47 | 4.28E-03 | 4.01E-02 |
| ENSG00000119314 | PTBP3 | -0.47 | 4.28E-03 | 4.01E-02 |
| ENSG00000131475 | VPS25 | -0.47 | 4.28E-03 | 4.01E-02 |
| ENSG00000148834 | GSTO1 | -0.47 | 4.28E-03 | 4.01E-02 |
| ENSG00000163159 | VPS72 | -0.47 | 4.28E-03 | 4.01E-02 |
| ENSG00000169020 | ATP5ME | -0.47 | 4.28E-03 | 4.01E-02 |
| ENSG00000082781 | ITGB5 | -0.47 | 4.31E-03 | 4.03E-02 |
| ENSG00000093144 | ECHDC1 | -0.47 | 4.31E-03 | 4.03E-02 |
| ENSG00000101353 | MROH8 | 0.47 | 4.31E-03 | 4.03E-02 |
| ENSG00000180616 | SSTR2 | 0.47 | 4.31E-03 | 4.03E-02 |
| ENSG00000185963 | BICD2 | -0.47 | 4.31E-03 | 4.03E-02 |
| ENSG00000260404 | AC110079.1 | 0.47 | 4.31E-03 | 4.03E-02 |
| ENSG00000273038 | AL365203.2 | -0.47 | 4.31E-03 | 4.03E-02 |
| ENSG00000255234 | AP000446.1 | 0.47 | 4.33E-03 | 4.04E-02 |
| ENSG00000145990 | GFOD1 | 0.47 | 4.33E-03 | 4.04E-02 |
| ENSG00000196498 | NCOR2 | 0.47 | 4.33E-03 | 4.04E-02 |
| ENSG00000267152 | AC093227.1 | 0.47 | 4.33E-03 | 4.04E-02 |
| ENSG00000010810 | FYN | -0.47 | 4.36E-03 | 4.06E-02 |
| ENSG00000137216 | TMEM63B | -0.47 | 4.36E-03 | 4.06E-02 |
| ENSG00000183648 | NDUFB1 | -0.47 | 4.36E-03 | 4.06E-02 |
| ENSG00000111737 | RAB35 | -0.47 | 4.38E-03 | 4.07E-02 |
| ENSG00000115310 | RTN4 | -0.47 | 4.38E-03 | 4.07E-02 |
| ENSG00000124299 | PEPD | -0.47 | 4.38E-03 | 4.07E-02 |
| ENSG00000145075 | CCDC39 | 0.47 | 4.38E-03 | 4.07E-02 |
| ENSG00000170270 | GON7 | -0.47 | 4.38E-03 | 4.07E-02 |
| ENSG00000183160 | TMEM119 | -0.47 | 4.38E-03 | 4.07E-02 |

| Ensembl ID | Gene Name | ρ | Pval | Padj |
|---|---|---|---|---|
| ENSG00000271918 | AC034236.2 | 0.47 | 4.38E-03 | 4.07E-02 |
| ENSG00000254206 | NPIPB11 | 0.47 | 4.40E-03 | 4.08E-02 |
| ENSG00000109118 | PHF12 | 0.47 | 4.41E-03 | 4.09E-02 |
| ENSG00000116883 | AL591845.1 | 0.47 | 4.41E-03 | 4.09E-02 |
| ENSG00000135723 | FHOD1 | -0.47 | 4.41E-03 | 4.09E-02 |
| ENSG00000272180 | AC011306.1 | 0.47 | 4.42E-03 | 4.10E-02 |
| ENSG00000012779 | ALOX5 | -0.47 | 4.43E-03 | 4.10E-02 |
| ENSG00000105426 | PTPRS | 0.47 | 4.44E-03 | 4.10E-02 |
| ENSG00000259577 | CERNA1 | 0.47 | 4.44E-03 | 4.10E-02 |
| ENSG00000263956 | NBPF11 | 0.47 | 4.44E-03 | 4.10E-02 |
| ENSG00000162779 | AXDND1 | 0.47 | 4.44E-03 | 4.10E-02 |
| ENSG00000164266 | SPINK1 | -0.47 | 4.45E-03 | 4.12E-02 |
| ENSG00000108039 | XPNPEP1 | -0.47 | 4.47E-03 | 4.12E-02 |
| ENSG00000142686 | C1orf216 | -0.47 | 4.47E-03 | 4.12E-02 |
| ENSG00000111445 | RFC5 | -0.47 | 4.49E-03 | 4.14E-02 |
| ENSG00000126870 | WDR60 | 0.47 | 4.49E-03 | 4.14E-02 |
| ENSG00000132326 | PER2 | 0.47 | 4.49E-03 | 4.14E-02 |
| ENSG00000166348 | USP54 | 0.47 | 4.49E-03 | 4.14E-02 |
| ENSG00000184381 | PLA2G6 | 0.47 | 4.49E-03 | 4.14E-02 |
| ENSG00000239857 | GET4 | 0.47 | 4.50E-03 | 4.15E-02 |
| ENSG00000095752 | IL11 | -0.47 | 4.52E-03 | 4.15E-02 |
| ENSG00000115226 | FNDC4 | -0.47 | 4.52E-03 | 4.15E-02 |
| ENSG00000128928 | IVD | 0.47 | 4.52E-03 | 4.15E-02 |
| ENSG00000143458 | GABPB2 | 0.47 | 4.52E-03 | 4.15E-02 |
| ENSG00000035403 | VCL | -0.47 | 4.55E-03 | 4.16E-02 |
| ENSG00000100206 | DMC1 | 0.47 | 4.55E-03 | 4.16E-02 |
| ENSG00000111481 | COPZ1 | -0.47 | 4.55E-03 | 4.16E-02 |
| ENSG00000164305 | CASP3 | -0.47 | 4.55E-03 | 4.16E-02 |
| ENSG00000180228 | PRKRA | -0.47 | 4.55E-03 | 4.16E-02 |
| ENSG00000204650 | LINC02210 | 0.47 | 4.55E-03 | 4.16E-02 |
| ENSG00000235703 | LINC00894 | 0.47 | 4.55E-03 | 4.16E-02 |
| ENSG00000150048 | CLEC1A | -0.47 | 4.55E-03 | 4.16E-02 |
| ENSG00000233977 | AC099681.3 | 0.47 | 4.57E-03 | 4.18E-02 |
| ENSG00000082212 | ME2 | -0.47 | 4.57E-03 | 4.18E-02 |
| ENSG00000106305 | AIMP2 | -0.47 | 4.57E-03 | 4.18E-02 |
| ENSG00000123080 | CDKN2C | -0.47 | 4.57E-03 | 4.18E-02 |
| ENSG00000138430 | OLA1 | -0.47 | 4.57E-03 | 4.18E-02 |
| ENSG00000169604 | ANTXR1 | -0.47 | 4.57E-03 | 4.18E-02 |
| ENSG00000123384 | LRP1 | 0.47 | 4.60E-03 | 4.20E-02 |
| ENSG00000145284 | SCD5 | -0.47 | 4.60E-03 | 4.20E-02 |
| ENSG00000197045 | GMFB | -0.47 | 4.60E-03 | 4.20E-02 |
| ENSG00000259884 | AC025259.3 | -0.47 | 4.61E-03 | 4.20E-02 |
| ENSG00000035115 | SH3YL1 | 0.47 | 4.63E-03 | 4.20E-02 |
| ENSG00000079246 | XRCC5 | -0.47 | 4.63E-03 | 4.20E-02 |
| ENSG00000088826 | SMOX | -0.47 | 4.63E-03 | 4.20E-02 |

| Ensembl ID | Gene Name | ρ | Pval | Padj |
|---|---|---|---|---|
| ENSG00000126775 | ATG14 | 0.47 | 4.63E-03 | 4.20E-02 |
| ENSG00000144134 | RABL2A | 0.47 | 4.63E-03 | 4.20E-02 |
| ENSG00000174989 | FBXW8 | 0.47 | 4.63E-03 | 4.20E-02 |
| ENSG00000178896 | EXOSC4 | -0.47 | 4.63E-03 | 4.20E-02 |
| ENSG00000205060 | SLC35B4 | -0.47 | 4.63E-03 | 4.20E-02 |
| ENSG00000268858 | AL118506.1 | 0.47 | 4.63E-03 | 4.20E-02 |
| ENSG00000259347 | AC087482.1 | 0.47 | 4.65E-03 | 4.22E-02 |
| ENSG00000095383 | TBC1D2 | -0.47 | 4.66E-03 | 4.22E-02 |
| ENSG00000158869 | FCER1G | -0.47 | 4.66E-03 | 4.22E-02 |
| ENSG00000184924 | PTRHD1 | -0.47 | 4.66E-03 | 4.22E-02 |
| ENSG00000271576 | AL359504.2 | 0.47 | 4.68E-03 | 4.23E-02 |
| ENSG00000110717 | NDUFS8 | -0.47 | 4.69E-03 | 4.23E-02 |
| ENSG00000141644 | MBD1 | 0.47 | 4.69E-03 | 4.23E-02 |
| ENSG00000163975 | MELTF | 0.47 | 4.69E-03 | 4.23E-02 |
| ENSG00000166797 | CIAO2A | -0.47 | 4.69E-03 | 4.23E-02 |
| ENSG00000275400 | AC006001.4 | -0.47 | 4.69E-03 | 4.23E-02 |
| ENSG00000260139 | CSPG4P13 | 0.47 | 4.71E-03 | 4.23E-02 |
| ENSG00000271980 | AC012640.4 | -0.47 | 4.71E-03 | 4.23E-02 |
| ENSG00000002586 | CD99 | -0.47 | 4.71E-03 | 4.23E-02 |
| ENSG00000071994 | PDCD2 | -0.47 | 4.71E-03 | 4.23E-02 |
| ENSG00000131467 | PSME3 | -0.47 | 4.71E-03 | 4.23E-02 |
| ENSG00000132964 | CDK8 | -0.47 | 4.71E-03 | 4.23E-02 |
| ENSG00000144810 | COL8A1 | -0.47 | 4.71E-03 | 4.23E-02 |
| ENSG00000152894 | PTPRK | -0.47 | 4.71E-03 | 4.23E-02 |
| ENSG00000156017 | CARNMT1 | -0.47 | 4.71E-03 | 4.23E-02 |
| ENSG00000160714 | UBE2Q1 | -0.47 | 4.71E-03 | 4.23E-02 |
| ENSG00000161149 | TUBA3FP | 0.47 | 4.71E-03 | 4.23E-02 |
| ENSG00000166105 | GLB1L3 | 0.47 | 4.71E-03 | 4.23E-02 |
| ENSG00000169062 | UPF3A | 0.47 | 4.71E-03 | 4.23E-02 |
| ENSG00000174442 | ZWILCH | -0.47 | 4.71E-03 | 4.23E-02 |
| ENSG00000214021 | TTLL3 | 0.47 | 4.71E-03 | 4.23E-02 |
| ENSG00000233622 | CYP2T1P | 0.47 | 4.71E-03 | 4.23E-02 |
| ENSG00000234911 | TEX21P | 0.47 | 4.72E-03 | 4.23E-02 |
| ENSG00000129946 | SHC2 | -0.47 | 4.74E-03 | 4.24E-02 |
| ENSG00000134440 | NARS | -0.47 | 4.74E-03 | 4.24E-02 |
| ENSG00000135406 | PRPH | 0.47 | 4.74E-03 | 4.24E-02 |
| ENSG00000138071 | ACTR2 | -0.47 | 4.74E-03 | 4.24E-02 |
| ENSG00000196878 | LAMB3 | -0.47 | 4.74E-03 | 4.24E-02 |
| ENSG00000122966 | CIT | -0.47 | 4.77E-03 | 4.26E-02 |
| ENSG00000124562 | SNRPC | -0.47 | 4.77E-03 | 4.26E-02 |
| ENSG00000143106 | PSMA5 | -0.47 | 4.77E-03 | 4.26E-02 |
| ENSG00000148688 | RPP30 | -0.47 | 4.77E-03 | 4.26E-02 |
| ENSG00000177839 | PCDHB9 | 0.47 | 4.77E-03 | 4.26E-02 |
| ENSG00000203760 | CENPW | -0.47 | 4.77E-03 | 4.26E-02 |
| ENSG00000007171 | NOS2 | 0.47 | 4.80E-03 | 4.27E-02 |

| Ensembl ID | Gene Name | ρ | Pval | Padj |
|---|---|---|---|---|
| ENSG00000054219 | LY75 | 0.47 | 4.80E-03 | 4.27E-02 |
| ENSG00000120149 | MSX2 | -0.47 | 4.80E-03 | 4.27E-02 |
| ENSG00000133216 | EPHB2 | -0.47 | 4.80E-03 | 4.27E-02 |
| ENSG00000162521 | RBBP4 | -0.47 | 4.80E-03 | 4.27E-02 |
| ENSG00000165168 | CYBB | -0.47 | 4.80E-03 | 4.27E-02 |
| ENSG00000001036 | FUCA2 | -0.47 | 4.83E-03 | 4.29E-02 |
| ENSG00000092964 | DPYSL2 | -0.47 | 4.83E-03 | 4.29E-02 |
| ENSG00000100030 | MAPK1 | -0.47 | 4.83E-03 | 4.29E-02 |
| ENSG00000137171 | KLC4 | 0.47 | 4.83E-03 | 4.29E-02 |
| ENSG00000066583 | ISOC1 | -0.47 | 4.86E-03 | 4.31E-02 |
| ENSG00000156171 | DRAM2 | -0.47 | 4.86E-03 | 4.31E-02 |
| ENSG00000164828 | SUN1 | 0.47 | 4.86E-03 | 4.31E-02 |
| ENSG00000184232 | OAF | 0.47 | 4.86E-03 | 4.31E-02 |
| ENSG00000212694 | LINC01089 | 0.47 | 4.86E-03 | 4.31E-02 |
| ENSG00000120280 | CXorf21 | -0.47 | 4.87E-03 | 4.31E-02 |
| ENSG00000169403 | PTAFR | -0.47 | 4.87E-03 | 4.31E-02 |
| ENSG00000279246 | AP003108.3 | 0.47 | 4.87E-03 | 4.31E-02 |
| ENSG00000099326 | MZF1 | 0.46 | 4.89E-03 | 4.32E-02 |
| ENSG00000132432 | SEC61G | -0.46 | 4.89E-03 | 4.32E-02 |
| ENSG00000214655 | ZSWIM8 | 0.46 | 4.89E-03 | 4.32E-02 |
| ENSG00000083814 | ZNF671 | -0.46 | 4.91E-03 | 4.34E-02 |
| ENSG00000005075 | POLR2J | -0.46 | 4.92E-03 | 4.34E-02 |
| ENSG00000164976 | MYORG | 0.46 | 4.92E-03 | 4.34E-02 |
| ENSG00000173915 | ATP5MD | -0.46 | 4.92E-03 | 4.34E-02 |
| ENSG00000163072 | NOSTRIN | -0.46 | 4.93E-03 | 4.35E-02 |
| ENSG00000178343 | SHISA3 | 0.46 | 4.93E-03 | 4.35E-02 |
| ENSG00000163376 | KBTBD8 | -0.46 | 4.94E-03 | 4.35E-02 |
| ENSG00000124541 | RRP36 | -0.46 | 4.95E-03 | 4.35E-02 |
| ENSG00000128534 | LSM8 | -0.46 | 4.95E-03 | 4.35E-02 |
| ENSG00000129515 | SNX6 | -0.46 | 4.95E-03 | 4.35E-02 |
| ENSG00000134030 | CTIF | 0.46 | 4.95E-03 | 4.35E-02 |
| ENSG00000140400 | MAN2C1 | 0.46 | 4.95E-03 | 4.35E-02 |
| ENSG00000167863 | ATP5PD | -0.46 | 4.95E-03 | 4.35E-02 |
| ENSG00000234380 | LINC01426 | -0.46 | 4.95E-03 | 4.35E-02 |
| ENSG00000134049 | IER3IP1 | -0.46 | 4.97E-03 | 4.36E-02 |
| ENSG00000142611 | PRDM16 | 0.46 | 4.97E-03 | 4.36E-02 |
| ENSG00000172794 | RAB37 | 0.46 | 4.97E-03 | 4.36E-02 |
| ENSG00000182544 | MFSD5 | -0.46 | 4.97E-03 | 4.36E-02 |
| ENSG00000283031 | AC009242.1 | 0.46 | 4.98E-03 | 4.37E-02 |
| ENSG00000104783 | KCNN4 | -0.46 | 5.00E-03 | 4.38E-02 |
| ENSG00000114737 | CISH | 0.46 | 5.00E-03 | 4.38E-02 |
| ENSG00000180881 | CAPS2 | 0.46 | 5.00E-03 | 4.38E-02 |
| ENSG00000177508 | IRX3 | -0.46 | 5.01E-03 | 4.38E-02 |
| ENSG00000235545 | AC103923.1 | 0.46 | 5.02E-03 | 4.39E-02 |
| ENSG00000085511 | MAP3K4 | 0.46 | 5.03E-03 | 4.40E-02 |

| Ensembl ID | Gene Name | ρ | Pval | Padj |
|---|---|---|---|---|
| ENSG00000101255 | TRIB3 | -0.46 | 5.03E-03 | 4.40E-02 |
| ENSG00000108590 | MED31 | -0.46 | 5.03E-03 | 4.40E-02 |
| ENSG00000152904 | GGPS1 | -0.46 | 5.03E-03 | 4.40E-02 |
| ENSG00000270015 | AC087481.3 | 0.46 | 5.03E-03 | 4.40E-02 |
| ENSG00000275374 | AC113385.3 | 0.46 | 5.06E-03 | 4.41E-02 |
| ENSG00000122565 | CBX3 | -0.46 | 5.06E-03 | 4.41E-02 |
| ENSG00000136930 | PSMB7 | -0.46 | 5.06E-03 | 4.41E-02 |
| ENSG00000198155 | ZNF876P | 0.46 | 5.06E-03 | 4.41E-02 |
| ENSG00000145703 | IQGAP2 | -0.46 | 5.08E-03 | 4.43E-02 |
| ENSG00000260796 | AC145285.3 | 0.46 | 5.09E-03 | 4.43E-02 |
| ENSG00000086666 | ZFAND6 | -0.46 | 5.09E-03 | 4.43E-02 |
| ENSG00000224078 | SNHG14 | 0.46 | 5.09E-03 | 4.43E-02 |
| ENSG00000260747 | AC022968.1 | -0.46 | 5.09E-03 | 4.43E-02 |
| ENSG00000187474 | FPR3 | -0.46 | 5.11E-03 | 4.44E-02 |
| ENSG00000125970 | RALY | -0.46 | 5.12E-03 | 4.45E-02 |
| ENSG00000165264 | NDUFB6 | -0.46 | 5.12E-03 | 4.45E-02 |
| ENSG00000215374 | FAM66B | 0.46 | 5.12E-03 | 4.45E-02 |
| ENSG00000261061 | AC092718.4 | -0.46 | 5.12E-03 | 4.45E-02 |
| ENSG00000049089 | COL9A2 | 0.46 | 5.16E-03 | 4.46E-02 |
| ENSG00000109536 | FRG1 | -0.46 | 5.16E-03 | 4.46E-02 |
| ENSG00000149100 | EIF3M | -0.46 | 5.16E-03 | 4.46E-02 |
| ENSG00000162695 | SLC30A7 | -0.46 | 5.16E-03 | 4.46E-02 |
| ENSG00000165609 | NUDT5 | -0.46 | 5.16E-03 | 4.46E-02 |
| ENSG00000188428 | BLOC1S5 | 0.46 | 5.16E-03 | 4.46E-02 |
| ENSG00000282826 | FRG1CP | -0.46 | 5.16E-03 | 4.46E-02 |
| ENSG00000267397 | AC090229.1 | -0.46 | 5.18E-03 | 4.48E-02 |
| ENSG00000267193 | AC023421.2 | 0.46 | 5.18E-03 | 4.48E-02 |
| ENSG00000078369 | GNB1 | -0.46 | 5.19E-03 | 4.48E-02 |
| ENSG00000100014 | SPECC1L | -0.46 | 5.19E-03 | 4.48E-02 |
| ENSG00000117280 | RAB29 | -0.46 | 5.19E-03 | 4.48E-02 |
| ENSG00000261123 | AC009065.5 | 0.46 | 5.19E-03 | 4.48E-02 |
| ENSG00000177990 | DPY19L2 | 0.46 | 5.21E-03 | 4.49E-02 |
| ENSG00000116729 | WLS | -0.46 | 5.22E-03 | 4.49E-02 |
| ENSG00000127314 | RAP1B | -0.46 | 5.22E-03 | 4.49E-02 |
| ENSG00000132603 | NIP7 | -0.46 | 5.22E-03 | 4.49E-02 |
| ENSG00000138756 | BMP2K | -0.46 | 5.22E-03 | 4.49E-02 |
| ENSG00000151611 | MMAA | -0.46 | 5.22E-03 | 4.49E-02 |
| ENSG00000223768 | LINC00205 | 0.46 | 5.22E-03 | 4.49E-02 |
| ENSG00000251364 | AC107884.1 | 0.46 | 5.22E-03 | 4.49E-02 |
| ENSG00000153558 | FBXL2 | 0.46 | 5.25E-03 | 4.50E-02 |
| ENSG00000198026 | ZNF335 | 0.46 | 5.25E-03 | 4.50E-02 |
| ENSG00000219023 | AL033519.2 | -0.46 | 5.25E-03 | 4.50E-02 |
| ENSG00000230043 | TMSB4XP6 | -0.46 | 5.25E-03 | 4.50E-02 |
| ENSG00000240857 | RDH14 | -0.46 | 5.25E-03 | 4.50E-02 |
| ENSG00000227482 | AL157702.2 | -0.46 | 5.26E-03 | 4.52E-02 |

| Ensembl ID | Gene Name | ρ | Pval | Padj |
|---|---|---|---|---|
| ENSG00000124787 | RPP40 | -0.46 | 5.28E-03 | 4.52E-02 |
| ENSG00000104691 | UBXN8 | -0.46 | 5.28E-03 | 4.52E-02 |
| ENSG00000124802 | EEF1E1 | -0.46 | 5.28E-03 | 4.52E-02 |
| ENSG00000148700 | ADD3 | -0.46 | 5.28E-03 | 4.52E-02 |
| ENSG00000158985 | CDC42SE2 | -0.46 | 5.28E-03 | 4.52E-02 |
| ENSG00000225868 | AC016582.1 | 0.46 | 5.28E-03 | 4.52E-02 |
| ENSG00000065621 | GSTO2 | 0.46 | 5.31E-03 | 4.53E-02 |
| ENSG00000126351 | THRA | 0.46 | 5.31E-03 | 4.53E-02 |
| ENSG00000137168 | PPIL1 | -0.46 | 5.31E-03 | 4.53E-02 |
| ENSG00000144445 | KANSL1L | 0.46 | 5.31E-03 | 4.53E-02 |
| ENSG00000160219 | GAB3 | -0.46 | 5.31E-03 | 4.53E-02 |
| ENSG00000175155 | YPEL2 | 0.46 | 5.31E-03 | 4.53E-02 |
| ENSG00000185052 | SLC24A3 | -0.46 | 5.31E-03 | 4.53E-02 |
| ENSG00000128512 | DOCK4 | -0.46 | 5.32E-03 | 4.54E-02 |
| ENSG00000051128 | HOMER3 | -0.46 | 5.34E-03 | 4.55E-02 |
| ENSG00000100632 | ERH | -0.46 | 5.34E-03 | 4.55E-02 |
| ENSG00000112659 | CUL9 | 0.46 | 5.34E-03 | 4.55E-02 |
| ENSG00000170629 | DPY19L2P2 | 0.46 | 5.34E-03 | 4.55E-02 |
| ENSG00000272106 | AL691432.2 | 0.46 | 5.34E-03 | 4.55E-02 |
| ENSG00000272509 | AC087752.4 | -0.46 | 5.37E-03 | 4.56E-02 |
| ENSG00000022976 | ZNF839 | 0.46 | 5.37E-03 | 4.56E-02 |
| ENSG00000064205 | CCN5 | -0.46 | 5.37E-03 | 4.56E-02 |
| ENSG00000106868 | SUSD1 | -0.46 | 5.37E-03 | 4.56E-02 |
| ENSG00000140990 | NDUFB10 | -0.46 | 5.37E-03 | 4.56E-02 |
| ENSG00000214013 | GANC | 0.46 | 5.37E-03 | 4.56E-02 |
| ENSG00000110025 | SNX15 | 0.46 | 5.38E-03 | 4.57E-02 |
| ENSG00000157542 | KCNJ6 | -0.46 | 5.39E-03 | 4.57E-02 |
| ENSG00000064199 | SPA17 | -0.46 | 5.41E-03 | 4.58E-02 |
| ENSG00000105254 | TBCB | -0.46 | 5.41E-03 | 4.58E-02 |
| ENSG00000155438 | NIFK | -0.46 | 5.41E-03 | 4.58E-02 |
| ENSG00000061794 | MRPS35 | -0.46 | 5.44E-03 | 4.60E-02 |
| ENSG00000103888 | CEMIP | -0.46 | 5.44E-03 | 4.60E-02 |
| ENSG00000174227 | PIGG | 0.46 | 5.44E-03 | 4.60E-02 |
| ENSG00000235162 | C12orf75 | -0.46 | 5.44E-03 | 4.60E-02 |
| ENSG00000247596 | TWF2 | -0.46 | 5.44E-03 | 4.60E-02 |
| ENSG00000230614 | DYNLL1P7 | -0.46 | 5.45E-03 | 4.60E-02 |
| ENSG00000235505 | CASP17P | -0.46 | 5.46E-03 | 4.61E-02 |
| ENSG00000023697 | DERA | -0.46 | 5.47E-03 | 4.61E-02 |
| ENSG00000078237 | TIGAR | -0.46 | 5.47E-03 | 4.61E-02 |
| ENSG00000136816 | TOR1B | -0.46 | 5.47E-03 | 4.61E-02 |
| ENSG00000151465 | CDC123 | -0.46 | 5.47E-03 | 4.61E-02 |
| ENSG00000105538 | RASIP1 | -0.46 | 5.50E-03 | 4.62E-02 |
| ENSG00000108559 | NUP88 | -0.46 | 5.50E-03 | 4.62E-02 |
| ENSG00000139826 | ABHD13 | -0.46 | 5.50E-03 | 4.62E-02 |
| ENSG00000148481 | MINDY3 | -0.46 | 5.50E-03 | 4.62E-02 |

| Ensembl ID | Gene Name | ρ | Pval | Padj |
|---|---|---|---|---|
| ENSG00000206053 | JPT2 | -0.46 | 5.50E-03 | 4.62E-02 |
| ENSG00000227372 | TP73-AS1 | 0.46 | 5.50E-03 | 4.62E-02 |
| ENSG00000256894 | AC022509.3 | 0.46 | 5.50E-03 | 4.62E-02 |
| ENSG00000124228 | DDX27 | 0.46 | 5.53E-03 | 4.65E-02 |
| ENSG00000269069 | AC007842.1 | 0.46 | 5.53E-03 | 4.65E-02 |
| ENSG00000156709 | AIFM1 | -0.46 | 5.57E-03 | 4.67E-02 |
| ENSG00000166405 | RIC3 | 0.46 | 5.57E-03 | 4.67E-02 |
| ENSG00000197872 | FAM49A | -0.46 | 5.57E-03 | 4.67E-02 |
| ENSG00000121022 | COPS5 | -0.46 | 5.60E-03 | 4.69E-02 |
| ENSG00000147144 | CCDC120 | 0.46 | 5.60E-03 | 4.69E-02 |
| ENSG00000249087 | ZNF436-AS1 | 0.46 | 5.60E-03 | 4.69E-02 |
| ENSG00000125868 | DSTN | -0.46 | 5.63E-03 | 4.72E-02 |
| ENSG00000163631 | ALB | 0.46 | 5.66E-03 | 4.73E-02 |
| ENSG00000249992 | TMEM158 | -0.46 | 5.66E-03 | 4.73E-02 |
| ENSG00000130962 | PRRG1 | -0.46 | 5.67E-03 | 4.73E-02 |
| ENSG00000166548 | TK2 | 0.46 | 5.67E-03 | 4.73E-02 |
| ENSG00000174231 | PRPF8 | 0.46 | 5.67E-03 | 4.73E-02 |
| ENSG00000213186 | TRIM59 | -0.46 | 5.67E-03 | 4.73E-02 |
| ENSG00000245149 | RNF139-AS1 | 0.46 | 5.67E-03 | 4.73E-02 |
| ENSG00000107796 | ACTA2 | -0.46 | 5.70E-03 | 4.75E-02 |
| ENSG00000137275 | RIPK1 | -0.46 | 5.70E-03 | 4.75E-02 |
| ENSG00000102032 | RENBP | -0.46 | 5.72E-03 | 4.77E-02 |
| ENSG00000269926 | DDIT4-AS1 | 0.46 | 5.72E-03 | 4.77E-02 |
| ENSG00000014641 | MDH1 | -0.46 | 5.77E-03 | 4.80E-02 |
| ENSG00000112584 | FAM120B | 0.46 | 5.77E-03 | 4.80E-02 |
| ENSG00000171943 | SRGAP2C | -0.46 | 5.77E-03 | 4.80E-02 |
| ENSG00000173818 | ENDOV | 0.46 | 5.77E-03 | 4.80E-02 |
| ENSG00000188613 | NANOS1 | -0.46 | 5.77E-03 | 4.80E-02 |
| ENSG00000279861 | AC073548.1 | 0.46 | 5.77E-03 | 4.80E-02 |
| ENSG00000272630 | AL731563.3 | 0.46 | 5.78E-03 | 4.80E-02 |
| ENSG00000140379 | BCL2A1 | -0.46 | 5.80E-03 | 4.81E-02 |
| ENSG00000001497 | LAS1L | 0.46 | 5.80E-03 | 4.81E-02 |
| ENSG00000049449 | RCN1 | -0.46 | 5.80E-03 | 4.81E-02 |
| ENSG00000070367 | EXOC5 | -0.46 | 5.80E-03 | 4.81E-02 |
| ENSG00000113368 | LMNB1 | -0.46 | 5.80E-03 | 4.81E-02 |
| ENSG00000166441 | RPL27A | -0.46 | 5.80E-03 | 4.81E-02 |
| ENSG00000197696 | NMB | -0.46 | 5.80E-03 | 4.81E-02 |
| ENSG00000160785 | SLC25A44 | -0.46 | 5.83E-03 | 4.83E-02 |
| ENSG00000174574 | AKIRIN1 | -0.46 | 5.83E-03 | 4.83E-02 |
| ENSG00000160883 | HK3 | -0.46 | 5.86E-03 | 4.85E-02 |
| ENSG00000135365 | PHF21A | 0.46 | 5.87E-03 | 4.85E-02 |
| ENSG00000198858 | R3HDM4 | -0.46 | 5.87E-03 | 4.85E-02 |
| ENSG00000232434 | AJM1 | 0.46 | 5.90E-03 | 4.88E-02 |
| ENSG00000278291 | AL161772.1 | 0.46 | 5.90E-03 | 4.88E-02 |
| ENSG00000108691 | CCL2 | -0.46 | 5.92E-03 | 4.89E-02 |

| Ensembl ID | Gene Name | ρ | Pval | Padj |
|---|---|---|---|---|
| ENSG00000161888 | SPC24 | -0.46 | 5.93E-03 | 4.89E-02 |
| ENSG00000084754 | HADHA | -0.46 | 5.94E-03 | 4.89E-02 |
| ENSG00000160323 | ADAMTS13 | 0.46 | 5.94E-03 | 4.89E-02 |
| ENSG00000236144 | TMEM147-AS1 | 0.46 | 5.94E-03 | 4.89E-02 |
| ENSG00000283635 | AC012485.3 | 0.46 | 5.95E-03 | 4.90E-02 |
| ENSG00000035141 | FAM136A | -0.46 | 5.97E-03 | 4.90E-02 |
| ENSG00000122694 | GLIPR2 | -0.46 | 5.97E-03 | 4.90E-02 |
| ENSG00000140937 | CDH11 | -0.46 | 5.97E-03 | 4.90E-02 |
| ENSG00000142949 | PTPRF | 0.46 | 5.97E-03 | 4.90E-02 |
| ENSG00000163686 | ABHD6 | -0.46 | 5.97E-03 | 4.90E-02 |
| ENSG00000168994 | PXDC1 | -0.46 | 5.97E-03 | 4.90E-02 |
| ENSG00000183114 | FAM43B | 0.46 | 5.97E-03 | 4.90E-02 |
| ENSG00000187498 | COL4A1 | -0.46 | 5.97E-03 | 4.90E-02 |
| ENSG00000228510 | AL359091.2 | 0.46 | 5.97E-03 | 4.90E-02 |
| ENSG00000234009 | RPL5P34 | -0.46 | 5.97E-03 | 4.90E-02 |
| ENSG00000270605 | AL353622.1 | 0.46 | 6.00E-03 | 4.92E-02 |
| ENSG00000015520 | NPC1L1 | 0.46 | 6.00E-03 | 4.92E-02 |
| ENSG00000100225 | FBXO7 | -0.46 | 6.01E-03 | 4.92E-02 |
| ENSG00000161558 | TMEM143 | 0.46 | 6.01E-03 | 4.92E-02 |
| ENSG00000173065 | FAM222B | 0.46 | 6.01E-03 | 4.92E-02 |
| ENSG00000181019 | NQO1 | -0.46 | 6.01E-03 | 4.92E-02 |
| ENSG00000272702 | AC010913.1 | 0.46 | 6.01E-03 | 4.92E-02 |
| ENSG00000276255 | AL136379.1 | 0.46 | 6.02E-03 | 4.93E-02 |
| ENSG00000048392 | RRM2B | -0.45 | 6.04E-03 | 4.93E-02 |
| ENSG00000164591 | MYOZ3 | 0.45 | 6.04E-03 | 4.93E-02 |
| ENSG00000182168 | UNC5C | 0.45 | 6.04E-03 | 4.93E-02 |
| ENSG00000182185 | RAD51B | 0.45 | 6.04E-03 | 4.93E-02 |
| ENSG00000279133 | AC018628.1 | 0.45 | 6.07E-03 | 4.95E-02 |
| ENSG00000131043 | AAR2 | -0.45 | 6.08E-03 | 4.95E-02 |
| ENSG00000175066 | GK5 | 0.45 | 6.08E-03 | 4.95E-02 |
| ENSG00000264112 | AC015813.1 | 0.45 | 6.08E-03 | 4.95E-02 |
| ENSG00000225264 | ZNRF2P2 | 0.45 | 6.09E-03 | 4.96E-02 |
| ENSG00000137968 | SLC44A5 | -0.45 | 6.11E-03 | 4.97E-02 |
| ENSG00000102900 | NUP93 | -0.45 | 6.11E-03 | 4.97E-02 |
| ENSG00000116455 | WDR77 | -0.45 | 6.11E-03 | 4.97E-02 |
| ENSG00000142784 | WDTC1 | 0.45 | 6.11E-03 | 4.97E-02 |
| ENSG00000197930 | ERO1A | -0.45 | 6.11E-03 | 4.97E-02 |
| ENSG00000226200 | SGMS1-AS1 | 0.45 | 6.11E-03 | 4.97E-02 |
| ENSG00000067177 | PHKA1 | 0.45 | 6.15E-03 | 4.99E-02 |
| ENSG00000154582 | ELOC | -0.45 | 6.15E-03 | 4.99E-02 |
| ENSG00000241553 | ARPC4 | -0.45 | 6.15E-03 | 4.99E-02 |
| ENSG00000175305 | CCNE2 | -0.45 | 6.18E-03 | 5.01E-02 |
| ENSG00000224743 | TEX26-AS1 | -0.45 | 6.18E-03 | 5.01E-02 |
| ENSG00000125656 | CLPP | -0.45 | 6.18E-03 | 5.01E-02 |
| ENSG00000126012 | KDM5C | 0.45 | 6.18E-03 | 5.01E-02 |

| Ensembl ID | Gene Name | ρ | Pval | Padj |
|---|---|---|---|---|
| ENSG00000157613 | CREB3L1 | -0.45 | 6.18E-03 | 5.01E-02 |
| ENSG00000243335 | KCTD7 | 0.45 | 6.18E-03 | 5.01E-02 |
| ENSG00000203808 | BVES-AS1 | 0.45 | 6.20E-03 | 5.02E-02 |
| ENSG00000167900 | TK1 | -0.45 | 6.22E-03 | 5.03E-02 |
| ENSG00000196440 | ARMCX4 | 0.45 | 6.22E-03 | 5.03E-02 |
| ENSG00000204815 | TTC25 | 0.45 | 6.22E-03 | 5.03E-02 |
| ENSG00000225300 | AL591623.1 | 0.45 | 6.22E-03 | 5.03E-02 |
| ENSG00000100060 | MFNG | -0.45 | 6.23E-03 | 5.04E-02 |
| ENSG00000274963 | RN7SL600P | 0.45 | 6.24E-03 | 5.04E-02 |
| ENSG00000280332 | AC020917.4 | 0.45 | 6.24E-03 | 5.04E-02 |
| ENSG00000235288 | AC099329.1 | 0.45 | 6.25E-03 | 5.04E-02 |
| ENSG00000134308 | YWHAQ | -0.45 | 6.25E-03 | 5.04E-02 |
| ENSG00000137492 | THAP12 | -0.45 | 6.25E-03 | 5.04E-02 |
| ENSG00000174173 | TRMT10C | -0.45 | 6.25E-03 | 5.04E-02 |
| ENSG00000185730 | ZNF696 | 0.45 | 6.25E-03 | 5.04E-02 |
| ENSG00000278434 | AC023830.3 | 0.45 | 6.26E-03 | 5.04E-02 |
| ENSG00000251661 | AC136475.1 | 0.45 | 6.26E-03 | 5.04E-02 |
| ENSG00000138463 | SLC49A4 | -0.45 | 6.29E-03 | 5.06E-02 |
| ENSG00000163812 | ZDHHC3 | -0.45 | 6.29E-03 | 5.06E-02 |
| ENSG00000178057 | NDUFAF3 | -0.45 | 6.29E-03 | 5.06E-02 |
| ENSG00000128596 | CCDC136 | 0.45 | 6.33E-03 | 5.08E-02 |
| ENSG00000137124 | ALDH1B1 | -0.45 | 6.33E-03 | 5.08E-02 |
| ENSG00000138640 | FAM13A | 0.45 | 6.33E-03 | 5.08E-02 |
| ENSG00000156222 | SLC28A1 | 0.45 | 6.33E-03 | 5.08E-02 |
| ENSG00000267125 | AC012615.3 | -0.45 | 6.34E-03 | 5.09E-02 |
| ENSG00000110906 | KCTD10 | -0.45 | 6.36E-03 | 5.10E-02 |
| ENSG00000147535 | PLPP5 | 0.45 | 6.36E-03 | 5.10E-02 |
| ENSG00000204161 | TMEM273 | -0.45 | 6.37E-03 | 5.11E-02 |
| ENSG00000185905 | C16orf54 | -0.45 | 6.39E-03 | 5.11E-02 |
| ENSG00000172236 | TPSAB1 | -0.45 | 6.39E-03 | 5.11E-02 |
| ENSG00000075702 | WDR62 | 0.45 | 6.40E-03 | 5.11E-02 |
| ENSG00000105750 | ZNF85 | -0.45 | 6.40E-03 | 5.11E-02 |
| ENSG00000110046 | ATG2A | 0.45 | 6.40E-03 | 5.11E-02 |
| ENSG00000119681 | LTBP2 | 0.45 | 6.40E-03 | 5.11E-02 |
| ENSG00000124207 | CSE1L | -0.45 | 6.40E-03 | 5.11E-02 |
| ENSG00000175283 | DOLK | -0.45 | 6.40E-03 | 5.11E-02 |
| ENSG00000186577 | SMIM29 | -0.45 | 6.40E-03 | 5.11E-02 |
| ENSG00000213983 | AP1G2 | 0.45 | 6.40E-03 | 5.11E-02 |
| ENSG00000229931 | AL137003.1 | 0.45 | 6.43E-03 | 5.13E-02 |
| ENSG00000077238 | IL4R | -0.45 | 6.44E-03 | 5.13E-02 |
| ENSG00000099282 | TSPAN15 | -0.45 | 6.44E-03 | 5.13E-02 |
| ENSG00000142166 | IFNAR1 | -0.45 | 6.44E-03 | 5.13E-02 |
| ENSG00000166855 | CLPX | 0.45 | 6.44E-03 | 5.13E-02 |
| ENSG00000260460 | AL365181.1 | 0.45 | 6.46E-03 | 5.15E-02 |
| ENSG00000097033 | SH3GLB1 | -0.45 | 6.47E-03 | 5.15E-02 |

| Ensembl ID | Gene Name | ρ | Pval | Padj |
|---|---|---|---|---|
| ENSG00000120278 | PLEKHG1 | -0.45 | 6.47E-03 | 5.15E-02 |
| ENSG00000138162 | TACC2 | 0.45 | 6.47E-03 | 5.15E-02 |
| ENSG00000240583 | AQP1 | -0.45 | 6.47E-03 | 5.15E-02 |
| ENSG00000250159 | AC106791.1 | 0.45 | 6.47E-03 | 5.15E-02 |
| ENSG00000267080 | ASB16-AS1 | 0.45 | 6.47E-03 | 5.15E-02 |
| ENSG00000273802 | HIST1H2BG | -0.45 | 6.48E-03 | 5.15E-02 |
| ENSG00000223849 | AL354893.1 | 0.45 | 6.49E-03 | 5.16E-02 |
| ENSG00000090006 | LTBP4 | 0.45 | 6.51E-03 | 5.16E-02 |
| ENSG00000103811 | CTSH | -0.45 | 6.51E-03 | 5.16E-02 |
| ENSG00000162889 | MAPKAPK2 | -0.45 | 6.51E-03 | 5.16E-02 |
| ENSG00000164032 | H2AFZ | -0.45 | 6.51E-03 | 5.16E-02 |
| ENSG00000112335 | SNX3 | -0.45 | 6.55E-03 | 5.18E-02 |
| ENSG00000139719 | VPS33A | -0.45 | 6.55E-03 | 5.18E-02 |
| ENSG00000214198 | TTC41P | 0.45 | 6.55E-03 | 5.18E-02 |
| ENSG00000224660 | SH3BP5-AS1 | 0.45 | 6.55E-03 | 5.18E-02 |
| ENSG00000267265 | AC011476.3 | 0.45 | 6.56E-03 | 5.19E-02 |
| ENSG00000116922 | C1orf109 | 0.45 | 6.59E-03 | 5.19E-02 |
| ENSG00000139190 | VAMP1 | 0.45 | 6.59E-03 | 5.19E-02 |
| ENSG00000139631 | CSAD | 0.45 | 6.59E-03 | 5.19E-02 |
| ENSG00000145743 | FBXL17 | 0.45 | 6.59E-03 | 5.19E-02 |
| ENSG00000155393 | HEATR3 | -0.45 | 6.59E-03 | 5.19E-02 |
| ENSG00000158669 | GPAT4 | -0.45 | 6.59E-03 | 5.19E-02 |
| ENSG00000171889 | MIR31HG | -0.45 | 6.59E-03 | 5.19E-02 |
| ENSG00000172831 | CES2 | 0.45 | 6.59E-03 | 5.19E-02 |
| ENSG00000196950 | SLC39A10 | -0.45 | 6.59E-03 | 5.19E-02 |
| ENSG00000233554 | B4GALT1-AS1 | -0.45 | 6.59E-03 | 5.19E-02 |
| ENSG00000228222 | AC073050.1 | 0.45 | 6.61E-03 | 5.21E-02 |
| ENSG00000013563 | DNASE1L1 | -0.45 | 6.62E-03 | 5.21E-02 |
| ENSG00000088543 | C3orf18 | 0.45 | 6.62E-03 | 5.21E-02 |
| ENSG00000141447 | OSBPL1A | 0.45 | 6.62E-03 | 5.21E-02 |
| ENSG00000142408 | CACNG8 | 0.45 | 6.63E-03 | 5.21E-02 |
| ENSG00000133574 | GIMAP4 | -0.45 | 6.64E-03 | 5.22E-02 |
| ENSG00000157368 | IL34 | -0.45 | 6.66E-03 | 5.23E-02 |
| ENSG00000057663 | ATG5 | -0.45 | 6.66E-03 | 5.23E-02 |
| ENSG00000134259 | NGF | -0.45 | 6.66E-03 | 5.23E-02 |
| ENSG00000237061 | AC100823.1 | 0.45 | 6.67E-03 | 5.23E-02 |
| ENSG00000205209 | SCGB2B2 | 0.45 | 6.69E-03 | 5.25E-02 |
| ENSG00000147443 | DOK2 | -0.45 | 6.70E-03 | 5.25E-02 |
| ENSG00000081985 | IL12RB2 | 0.45 | 6.70E-03 | 5.25E-02 |
| ENSG00000102878 | HSF4 | 0.45 | 6.70E-03 | 5.25E-02 |
| ENSG00000166963 | MAP1A | -0.45 | 6.70E-03 | 5.25E-02 |
| ENSG00000169871 | TRIM56 | 0.45 | 6.70E-03 | 5.25E-02 |
| ENSG00000237037 | NDUFA6-DT | 0.45 | 6.70E-03 | 5.25E-02 |
| ENSG00000112414 | ADGRG6 | 0.45 | 6.74E-03 | 5.27E-02 |
| ENSG00000163389 | POGLUT1 | -0.45 | 6.74E-03 | 5.27E-02 |

| Ensembl ID | Gene Name | ρ | Pval | Padj |
|---|---|---|---|---|
| ENSG00000167085 | PHB | -0.45 | 6.74E-03 | 5.27E-02 |
| ENSG00000213411 | RBM22P2 | 0.45 | 6.74E-03 | 5.27E-02 |
| ENSG00000126860 | EVI2A | -0.45 | 6.78E-03 | 5.29E-02 |
| ENSG00000135047 | CTSL | -0.45 | 6.78E-03 | 5.29E-02 |
| ENSG00000145476 | CYP4V2 | 0.45 | 6.78E-03 | 5.29E-02 |
| ENSG00000237668 | RPS15AP38 | -0.45 | 6.78E-03 | 5.29E-02 |
| ENSG00000235370 | DNM1P51 | 0.45 | 6.78E-03 | 5.29E-02 |
| ENSG00000281538 | AC254562.3 | 0.45 | 6.79E-03 | 5.29E-02 |
| ENSG00000109501 | WFS1 | 0.45 | 6.82E-03 | 5.31E-02 |
| ENSG00000204256 | BRD2 | 0.45 | 6.82E-03 | 5.31E-02 |
| ENSG00000260517 | AC009093.2 | 0.45 | 6.84E-03 | 5.33E-02 |
| ENSG00000000938 | FGR | -0.45 | 6.85E-03 | 5.33E-02 |
| ENSG00000021300 | PLEKHB1 | 0.45 | 6.85E-03 | 5.33E-02 |
| ENSG00000066294 | CD84 | -0.45 | 6.85E-03 | 5.33E-02 |
| ENSG00000152558 | TMEM123 | -0.45 | 6.85E-03 | 5.33E-02 |
| ENSG00000227953 | LINC01341 | 0.45 | 6.85E-03 | 5.33E-02 |
| ENSG00000261113 | AC009034.1 | -0.45 | 6.86E-03 | 5.33E-02 |
| ENSG00000248712 | CCDC153 | -0.45 | 6.88E-03 | 5.35E-02 |
| ENSG00000130176 | CNN1 | -0.45 | 6.89E-03 | 5.35E-02 |
| ENSG00000204352 | C9orf129 | 0.45 | 6.90E-03 | 5.36E-02 |
| ENSG00000272779 | AC245060.4 | -0.45 | 6.91E-03 | 5.36E-02 |
| ENSG00000264635 | AP001020.2 | 0.45 | 6.92E-03 | 5.36E-02 |
| ENSG00000170290 | SLN | -0.45 | 6.92E-03 | 5.36E-02 |
| ENSG00000032389 | EIPR1 | -0.45 | 6.93E-03 | 5.36E-02 |
| ENSG00000100242 | SUN2 | 0.45 | 6.93E-03 | 5.36E-02 |
| ENSG00000137409 | MTCH1 | -0.45 | 6.93E-03 | 5.36E-02 |
| ENSG00000169116 | PARM1 | 0.45 | 6.93E-03 | 5.36E-02 |
| ENSG00000170043 | TRAPPC1 | -0.45 | 6.93E-03 | 5.36E-02 |
| ENSG00000181982 | CCDC149 | 0.45 | 6.93E-03 | 5.36E-02 |
| ENSG00000256356 | HSPA8P5 | -0.45 | 6.93E-03 | 5.36E-02 |
| ENSG00000000419 | DPM1 | -0.45 | 6.97E-03 | 5.37E-02 |
| ENSG00000033050 | ABCF2 | -0.45 | 6.97E-03 | 5.37E-02 |
| ENSG00000111077 | TNS2 | 0.45 | 6.97E-03 | 5.37E-02 |
| ENSG00000138316 | ADAMTS14 | -0.45 | 6.97E-03 | 5.37E-02 |
| ENSG00000164077 | MON1A | -0.45 | 6.97E-03 | 5.37E-02 |
| ENSG00000170035 | UBE2E3 | -0.45 | 6.97E-03 | 5.37E-02 |
| ENSG00000198612 | COPS8 | -0.45 | 6.97E-03 | 5.37E-02 |
| ENSG00000198840 | MT-ND3 | -0.45 | 6.97E-03 | 5.37E-02 |
| ENSG00000204138 | PHACTR4 | 0.45 | 6.97E-03 | 5.37E-02 |
| ENSG00000089356 | FXYD3 | 0.45 | 6.99E-03 | 5.38E-02 |
| ENSG00000258745 | AL358334.3 | 0.45 | 7.00E-03 | 5.39E-02 |
| ENSG00000255458 | AC108471.2 | 0.45 | 7.01E-03 | 5.39E-02 |
| ENSG00000266967 | AARSD1 | 0.45 | 7.01E-03 | 5.39E-02 |
| ENSG00000146574 | CCZ1B | -0.45 | 7.01E-03 | 5.39E-02 |
| ENSG00000204128 | C2orf72 | 0.45 | 7.01E-03 | 5.39E-02 |

| Ensembl ID | Gene Name | ρ | Pval | Padj |
|---|---|---|---|---|
| ENSG00000257354 | AC048341.1 | 0.45 | 7.01E-03 | 5.39E-02 |
| ENSG00000129167 | TPH1 | 0.45 | 7.03E-03 | 5.40E-02 |
| ENSG00000176222 | ZNF404 | 0.45 | 7.03E-03 | 5.40E-02 |
| ENSG00000054803 | CBLN4 | -0.45 | 7.05E-03 | 5.41E-02 |
| ENSG00000215067 | ALOX12-AS1 | 0.45 | 7.05E-03 | 5.41E-02 |
| ENSG00000198221 | AFDN-DT | 0.45 | 7.06E-03 | 5.41E-02 |
| ENSG00000130032 | PRRG3 | -0.45 | 7.07E-03 | 5.42E-02 |
| ENSG00000277299 | AC084876.1 | -0.45 | 7.07E-03 | 5.42E-02 |
| ENSG00000155368 | DBI | -0.45 | 7.09E-03 | 5.43E-02 |
| ENSG00000161010 | MRNIP | 0.45 | 7.09E-03 | 5.43E-02 |
| ENSG00000269378 | AC022149.1 | -0.45 | 7.09E-03 | 5.43E-02 |
| ENSG00000107819 | SFXN3 | -0.45 | 7.13E-03 | 5.46E-02 |
| ENSG00000109805 | NCAPG | -0.45 | 7.13E-03 | 5.46E-02 |
| ENSG00000168077 | SCARA3 | 0.45 | 7.13E-03 | 5.46E-02 |
| ENSG00000115919 | KYNU | -0.45 | 7.15E-03 | 5.47E-02 |
| ENSG00000067365 | METTL22 | 0.45 | 7.17E-03 | 5.47E-02 |
| ENSG00000107562 | CXCL12 | -0.45 | 7.17E-03 | 5.47E-02 |
| ENSG00000108469 | RECQL5 | 0.45 | 7.17E-03 | 5.47E-02 |
| ENSG00000123144 | TRIR | -0.45 | 7.17E-03 | 5.47E-02 |
| ENSG00000164692 | COL1A2 | -0.45 | 7.17E-03 | 5.47E-02 |
| ENSG00000176853 | FAM91A1 | -0.45 | 7.17E-03 | 5.47E-02 |
| ENSG00000282772 | AL358790.1 | 0.45 | 7.20E-03 | 5.49E-02 |
| ENSG00000100567 | PSMA3 | -0.45 | 7.21E-03 | 5.49E-02 |
| ENSG00000116161 | CACYBP | -0.45 | 7.21E-03 | 5.49E-02 |
| ENSG00000139998 | RAB15 | -0.45 | 7.21E-03 | 5.49E-02 |
| ENSG00000166401 | SERPINB8 | -0.45 | 7.21E-03 | 5.49E-02 |
| ENSG00000198729 | PPP1R14C | -0.45 | 7.21E-03 | 5.49E-02 |
| ENSG00000214176 | PLEKHM1P1 | 0.45 | 7.21E-03 | 5.49E-02 |
| ENSG00000274128 | AC027807.2 | 0.45 | 7.24E-03 | 5.50E-02 |
| ENSG00000197061 | HIST1H4C | -0.45 | 7.25E-03 | 5.51E-02 |
| ENSG00000110880 | CORO1C | -0.45 | 7.25E-03 | 5.51E-02 |
| ENSG00000126464 | PRR12 | 0.45 | 7.25E-03 | 5.51E-02 |
| ENSG00000256269 | HMBS | -0.45 | 7.25E-03 | 5.51E-02 |
| ENSG00000100591 | AHSA1 | -0.45 | 7.30E-03 | 5.53E-02 |
| ENSG00000101213 | PTK6 | 0.45 | 7.30E-03 | 5.53E-02 |
| ENSG00000128692 | EIF2S2P4 | -0.45 | 7.30E-03 | 5.53E-02 |
| ENSG00000134871 | COL4A2 | -0.45 | 7.30E-03 | 5.53E-02 |
| ENSG00000136950 | ARPC5L | -0.45 | 7.30E-03 | 5.53E-02 |
| ENSG00000153395 | LPCAT1 | -0.45 | 7.30E-03 | 5.53E-02 |
| ENSG00000187678 | SPRY4 | 0.45 | 7.30E-03 | 5.53E-02 |
| ENSG00000125848 | FLRT3 | -0.45 | 7.32E-03 | 5.54E-02 |
| ENSG00000066427 | ATXN3 | 0.45 | 7.34E-03 | 5.54E-02 |
| ENSG00000160439 | RDH13 | 0.45 | 7.34E-03 | 5.54E-02 |
| ENSG00000173660 | UQCRH | -0.45 | 7.34E-03 | 5.54E-02 |
| ENSG00000197226 | TBC1D9B | 0.45 | 7.34E-03 | 5.54E-02 |

| Ensembl ID | Gene Name | ρ | Pval | Padj |
|---|---|---|---|---|
| ENSG00000214140 | PRCD | 0.45 | 7.34E-03 | 5.54E-02 |
| ENSG00000230424 | EMC1-AS1 | 0.45 | 7.34E-03 | 5.54E-02 |
| ENSG00000159231 | CBR3 | -0.45 | 7.38E-03 | 5.56E-02 |
| ENSG00000164163 | ABCE1 | -0.45 | 7.38E-03 | 5.56E-02 |
| ENSG00000166250 | CLMP | -0.45 | 7.38E-03 | 5.56E-02 |
| ENSG00000189184 | PCDH18 | -0.45 | 7.38E-03 | 5.56E-02 |
| ENSG00000214279 | SCART1 | 0.45 | 7.38E-03 | 5.56E-02 |
| ENSG00000242951 | AC007182.2 | -0.45 | 7.38E-03 | 5.56E-02 |
| ENSG00000271614 | ATP2B1-AS1 | 0.45 | 7.38E-03 | 5.56E-02 |
| ENSG00000272983 | AL117339.4 | 0.45 | 7.38E-03 | 5.56E-02 |
| ENSG00000266002 | AC091059.1 | 0.44 | 7.40E-03 | 5.57E-02 |
| ENSG00000085449 | WDFY1 | -0.44 | 7.42E-03 | 5.57E-02 |
| ENSG00000106546 | AHR | -0.44 | 7.42E-03 | 5.57E-02 |
| ENSG00000135077 | HAVCR2 | -0.44 | 7.42E-03 | 5.57E-02 |
| ENSG00000136982 | DSCC1 | -0.44 | 7.42E-03 | 5.57E-02 |
| ENSG00000150281 | CTF1 | 0.44 | 7.42E-03 | 5.57E-02 |
| ENSG00000235257 | ITGA9-AS1 | 0.44 | 7.42E-03 | 5.57E-02 |
| ENSG00000241511 | RPS15AP24 | -0.44 | 7.42E-03 | 5.57E-02 |
| ENSG00000250251 | PKD1P6 | 0.44 | 7.42E-03 | 5.57E-02 |
| ENSG00000235358 | AC093151.3 | 0.44 | 7.42E-03 | 5.57E-02 |
| ENSG00000279344 | AC007342.6 | 0.44 | 7.45E-03 | 5.58E-02 |
| ENSG00000120533 | ENY2 | -0.44 | 7.46E-03 | 5.58E-02 |
| ENSG00000167397 | VKORC1 | -0.44 | 7.46E-03 | 5.58E-02 |
| ENSG00000168724 | DNAJC21 | -0.44 | 7.46E-03 | 5.58E-02 |
| ENSG00000176945 | MUC20 | 0.44 | 7.46E-03 | 5.58E-02 |
| ENSG00000234231 | AC093616.1 | -0.44 | 7.46E-03 | 5.58E-02 |
| ENSG00000241837 | ATP5PO | -0.44 | 7.46E-03 | 5.58E-02 |
| ENSG00000246067 | RAB30-AS1 | 0.44 | 7.46E-03 | 5.58E-02 |
| ENSG00000270344 | POC1B-AS1 | 0.44 | 7.46E-03 | 5.58E-02 |
| ENSG00000002834 | LASP1 | -0.44 | 7.50E-03 | 5.60E-02 |
| ENSG00000090905 | TNRC6A | 0.44 | 7.50E-03 | 5.60E-02 |
| ENSG00000135439 | AGAP2 | -0.44 | 7.50E-03 | 5.60E-02 |
| ENSG00000271335 | AL117336.2 | 0.44 | 7.50E-03 | 5.60E-02 |
| ENSG00000137269 | LRRC1 | -0.44 | 7.55E-03 | 5.63E-02 |
| ENSG00000151726 | ACSL1 | -0.44 | 7.55E-03 | 5.63E-02 |
| ENSG00000166123 | GPT2 | 0.44 | 7.55E-03 | 5.63E-02 |
| ENSG00000148735 | PLEKHS1 | -0.44 | 7.55E-03 | 5.63E-02 |
| ENSG00000117091 | CD48 | -0.44 | 7.58E-03 | 5.65E-02 |
| ENSG00000267470 | ZNF571-AS1 | 0.44 | 7.58E-03 | 5.65E-02 |
| ENSG00000087245 | MMP2 | -0.44 | 7.59E-03 | 5.65E-02 |
| ENSG00000139726 | DENR | -0.44 | 7.59E-03 | 5.65E-02 |
| ENSG00000163257 | DCAF16 | 0.44 | 7.59E-03 | 5.65E-02 |
| ENSG00000175550 | DRAP1 | -0.44 | 7.59E-03 | 5.65E-02 |
| ENSG00000049249 | TNFRSF9 | -0.44 | 7.62E-03 | 5.67E-02 |
| ENSG00000117010 | ZNF684 | -0.44 | 7.63E-03 | 5.67E-02 |

| Ensembl ID | Gene Name | ρ | Pval | Padj |
|---|---|---|---|---|
| ENSG00000136518 | ACTL6A | -0.44 | 7.63E-03 | 5.67E-02 |
| ENSG00000154118 | JPH3 | 0.44 | 7.63E-03 | 5.67E-02 |
| ENSG00000211459 | MT-RNR1 | -0.44 | 7.63E-03 | 5.67E-02 |
| ENSG00000239763 | AC009120.1 | 0.44 | 7.66E-03 | 5.68E-02 |
| ENSG00000125743 | SNRPD2 | -0.44 | 7.67E-03 | 5.69E-02 |
| ENSG00000130935 | NOL11 | -0.44 | 7.67E-03 | 5.69E-02 |
| ENSG00000132965 | ALOX5AP | -0.44 | 7.67E-03 | 5.69E-02 |
| ENSG00000174547 | MRPL11 | -0.44 | 7.67E-03 | 5.69E-02 |
| ENSG00000105609 | LILRB5 | -0.44 | 7.71E-03 | 5.70E-02 |
| ENSG00000138326 | RPS24 | -0.44 | 7.72E-03 | 5.70E-02 |
| ENSG00000146830 | GIGYF1 | 0.44 | 7.72E-03 | 5.70E-02 |
| ENSG00000147669 | POLR2K | -0.44 | 7.72E-03 | 5.70E-02 |
| ENSG00000160588 | MPZL3 | -0.44 | 7.72E-03 | 5.70E-02 |
| ENSG00000162063 | CCNF | -0.44 | 7.72E-03 | 5.70E-02 |
| ENSG00000166986 | MARS | -0.44 | 7.72E-03 | 5.70E-02 |
| ENSG00000169976 | SF3B5 | -0.44 | 7.72E-03 | 5.70E-02 |
| ENSG00000173530 | TNFRSF10D | -0.44 | 7.72E-03 | 5.70E-02 |
| ENSG00000181610 | MRPS23 | -0.44 | 7.72E-03 | 5.70E-02 |
| ENSG00000138778 | CENPE | -0.44 | 7.76E-03 | 5.72E-02 |
| ENSG00000185811 | IKZF1 | -0.44 | 7.76E-03 | 5.72E-02 |
| ENSG00000102390 | PBDC1 | -0.44 | 7.76E-03 | 5.72E-02 |
| ENSG00000168209 | DDIT4 | 0.44 | 7.76E-03 | 5.72E-02 |
| ENSG00000237350 | CDC42P6 | -0.44 | 7.76E-03 | 5.72E-02 |
| ENSG00000070193 | FGF10 | -0.44 | 7.77E-03 | 5.73E-02 |
| ENSG00000261175 | LINC02188 | 0.44 | 7.79E-03 | 5.74E-02 |
| ENSG00000118855 | MFSD1 | -0.44 | 7.80E-03 | 5.74E-02 |
| ENSG00000161714 | PLCD3 | 0.44 | 7.80E-03 | 5.74E-02 |
| ENSG00000231770 | TMEM44-AS1 | 0.44 | 7.80E-03 | 5.74E-02 |
| ENSG00000234287 | AC099560.2 | -0.44 | 7.80E-03 | 5.74E-02 |
| ENSG00000231873 | AC099560.1 | -0.44 | 7.82E-03 | 5.75E-02 |
| ENSG00000276966 | HIST1H4E | -0.44 | 7.83E-03 | 5.75E-02 |
| ENSG00000234964 | FABP5P7 | -0.44 | 7.84E-03 | 5.76E-02 |
| ENSG00000006704 | GTF2IRD1 | 0.44 | 7.85E-03 | 5.76E-02 |
| ENSG00000105185 | PDCD5 | -0.44 | 7.85E-03 | 5.76E-02 |
| ENSG00000198198 | SZT2 | 0.44 | 7.85E-03 | 5.76E-02 |
| ENSG00000132581 | SDF2 | -0.44 | 7.89E-03 | 5.77E-02 |
| ENSG00000134330 | IAH1 | -0.44 | 7.89E-03 | 5.77E-02 |
| ENSG00000139636 | LMBR1L | 0.44 | 7.89E-03 | 5.77E-02 |
| ENSG00000140740 | UQCRC2 | -0.44 | 7.89E-03 | 5.77E-02 |
| ENSG00000144824 | PHLDB2 | -0.44 | 7.89E-03 | 5.77E-02 |
| ENSG00000165527 | ARF6 | -0.44 | 7.89E-03 | 5.77E-02 |
| ENSG00000169230 | PRELID1 | -0.44 | 7.89E-03 | 5.77E-02 |
| ENSG00000169756 | LIMS1 | -0.44 | 7.89E-03 | 5.77E-02 |
| ENSG00000185163 | DDX51 | 0.44 | 7.89E-03 | 5.77E-02 |
| ENSG00000213888 | LINC01521 | 0.44 | 7.89E-03 | 5.77E-02 |

| Ensembl ID | Gene Name | ρ | Pval | Padj |
|---|---|---|---|---|
| ENSG00000121966 | CXCR4 | -0.44 | 7.91E-03 | 5.78E-02 |
| ENSG00000267370 | AC008752.3 | 0.44 | 7.93E-03 | 5.79E-02 |
| ENSG00000105875 | WDR91 | 0.44 | 7.94E-03 | 5.79E-02 |
| ENSG00000136161 | RCBTB2 | -0.44 | 7.94E-03 | 5.79E-02 |
| ENSG00000149187 | CELF1 | -0.44 | 7.94E-03 | 5.79E-02 |
| ENSG00000163596 | ICA1L | 0.44 | 7.98E-03 | 5.82E-02 |
| ENSG00000163945 | UVSSA | 0.44 | 7.98E-03 | 5.82E-02 |
| ENSG00000166788 | SAAL1 | -0.44 | 7.98E-03 | 5.82E-02 |
| ENSG00000180044 | C3orf80 | -0.44 | 8.00E-03 | 5.83E-02 |
| ENSG00000185189 | NRBP2 | 0.44 | 8.03E-03 | 5.85E-02 |
| ENSG00000175489 | LRRC25 | -0.44 | 8.05E-03 | 5.86E-02 |
| ENSG00000127526 | SLC35E1 | 0.44 | 8.07E-03 | 5.87E-02 |
| ENSG00000147872 | PLIN2 | -0.44 | 8.07E-03 | 5.87E-02 |
| ENSG00000156795 | WDYHV1 | -0.44 | 8.07E-03 | 5.87E-02 |
| ENSG00000166343 | MSS51 | 0.44 | 8.07E-03 | 5.87E-02 |
| ENSG00000260314 | MRC1 | -0.44 | 8.07E-03 | 5.87E-02 |
| ENSG00000254192 | AC011365.2 | -0.44 | 8.07E-03 | 5.87E-02 |
| ENSG00000146521 | LINC01558 | 0.44 | 8.11E-03 | 5.88E-02 |
| ENSG00000106483 | SFRP4 | -0.44 | 8.12E-03 | 5.88E-02 |
| ENSG00000125870 | SNRPB2 | -0.44 | 8.12E-03 | 5.88E-02 |
| ENSG00000137055 | PLAA | -0.44 | 8.12E-03 | 5.88E-02 |
| ENSG00000168487 | BMP1 | -0.44 | 8.12E-03 | 5.88E-02 |
| ENSG00000188612 | SUMO2 | -0.44 | 8.12E-03 | 5.88E-02 |
| ENSG00000245552 | AP000787.1 | 0.44 | 8.12E-03 | 5.88E-02 |
| ENSG00000149571 | KIRREL3 | -0.44 | 8.14E-03 | 5.89E-02 |
| ENSG00000102265 | TIMP1 | -0.44 | 8.16E-03 | 5.90E-02 |
| ENSG00000159377 | PSMB4 | -0.44 | 8.16E-03 | 5.90E-02 |
| ENSG00000161057 | PSMC2 | -0.44 | 8.16E-03 | 5.90E-02 |
| ENSG00000164938 | TP53INP1 | -0.44 | 8.16E-03 | 5.90E-02 |
| ENSG00000198937 | CCDC167 | -0.44 | 8.16E-03 | 5.90E-02 |
| ENSG00000242086 | MUC20-OT1 | 0.44 | 8.16E-03 | 5.90E-02 |
| ENSG00000110274 | CEP164 | 0.44 | 8.21E-03 | 5.92E-02 |
| ENSG00000119938 | PPP1R3C | 0.44 | 8.21E-03 | 5.92E-02 |
| ENSG00000155366 | RHOC | -0.44 | 8.21E-03 | 5.92E-02 |
| ENSG00000173209 | AHSA2P | 0.44 | 8.21E-03 | 5.92E-02 |
| ENSG00000272154 | AC244517.2 | 0.44 | 8.23E-03 | 5.94E-02 |
| ENSG00000103067 | ESRP2 | 0.44 | 8.23E-03 | 5.94E-02 |
| ENSG00000042753 | AP2S1 | -0.44 | 8.25E-03 | 5.94E-02 |
| ENSG00000166503 | HDGFL3 | -0.44 | 8.25E-03 | 5.94E-02 |
| ENSG00000170315 | UBB | -0.44 | 8.25E-03 | 5.94E-02 |
| ENSG00000204619 | PPP1R11 | -0.44 | 8.25E-03 | 5.94E-02 |
| ENSG00000132205 | EMILIN2 | -0.44 | 8.30E-03 | 5.96E-02 |
| ENSG00000135905 | DOCK10 | -0.44 | 8.30E-03 | 5.96E-02 |
| ENSG00000136999 | CCN3 | -0.44 | 8.30E-03 | 5.96E-02 |
| ENSG00000149131 | SERPING1 | 0.44 | 8.30E-03 | 5.96E-02 |

| Ensembl ID | Gene Name | ρ | Pval | Padj |
|---|---|---|---|---|
| ENSG00000204842 | ATXN2 | 0.44 | 8.30E-03 | 5.96E-02 |
| ENSG00000214029 | ZNF891 | 0.44 | 8.30E-03 | 5.96E-02 |
| ENSG00000095970 | TREM2 | -0.44 | 8.31E-03 | 5.97E-02 |
| ENSG00000224577 | LINC01117 | 0.44 | 8.32E-03 | 5.97E-02 |
| ENSG00000104044 | OCA2 | 0.44 | 8.33E-03 | 5.98E-02 |
| ENSG00000135414 | GDF11 | -0.44 | 8.34E-03 | 5.98E-02 |
| ENSG00000170959 | DCDC1 | 0.44 | 8.34E-03 | 5.98E-02 |
| ENSG00000205476 | CCDC85C | 0.44 | 8.34E-03 | 5.98E-02 |
| ENSG00000172578 | KLHL6 | -0.44 | 8.36E-03 | 6.00E-02 |
| ENSG00000182463 | TSHZ2 | -0.44 | 8.38E-03 | 6.00E-02 |
| ENSG00000104325 | DECR1 | -0.44 | 8.39E-03 | 6.00E-02 |
| ENSG00000106012 | IQCE | 0.44 | 8.39E-03 | 6.00E-02 |
| ENSG00000123610 | TNFAIP6 | -0.44 | 8.39E-03 | 6.00E-02 |
| ENSG00000164167 | LSM6 | -0.44 | 8.39E-03 | 6.00E-02 |
| ENSG00000172379 | ARNT2 | -0.44 | 8.39E-03 | 6.00E-02 |
| ENSG00000187609 | EXD3 | 0.44 | 8.39E-03 | 6.00E-02 |
| ENSG00000184357 | HIST1H1B | -0.44 | 8.43E-03 | 6.01E-02 |
| ENSG00000112511 | PHF1 | 0.44 | 8.44E-03 | 6.01E-02 |
| ENSG00000112902 | SEMA5A | -0.44 | 8.44E-03 | 6.01E-02 |
| ENSG00000125384 | PTGER2 | -0.44 | 8.44E-03 | 6.01E-02 |
| ENSG00000132680 | KHDC4 | 0.44 | 8.44E-03 | 6.01E-02 |
| ENSG00000171155 | C1GALT1C1 | -0.44 | 8.44E-03 | 6.01E-02 |
| ENSG00000180758 | GPR157 | 0.44 | 8.44E-03 | 6.01E-02 |
| ENSG00000186409 | CCDC30 | 0.44 | 8.44E-03 | 6.01E-02 |
| ENSG00000204392 | LSM2 | -0.44 | 8.44E-03 | 6.01E-02 |
| ENSG00000111245 | MYL2 | -0.44 | 8.44E-03 | 6.01E-02 |
| ENSG00000134996 | OSTF1 | -0.44 | 8.48E-03 | 6.04E-02 |
| ENSG00000137478 | FCHSD2 | 0.44 | 8.48E-03 | 6.04E-02 |
| ENSG00000168542 | COL3A1 | -0.44 | 8.48E-03 | 6.04E-02 |
| ENSG00000226976 | COX6A1P2 | -0.44 | 8.48E-03 | 6.04E-02 |
| ENSG00000178150 | ZNF114 | 0.44 | 8.50E-03 | 6.05E-02 |
| ENSG00000248671 | AP002495.1 | 0.44 | 8.51E-03 | 6.05E-02 |
| ENSG00000072195 | SPEG | 0.44 | 8.53E-03 | 6.06E-02 |
| ENSG00000145247 | OCIAD2 | -0.44 | 8.53E-03 | 6.06E-02 |
| ENSG00000182175 | RGMA | 0.44 | 8.53E-03 | 6.06E-02 |
| ENSG00000246662 | LINC00535 | 0.44 | 8.53E-03 | 6.06E-02 |
| ENSG00000262902 | MTCO1P40 | -0.44 | 8.54E-03 | 6.06E-02 |
| ENSG00000123352 | SPATS2 | 0.44 | 8.58E-03 | 6.08E-02 |
| ENSG00000135457 | TFCP2 | -0.44 | 8.58E-03 | 6.08E-02 |
| ENSG00000227039 | ITGB2-AS1 | -0.44 | 8.59E-03 | 6.09E-02 |
| ENSG00000277728 | AC097641.2 | 0.44 | 8.59E-03 | 6.09E-02 |
| ENSG00000267419 | AC011477.1 | 0.44 | 8.60E-03 | 6.09E-02 |
| ENSG00000109819 | PPARGC1A | 0.44 | 8.62E-03 | 6.10E-02 |
| ENSG00000132423 | COQ3 | -0.44 | 8.62E-03 | 6.10E-02 |
| ENSG00000132688 | NES | -0.44 | 8.62E-03 | 6.10E-02 |

| Ensembl ID | Gene Name | ρ | Pval | Padj |
|---|---|---|---|---|
| ENSG00000135931 | ARMC9 | 0.44 | 8.62E-03 | 6.10E-02 |
| ENSG00000172331 | BPGM | -0.44 | 8.62E-03 | 6.10E-02 |
| ENSG00000133561 | GIMAP6 | -0.44 | 8.63E-03 | 6.10E-02 |
| ENSG00000109265 | KIAA1211 | 0.44 | 8.64E-03 | 6.11E-02 |
| ENSG00000182612 | TSPAN10 | 0.44 | 8.67E-03 | 6.13E-02 |
| ENSG00000112290 | WASF1 | -0.44 | 8.69E-03 | 6.14E-02 |
| ENSG00000254941 | AP000866.4 | 0.44 | 8.70E-03 | 6.14E-02 |
| ENSG00000138190 | EXOC6 | -0.44 | 8.72E-03 | 6.15E-02 |
| ENSG00000141376 | BCAS3 | 0.44 | 8.72E-03 | 6.15E-02 |
| ENSG00000143515 | ATP8B2 | 0.44 | 8.72E-03 | 6.15E-02 |
| ENSG00000146729 | NIPSNAP2 | -0.44 | 8.72E-03 | 6.15E-02 |
| ENSG00000132182 | NUP210 | -0.44 | 8.75E-03 | 6.17E-02 |
| ENSG00000100441 | KHNYN | 0.44 | 8.77E-03 | 6.17E-02 |
| ENSG00000111863 | ADTRP | -0.44 | 8.77E-03 | 6.17E-02 |
| ENSG00000158301 | GPRASP2 | 0.44 | 8.77E-03 | 6.17E-02 |
| ENSG00000163694 | RBM47 | -0.44 | 8.77E-03 | 6.17E-02 |
| ENSG00000236723 | AL606760.2 | 0.44 | 8.81E-03 | 6.19E-02 |
| ENSG00000138660 | AP1AR | -0.44 | 8.82E-03 | 6.19E-02 |
| ENSG00000162517 | PEF1 | -0.44 | 8.82E-03 | 6.19E-02 |
| ENSG00000174238 | PITPNA | -0.44 | 8.82E-03 | 6.19E-02 |
| ENSG00000185238 | PRMT3 | -0.44 | 8.82E-03 | 6.19E-02 |
| ENSG00000272734 | ADIRF-AS1 | 0.44 | 8.82E-03 | 6.19E-02 |
| ENSG00000114209 | PDCD10 | -0.44 | 8.86E-03 | 6.22E-02 |
| ENSG00000135241 | PNPLA8 | -0.44 | 8.86E-03 | 6.22E-02 |
| ENSG00000164414 | SLC35A1 | -0.44 | 8.86E-03 | 6.22E-02 |
| ENSG00000204304 | PBX2 | 0.44 | 8.86E-03 | 6.22E-02 |
| ENSG00000154065 | ANKRD29 | -0.44 | 8.88E-03 | 6.23E-02 |
| ENSG00000115946 | PNO1 | -0.44 | 8.91E-03 | 6.23E-02 |
| ENSG00000137094 | DNAJB5 | -0.44 | 8.91E-03 | 6.23E-02 |
| ENSG00000157322 | CLEC18A | 0.44 | 8.91E-03 | 6.23E-02 |
| ENSG00000165282 | PIGO | 0.44 | 8.91E-03 | 6.23E-02 |
| ENSG00000165916 | PSMC3 | -0.44 | 8.91E-03 | 6.23E-02 |
| ENSG00000169446 | MMGT1 | -0.44 | 8.91E-03 | 6.23E-02 |
| ENSG00000169991 | IFFO2 | -0.44 | 8.91E-03 | 6.23E-02 |
| ENSG00000188493 | C19orf54 | 0.44 | 8.91E-03 | 6.23E-02 |
| ENSG00000218809 | AL391903.1 | -0.44 | 8.93E-03 | 6.24E-02 |
| ENSG00000012963 | UBR7 | -0.44 | 8.96E-03 | 6.25E-02 |
| ENSG00000108001 | EBF3 | -0.44 | 8.96E-03 | 6.25E-02 |
| ENSG00000129353 | SLC44A2 | 0.44 | 8.96E-03 | 6.25E-02 |
| ENSG00000131495 | NDUFA2 | -0.44 | 8.96E-03 | 6.25E-02 |
| ENSG00000135547 | HEY2 | -0.44 | 8.96E-03 | 6.25E-02 |
| ENSG00000167371 | PRRT2 | 0.44 | 8.96E-03 | 6.25E-02 |
| ENSG00000230673 | PABPC1P3 | -0.44 | 8.96E-03 | 6.25E-02 |
| ENSG00000181885 | CLDN7 | -0.44 | 9.01E-03 | 6.27E-02 |
| ENSG00000165457 | FOLR2 | -0.44 | 9.01E-03 | 6.27E-02 |

| Ensembl ID | Gene Name | ρ | Pval | Padj |
|---|---|---|---|---|
| ENSG00000121900 | TMEM54 | -0.44 | 9.01E-03 | 6.27E-02 |
| ENSG00000133313 | CNDP2 | -0.44 | 9.01E-03 | 6.27E-02 |
| ENSG00000196581 | AJAP1 | -0.44 | 9.01E-03 | 6.27E-02 |
| ENSG00000204231 | RXRB | 0.44 | 9.01E-03 | 6.27E-02 |
| ENSG00000008517 | IL32 | -0.43 | 9.06E-03 | 6.29E-02 |
| ENSG00000103121 | CMC2 | -0.43 | 9.06E-03 | 6.29E-02 |
| ENSG00000125863 | MKKS | -0.43 | 9.06E-03 | 6.29E-02 |
| ENSG00000134697 | GNL2 | -0.43 | 9.06E-03 | 6.29E-02 |
| ENSG00000146233 | CYP39A1 | 0.43 | 9.06E-03 | 6.29E-02 |
| ENSG00000173253 | DMRT2 | 0.43 | 9.06E-03 | 6.29E-02 |
| ENSG00000204103 | MAFB | -0.43 | 9.06E-03 | 6.29E-02 |
| ENSG00000214973 | CHCHD3P3 | -0.43 | 9.06E-03 | 6.29E-02 |
| ENSG00000171462 | DLK2 | -0.43 | 9.10E-03 | 6.31E-02 |
| ENSG00000183889 | AC138969.1 | 0.43 | 9.11E-03 | 6.31E-02 |
| ENSG00000131969 | ABHD12B | 0.43 | 9.11E-03 | 6.31E-02 |
| ENSG00000198843 | SELENOT | -0.43 | 9.11E-03 | 6.31E-02 |
| ENSG00000257243 | AC020612.1 | 0.43 | 9.16E-03 | 6.33E-02 |
| ENSG00000064652 | SNX24 | -0.43 | 9.16E-03 | 6.33E-02 |
| ENSG00000136986 | DERL1 | -0.43 | 9.16E-03 | 6.33E-02 |
| ENSG00000204604 | ZNF468 | -0.43 | 9.16E-03 | 6.33E-02 |
| ENSG00000210082 | MT-RNR2 | -0.43 | 9.16E-03 | 6.33E-02 |
| ENSG00000214784 | AC010468.1 | -0.43 | 9.16E-03 | 6.33E-02 |
| ENSG00000248932 | AC097103.2 | 0.43 | 9.16E-03 | 6.33E-02 |
| ENSG00000100941 | PNN | 0.43 | 9.21E-03 | 6.35E-02 |
| ENSG00000109861 | CTSC | -0.43 | 9.21E-03 | 6.35E-02 |
| ENSG00000116044 | NFE2L2 | -0.43 | 9.21E-03 | 6.35E-02 |
| ENSG00000131378 | RFTN1 | -0.43 | 9.21E-03 | 6.35E-02 |
| ENSG00000132570 | PCBD2 | 0.43 | 9.21E-03 | 6.35E-02 |
| ENSG00000149636 | DSN1 | -0.43 | 9.21E-03 | 6.35E-02 |
| ENSG00000163866 | SMIM12 | -0.43 | 9.21E-03 | 6.35E-02 |
| ENSG00000186976 | EFCAB6 | 0.43 | 9.21E-03 | 6.35E-02 |
| ENSG00000218426 | AL590867.2 | -0.43 | 9.21E-03 | 6.35E-02 |
| ENSG00000273448 | AC006480.2 | 0.43 | 9.24E-03 | 6.36E-02 |
| ENSG00000183484 | GPR132 | -0.43 | 9.24E-03 | 6.36E-02 |
| ENSG00000251417 | AC145285.2 | 0.43 | 9.25E-03 | 6.37E-02 |
| ENSG00000011021 | CLCN6 | 0.43 | 9.26E-03 | 6.37E-02 |
| ENSG00000100220 | RTCB | -0.43 | 9.26E-03 | 6.37E-02 |
| ENSG00000109920 | FNBP4 | 0.43 | 9.26E-03 | 6.37E-02 |
| ENSG00000115884 | SDC1 | 0.43 | 9.26E-03 | 6.37E-02 |
| ENSG00000131143 | COX4I1 | -0.43 | 9.31E-03 | 6.39E-02 |
| ENSG00000134056 | MRPS36 | -0.43 | 9.31E-03 | 6.39E-02 |
| ENSG00000144848 | ATG3 | -0.43 | 9.31E-03 | 6.39E-02 |
| ENSG00000167978 | SRRM2 | 0.43 | 9.31E-03 | 6.39E-02 |
| ENSG00000168385 | SEPT2 | -0.43 | 9.31E-03 | 6.39E-02 |
| ENSG00000170088 | TMEM192 | 0.43 | 9.31E-03 | 6.39E-02 |

| Ensembl ID | Gene Name | ρ | Pval | Padj |
|---|---|---|---|---|
| ENSG00000166927 | MS4A7 | -0.43 | 9.34E-03 | 6.41E-02 |
| ENSG00000226377 | AC084809.1 | 0.43 | 9.35E-03 | 6.41E-02 |
| ENSG00000125845 | BMP2 | -0.43 | 9.36E-03 | 6.41E-02 |
| ENSG00000139324 | TMTC3 | -0.43 | 9.36E-03 | 6.41E-02 |
| ENSG00000149196 | HIKESHI | -0.43 | 9.36E-03 | 6.41E-02 |
| ENSG00000167394 | ZNF668 | -0.43 | 9.36E-03 | 6.41E-02 |
| ENSG00000204525 | HLA-C | 0.43 | 9.36E-03 | 6.41E-02 |
| ENSG00000263873 | AP003396.5 | -0.43 | 9.37E-03 | 6.41E-02 |
| ENSG00000188290 | HES4 | -0.43 | 9.40E-03 | 6.42E-02 |
| ENSG00000073331 | ALPK1 | 0.43 | 9.41E-03 | 6.42E-02 |
| ENSG00000086065 | CHMP5 | -0.43 | 9.41E-03 | 6.42E-02 |
| ENSG00000100266 | PACSIN2 | 0.43 | 9.41E-03 | 6.42E-02 |
| ENSG00000100916 | BRMS1L | -0.43 | 9.41E-03 | 6.42E-02 |
| ENSG00000178537 | SLC25A20 | -0.43 | 9.41E-03 | 6.42E-02 |
| ENSG00000230590 | FTX | 0.43 | 9.41E-03 | 6.42E-02 |
| ENSG00000234036 | TXNP6 | -0.43 | 9.41E-03 | 6.42E-02 |
| ENSG00000264278 | ZNF236-DT | 0.43 | 9.41E-03 | 6.42E-02 |
| ENSG00000251314 | AC104123.1 | 0.43 | 9.42E-03 | 6.43E-02 |
| ENSG00000080166 | DCT | 0.43 | 9.47E-03 | 6.44E-02 |
| ENSG00000100324 | TAB1 | 0.43 | 9.47E-03 | 6.44E-02 |
| ENSG00000107249 | GLIS3 | -0.43 | 9.47E-03 | 6.44E-02 |
| ENSG00000125445 | MRPS7 | -0.43 | 9.47E-03 | 6.44E-02 |
| ENSG00000154473 | BUB3 | -0.43 | 9.47E-03 | 6.44E-02 |
| ENSG00000172301 | COPRS | -0.43 | 9.47E-03 | 6.44E-02 |
| ENSG00000173258 | ZNF483 | 0.43 | 9.47E-03 | 6.44E-02 |
| ENSG00000175691 | ZNF77 | -0.43 | 9.47E-03 | 6.44E-02 |
| ENSG00000227908 | AC016596.1 | 0.43 | 9.48E-03 | 6.45E-02 |
| ENSG00000272568 | AC005162.3 | 0.43 | 9.51E-03 | 6.45E-02 |
| ENSG00000255026 | AC136475.3 | 0.43 | 9.51E-03 | 6.45E-02 |
| ENSG00000099953 | MMP11 | -0.43 | 9.52E-03 | 6.45E-02 |
| ENSG00000132716 | DCAF8 | 0.43 | 9.52E-03 | 6.45E-02 |
| ENSG00000140280 | LYSMD2 | -0.43 | 9.52E-03 | 6.45E-02 |
| ENSG00000150977 | RILPL2 | 0.43 | 9.52E-03 | 6.45E-02 |
| ENSG00000166452 | AKIP1 | -0.43 | 9.52E-03 | 6.45E-02 |
| ENSG00000188215 | DCUN1D3 | -0.43 | 9.52E-03 | 6.45E-02 |
| ENSG00000231721 | LINC-PINT | 0.43 | 9.52E-03 | 6.45E-02 |
| ENSG00000002549 | LAP3 | -0.43 | 9.57E-03 | 6.47E-02 |
| ENSG00000109586 | GALNT7 | -0.43 | 9.57E-03 | 6.47E-02 |
| ENSG00000111832 | RWDD1 | -0.43 | 9.57E-03 | 6.47E-02 |
| ENSG00000145860 | RNF145 | -0.43 | 9.57E-03 | 6.47E-02 |
| ENSG00000158023 | WDR66 | 0.43 | 9.57E-03 | 6.47E-02 |
| ENSG00000241468 | ATP5MF | -0.43 | 9.57E-03 | 6.47E-02 |
| ENSG00000101104 | PABPC1L | 0.43 | 9.62E-03 | 6.49E-02 |
| ENSG00000115935 | WIPF1 | -0.43 | 9.62E-03 | 6.49E-02 |
| ENSG00000143643 | TTC13 | -0.43 | 9.62E-03 | 6.49E-02 |

| Ensembl ID | Gene Name | ρ | Pval | Padj |
|---|---|---|---|---|
| ENSG00000158882 | TOMM40L | -0.43 | 9.62E-03 | 6.49E-02 |
| ENSG00000170891 | CYTL1 | 0.43 | 9.62E-03 | 6.49E-02 |
| ENSG00000188185 | LINC00265 | 0.43 | 9.62E-03 | 6.49E-02 |
| ENSG00000234465 | PINLYP | -0.43 | 9.62E-03 | 6.49E-02 |
| ENSG00000021488 | SLC7A9 | 0.43 | 9.67E-03 | 6.51E-02 |
| ENSG00000105852 | PON3 | 0.43 | 9.67E-03 | 6.51E-02 |
| ENSG00000173889 | PHC3 | 0.43 | 9.67E-03 | 6.51E-02 |
| ENSG00000182400 | TRAPPC6B | -0.43 | 9.67E-03 | 6.51E-02 |
| ENSG00000196313 | POM121 | 0.43 | 9.67E-03 | 6.51E-02 |
| ENSG00000215845 | TSTD1 | -0.43 | 9.67E-03 | 6.51E-02 |
| ENSG00000267305 | AL592148.2 | 0.43 | 9.69E-03 | 6.53E-02 |
| ENSG00000186871 | ERCC6L | -0.43 | 9.72E-03 | 6.54E-02 |
| ENSG00000112992 | NNT | -0.43 | 9.73E-03 | 6.54E-02 |
| ENSG00000161813 | LARP4 | -0.43 | 9.73E-03 | 6.54E-02 |
| ENSG00000172901 | LVRN | 0.43 | 9.73E-03 | 6.54E-02 |
| ENSG00000240694 | PNMA2 | -0.43 | 9.73E-03 | 6.54E-02 |
| ENSG00000115657 | ABCB6 | 0.43 | 9.75E-03 | 6.55E-02 |
| ENSG00000145740 | SLC30A5 | -0.43 | 9.78E-03 | 6.56E-02 |
| ENSG00000204316 | MRPL38 | 0.43 | 9.78E-03 | 6.56E-02 |
| ENSG00000276710 | CSPG4P10 | 0.43 | 9.78E-03 | 6.56E-02 |
| ENSG00000261889 | AC108134.2 | 0.43 | 9.80E-03 | 6.57E-02 |
| ENSG00000171596 | NMUR1 | 0.43 | 9.81E-03 | 6.58E-02 |
| ENSG00000269997 | AC068790.3 | 0.43 | 9.82E-03 | 6.58E-02 |
| ENSG00000254602 | AP000662.1 | 0.43 | 9.82E-03 | 6.58E-02 |
| ENSG00000199038 | MIR210 | 0.43 | 9.83E-03 | 6.58E-02 |
| ENSG00000236852 | BX322784.1 | -0.43 | 9.83E-03 | 6.58E-02 |
| ENSG00000112294 | ALDH5A1 | 0.43 | 9.83E-03 | 6.58E-02 |
| ENSG00000168439 | STIP1 | -0.43 | 9.83E-03 | 6.58E-02 |
| ENSG00000070761 | CFAP20 | -0.43 | 9.88E-03 | 6.61E-02 |
| ENSG00000100982 | PCIF1 | 0.43 | 9.88E-03 | 6.61E-02 |
| ENSG00000125810 | CD93 | -0.43 | 9.88E-03 | 6.61E-02 |
| ENSG00000144635 | DYNC1LI1 | -0.43 | 9.88E-03 | 6.61E-02 |
| ENSG00000166181 | API5 | -0.43 | 9.88E-03 | 6.61E-02 |
| ENSG00000171051 | FPR1 | -0.43 | 9.90E-03 | 6.61E-02 |
| ENSG00000259627 | AC079328.2 | -0.43 | 9.91E-03 | 6.62E-02 |
| ENSG00000147408 | CSGALNACT1 | 0.43 | 9.94E-03 | 6.63E-02 |
| ENSG00000005194 | CIAPIN1 | -0.43 | 9.99E-03 | 6.66E-02 |
| ENSG00000065970 | FOXJ2 | 0.43 | 9.99E-03 | 6.66E-02 |
| ENSG00000122912 | SLC25A16 | -0.43 | 9.99E-03 | 6.66E-02 |
| ENSG00000230630 | DNM3OS | -0.43 | 9.99E-03 | 6.66E-02 |
| ENSG00000233270 | SNRPEP4 | -0.43 | 9.99E-03 | 6.66E-02 |
| ENSG00000277369 | AC010654.1 | 0.43 | 9.99E-03 | 6.66E-02 |
| ENSG00000018236 | CNTN1 | -0.43 | 1.00E-02 | 6.68E-02 |
| ENSG00000226284 | ARPC3P1 | -0.43 | 1.00E-02 | 6.68E-02 |
| ENSG00000168395 | ING5 | 0.43 | 1.00E-02 | 6.69E-02 |

| Ensembl ID | Gene Name | ρ | Pval | Padj |
|---|---|---|---|---|
| ENSG00000236411 | NDUFAF4P3 | 0.43 | 1.01E-02 | 6.70E-02 |
| ENSG00000178631 | ACTG1P1 | 0.43 | 1.01E-02 | 6.70E-02 |
| ENSG00000101773 | RBBP8 | -0.43 | 1.01E-02 | 6.70E-02 |
| ENSG00000113070 | HBEGF | -0.43 | 1.01E-02 | 6.70E-02 |
| ENSG00000115241 | PPM1G | -0.43 | 1.01E-02 | 6.70E-02 |
| ENSG00000115761 | NOL10 | -0.43 | 1.01E-02 | 6.70E-02 |
| ENSG00000144597 | EAF1 | -0.43 | 1.01E-02 | 6.70E-02 |
| ENSG00000256667 | KLRA1P | 0.43 | 1.01E-02 | 6.70E-02 |
| ENSG00000277159 | AL139384.2 | 0.43 | 1.01E-02 | 6.70E-02 |
| ENSG00000168685 | IL7R | -0.43 | 1.01E-02 | 6.73E-02 |
| ENSG00000172824 | CES4A | 0.43 | 1.01E-02 | 6.73E-02 |
| ENSG00000100387 | RBX1 | -0.43 | 1.02E-02 | 6.73E-02 |
| ENSG00000125611 | CHCHD5 | -0.43 | 1.02E-02 | 6.73E-02 |
| ENSG00000273117 | AC144652.1 | 0.43 | 1.02E-02 | 6.73E-02 |
| ENSG00000270228 | AC079880.1 | 0.43 | 1.02E-02 | 6.74E-02 |
| ENSG00000266677 | AC087164.1 | 0.43 | 1.02E-02 | 6.74E-02 |
| ENSG00000122420 | PTGFR | -0.43 | 1.02E-02 | 6.74E-02 |
| ENSG00000145741 | BTF3 | -0.43 | 1.02E-02 | 6.74E-02 |
| ENSG00000162341 | TPCN2 | 0.43 | 1.02E-02 | 6.74E-02 |
| ENSG00000166839 | ANKDD1A | 0.43 | 1.02E-02 | 6.74E-02 |
| ENSG00000172409 | CLP1 | -0.43 | 1.02E-02 | 6.74E-02 |
| ENSG00000180211 | FO393411.1 | -0.43 | 1.02E-02 | 6.74E-02 |
| ENSG00000227036 | LINC00511 | -0.43 | 1.02E-02 | 6.74E-02 |
| ENSG00000010671 | BTK | -0.43 | 1.02E-02 | 6.76E-02 |
| ENSG00000100036 | SLC35E4 | -0.43 | 1.03E-02 | 6.76E-02 |
| ENSG00000103642 | LACTB | -0.43 | 1.03E-02 | 6.76E-02 |
| ENSG00000104067 | TJP1 | -0.43 | 1.03E-02 | 6.76E-02 |
| ENSG00000163818 | LZTFL1 | -0.43 | 1.03E-02 | 6.76E-02 |
| ENSG00000234337 | AC026462.1 | -0.43 | 1.03E-02 | 6.76E-02 |
| ENSG00000249740 | OSMR-AS1 | 0.43 | 1.03E-02 | 6.76E-02 |
| ENSG00000257038 | AP002761.3 | 0.43 | 1.03E-02 | 6.76E-02 |
| ENSG00000101019 | UQCC1 | 0.43 | 1.03E-02 | 6.79E-02 |
| ENSG00000106245 | BUD31 | -0.43 | 1.03E-02 | 6.79E-02 |
| ENSG00000118402 | ELOVL4 | -0.43 | 1.03E-02 | 6.79E-02 |
| ENSG00000126705 | AHDC1 | 0.43 | 1.03E-02 | 6.79E-02 |
| ENSG00000260496 | AC009041.1 | 0.43 | 1.03E-02 | 6.79E-02 |
| ENSG00000279981 | AC018445.4 | 0.43 | 1.03E-02 | 6.79E-02 |
| ENSG00000267078 | AC015802.1 | 0.43 | 1.04E-02 | 6.80E-02 |
| ENSG00000258645 | HSPE1P2 | -0.43 | 1.04E-02 | 6.80E-02 |
| ENSG00000100519 | PSMC6 | -0.43 | 1.04E-02 | 6.80E-02 |
| ENSG00000112769 | LAMA4 | -0.43 | 1.04E-02 | 6.80E-02 |
| ENSG00000134852 | CLOCK | -0.43 | 1.04E-02 | 6.80E-02 |
| ENSG00000146966 | DENND2A | -0.43 | 1.04E-02 | 6.80E-02 |
| ENSG00000182253 | SYNM | 0.43 | 1.04E-02 | 6.80E-02 |
| ENSG00000269514 | AC024257.3 | 0.43 | 1.04E-02 | 6.80E-02 |

| Ensembl ID | Gene Name | ρ | Pval | Padj |
|---|---|---|---|---|
| ENSG00000163995 | ABLIM2 | -0.43 | 1.04E-02 | 6.82E-02 |
| ENSG00000278464 | AC068506.1 | 0.43 | 1.04E-02 | 6.82E-02 |
| ENSG00000005108 | THSD7A | 0.43 | 1.04E-02 | 6.83E-02 |
| ENSG00000238086 | PPP1R26P1 | 0.43 | 1.04E-02 | 6.83E-02 |
| ENSG00000272754 | AL133245.1 | 0.43 | 1.04E-02 | 6.83E-02 |
| ENSG00000105173 | CCNE1 | -0.43 | 1.05E-02 | 6.85E-02 |
| ENSG00000110324 | IL10RA | -0.43 | 1.05E-02 | 6.85E-02 |
| ENSG00000137876 | RSL24D1 | -0.43 | 1.05E-02 | 6.85E-02 |
| ENSG00000198270 | TMEM116 | 0.43 | 1.05E-02 | 6.85E-02 |
| ENSG00000241058 | NSUN6 | 0.43 | 1.05E-02 | 6.85E-02 |
| ENSG00000270878 | AL136038.4 | 0.43 | 1.05E-02 | 6.88E-02 |
| ENSG00000116661 | FBXO2 | 0.43 | 1.05E-02 | 6.88E-02 |
| ENSG00000127337 | YEATS4 | -0.43 | 1.05E-02 | 6.88E-02 |
| ENSG00000132386 | SERPINF1 | -0.43 | 1.05E-02 | 6.88E-02 |
| ENSG00000132613 | MTSS1L | 0.43 | 1.05E-02 | 6.88E-02 |
| ENSG00000143653 | SCCPDH | -0.43 | 1.05E-02 | 6.88E-02 |
| ENSG00000158428 | CATIP | 0.43 | 1.06E-02 | 6.90E-02 |
| ENSG00000104064 | GABPB1 | 0.43 | 1.06E-02 | 6.90E-02 |
| ENSG00000164221 | CCDC112 | -0.43 | 1.06E-02 | 6.90E-02 |
| ENSG00000177576 | C18orf32 | 0.43 | 1.06E-02 | 6.90E-02 |
| ENSG00000184216 | IRAK1 | -0.43 | 1.06E-02 | 6.90E-02 |
| ENSG00000229214 | LINC00242 | 0.43 | 1.06E-02 | 6.90E-02 |
| ENSG00000260808 | AP003096.1 | 0.43 | 1.06E-02 | 6.91E-02 |
| ENSG00000073712 | FERMT2 | -0.43 | 1.07E-02 | 6.92E-02 |
| ENSG00000079432 | CIC | 0.43 | 1.07E-02 | 6.92E-02 |
| ENSG00000136379 | ABHD17C | -0.43 | 1.07E-02 | 6.92E-02 |
| ENSG00000140374 | ETFA | -0.43 | 1.07E-02 | 6.92E-02 |
| ENSG00000164916 | FOXK1 | 0.43 | 1.07E-02 | 6.92E-02 |
| ENSG00000169288 | MRPL1 | -0.43 | 1.07E-02 | 6.92E-02 |
| ENSG00000197766 | CFD | -0.43 | 1.07E-02 | 6.92E-02 |
| ENSG00000231652 | AL590428.1 | 0.43 | 1.07E-02 | 6.92E-02 |
| ENSG00000108798 | ABI3 | -0.43 | 1.07E-02 | 6.94E-02 |
| ENSG00000104915 | STX10 | -0.43 | 1.07E-02 | 6.94E-02 |
| ENSG00000125746 | EML2 | 0.43 | 1.07E-02 | 6.94E-02 |
| ENSG00000131471 | AOC3 | -0.43 | 1.07E-02 | 6.94E-02 |
| ENSG00000135624 | CCT7 | -0.43 | 1.07E-02 | 6.94E-02 |
| ENSG00000143499 | SMYD2 | -0.43 | 1.07E-02 | 6.94E-02 |
| ENSG00000228594 | FNDC10 | -0.43 | 1.07E-02 | 6.94E-02 |
| ENSG00000188452 | CERKL | -0.43 | 1.08E-02 | 6.96E-02 |
| ENSG00000110876 | SELPLG | -0.43 | 1.08E-02 | 6.97E-02 |
| ENSG00000261371 | PECAM1 | -0.43 | 1.08E-02 | 6.97E-02 |
| ENSG00000234093 | RPS15AP11 | -0.43 | 1.08E-02 | 6.98E-02 |
| ENSG00000099377 | HSD3B7 | -0.43 | 1.08E-02 | 6.98E-02 |
| ENSG00000100304 | TTLL12 | -0.43 | 1.08E-02 | 6.98E-02 |
| ENSG00000103051 | COG4 | -0.43 | 1.08E-02 | 6.98E-02 |

| Ensembl ID | Gene Name | ρ | Pval | Padj |
|---|---|---|---|---|
| ENSG00000124688 | MAD2L1BP | -0.43 | 1.08E-02 | 6.98E-02 |
| ENSG00000154263 | ABCA10 | 0.43 | 1.08E-02 | 6.98E-02 |
| ENSG00000158006 | PAFAH2 | 0.43 | 1.08E-02 | 6.98E-02 |
| ENSG00000164346 | NSA2 | -0.43 | 1.08E-02 | 6.98E-02 |
| ENSG00000173706 | HEG1 | -0.43 | 1.08E-02 | 6.98E-02 |
| ENSG00000253485 | PCDHGA5 | 0.43 | 1.08E-02 | 6.98E-02 |
| ENSG00000086598 | TMED2 | -0.43 | 1.09E-02 | 7.01E-02 |
| ENSG00000100104 | SRRD | -0.43 | 1.09E-02 | 7.01E-02 |
| ENSG00000149575 | SCN2B | 0.43 | 1.09E-02 | 7.01E-02 |
| ENSG00000263766 | AC025682.1 | 0.43 | 1.09E-02 | 7.02E-02 |
| ENSG00000260751 | AC008870.2 | 0.43 | 1.09E-02 | 7.02E-02 |
| ENSG00000280152 | AC009078.3 | 0.43 | 1.09E-02 | 7.02E-02 |
| ENSG00000272711 | AC019069.1 | -0.43 | 1.09E-02 | 7.03E-02 |
| ENSG00000008838 | MED24 | 0.42 | 1.09E-02 | 7.04E-02 |
| ENSG00000136861 | CDK5RAP2 | 0.42 | 1.09E-02 | 7.04E-02 |
| ENSG00000275302 | CCL4 | -0.42 | 1.10E-02 | 7.05E-02 |
| ENSG00000074657 | ZNF532 | 0.42 | 1.10E-02 | 7.07E-02 |
| ENSG00000120437 | ACAT2 | -0.42 | 1.10E-02 | 7.07E-02 |
| ENSG00000204311 | PJVK | 0.42 | 1.10E-02 | 7.07E-02 |
| ENSG00000105967 | TFEC | -0.42 | 1.10E-02 | 7.08E-02 |
| ENSG00000232098 | AC012313.1 | 0.42 | 1.10E-02 | 7.08E-02 |
| ENSG00000116918 | TSNAX | -0.42 | 1.11E-02 | 7.09E-02 |
| ENSG00000155629 | PIK3AP1 | -0.42 | 1.11E-02 | 7.09E-02 |
| ENSG00000187713 | TMEM203 | -0.42 | 1.11E-02 | 7.09E-02 |
| ENSG00000228065 | LINC01515 | 0.42 | 1.11E-02 | 7.09E-02 |
| ENSG00000137752 | CASP1 | -0.42 | 1.11E-02 | 7.10E-02 |
| ENSG00000225177 | AL590617.2 | -0.42 | 1.11E-02 | 7.10E-02 |
| ENSG00000067334 | DNTTIP2 | -0.42 | 1.11E-02 | 7.11E-02 |
| ENSG00000083844 | ZNF264 | 0.42 | 1.11E-02 | 7.11E-02 |
| ENSG00000101463 | SYNDIG1 | -0.42 | 1.11E-02 | 7.11E-02 |
| ENSG00000134453 | RBM17 | -0.42 | 1.11E-02 | 7.11E-02 |
| ENSG00000137692 | DCUN1D5 | -0.42 | 1.11E-02 | 7.11E-02 |
| ENSG00000170448 | NFXL1 | 0.42 | 1.11E-02 | 7.11E-02 |
| ENSG00000271913 | AL035530.2 | 0.42 | 1.11E-02 | 7.11E-02 |
| ENSG00000162711 | NLRP3 | -0.42 | 1.11E-02 | 7.13E-02 |
| ENSG00000224543 | SNRPGP15 | -0.42 | 1.12E-02 | 7.13E-02 |
| ENSG00000122786 | CALD1 | -0.42 | 1.12E-02 | 7.13E-02 |
| ENSG00000154258 | ABCA9 | 0.42 | 1.12E-02 | 7.13E-02 |
| ENSG00000167528 | ZNF641 | -0.42 | 1.12E-02 | 7.13E-02 |
| ENSG00000276672 | AL161891.1 | 0.42 | 1.12E-02 | 7.14E-02 |
| ENSG00000230337 | EXOSC10-AS1 | 0.42 | 1.12E-02 | 7.14E-02 |
| ENSG00000132196 | HSD17B7 | 0.42 | 1.12E-02 | 7.16E-02 |
| ENSG00000139146 | SINHCAF | -0.42 | 1.12E-02 | 7.16E-02 |
| ENSG00000164048 | ZNF589 | 0.42 | 1.12E-02 | 7.16E-02 |
| ENSG00000197461 | PDGFA | -0.42 | 1.12E-02 | 7.16E-02 |

| Ensembl ID | Gene Name | ρ | Pval | Padj |
|---|---|---|---|---|
| ENSG00000186818 | LILRB4 | -0.42 | 1.12E-02 | 7.16E-02 |
| ENSG00000165490 | DDIAS | -0.42 | 1.13E-02 | 7.16E-02 |
| ENSG00000122705 | CLTA | -0.42 | 1.13E-02 | 7.18E-02 |
| ENSG00000135316 | SYNCRIP | -0.42 | 1.13E-02 | 7.18E-02 |
| ENSG00000145293 | ENOPH1 | -0.42 | 1.13E-02 | 7.18E-02 |
| ENSG00000166398 | KIAA0355 | 0.42 | 1.13E-02 | 7.18E-02 |
| ENSG00000178971 | CTC1 | 0.42 | 1.13E-02 | 7.18E-02 |
| ENSG00000189164 | ZNF527 | 0.42 | 1.13E-02 | 7.18E-02 |
| ENSG00000251002 | AC244502.1 | -0.42 | 1.13E-02 | 7.19E-02 |
| ENSG00000275221 | HIST1H2AK | -0.42 | 1.13E-02 | 7.19E-02 |
| ENSG00000254230 | AC037459.4 | 0.42 | 1.14E-02 | 7.19E-02 |
| ENSG00000154822 | PLCL2 | -0.42 | 1.14E-02 | 7.19E-02 |
| ENSG00000160746 | ANO10 | -0.42 | 1.14E-02 | 7.19E-02 |
| ENSG00000173875 | ZNF791 | 0.42 | 1.14E-02 | 7.19E-02 |
| ENSG00000197451 | HNRNPAB | -0.42 | 1.14E-02 | 7.19E-02 |
| ENSG00000206573 | THUMPD3-AS1 | 0.42 | 1.14E-02 | 7.19E-02 |
| ENSG00000224905 | AP001347.1 | 0.42 | 1.14E-02 | 7.19E-02 |
| ENSG00000228782 | MRPL45P2 | 0.42 | 1.14E-02 | 7.19E-02 |
| ENSG00000283201 | AC092329.3 | 0.42 | 1.14E-02 | 7.22E-02 |
| ENSG00000282608 | ADORA3 | -0.42 | 1.14E-02 | 7.22E-02 |
| ENSG00000095539 | SEMA4G | 0.42 | 1.14E-02 | 7.22E-02 |
| ENSG00000137996 | RTCA | -0.42 | 1.14E-02 | 7.22E-02 |
| ENSG00000139626 | ITGB7 | 0.42 | 1.14E-02 | 7.22E-02 |
| ENSG00000183735 | TBK1 | -0.42 | 1.14E-02 | 7.22E-02 |
| ENSG00000273084 | AC092171.5 | 0.42 | 1.14E-02 | 7.22E-02 |
| ENSG00000236204 | LINC01376 | 0.42 | 1.15E-02 | 7.24E-02 |
| ENSG00000100483 | VCPKMT | -0.42 | 1.15E-02 | 7.24E-02 |
| ENSG00000143578 | CREB3L4 | 0.42 | 1.15E-02 | 7.24E-02 |
| ENSG00000156299 | TIAM1 | -0.42 | 1.15E-02 | 7.24E-02 |
| ENSG00000234869 | AL021392.1 | 0.42 | 1.15E-02 | 7.24E-02 |
| ENSG00000247746 | USP51 | 0.42 | 1.15E-02 | 7.24E-02 |
| ENSG00000104774 | MAN2B1 | -0.42 | 1.15E-02 | 7.27E-02 |
| ENSG00000170275 | CRTAP | 0.42 | 1.15E-02 | 7.27E-02 |
| ENSG00000204922 | UQCC3 | -0.42 | 1.15E-02 | 7.27E-02 |
| ENSG00000235706 | DICER1-AS1 | 0.42 | 1.15E-02 | 7.27E-02 |
| ENSG00000161328 | LRRC56 | 0.42 | 1.16E-02 | 7.30E-02 |
| ENSG00000169896 | ITGAM | -0.42 | 1.16E-02 | 7.30E-02 |
| ENSG00000188277 | C15orf62 | 0.42 | 1.16E-02 | 7.30E-02 |
| ENSG00000154027 | AK5 | -0.42 | 1.16E-02 | 7.30E-02 |
| ENSG00000279467 | AP000350.7 | 0.42 | 1.16E-02 | 7.31E-02 |
| ENSG00000100558 | PLEK2 | -0.42 | 1.17E-02 | 7.31E-02 |
| ENSG00000069702 | TGFBR3 | 0.42 | 1.17E-02 | 7.31E-02 |
| ENSG00000109220 | CHIC2 | -0.42 | 1.17E-02 | 7.31E-02 |
| ENSG00000116138 | DNAJC16 | -0.42 | 1.17E-02 | 7.31E-02 |
| ENSG00000135617 | PRADC1 | -0.42 | 1.17E-02 | 7.31E-02 |

| Ensembl ID | Gene Name | ρ | Pval | Padj |
|---|---|---|---|---|
| ENSG00000180818 | HOXC10 | 0.42 | 1.17E-02 | 7.31E-02 |
| ENSG00000185803 | SLC52A2 | -0.42 | 1.17E-02 | 7.31E-02 |
| ENSG00000196372 | ASB13 | 0.42 | 1.17E-02 | 7.31E-02 |
| ENSG00000217130 | AL139100.1 | -0.42 | 1.17E-02 | 7.31E-02 |
| ENSG00000245498 | AP000866.1 | 0.42 | 1.17E-02 | 7.31E-02 |
| ENSG00000240591 | AL096701.3 | 0.42 | 1.17E-02 | 7.31E-02 |
| ENSG00000112837 | TBX18 | 0.42 | 1.17E-02 | 7.34E-02 |
| ENSG00000278969 | AC026310.3 | 0.42 | 1.17E-02 | 7.34E-02 |
| ENSG00000130513 | GDF15 | -0.42 | 1.18E-02 | 7.40E-02 |
| ENSG00000067445 | TRO | 0.42 | 1.18E-02 | 7.40E-02 |
| ENSG00000067955 | CBFB | -0.42 | 1.18E-02 | 7.40E-02 |
| ENSG00000080947 | CROCCP3 | 0.42 | 1.18E-02 | 7.40E-02 |
| ENSG00000085872 | CHERP | 0.42 | 1.18E-02 | 7.40E-02 |
| ENSG00000111785 | RIC8B | -0.42 | 1.18E-02 | 7.40E-02 |
| ENSG00000132256 | TRIM5 | -0.42 | 1.18E-02 | 7.40E-02 |
| ENSG00000156469 | MTERF3 | -0.42 | 1.18E-02 | 7.40E-02 |
| ENSG00000164061 | BSN | 0.42 | 1.18E-02 | 7.40E-02 |
| ENSG00000173376 | NDNF | 0.42 | 1.18E-02 | 7.40E-02 |
| ENSG00000163106 | HPGDS | -0.42 | 1.19E-02 | 7.40E-02 |
| ENSG00000237301 | AL121992.1 | 0.42 | 1.19E-02 | 7.41E-02 |
| ENSG00000131100 | ATP6V1E1 | -0.42 | 1.19E-02 | 7.42E-02 |
| ENSG00000158458 | NRG2 | 0.42 | 1.19E-02 | 7.42E-02 |
| ENSG00000163125 | RPRD2 | 0.42 | 1.19E-02 | 7.42E-02 |
| ENSG00000171311 | EXOSC1 | -0.42 | 1.19E-02 | 7.42E-02 |
| ENSG00000196083 | IL1RAP | -0.42 | 1.19E-02 | 7.42E-02 |
| ENSG00000166359 | WDR88 | 0.42 | 1.19E-02 | 7.43E-02 |
| ENSG00000225099 | ATP6V1E1P1 | -0.42 | 1.19E-02 | 7.44E-02 |
| ENSG00000186301 | MST1P2 | 0.42 | 1.20E-02 | 7.45E-02 |
| ENSG00000128578 | STRIP2 | -0.42 | 1.20E-02 | 7.45E-02 |
| ENSG00000197608 | ZNF841 | 0.42 | 1.20E-02 | 7.45E-02 |
| ENSG00000215021 | PHB2 | -0.42 | 1.20E-02 | 7.45E-02 |
| ENSG00000163814 | CDCP1 | -0.42 | 1.20E-02 | 7.46E-02 |
| ENSG00000142233 | NTN5 | 0.42 | 1.20E-02 | 7.47E-02 |
| ENSG00000109917 | ZPR1 | -0.42 | 1.20E-02 | 7.47E-02 |
| ENSG00000122824 | NUDT10 | -0.42 | 1.20E-02 | 7.47E-02 |
| ENSG00000139718 | SETD1B | 0.42 | 1.20E-02 | 7.47E-02 |
| ENSG00000143815 | LBR | -0.42 | 1.20E-02 | 7.47E-02 |
| ENSG00000258461 | AC012651.1 | 0.42 | 1.20E-02 | 7.48E-02 |
| ENSG00000126267 | COX6B1 | -0.42 | 1.21E-02 | 7.49E-02 |
| ENSG00000144712 | CAND2 | 0.42 | 1.21E-02 | 7.49E-02 |
| ENSG00000170949 | ZNF160 | 0.42 | 1.21E-02 | 7.49E-02 |
| ENSG00000180353 | HCLS1 | -0.42 | 1.21E-02 | 7.49E-02 |
| ENSG00000188783 | PRELP | 0.42 | 1.21E-02 | 7.49E-02 |
| ENSG00000240291 | AL450384.2 | 0.42 | 1.21E-02 | 7.49E-02 |
| ENSG00000120910 | PPP3CC | -0.42 | 1.22E-02 | 7.51E-02 |

| Ensembl ID | Gene Name | ρ | Pval | Padj |
|---|---|---|---|---|
| ENSG00000135297 | MTO1 | 0.42 | 1.22E-02 | 7.51E-02 |
| ENSG00000136521 | NDUFB5 | -0.42 | 1.22E-02 | 7.51E-02 |
| ENSG00000159199 | ATP5MC1 | -0.42 | 1.22E-02 | 7.51E-02 |
| ENSG00000163961 | RNF168 | 0.42 | 1.22E-02 | 7.51E-02 |
| ENSG00000171408 | PDE7B | 0.42 | 1.22E-02 | 7.51E-02 |
| ENSG00000204574 | ABCF1 | -0.42 | 1.22E-02 | 7.51E-02 |
| ENSG00000232119 | MCTS1 | -0.42 | 1.22E-02 | 7.51E-02 |
| ENSG00000249115 | HAUS5 | 0.42 | 1.22E-02 | 7.51E-02 |
| ENSG00000104388 | RAB2A | -0.42 | 1.22E-02 | 7.54E-02 |
| ENSG00000121481 | RNF2 | -0.42 | 1.22E-02 | 7.54E-02 |
| ENSG00000167077 | MEI1 | 0.42 | 1.22E-02 | 7.54E-02 |
| ENSG00000237506 | RPSAP15 | -0.42 | 1.22E-02 | 7.54E-02 |
| ENSG00000178860 | MSC | -0.42 | 1.22E-02 | 7.55E-02 |
| ENSG00000234432 | AC092171.3 | 0.42 | 1.22E-02 | 7.55E-02 |
| ENSG00000172543 | CTSW | -0.42 | 1.23E-02 | 7.56E-02 |
| ENSG00000137312 | FLOT1 | -0.42 | 1.23E-02 | 7.57E-02 |
| ENSG00000172428 | COPS9 | -0.42 | 1.23E-02 | 7.57E-02 |
| ENSG00000196739 | COL27A1 | 0.42 | 1.23E-02 | 7.57E-02 |
| ENSG00000228474 | OST4 | -0.42 | 1.23E-02 | 7.57E-02 |
| ENSG00000225849 | MKRN7P | 0.42 | 1.23E-02 | 7.57E-02 |
| ENSG00000233967 | AL359715.1 | 0.42 | 1.23E-02 | 7.58E-02 |
| ENSG00000132514 | CLEC10A | -0.42 | 1.23E-02 | 7.59E-02 |
| ENSG00000106477 | CEP41 | -0.42 | 1.24E-02 | 7.59E-02 |
| ENSG00000130827 | PLXNA3 | 0.42 | 1.24E-02 | 7.59E-02 |
| ENSG00000177189 | RPS6KA3 | -0.42 | 1.24E-02 | 7.59E-02 |
| ENSG00000196422 | PPP1R26 | 0.42 | 1.24E-02 | 7.59E-02 |
| ENSG00000129625 | REEP5 | -0.42 | 1.24E-02 | 7.62E-02 |
| ENSG00000196810 | CTBP1-DT | 0.42 | 1.24E-02 | 7.62E-02 |
| ENSG00000244041 | LINC01011 | 0.42 | 1.25E-02 | 7.64E-02 |
| ENSG00000254676 | AP000873.4 | 0.42 | 1.25E-02 | 7.64E-02 |
| ENSG00000227704 | AL354892.1 | 0.42 | 1.25E-02 | 7.64E-02 |
| ENSG00000025293 | PHF20 | 0.42 | 1.25E-02 | 7.64E-02 |
| ENSG00000118961 | LDAH | -0.42 | 1.25E-02 | 7.64E-02 |
| ENSG00000131848 | ZSCAN5A | 0.42 | 1.25E-02 | 7.64E-02 |
| ENSG00000156411 | ATP5MPL | -0.42 | 1.25E-02 | 7.64E-02 |
| ENSG00000203747 | FCGR3A | -0.42 | 1.25E-02 | 7.64E-02 |
| ENSG00000253683 | AC027309.2 | -0.42 | 1.25E-02 | 7.64E-02 |
| ENSG00000276043 | UHRF1 | -0.42 | 1.25E-02 | 7.64E-02 |
| ENSG00000114646 | CSPG5 | 0.42 | 1.25E-02 | 7.67E-02 |
| ENSG00000072134 | EPN2 | 0.42 | 1.25E-02 | 7.67E-02 |
| ENSG00000116473 | RAP1A | -0.42 | 1.25E-02 | 7.67E-02 |
| ENSG00000226084 | AC113935.1 | -0.42 | 1.25E-02 | 7.67E-02 |
| ENSG00000282164 | PEG13 | 0.42 | 1.25E-02 | 7.67E-02 |
| ENSG00000198075 | SULT1C4 | 0.42 | 1.26E-02 | 7.68E-02 |
| ENSG00000225969 | ABHD11-AS1 | -0.42 | 1.26E-02 | 7.68E-02 |

| Ensembl ID | Gene Name | ρ | Pval | Padj |
|---|---|---|---|---|
| ENSG00000184786 | TCTE3 | 0.42 | 1.26E-02 | 7.68E-02 |
| ENSG00000104164 | BLOC1S6 | -0.42 | 1.26E-02 | 7.68E-02 |
| ENSG00000104361 | NIPAL2 | -0.42 | 1.26E-02 | 7.68E-02 |
| ENSG00000129116 | PALLD | -0.42 | 1.26E-02 | 7.68E-02 |
| ENSG00000146007 | ZMAT2 | -0.42 | 1.26E-02 | 7.68E-02 |
| ENSG00000146833 | TRIM4 | 0.42 | 1.26E-02 | 7.68E-02 |
| ENSG00000163913 | IFT122 | 0.42 | 1.26E-02 | 7.68E-02 |
| ENSG00000176731 | C8orf59 | -0.42 | 1.26E-02 | 7.68E-02 |
| ENSG00000205746 | AC126755.1 | 0.42 | 1.26E-02 | 7.68E-02 |
| ENSG00000229521 | MYCBP2-AS2 | 0.42 | 1.26E-02 | 7.68E-02 |
| ENSG00000213860 | RPL21P75 | -0.42 | 1.26E-02 | 7.69E-02 |
| ENSG00000187105 | HEATR4 | 0.42 | 1.27E-02 | 7.70E-02 |
| ENSG00000118245 | TNP1 | -0.42 | 1.27E-02 | 7.70E-02 |
| ENSG00000205562 | AL049775.1 | -0.42 | 1.27E-02 | 7.70E-02 |
| ENSG00000168288 | MMADHC | -0.42 | 1.27E-02 | 7.70E-02 |
| ENSG00000204172 | AGAP9 | 0.42 | 1.27E-02 | 7.70E-02 |
| ENSG00000231389 | HLA-DPA1 | -0.42 | 1.27E-02 | 7.70E-02 |
| ENSG00000261243 | AC022165.1 | 0.42 | 1.27E-02 | 7.73E-02 |
| ENSG00000104142 | VPS18 | -0.42 | 1.27E-02 | 7.73E-02 |
| ENSG00000156875 | MFSD14A | -0.42 | 1.27E-02 | 7.73E-02 |
| ENSG00000167315 | ACAA2 | -0.42 | 1.27E-02 | 7.73E-02 |
| ENSG00000168701 | TMEM208 | -0.42 | 1.27E-02 | 7.73E-02 |
| ENSG00000214770 | AL161756.1 | 0.42 | 1.28E-02 | 7.75E-02 |
| ENSG00000256806 | C17orf100 | -0.42 | 1.28E-02 | 7.75E-02 |
| ENSG00000073756 | PTGS2 | -0.42 | 1.28E-02 | 7.75E-02 |
| ENSG00000105810 | CDK6 | -0.42 | 1.28E-02 | 7.75E-02 |
| ENSG00000169375 | SIN3A | 0.42 | 1.28E-02 | 7.75E-02 |
| ENSG00000184634 | MED12 | 0.42 | 1.28E-02 | 7.75E-02 |
| ENSG00000204381 | LAYN | 0.42 | 1.28E-02 | 7.75E-02 |
| ENSG00000247400 | DNAJC3-DT | 0.42 | 1.28E-02 | 7.75E-02 |
| ENSG00000152402 | GUCY1A2 | -0.42 | 1.28E-02 | 7.76E-02 |
| ENSG00000261604 | AC114947.2 | -0.42 | 1.29E-02 | 7.78E-02 |
| ENSG00000101337 | TM9SF4 | 0.42 | 1.29E-02 | 7.78E-02 |
| ENSG00000166762 | CATSPER2 | 0.42 | 1.29E-02 | 7.78E-02 |
| ENSG00000172830 | SSH3 | 0.42 | 1.29E-02 | 7.78E-02 |
| ENSG00000235244 | DANT2 | 0.42 | 1.29E-02 | 7.79E-02 |
| ENSG00000108106 | UBE2S | -0.42 | 1.29E-02 | 7.80E-02 |
| ENSG00000130429 | ARPC1B | -0.42 | 1.29E-02 | 7.80E-02 |
| ENSG00000143442 | POGZ | 0.42 | 1.29E-02 | 7.80E-02 |
| ENSG00000172113 | NME6 | -0.42 | 1.29E-02 | 7.80E-02 |
| ENSG00000180370 | PAK2 | -0.42 | 1.29E-02 | 7.80E-02 |
| ENSG00000185627 | PSMD13 | -0.42 | 1.29E-02 | 7.80E-02 |
| ENSG00000204272 | NBDY | -0.42 | 1.29E-02 | 7.80E-02 |
| ENSG00000249715 | FER1L5 | 0.42 | 1.30E-02 | 7.83E-02 |
| ENSG00000186185 | KIF18B | -0.42 | 1.30E-02 | 7.83E-02 |

| Ensembl ID | Gene Name | ρ | Pval | Padj |
|---|---|---|---|---|
| ENSG00000088035 | ALG6 | -0.42 | 1.30E-02 | 7.83E-02 |
| ENSG00000170624 | SGCD | -0.42 | 1.30E-02 | 7.83E-02 |
| ENSG00000181472 | ZBTB2 | -0.42 | 1.30E-02 | 7.83E-02 |
| ENSG00000185946 | RNPC3 | 0.42 | 1.30E-02 | 7.83E-02 |
| ENSG00000270820 | AC016727.1 | 0.42 | 1.30E-02 | 7.83E-02 |
| ENSG00000110848 | CD69 | -0.42 | 1.30E-02 | 7.83E-02 |
| ENSG00000011275 | RNF216 | 0.42 | 1.31E-02 | 7.85E-02 |
| ENSG00000101412 | E2F1 | -0.42 | 1.31E-02 | 7.85E-02 |
| ENSG00000102699 | PARP4 | -0.42 | 1.31E-02 | 7.85E-02 |
| ENSG00000135272 | MDFIC | -0.42 | 1.31E-02 | 7.85E-02 |
| ENSG00000140092 | FBLN5 | -0.42 | 1.31E-02 | 7.85E-02 |
| ENSG00000254469 | AP002495.2 | 0.42 | 1.31E-02 | 7.85E-02 |
| ENSG00000107937 | GTPBP4 | -0.42 | 1.31E-02 | 7.88E-02 |
| ENSG00000116774 | OLFML3 | -0.42 | 1.31E-02 | 7.88E-02 |
| ENSG00000136104 | RNASEH2B | -0.42 | 1.31E-02 | 7.88E-02 |
| ENSG00000138356 | AOX1 | 0.42 | 1.31E-02 | 7.88E-02 |
| ENSG00000146282 | RARS2 | -0.42 | 1.31E-02 | 7.88E-02 |
| ENSG00000184402 | SS18L1 | 0.42 | 1.31E-02 | 7.88E-02 |
| ENSG00000197724 | PHF2 | 0.42 | 1.31E-02 | 7.88E-02 |
| ENSG00000148541 | FAM13C | -0.41 | 1.32E-02 | 7.90E-02 |
| ENSG00000186510 | CLCNKA | 0.41 | 1.32E-02 | 7.90E-02 |
| ENSG00000256139 | AC007637.1 | 0.41 | 1.32E-02 | 7.90E-02 |
| ENSG00000183597 | TANGO2 | -0.41 | 1.32E-02 | 7.91E-02 |
| ENSG00000231340 | ACTG1P10 | 0.41 | 1.32E-02 | 7.92E-02 |
| ENSG00000253744 | AC025442.1 | 0.41 | 1.32E-02 | 7.92E-02 |
| ENSG00000150782 | IL18 | -0.41 | 1.33E-02 | 7.92E-02 |
| ENSG00000256940 | AP001453.2 | -0.41 | 1.33E-02 | 7.92E-02 |
| ENSG00000090013 | BLVRB | -0.41 | 1.33E-02 | 7.92E-02 |
| ENSG00000100478 | AP4S1 | 0.41 | 1.33E-02 | 7.92E-02 |
| ENSG00000130165 | ELOF1 | -0.41 | 1.33E-02 | 7.92E-02 |
| ENSG00000176973 | FAM89B | -0.41 | 1.33E-02 | 7.92E-02 |
| ENSG00000196262 | PPIA | -0.41 | 1.33E-02 | 7.92E-02 |
| ENSG00000213139 | CRYGS | -0.41 | 1.33E-02 | 7.92E-02 |
| ENSG00000255112 | CHMP1B | -0.41 | 1.33E-02 | 7.92E-02 |
| ENSG00000202515 | VTRNA1-3 | -0.41 | 1.33E-02 | 7.94E-02 |
| ENSG00000131037 | EPS8L1 | -0.41 | 1.33E-02 | 7.94E-02 |
| ENSG00000013275 | PSMC4 | -0.41 | 1.34E-02 | 7.94E-02 |
| ENSG00000105825 | TFPI2 | -0.41 | 1.34E-02 | 7.94E-02 |
| ENSG00000165502 | RPL36AL | -0.41 | 1.34E-02 | 7.94E-02 |
| ENSG00000196943 | NOP9 | 0.41 | 1.34E-02 | 7.94E-02 |
| ENSG00000249915 | PDCD6 | -0.41 | 1.34E-02 | 7.94E-02 |
| ENSG00000254332 | AF201337.1 | -0.41 | 1.34E-02 | 7.94E-02 |
| ENSG00000267871 | ZNF460-AS1 | 0.41 | 1.34E-02 | 7.98E-02 |
| ENSG00000104903 | LYL1 | -0.41 | 1.34E-02 | 7.98E-02 |
| ENSG00000175701 | MTLN | -0.41 | 1.34E-02 | 7.98E-02 |

| Ensembl ID | Gene Name | ρ | Pval | Padj |
|---|---|---|---|---|
| ENSG00000260173 | AC020779.2 | 0.41 | 1.34E-02 | 7.98E-02 |
| ENSG00000156467 | UQCRB | -0.41 | 1.35E-02 | 8.01E-02 |
| ENSG00000204899 | MZT1 | -0.41 | 1.35E-02 | 8.01E-02 |
| ENSG00000262468 | LINC01569 | 0.41 | 1.35E-02 | 8.01E-02 |
| ENSG00000181104 | F2R | -0.41 | 1.35E-02 | 8.02E-02 |
| ENSG00000228639 | ROCR | 0.41 | 1.36E-02 | 8.04E-02 |
| ENSG00000061936 | SFSWAP | 0.41 | 1.36E-02 | 8.04E-02 |
| ENSG00000075856 | SART3 | 0.41 | 1.36E-02 | 8.04E-02 |
| ENSG00000106268 | NUDT1 | -0.41 | 1.36E-02 | 8.04E-02 |
| ENSG00000186625 | KATNA1 | -0.41 | 1.36E-02 | 8.04E-02 |
| ENSG00000135709 | KIAA0513 | 0.41 | 1.36E-02 | 8.07E-02 |
| ENSG00000171302 | CANT1 | -0.41 | 1.36E-02 | 8.07E-02 |
| ENSG00000182979 | MTA1 | 0.41 | 1.36E-02 | 8.07E-02 |
| ENSG00000185950 | IRS2 | 0.41 | 1.36E-02 | 8.07E-02 |
| ENSG00000233954 | UQCRHL | -0.41 | 1.36E-02 | 8.07E-02 |
| ENSG00000250220 | AC053527.1 | 0.41 | 1.37E-02 | 8.08E-02 |
| ENSG00000111676 | ATN1 | 0.41 | 1.37E-02 | 8.10E-02 |
| ENSG00000154265 | ABCA5 | 0.41 | 1.37E-02 | 8.10E-02 |
| ENSG00000134007 | ADAM20 | 0.41 | 1.37E-02 | 8.11E-02 |
| ENSG00000251600 | AC107223.1 | 0.41 | 1.37E-02 | 8.11E-02 |
| ENSG00000004897 | CDC27 | -0.41 | 1.38E-02 | 8.13E-02 |
| ENSG00000128708 | HAT1 | -0.41 | 1.38E-02 | 8.13E-02 |
| ENSG00000162009 | SSTR5 | 0.41 | 1.38E-02 | 8.13E-02 |
| ENSG00000261126 | RBFADN | 0.41 | 1.38E-02 | 8.13E-02 |
| ENSG00000244619 | AC243547.1 | 0.41 | 1.38E-02 | 8.15E-02 |
| ENSG00000256482 | AC023796.2 | 0.41 | 1.38E-02 | 8.15E-02 |
| ENSG00000078140 | UBE2K | -0.41 | 1.38E-02 | 8.15E-02 |
| ENSG00000101161 | PRPF6 | 0.41 | 1.38E-02 | 8.15E-02 |
| ENSG00000139990 | DCAF5 | 0.41 | 1.38E-02 | 8.15E-02 |
| ENSG00000146122 | DAAM2 | 0.41 | 1.38E-02 | 8.15E-02 |
| ENSG00000166275 | BORCS7 | -0.41 | 1.38E-02 | 8.15E-02 |
| ENSG00000170310 | STX8 | -0.41 | 1.38E-02 | 8.15E-02 |
| ENSG00000225518 | LINC01703 | 0.41 | 1.38E-02 | 8.15E-02 |
| ENSG00000004961 | HCCS | -0.41 | 1.39E-02 | 8.17E-02 |
| ENSG00000109680 | TBC1D19 | -0.41 | 1.39E-02 | 8.17E-02 |
| ENSG00000116574 | RHOU | 0.41 | 1.39E-02 | 8.17E-02 |
| ENSG00000120889 | TNFRSF10B | -0.41 | 1.39E-02 | 8.17E-02 |
| ENSG00000129691 | ASH2L | -0.41 | 1.39E-02 | 8.17E-02 |
| ENSG00000138031 | ADCY3 | 0.41 | 1.39E-02 | 8.17E-02 |
| ENSG00000179698 | WDR97 | 0.41 | 1.39E-02 | 8.17E-02 |
| ENSG00000213906 | LTB4R2 | 0.41 | 1.39E-02 | 8.17E-02 |
| ENSG00000245651 | AC083805.1 | -0.41 | 1.39E-02 | 8.17E-02 |
| ENSG00000281195 | AC007878.1 | 0.41 | 1.39E-02 | 8.18E-02 |
| ENSG00000271806 | AL590822.2 | 0.41 | 1.40E-02 | 8.18E-02 |
| ENSG00000095303 | PTGS1 | 0.41 | 1.40E-02 | 8.18E-02 |

| Ensembl ID | Gene Name | ρ | Pval | Padj |
|---|---|---|---|---|
| ENSG00000108826 | MRPL27 | -0.41 | 1.40E-02 | 8.18E-02 |
| ENSG00000111843 | TMEM14C | -0.41 | 1.40E-02 | 8.18E-02 |
| ENSG00000123737 | EXOSC9 | -0.41 | 1.40E-02 | 8.18E-02 |
| ENSG00000125877 | ITPA | -0.41 | 1.40E-02 | 8.18E-02 |
| ENSG00000143013 | LMO4 | -0.41 | 1.40E-02 | 8.18E-02 |
| ENSG00000150540 | HNMT | -0.41 | 1.40E-02 | 8.18E-02 |
| ENSG00000160213 | CSTB | -0.41 | 1.40E-02 | 8.18E-02 |
| ENSG00000175895 | PLEKHF2 | -0.41 | 1.40E-02 | 8.18E-02 |
| ENSG00000187650 | VMAC | 0.41 | 1.40E-02 | 8.18E-02 |
| ENSG00000196890 | HIST3H2BB | -0.41 | 1.40E-02 | 8.19E-02 |
| ENSG00000276449 | AC004076.2 | 0.41 | 1.40E-02 | 8.19E-02 |
| ENSG00000088827 | SIGLEC1 | -0.41 | 1.40E-02 | 8.20E-02 |
| ENSG00000239791 | AC002310.2 | 0.41 | 1.40E-02 | 8.21E-02 |
| ENSG00000214535 | RPS15AP1 | -0.41 | 1.40E-02 | 8.21E-02 |
| ENSG00000166352 | C11orf74 | -0.41 | 1.41E-02 | 8.21E-02 |
| ENSG00000283491 | AC017104.5 | 0.41 | 1.41E-02 | 8.21E-02 |
| ENSG00000282961 | PRNCR1 | 0.41 | 1.41E-02 | 8.21E-02 |
| ENSG00000117154 | IGSF21 | -0.41 | 1.41E-02 | 8.22E-02 |
| ENSG00000250240 | AC008840.1 | 0.41 | 1.41E-02 | 8.23E-02 |
| ENSG00000103319 | EEF2K | 0.41 | 1.41E-02 | 8.23E-02 |
| ENSG00000117448 | AKR1A1 | -0.41 | 1.41E-02 | 8.23E-02 |
| ENSG00000135387 | CAPRIN1 | -0.41 | 1.41E-02 | 8.23E-02 |
| ENSG00000167642 | SPINT2 | 0.41 | 1.41E-02 | 8.23E-02 |
| ENSG00000274211 | SOCS7 | 0.41 | 1.41E-02 | 8.23E-02 |
| ENSG00000227751 | RCC2-AS1 | -0.41 | 1.41E-02 | 8.24E-02 |
| ENSG00000186766 | FOXI2 | 0.41 | 1.42E-02 | 8.24E-02 |
| ENSG00000274312 | AC015722.2 | -0.41 | 1.42E-02 | 8.24E-02 |
| ENSG00000128262 | POM121L9P | -0.41 | 1.42E-02 | 8.24E-02 |
| ENSG00000088888 | MAVS | 0.41 | 1.42E-02 | 8.25E-02 |
| ENSG00000137501 | SYTL2 | -0.41 | 1.42E-02 | 8.25E-02 |
| ENSG00000146414 | SHPRH | 0.41 | 1.42E-02 | 8.25E-02 |
| ENSG00000153064 | BANK1 | 0.41 | 1.42E-02 | 8.25E-02 |
| ENSG00000163815 | CLEC3B | -0.41 | 1.42E-02 | 8.25E-02 |
| ENSG00000164989 | CCDC171 | 0.41 | 1.42E-02 | 8.25E-02 |
| ENSG00000227218 | AL157935.1 | -0.41 | 1.42E-02 | 8.25E-02 |
| ENSG00000229044 | AL451070.1 | 0.41 | 1.42E-02 | 8.26E-02 |
| ENSG00000250900 | AC008443.6 | 0.41 | 1.42E-02 | 8.26E-02 |
| ENSG00000128285 | MCHR1 | -0.41 | 1.43E-02 | 8.27E-02 |
| ENSG00000278873 | AC005495.1 | 0.41 | 1.43E-02 | 8.27E-02 |
| ENSG00000005339 | CREBBP | 0.41 | 1.43E-02 | 8.27E-02 |
| ENSG00000163132 | MSX1 | -0.41 | 1.43E-02 | 8.27E-02 |
| ENSG00000179967 | PPP1R14BP3 | -0.41 | 1.43E-02 | 8.27E-02 |
| ENSG00000206341 | HLA-H | 0.41 | 1.43E-02 | 8.27E-02 |
| ENSG00000278962 | AC092645.1 | 0.41 | 1.43E-02 | 8.27E-02 |
| ENSG00000154133 | ROBO4 | -0.41 | 1.43E-02 | 8.28E-02 |

| Ensembl ID | Gene Name | ρ | Pval | Padj |
|---|---|---|---|---|
| ENSG00000270540 | AC093915.1 | 0.41 | 1.43E-02 | 8.28E-02 |
| ENSG00000283341 | AC068205.2 | 0.41 | 1.43E-02 | 8.28E-02 |
| ENSG00000278576 | AL162171.4 | 0.41 | 1.43E-02 | 8.29E-02 |
| ENSG00000138131 | LOXL4 | -0.41 | 1.43E-02 | 8.29E-02 |
| ENSG00000147684 | NDUFB9 | -0.41 | 1.43E-02 | 8.29E-02 |
| ENSG00000159403 | C1R | -0.41 | 1.43E-02 | 8.29E-02 |
| ENSG00000181481 | RNF135 | -0.41 | 1.43E-02 | 8.29E-02 |
| ENSG00000239672 | NME1 | -0.41 | 1.43E-02 | 8.29E-02 |
| ENSG00000057704 | TMCC3 | -0.41 | 1.44E-02 | 8.30E-02 |
| ENSG00000134516 | DOCK2 | -0.41 | 1.44E-02 | 8.31E-02 |
| ENSG00000091972 | CD200 | -0.41 | 1.44E-02 | 8.31E-02 |
| ENSG00000120798 | NR2C1 | 0.41 | 1.44E-02 | 8.31E-02 |
| ENSG00000154153 | RETREG1 | -0.41 | 1.44E-02 | 8.31E-02 |
| ENSG00000157625 | TAB3 | 0.41 | 1.44E-02 | 8.31E-02 |
| ENSG00000225573 | RPL35P5 | -0.41 | 1.44E-02 | 8.31E-02 |
| ENSG00000232362 | ATP5MGP2 | -0.41 | 1.45E-02 | 8.33E-02 |
| ENSG00000108094 | CUL2 | -0.41 | 1.45E-02 | 8.33E-02 |
| ENSG00000135749 | PCNX2 | 0.41 | 1.45E-02 | 8.33E-02 |
| ENSG00000144909 | OSBPL11 | -0.41 | 1.45E-02 | 8.33E-02 |
| ENSG00000152455 | SUV39H2 | -0.41 | 1.45E-02 | 8.33E-02 |
| ENSG00000170855 | TRIAP1 | -0.41 | 1.45E-02 | 8.33E-02 |
| ENSG00000186897 | C1QL4 | 0.41 | 1.45E-02 | 8.33E-02 |
| ENSG00000235912 | AL031729.1 | -0.41 | 1.45E-02 | 8.33E-02 |
| ENSG00000249348 | UGDH-AS1 | 0.41 | 1.45E-02 | 8.33E-02 |
| ENSG00000269313 | MAGIX | 0.41 | 1.45E-02 | 8.33E-02 |
| ENSG00000012171 | SEMA3B | 0.41 | 1.46E-02 | 8.36E-02 |
| ENSG00000120694 | HSPH1 | -0.41 | 1.46E-02 | 8.36E-02 |
| ENSG00000138614 | INTS14 | -0.41 | 1.46E-02 | 8.36E-02 |
| ENSG00000142684 | ZNF593 | -0.41 | 1.46E-02 | 8.36E-02 |
| ENSG00000176842 | IRX5 | -0.41 | 1.46E-02 | 8.36E-02 |
| ENSG00000237989 | LINC01679 | 0.41 | 1.46E-02 | 8.36E-02 |
| ENSG00000271848 | AC073389.2 | 0.41 | 1.46E-02 | 8.36E-02 |
| ENSG00000228204 | AC004830.1 | -0.41 | 1.46E-02 | 8.38E-02 |
| ENSG00000275379 | HIST1H3I | -0.41 | 1.46E-02 | 8.39E-02 |
| ENSG00000117118 | SDHB | -0.41 | 1.46E-02 | 8.39E-02 |
| ENSG00000184677 | ZBTB40 | 0.41 | 1.46E-02 | 8.39E-02 |
| ENSG00000185986 | SDHAP3 | 0.41 | 1.46E-02 | 8.39E-02 |
| ENSG00000276903 | HIST1H2AL | -0.41 | 1.47E-02 | 8.41E-02 |
| ENSG00000150773 | PIH1D2 | -0.41 | 1.47E-02 | 8.42E-02 |
| ENSG00000189337 | KAZN | -0.41 | 1.47E-02 | 8.42E-02 |
| ENSG00000198205 | ZXDA | 0.41 | 1.47E-02 | 8.42E-02 |
| ENSG00000279077 | AC023090.2 | 0.41 | 1.48E-02 | 8.45E-02 |
| ENSG00000091947 | TMEM101 | -0.41 | 1.48E-02 | 8.45E-02 |
| ENSG00000161509 | GRIN2C | 0.41 | 1.48E-02 | 8.45E-02 |
| ENSG00000161847 | RAVER1 | 0.41 | 1.48E-02 | 8.45E-02 |

| Ensembl ID | Gene Name | ρ | Pval | Padj |
|---|---|---|---|---|
| ENSG00000163328 | GPR155 | -0.41 | 1.48E-02 | 8.45E-02 |
| ENSG00000224165 | DNAJC27-AS1 | 0.41 | 1.48E-02 | 8.45E-02 |
| ENSG00000207721 | MIR186 | 0.41 | 1.48E-02 | 8.46E-02 |
| ENSG00000228043 | AC114763.1 | 0.41 | 1.49E-02 | 8.46E-02 |
| ENSG00000237991 | RPL35P1 | -0.41 | 1.49E-02 | 8.46E-02 |
| ENSG00000105438 | KDELR1 | -0.41 | 1.49E-02 | 8.46E-02 |
| ENSG00000106617 | PRKAG2 | 0.41 | 1.49E-02 | 8.46E-02 |
| ENSG00000122376 | SHLD2 | -0.41 | 1.49E-02 | 8.46E-02 |
| ENSG00000138411 | HECW2 | -0.41 | 1.49E-02 | 8.46E-02 |
| ENSG00000144485 | HES6 | -0.41 | 1.49E-02 | 8.46E-02 |
| ENSG00000155868 | MED7 | -0.41 | 1.49E-02 | 8.46E-02 |
| ENSG00000175110 | MRPS22 | -0.41 | 1.49E-02 | 8.46E-02 |
| ENSG00000243317 | STMP1 | -0.41 | 1.49E-02 | 8.46E-02 |
| ENSG00000255468 | AP001107.9 | 0.41 | 1.49E-02 | 8.46E-02 |
| ENSG00000015475 | BID | -0.41 | 1.49E-02 | 8.48E-02 |
| ENSG00000102007 | PLP2 | -0.41 | 1.49E-02 | 8.48E-02 |
| ENSG00000104671 | DCTN6 | -0.41 | 1.49E-02 | 8.48E-02 |
| ENSG00000117395 | EBNA1BP2 | -0.41 | 1.49E-02 | 8.48E-02 |
| ENSG00000129071 | MBD4 | -0.41 | 1.49E-02 | 8.48E-02 |
| ENSG00000141349 | G6PC3 | -0.41 | 1.49E-02 | 8.48E-02 |
| ENSG00000173295 | FAM86B3P | 0.41 | 1.49E-02 | 8.48E-02 |
| ENSG00000179344 | HLA-DQB1 | -0.41 | 1.49E-02 | 8.48E-02 |
| ENSG00000214113 | LYRM4 | -0.41 | 1.49E-02 | 8.48E-02 |
| ENSG00000223396 | RPS10P7 | 0.41 | 1.49E-02 | 8.48E-02 |
| ENSG00000264538 | SUZ12P1 | 0.41 | 1.49E-02 | 8.48E-02 |
| ENSG00000101049 | SGK2 | 0.41 | 1.50E-02 | 8.51E-02 |
| ENSG00000026025 | VIM | -0.41 | 1.50E-02 | 8.51E-02 |
| ENSG00000102471 | NDFIP2 | -0.41 | 1.50E-02 | 8.51E-02 |
| ENSG00000126581 | BECN1 | -0.41 | 1.50E-02 | 8.51E-02 |
| ENSG00000157570 | TSPAN18 | -0.41 | 1.50E-02 | 8.51E-02 |
| ENSG00000133488 | SEC14L4 | 0.41 | 1.51E-02 | 8.54E-02 |
| ENSG00000139684 | ESD | -0.41 | 1.51E-02 | 8.54E-02 |
| ENSG00000171819 | ANGPTL7 | 0.41 | 1.51E-02 | 8.54E-02 |
| ENSG00000178222 | RNF212 | 0.41 | 1.51E-02 | 8.54E-02 |
| ENSG00000185485 | SDHAP1 | 0.41 | 1.51E-02 | 8.54E-02 |
| ENSG00000213523 | SRA1 | -0.41 | 1.51E-02 | 8.54E-02 |
| ENSG00000260804 | LINC01963 | -0.41 | 1.51E-02 | 8.54E-02 |
| ENSG00000274267 | HIST1H3B | -0.41 | 1.52E-02 | 8.56E-02 |
| ENSG00000116761 | CTH | 0.41 | 1.52E-02 | 8.56E-02 |
| ENSG00000130600 | H19 | 0.41 | 1.52E-02 | 8.56E-02 |
| ENSG00000136383 | ALPK3 | 0.41 | 1.52E-02 | 8.56E-02 |
| ENSG00000136960 | ENPP2 | -0.41 | 1.52E-02 | 8.56E-02 |
| ENSG00000163947 | ARHGEF3 | -0.41 | 1.52E-02 | 8.56E-02 |
| ENSG00000196453 | ZNF777 | 0.41 | 1.52E-02 | 8.56E-02 |
| ENSG00000255769 | GOLGA2P10 | 0.41 | 1.52E-02 | 8.56E-02 |

| Ensembl ID | Gene Name | ρ | Pval | Padj |
|---|---|---|---|---|
| ENSG00000277998 | AC107075.1 | -0.41 | 1.52E-02 | 8.56E-02 |
| ENSG00000135454 | B4GALNT1 | -0.41 | 1.52E-02 | 8.58E-02 |
| ENSG00000232528 | AL109809.1 | 0.41 | 1.52E-02 | 8.58E-02 |
| ENSG00000124479 | NDP | -0.41 | 1.52E-02 | 8.58E-02 |
| ENSG00000130755 | GMFG | -0.41 | 1.52E-02 | 8.58E-02 |
| ENSG00000183087 | GAS6 | 0.41 | 1.52E-02 | 8.58E-02 |
| ENSG00000188242 | AC010442.1 | 0.41 | 1.52E-02 | 8.58E-02 |
| ENSG00000273841 | TAF9 | -0.41 | 1.52E-02 | 8.58E-02 |
| ENSG00000115361 | ACADL | 0.41 | 1.53E-02 | 8.60E-02 |
| ENSG00000115163 | CENPA | -0.41 | 1.53E-02 | 8.60E-02 |
| ENSG00000069345 | DNAJA2 | -0.41 | 1.53E-02 | 8.60E-02 |
| ENSG00000111639 | MRPL51 | -0.41 | 1.53E-02 | 8.60E-02 |
| ENSG00000118523 | CCN2 | 0.41 | 1.53E-02 | 8.60E-02 |
| ENSG00000140450 | ARRDC4 | -0.41 | 1.53E-02 | 8.60E-02 |
| ENSG00000143553 | SNAPIN | -0.41 | 1.53E-02 | 8.60E-02 |
| ENSG00000146143 | PRIM2 | -0.41 | 1.53E-02 | 8.60E-02 |
| ENSG00000188002 | AC026412.1 | 0.41 | 1.53E-02 | 8.60E-02 |
| ENSG00000198466 | ZNF587 | 0.41 | 1.53E-02 | 8.60E-02 |
| ENSG00000261286 | ATP2C2-AS1 | 0.41 | 1.54E-02 | 8.62E-02 |
| ENSG00000147889 | CDKN2A | -0.41 | 1.54E-02 | 8.63E-02 |
| ENSG00000005249 | PRKAR2B | -0.41 | 1.54E-02 | 8.63E-02 |
| ENSG00000084207 | GSTP1 | -0.41 | 1.54E-02 | 8.63E-02 |
| ENSG00000103199 | ZNF500 | 0.41 | 1.54E-02 | 8.63E-02 |
| ENSG00000108671 | PSMD11 | -0.41 | 1.54E-02 | 8.63E-02 |
| ENSG00000133065 | SLC41A1 | 0.41 | 1.54E-02 | 8.63E-02 |
| ENSG00000144792 | ZNF660 | 0.41 | 1.54E-02 | 8.63E-02 |
| ENSG00000197961 | ZNF121 | -0.41 | 1.54E-02 | 8.63E-02 |
| ENSG00000256650 | RERG-IT1 | 0.41 | 1.54E-02 | 8.64E-02 |
| ENSG00000124915 | AP002380.1 | -0.41 | 1.55E-02 | 8.66E-02 |
| ENSG00000158246 | TENT5B | 0.41 | 1.55E-02 | 8.66E-02 |
| ENSG00000162882 | HAAO | 0.41 | 1.55E-02 | 8.66E-02 |
| ENSG00000170322 | NFRKB | 0.41 | 1.55E-02 | 8.66E-02 |
| ENSG00000027001 | MIPEP | -0.41 | 1.56E-02 | 8.68E-02 |
| ENSG00000114125 | RNF7 | -0.41 | 1.56E-02 | 8.68E-02 |
| ENSG00000162244 | RPL29 | -0.41 | 1.56E-02 | 8.68E-02 |
| ENSG00000164970 | FAM219A | 0.41 | 1.56E-02 | 8.68E-02 |
| ENSG00000167615 | LENG8 | 0.41 | 1.56E-02 | 8.68E-02 |
| ENSG00000188385 | JAKMIP3 | 0.41 | 1.56E-02 | 8.68E-02 |
| ENSG00000258708 | SLC25A21-AS1 | 0.41 | 1.56E-02 | 8.68E-02 |
| ENSG00000278784 | AL136295.7 | 0.41 | 1.56E-02 | 8.68E-02 |
| ENSG00000079215 | SLC1A3 | -0.41 | 1.56E-02 | 8.70E-02 |
| ENSG00000237004 | ZNRF2P1 | 0.41 | 1.56E-02 | 8.71E-02 |
| ENSG00000092208 | GEMIN2 | -0.41 | 1.56E-02 | 8.71E-02 |
| ENSG00000109133 | TMEM33 | -0.41 | 1.56E-02 | 8.71E-02 |
| ENSG00000129535 | NRL | 0.41 | 1.56E-02 | 8.71E-02 |

| Ensembl ID | Gene Name | ρ | Pval | Padj |
|---|---|---|---|---|
| ENSG00000137337 | MDC1 | 0.41 | 1.56E-02 | 8.71E-02 |
| ENSG00000177700 | POLR2L | -0.41 | 1.56E-02 | 8.71E-02 |
| ENSG00000188735 | TMEM120B | 0.41 | 1.56E-02 | 8.71E-02 |
| ENSG00000225450 | AL021707.1 | 0.41 | 1.57E-02 | 8.72E-02 |
| ENSG00000261759 | AC099518.5 | -0.41 | 1.57E-02 | 8.72E-02 |
| ENSG00000198125 | MB | -0.41 | 1.57E-02 | 8.72E-02 |
| ENSG00000166106 | ADAMTS15 | -0.41 | 1.57E-02 | 8.72E-02 |
| ENSG00000180921 | FAM83H | -0.41 | 1.57E-02 | 8.72E-02 |
| ENSG00000057935 | MTA3 | 0.41 | 1.57E-02 | 8.72E-02 |
| ENSG00000131242 | RAB11FIP4 | 0.41 | 1.57E-02 | 8.72E-02 |
| ENSG00000134255 | CEPT1 | -0.41 | 1.57E-02 | 8.72E-02 |
| ENSG00000151176 | PLBD2 | 0.41 | 1.57E-02 | 8.72E-02 |
| ENSG00000170852 | KBTBD2 | -0.41 | 1.57E-02 | 8.72E-02 |
| ENSG00000174684 | B4GAT1 | -0.41 | 1.57E-02 | 8.72E-02 |
| ENSG00000204622 | HLA-J | 0.41 | 1.57E-02 | 8.72E-02 |
| ENSG00000267007 | AC012615.2 | -0.41 | 1.57E-02 | 8.72E-02 |
| ENSG00000271581 | AL671883.2 | 0.41 | 1.58E-02 | 8.75E-02 |
| ENSG00000004975 | DVL2 | 0.41 | 1.58E-02 | 8.75E-02 |
| ENSG00000137656 | BUD13 | -0.41 | 1.58E-02 | 8.75E-02 |
| ENSG00000184863 | RBM33 | 0.41 | 1.58E-02 | 8.75E-02 |
| ENSG00000215218 | UBE2QL1 | -0.41 | 1.58E-02 | 8.75E-02 |
| ENSG00000235916 | AC233279.1 | -0.40 | 1.59E-02 | 8.78E-02 |
| ENSG00000228010 | AC073343.2 | 0.40 | 1.59E-02 | 8.78E-02 |
| ENSG00000061938 | TNK2 | 0.40 | 1.59E-02 | 8.78E-02 |
| ENSG00000110237 | ARHGEF17 | 0.40 | 1.59E-02 | 8.78E-02 |
| ENSG00000156650 | KAT6B | 0.40 | 1.59E-02 | 8.78E-02 |
| ENSG00000213551 | DNAJC9 | -0.40 | 1.59E-02 | 8.78E-02 |
| ENSG00000229689 | AC009237.3 | 0.40 | 1.59E-02 | 8.78E-02 |
| ENSG00000114757 | PEX5L | 0.40 | 1.59E-02 | 8.80E-02 |
| ENSG00000274341 | AC005899.6 | 0.40 | 1.59E-02 | 8.80E-02 |
| ENSG00000137648 | TMPRSS4 | -0.40 | 1.59E-02 | 8.80E-02 |
| ENSG00000205791 | LOH12CR2 | 0.40 | 1.59E-02 | 8.80E-02 |
| ENSG00000115540 | MOB4 | -0.40 | 1.60E-02 | 8.80E-02 |
| ENSG00000179981 | TSHZ1 | 0.40 | 1.60E-02 | 8.80E-02 |
| ENSG00000183828 | NUDT14 | -0.40 | 1.60E-02 | 8.80E-02 |
| ENSG00000186566 | GPATCH8 | 0.40 | 1.60E-02 | 8.80E-02 |
| ENSG00000276248 | AL442125.1 | 0.40 | 1.60E-02 | 8.80E-02 |
| ENSG00000272506 | AL357078.3 | 0.40 | 1.60E-02 | 8.82E-02 |
| ENSG00000188282 | RUFY4 | -0.40 | 1.60E-02 | 8.83E-02 |
| ENSG00000120332 | TNN | -0.40 | 1.60E-02 | 8.83E-02 |
| ENSG00000205464 | ATP6AP1L | 0.40 | 1.60E-02 | 8.83E-02 |
| ENSG00000224722 | AC020688.1 | 0.40 | 1.60E-02 | 8.83E-02 |
| ENSG00000185187 | SIGIRR | -0.40 | 1.61E-02 | 8.87E-02 |
| ENSG00000272320 | AL445309.1 | 0.40 | 1.61E-02 | 8.89E-02 |
| ENSG00000120093 | HOXB3 | -0.40 | 1.62E-02 | 8.89E-02 |

| Ensembl ID | Gene Name | ρ | Pval | Padj |
|---|---|---|---|---|
| ENSG00000070444 | MNT | 0.40 | 1.62E-02 | 8.89E-02 |
| ENSG00000082074 | FYB1 | 0.40 | 1.62E-02 | 8.89E-02 |
| ENSG00000103150 | MLYCD | 0.40 | 1.62E-02 | 8.89E-02 |
| ENSG00000120071 | KANSL1 | 0.40 | 1.62E-02 | 8.89E-02 |
| ENSG00000137100 | DCTN3 | -0.40 | 1.62E-02 | 8.89E-02 |
| ENSG00000157510 | AFAP1L1 | -0.40 | 1.62E-02 | 8.89E-02 |
| ENSG00000184983 | NDUFA6 | -0.40 | 1.62E-02 | 8.89E-02 |
| ENSG00000228274 | AL021707.2 | 0.40 | 1.62E-02 | 8.89E-02 |
| ENSG00000235605 | AL355472.1 | -0.40 | 1.62E-02 | 8.89E-02 |
| ENSG00000235436 | DPY19L2P4 | 0.40 | 1.62E-02 | 8.91E-02 |
| ENSG00000110079 | MS4A4A | -0.40 | 1.62E-02 | 8.91E-02 |
| ENSG00000260920 | AL031985.3 | -0.40 | 1.63E-02 | 8.91E-02 |
| ENSG00000105854 | PON2 | -0.40 | 1.63E-02 | 8.91E-02 |
| ENSG00000115423 | DNAH6 | 0.40 | 1.63E-02 | 8.91E-02 |
| ENSG00000123106 | CCDC91 | -0.40 | 1.63E-02 | 8.91E-02 |
| ENSG00000137073 | UBAP2 | 0.40 | 1.63E-02 | 8.91E-02 |
| ENSG00000165678 | GHITM | -0.40 | 1.63E-02 | 8.91E-02 |
| ENSG00000179818 | PCBP1-AS1 | 0.40 | 1.63E-02 | 8.91E-02 |
| ENSG00000263266 | RPS7P1 | -0.40 | 1.63E-02 | 8.91E-02 |
| ENSG00000150627 | WDR17 | 0.40 | 1.63E-02 | 8.93E-02 |
| ENSG00000166793 | YPEL4 | -0.40 | 1.63E-02 | 8.93E-02 |
| ENSG00000072849 | DERL2 | -0.40 | 1.64E-02 | 8.94E-02 |
| ENSG00000182957 | SPATA13 | 0.40 | 1.64E-02 | 8.94E-02 |
| ENSG00000232160 | RAP2C-AS1 | 0.40 | 1.64E-02 | 8.94E-02 |
| ENSG00000237637 | FRY-AS1 | 0.40 | 1.64E-02 | 8.96E-02 |
| ENSG00000264290 | AC104564.3 | 0.40 | 1.64E-02 | 8.96E-02 |
| ENSG00000126249 | PDCD2L | -0.40 | 1.64E-02 | 8.96E-02 |
| ENSG00000099250 | NRP1 | -0.40 | 1.64E-02 | 8.96E-02 |
| ENSG00000106799 | TGFBR1 | -0.40 | 1.64E-02 | 8.96E-02 |
| ENSG00000136235 | GPNMB | -0.40 | 1.64E-02 | 8.96E-02 |
| ENSG00000144959 | NCEH1 | -0.40 | 1.64E-02 | 8.96E-02 |
| ENSG00000152422 | XRCC4 | -0.40 | 1.64E-02 | 8.96E-02 |
| ENSG00000159214 | CCDC24 | 0.40 | 1.64E-02 | 8.96E-02 |
| ENSG00000170500 | LONRF2 | 0.40 | 1.64E-02 | 8.96E-02 |
| ENSG00000177156 | TALDO1 | -0.40 | 1.64E-02 | 8.96E-02 |
| ENSG00000223878 | PPIAP53 | 0.40 | 1.65E-02 | 8.98E-02 |
| ENSG00000121621 | KIF18A | -0.40 | 1.65E-02 | 8.98E-02 |
| ENSG00000279259 | AC087741.3 | 0.40 | 1.65E-02 | 8.98E-02 |
| ENSG00000103042 | SLC38A7 | -0.40 | 1.65E-02 | 8.98E-02 |
| ENSG00000106123 | EPHB6 | 0.40 | 1.65E-02 | 8.98E-02 |
| ENSG00000138663 | COPS4 | -0.40 | 1.65E-02 | 8.98E-02 |
| ENSG00000189376 | C8orf76 | -0.40 | 1.65E-02 | 8.98E-02 |
| ENSG00000270069 | MIR222HG | -0.40 | 1.65E-02 | 8.98E-02 |
| ENSG00000276023 | DUSP14 | -0.40 | 1.65E-02 | 8.98E-02 |
| ENSG00000000971 | CFH | 0.40 | 1.66E-02 | 9.02E-02 |

| Ensembl ID | Gene Name | ρ | Pval | Padj |
|---|---|---|---|---|
| ENSG00000121289 | CEP89 | -0.40 | 1.66E-02 | 9.02E-02 |
| ENSG00000272168 | CASC15 | -0.40 | 1.66E-02 | 9.02E-02 |
| ENSG00000120279 | MYCT1 | -0.40 | 1.67E-02 | 9.05E-02 |
| ENSG00000204052 | LRRC73 | -0.40 | 1.67E-02 | 9.05E-02 |
| ENSG00000070047 | PHRF1 | 0.40 | 1.67E-02 | 9.05E-02 |
| ENSG00000106080 | FKBP14 | -0.40 | 1.67E-02 | 9.05E-02 |
| ENSG00000106615 | RHEB | -0.40 | 1.67E-02 | 9.05E-02 |
| ENSG00000153363 | LINC00467 | 0.40 | 1.67E-02 | 9.05E-02 |
| ENSG00000160255 | ITGB2 | -0.40 | 1.67E-02 | 9.05E-02 |
| ENSG00000183337 | BCOR | 0.40 | 1.67E-02 | 9.05E-02 |
| ENSG00000145431 | PDGFC | -0.40 | 1.68E-02 | 9.08E-02 |
| ENSG00000145675 | PIK3R1 | 0.40 | 1.68E-02 | 9.08E-02 |
| ENSG00000166925 | TSC22D4 | 0.40 | 1.68E-02 | 9.08E-02 |
| ENSG00000172590 | MRPL52 | -0.40 | 1.68E-02 | 9.08E-02 |
| ENSG00000173548 | SNX33 | 0.40 | 1.68E-02 | 9.08E-02 |
| ENSG00000198171 | DDRGK1 | -0.40 | 1.68E-02 | 9.08E-02 |
| ENSG00000042980 | ADAM28 | -0.40 | 1.68E-02 | 9.08E-02 |
| ENSG00000233144 | AL512656.1 | 0.40 | 1.68E-02 | 9.10E-02 |
| ENSG00000039123 | MTREX | -0.40 | 1.69E-02 | 9.10E-02 |
| ENSG00000080824 | HSP90AA1 | -0.40 | 1.69E-02 | 9.10E-02 |
| ENSG00000092010 | PSME1 | -0.40 | 1.69E-02 | 9.10E-02 |
| ENSG00000116885 | OSCP1 | -0.40 | 1.69E-02 | 9.10E-02 |
| ENSG00000134020 | PEBP4 | 0.40 | 1.69E-02 | 9.10E-02 |
| ENSG00000182890 | GLUD2 | -0.40 | 1.69E-02 | 9.10E-02 |
| ENSG00000196700 | ZNF512B | 0.40 | 1.69E-02 | 9.10E-02 |
| ENSG00000269054 | AC012313.6 | 0.40 | 1.69E-02 | 9.12E-02 |
| ENSG00000129696 | TTI2 | -0.40 | 1.69E-02 | 9.13E-02 |
| ENSG00000168056 | LTBP3 | 0.40 | 1.69E-02 | 9.13E-02 |
| ENSG00000171863 | RPS7 | -0.40 | 1.69E-02 | 9.13E-02 |
| ENSG00000174137 | FAM53A | 0.40 | 1.69E-02 | 9.13E-02 |
| ENSG00000175322 | ZNF519 | 0.40 | 1.69E-02 | 9.13E-02 |
| ENSG00000187098 | MITF | -0.40 | 1.69E-02 | 9.13E-02 |
| ENSG00000254614 | AP003068.2 | 0.40 | 1.69E-02 | 9.13E-02 |
| ENSG00000196209 | SIRPB2 | -0.40 | 1.69E-02 | 9.13E-02 |
| ENSG00000259007 | AL358333.3 | -0.40 | 1.70E-02 | 9.15E-02 |
| ENSG00000073969 | NSF | -0.40 | 1.70E-02 | 9.15E-02 |
| ENSG00000104497 | SNX16 | -0.40 | 1.70E-02 | 9.15E-02 |
| ENSG00000127184 | COX7C | -0.40 | 1.70E-02 | 9.15E-02 |
| ENSG00000134461 | ANKRD16 | 0.40 | 1.70E-02 | 9.15E-02 |
| ENSG00000163659 | TIPARP | 0.40 | 1.70E-02 | 9.15E-02 |
| ENSG00000196136 | SERPINA3 | 0.40 | 1.70E-02 | 9.15E-02 |
| ENSG00000213722 | DDAH2 | -0.40 | 1.70E-02 | 9.15E-02 |
| ENSG00000268985 | AC020908.2 | 0.40 | 1.71E-02 | 9.17E-02 |
| ENSG00000112851 | ERBIN | -0.40 | 1.71E-02 | 9.18E-02 |
| ENSG00000152433 | ZNF547 | -0.40 | 1.71E-02 | 9.18E-02 |

| Ensembl ID | Gene Name | ρ | Pval | Padj |
|---|---|---|---|---|
| ENSG00000175592 | FOSL1 | -0.40 | 1.71E-02 | 9.18E-02 |
| ENSG00000180011 | ZADH2 | 0.40 | 1.71E-02 | 9.18E-02 |
| ENSG00000247572 | CKMT2-AS1 | 0.40 | 1.71E-02 | 9.18E-02 |
| ENSG00000259330 | INAFM2 | -0.40 | 1.72E-02 | 9.20E-02 |
| ENSG00000254780 | AC023232.1 | -0.40 | 1.72E-02 | 9.20E-02 |
| ENSG00000114520 | SNX4 | -0.40 | 1.72E-02 | 9.20E-02 |
| ENSG00000153531 | ADPRHL1 | 0.40 | 1.72E-02 | 9.20E-02 |
| ENSG00000166595 | CIAO2B | -0.40 | 1.72E-02 | 9.20E-02 |
| ENSG00000175984 | DENND2C | 0.40 | 1.72E-02 | 9.20E-02 |
| ENSG00000182871 | COL18A1 | -0.40 | 1.72E-02 | 9.20E-02 |
| ENSG00000198712 | MT-CO2 | -0.40 | 1.72E-02 | 9.20E-02 |
| ENSG00000226419 | SLC16A1-AS1 | 0.40 | 1.72E-02 | 9.20E-02 |
| ENSG00000224195 | AC022400.1 | 0.40 | 1.72E-02 | 9.21E-02 |
| ENSG00000278376 | AP004609.3 | 0.40 | 1.72E-02 | 9.23E-02 |
| ENSG00000111670 | GNPTAB | -0.40 | 1.73E-02 | 9.23E-02 |
| ENSG00000152240 | HAUS1 | -0.40 | 1.73E-02 | 9.23E-02 |
| ENSG00000164951 | PDP1 | -0.40 | 1.73E-02 | 9.23E-02 |
| ENSG00000182600 | SNORC | 0.40 | 1.73E-02 | 9.23E-02 |
| ENSG00000187391 | MAGI2 | 0.40 | 1.73E-02 | 9.23E-02 |
| ENSG00000226174 | TEX22 | 0.40 | 1.74E-02 | 9.25E-02 |
| ENSG00000087088 | BAX | -0.40 | 1.74E-02 | 9.25E-02 |
| ENSG00000101888 | NXT2 | -0.40 | 1.74E-02 | 9.25E-02 |
| ENSG00000106258 | CYP3A5 | 0.40 | 1.74E-02 | 9.25E-02 |
| ENSG00000118804 | STBD1 | 0.40 | 1.74E-02 | 9.25E-02 |
| ENSG00000122257 | RBBP6 | 0.40 | 1.74E-02 | 9.25E-02 |
| ENSG00000136146 | MED4 | -0.40 | 1.74E-02 | 9.25E-02 |
| ENSG00000143207 | COP1 | -0.40 | 1.74E-02 | 9.25E-02 |
| ENSG00000164023 | SGMS2 | -0.40 | 1.74E-02 | 9.25E-02 |
| ENSG00000188257 | PLA2G2A | 0.40 | 1.74E-02 | 9.25E-02 |
| ENSG00000219487 | AL603766.1 | -0.40 | 1.74E-02 | 9.29E-02 |
| ENSG00000148572 | NRBF2 | -0.40 | 1.74E-02 | 9.29E-02 |
| ENSG00000149923 | PPP4C | -0.40 | 1.74E-02 | 9.29E-02 |
| ENSG00000165775 | FUNDC2 | -0.40 | 1.74E-02 | 9.29E-02 |
| ENSG00000226989 | AL049758.1 | 0.40 | 1.75E-02 | 9.32E-02 |
| ENSG00000086289 | EPDR1 | -0.40 | 1.75E-02 | 9.32E-02 |
| ENSG00000185787 | MORF4L1 | -0.40 | 1.75E-02 | 9.32E-02 |
| ENSG00000256006 | AC084117.1 | 0.40 | 1.75E-02 | 9.32E-02 |
| ENSG00000267379 | AC008569.1 | -0.40 | 1.75E-02 | 9.33E-02 |
| ENSG00000133983 | COX16 | -0.40 | 1.76E-02 | 9.35E-02 |
| ENSG00000078246 | TULP3 | 0.40 | 1.76E-02 | 9.35E-02 |
| ENSG00000108799 | EZH1 | 0.40 | 1.76E-02 | 9.35E-02 |
| ENSG00000141905 | NFIC | 0.40 | 1.76E-02 | 9.35E-02 |
| ENSG00000161203 | AP2M1 | -0.40 | 1.76E-02 | 9.35E-02 |
| ENSG00000166900 | STX3 | -0.40 | 1.76E-02 | 9.35E-02 |
| ENSG00000268573 | AC011815.1 | 0.40 | 1.76E-02 | 9.35E-02 |

| Ensembl ID | Gene Name | ρ | Pval | Padj |
|---|---|---|---|---|
| ENSG00000205838 | TTC23L | 0.40 | 1.76E-02 | 9.35E-02 |
| ENSG00000110080 | ST3GAL4 | -0.40 | 1.77E-02 | 9.37E-02 |
| ENSG00000110200 | ANAPC15 | -0.40 | 1.77E-02 | 9.37E-02 |
| ENSG00000110514 | MADD | 0.40 | 1.77E-02 | 9.37E-02 |
| ENSG00000132740 | IGHMBP2 | 0.40 | 1.77E-02 | 9.37E-02 |
| ENSG00000157916 | RER1 | -0.40 | 1.77E-02 | 9.37E-02 |
| ENSG00000166887 | VPS39 | 0.40 | 1.77E-02 | 9.37E-02 |
| ENSG00000250539 | KRT8P33 | 0.40 | 1.77E-02 | 9.37E-02 |
| ENSG00000256591 | AP003108.2 | 0.40 | 1.77E-02 | 9.37E-02 |
| ENSG00000262879 | AC068152.1 | 0.40 | 1.77E-02 | 9.37E-02 |
| ENSG00000268006 | PTOV1-AS1 | 0.40 | 1.77E-02 | 9.37E-02 |
| ENSG00000174371 | EXO1 | -0.40 | 1.77E-02 | 9.39E-02 |
| ENSG00000186907 | RTN4RL2 | -0.40 | 1.78E-02 | 9.39E-02 |
| ENSG00000106028 | SSBP1 | -0.40 | 1.78E-02 | 9.39E-02 |
| ENSG00000120675 | DNAJC15 | -0.40 | 1.78E-02 | 9.39E-02 |
| ENSG00000154001 | PPP2R5E | -0.40 | 1.78E-02 | 9.39E-02 |
| ENSG00000159228 | CBR1 | -0.40 | 1.78E-02 | 9.39E-02 |
| ENSG00000162976 | PQLC3 | -0.40 | 1.78E-02 | 9.39E-02 |
| ENSG00000180573 | HIST1H2AC | -0.40 | 1.78E-02 | 9.39E-02 |
| ENSG00000187951 | AC091057.1 | -0.40 | 1.78E-02 | 9.39E-02 |
| ENSG00000221886 | ZBED8 | 0.40 | 1.78E-02 | 9.39E-02 |
| ENSG00000228506 | AL513550.1 | 0.40 | 1.78E-02 | 9.39E-02 |
| ENSG00000123545 | NDUFAF4 | -0.40 | 1.79E-02 | 9.42E-02 |
| ENSG00000135069 | PSAT1 | -0.40 | 1.79E-02 | 9.42E-02 |
| ENSG00000167779 | IGFBP6 | -0.40 | 1.79E-02 | 9.42E-02 |
| ENSG00000171055 | FEZ2 | -0.40 | 1.79E-02 | 9.42E-02 |
| ENSG00000176928 | GCNT4 | -0.40 | 1.79E-02 | 9.42E-02 |
| ENSG00000179361 | ARID3B | 0.40 | 1.79E-02 | 9.42E-02 |
| ENSG00000242028 | HYPK | 0.40 | 1.79E-02 | 9.42E-02 |
| ENSG00000115232 | ITGA4 | -0.40 | 1.79E-02 | 9.43E-02 |
| ENSG00000146192 | FGD2 | -0.40 | 1.79E-02 | 9.44E-02 |
| ENSG00000074695 | LMAN1 | -0.40 | 1.80E-02 | 9.44E-02 |
| ENSG00000158042 | MRPL17 | -0.40 | 1.80E-02 | 9.44E-02 |
| ENSG00000161243 | FBXO27 | 0.40 | 1.80E-02 | 9.44E-02 |
| ENSG00000169189 | NSMCE1 | -0.40 | 1.80E-02 | 9.44E-02 |
| ENSG00000173113 | TRMT112 | -0.40 | 1.80E-02 | 9.44E-02 |
| ENSG00000241494 | AL355032.1 | -0.40 | 1.80E-02 | 9.44E-02 |
| ENSG00000276410 | HIST1H2BB | -0.40 | 1.80E-02 | 9.45E-02 |
| ENSG00000267573 | KRT8P5 | 0.40 | 1.80E-02 | 9.47E-02 |
| ENSG00000137563 | GGH | -0.40 | 1.81E-02 | 9.47E-02 |
| ENSG00000140263 | SORD | -0.40 | 1.81E-02 | 9.47E-02 |
| ENSG00000142039 | CCDC97 | 0.40 | 1.81E-02 | 9.47E-02 |
| ENSG00000170860 | LSM3 | -0.40 | 1.81E-02 | 9.47E-02 |
| ENSG00000271034 | AC139700.1 | 0.40 | 1.81E-02 | 9.47E-02 |
| ENSG00000041880 | PARP3 | 0.40 | 1.81E-02 | 9.51E-02 |

| Ensembl ID | Gene Name | ρ | Pval | Padj |
|---|---|---|---|---|
| ENSG00000108883 | EFTUD2 | -0.40 | 1.81E-02 | 9.51E-02 |
| ENSG00000147536 | GINS4 | -0.40 | 1.81E-02 | 9.51E-02 |
| ENSG00000168961 | LGALS9 | -0.40 | 1.81E-02 | 9.51E-02 |
| ENSG00000196502 | SULT1A1 | 0.40 | 1.81E-02 | 9.51E-02 |
| ENSG00000250602 | AC093535.1 | 0.40 | 1.82E-02 | 9.52E-02 |
| ENSG00000137955 | RABGGTB | -0.40 | 1.82E-02 | 9.54E-02 |
| ENSG00000144468 | RHBDD1 | -0.40 | 1.82E-02 | 9.54E-02 |
| ENSG00000163558 | PRKCI | -0.40 | 1.82E-02 | 9.54E-02 |
| ENSG00000078487 | ZCWPW1 | 0.40 | 1.83E-02 | 9.58E-02 |
| ENSG00000118058 | KMT2A | 0.40 | 1.83E-02 | 9.58E-02 |
| ENSG00000153767 | GTF2E1 | -0.40 | 1.83E-02 | 9.58E-02 |
| ENSG00000162545 | CAMK2N1 | -0.40 | 1.83E-02 | 9.58E-02 |
| ENSG00000259683 | AC243562.2 | 0.40 | 1.83E-02 | 9.58E-02 |
| ENSG00000269371 | AC008878.4 | -0.40 | 1.84E-02 | 9.59E-02 |
| ENSG00000214188 | ST7-OT4 | 0.40 | 1.84E-02 | 9.59E-02 |
| ENSG00000072135 | PTPN18 | 0.40 | 1.84E-02 | 9.60E-02 |
| ENSG00000076003 | MCM6 | -0.40 | 1.84E-02 | 9.60E-02 |
| ENSG00000090382 | LYZ | -0.40 | 1.84E-02 | 9.60E-02 |
| ENSG00000119522 | DENND1A | 0.40 | 1.84E-02 | 9.60E-02 |
| ENSG00000134884 | ARGLU1 | 0.40 | 1.84E-02 | 9.60E-02 |
| ENSG00000169826 | CSGALNACT2 | -0.40 | 1.84E-02 | 9.60E-02 |
| ENSG00000254999 | BRK1 | -0.40 | 1.84E-02 | 9.60E-02 |
| ENSG00000280383 | Z95331.1 | 0.40 | 1.84E-02 | 9.60E-02 |
| ENSG00000119446 | RBM18 | -0.40 | 1.85E-02 | 9.64E-02 |
| ENSG00000136040 | PLXNC1 | -0.40 | 1.85E-02 | 9.64E-02 |
| ENSG00000225606 | AC005281.1 | 0.40 | 1.85E-02 | 9.64E-02 |
| ENSG00000100365 | NCF4 | -0.40 | 1.86E-02 | 9.67E-02 |
| ENSG00000128654 | MTX2 | -0.40 | 1.86E-02 | 9.67E-02 |
| ENSG00000139263 | LRIG3 | 0.40 | 1.86E-02 | 9.67E-02 |
| ENSG00000157653 | C9orf43 | 0.40 | 1.86E-02 | 9.67E-02 |
| ENSG00000163938 | GNL3 | -0.40 | 1.86E-02 | 9.67E-02 |
| ENSG00000170917 | NUDT6 | 0.40 | 1.86E-02 | 9.67E-02 |
| ENSG00000105664 | COMP | 0.40 | 1.87E-02 | 9.71E-02 |
| ENSG00000121073 | SLC35B1 | -0.40 | 1.87E-02 | 9.71E-02 |
| ENSG00000171791 | BCL2 | 0.40 | 1.87E-02 | 9.71E-02 |
| ENSG00000171960 | PPIH | -0.40 | 1.87E-02 | 9.71E-02 |
| ENSG00000235655 | H3F3AP4 | -0.40 | 1.87E-02 | 9.71E-02 |
| ENSG00000276727 | AC137834.2 | 0.40 | 1.88E-02 | 9.73E-02 |
| ENSG00000082213 | C5orf22 | -0.40 | 1.88E-02 | 9.73E-02 |
| ENSG00000100479 | POLE2 | -0.40 | 1.88E-02 | 9.73E-02 |
| ENSG00000115289 | PCGF1 | -0.40 | 1.88E-02 | 9.73E-02 |
| ENSG00000162971 | TYW5 | 0.40 | 1.88E-02 | 9.73E-02 |
| ENSG00000172572 | PDE3A | -0.40 | 1.88E-02 | 9.73E-02 |
| ENSG00000198899 | MT-ATP6 | -0.40 | 1.88E-02 | 9.73E-02 |
| ENSG00000221890 | NPTXR | 0.40 | 1.88E-02 | 9.73E-02 |

| Ensembl ID | Gene Name | ρ | Pval | Padj |
|---|---|---|---|---|
| ENSG00000246082 | NUDT16P1 | 0.40 | 1.88E-02 | 9.73E-02 |
| ENSG00000134297 | PLEKHA8P1 | -0.39 | 1.89E-02 | 9.77E-02 |
| ENSG00000198918 | RPL39 | -0.39 | 1.89E-02 | 9.77E-02 |
| ENSG00000181097 | AC105219.1 | 0.39 | 1.89E-02 | 9.79E-02 |
| ENSG00000083123 | BCKDHB | 0.39 | 1.90E-02 | 9.80E-02 |
| ENSG00000179241 | LDLRAD3 | 0.39 | 1.90E-02 | 9.80E-02 |
| ENSG00000205277 | MUC12 | -0.39 | 1.90E-02 | 9.80E-02 |
| ENSG00000217791 | ASS1P9 | 0.39 | 1.90E-02 | 9.80E-02 |
| ENSG00000272419 | LINC01145 | 0.39 | 1.90E-02 | 9.80E-02 |
| ENSG00000277791 | PSMB3 | -0.39 | 1.90E-02 | 9.80E-02 |
| ENSG00000260910 | LINC00565 | -0.39 | 1.90E-02 | 9.83E-02 |
| ENSG00000067248 | DHX29 | -0.39 | 1.91E-02 | 9.83E-02 |
| ENSG00000100749 | VRK1 | -0.39 | 1.91E-02 | 9.83E-02 |
| ENSG00000111145 | ELK3 | -0.39 | 1.91E-02 | 9.83E-02 |
| ENSG00000147400 | CETN2 | -0.39 | 1.91E-02 | 9.83E-02 |
| ENSG00000159200 | RCAN1 | -0.39 | 1.91E-02 | 9.83E-02 |
| ENSG00000161381 | PLXDC1 | -0.39 | 1.91E-02 | 9.83E-02 |
| ENSG00000198618 | PPIAP22 | -0.39 | 1.91E-02 | 9.83E-02 |
| ENSG00000214135 | AC132008.2 | 0.39 | 1.91E-02 | 9.83E-02 |
| ENSG00000235438 | ESRRAP2 | 0.39 | 1.91E-02 | 9.83E-02 |
| ENSG00000279520 | AC093525.8 | 0.39 | 1.91E-02 | 9.84E-02 |
| ENSG00000262766 | AC135050.5 | 0.39 | 1.91E-02 | 9.85E-02 |
| ENSG00000013503 | POLR3B | -0.39 | 1.91E-02 | 9.85E-02 |
| ENSG00000033030 | ZCCHC8 | 0.39 | 1.91E-02 | 9.85E-02 |
| ENSG00000105755 | ETHE1 | -0.39 | 1.91E-02 | 9.85E-02 |
| ENSG00000164818 | DNAAF5 | 0.39 | 1.91E-02 | 9.85E-02 |
| ENSG00000169139 | UBE2V2 | -0.39 | 1.91E-02 | 9.85E-02 |
| ENSG00000170873 | MTSS1 | -0.39 | 1.91E-02 | 9.85E-02 |
| ENSG00000188064 | WNT7B | -0.39 | 1.91E-02 | 9.85E-02 |
| ENSG00000246982 | Z84485.1 | 0.39 | 1.91E-02 | 9.85E-02 |
| ENSG00000066230 | SLC9A3 | 0.39 | 1.92E-02 | 9.85E-02 |
| ENSG00000080854 | IGSF9B | 0.39 | 1.92E-02 | 9.88E-02 |
| ENSG00000112782 | CLIC5 | -0.39 | 1.92E-02 | 9.88E-02 |
| ENSG00000122952 | ZWINT | -0.39 | 1.92E-02 | 9.88E-02 |
| ENSG00000145569 | OTULINL | -0.39 | 1.92E-02 | 9.88E-02 |
| ENSG00000227374 | AL157832.1 | 0.39 | 1.93E-02 | 9.92E-02 |
| ENSG00000085840 | ORC1 | -0.39 | 1.93E-02 | 9.92E-02 |
| ENSG00000110497 | AMBRA1 | 0.39 | 1.93E-02 | 9.92E-02 |
| ENSG00000133195 | SLC39A11 | 0.39 | 1.93E-02 | 9.92E-02 |
| ENSG00000179869 | ABCA13 | -0.39 | 1.93E-02 | 9.92E-02 |
| ENSG00000260007 | AC107871.1 | 0.39 | 1.93E-02 | 9.92E-02 |
| ENSG00000236654 | AC079780.1 | -0.39 | 1.94E-02 | 9.94E-02 |
| ENSG00000076662 | ICAM3 | -0.39 | 1.94E-02 | 9.95E-02 |
| ENSG00000104522 | TSTA3 | -0.39 | 1.94E-02 | 9.95E-02 |
| ENSG00000119392 | GLE1 | -0.39 | 1.94E-02 | 9.95E-02 |

| Ensembl ID | Gene Name | ρ | Pval | Padj |
|---|---|---|---|---|
| ENSG00000156521 | TYSND1 | 0.39 | 1.94E-02 | 9.95E-02 |
| ENSG00000204370 | SDHD | -0.39 | 1.94E-02 | 9.95E-02 |
| ENSG00000100314 | CABP7 | -0.39 | 1.95E-02 | 9.98E-02 |
| ENSG00000099331 | MYO9B | 0.39 | 1.95E-02 | 9.98E-02 |
| ENSG00000105518 | TMEM205 | -0.39 | 1.95E-02 | 9.98E-02 |
| ENSG00000159063 | ALG8 | -0.39 | 1.95E-02 | 9.98E-02 |
| ENSG00000167325 | RRM1 | -0.39 | 1.95E-02 | 9.98E-02 |
| ENSG00000182366 | FAM87A | -0.39 | 1.95E-02 | 9.98E-02 |
| ENSG00000205832 | C16orf96 | 0.39 | 1.95E-02 | 9.98E-02 |
| ENSG00000253540 | FAM86HP | 0.39 | 1.96E-02 | 1.00E-01 |
| ENSG00000119421 | NDUFA8 | -0.39 | 1.96E-02 | 1.00E-01 |
| ENSG00000120733 | KDM3B | 0.39 | 1.96E-02 | 1.00E-01 |
| ENSG00000141198 | TOM1L1 | -0.39 | 1.96E-02 | 1.00E-01 |
| ENSG00000141873 | SLC39A3 | -0.39 | 1.96E-02 | 1.00E-01 |
| ENSG00000144791 | LIMD1 | 0.39 | 1.96E-02 | 1.00E-01 |
| ENSG00000148429 | USP6NL | 0.39 | 1.96E-02 | 1.00E-01 |
| ENSG00000187535 | IFT140 | 0.39 | 1.96E-02 | 1.00E-01 |
| ENSG00000211451 | GNRHR2 | 0.39 | 1.96E-02 | 1.00E-01 |
| ENSG00000213928 | IRF9 | 0.39 | 1.96E-02 | 1.00E-01 |
| ENSG00000256060 | TRAPPC2B | -0.39 | 1.96E-02 | 1.00E-01 |
| ENSG00000152056 | AP1S3 | -0.39 | 1.96E-02 | 1.00E-01 |
| ENSG00000231848 | AC012354.2 | -0.39 | 1.97E-02 | 1.00E-01 |
| ENSG00000126522 | ASL | 0.39 | 1.97E-02 | 1.00E-01 |
| ENSG00000146112 | PPP1R18 | -0.39 | 1.97E-02 | 1.00E-01 |
| ENSG00000148824 | MTG1 | 0.39 | 1.97E-02 | 1.00E-01 |
| ENSG00000167524 | RSKR | 0.39 | 1.97E-02 | 1.00E-01 |
| ENSG00000230777 | RPS29P5 | -0.39 | 1.98E-02 | 1.01E-01 |
| ENSG00000271870 | AC024060.1 | -0.39 | 1.98E-02 | 1.01E-01 |
| ENSG00000144161 | ZC3H8 | 0.39 | 1.98E-02 | 1.01E-01 |
| ENSG00000168137 | SETD5 | 0.39 | 1.98E-02 | 1.01E-01 |
| ENSG00000183283 | DAZAP2 | -0.39 | 1.98E-02 | 1.01E-01 |
| ENSG00000250903 | GMDS-DT | 0.39 | 1.98E-02 | 1.01E-01 |
| ENSG00000154639 | CXADR | -0.39 | 1.98E-02 | 1.01E-01 |
| ENSG00000227709 | AC091607.1 | -0.39 | 1.99E-02 | 1.01E-01 |
| ENSG00000136883 | KIF12 | 0.39 | 1.99E-02 | 1.01E-01 |
| ENSG00000065491 | TBC1D22B | -0.39 | 1.99E-02 | 1.01E-01 |
| ENSG00000076108 | BAZ2A | 0.39 | 1.99E-02 | 1.01E-01 |
| ENSG00000100138 | SNU13 | -0.39 | 1.99E-02 | 1.01E-01 |
| ENSG00000136718 | IMP4 | -0.39 | 1.99E-02 | 1.01E-01 |
| ENSG00000152223 | EPG5 | 0.39 | 1.99E-02 | 1.01E-01 |
| ENSG00000153989 | NUS1 | -0.39 | 1.99E-02 | 1.01E-01 |
| ENSG00000172020 | GAP43 | -0.39 | 1.99E-02 | 1.01E-01 |
| ENSG00000185049 | NELFA | 0.39 | 1.99E-02 | 1.01E-01 |
| ENSG00000197063 | MAFG | -0.39 | 1.99E-02 | 1.01E-01 |
| ENSG00000204514 | ZNF814 | 0.39 | 1.99E-02 | 1.01E-01 |

| Ensembl ID | Gene Name | ρ | Pval | Padj |
|---|---|---|---|---|
| ENSG00000226803 | ZNF451-AS1 | 0.39 | 1.99E-02 | 1.01E-01 |
| ENSG00000268043 | NBPF12 | 0.39 | 1.99E-02 | 1.01E-01 |
| ENSG00000278053 | DDX52 | -0.39 | 1.99E-02 | 1.01E-01 |
| ENSG00000184730 | APOBR | -0.39 | 1.99E-02 | 1.01E-01 |
| ENSG00000066735 | KIF26A | 0.39 | 2.00E-02 | 1.01E-01 |
| ENSG00000118369 | USP35 | 0.39 | 2.00E-02 | 1.01E-01 |
| ENSG00000130203 | APOE | -0.39 | 2.00E-02 | 1.01E-01 |
| ENSG00000164185 | ZNF474 | -0.39 | 2.00E-02 | 1.01E-01 |
| ENSG00000196449 | YRDC | -0.39 | 2.00E-02 | 1.01E-01 |
| ENSG00000247137 | AP000873.2 | 0.39 | 2.00E-02 | 1.01E-01 |
| ENSG00000177994 | C2orf73 | 0.39 | 2.00E-02 | 1.01E-01 |
| ENSG00000118432 | CNR1 | 0.39 | 2.00E-02 | 1.01E-01 |
| ENSG00000274618 | HIST1H4F | -0.39 | 2.01E-02 | 1.01E-01 |
| ENSG00000015133 | CCDC88C | -0.39 | 2.01E-02 | 1.01E-01 |
| ENSG00000053372 | MRTO4 | -0.39 | 2.01E-02 | 1.01E-01 |
| ENSG00000082153 | BZW1 | -0.39 | 2.01E-02 | 1.01E-01 |
| ENSG00000119943 | PYROXD2 | 0.39 | 2.01E-02 | 1.01E-01 |
| ENSG00000127946 | HIP1 | 0.39 | 2.01E-02 | 1.01E-01 |
| ENSG00000134419 | RPS15A | -0.39 | 2.01E-02 | 1.01E-01 |
| ENSG00000143878 | RHOB | 0.39 | 2.01E-02 | 1.01E-01 |
| ENSG00000151247 | EIF4E | -0.39 | 2.01E-02 | 1.01E-01 |
| ENSG00000171314 | PGAM1 | -0.39 | 2.01E-02 | 1.01E-01 |
| ENSG00000228939 | AKT3-IT1 | 0.39 | 2.01E-02 | 1.01E-01 |
| ENSG00000073605 | GSDMB | 0.39 | 2.02E-02 | 1.02E-01 |
| ENSG00000171135 | JAGN1 | -0.39 | 2.02E-02 | 1.02E-01 |
| ENSG00000234028 | EIF2AK3-DT | 0.39 | 2.02E-02 | 1.02E-01 |
| ENSG00000113068 | PFDN1 | -0.39 | 2.03E-02 | 1.02E-01 |
| ENSG00000114491 | UMPS | -0.39 | 2.03E-02 | 1.02E-01 |
| ENSG00000120860 | WASHC3 | -0.39 | 2.03E-02 | 1.02E-01 |
| ENSG00000131149 | GSE1 | 0.39 | 2.03E-02 | 1.02E-01 |
| ENSG00000137509 | PRCP | -0.39 | 2.03E-02 | 1.02E-01 |
| ENSG00000156110 | ADK | -0.39 | 2.03E-02 | 1.02E-01 |
| ENSG00000165801 | ARHGEF40 | 0.39 | 2.03E-02 | 1.02E-01 |
| ENSG00000182621 | PLCB1 | -0.39 | 2.03E-02 | 1.02E-01 |
| ENSG00000240445 | FOXO3B | 0.39 | 2.03E-02 | 1.02E-01 |
| ENSG00000266282 | UBL5P2 | -0.39 | 2.03E-02 | 1.02E-01 |
| ENSG00000270049 | AC009061.2 | 0.39 | 2.03E-02 | 1.02E-01 |
| ENSG00000274565 | AC080038.2 | 0.39 | 2.03E-02 | 1.02E-01 |
| ENSG00000163221 | S100A12 | -0.39 | 2.03E-02 | 1.02E-01 |
| ENSG00000196839 | ADA | -0.39 | 2.04E-02 | 1.02E-01 |
| ENSG00000065559 | MAP2K4 | -0.39 | 2.04E-02 | 1.02E-01 |
| ENSG00000115526 | CHST10 | 0.39 | 2.04E-02 | 1.02E-01 |
| ENSG00000128284 | APOL3 | 0.39 | 2.04E-02 | 1.02E-01 |
| ENSG00000139505 | MTMR6 | -0.39 | 2.04E-02 | 1.02E-01 |
| ENSG00000174306 | ZHX3 | 0.39 | 2.04E-02 | 1.02E-01 |

| Ensembl ID | Gene Name | ρ | Pval | Padj |
|---|---|---|---|---|
| ENSG00000232346 | Z74021.1 | -0.39 | 2.04E-02 | 1.02E-01 |
| ENSG00000273270 | AC090114.2 | 0.39 | 2.04E-02 | 1.02E-01 |
| ENSG00000279410 | AC099494.1 | 0.39 | 2.04E-02 | 1.02E-01 |
| ENSG00000146722 | AC211486.1 | 0.39 | 2.05E-02 | 1.02E-01 |
| ENSG00000140939 | AC074143.1 | 0.39 | 2.05E-02 | 1.02E-01 |
| ENSG00000171992 | SYNPO | 0.39 | 2.05E-02 | 1.02E-01 |
| ENSG00000174307 | PHLDA3 | -0.39 | 2.05E-02 | 1.02E-01 |
| ENSG00000185436 | IFNLR1 | 0.39 | 2.05E-02 | 1.02E-01 |
| ENSG00000267436 | AC005786.3 | -0.39 | 2.05E-02 | 1.02E-01 |
| ENSG00000269937 | AC093525.7 | 0.39 | 2.05E-02 | 1.02E-01 |
| ENSG00000247774 | PCED1B-AS1 | -0.39 | 2.05E-02 | 1.02E-01 |
| ENSG00000004776 | HSPB6 | 0.39 | 2.06E-02 | 1.03E-01 |
| ENSG00000091879 | ANGPT2 | -0.39 | 2.06E-02 | 1.03E-01 |
| ENSG00000105997 | HOXA3 | 0.39 | 2.06E-02 | 1.03E-01 |
| ENSG00000124767 | GLO1 | -0.39 | 2.06E-02 | 1.03E-01 |
| ENSG00000128652 | HOXD3 | 0.39 | 2.06E-02 | 1.03E-01 |
| ENSG00000148110 | MFSD14B | -0.39 | 2.06E-02 | 1.03E-01 |
| ENSG00000181126 | HLA-V | 0.39 | 2.06E-02 | 1.03E-01 |
| ENSG00000179772 | FOXS1 | -0.39 | 2.06E-02 | 1.03E-01 |
| ENSG00000175785 | PRIMA1 | 0.39 | 2.06E-02 | 1.03E-01 |
| ENSG00000009844 | VTA1 | -0.39 | 2.07E-02 | 1.03E-01 |
| ENSG00000017427 | IGF1 | -0.39 | 2.07E-02 | 1.03E-01 |
| ENSG00000136021 | SCYL2 | -0.39 | 2.07E-02 | 1.03E-01 |
| ENSG00000185220 | PGBD2 | -0.39 | 2.07E-02 | 1.03E-01 |
| ENSG00000231128 | AL137856.1 | 0.39 | 2.07E-02 | 1.03E-01 |
| ENSG00000156869 | FRRS1 | -0.39 | 2.07E-02 | 1.03E-01 |
| ENSG00000130045 | NXNL2 | 0.39 | 2.07E-02 | 1.03E-01 |
| ENSG00000237651 | C2orf74 | 0.39 | 2.07E-02 | 1.03E-01 |
| ENSG00000004487 | KDM1A | -0.39 | 2.08E-02 | 1.03E-01 |
| ENSG00000008256 | CYTH3 | -0.39 | 2.08E-02 | 1.03E-01 |
| ENSG00000077782 | FGFR1 | 0.39 | 2.08E-02 | 1.03E-01 |
| ENSG00000115539 | PDCL3 | -0.39 | 2.08E-02 | 1.03E-01 |
| ENSG00000120656 | TAF12 | -0.39 | 2.08E-02 | 1.03E-01 |
| ENSG00000147164 | SNX12 | -0.39 | 2.08E-02 | 1.03E-01 |
| ENSG00000150093 | ITGB1 | -0.39 | 2.08E-02 | 1.03E-01 |
| ENSG00000166987 | MBD6 | 0.39 | 2.08E-02 | 1.03E-01 |
| ENSG00000261423 | TMEM202-AS1 | 0.39 | 2.08E-02 | 1.03E-01 |
| ENSG00000204482 | LST1 | -0.39 | 2.08E-02 | 1.03E-01 |
| ENSG00000160200 | CBS | 0.39 | 2.08E-02 | 1.03E-01 |
| ENSG00000101336 | HCK | -0.39 | 2.08E-02 | 1.03E-01 |
| ENSG00000204745 | AC083899.1 | -0.39 | 2.08E-02 | 1.03E-01 |
| ENSG00000033100 | CHPF2 | -0.39 | 2.09E-02 | 1.03E-01 |
| ENSG00000049540 | ELN | -0.39 | 2.09E-02 | 1.03E-01 |
| ENSG00000115902 | SLC1A4 | 0.39 | 2.09E-02 | 1.03E-01 |
| ENSG00000153214 | TMEM87B | -0.39 | 2.09E-02 | 1.03E-01 |

| Ensembl ID | Gene Name | ρ | Pval | Padj |
|---|---|---|---|---|
| ENSG00000163597 | SNHG16 | -0.39 | 2.09E-02 | 1.03E-01 |
| ENSG00000171298 | GAA | 0.39 | 2.09E-02 | 1.03E-01 |
| ENSG00000141968 | VAV1 | -0.39 | 2.09E-02 | 1.03E-01 |
| ENSG00000177570 | SAMD12 | 0.39 | 2.09E-02 | 1.03E-01 |
| ENSG00000279748 | AC008764.9 | 0.39 | 2.09E-02 | 1.04E-01 |
| ENSG00000164669 | INTS4P1 | 0.39 | 2.10E-02 | 1.04E-01 |
| ENSG00000135677 | GNS | -0.39 | 2.10E-02 | 1.04E-01 |
| ENSG00000145050 | MANF | -0.39 | 2.10E-02 | 1.04E-01 |
| ENSG00000146416 | AIG1 | -0.39 | 2.10E-02 | 1.04E-01 |
| ENSG00000170266 | GLB1 | -0.39 | 2.10E-02 | 1.04E-01 |
| ENSG00000183495 | EP400 | 0.39 | 2.10E-02 | 1.04E-01 |
| ENSG00000186300 | ZNF555 | 0.39 | 2.10E-02 | 1.04E-01 |
| ENSG00000270194 | AC097359.2 | 0.39 | 2.10E-02 | 1.04E-01 |
| ENSG00000260350 | AC012173.1 | -0.39 | 2.10E-02 | 1.04E-01 |
| ENSG00000091164 | TXNL1 | -0.39 | 2.11E-02 | 1.04E-01 |
| ENSG00000105677 | TMEM147 | -0.39 | 2.11E-02 | 1.04E-01 |
| ENSG00000125691 | RPL23 | -0.39 | 2.11E-02 | 1.04E-01 |
| ENSG00000179152 | TCAIM | -0.39 | 2.11E-02 | 1.04E-01 |
| ENSG00000198860 | TSEN15 | -0.39 | 2.11E-02 | 1.04E-01 |
| ENSG00000239887 | C1orf226 | 0.39 | 2.11E-02 | 1.04E-01 |
| ENSG00000269959 | SPACA6P-AS | 0.39 | 2.11E-02 | 1.04E-01 |
| ENSG00000231878 | SNRPFP1 | -0.39 | 2.11E-02 | 1.04E-01 |
| ENSG00000108423 | TUBD1 | 0.39 | 2.12E-02 | 1.04E-01 |
| ENSG00000140319 | SRP14 | -0.39 | 2.12E-02 | 1.04E-01 |
| ENSG00000172292 | CERS6 | -0.39 | 2.12E-02 | 1.04E-01 |
| ENSG00000105146 | AURKC | -0.39 | 2.12E-02 | 1.04E-01 |
| ENSG00000204620 | AC115618.1 | 0.39 | 2.12E-02 | 1.05E-01 |
| ENSG00000028528 | SNX1 | 0.39 | 2.13E-02 | 1.05E-01 |
| ENSG00000103066 | PLA2G15 | -0.39 | 2.13E-02 | 1.05E-01 |
| ENSG00000143570 | SLC39A1 | -0.39 | 2.13E-02 | 1.05E-01 |
| ENSG00000205517 | RGL3 | 0.39 | 2.13E-02 | 1.05E-01 |
| ENSG00000171766 | GATM | -0.39 | 2.14E-02 | 1.05E-01 |
| ENSG00000108641 | B9D1 | 0.39 | 2.14E-02 | 1.05E-01 |
| ENSG00000112297 | CRYBG1 | -0.39 | 2.14E-02 | 1.05E-01 |
| ENSG00000131094 | C1QL1 | -0.39 | 2.14E-02 | 1.05E-01 |
| ENSG00000157021 | FAM92A1P1 | -0.39 | 2.14E-02 | 1.05E-01 |
| ENSG00000171262 | FAM98B | -0.39 | 2.14E-02 | 1.05E-01 |
| ENSG00000228804 | AC072022.1 | 0.39 | 2.15E-02 | 1.05E-01 |
| ENSG00000262155 | LINC02175 | 0.39 | 2.15E-02 | 1.05E-01 |
| ENSG00000070540 | WIPI1 | -0.39 | 2.15E-02 | 1.05E-01 |
| ENSG00000103145 | HCFC1R1 | -0.39 | 2.15E-02 | 1.05E-01 |
| ENSG00000114993 | RTKN | 0.39 | 2.15E-02 | 1.05E-01 |
| ENSG00000146856 | AGBL3 | 0.39 | 2.15E-02 | 1.05E-01 |
| ENSG00000149922 | TBX6 | 0.39 | 2.15E-02 | 1.05E-01 |
| ENSG00000229091 | HSPA8P8 | -0.39 | 2.15E-02 | 1.05E-01 |

| Ensembl ID | Gene Name | ρ | Pval | Padj |
|---|---|---|---|---|
| ENSG00000227179 | PGCP1 | -0.39 | 2.15E-02 | 1.05E-01 |
| ENSG00000106484 | MEST | -0.39 | 2.16E-02 | 1.06E-01 |
| ENSG00000008311 | AASS | 0.39 | 2.16E-02 | 1.06E-01 |
| ENSG00000048140 | TSPAN17 | -0.39 | 2.16E-02 | 1.06E-01 |
| ENSG00000129083 | COPB1 | -0.39 | 2.16E-02 | 1.06E-01 |
| ENSG00000141367 | CLTC | -0.39 | 2.16E-02 | 1.06E-01 |
| ENSG00000153006 | SREK1IP1 | -0.39 | 2.16E-02 | 1.06E-01 |
| ENSG00000165731 | RET | 0.39 | 2.16E-02 | 1.06E-01 |
| ENSG00000173171 | MTX1 | -0.39 | 2.16E-02 | 1.06E-01 |
| ENSG00000228192 | AL512353.1 | 0.39 | 2.16E-02 | 1.06E-01 |
| ENSG00000206028 | Z99774.1 | -0.39 | 2.16E-02 | 1.06E-01 |
| ENSG00000182487 | NCF1B | -0.39 | 2.16E-02 | 1.06E-01 |
| ENSG00000162062 | TEDC2 | -0.39 | 2.17E-02 | 1.06E-01 |
| ENSG00000100154 | TTC28 | 0.39 | 2.17E-02 | 1.06E-01 |
| ENSG00000127774 | EMC6 | -0.39 | 2.17E-02 | 1.06E-01 |
| ENSG00000130560 | UBAC1 | -0.39 | 2.17E-02 | 1.06E-01 |
| ENSG00000142661 | MYOM3 | -0.39 | 2.17E-02 | 1.06E-01 |
| ENSG00000165416 | SUGT1 | -0.39 | 2.17E-02 | 1.06E-01 |
| ENSG00000180385 | EMC3-AS1 | 0.39 | 2.17E-02 | 1.06E-01 |
| ENSG00000103569 | AQP9 | -0.39 | 2.17E-02 | 1.06E-01 |
| ENSG00000270021 | AC026691.1 | 0.39 | 2.18E-02 | 1.06E-01 |
| ENSG00000279555 | AC091181.1 | 0.39 | 2.18E-02 | 1.06E-01 |
| ENSG00000226054 | MEMO1P1 | -0.39 | 2.18E-02 | 1.06E-01 |
| ENSG00000108384 | RAD51C | -0.39 | 2.18E-02 | 1.06E-01 |
| ENSG00000125753 | VASP | -0.39 | 2.18E-02 | 1.06E-01 |
| ENSG00000136280 | CCM2 | -0.39 | 2.18E-02 | 1.06E-01 |
| ENSG00000144401 | METTL21A | -0.39 | 2.18E-02 | 1.06E-01 |
| ENSG00000145386 | CCNA2 | -0.39 | 2.18E-02 | 1.06E-01 |
| ENSG00000128713 | HOXD11 | 0.39 | 2.19E-02 | 1.06E-01 |
| ENSG00000089280 | FUS | 0.39 | 2.19E-02 | 1.06E-01 |
| ENSG00000132475 | H3F3B | -0.39 | 2.19E-02 | 1.06E-01 |
| ENSG00000137504 | CREBZF | 0.39 | 2.19E-02 | 1.06E-01 |
| ENSG00000149257 | SERPINH1 | -0.39 | 2.19E-02 | 1.06E-01 |
| ENSG00000164142 | FAM160A1 | 0.39 | 2.19E-02 | 1.06E-01 |
| ENSG00000233426 | EIF3FP3 | -0.39 | 2.19E-02 | 1.06E-01 |
| ENSG00000235852 | AC005540.1 | 0.39 | 2.20E-02 | 1.07E-01 |
| ENSG00000169764 | UGP2 | -0.39 | 2.20E-02 | 1.07E-01 |
| ENSG00000177613 | CSTF2T | -0.39 | 2.20E-02 | 1.07E-01 |
| ENSG00000185721 | DRG1 | -0.39 | 2.20E-02 | 1.07E-01 |
| ENSG00000260400 | AL513534.2 | 0.39 | 2.20E-02 | 1.07E-01 |
| ENSG00000110934 | BIN2 | -0.39 | 2.21E-02 | 1.07E-01 |
| ENSG00000105855 | ITGB8 | -0.39 | 2.21E-02 | 1.07E-01 |
| ENSG00000108176 | DNAJC12 | -0.39 | 2.21E-02 | 1.07E-01 |
| ENSG00000114395 | CYB561D2 | -0.39 | 2.21E-02 | 1.07E-01 |
| ENSG00000121691 | CAT | -0.39 | 2.21E-02 | 1.07E-01 |

| Ensembl ID | Gene Name | ρ | Pval | Padj |
|---|---|---|---|---|
| ENSG00000161980 | POLR3K | -0.39 | 2.21E-02 | 1.07E-01 |
| ENSG00000171720 | HDAC3 | -0.39 | 2.21E-02 | 1.07E-01 |
| ENSG00000171817 | ZNF540 | 0.39 | 2.21E-02 | 1.07E-01 |
| ENSG00000186395 | KRT10 | -0.39 | 2.21E-02 | 1.07E-01 |
| ENSG00000228205 | AC131235.1 | -0.39 | 2.21E-02 | 1.07E-01 |
| ENSG00000260792 | LINC02280 | 0.39 | 2.21E-02 | 1.07E-01 |
| ENSG00000072694 | FCGR2B | -0.39 | 2.21E-02 | 1.07E-01 |
| ENSG00000253768 | AC008663.1 | 0.39 | 2.21E-02 | 1.07E-01 |
| ENSG00000271334 | LINC02104 | -0.39 | 2.22E-02 | 1.07E-01 |
| ENSG00000141506 | PIK3R5 | -0.39 | 2.22E-02 | 1.07E-01 |
| ENSG00000101311 | FERMT1 | -0.39 | 2.22E-02 | 1.07E-01 |
| ENSG00000127955 | GNAI1 | -0.39 | 2.22E-02 | 1.07E-01 |
| ENSG00000130669 | PAK4 | 0.39 | 2.22E-02 | 1.07E-01 |
| ENSG00000158321 | AUTS2 | 0.39 | 2.22E-02 | 1.07E-01 |
| ENSG00000164161 | HHIP | -0.39 | 2.22E-02 | 1.07E-01 |
| ENSG00000177054 | ZDHHC13 | -0.39 | 2.22E-02 | 1.07E-01 |
| ENSG00000181649 | PHLDA2 | -0.39 | 2.22E-02 | 1.07E-01 |
| ENSG00000213246 | SUPT4H1 | -0.39 | 2.22E-02 | 1.07E-01 |
| ENSG00000232112 | TMA7 | -0.39 | 2.22E-02 | 1.07E-01 |
| ENSG00000236671 | PRKG1-AS1 | 0.39 | 2.23E-02 | 1.07E-01 |
| ENSG00000139197 | PEX5 | 0.39 | 2.23E-02 | 1.08E-01 |
| ENSG00000164091 | WDR82 | -0.39 | 2.23E-02 | 1.08E-01 |
| ENSG00000176658 | MYO1D | 0.39 | 2.23E-02 | 1.08E-01 |
| ENSG00000231889 | TRAF3IP2-AS1 | 0.39 | 2.23E-02 | 1.08E-01 |
| ENSG00000235174 | RPL39P3 | -0.39 | 2.23E-02 | 1.08E-01 |
| ENSG00000253704 | AC023632.2 | 0.39 | 2.23E-02 | 1.08E-01 |
| ENSG00000131381 | RBSN | 0.38 | 2.24E-02 | 1.08E-01 |
| ENSG00000133937 | GSC | -0.38 | 2.24E-02 | 1.08E-01 |
| ENSG00000163535 | SGO2 | -0.38 | 2.24E-02 | 1.08E-01 |
| ENSG00000169299 | PGM2 | -0.38 | 2.24E-02 | 1.08E-01 |
| ENSG00000175182 | FAM131A | 0.38 | 2.24E-02 | 1.08E-01 |
| ENSG00000004700 | RECQL | -0.38 | 2.25E-02 | 1.08E-01 |
| ENSG00000005102 | MEOX1 | -0.38 | 2.25E-02 | 1.08E-01 |
| ENSG00000135211 | TMEM60 | -0.38 | 2.25E-02 | 1.08E-01 |
| ENSG00000139613 | SMARCC2 | 0.38 | 2.25E-02 | 1.08E-01 |
| ENSG00000155304 | HSPA13 | -0.38 | 2.25E-02 | 1.08E-01 |
| ENSG00000163634 | THOC7 | -0.38 | 2.25E-02 | 1.08E-01 |
| ENSG00000183386 | FHL3 | -0.38 | 2.25E-02 | 1.08E-01 |
| ENSG00000187091 | PLCD1 | 0.38 | 2.25E-02 | 1.08E-01 |
| ENSG00000272333 | KMT2B | 0.38 | 2.25E-02 | 1.08E-01 |
| ENSG00000272551 | AC017048.3 | 0.38 | 2.25E-02 | 1.08E-01 |
| ENSG00000070961 | ATP2B1 | -0.38 | 2.26E-02 | 1.08E-01 |
| ENSG00000086159 | AQP6 | 0.38 | 2.26E-02 | 1.08E-01 |
| ENSG00000139372 | TDG | -0.38 | 2.26E-02 | 1.08E-01 |
| ENSG00000149016 | TUT1 | 0.38 | 2.26E-02 | 1.08E-01 |

| Ensembl ID | Gene Name | ρ | Pval | Padj |
|---|---|---|---|---|
| ENSG00000154380 | ENAH | -0.38 | 2.26E-02 | 1.08E-01 |
| ENSG00000176971 | FIBIN | 0.38 | 2.26E-02 | 1.08E-01 |
| ENSG00000250317 | SMIM20 | -0.38 | 2.26E-02 | 1.08E-01 |
| ENSG00000266714 | MYO15B | 0.38 | 2.26E-02 | 1.08E-01 |
| ENSG00000238390 | RF01241 | 0.38 | 2.26E-02 | 1.08E-01 |
| ENSG00000157654 | PALM2-AKAP2 | -0.38 | 2.27E-02 | 1.08E-01 |
| ENSG00000108352 | RAPGEFL1 | 0.38 | 2.28E-02 | 1.09E-01 |
| ENSG00000113212 | PCDHB7 | 0.38 | 2.28E-02 | 1.09E-01 |
| ENSG00000129038 | LOXL1 | -0.38 | 2.28E-02 | 1.09E-01 |
| ENSG00000133872 | SARAF | -0.38 | 2.28E-02 | 1.09E-01 |
| ENSG00000163584 | RPL22L1 | -0.38 | 2.28E-02 | 1.09E-01 |
| ENSG00000167216 | KATNAL2 | 0.38 | 2.28E-02 | 1.09E-01 |
| ENSG00000170745 | KCNS3 | -0.38 | 2.28E-02 | 1.09E-01 |
| ENSG00000256293 | AC135279.2 | -0.38 | 2.28E-02 | 1.09E-01 |
| ENSG00000169385 | RNASE2 | -0.38 | 2.28E-02 | 1.09E-01 |
| ENSG00000105722 | ERF | 0.38 | 2.29E-02 | 1.09E-01 |
| ENSG00000167625 | ZNF526 | 0.38 | 2.29E-02 | 1.09E-01 |
| ENSG00000173011 | TADA2B | 0.38 | 2.29E-02 | 1.09E-01 |
| ENSG00000188732 | FAM221A | 0.38 | 2.29E-02 | 1.09E-01 |
| ENSG00000162888 | C1orf147 | 0.38 | 2.29E-02 | 1.09E-01 |
| ENSG00000012232 | EXTL3 | -0.38 | 2.30E-02 | 1.10E-01 |
| ENSG00000161692 | DBF4B | 0.38 | 2.30E-02 | 1.10E-01 |
| ENSG00000115896 | PLCL1 | -0.38 | 2.30E-02 | 1.10E-01 |
| ENSG00000118777 | ABCG2 | -0.38 | 2.30E-02 | 1.10E-01 |
| ENSG00000274220 | AC009163.6 | 0.38 | 2.31E-02 | 1.10E-01 |
| ENSG00000108823 | SGCA | 0.38 | 2.31E-02 | 1.10E-01 |
| ENSG00000165283 | STOML2 | -0.38 | 2.31E-02 | 1.10E-01 |
| ENSG00000204947 | ZNF425 | 0.38 | 2.31E-02 | 1.10E-01 |
| ENSG00000243646 | IL10RB | -0.38 | 2.31E-02 | 1.10E-01 |
| ENSG00000118849 | RARRES1 | -0.38 | 2.32E-02 | 1.10E-01 |
| ENSG00000122873 | CISD1 | -0.38 | 2.32E-02 | 1.10E-01 |
| ENSG00000126247 | CAPNS1 | -0.38 | 2.32E-02 | 1.10E-01 |
| ENSG00000143222 | UFC1 | -0.38 | 2.32E-02 | 1.10E-01 |
| ENSG00000214694 | ARHGEF33 | 0.38 | 2.32E-02 | 1.10E-01 |
| ENSG00000197614 | MFAP5 | -0.38 | 2.32E-02 | 1.10E-01 |
| ENSG00000158104 | HPD | -0.38 | 2.32E-02 | 1.10E-01 |
| ENSG00000127578 | WFIKKN1 | 0.38 | 2.32E-02 | 1.10E-01 |
| ENSG00000128710 | HOXD10 | 0.38 | 2.33E-02 | 1.10E-01 |
| ENSG00000197847 | SLC22A20P | 0.38 | 2.33E-02 | 1.10E-01 |
| ENSG00000100243 | CYB5R3 | 0.38 | 2.33E-02 | 1.10E-01 |
| ENSG00000141458 | NPC1 | -0.38 | 2.33E-02 | 1.10E-01 |
| ENSG00000145979 | TBC1D7 | -0.38 | 2.33E-02 | 1.10E-01 |
| ENSG00000171097 | KYAT1 | 0.38 | 2.33E-02 | 1.10E-01 |
| ENSG00000189221 | MAOA | -0.38 | 2.33E-02 | 1.10E-01 |
| ENSG00000130208 | APOC1 | -0.38 | 2.33E-02 | 1.10E-01 |

| Ensembl ID | Gene Name | ρ | Pval | Padj |
|---|---|---|---|---|
| ENSG00000283064 | AL353759.1 | 0.38 | 2.33E-02 | 1.10E-01 |
| ENSG00000051180 | RAD51 | -0.38 | 2.33E-02 | 1.11E-01 |
| ENSG00000235618 | FAM21EP | 0.38 | 2.33E-02 | 1.11E-01 |
| ENSG00000167680 | SEMA6B | -0.38 | 2.34E-02 | 1.11E-01 |
| ENSG00000100347 | SAMM50 | -0.38 | 2.34E-02 | 1.11E-01 |
| ENSG00000105072 | C19orf44 | 0.38 | 2.34E-02 | 1.11E-01 |
| ENSG00000112640 | PPP2R5D | -0.38 | 2.34E-02 | 1.11E-01 |
| ENSG00000166819 | PLIN1 | 0.38 | 2.34E-02 | 1.11E-01 |
| ENSG00000184347 | SLIT3 | -0.38 | 2.34E-02 | 1.11E-01 |
| ENSG00000189423 | USP32P3 | 0.38 | 2.34E-02 | 1.11E-01 |
| ENSG00000197816 | CCDC180 | 0.38 | 2.34E-02 | 1.11E-01 |
| ENSG00000225791 | TRAM2-AS1 | 0.38 | 2.34E-02 | 1.11E-01 |
| ENSG00000279730 | KMT5AP1 | 0.38 | 2.34E-02 | 1.11E-01 |
| ENSG00000228434 | AC004951.1 | 0.38 | 2.34E-02 | 1.11E-01 |
| ENSG00000132639 | SNAP25 | 0.38 | 2.34E-02 | 1.11E-01 |
| ENSG00000256069 | A2MP1 | 0.38 | 2.34E-02 | 1.11E-01 |
| ENSG00000280721 | LINC01943 | -0.38 | 2.35E-02 | 1.11E-01 |
| ENSG00000104972 | LILRB1 | -0.38 | 2.35E-02 | 1.11E-01 |
| ENSG00000130749 | AC008755.1 | 0.38 | 2.35E-02 | 1.11E-01 |
| ENSG00000144028 | SNRNP200 | 0.38 | 2.35E-02 | 1.11E-01 |
| ENSG00000157680 | DGKI | -0.38 | 2.35E-02 | 1.11E-01 |
| ENSG00000158850 | B4GALT3 | -0.38 | 2.35E-02 | 1.11E-01 |
| ENSG00000237748 | UQCRBP1 | -0.38 | 2.35E-02 | 1.11E-01 |
| ENSG00000269151 | AC007193.3 | -0.38 | 2.36E-02 | 1.11E-01 |
| ENSG00000119203 | CPSF3 | -0.38 | 2.36E-02 | 1.11E-01 |
| ENSG00000123338 | NCKAP1L | -0.38 | 2.36E-02 | 1.11E-01 |
| ENSG00000131469 | RPL27 | -0.38 | 2.36E-02 | 1.11E-01 |
| ENSG00000145861 | C1QTNF2 | -0.38 | 2.36E-02 | 1.11E-01 |
| ENSG00000157800 | SLC37A3 | -0.38 | 2.36E-02 | 1.11E-01 |
| ENSG00000163605 | PPP4R2 | -0.38 | 2.36E-02 | 1.11E-01 |
| ENSG00000172197 | MBOAT1 | 0.38 | 2.36E-02 | 1.11E-01 |
| ENSG00000279192 | PWAR5 | 0.38 | 2.36E-02 | 1.11E-01 |
| ENSG00000023839 | ABCC2 | 0.38 | 2.37E-02 | 1.11E-01 |
| ENSG00000111252 | SH2B3 | -0.38 | 2.37E-02 | 1.11E-01 |
| ENSG00000123213 | NLN | -0.38 | 2.37E-02 | 1.11E-01 |
| ENSG00000134285 | FKBP11 | -0.38 | 2.37E-02 | 1.11E-01 |
| ENSG00000143436 | MRPL9 | -0.38 | 2.37E-02 | 1.11E-01 |
| ENSG00000166272 | WBP1L | -0.38 | 2.37E-02 | 1.11E-01 |
| ENSG00000168000 | BSCL2 | 0.38 | 2.37E-02 | 1.11E-01 |
| ENSG00000183458 | AC138932.1 | 0.38 | 2.37E-02 | 1.11E-01 |
| ENSG00000244945 | AC136604.2 | 0.38 | 2.37E-02 | 1.11E-01 |
| ENSG00000279672 | AP006621.5 | 0.38 | 2.37E-02 | 1.11E-01 |
| ENSG00000185904 | LINC00839 | -0.38 | 2.37E-02 | 1.11E-01 |
| ENSG00000250938 | AC108866.1 | 0.38 | 2.37E-02 | 1.11E-01 |
| ENSG00000280239 | AC011498.7 | 0.38 | 2.37E-02 | 1.11E-01 |

| Ensembl ID | Gene Name | ρ | Pval | Padj |
|---|---|---|---|---|
| ENSG00000205423 | CNEP1R1 | -0.38 | 2.38E-02 | 1.12E-01 |
| ENSG00000229852 | AC019205.1 | 0.38 | 2.38E-02 | 1.12E-01 |
| ENSG00000181444 | ZNF467 | -0.38 | 2.38E-02 | 1.12E-01 |
| ENSG00000283156 | AC068620.3 | 0.38 | 2.39E-02 | 1.12E-01 |
| ENSG00000101464 | PIGU | -0.38 | 2.40E-02 | 1.12E-01 |
| ENSG00000108424 | KPNB1 | -0.38 | 2.40E-02 | 1.12E-01 |
| ENSG00000135317 | SNX14 | -0.38 | 2.40E-02 | 1.12E-01 |
| ENSG00000165637 | VDAC2 | -0.38 | 2.40E-02 | 1.12E-01 |
| ENSG00000184867 | ARMCX2 | 0.38 | 2.40E-02 | 1.12E-01 |
| ENSG00000198515 | CNGA1 | 0.38 | 2.40E-02 | 1.12E-01 |
| ENSG00000237550 | RPL9P9 | -0.38 | 2.40E-02 | 1.12E-01 |
| ENSG00000102309 | PIN4 | -0.38 | 2.41E-02 | 1.13E-01 |
| ENSG00000115598 | IL1RL2 | 0.38 | 2.41E-02 | 1.13E-01 |
| ENSG00000143614 | GATAD2B | 0.38 | 2.41E-02 | 1.13E-01 |
| ENSG00000272674 | PCDHB16 | 0.38 | 2.41E-02 | 1.13E-01 |
| ENSG00000168004 | HRASLS5 | 0.38 | 2.41E-02 | 1.13E-01 |
| ENSG00000226026 | AC092802.1 | 0.38 | 2.42E-02 | 1.13E-01 |
| ENSG00000006740 | ARHGAP44 | -0.38 | 2.42E-02 | 1.13E-01 |
| ENSG00000120549 | KIAA1217 | -0.38 | 2.42E-02 | 1.13E-01 |
| ENSG00000134198 | TSPAN2 | -0.38 | 2.42E-02 | 1.13E-01 |
| ENSG00000136504 | KAT7 | 0.38 | 2.42E-02 | 1.13E-01 |
| ENSG00000161956 | SENP3 | -0.38 | 2.42E-02 | 1.13E-01 |
| ENSG00000198934 | MAGEE1 | 0.38 | 2.42E-02 | 1.13E-01 |
| ENSG00000203930 | LINC00632 | 0.38 | 2.42E-02 | 1.13E-01 |
| ENSG00000229807 | XIST | 0.38 | 2.42E-02 | 1.13E-01 |
| ENSG00000234420 | ZNF37BP | 0.38 | 2.42E-02 | 1.13E-01 |
| ENSG00000269044 | AC024075.2 | 0.38 | 2.42E-02 | 1.13E-01 |
| ENSG00000280400 | AC127459.3 | -0.38 | 2.43E-02 | 1.13E-01 |
| ENSG00000165194 | PCDH19 | 0.38 | 2.43E-02 | 1.13E-01 |
| ENSG00000131196 | NFATC1 | 0.38 | 2.43E-02 | 1.13E-01 |
| ENSG00000167232 | ZNF91 | 0.38 | 2.43E-02 | 1.13E-01 |
| ENSG00000204469 | PRRC2A | 0.38 | 2.43E-02 | 1.13E-01 |
| ENSG00000261971 | MMP25-AS1 | 0.38 | 2.43E-02 | 1.13E-01 |
| ENSG00000273090 | AC007378.1 | 0.38 | 2.43E-02 | 1.13E-01 |
| ENSG00000134775 | FHOD3 | -0.38 | 2.44E-02 | 1.13E-01 |
| ENSG00000228791 | THRB-AS1 | 0.38 | 2.44E-02 | 1.13E-01 |
| ENSG00000090530 | P3H2 | -0.38 | 2.44E-02 | 1.13E-01 |
| ENSG00000111237 | VPS29 | -0.38 | 2.44E-02 | 1.13E-01 |
| ENSG00000130723 | PRRC2B | 0.38 | 2.44E-02 | 1.13E-01 |
| ENSG00000146350 | TBC1D32 | 0.38 | 2.44E-02 | 1.13E-01 |
| ENSG00000152818 | UTRN | 0.38 | 2.44E-02 | 1.13E-01 |
| ENSG00000204406 | MBD5 | 0.38 | 2.44E-02 | 1.13E-01 |
| ENSG00000213171 | LINGO4 | 0.38 | 2.44E-02 | 1.13E-01 |
| ENSG00000261441 | AC124068.2 | 0.38 | 2.44E-02 | 1.13E-01 |
| ENSG00000242866 | STRC | 0.38 | 2.45E-02 | 1.14E-01 |

| Ensembl ID | Gene Name | ρ | Pval | Padj |
|---|---|---|---|---|
| ENSG00000025772 | TOMM34 | -0.38 | 2.45E-02 | 1.14E-01 |
| ENSG00000059122 | FLYWCH1 | 0.38 | 2.45E-02 | 1.14E-01 |
| ENSG00000138615 | CILP | 0.38 | 2.45E-02 | 1.14E-01 |
| ENSG00000224550 | AC114491.1 | -0.38 | 2.45E-02 | 1.14E-01 |
| ENSG00000164283 | ESM1 | -0.38 | 2.46E-02 | 1.14E-01 |
| ENSG00000250471 | GMPSP1 | -0.38 | 2.46E-02 | 1.14E-01 |
| ENSG00000013306 | SLC25A39 | -0.38 | 2.46E-02 | 1.14E-01 |
| ENSG00000095066 | HOOK2 | 0.38 | 2.46E-02 | 1.14E-01 |
| ENSG00000124813 | RUNX2 | -0.38 | 2.46E-02 | 1.14E-01 |
| ENSG00000188338 | SLC38A3 | 0.38 | 2.46E-02 | 1.14E-01 |
| ENSG00000197771 | MCMBP | -0.38 | 2.46E-02 | 1.14E-01 |
| ENSG00000218336 | TENM3 | -0.38 | 2.46E-02 | 1.14E-01 |
| ENSG00000280047 | AC091825.1 | 0.38 | 2.46E-02 | 1.14E-01 |
| ENSG00000166896 | ATP23 | -0.38 | 2.48E-02 | 1.14E-01 |
| ENSG00000173947 | PIFO | -0.38 | 2.48E-02 | 1.14E-01 |
| ENSG00000178177 | LCORL | -0.38 | 2.48E-02 | 1.14E-01 |
| ENSG00000186017 | ZNF566 | 0.38 | 2.48E-02 | 1.14E-01 |
| ENSG00000205485 | AC004980.1 | 0.38 | 2.48E-02 | 1.14E-01 |
| ENSG00000213977 | TAX1BP3 | -0.38 | 2.48E-02 | 1.14E-01 |
| ENSG00000102981 | PARD6A | -0.38 | 2.48E-02 | 1.15E-01 |
| ENSG00000270641 | TSIX | 0.38 | 2.49E-02 | 1.15E-01 |
| ENSG00000169220 | RGS14 | -0.38 | 2.49E-02 | 1.15E-01 |
| ENSG00000187866 | FAM122A | -0.38 | 2.49E-02 | 1.15E-01 |
| ENSG00000196150 | ZNF250 | 0.38 | 2.49E-02 | 1.15E-01 |
| ENSG00000043514 | TRIT1 | -0.38 | 2.50E-02 | 1.15E-01 |
| ENSG00000104881 | PPP1R13L | 0.38 | 2.50E-02 | 1.15E-01 |
| ENSG00000156966 | B3GNT7 | 0.38 | 2.50E-02 | 1.15E-01 |
| ENSG00000187187 | ZNF546 | 0.38 | 2.50E-02 | 1.15E-01 |
| ENSG00000253305 | PCDHGB6 | 0.38 | 2.50E-02 | 1.15E-01 |
| ENSG00000105357 | MYH14 | 0.38 | 2.51E-02 | 1.15E-01 |
| ENSG00000112877 | CEP72 | 0.38 | 2.51E-02 | 1.15E-01 |
| ENSG00000125166 | GOT2 | -0.38 | 2.51E-02 | 1.15E-01 |
| ENSG00000136167 | LCP1 | -0.38 | 2.51E-02 | 1.15E-01 |
| ENSG00000136754 | ABI1 | -0.38 | 2.51E-02 | 1.15E-01 |
| ENSG00000139668 | WDFY2 | 0.38 | 2.51E-02 | 1.15E-01 |
| ENSG00000143493 | INTS7 | -0.38 | 2.51E-02 | 1.15E-01 |
| ENSG00000166333 | ILK | -0.38 | 2.51E-02 | 1.15E-01 |
| ENSG00000181222 | POLR2A | 0.38 | 2.51E-02 | 1.15E-01 |
| ENSG00000197748 | CFAP43 | 0.38 | 2.51E-02 | 1.15E-01 |
| ENSG00000204314 | PRRT1 | 0.38 | 2.51E-02 | 1.15E-01 |
| ENSG00000223865 | HLA-DPB1 | -0.38 | 2.51E-02 | 1.15E-01 |
| ENSG00000224281 | SLC25A5-AS1 | 0.38 | 2.51E-02 | 1.15E-01 |
| ENSG00000197815 | AC122129.1 | 0.38 | 2.51E-02 | 1.15E-01 |
| ENSG00000278588 | HIST1H2BI | -0.38 | 2.52E-02 | 1.16E-01 |
| ENSG00000135951 | TSGA10 | 0.38 | 2.52E-02 | 1.16E-01 |

| Ensembl ID | Gene Name | ρ | Pval | Padj |
|---|---|---|---|---|
| ENSG00000138795 | LEF1 | -0.38 | 2.52E-02 | 1.16E-01 |
| ENSG00000143630 | HCN3 | 0.38 | 2.52E-02 | 1.16E-01 |
| ENSG00000282508 | LINC01002 | 0.38 | 2.52E-02 | 1.16E-01 |
| ENSG00000248873 | SERBP1P6 | -0.38 | 2.53E-02 | 1.16E-01 |
| ENSG00000181458 | TMEM45A | -0.38 | 2.53E-02 | 1.16E-01 |
| ENSG00000181754 | AMIGO1 | 0.38 | 2.53E-02 | 1.16E-01 |
| ENSG00000253200 | AC037459.3 | 0.38 | 2.53E-02 | 1.16E-01 |
| ENSG00000065883 | CDK13 | 0.38 | 2.55E-02 | 1.17E-01 |
| ENSG00000104946 | TBC1D17 | 0.38 | 2.55E-02 | 1.17E-01 |
| ENSG00000106591 | MRPL32 | -0.38 | 2.55E-02 | 1.17E-01 |
| ENSG00000127445 | PIN1 | -0.38 | 2.55E-02 | 1.17E-01 |
| ENSG00000128731 | HERC2 | 0.38 | 2.55E-02 | 1.17E-01 |
| ENSG00000159399 | HK2 | 0.38 | 2.55E-02 | 1.17E-01 |
| ENSG00000225178 | RPSAP58 | -0.38 | 2.55E-02 | 1.17E-01 |
| ENSG00000234797 | RPS3AP6 | -0.38 | 2.55E-02 | 1.17E-01 |
| ENSG00000136160 | EDNRB | -0.38 | 2.55E-02 | 1.17E-01 |
| ENSG00000224992 | AL445645.1 | 0.38 | 2.55E-02 | 1.17E-01 |
| ENSG00000145850 | TIMD4 | -0.38 | 2.55E-02 | 1.17E-01 |
| ENSG00000051341 | POLQ | -0.38 | 2.55E-02 | 1.17E-01 |
| ENSG00000154493 | C10orf90 | -0.38 | 2.55E-02 | 1.17E-01 |
| ENSG00000100292 | HMOX1 | -0.38 | 2.56E-02 | 1.17E-01 |
| ENSG00000100600 | LGMN | -0.38 | 2.56E-02 | 1.17E-01 |
| ENSG00000115756 | HPCAL1 | -0.38 | 2.56E-02 | 1.17E-01 |
| ENSG00000148950 | IMMP1L | -0.38 | 2.56E-02 | 1.17E-01 |
| ENSG00000163382 | NAXE | -0.38 | 2.56E-02 | 1.17E-01 |
| ENSG00000167522 | ANKRD11 | 0.38 | 2.56E-02 | 1.17E-01 |
| ENSG00000204634 | TBC1D8 | 0.38 | 2.56E-02 | 1.17E-01 |
| ENSG00000119042 | SATB2 | -0.38 | 2.57E-02 | 1.17E-01 |
| ENSG00000122033 | MTIF3 | -0.38 | 2.57E-02 | 1.17E-01 |
| ENSG00000159210 | SNF8 | -0.38 | 2.57E-02 | 1.17E-01 |
| ENSG00000173153 | ESRRA | -0.38 | 2.57E-02 | 1.17E-01 |
| ENSG00000271895 | AL109811.2 | 0.38 | 2.57E-02 | 1.17E-01 |
| ENSG00000229142 | HCG4P8 | 0.38 | 2.57E-02 | 1.17E-01 |
| ENSG00000254815 | AP006284.1 | 0.38 | 2.57E-02 | 1.17E-01 |
| ENSG00000196388 | INCA1 | 0.38 | 2.58E-02 | 1.17E-01 |
| ENSG00000101230 | ISM1 | 0.38 | 2.58E-02 | 1.18E-01 |
| ENSG00000115520 | COQ10B | -0.38 | 2.58E-02 | 1.18E-01 |
| ENSG00000125247 | TMTC4 | -0.38 | 2.58E-02 | 1.18E-01 |
| ENSG00000149269 | PAK1 | -0.38 | 2.58E-02 | 1.18E-01 |
| ENSG00000206417 | H1FX-AS1 | 0.38 | 2.58E-02 | 1.18E-01 |
| ENSG00000248485 | PCP4L1 | -0.38 | 2.58E-02 | 1.18E-01 |
| ENSG00000273289 | AL121672.3 | 0.38 | 2.58E-02 | 1.18E-01 |
| ENSG00000254452 | AP001107.2 | -0.38 | 2.58E-02 | 1.18E-01 |
| ENSG00000230729 | AL160270.1 | 0.38 | 2.59E-02 | 1.18E-01 |
| ENSG00000066336 | SPI1 | -0.38 | 2.59E-02 | 1.18E-01 |

| Ensembl ID | Gene Name | ρ | Pval | Padj |
|---|---|---|---|---|
| ENSG00000081181 | ARG2 | -0.38 | 2.59E-02 | 1.18E-01 |
| ENSG00000163347 | CLDN1 | -0.38 | 2.59E-02 | 1.18E-01 |
| ENSG00000274177 | AC004528.2 | -0.38 | 2.60E-02 | 1.18E-01 |
| ENSG00000270392 | PFN1P2 | 0.38 | 2.60E-02 | 1.18E-01 |
| ENSG00000186431 | FCAR | -0.38 | 2.60E-02 | 1.18E-01 |
| ENSG00000119147 | C2orf40 | 0.38 | 2.60E-02 | 1.18E-01 |
| ENSG00000120875 | DUSP4 | -0.38 | 2.60E-02 | 1.18E-01 |
| ENSG00000124733 | MEA1 | -0.38 | 2.60E-02 | 1.18E-01 |
| ENSG00000125952 | MAX | -0.38 | 2.60E-02 | 1.18E-01 |
| ENSG00000139180 | NDUFA9 | -0.38 | 2.60E-02 | 1.18E-01 |
| ENSG00000140451 | PIF1 | -0.38 | 2.60E-02 | 1.18E-01 |
| ENSG00000153975 | ZUP1 | -0.38 | 2.60E-02 | 1.18E-01 |
| ENSG00000170677 | SOCS6 | -0.38 | 2.60E-02 | 1.18E-01 |
| ENSG00000183891 | TTC32 | 0.38 | 2.60E-02 | 1.18E-01 |
| ENSG00000185483 | ROR1 | -0.38 | 2.60E-02 | 1.18E-01 |
| ENSG00000213593 | TMX2 | -0.38 | 2.60E-02 | 1.18E-01 |
| ENSG00000257621 | PSMA3-AS1 | 0.38 | 2.60E-02 | 1.18E-01 |
| ENSG00000258890 | CEP95 | 0.38 | 2.60E-02 | 1.18E-01 |
| ENSG00000266302 | AC098850.3 | 0.38 | 2.61E-02 | 1.18E-01 |
| ENSG00000124003 | MOGAT1 | 0.38 | 2.61E-02 | 1.18E-01 |
| ENSG00000127084 | FGD3 | -0.38 | 2.61E-02 | 1.18E-01 |
| ENSG00000277003 | AC026468.1 | 0.38 | 2.61E-02 | 1.18E-01 |
| ENSG00000127074 | RGS13 | -0.38 | 2.61E-02 | 1.18E-01 |
| ENSG00000068438 | FTSJ1 | -0.38 | 2.62E-02 | 1.18E-01 |
| ENSG00000080819 | CPOX | -0.38 | 2.62E-02 | 1.18E-01 |
| ENSG00000089157 | RPLP0 | -0.38 | 2.62E-02 | 1.18E-01 |
| ENSG00000092470 | WDR76 | -0.38 | 2.62E-02 | 1.18E-01 |
| ENSG00000151006 | PRSS53 | 0.38 | 2.62E-02 | 1.18E-01 |
| ENSG00000162654 | GBP4 | 0.38 | 2.62E-02 | 1.18E-01 |
| ENSG00000185219 | ZNF445 | 0.38 | 2.62E-02 | 1.18E-01 |
| ENSG00000189339 | SLC35E2B | 0.38 | 2.62E-02 | 1.18E-01 |
| ENSG00000198947 | DMD | -0.38 | 2.62E-02 | 1.18E-01 |
| ENSG00000260740 | AC026471.3 | 0.38 | 2.62E-02 | 1.18E-01 |
| ENSG00000225912 | AL121871.1 | -0.38 | 2.62E-02 | 1.18E-01 |
| ENSG00000119608 | PROX2 | 0.38 | 2.63E-02 | 1.19E-01 |
| ENSG00000002016 | RAD52 | 0.38 | 2.63E-02 | 1.19E-01 |
| ENSG00000081913 | PHLPP1 | 0.38 | 2.63E-02 | 1.19E-01 |
| ENSG00000119782 | FKBP1B | -0.38 | 2.63E-02 | 1.19E-01 |
| ENSG00000147394 | ZNF185 | 0.38 | 2.63E-02 | 1.19E-01 |
| ENSG00000253144 | AC104117.2 | -0.38 | 2.63E-02 | 1.19E-01 |
| ENSG00000076513 | ANKRD13A | -0.38 | 2.64E-02 | 1.19E-01 |
| ENSG00000090061 | CCNK | 0.38 | 2.64E-02 | 1.19E-01 |
| ENSG00000135048 | CEMIP2 | 0.38 | 2.64E-02 | 1.19E-01 |
| ENSG00000143545 | RAB13 | -0.38 | 2.64E-02 | 1.19E-01 |
| ENSG00000144645 | OSBPL10 | 0.38 | 2.64E-02 | 1.19E-01 |

| Ensembl ID | Gene Name | ρ | Pval | Padj |
|---|---|---|---|---|
| ENSG00000179776 | CDH5 | -0.38 | 2.64E-02 | 1.19E-01 |
| ENSG00000107317 | PTGDS | -0.37 | 2.65E-02 | 1.19E-01 |
| ENSG00000112276 | BVES | 0.37 | 2.65E-02 | 1.19E-01 |
| ENSG00000151502 | VPS26B | 0.37 | 2.65E-02 | 1.19E-01 |
| ENSG00000173638 | SLC19A1 | -0.37 | 2.65E-02 | 1.19E-01 |
| ENSG00000196586 | MYO6 | -0.37 | 2.65E-02 | 1.19E-01 |
| ENSG00000228748 | AL450306.1 | -0.37 | 2.66E-02 | 1.19E-01 |
| ENSG00000205622 | AP001043.1 | 0.37 | 2.66E-02 | 1.19E-01 |
| ENSG00000130948 | HSD17B3 | 0.37 | 2.66E-02 | 1.19E-01 |
| ENSG00000283633 | AP000547.3 | -0.37 | 2.66E-02 | 1.19E-01 |
| ENSG00000072506 | HSD17B10 | -0.37 | 2.67E-02 | 1.19E-01 |
| ENSG00000091136 | LAMB1 | -0.37 | 2.67E-02 | 1.19E-01 |
| ENSG00000145220 | LYAR | -0.37 | 2.67E-02 | 1.19E-01 |
| ENSG00000168763 | CNNM3 | 0.37 | 2.67E-02 | 1.19E-01 |
| ENSG00000171084 | FAM86JP | 0.37 | 2.67E-02 | 1.19E-01 |
| ENSG00000188725 | SMIM15 | -0.37 | 2.67E-02 | 1.19E-01 |
| ENSG00000215417 | MIR17HG | -0.37 | 2.67E-02 | 1.19E-01 |
| ENSG00000225828 | FAM229A | 0.37 | 2.67E-02 | 1.19E-01 |
| ENSG00000280195 | AC245140.2 | 0.37 | 2.67E-02 | 1.19E-01 |
| ENSG00000041353 | RAB27B | -0.37 | 2.67E-02 | 1.19E-01 |
| ENSG00000121797 | CCRL2 | -0.37 | 2.67E-02 | 1.19E-01 |
| ENSG00000244682 | FCGR2C | -0.37 | 2.67E-02 | 1.20E-01 |
| ENSG00000241749 | RPSAP52 | -0.37 | 2.67E-02 | 1.20E-01 |
| ENSG00000063854 | HAGH | -0.37 | 2.68E-02 | 1.20E-01 |
| ENSG00000069188 | SDK2 | 0.37 | 2.68E-02 | 1.20E-01 |
| ENSG00000154144 | TBRG1 | -0.37 | 2.68E-02 | 1.20E-01 |
| ENSG00000156113 | KCNMA1 | 0.37 | 2.68E-02 | 1.20E-01 |
| ENSG00000156603 | MED19 | -0.37 | 2.68E-02 | 1.20E-01 |
| ENSG00000171603 | CLSTN1 | 0.37 | 2.68E-02 | 1.20E-01 |
| ENSG00000185339 | TCN2 | -0.37 | 2.68E-02 | 1.20E-01 |
| ENSG00000237522 | NONOP2 | 0.37 | 2.68E-02 | 1.20E-01 |
| ENSG00000173083 | HPSE | -0.37 | 2.69E-02 | 1.20E-01 |
| ENSG00000120659 | TNFSF11 | -0.37 | 2.69E-02 | 1.20E-01 |
| ENSG00000125703 | ATG4C | -0.37 | 2.69E-02 | 1.20E-01 |
| ENSG00000136630 | HLX | -0.37 | 2.69E-02 | 1.20E-01 |
| ENSG00000151929 | BAG3 | 0.37 | 2.69E-02 | 1.20E-01 |
| ENSG00000174744 | BRMS1 | -0.37 | 2.69E-02 | 1.20E-01 |
| ENSG00000175193 | PARL | -0.37 | 2.69E-02 | 1.20E-01 |
| ENSG00000197283 | SYNGAP1 | 0.37 | 2.69E-02 | 1.20E-01 |
| ENSG00000226243 | RPL37AP1 | -0.37 | 2.69E-02 | 1.20E-01 |
| ENSG00000281357 | ARRDC3-AS1 | 0.37 | 2.70E-02 | 1.20E-01 |
| ENSG00000126883 | NUP214 | 0.37 | 2.70E-02 | 1.20E-01 |
| ENSG00000145907 | G3BP1 | -0.37 | 2.70E-02 | 1.20E-01 |
| ENSG00000163319 | MRPS18C | -0.37 | 2.70E-02 | 1.20E-01 |
| ENSG00000185909 | KLHDC8B | -0.37 | 2.70E-02 | 1.20E-01 |

| Ensembl ID | Gene Name | ρ | Pval | Padj |
|---|---|---|---|---|
| ENSG00000242588 | AC108010.1 | 0.37 | 2.70E-02 | 1.21E-01 |
| ENSG00000166183 | ASPG | 0.37 | 2.70E-02 | 1.21E-01 |
| ENSG00000084070 | SMAP2 | -0.37 | 2.71E-02 | 1.21E-01 |
| ENSG00000112893 | MAN2A1 | -0.37 | 2.71E-02 | 1.21E-01 |
| ENSG00000122691 | TWIST1 | -0.37 | 2.71E-02 | 1.21E-01 |
| ENSG00000134602 | STK26 | 0.37 | 2.71E-02 | 1.21E-01 |
| ENSG00000135766 | EGLN1 | 0.37 | 2.71E-02 | 1.21E-01 |
| ENSG00000173068 | BNC2 | -0.37 | 2.71E-02 | 1.21E-01 |
| ENSG00000185201 | IFITM2 | -0.37 | 2.71E-02 | 1.21E-01 |
| ENSG00000185340 | GAS2L1 | -0.37 | 2.71E-02 | 1.21E-01 |
| ENSG00000239556 | AC004951.2 | 0.37 | 2.72E-02 | 1.21E-01 |
| ENSG00000251348 | HSPD1P11 | 0.37 | 2.72E-02 | 1.21E-01 |
| ENSG00000122122 | SASH3 | -0.37 | 2.72E-02 | 1.21E-01 |
| ENSG00000057019 | DCBLD2 | -0.37 | 2.73E-02 | 1.21E-01 |
| ENSG00000148154 | UGCG | -0.37 | 2.73E-02 | 1.21E-01 |
| ENSG00000155100 | OTUD6B | -0.37 | 2.73E-02 | 1.21E-01 |
| ENSG00000196873 | CBWD3 | 0.37 | 2.73E-02 | 1.21E-01 |
| ENSG00000276278 | AC048382.6 | 0.37 | 2.73E-02 | 1.21E-01 |
| ENSG00000126882 | FAM78A | -0.37 | 2.73E-02 | 1.21E-01 |
| ENSG00000034239 | EFCAB1 | 0.37 | 2.74E-02 | 1.21E-01 |
| ENSG00000119686 | FLVCR2 | 0.37 | 2.74E-02 | 1.21E-01 |
| ENSG00000124333 | VAMP7 | -0.37 | 2.74E-02 | 1.21E-01 |
| ENSG00000126756 | UXT | -0.37 | 2.74E-02 | 1.21E-01 |
| ENSG00000177084 | POLE | 0.37 | 2.74E-02 | 1.21E-01 |
| ENSG00000198185 | ZNF334 | 0.37 | 2.74E-02 | 1.21E-01 |
| ENSG00000254481 | PTP4A2P2 | -0.37 | 2.74E-02 | 1.22E-01 |
| ENSG00000186854 | TRABD2A | -0.37 | 2.75E-02 | 1.22E-01 |
| ENSG00000186377 | CYP4X1 | 0.37 | 2.75E-02 | 1.22E-01 |
| ENSG00000134046 | MBD2 | -0.37 | 2.75E-02 | 1.22E-01 |
| ENSG00000154222 | CC2D1B | 0.37 | 2.75E-02 | 1.22E-01 |
| ENSG00000197181 | PIWIL2 | 0.37 | 2.75E-02 | 1.22E-01 |
| ENSG00000204977 | TRIM13 | -0.37 | 2.75E-02 | 1.22E-01 |
| ENSG00000237298 | TTN-AS1 | 0.37 | 2.75E-02 | 1.22E-01 |
| ENSG00000268941 | LINC01711 | -0.37 | 2.75E-02 | 1.22E-01 |
| ENSG00000182866 | LCK | -0.37 | 2.75E-02 | 1.22E-01 |
| ENSG00000236675 | MTX1P1 | -0.37 | 2.75E-02 | 1.22E-01 |
| ENSG00000232814 | COL4A2-AS1 | -0.37 | 2.76E-02 | 1.22E-01 |
| ENSG00000106397 | PLOD3 | -0.37 | 2.76E-02 | 1.22E-01 |
| ENSG00000119537 | KDSR | -0.37 | 2.76E-02 | 1.22E-01 |
| ENSG00000141543 | EIF4A3 | -0.37 | 2.76E-02 | 1.22E-01 |
| ENSG00000167632 | TRAPPC9 | 0.37 | 2.76E-02 | 1.22E-01 |
| ENSG00000215712 | TMEM242 | -0.37 | 2.76E-02 | 1.22E-01 |
| ENSG00000226564 | FTH1P20 | -0.37 | 2.76E-02 | 1.22E-01 |
| ENSG00000245532 | NEAT1 | 0.37 | 2.76E-02 | 1.22E-01 |
| ENSG00000231120 | BTF3P10 | -0.37 | 2.77E-02 | 1.22E-01 |

| Ensembl ID | Gene Name | ρ | Pval | Padj |
|---|---|---|---|---|
| ENSG00000151062 | CACNA2D4 | -0.37 | 2.78E-02 | 1.22E-01 |
| ENSG00000165606 | DRGX | -0.37 | 2.78E-02 | 1.22E-01 |
| ENSG00000137106 | GRHPR | -0.37 | 2.78E-02 | 1.22E-01 |
| ENSG00000147457 | CHMP7 | -0.37 | 2.78E-02 | 1.22E-01 |
| ENSG00000159110 | IFNAR2 | -0.37 | 2.78E-02 | 1.22E-01 |
| ENSG00000162298 | SYVN1 | 0.37 | 2.78E-02 | 1.22E-01 |
| ENSG00000162894 | FCMR | 0.37 | 2.78E-02 | 1.22E-01 |
| ENSG00000171130 | ATP6V0E2 | -0.37 | 2.78E-02 | 1.22E-01 |
| ENSG00000172339 | ALG14 | -0.37 | 2.78E-02 | 1.22E-01 |
| ENSG00000280063 | AC012676.5 | 0.37 | 2.78E-02 | 1.22E-01 |
| ENSG00000235919 | ASH1L-AS1 | 0.37 | 2.78E-02 | 1.23E-01 |
| ENSG00000271933 | AL603756.1 | 0.37 | 2.79E-02 | 1.23E-01 |
| ENSG00000118096 | IFT46 | -0.37 | 2.79E-02 | 1.23E-01 |
| ENSG00000138758 | SEPT11 | -0.37 | 2.79E-02 | 1.23E-01 |
| ENSG00000184831 | APOO | -0.37 | 2.79E-02 | 1.23E-01 |
| ENSG00000237676 | RPL30P4 | -0.37 | 2.79E-02 | 1.23E-01 |
| ENSG00000243449 | C4orf48 | -0.37 | 2.79E-02 | 1.23E-01 |
| ENSG00000256448 | AP000763.3 | 0.37 | 2.79E-02 | 1.23E-01 |
| ENSG00000276368 | HIST1H2AJ | -0.37 | 2.80E-02 | 1.23E-01 |
| ENSG00000135318 | NT5E | -0.37 | 2.80E-02 | 1.23E-01 |
| ENSG00000197943 | PLCG2 | 0.37 | 2.80E-02 | 1.23E-01 |
| ENSG00000224594 | RPL29P19 | 0.37 | 2.80E-02 | 1.23E-01 |
| ENSG00000230550 | ERLNC1 | 0.37 | 2.81E-02 | 1.24E-01 |
| ENSG00000166912 | MTMR10 | 0.37 | 2.82E-02 | 1.24E-01 |
| ENSG00000168938 | PPIC | -0.37 | 2.82E-02 | 1.24E-01 |
| ENSG00000185298 | CCDC137 | -0.37 | 2.82E-02 | 1.24E-01 |
| ENSG00000231952 | DPY19L1P2 | -0.37 | 2.82E-02 | 1.24E-01 |
| ENSG00000260633 | AC010207.1 | 0.37 | 2.83E-02 | 1.24E-01 |
| ENSG00000128626 | MRPS12 | -0.37 | 2.83E-02 | 1.24E-01 |
| ENSG00000142347 | MYO1F | -0.37 | 2.83E-02 | 1.24E-01 |
| ENSG00000152082 | MZT2B | -0.37 | 2.83E-02 | 1.24E-01 |
| ENSG00000175606 | TMEM70 | -0.37 | 2.83E-02 | 1.24E-01 |
| ENSG00000185798 | WDR53 | -0.37 | 2.83E-02 | 1.24E-01 |
| ENSG00000196456 | ZNF775 | 0.37 | 2.83E-02 | 1.24E-01 |
| ENSG00000233682 | AL356417.2 | -0.37 | 2.83E-02 | 1.24E-01 |
| ENSG00000095596 | CYP26A1 | -0.37 | 2.83E-02 | 1.24E-01 |
| ENSG00000275708 | MIR3648-1 | 0.37 | 2.83E-02 | 1.24E-01 |
| ENSG00000273300 | AC000068.3 | 0.37 | 2.83E-02 | 1.24E-01 |
| ENSG00000148204 | CRB2 | 0.37 | 2.84E-02 | 1.24E-01 |
| ENSG00000272858 | Z93930.3 | 0.37 | 2.84E-02 | 1.24E-01 |
| ENSG00000159905 | AC006213.1 | 0.37 | 2.84E-02 | 1.24E-01 |
| ENSG00000159128 | IFNGR2 | -0.37 | 2.84E-02 | 1.24E-01 |
| ENSG00000162736 | NCSTN | -0.37 | 2.84E-02 | 1.24E-01 |
| ENSG00000187554 | TLR5 | -0.37 | 2.84E-02 | 1.24E-01 |
| ENSG00000196923 | PDLIM7 | -0.37 | 2.84E-02 | 1.24E-01 |

| Ensembl ID | Gene Name | ρ | Pval | Padj |
|---|---|---|---|---|
| ENSG00000231064 | AC234582.1 | 0.37 | 2.84E-02 | 1.24E-01 |
| ENSG00000160472 | TMEM190 | -0.37 | 2.84E-02 | 1.24E-01 |
| ENSG00000274026 | FAM27E3 | 0.37 | 2.85E-02 | 1.24E-01 |
| ENSG00000279080 | AL022322.2 | 0.37 | 2.85E-02 | 1.24E-01 |
| ENSG00000154654 | NCAM2 | -0.37 | 2.85E-02 | 1.24E-01 |
| ENSG00000183662 | FAM19A1 | 0.37 | 2.85E-02 | 1.24E-01 |
| ENSG00000165071 | TMEM71 | -0.37 | 2.85E-02 | 1.24E-01 |
| ENSG00000146063 | TRIM41 | 0.37 | 2.85E-02 | 1.24E-01 |
| ENSG00000149480 | MTA2 | -0.37 | 2.85E-02 | 1.24E-01 |
| ENSG00000166246 | C16orf71 | 0.37 | 2.85E-02 | 1.24E-01 |
| ENSG00000169193 | CCDC126 | -0.37 | 2.85E-02 | 1.24E-01 |
| ENSG00000167550 | RHEBL1 | -0.37 | 2.86E-02 | 1.24E-01 |
| ENSG00000204837 | FGF7P3 | 0.37 | 2.86E-02 | 1.24E-01 |
| ENSG00000225342 | AC079630.1 | 0.37 | 2.86E-02 | 1.24E-01 |
| ENSG00000117009 | KMO | -0.37 | 2.86E-02 | 1.25E-01 |
| ENSG00000054523 | KIF1B | -0.37 | 2.87E-02 | 1.25E-01 |
| ENSG00000140612 | SEC11A | -0.37 | 2.87E-02 | 1.25E-01 |
| ENSG00000163590 | PPM1L | -0.37 | 2.87E-02 | 1.25E-01 |
| ENSG00000272655 | POLR2J4 | 0.37 | 2.87E-02 | 1.25E-01 |
| ENSG00000273344 | PAXIP1-AS1 | 0.37 | 2.87E-02 | 1.25E-01 |
| ENSG00000101938 | CHRDL1 | -0.37 | 2.87E-02 | 1.25E-01 |
| ENSG00000267457 | AC004223.2 | 0.37 | 2.87E-02 | 1.25E-01 |
| ENSG00000106031 | HOXA13 | 0.37 | 2.88E-02 | 1.25E-01 |
| ENSG00000111275 | ALDH2 | 0.37 | 2.88E-02 | 1.25E-01 |
| ENSG00000115828 | QPCT | -0.37 | 2.88E-02 | 1.25E-01 |
| ENSG00000122965 | RBM19 | 0.37 | 2.88E-02 | 1.25E-01 |
| ENSG00000143183 | TMCO1 | -0.37 | 2.88E-02 | 1.25E-01 |
| ENSG00000162496 | DHRS3 | 0.37 | 2.88E-02 | 1.25E-01 |
| ENSG00000172799 | ZBTB8OSP2 | -0.37 | 2.89E-02 | 1.25E-01 |
| ENSG00000130005 | GAMT | -0.37 | 2.89E-02 | 1.26E-01 |
| ENSG00000149380 | P4HA3 | -0.37 | 2.89E-02 | 1.26E-01 |
| ENSG00000167595 | PROSER3 | 0.37 | 2.89E-02 | 1.26E-01 |
| ENSG00000225978 | HAR1A | 0.37 | 2.89E-02 | 1.26E-01 |
| ENSG00000230844 | ZNF674-AS1 | 0.37 | 2.89E-02 | 1.26E-01 |
| ENSG00000272156 | AC008280.3 | 0.37 | 2.89E-02 | 1.26E-01 |
| ENSG00000265618 | AC002094.2 | -0.37 | 2.90E-02 | 1.26E-01 |
| ENSG00000225770 | RPS29P3 | -0.37 | 2.90E-02 | 1.26E-01 |
| ENSG00000100399 | CHADL | 0.37 | 2.91E-02 | 1.26E-01 |
| ENSG00000140471 | LINS1 | -0.37 | 2.91E-02 | 1.26E-01 |
| ENSG00000152580 | IGSF10 | -0.37 | 2.91E-02 | 1.26E-01 |
| ENSG00000171204 | TMEM126B | -0.37 | 2.91E-02 | 1.26E-01 |
| ENSG00000237493 | AC034102.1 | -0.37 | 2.91E-02 | 1.26E-01 |
| ENSG00000127418 | FGFRL1 | 0.37 | 2.92E-02 | 1.26E-01 |
| ENSG00000128604 | IRF5 | -0.37 | 2.92E-02 | 1.26E-01 |
| ENSG00000135521 | LTV1 | -0.37 | 2.92E-02 | 1.26E-01 |

| Ensembl ID | Gene Name | ρ | Pval | Padj |
|---|---|---|---|---|
| ENSG00000162390 | ACOT11 | 0.37 | 2.92E-02 | 1.26E-01 |
| ENSG00000243649 | CFB | 0.37 | 2.92E-02 | 1.26E-01 |
| ENSG00000276070 | CCL4L2 | -0.37 | 2.93E-02 | 1.27E-01 |
| ENSG00000272610 | MAGI1-IT1 | 0.37 | 2.93E-02 | 1.27E-01 |
| ENSG00000255561 | FDXACB1 | -0.37 | 2.93E-02 | 1.27E-01 |
| ENSG00000013561 | RNF14 | -0.37 | 2.93E-02 | 1.27E-01 |
| ENSG00000103264 | FBXO31 | 0.37 | 2.93E-02 | 1.27E-01 |
| ENSG00000123124 | WWP1 | -0.37 | 2.93E-02 | 1.27E-01 |
| ENSG00000166135 | HIF1AN | 0.37 | 2.93E-02 | 1.27E-01 |
| ENSG00000198055 | GRK6 | -0.37 | 2.93E-02 | 1.27E-01 |
| ENSG00000259431 | THTPA | 0.37 | 2.93E-02 | 1.27E-01 |
| ENSG00000040275 | SPDL1 | -0.37 | 2.95E-02 | 1.27E-01 |
| ENSG00000127947 | PTPN12 | -0.37 | 2.95E-02 | 1.27E-01 |
| ENSG00000181513 | ACBD4 | 0.37 | 2.95E-02 | 1.27E-01 |
| ENSG00000183431 | SF3A3 | -0.37 | 2.95E-02 | 1.27E-01 |
| ENSG00000264522 | OTUD7B | 0.37 | 2.95E-02 | 1.27E-01 |
| ENSG00000258985 | AL352979.2 | -0.37 | 2.95E-02 | 1.27E-01 |
| ENSG00000224822 | THRB-IT1 | 0.37 | 2.95E-02 | 1.27E-01 |
| ENSG00000128242 | GAL3ST1 | 0.37 | 2.95E-02 | 1.27E-01 |
| ENSG00000233382 | NKAPP1 | 0.37 | 2.96E-02 | 1.27E-01 |
| ENSG00000065615 | CYB5R4 | -0.37 | 2.96E-02 | 1.27E-01 |
| ENSG00000102554 | KLF5 | 0.37 | 2.96E-02 | 1.27E-01 |
| ENSG00000172006 | ZNF554 | 0.37 | 2.96E-02 | 1.27E-01 |
| ENSG00000225663 | MCRIP1 | -0.37 | 2.96E-02 | 1.27E-01 |
| ENSG00000232611 | AL683813.1 | 0.37 | 2.96E-02 | 1.27E-01 |
| ENSG00000244687 | UBE2V1 | 0.37 | 2.96E-02 | 1.27E-01 |
| ENSG00000269001 | AC092070.2 | 0.37 | 2.96E-02 | 1.27E-01 |
| ENSG00000101342 | TLDC2 | -0.37 | 2.96E-02 | 1.27E-01 |
| ENSG00000134757 | DSG3 | -0.37 | 2.96E-02 | 1.27E-01 |
| ENSG00000129450 | SIGLEC9 | -0.37 | 2.96E-02 | 1.27E-01 |
| ENSG00000172243 | CLEC7A | -0.37 | 2.97E-02 | 1.28E-01 |
| ENSG00000196664 | TLR7 | -0.37 | 2.97E-02 | 1.28E-01 |
| ENSG00000171566 | PLRG1 | -0.37 | 2.97E-02 | 1.28E-01 |
| ENSG00000182054 | IDH2 | -0.37 | 2.97E-02 | 1.28E-01 |
| ENSG00000213015 | ZNF580 | 0.37 | 2.97E-02 | 1.28E-01 |
| ENSG00000270075 | AL162742.1 | -0.37 | 2.98E-02 | 1.28E-01 |
| ENSG00000100311 | PDGFB | -0.37 | 2.98E-02 | 1.28E-01 |
| ENSG00000225381 | RPS5P7 | -0.37 | 2.98E-02 | 1.28E-01 |
| ENSG00000100216 | TOMM22 | -0.37 | 2.99E-02 | 1.28E-01 |
| ENSG00000107021 | TBC1D13 | 0.37 | 2.99E-02 | 1.28E-01 |
| ENSG00000108389 | MTMR4 | 0.37 | 2.99E-02 | 1.28E-01 |
| ENSG00000132463 | GRSF1 | -0.37 | 2.99E-02 | 1.28E-01 |
| ENSG00000135722 | FBXL8 | 0.37 | 2.99E-02 | 1.28E-01 |
| ENSG00000170881 | RNF139 | -0.37 | 2.99E-02 | 1.28E-01 |
| ENSG00000172974 | AC007318.1 | -0.37 | 2.99E-02 | 1.28E-01 |

| Ensembl ID | Gene Name | ρ | Pval | Padj |
|---|---|---|---|---|
| ENSG00000173064 | HECTD4 | 0.37 | 2.99E-02 | 1.28E-01 |
| ENSG00000205208 | C4orf46 | -0.37 | 2.99E-02 | 1.28E-01 |
| ENSG00000223799 | IL10RB-DT | -0.37 | 2.99E-02 | 1.28E-01 |
| ENSG00000234488 | AC096664.2 | -0.37 | 2.99E-02 | 1.28E-01 |
| ENSG00000236883 | AP001615.1 | -0.37 | 2.99E-02 | 1.28E-01 |
| ENSG00000198759 | EGFL6 | -0.37 | 3.00E-02 | 1.28E-01 |
| ENSG00000142185 | TRPM2 | -0.37 | 3.00E-02 | 1.28E-01 |
| ENSG00000108479 | GALK1 | -0.37 | 3.00E-02 | 1.28E-01 |
| ENSG00000118495 | PLAGL1 | 0.37 | 3.00E-02 | 1.28E-01 |
| ENSG00000123977 | DAW1 | -0.37 | 3.00E-02 | 1.28E-01 |
| ENSG00000177410 | ZFAS1 | -0.37 | 3.00E-02 | 1.28E-01 |
| ENSG00000181856 | SLC2A4 | 0.37 | 3.00E-02 | 1.28E-01 |
| ENSG00000258701 | LINC00638 | 0.37 | 3.00E-02 | 1.28E-01 |
| ENSG00000148948 | LRRC4C | -0.37 | 3.01E-02 | 1.28E-01 |
| ENSG00000254681 | PKD1P5 | 0.37 | 3.01E-02 | 1.29E-01 |
| ENSG00000267698 | AC002116.2 | 0.37 | 3.01E-02 | 1.29E-01 |
| ENSG00000036530 | CYP46A1 | 0.37 | 3.01E-02 | 1.29E-01 |
| ENSG00000106025 | TSPAN12 | -0.37 | 3.01E-02 | 1.29E-01 |
| ENSG00000131697 | NPHP4 | 0.37 | 3.01E-02 | 1.29E-01 |
| ENSG00000139154 | AEBP2 | 0.37 | 3.01E-02 | 1.29E-01 |
| ENSG00000156218 | ADAMTSL3 | 0.37 | 3.01E-02 | 1.29E-01 |
| ENSG00000170412 | GPRC5C | 0.37 | 3.01E-02 | 1.29E-01 |
| ENSG00000267809 | NDUFV2P1 | -0.37 | 3.01E-02 | 1.29E-01 |
| ENSG00000178568 | ERBB4 | 0.37 | 3.01E-02 | 1.29E-01 |
| ENSG00000243319 | FGF14-IT1 | 0.37 | 3.02E-02 | 1.29E-01 |
| ENSG00000163687 | DNASE1L3 | -0.37 | 3.02E-02 | 1.29E-01 |
| ENSG00000151023 | ENKUR | -0.37 | 3.03E-02 | 1.29E-01 |
| ENSG00000061918 | GUCY1B1 | -0.37 | 3.03E-02 | 1.29E-01 |
| ENSG00000129292 | PHF20L1 | -0.37 | 3.03E-02 | 1.29E-01 |
| ENSG00000143727 | ACP1 | -0.37 | 3.03E-02 | 1.29E-01 |
| ENSG00000140254 | DUOXA1 | -0.37 | 3.04E-02 | 1.29E-01 |
| ENSG00000079156 | OSBPL6 | -0.37 | 3.04E-02 | 1.29E-01 |
| ENSG00000099203 | TMED1 | -0.37 | 3.04E-02 | 1.29E-01 |
| ENSG00000102786 | INTS6 | 0.37 | 3.04E-02 | 1.29E-01 |
| ENSG00000105355 | PLIN3 | -0.37 | 3.04E-02 | 1.29E-01 |
| ENSG00000109016 | DHRS7B | -0.37 | 3.04E-02 | 1.29E-01 |
| ENSG00000109046 | WSB1 | 0.37 | 3.04E-02 | 1.29E-01 |
| ENSG00000125430 | HS3ST3B1 | 0.37 | 3.04E-02 | 1.29E-01 |
| ENSG00000125991 | ERGIC3 | -0.37 | 3.04E-02 | 1.29E-01 |
| ENSG00000166750 | SLFN5 | -0.37 | 3.04E-02 | 1.29E-01 |
| ENSG00000276116 | FUT8-AS1 | 0.37 | 3.04E-02 | 1.29E-01 |
| ENSG00000237054 | PRMT5-AS1 | -0.37 | 3.04E-02 | 1.29E-01 |
| ENSG00000229431 | AL139289.1 | 0.37 | 3.04E-02 | 1.29E-01 |
| ENSG00000067191 | CACNB1 | 0.37 | 3.05E-02 | 1.30E-01 |
| ENSG00000116670 | MAD2L2 | -0.37 | 3.05E-02 | 1.30E-01 |

| Ensembl ID | Gene Name | ρ | Pval | Padj |
|---|---|---|---|---|
| ENSG00000177105 | RHOG | -0.37 | 3.05E-02 | 1.30E-01 |
| ENSG00000244045 | TMEM199 | -0.37 | 3.05E-02 | 1.30E-01 |
| ENSG00000244716 | BX679664.3 | -0.37 | 3.05E-02 | 1.30E-01 |
| ENSG00000270696 | AC005034.3 | -0.37 | 3.05E-02 | 1.30E-01 |
| ENSG00000272455 | AL391244.2 | 0.37 | 3.05E-02 | 1.30E-01 |
| ENSG00000280407 | AC132872.4 | 0.37 | 3.05E-02 | 1.30E-01 |
| ENSG00000279631 | AL158211.5 | 0.37 | 3.06E-02 | 1.30E-01 |
| ENSG00000260441 | AC020978.2 | -0.37 | 3.06E-02 | 1.30E-01 |
| ENSG00000177300 | CLDN22 | -0.37 | 3.06E-02 | 1.30E-01 |
| ENSG00000128165 | ADM2 | -0.37 | 3.07E-02 | 1.30E-01 |
| ENSG00000137221 | TJAP1 | 0.37 | 3.07E-02 | 1.30E-01 |
| ENSG00000144746 | ARL6IP5 | -0.37 | 3.07E-02 | 1.30E-01 |
| ENSG00000177283 | FZD8 | 0.37 | 3.07E-02 | 1.30E-01 |
| ENSG00000177697 | CD151 | -0.37 | 3.07E-02 | 1.30E-01 |
| ENSG00000255750 | AC022509.1 | 0.37 | 3.08E-02 | 1.30E-01 |
| ENSG00000002330 | BAD | -0.37 | 3.08E-02 | 1.30E-01 |
| ENSG00000086475 | SEPHS1 | -0.37 | 3.08E-02 | 1.30E-01 |
| ENSG00000127481 | UBR4 | 0.37 | 3.08E-02 | 1.30E-01 |
| ENSG00000148926 | ADM | 0.37 | 3.08E-02 | 1.30E-01 |
| ENSG00000167548 | KMT2D | 0.37 | 3.08E-02 | 1.30E-01 |
| ENSG00000172380 | GNG12 | -0.37 | 3.08E-02 | 1.30E-01 |
| ENSG00000117385 | P3H1 | -0.37 | 3.10E-02 | 1.31E-01 |
| ENSG00000184557 | SOCS3 | -0.37 | 3.10E-02 | 1.31E-01 |
| ENSG00000206149 | HERC2P9 | 0.37 | 3.10E-02 | 1.31E-01 |
| ENSG00000263400 | TMEM220-AS1 | 0.37 | 3.10E-02 | 1.31E-01 |
| ENSG00000267855 | NDUFA7 | 0.37 | 3.10E-02 | 1.31E-01 |
| ENSG00000124177 | CHD6 | 0.36 | 3.11E-02 | 1.31E-01 |
| ENSG00000146072 | TNFRSF21 | -0.36 | 3.11E-02 | 1.31E-01 |
| ENSG00000174695 | TMEM167A | -0.36 | 3.11E-02 | 1.31E-01 |
| ENSG00000182973 | CNOT10 | -0.36 | 3.11E-02 | 1.31E-01 |
| ENSG00000205978 | NYNRIN | 0.36 | 3.11E-02 | 1.31E-01 |
| ENSG00000099866 | MADCAM1 | 0.36 | 3.11E-02 | 1.31E-01 |
| ENSG00000259322 | AC090607.1 | 0.36 | 3.11E-02 | 1.31E-01 |
| ENSG00000089723 | OTUB2 | -0.36 | 3.12E-02 | 1.31E-01 |
| ENSG00000231747 | AC079922.1 | -0.36 | 3.12E-02 | 1.31E-01 |
| ENSG00000096654 | ZNF184 | -0.36 | 3.12E-02 | 1.32E-01 |
| ENSG00000099769 | IGFALS | 0.36 | 3.13E-02 | 1.32E-01 |
| ENSG00000215482 | CALM2P3 | -0.36 | 3.13E-02 | 1.32E-01 |
| ENSG00000198844 | ARHGEF15 | -0.36 | 3.14E-02 | 1.32E-01 |
| ENSG00000047056 | WDR37 | 0.36 | 3.14E-02 | 1.32E-01 |
| ENSG00000081237 | PTPRC | -0.36 | 3.14E-02 | 1.32E-01 |
| ENSG00000179958 | DCTPP1 | -0.36 | 3.14E-02 | 1.32E-01 |
| ENSG00000197442 | MAP3K5 | 0.36 | 3.14E-02 | 1.32E-01 |
| ENSG00000269583 | L34079.3 | 0.36 | 3.14E-02 | 1.32E-01 |
| ENSG00000050393 | MCUR1 | -0.36 | 3.15E-02 | 1.32E-01 |

| Ensembl ID | Gene Name | ρ | Pval | Padj |
|---|---|---|---|---|
| ENSG00000076555 | ACACB | 0.36 | 3.15E-02 | 1.32E-01 |
| ENSG00000106605 | BLVRA | -0.36 | 3.15E-02 | 1.32E-01 |
| ENSG00000134825 | TMEM258 | -0.36 | 3.15E-02 | 1.32E-01 |
| ENSG00000164850 | GPER1 | 0.36 | 3.15E-02 | 1.32E-01 |
| ENSG00000213087 | AL356535.1 | -0.36 | 3.15E-02 | 1.32E-01 |
| ENSG00000259746 | HSPE1P3 | -0.36 | 3.15E-02 | 1.32E-01 |
| ENSG00000106069 | CHN2 | -0.36 | 3.16E-02 | 1.33E-01 |
| ENSG00000162738 | VANGL2 | -0.36 | 3.16E-02 | 1.33E-01 |
| ENSG00000106355 | LSM5 | -0.36 | 3.17E-02 | 1.33E-01 |
| ENSG00000108797 | CNTNAP1 | 0.36 | 3.17E-02 | 1.33E-01 |
| ENSG00000168066 | SF1 | 0.36 | 3.17E-02 | 1.33E-01 |
| ENSG00000171466 | ZNF562 | -0.36 | 3.17E-02 | 1.33E-01 |
| ENSG00000225793 | AL080250.1 | 0.36 | 3.17E-02 | 1.33E-01 |
| ENSG00000226715 | LINC01709 | -0.36 | 3.17E-02 | 1.33E-01 |
| ENSG00000183423 | LRIT3 | 0.36 | 3.17E-02 | 1.33E-01 |
| ENSG00000179639 | FCER1A | -0.36 | 3.17E-02 | 1.33E-01 |
| ENSG00000262477 | AC021224.1 | 0.36 | 3.17E-02 | 1.33E-01 |
| ENSG00000100319 | ZMAT5 | -0.36 | 3.18E-02 | 1.33E-01 |
| ENSG00000100983 | GSS | -0.36 | 3.18E-02 | 1.33E-01 |
| ENSG00000110047 | EHD1 | -0.36 | 3.18E-02 | 1.33E-01 |
| ENSG00000110395 | CBL | -0.36 | 3.18E-02 | 1.33E-01 |
| ENSG00000145779 | TNFAIP8 | -0.36 | 3.18E-02 | 1.33E-01 |
| ENSG00000146433 | TMEM181 | -0.36 | 3.18E-02 | 1.33E-01 |
| ENSG00000251615 | AC104825.1 | 0.36 | 3.18E-02 | 1.33E-01 |
| ENSG00000099985 | OSM | -0.36 | 3.18E-02 | 1.33E-01 |
| ENSG00000128917 | DLL4 | -0.36 | 3.19E-02 | 1.33E-01 |
| ENSG00000243566 | UPK3B | 0.36 | 3.19E-02 | 1.33E-01 |
| ENSG00000267251 | AC139100.1 | 0.36 | 3.19E-02 | 1.33E-01 |
| ENSG00000049618 | ARID1B | 0.36 | 3.19E-02 | 1.33E-01 |
| ENSG00000104643 | MTMR9 | -0.36 | 3.19E-02 | 1.33E-01 |
| ENSG00000198554 | WDHD1 | -0.36 | 3.19E-02 | 1.33E-01 |
| ENSG00000244055 | AC007566.1 | 0.36 | 3.19E-02 | 1.33E-01 |
| ENSG00000273372 | SFTPD-AS1 | 0.36 | 3.20E-02 | 1.33E-01 |
| ENSG00000261003 | AL008628.1 | 0.36 | 3.20E-02 | 1.34E-01 |
| ENSG00000104147 | OIP5 | -0.36 | 3.20E-02 | 1.34E-01 |
| ENSG00000163075 | CFAP221 | 0.36 | 3.21E-02 | 1.34E-01 |
| ENSG00000116580 | GON4L | 0.36 | 3.21E-02 | 1.34E-01 |
| ENSG00000141179 | PCTP | -0.36 | 3.21E-02 | 1.34E-01 |
| ENSG00000185453 | ZSWIM9 | 0.36 | 3.21E-02 | 1.34E-01 |
| ENSG00000260645 | AL359715.2 | 0.36 | 3.21E-02 | 1.34E-01 |
| ENSG00000208017 | MIR140 | 0.36 | 3.21E-02 | 1.34E-01 |
| ENSG00000280351 | AC127496.7 | 0.36 | 3.22E-02 | 1.34E-01 |
| ENSG00000166415 | WDR72 | -0.36 | 3.22E-02 | 1.34E-01 |
| ENSG00000119705 | SLIRP | -0.36 | 3.22E-02 | 1.34E-01 |
| ENSG00000148908 | RGS10 | -0.36 | 3.22E-02 | 1.34E-01 |

| Ensembl ID | Gene Name | ρ | Pval | Padj |
|---|---|---|---|---|
| ENSG00000165487 | MICU2 | -0.36 | 3.22E-02 | 1.34E-01 |
| ENSG00000172469 | MANEA | -0.36 | 3.22E-02 | 1.34E-01 |
| ENSG00000177556 | ATOX1 | -0.36 | 3.22E-02 | 1.34E-01 |
| ENSG00000188779 | SKOR1 | 0.36 | 3.22E-02 | 1.34E-01 |
| ENSG00000267272 | LINC01140 | 0.36 | 3.22E-02 | 1.34E-01 |
| ENSG00000196812 | ZSCAN16 | -0.36 | 3.24E-02 | 1.34E-01 |
| ENSG00000075790 | BCAP29 | -0.36 | 3.24E-02 | 1.34E-01 |
| ENSG00000106991 | ENG | -0.36 | 3.24E-02 | 1.34E-01 |
| ENSG00000158220 | ESYT3 | 0.36 | 3.24E-02 | 1.34E-01 |
| ENSG00000180644 | PRF1 | -0.36 | 3.24E-02 | 1.34E-01 |
| ENSG00000180964 | TCEAL8 | -0.36 | 3.24E-02 | 1.34E-01 |
| ENSG00000182606 | TRAK1 | -0.36 | 3.24E-02 | 1.34E-01 |
| ENSG00000186998 | EMID1 | -0.36 | 3.24E-02 | 1.34E-01 |
| ENSG00000221983 | UBA52 | -0.36 | 3.24E-02 | 1.34E-01 |
| ENSG00000070087 | PFN2 | -0.36 | 3.25E-02 | 1.35E-01 |
| ENSG00000082482 | KCNK2 | -0.36 | 3.25E-02 | 1.35E-01 |
| ENSG00000124641 | MED20 | -0.36 | 3.25E-02 | 1.35E-01 |
| ENSG00000135823 | STX6 | -0.36 | 3.25E-02 | 1.35E-01 |
| ENSG00000136463 | TACO1 | -0.36 | 3.25E-02 | 1.35E-01 |
| ENSG00000163346 | PBXIP1 | 0.36 | 3.25E-02 | 1.35E-01 |
| ENSG00000163811 | WDR43 | -0.36 | 3.25E-02 | 1.35E-01 |
| ENSG00000168393 | DTYMK | -0.36 | 3.25E-02 | 1.35E-01 |
| ENSG00000213859 | KCTD11 | -0.36 | 3.25E-02 | 1.35E-01 |
| ENSG00000250548 | LINC01303 | -0.36 | 3.25E-02 | 1.35E-01 |
| ENSG00000100991 | TRPC4AP | 0.36 | 3.27E-02 | 1.35E-01 |
| ENSG00000106993 | CDC37L1 | -0.36 | 3.27E-02 | 1.35E-01 |
| ENSG00000112245 | PTP4A1 | -0.36 | 3.27E-02 | 1.35E-01 |
| ENSG00000112977 | DAP | 0.36 | 3.27E-02 | 1.35E-01 |
| ENSG00000115364 | MRPL19 | -0.36 | 3.27E-02 | 1.35E-01 |
| ENSG00000136869 | TLR4 | -0.36 | 3.27E-02 | 1.35E-01 |
| ENSG00000162522 | KIAA1522 | -0.36 | 3.27E-02 | 1.35E-01 |
| ENSG00000253676 | TAGLN2P1 | -0.36 | 3.27E-02 | 1.35E-01 |
| ENSG00000269439 | AC010618.4 | 0.36 | 3.27E-02 | 1.35E-01 |
| ENSG00000270933 | AC010719.1 | 0.36 | 3.27E-02 | 1.35E-01 |
| ENSG00000070404 | FSTL3 | -0.36 | 3.28E-02 | 1.35E-01 |
| ENSG00000106351 | AGFG2 | 0.36 | 3.28E-02 | 1.35E-01 |
| ENSG00000143977 | SNRPG | -0.36 | 3.28E-02 | 1.35E-01 |
| ENSG00000172493 | AFF1 | 0.36 | 3.28E-02 | 1.35E-01 |
| ENSG00000185495 | AC138393.1 | 0.36 | 3.28E-02 | 1.35E-01 |
| ENSG00000259728 | LINC00933 | 0.36 | 3.28E-02 | 1.35E-01 |
| ENSG00000260236 | AC099778.1 | 0.36 | 3.28E-02 | 1.35E-01 |
| ENSG00000203761 | MSTO2P | 0.36 | 3.29E-02 | 1.36E-01 |
| ENSG00000124786 | SLC35B3 | -0.36 | 3.29E-02 | 1.36E-01 |
| ENSG00000133302 | SLF1 | -0.36 | 3.29E-02 | 1.36E-01 |
| ENSG00000141564 | RPTOR | 0.36 | 3.29E-02 | 1.36E-01 |

| Ensembl ID | Gene Name | ρ | Pval | Padj |
|---|---|---|---|---|
| ENSG00000184076 | UQCR10 | -0.36 | 3.29E-02 | 1.36E-01 |
| ENSG00000064042 | LIMCH1 | 0.36 | 3.31E-02 | 1.36E-01 |
| ENSG00000180667 | YOD1 | -0.36 | 3.31E-02 | 1.36E-01 |
| ENSG00000228049 | POLR2J2 | 0.36 | 3.31E-02 | 1.37E-01 |
| ENSG00000226833 | AC092164.1 | 0.36 | 3.32E-02 | 1.37E-01 |
| ENSG00000009724 | MASP2 | 0.36 | 3.32E-02 | 1.37E-01 |
| ENSG00000117500 | TMED5 | -0.36 | 3.32E-02 | 1.37E-01 |
| ENSG00000134324 | LPIN1 | 0.36 | 3.32E-02 | 1.37E-01 |
| ENSG00000164024 | METAP1 | -0.36 | 3.32E-02 | 1.37E-01 |
| ENSG00000173120 | KDM2A | 0.36 | 3.32E-02 | 1.37E-01 |
| ENSG00000173482 | PTPRM | 0.36 | 3.32E-02 | 1.37E-01 |
| ENSG00000249790 | AC092490.1 | -0.36 | 3.33E-02 | 1.37E-01 |
| ENSG00000184414 | IRS3P | 0.36 | 3.33E-02 | 1.37E-01 |
| ENSG00000074755 | ZZEF1 | 0.36 | 3.34E-02 | 1.37E-01 |
| ENSG00000124374 | PAIP2B | 0.36 | 3.34E-02 | 1.37E-01 |
| ENSG00000136868 | SLC31A1 | -0.36 | 3.34E-02 | 1.37E-01 |
| ENSG00000143256 | PFDN2 | -0.36 | 3.34E-02 | 1.37E-01 |
| ENSG00000166847 | DCTN5 | -0.36 | 3.34E-02 | 1.37E-01 |
| ENSG00000169905 | TOR1AIP2 | -0.36 | 3.34E-02 | 1.37E-01 |
| ENSG00000178988 | MRFAP1L1 | -0.36 | 3.34E-02 | 1.37E-01 |
| ENSG00000231365 | WARS2-AS1 | 0.36 | 3.34E-02 | 1.37E-01 |
| ENSG00000228906 | AL353804.1 | 0.36 | 3.34E-02 | 1.37E-01 |
| ENSG00000254985 | RSF1-IT2 | 0.36 | 3.34E-02 | 1.37E-01 |
| ENSG00000186019 | AC021092.1 | 0.36 | 3.34E-02 | 1.37E-01 |
| ENSG00000279484 | KLHL30-AS1 | -0.36 | 3.34E-02 | 1.37E-01 |
| ENSG00000165443 | PHYHIPL | -0.36 | 3.35E-02 | 1.37E-01 |
| ENSG00000135821 | GLUL | -0.36 | 3.35E-02 | 1.38E-01 |
| ENSG00000163684 | RPP14 | 0.36 | 3.35E-02 | 1.38E-01 |
| ENSG00000260361 | AC106028.3 | 0.36 | 3.35E-02 | 1.38E-01 |
| ENSG00000103485 | QPRT | -0.36 | 3.37E-02 | 1.38E-01 |
| ENSG00000100575 | TIMM9 | -0.36 | 3.37E-02 | 1.38E-01 |
| ENSG00000140511 | HAPLN3 | -0.36 | 3.37E-02 | 1.38E-01 |
| ENSG00000143748 | NVL | 0.36 | 3.37E-02 | 1.38E-01 |
| ENSG00000147642 | SYBU | 0.36 | 3.37E-02 | 1.38E-01 |
| ENSG00000148303 | RPL7A | -0.36 | 3.37E-02 | 1.38E-01 |
| ENSG00000176903 | PNMA1 | -0.36 | 3.37E-02 | 1.38E-01 |
| ENSG00000186462 | NAP1L2 | 0.36 | 3.37E-02 | 1.38E-01 |
| ENSG00000254505 | CHMP4A | 0.36 | 3.37E-02 | 1.38E-01 |
| ENSG00000164821 | DEFA4 | -0.36 | 3.38E-02 | 1.38E-01 |
| ENSG00000283498 | MIR1244-2 | -0.36 | 3.38E-02 | 1.38E-01 |
| ENSG00000215022 | AL008729.1 | 0.36 | 3.38E-02 | 1.38E-01 |
| ENSG00000054598 | FOXC1 | 0.36 | 3.38E-02 | 1.38E-01 |
| ENSG00000128655 | PDE11A | 0.36 | 3.38E-02 | 1.38E-01 |
| ENSG00000141552 | ANAPC11 | -0.36 | 3.38E-02 | 1.38E-01 |
| ENSG00000163291 | PAQR3 | -0.36 | 3.38E-02 | 1.38E-01 |

| Ensembl ID | Gene Name | ρ | Pval | Padj |
|---|---|---|---|---|
| ENSG00000170540 | ARL6IP1 | -0.36 | 3.38E-02 | 1.38E-01 |
| ENSG00000171962 | DRC3 | 0.36 | 3.38E-02 | 1.38E-01 |
| ENSG00000163357 | DCST1 | 0.36 | 3.38E-02 | 1.38E-01 |
| ENSG00000072041 | SLC6A15 | 0.36 | 3.38E-02 | 1.38E-01 |
| ENSG00000280106 | AC008555.7 | -0.36 | 3.39E-02 | 1.38E-01 |
| ENSG00000271966 | AC021321.1 | 0.36 | 3.39E-02 | 1.38E-01 |
| ENSG00000263818 | RDM1P5 | -0.36 | 3.39E-02 | 1.38E-01 |
| ENSG00000255503 | AP001767.3 | 0.36 | 3.40E-02 | 1.39E-01 |
| ENSG00000103429 | BFAR | -0.36 | 3.40E-02 | 1.39E-01 |
| ENSG00000131845 | ZNF304 | -0.36 | 3.40E-02 | 1.39E-01 |
| ENSG00000188177 | ZC3H6 | 0.36 | 3.40E-02 | 1.39E-01 |
| ENSG00000198015 | MRPL42 | -0.36 | 3.40E-02 | 1.39E-01 |
| ENSG00000227220 | AL133346.1 | 0.36 | 3.40E-02 | 1.39E-01 |
| ENSG00000261716 | AC239868.1 | 0.36 | 3.40E-02 | 1.39E-01 |
| ENSG00000269890 | AL353593.1 | 0.36 | 3.40E-02 | 1.39E-01 |
| ENSG00000233922 | LINC01694 | 0.36 | 3.41E-02 | 1.39E-01 |
| ENSG00000188985 | DHFRP1 | -0.36 | 3.41E-02 | 1.39E-01 |
| ENSG00000040633 | PHF23 | -0.36 | 3.41E-02 | 1.39E-01 |
| ENSG00000102879 | CORO1A | -0.36 | 3.41E-02 | 1.39E-01 |
| ENSG00000143546 | S100A8 | -0.36 | 3.41E-02 | 1.39E-01 |
| ENSG00000205054 | LINC01121 | -0.36 | 3.42E-02 | 1.39E-01 |
| ENSG00000141574 | SECTM1 | -0.36 | 3.42E-02 | 1.39E-01 |
| ENSG00000072163 | LIMS2 | -0.36 | 3.43E-02 | 1.39E-01 |
| ENSG00000087095 | NLK | -0.36 | 3.43E-02 | 1.39E-01 |
| ENSG00000090861 | AARS | -0.36 | 3.43E-02 | 1.39E-01 |
| ENSG00000131127 | ZNF141 | 0.36 | 3.43E-02 | 1.39E-01 |
| ENSG00000179010 | MRFAP1 | -0.36 | 3.43E-02 | 1.39E-01 |
| ENSG00000230896 | AL604028.1 | 0.36 | 3.43E-02 | 1.39E-01 |
| ENSG00000231167 | YBX1P2 | -0.36 | 3.43E-02 | 1.39E-01 |
| ENSG00000259877 | AC009113.1 | 0.36 | 3.43E-02 | 1.39E-01 |
| ENSG00000151490 | PTPRO | -0.36 | 3.44E-02 | 1.40E-01 |
| ENSG00000226314 | ZNF192P1 | 0.36 | 3.44E-02 | 1.40E-01 |
| ENSG00000022277 | RTF2 | -0.36 | 3.44E-02 | 1.40E-01 |
| ENSG00000136942 | RPL35 | -0.36 | 3.44E-02 | 1.40E-01 |
| ENSG00000166510 | CCDC68 | 0.36 | 3.44E-02 | 1.40E-01 |
| ENSG00000173960 | UBXN2A | 0.36 | 3.44E-02 | 1.40E-01 |
| ENSG00000184271 | AC139768.1 | 0.36 | 3.44E-02 | 1.40E-01 |
| ENSG00000231133 | HAR1B | 0.36 | 3.45E-02 | 1.40E-01 |
| ENSG00000046651 | OFD1 | 0.36 | 3.46E-02 | 1.40E-01 |
| ENSG00000055211 | GINM1 | -0.36 | 3.46E-02 | 1.40E-01 |
| ENSG00000105939 | ZC3HAV1 | -0.36 | 3.46E-02 | 1.40E-01 |
| ENSG00000160789 | LMNA | -0.36 | 3.46E-02 | 1.40E-01 |
| ENSG00000164109 | MAD2L1 | -0.36 | 3.46E-02 | 1.40E-01 |
| ENSG00000169567 | HINT1 | -0.36 | 3.46E-02 | 1.40E-01 |
| ENSG00000267356 | AC006557.3 | 0.36 | 3.47E-02 | 1.41E-01 |

| Ensembl ID | Gene Name | ρ | Pval | Padj |
|---|---|---|---|---|
| ENSG00000087274 | ADD1 | 0.36 | 3.47E-02 | 1.41E-01 |
| ENSG00000104738 | MCM4 | -0.36 | 3.47E-02 | 1.41E-01 |
| ENSG00000109084 | TMEM97 | -0.36 | 3.47E-02 | 1.41E-01 |
| ENSG00000135090 | TAOK3 | -0.36 | 3.47E-02 | 1.41E-01 |
| ENSG00000163728 | TTC14 | 0.36 | 3.47E-02 | 1.41E-01 |
| ENSG00000175538 | KCNE3 | -0.36 | 3.47E-02 | 1.41E-01 |
| ENSG00000197345 | MRPL21 | -0.36 | 3.47E-02 | 1.41E-01 |
| ENSG00000247828 | TMEM161B-AS1 | 0.36 | 3.47E-02 | 1.41E-01 |
| ENSG00000203814 | HIST2H2BF | -0.36 | 3.48E-02 | 1.41E-01 |
| ENSG00000153012 | LGI2 | -0.36 | 3.49E-02 | 1.41E-01 |
| ENSG00000158406 | HIST1H4H | -0.36 | 3.49E-02 | 1.41E-01 |
| ENSG00000167785 | ZNF558 | 0.36 | 3.49E-02 | 1.41E-01 |
| ENSG00000198793 | MTOR | 0.36 | 3.49E-02 | 1.41E-01 |
| ENSG00000237788 | AL162615.1 | 0.36 | 3.50E-02 | 1.41E-01 |
| ENSG00000272600 | AC007308.1 | -0.36 | 3.50E-02 | 1.41E-01 |
| ENSG00000004142 | POLDIP2 | -0.36 | 3.50E-02 | 1.41E-01 |
| ENSG00000071243 | ING3 | -0.36 | 3.50E-02 | 1.41E-01 |
| ENSG00000105717 | PBX4 | 0.36 | 3.50E-02 | 1.41E-01 |
| ENSG00000127419 | TMEM175 | 0.36 | 3.50E-02 | 1.41E-01 |
| ENSG00000156136 | DCK | -0.36 | 3.50E-02 | 1.41E-01 |
| ENSG00000161281 | COX7A1 | -0.36 | 3.50E-02 | 1.41E-01 |
| ENSG00000172977 | KAT5 | 0.36 | 3.50E-02 | 1.41E-01 |
| ENSG00000236060 | HSPB1P1 | -0.36 | 3.51E-02 | 1.41E-01 |
| ENSG00000261093 | AC141586.3 | 0.36 | 3.51E-02 | 1.41E-01 |
| ENSG00000274210 | RF00003 | -0.36 | 3.51E-02 | 1.42E-01 |
| ENSG00000003249 | DBNDD1 | -0.36 | 3.52E-02 | 1.42E-01 |
| ENSG00000034693 | PEX3 | 0.36 | 3.52E-02 | 1.42E-01 |
| ENSG00000105669 | COPE | -0.36 | 3.52E-02 | 1.42E-01 |
| ENSG00000115216 | NRBP1 | -0.36 | 3.52E-02 | 1.42E-01 |
| ENSG00000152380 | FAM151B | -0.36 | 3.52E-02 | 1.42E-01 |
| ENSG00000160445 | ZER1 | 0.36 | 3.52E-02 | 1.42E-01 |
| ENSG00000180891 | CUEDC1 | -0.36 | 3.52E-02 | 1.42E-01 |
| ENSG00000268575 | AL031282.2 | 0.36 | 3.52E-02 | 1.42E-01 |
| ENSG00000137343 | ATAT1 | 0.36 | 3.53E-02 | 1.42E-01 |
| ENSG00000144736 | SHQ1 | -0.36 | 3.53E-02 | 1.42E-01 |
| ENSG00000176624 | MEX3C | -0.36 | 3.53E-02 | 1.42E-01 |
| ENSG00000246922 | UBAP1L | 0.36 | 3.53E-02 | 1.42E-01 |
| ENSG00000273295 | AP000350.6 | 0.36 | 3.53E-02 | 1.42E-01 |
| ENSG00000135436 | FAM186B | 0.36 | 3.54E-02 | 1.42E-01 |
| ENSG00000204577 | LILRB3 | -0.36 | 3.54E-02 | 1.42E-01 |
| ENSG00000231313 | CLIC1P1 | -0.36 | 3.54E-02 | 1.42E-01 |
| ENSG00000049541 | RFC2 | -0.36 | 3.55E-02 | 1.42E-01 |
| ENSG00000101343 | CRNKL1 | -0.36 | 3.55E-02 | 1.42E-01 |
| ENSG00000143321 | HDGF | -0.36 | 3.55E-02 | 1.42E-01 |

| Ensembl ID | Gene Name | ρ | Pval | Padj |
|---|---|---|---|---|
| ENSG00000155363 | MOV10 | 0.36 | 3.55E-02 | 1.42E-01 |
| ENSG00000228847 | ATP5MC2P4 | -0.36 | 3.55E-02 | 1.42E-01 |
| ENSG00000256628 | ZBTB11-AS1 | 0.36 | 3.55E-02 | 1.42E-01 |
| ENSG00000251442 | LINC01094 | -0.36 | 3.55E-02 | 1.42E-01 |
| ENSG00000105383 | CD33 | -0.36 | 3.55E-02 | 1.43E-01 |
| ENSG00000276476 | AL136962.1 | -0.36 | 3.55E-02 | 1.43E-01 |
| ENSG00000260369 | AC120024.1 | 0.36 | 3.56E-02 | 1.43E-01 |
| ENSG00000255334 | AP000907.2 | 0.36 | 3.56E-02 | 1.43E-01 |
| ENSG00000106635 | BCL7B | -0.36 | 3.57E-02 | 1.43E-01 |
| ENSG00000126432 | PRDX5 | -0.36 | 3.57E-02 | 1.43E-01 |
| ENSG00000176340 | COX8A | -0.36 | 3.57E-02 | 1.43E-01 |
| ENSG00000214389 | RPS3AP26 | -0.36 | 3.57E-02 | 1.43E-01 |
| ENSG00000240972 | MIF | -0.36 | 3.57E-02 | 1.43E-01 |
| ENSG00000168497 | CAVIN2 | -0.36 | 3.57E-02 | 1.43E-01 |
| ENSG00000237596 | AL138828.1 | 0.36 | 3.57E-02 | 1.43E-01 |
| ENSG00000181790 | ADGRB1 | 0.36 | 3.58E-02 | 1.43E-01 |
| ENSG00000028137 | TNFRSF1B | -0.36 | 3.58E-02 | 1.43E-01 |
| ENSG00000120837 | NFYB | -0.36 | 3.58E-02 | 1.43E-01 |
| ENSG00000126500 | FLRT1 | 0.36 | 3.58E-02 | 1.43E-01 |
| ENSG00000130702 | LAMA5 | -0.36 | 3.58E-02 | 1.43E-01 |
| ENSG00000144115 | THNSL2 | 0.36 | 3.58E-02 | 1.43E-01 |

**Table S3** - Significant gene enrichment of 98 genes correlating to WWP2

| GO-term | Count | Pval | FDR | Genes |
|---|---|---|---|---|
| GO:0070062~extracellular exosome | 36 | 1.94E-07 | 2.26E-05 | RALA, SCIN, CDC42, WDR1, CMTM6, MYL12A, CHCHD3, LIPA, HSPA8, STX2, RAB23, CD58, TGFBI, RAB22A, SMO, CAP1, PATJ, ACTR1A, TPM3, ALDH1L1, ATP6V1B2, PRSS23, CCT5, GJA1, XPC, TAGLN2, SPTBN4, ARPC2, YWHAZ, CCT2, YWHAB, TUBA1A, CDK1, ACTG1, AKR1C3, TRABD2B |
| GO:0043209~myelin sheath | 10 | 2.43E-07 | 2.26E-05 | RALA, CDC42, WDR1, HSPA8, ACTR1A, ATP6V1B2, CCT5, CCT2, TUBA1A, ACTG1 |
| GO:0005829~cytosol | 35 | 2.56E-04 | 1.59E-02 | TSPOAP1, TRAF1, CDC42, WDR1, TPX2, MYL12A, HSPA8, BAG2, RGS2, CENPF, HIF3A, WNK4, ACTR1A, TPM3, ALDH1L1, SH3KBP1, ATP6V1B2, CCT5, GJA1, OBSCN, SPTBN4, ARPC2, YWHAZ, WNK2, CCT2, YWHAB, SGSM1, TUBA1A, CDK1, PFKFB3, IL16, KPNA2, ACTG1, AKR1C3, ARHGAP11A |
| GO:0005925~focal adhesion | 10 | 6.46E-04 | 3.00E-02 | RALA, CDC42, HSPA8, CAP1, SH3KBP1, GJA1, ARPC2, YWHAZ, YWHAB, ACTG1 |

**Table S4** - Differential protein expression on day three and day seven relative to day zero of chondrogenesis in our 3D chondrocyte pellet cultures.

| Accession | Name | # Peptides | # Unique Peptides | Abundance Ratio: day 3 / day 0 | Abundance Ratio: day 7/ day 0 |
|---|---|---|---|---|---|
| Q92845 | KIFAP3 | 2 | 1 | 0.01 | 0.97 |
| Q96BM9 | ARL8A | 2 | 1 | 0.01 | 0.76 |
| Q8N138 | ORMDL3 | 2 | 1 | 0.01 | 0.01 |
| Q96T58 | SPEN | 2 | 1 | 0.01 | 0.01 |
| P78352 | DLG4 | 2 | 1 | 0.01 | 0.86 |
| Q8ND76 | CCNY | 2 | 1 | 0.01 | 1.15 |
| Q9NRF8 | CTPS2 | 2 | 1 | 0.01 | 0.01 |
| P05783 | KRT18 | 2 | 1 | 0.01 | 0.01 |
| Q0VDG4 | SCRN3 | 2 | 1 | 0.01 | 0.01 |
| Q5VTR2 | RNF20 | 2 | 1 | 0.01 | 0.01 |
| Q96C36 | PYCR2 | 2 | 1 | 0.01 | 0.01 |
| Q9UK45 | LSM7 | 2 | 1 | 0.01 | 0.95 |
| Q8N4A0 | GALNT4 | 2 | 1 | 0.01 | 0.01 |
| Q9HCN4 | GPN1 | 2 | 2 | 0.01 | 0.01 |
| Q7LBC6 | KDM3B | 2 | 2 | 0.01 | 1.12 |
| Q96EK9 | KTI12 | 2 | 2 | 0.01 | 0.01 |
| Q9UBN7 | HDAC6 | 2 | 2 | 0.01 | 0.01 |
| Q9UI42 | CPA4 | 2 | 2 | 0.01 | 0.01 |
| O14734 | ACOT8 | 2 | 2 | 0.01 | 0.01 |
| Q6IN85 | PPP4R3A | 2 | 2 | 0.01 | 0.01 |
| O95249 | GOSR1 | 2 | 2 | 0.01 | 0.87 |
| Q9H0U9 | TSPYL1 | 2 | 2 | 0.01 | 0.01 |
| Q68CQ7 | GLT8D1 | 2 | 2 | 0.01 | 1.58 |
| P45877 | PPIC | 2 | 2 | 0.01 | 2.13 |
| Q7LGC8 | CHST3 | 2 | 2 | 0.01 | 0.01 |
| O95183 | VAMP5 | 2 | 2 | 0.01 | 0.01 |
| Q96A35 | MRPL24 | 2 | 2 | 0.01 | 0.01 |
| Q8N4V1 | MMGT1 | 2 | 2 | 0.01 | 0.01 |
| Q8ND04 | SMG8 | 2 | 2 | 0.01 | 0.01 |
| Q9Y5K6 | CD2AP | 2 | 2 | 0.01 | 0.01 |
| Q7Z7L8 | C11orf96 | 2 | 2 | 0.01 | 0.01 |
| Q5VZ89 | DENND4C | 2 | 2 | 0.01 | 0.01 |
| P15954 | COX7C | 2 | 2 | 0.01 | 0.01 |
| P55789 | GFER | 2 | 2 | 0.01 | 0.01 |
| O60888 | CUTA | 2 | 2 | 0.01 | 0.01 |
| Q99685 | MGLL | 2 | 2 | 0.01 | 0.01 |
| Q14644 | RASA3 | 2 | 2 | 0.01 | 0.01 |
| Q96AB3 | ISOC2 | 2 | 2 | 0.01 | 0.01 |
| Q9NY26 | SLC39A1 | 2 | 2 | 0.01 | 0.01 |
| Q9Y2Q9 | MRPS28 | 2 | 2 | 0.01 | 0.01 |

| Accession | Name | # Peptides | # Unique Peptides | Abundance Ratio: day 3 / day 0 | Abundance Ratio: day 7/ day 0 |
|---|---|---|---|---|---|
| Q92845 | KIFAP3 | 2 | 1 | 0.01 | 0.97 |
| Q96BM9 | ARL8A | 2 | 1 | 0.01 | 0.76 |
| Q86X95 | CIR1 | 2 | 2 | 0.01 | 0.01 |
| Q9BXJ8 | TMEM120A | 2 | 2 | 0.01 | 0.01 |
| Q53HC9 | EIPR1 | 2 | 2 | 0.01 | 0.01 |
| Q9BXW7 | HDHD5 | 2 | 2 | 0.01 | 0.01 |
| Q96ER9 | CCDC51 | 2 | 2 | 0.01 | 0.01 |
| Q9UHB7 | AFF4 | 2 | 2 | 0.01 | 0.01 |
| Q9H000 | MKRN2 | 2 | 2 | 0.01 | 0.01 |
| Q13485 | SMAD4 | 2 | 2 | 0.01 | 0.01 |
| Q9UIV1 | CNOT7 | 2 | 2 | 0.01 | 0.01 |
| Q9H461 | FZD8 | 2 | 2 | 0.01 | 0.01 |
| Q96RL1 | UIMC1 | 2 | 2 | 0.01 | 0.99 |
| Q9Y3B7 | MRPL11 | 2 | 2 | 0.01 | 0.01 |
| O43464 | HTRA2 | 2 | 2 | 0.01 | 0.01 |
| O43402 | EMC8 | 2 | 2 | 0.01 | 0.01 |
| Q6ZS17 | RIPOR1 | 2 | 2 | 0.01 | 0.01 |
| P35251 | RFC1 | 2 | 2 | 0.01 | 0.01 |
| Q9HCE0 | EPG5 | 2 | 2 | 0.01 | 0.01 |
| Q14137 | BOP1 | 2 | 2 | 0.01 | 0.01 |
| Q13685 | AAMP | 2 | 2 | 0.01 | 0.01 |
| Q8WW12 | PCNP | 2 | 2 | 0.01 | 1.84 |
| O15056 | SYNJ2 | 2 | 2 | 0.01 | 0.01 |
| P27707 | DCK | 2 | 2 | 0.01 | 0.01 |
| P38435 | GGCX | 2 | 2 | 0.01 | 1.46 |
| Q8TEQ8 | PIGO | 2 | 2 | 0.01 | 0.01 |
| Q13332 | PTPRS | 2 | 2 | 0.01 | 0.01 |
| P11274 | BCR | 2 | 2 | 0.01 | 0.01 |
| O14782 | KIF3C | 2 | 2 | 0.01 | 0.76 |
| Q9NXW2 | DNAJB12 | 2 | 2 | 0.01 | 0.01 |
| Q8NBU5 | ATAD1 | 2 | 2 | 0.01 | 0.01 |
| Q9C0H2 | TTYH3 | 2 | 2 | 0.01 | 0.01 |
| P08174 | CD55 | 2 | 2 | 0.01 | 0.76 |
| P05230 | FGF1 | 2 | 2 | 0.01 | 0.01 |
| O75153 | CLUH | 2 | 2 | 0.01 | 0.01 |
| Q15036 | SNX17 | 2 | 2 | 0.01 | 1.69 |
| Q6BCY4 | CYB5R2 | 2 | 2 | 0.01 | 0.01 |
| P78536 | ADAM17 | 2 | 2 | 0.01 | 0.01 |
| Q8IY37 | DHX37 | 2 | 2 | 0.01 | 0.01 |
| Q53T59 | HS1BP3 | 2 | 2 | 0.01 | 0.01 |
| Q9Y6G5 | COMMD10 | 2 | 2 | 0.01 | 0.01 |
| Q8NB90 | SPATA5 | 2 | 2 | 0.01 | 0.01 |
| Q8TEQ0 | SNX29 | 2 | 2 | 0.01 | 0.01 |

| Accession | Name | # Peptides | # Unique Peptides | Abundance Ratio: day 3 / day 0 | Abundance Ratio: day 7/ day 0 |
|---|---|---|---|---|---|
| Q92845 | KIFAP3 | 2 | 1 | 0.01 | 0.97 |
| Q96BM9 | ARL8A | 2 | 1 | 0.01 | 0.76 |
| Q6P4E1 | GOLM2 | 2 | 2 | 0.01 | 0.01 |
| Q14197 | MRPL58 | 2 | 2 | 0.01 | 0.01 |
| Q01650 | SLC7A5 | 2 | 2 | 0.01 | 1.01 |
| O60306 | AQR | 2 | 2 | 0.01 | 0.01 |
| Q9UMZ2 | SYNRG | 2 | 2 | 0.01 | 0.01 |
| Q9NXR7 | BABAM2 | 2 | 2 | 0.01 | 0.01 |
| P40855 | PEX19 | 2 | 2 | 0.01 | 0.01 |
| P47989 | XDH | 2 | 2 | 0.01 | 0.70 |
| Q658Y4 | FAM91A1 | 2 | 2 | 0.01 | 1.02 |
| Q96QG7 | MTMR9 | 2 | 2 | 0.01 | 0.01 |
| Q5EB52 | MEST | 2 | 2 | 0.01 | 0.01 |
| Q8N511 | TMEM199 | 2 | 2 | 0.01 | 0.01 |
| Q9UK59 | DBR1 | 2 | 2 | 0.01 | 0.80 |
| Q53GG5 | PDLIM3 | 2 | 2 | 0.01 | 0.01 |
| O60711 | LPXN | 2 | 2 | 0.01 | 0.01 |
| Q96DV4 | MRPL38 | 2 | 2 | 0.01 | 0.01 |
| P57105 | SYNJ2BP | 2 | 2 | 0.01 | 1.21 |
| Q9Y5K5 | UCHL5 | 2 | 2 | 0.01 | 0.01 |
| Q9UKG9 | CROT | 2 | 2 | 0.01 | 0.01 |
| Q96SZ5 | ADO | 2 | 2 | 0.01 | 0.84 |
| Q92643 | PIGK | 2 | 2 | 0.01 | 1.03 |
| P15735 | PHKG2 | 2 | 2 | 0.01 | 0.01 |
| Q9Y496 | KIF3A | 2 | 2 | 0.01 | 0.01 |
| Q6ZUX7 | LHFPL2 | 2 | 2 | 0.01 | 0.01 |
| Q0VGL1 | LAMTOR4 | 2 | 2 | 0.01 | 0.01 |
| Q99704 | DOK1 | 2 | 2 | 0.01 | 0.54 |
| P11441 | UBL4A | 2 | 2 | 0.01 | 0.79 |
| Q5T5Y3 | CAMSAP1 | 2 | 2 | 0.01 | 0.77 |
| P35354 | PTGS2 | 2 | 2 | 0.01 | 0.01 |
| Q6IA86 | ELP2 | 2 | 2 | 0.01 | 0.01 |
| O75208 | COQ9 | 2 | 2 | 0.01 | 0.01 |
| Q9NRF9 | POLE3 | 2 | 2 | 0.01 | 0.01 |
| Q14432 | PDE3A | 2 | 2 | 0.01 | 0.01 |
| Q6NXE6 | ARMC6 | 2 | 2 | 0.01 | 0.01 |
| P43246 | MSH2 | 2 | 2 | 0.01 | 0.01 |
| Q96JH7 | VCPIP1 | 2 | 2 | 0.01 | 0.01 |
| Q8N7H5 | PAF1 | 2 | 2 | 0.01 | 0.90 |
| Q7Z7F7 | MRPL55 | 2 | 2 | 0.01 | 0.01 |
| Q9H4L5 | OSBPL3 | 2 | 2 | 0.01 | 0.01 |
| Q9BW62 | KATNAL1 | 2 | 2 | 0.01 | 0.01 |
| Q5JVS0 | HABP4 | 2 | 2 | 0.01 | 0.01 |

| Accession | Name | # Peptides | # Unique Peptides | Abundance Ratio: day 3 / day 0 | Abundance Ratio: day 7/ day 0 |
|---|---|---|---|---|---|
| Q92845 | KIFAP3 | 2 | 1 | 0.01 | 0.97 |
| Q96BM9 | ARL8A | 2 | 1 | 0.01 | 0.76 |
| Q9NPD3 | EXOSC4 | 2 | 2 | 0.01 | 0.01 |
| Q9H2W6 | MRPL46 | 2 | 2 | 0.01 | 0.01 |
| Q96KA5 | CLPTM1L | 2 | 2 | 0.01 | 0.82 |
| O15382 | BCAT2 | 2 | 2 | 0.01 | 0.01 |
| C9JLW8 | MCRIP1 | 2 | 2 | 0.01 | 0.01 |
| Q69YN4 | VIRMA | 2 | 2 | 0.01 | 0.01 |
| Q9ULX3 | NOB1 | 2 | 2 | 0.01 | 0.86 |
| Q15388 | TOMM20 | 2 | 2 | 0.01 | 0.56 |
| Q9Y2D0 | CA5B | 2 | 2 | 0.01 | 0.01 |
| A0MZ66 | SHTN1 | 2 | 2 | 0.01 | 0.01 |
| Q5KU26 | COLEC12 | 2 | 2 | 0.01 | 1.74 |
| O60331 | PIP5K1C | 2 | 2 | 0.01 | 0.01 |
| Q969Q5 | RAB24 | 2 | 2 | 0.01 | 0.01 |
| Q9HCE5 | METTL14 | 2 | 2 | 0.01 | 0.01 |
| Q8WU79 | SMAP2 | 2 | 2 | 0.01 | 0.56 |
| Q5T2E6 | ARMH3 | 2 | 2 | 0.01 | 0.01 |
| Q9UBP4 | DKK3 | 2 | 2 | 0.01 | 0.01 |
| Q9NRX2 | MRPL17 | 2 | 2 | 0.01 | 0.01 |
| Q8TDH9 | BLOC1S5 | 2 | 2 | 0.01 | 0.01 |
| P24386 | CHM | 2 | 2 | 0.01 | 0.90 |
| Q8NDT2 | RBM15B | 2 | 2 | 0.01 | 0.94 |
| Q13049 | TRIM32 | 2 | 2 | 0.01 | 0.01 |
| Q9H6R4 | NOL6 | 2 | 2 | 0.01 | 0.93 |
| P80217 | IFI35 | 2 | 2 | 0.01 | 0.01 |
| O43920 | NDUFS5 | 2 | 2 | 0.01 | 0.01 |
| P08603 | CFH | 2 | 2 | 0.01 | 0.01 |
| P03956 | MMP1 | 2 | 2 | 0.01 | 0.01 |
| Q9NWT6 | HIF1AN | 2 | 2 | 0.01 | 0.01 |
| Q5TDH0 | DDI2 | 2 | 2 | 0.01 | 0.01 |
| Q8IXM6 | NRM | 2 | 2 | 0.01 | 0.01 |
| O60462 | NRP2 | 2 | 2 | 0.01 | 0.01 |
| O15397 | IPO8 | 2 | 2 | 0.01 | 0.01 |
| Q6UW02 | CYP20A1 | 2 | 2 | 0.01 | 0.91 |
| Q9NYK5 | MRPL39 | 2 | 2 | 0.01 | 0.01 |
| Q7Z7H8 | MRPL10 | 2 | 2 | 0.01 | 0.01 |
| Q9Y2V7 | COG6 | 2 | 2 | 0.01 | 1.01 |
| P05107 | ITGB2 | 2 | 2 | 0.01 | 0.01 |
| Q6ZVM7 | TOM1L2 | 2 | 2 | 0.01 | 0.46 |
| O43181 | NDUFS4 | 2 | 2 | 0.01 | 0.59 |
| Q9Y606 | PUS1 | 2 | 2 | 0.01 | 0.01 |
| Q676U5 | ATG16L1 | 2 | 2 | 0.01 | 0.01 |

| Accession | Name | # Peptides | # Unique Peptides | Abundance Ratio: day 3 / day 0 | Abundance Ratio: day 7/ day 0 |
|---|---|---|---|---|---|
| Q92845 | KIFAP3 | 2 | 1 | 0.01 | 0.97 |
| Q96BM9 | ARL8A | 2 | 1 | 0.01 | 0.76 |
| Q96EY8 | MMAB | 2 | 2 | 0.01 | 0.01 |
| O14981 | BTAF1 | 2 | 2 | 0.01 | 0.01 |
| Q8WUJ3 | CEMIP | 2 | 2 | 0.01 | 0.01 |
| O75122 | CLASP2 | 2 | 2 | 0.01 | 1.06 |
| Q96PU5 | NEDD4L | 2 | 2 | 0.01 | 0.01 |
| Q9NYL2 | MAP3K20 | 2 | 2 | 0.01 | 1.02 |
| Q6PD62 | CTR9 | 2 | 2 | 0.01 | 0.01 |
| Q99571 | P2RX4 | 2 | 2 | 0.01 | 0.57 |
| Q6IA69 | NADSYN1 | 2 | 2 | 0.01 | 0.01 |
| Q9H920 | RNF121 | 2 | 2 | 0.01 | 1.13 |
| P14854 | COX6B1 | 2 | 2 | 0.01 | 0.01 |
| Q6NXT6 | TAPT1 | 2 | 2 | 0.01 | 0.01 |
| P78316 | NOP14 | 2 | 2 | 0.01 | 0.01 |
| Q15796 | SMAD2 | 3 | 1 | 0.01 | 0.01 |
| Q13636 | RAB31 | 3 | 1 | 0.01 | 0.01 |
| Q9UL26 | RAB22A | 3 | 1 | 0.01 | 0.78 |
| P51911 | CNN1 | 3 | 1 | 0.01 | 1.02 |
| Q8IU85 | CAMK1D | 3 | 1 | 0.01 | 0.01 |
| O15439 | ABCC4 | 3 | 2 | 0.01 | 0.01 |
| Q96K21 | ZFYVE19 | 3 | 2 | 0.01 | 0.01 |
| Q5JSZ5 | PRRC2B | 3 | 2 | 0.01 | 1.14 |
| O43447 | PPIH | 3 | 2 | 0.01 | 0.93 |
| P45984 | MAPK9 | 3 | 2 | 0.01 | 0.01 |
| O14641 | DVL2 | 3 | 2 | 0.01 | 0.87 |
| Q9NR46 | SH3GLB2 | 3 | 2 | 0.01 | 0.01 |
| O60231 | DHX16 | 3 | 2 | 0.01 | 0.01 |
| P30711 | GSTT1 | 3 | 2 | 0.01 | 1.01 |
| Q8WWC4 | MAIP1 | 3 | 3 | 0.01 | 0.01 |
| Q6P6C2 | ALKBH5 | 3 | 3 | 0.01 | 0.01 |
| Q96S66 | CLCC1 | 3 | 3 | 0.01 | 0.01 |
| Q96GD0 | PDXP | 3 | 3 | 0.01 | 0.01 |
| Q99719 | SEPTIN5 | 3 | 3 | 0.01 | 1.29 |
| Q9Y676 | MRPS18B | 3 | 3 | 0.01 | 0.01 |
| Q8NI27 | THOC2 | 3 | 3 | 0.01 | 0.01 |
| P16035 | TIMP2 | 3 | 3 | 0.01 | 0.01 |
| Q92785 | DPF2 | 3 | 3 | 0.01 | 0.01 |
| Q9UBS8 | RNF14 | 3 | 3 | 0.01 | 0.80 |
| Q8IX01 | SUGP2 | 3 | 3 | 0.01 | 0.01 |
| Q96EY7 | PTCD3 | 3 | 3 | 0.01 | 0.01 |
| P48723 | HSPA13 | 3 | 3 | 0.01 | 0.73 |
| O14653 | GOSR2 | 3 | 3 | 0.01 | 1.09 |

| Accession | Name | # Peptides | # Unique Peptides | Abundance Ratio: day 3 / day 0 | Abundance Ratio: day 7/ day 0 |
|---|---|---|---|---|---|
| Q92845 | KIFAP3 | 2 | 1 | 0.01 | 0.97 |
| Q96BM9 | ARL8A | 2 | 1 | 0.01 | 0.76 |
| Q8N5I4 | DHRSX | 3 | 3 | 0.01 | 0.80 |
| Q9BYN8 | MRPS26 | 3 | 3 | 0.01 | 0.01 |
| P52815 | MRPL12 | 3 | 3 | 0.01 | 0.48 |
| Q96LJ7 | DHRS1 | 3 | 3 | 0.01 | 0.74 |
| Q9BVV7 | TIMM21 | 3 | 3 | 0.01 | 0.01 |
| Q7Z417 | NUFIP2 | 3 | 3 | 0.01 | 1.11 |
| Q13405 | MRPL49 | 3 | 3 | 0.01 | 0.01 |
| O00400 | SLC33A1 | 3 | 3 | 0.01 | 0.95 |
| P82932 | MRPS6 | 3 | 3 | 0.01 | 0.01 |
| Q01970 | PLCB3 | 3 | 3 | 0.01 | 0.01 |
| Q9H2F3 | HSD3B7 | 3 | 3 | 0.01 | 0.38 |
| Q29RF7 | PDS5A | 3 | 3 | 0.01 | 0.60 |
| Q8TER5 | ARHGEF40 | 3 | 3 | 0.01 | 0.01 |
| O95980 | RECK | 3 | 3 | 0.01 | 0.01 |
| Q15645 | TRIP13 | 3 | 3 | 0.01 | 0.47 |
| Q9P2W9 | STX18 | 3 | 3 | 0.01 | 0.01 |
| Q92917 | GPKOW | 3 | 3 | 0.01 | 0.85 |
| Q8TCS8 | PNPT1 | 3 | 3 | 0.01 | 0.01 |
| Q9UPQ0 | LIMCH1 | 3 | 3 | 0.01 | 1.46 |
| Q13523 | PRPF4B | 3 | 3 | 0.01 | 0.73 |
| Q9BXI6 | TBC1D10A | 3 | 3 | 0.01 | 0.01 |
| Q07889 | SOS1 | 3 | 3 | 0.01 | 0.01 |
| Q8NCC3 | PLA2G15 | 3 | 3 | 0.01 | 0.01 |
| O14972 | VPS26C | 3 | 3 | 0.01 | 0.01 |
| Q9NPF4 | OSGEP | 3 | 3 | 0.01 | 0.01 |
| Q9UJ68 | MSRA | 3 | 3 | 0.01 | 0.01 |
| O95347 | SMC2 | 3 | 3 | 0.01 | 0.01 |
| Q8WXX5 | DNAJC9 | 3 | 3 | 0.01 | 0.01 |
| Q8N129 | CNPY4 | 3 | 3 | 0.01 | 1.29 |
| Q53H96 | PYCR3 | 3 | 3 | 0.01 | 0.99 |
| P82933 | MRPS9 | 3 | 3 | 0.01 | 0.65 |
| Q9NRW7 | VPS45 | 3 | 3 | 0.01 | 1.52 |
| Q92805 | GOLGA1 | 3 | 3 | 0.01 | 0.01 |
| Q9H583 | HEATR1 | 3 | 3 | 0.01 | 1.11 |
| Q5T5U3 | ARHGAP21 | 3 | 3 | 0.01 | 0.01 |
| Q9UKX7 | NUP50 | 3 | 3 | 0.01 | 0.01 |
| Q9Y673 | ALG5 | 3 | 3 | 0.01 | 0.90 |
| Q9BST9 | RTKN | 3 | 3 | 0.01 | 2.32 |
| Q13042 | CDC16 | 3 | 3 | 0.01 | 0.69 |
| Q9H061 | TMEM126A | 3 | 3 | 0.01 | 0.01 |
| Q8WV41 | SNX33 | 3 | 3 | 0.01 | 0.01 |

| Accession | Name | # Peptides | # Unique Peptides | Abundance Ratio: day 3 / day 0 | Abundance Ratio: day 7/ day 0 |
|---|---|---|---|---|---|
| Q92845 | KIFAP3 | 2 | 1 | 0.01 | 0.97 |
| Q96BM9 | ARL8A | 2 | 1 | 0.01 | 0.76 |
| P51553 | IDH3G | 3 | 3 | 0.01 | 0.39 |
| Q9NS00 | C1GALT1 | 3 | 3 | 0.01 | 0.01 |
| Q96EY1 | DNAJA3 | 3 | 3 | 0.01 | 0.60 |
| Q8TD16 | BICD2 | 3 | 3 | 0.01 | 0.01 |
| P04150 | NR3C1 | 3 | 3 | 0.01 | 0.01 |
| O00391 | QSOX1 | 3 | 3 | 0.01 | 0.01 |
| Q8IWU6 | SULF1 | 3 | 3 | 0.01 | 0.01 |
| Q9UKJ3 | GPATCH8 | 3 | 3 | 0.01 | 0.01 |
| Q9UJM3 | ERRFI1 | 3 | 3 | 0.01 | 0.01 |
| Q3ZCQ8 | TIMM50 | 3 | 3 | 0.01 | 0.01 |
| O15126 | SCAMP1 | 3 | 3 | 0.01 | 0.79 |
| Q9NQ50 | MRPL40 | 3 | 3 | 0.01 | 0.01 |
| Q9UBP9 | GULP1 | 3 | 3 | 0.01 | 0.01 |
| Q8TBX8 | PIP4K2C | 3 | 3 | 0.01 | 0.76 |
| P35269 | GTF2F1 | 3 | 3 | 0.01 | 0.01 |
| Q9BW27 | NUP85 | 3 | 3 | 0.01 | 0.01 |
| Q16795 | NDUFA9 | 3 | 3 | 0.01 | 0.01 |
| Q02040 | AKAP17A | 3 | 3 | 0.01 | 0.01 |
| Q15427 | SF3B4 | 3 | 3 | 0.01 | 0.01 |
| P19320 | VCAM1 | 3 | 3 | 0.01 | 0.01 |
| Q9C0C9 | UBE2O | 3 | 3 | 0.01 | 0.90 |
| Q9UKF6 | CPSF3 | 3 | 3 | 0.01 | 0.01 |
| Q9Y4K4 | MAP4K5 | 3 | 3 | 0.01 | 0.80 |
| Q9BYD1 | MRPL13 | 3 | 3 | 0.01 | 0.01 |
| Q9UNK0 | STX8 | 3 | 3 | 0.01 | 0.01 |
| Q9H8M7 | MINDY3 | 3 | 3 | 0.01 | 0.01 |
| P82650 | MRPS22 | 3 | 3 | 0.01 | 0.01 |
| Q96SY0 | INTS14 | 3 | 3 | 0.01 | 0.01 |
| Q7Z6M1 | RABEPK | 3 | 3 | 0.01 | 0.01 |
| P78345 | RPP38 | 3 | 3 | 0.01 | 0.45 |
| Q16740 | CLPP | 3 | 3 | 0.01 | 0.01 |
| Q15773 | MLF2 | 3 | 3 | 0.01 | 0.01 |
| Q8WTW3 | COG1 | 3 | 3 | 0.01 | 1.12 |
| Q8TBM8 | DNAJB14 | 3 | 3 | 0.01 | 0.01 |
| Q16644 | MAPKAPK3 | 4 | 1 | 0.01 | 0.01 |
| Q12873 | CHD3 | 4 | 1 | 0.01 | 0.01 |
| P04049 | RAF1 | 4 | 2 | 0.01 | 0.01 |
| Q13884 | SNTB1 | 4 | 2 | 0.01 | 0.01 |
| P11233 | RALA | 4 | 3 | 0.01 | 0.64 |
| Q92871 | PMM1 | 4 | 3 | 0.01 | 0.01 |
| Q9Y5V3 | MAGED1 | 4 | 3 | 0.01 | 1.37 |

| Accession | Name | # Peptides | # Unique Peptides | Abundance Ratio: day 3 / day 0 | Abundance Ratio: day 7/ day 0 |
|---|---|---|---|---|---|
| Q92845 | KIFAP3 | 2 | 1 | 0.01 | 0.97 |
| Q96BM9 | ARL8A | 2 | 1 | 0.01 | 0.76 |
| P42285 | MTREX | 4 | 3 | 0.01 | 0.01 |
| Q7Z7M9 | GALNT5 | 4 | 3 | 0.01 | 0.01 |
| Q13033 | STRN3 | 4 | 3 | 0.01 | 1.07 |
| Q9BZ67 | FRMD8 | 4 | 4 | 0.01 | 0.01 |
| Q96S55 | WRNIP1 | 4 | 4 | 0.01 | 0.01 |
| Q13546 | RIPK1 | 4 | 4 | 0.01 | 1.28 |
| P42356 | PI4KA | 4 | 4 | 0.01 | 0.79 |
| O60343 | TBC1D4 | 4 | 4 | 0.01 | 0.01 |
| P30260 | CDC27 | 4 | 4 | 0.01 | 0.01 |
| P23368 | ME2 | 4 | 4 | 0.01 | 0.01 |
| Q96PU4 | UHRF2 | 4 | 4 | 0.01 | 0.01 |
| Q9GZT3 | SLIRP | 4 | 4 | 0.01 | 0.01 |
| Q7L5D6 | GET4 | 4 | 4 | 0.01 | 0.01 |
| P23786 | CPT2 | 4 | 4 | 0.01 | 0.01 |
| Q9H845 | ACAD9 | 4 | 4 | 0.01 | 0.68 |
| P11166 | SLC2A1 | 4 | 4 | 0.01 | 0.01 |
| Q14393 | GAS6 | 4 | 4 | 0.01 | 0.01 |
| Q96JJ3 | ELMO2 | 4 | 4 | 0.01 | 0.01 |
| Q9Y315 | DERA | 4 | 4 | 0.01 | 0.99 |
| Q13443 | ADAM9 | 4 | 4 | 0.01 | 0.01 |
| P53814 | SMTN | 4 | 4 | 0.01 | 1.10 |
| O14786 | NRP1 | 4 | 4 | 0.01 | 0.01 |
| Q9UKU9 | ANGPTL2 | 4 | 4 | 0.01 | 1.02 |
| Q13683 | ITGA7 | 4 | 4 | 0.01 | 0.01 |
| Q86XA9 | HEATR5A | 4 | 4 | 0.01 | 0.01 |
| O96005 | CLPTM1 | 4 | 4 | 0.01 | 1.10 |
| O14672 | ADAM10 | 4 | 4 | 0.01 | 0.67 |
| P01137 | TGFB1 | 4 | 4 | 0.01 | 0.01 |
| O76074 | PDE5A | 4 | 4 | 0.01 | 0.01 |
| Q03518 | TAP1 | 4 | 4 | 0.01 | 0.52 |
| Q96E11 | MRRF | 4 | 4 | 0.01 | 0.01 |
| Q8TDD1 | DDX54 | 4 | 4 | 0.01 | 1.07 |
| Q9UKM7 | MAN1B1 | 4 | 4 | 0.01 | 0.72 |
| Q6P9B6 | MEAK7 | 4 | 4 | 0.01 | 0.95 |
| Q12983 | BNIP3 | 4 | 4 | 0.01 | 0.01 |
| Q86U38 | NOP9 | 4 | 4 | 0.01 | 1.08 |
| Q5JSH3 | WDR44 | 4 | 4 | 0.01 | 0.01 |
| Q5T5P2 | KIAA1217 | 4 | 4 | 0.01 | 0.01 |
| Q9UQB8 | BAIAP2 | 4 | 4 | 0.01 | 0.01 |
| P46087 | NOP2 | 4 | 4 | 0.01 | 0.01 |
| P58335 | ANTXR2 | 4 | 4 | 0.01 | 0.01 |

| Accession | Name | # Peptides | # Unique Peptides | Abundance Ratio: day 3 / day 0 | Abundance Ratio: day 7/ day 0 |
|---|---|---|---|---|---|
| Q92845 | KIFAP3 | 2 | 1 | 0.01 | 0.97 |
| Q96BM9 | ARL8A | 2 | 1 | 0.01 | 0.76 |
| O75695 | RP2 | 4 | 4 | 0.01 | 0.01 |
| Q9C0B1 | FTO | 4 | 4 | 0.01 | 0.93 |
| P30038 | ALDH4A1 | 4 | 4 | 0.01 | 0.01 |
| P49768 | PSEN1 | 4 | 4 | 0.01 | 0.01 |
| P56556 | NDUFA6 | 4 | 4 | 0.01 | 0.01 |
| O14920 | IKBKB | 4 | 4 | 0.01 | 0.01 |
| Q9UHW9 | SLC12A6 | 5 | 1 | 0.01 | 0.01 |
| Q9UPY8 | MAPRE3 | 5 | 2 | 0.01 | 0.01 |
| Q9Y5A9 | YTHDF2 | 5 | 2 | 0.01 | 1.33 |
| Q15437 | SEC23B | 5 | 3 | 0.01 | 0.96 |
| P08727 | KRT19 | 5 | 3 | 0.01 | 0.01 |
| P41221 | WNT5A | 5 | 3 | 0.01 | 0.01 |
| Q7L7X3 | TAOK1 | 5 | 4 | 0.01 | 0.97 |
| P29317 | EPHA2 | 5 | 4 | 0.01 | 0.01 |
| Q05655 | PRKCD | 5 | 4 | 0.01 | 0.71 |
| Q8N392 | ARHGAP18 | 5 | 5 | 0.01 | 0.01 |
| Q03426 | MVK | 5 | 5 | 0.01 | 0.01 |
| Q12849 | GRSF1 | 5 | 5 | 0.01 | 0.01 |
| Q9NS15 | LTBP3 | 5 | 5 | 0.01 | 1.17 |
| Q9H2U1 | DHX36 | 5 | 5 | 0.01 | 1.01 |
| O43896 | KIF1C | 5 | 5 | 0.01 | 0.01 |
| Q9Y223 | GNE | 5 | 5 | 0.01 | 0.81 |
| Q9BW91 | NUDT9 | 5 | 5 | 0.01 | 0.94 |
| Q13136 | PPFIA1 | 5 | 5 | 0.01 | 0.74 |
| Q5JVF3 | PCID2 | 5 | 5 | 0.01 | 0.01 |
| Q14669 | TRIP12 | 5 | 5 | 0.01 | 0.01 |
| Q7Z4N8 | P4HA3 | 5 | 5 | 0.01 | 0.88 |
| Q12788 | TBL3 | 5 | 5 | 0.01 | 0.01 |
| Q9NXF1 | TEX10 | 5 | 5 | 0.01 | 0.01 |
| P10398 | ARAF | 6 | 4 | 0.01 | 0.87 |
| Q96SB4 | SRPK1 | 6 | 5 | 0.01 | 0.01 |
| Q5T5C0 | STXBP5 | 6 | 6 | 0.01 | 0.01 |
| Q9Y282 | ERGIC3 | 6 | 6 | 0.01 | 0.83 |
| Q9H936 | SLC25A22 | 6 | 6 | 0.01 | 0.57 |
| P33992 | MCM5 | 6 | 6 | 0.01 | 0.01 |
| O15321 | TM9SF1 | 6 | 6 | 0.01 | 1.00 |
| Q53H12 | AGK | 6 | 6 | 0.01 | 0.01 |
| Q8IZH2 | XRN1 | 6 | 6 | 0.01 | 0.01 |
| Q9H7D0 | DOCK5 | 6 | 6 | 0.01 | 0.01 |
| Q9Y6E0 | STK24 | 7 | 2 | 0.01 | 0.84 |
| P31947 | SFN | 7 | 4 | 0.01 | 0.01 |

| Accession | Name | # Peptides | # Unique Peptides | Abundance Ratio: day 3 / day 0 | Abundance Ratio: day 7/ day 0 |
|---|---|---|---|---|---|
| Q92845 | KIFAP3 | 2 | 1 | 0.01 | 0.97 |
| Q96BM9 | ARL8A | 2 | 1 | 0.01 | 0.76 |
| Q86VI3 | IQGAP3 | 7 | 4 | 0.01 | 0.01 |
| Q13595 | TRA2A | 7 | 5 | 0.01 | 0.01 |
| O60826 | CCDC22 | 7 | 6 | 0.01 | 0.68 |
| Q15599 | SLC9A3R2 | 7 | 7 | 0.01 | 0.53 |
| P56199 | ITGA1 | 7 | 7 | 0.01 | 0.95 |
| Q14978 | NOLC1 | 7 | 7 | 0.01 | 1.84 |
| P16298 | PPP3CB | 8 | 2 | 0.01 | 0.01 |
| P51116 | FXR2 | 8 | 4 | 0.01 | 0.70 |
| Q06787 | FMR1 | 9 | 6 | 0.01 | 0.79 |
| P33993 | MCM7 | 9 | 9 | 0.01 | 0.01 |
| P36873 | PPP1CC | 10 | 1 | 0.01 | 1.01 |
| Q86TI2 | DPP9 | 10 | 10 | 0.01 | 0.83 |
| Q14690 | PDCD11 | 10 | 10 | 0.01 | 1.02 |
| Q9UNX3 | RPL26L1 | 11 | 1 | 0.01 | 0.01 |
| P22090 | RPS4Y1 | 11 | 2 | 0.01 | 0.01 |
| Q63HN8 | RNF213 | 11 | 11 | 0.01 | 0.46 |
| P00533 | EGFR | 12 | 11 | 0.01 | 0.01 |
| Q5T9A4 | ATAD3B | 16 | 2 | 0.01 | 0.01 |
| P17936 | IGFBP3 | 17 | 17 | 0.10 | 0.10 |
| P01023 | A2M | 6 | 6 | 0.25 | 0.12 |
| Q9HCJ1 | ANKH | 4 | 4 | 0.36 | 0.28 |
| Q8IVL5 | P3H2 | 11 | 11 | 0.41 | 0.44 |
| Q9Y572 | RIPK3 | 2 | 2 | 0.41 | 0.22 |
| Q01995 | TAGLN | 17 | 17 | 0.41 | 0.33 |
| P49736 | MCM2 | 3 | 3 | 0.42 | 0.32 |
| P26006 | ITGA3 | 6 | 6 | 0.42 | 0.01 |
| P50416 | CPT1A | 11 | 11 | 0.46 | 0.31 |
| P29966 | MARCKS | 10 | 10 | 0.47 | 0.67 |
| Q9H694 | BICC1 | 2 | 2 | 0.47 | 0.46 |
| Q10713 | PMPCA | 4 | 4 | 0.48 | 0.40 |
| Q14566 | MCM6 | 6 | 6 | 0.48 | 0.01 |
| O60637 | TSPAN3 | 2 | 2 | 0.48 | 0.36 |
| P28300 | LOX | 5 | 5 | 0.48 | 0.01 |
| Q8WWI1 | LMO7 | 52 | 52 | 0.48 | 0.28 |
| P05362 | ICAM1 | 15 | 15 | 0.49 | 0.63 |
| Q9BX68 | HINT2 | 3 | 3 | 0.49 | 0.52 |
| Q93062 | RBPMS | 3 | 3 | 0.49 | 0.01 |
| O94885 | SASH1 | 6 | 6 | 0.49 | 0.53 |
| Q8IVF2 | AHNAK2 | 41 | 39 | 0.49 | 0.37 |
| Q9HD67 | MYO10 | 2 | 2 | 0.50 | 0.59 |
| P26440 | IVD | 8 | 8 | 0.50 | 0.31 |

| Accession | Name | # Peptides | # Unique Peptides | Abundance Ratio: day 3 / day 0 | Abundance Ratio: day 7/ day 0 |
|---|---|---|---|---|---|
| Q92845 | KIFAP3 | 2 | 1 | 0.01 | 0.97 |
| Q96BM9 | ARL8A | 2 | 1 | 0.01 | 0.76 |
| Q9H1K1 | ISCU | 3 | 3 | 0.50 | 0.01 |
| P58215 | LOXL3 | 8 | 8 | 0.50 | 0.72 |
| P06493 | CDK1 | 3 | 2 | 0.51 | 0.01 |
| P27449 | ATP6V0C | 2 | 2 | 0.51 | 0.36 |
| O94925 | GLS | 11 | 11 | 0.51 | 0.51 |
| Q9NRX5 | SERINC1 | 3 | 3 | 0.52 | 0.43 |
| Q9Y4K0 | LOXL2 | 11 | 11 | 0.52 | 0.91 |
| P22692 | IGFBP4 | 3 | 3 | 0.52 | 0.01 |
| P15586 | GNS | 9 | 9 | 0.53 | 0.34 |
| Q5T653 | MRPL2 | 2 | 2 | 0.53 | 0.01 |
| P27701 | CD82 | 2 | 2 | 0.53 | 0.46 |
| Q9BRK3 | MXRA8 | 8 | 8 | 0.53 | 0.64 |
| Q15417 | CNN3 | 12 | 9 | 0.53 | 0.57 |
| P08574 | CYC1 | 4 | 4 | 0.54 | 0.41 |
| Q9P2R7 | SUCLA2 | 12 | 12 | 0.54 | 0.40 |
| Q9NUM4 | TMEM106B | 3 | 3 | 0.54 | 0.01 |
| P00367 | GLUD1 | 24 | 24 | 0.54 | 0.38 |
| Q12805 | EFEMP1 | 8 | 8 | 0.54 | 0.45 |
| Q8IY17 | PNPLA6 | 5 | 5 | 0.54 | 0.76 |
| Q9NTJ3 | SMC4 | 2 | 2 | 0.55 | 0.49 |
| Q9H098 | FAM107B | 5 | 5 | 0.55 | 0.71 |
| P09914 | IFIT1 | 6 | 5 | 0.55 | 0.01 |
| Q460N5 | PARP14 | 2 | 2 | 0.55 | 0.57 |
| P53597 | SUCLG1 | 8 | 8 | 0.55 | 0.45 |
| O75390 | CS | 12 | 12 | 0.56 | 0.39 |
| P11117 | ACP2 | 4 | 4 | 0.56 | 0.47 |
| P30048 | PRDX3 | 7 | 7 | 0.56 | 0.37 |
| P52789 | HK2 | 16 | 12 | 0.56 | 0.51 |
| Q02952 | AKAP12 | 50 | 50 | 0.56 | 0.26 |
| P10809 | HSPD1 | 26 | 26 | 0.56 | 0.49 |
| P25205 | MCM3 | 7 | 7 | 0.57 | 0.01 |
| Q8TAE8 | GADD45GIP1 | 2 | 2 | 0.57 | 0.01 |
| Q9UGV2 | NDRG3 | 2 | 2 | 0.57 | 0.01 |
| Q9Y6N5 | SQOR | 19 | 19 | 0.57 | 0.29 |
| Q6QNY0 | BLOC1S3 | 2 | 2 | 0.57 | 0.01 |
| Q9H6B4 | CLMP | 4 | 4 | 0.57 | 0.01 |
| P05161 | ISG15 | 3 | 3 | 0.57 | 0.01 |
| P24752 | ACAT1 | 11 | 11 | 0.57 | 0.33 |
| Q99439 | CNN2 | 13 | 11 | 0.58 | 0.45 |
| Q96N67 | DOCK7 | 10 | 9 | 0.58 | 0.62 |
| Q9BV86 | NTMT1 | 2 | 2 | 0.58 | 0.47 |

| Accession | Name | # Peptides | # Unique Peptides | Abundance Ratio: day 3 / day 0 | Abundance Ratio: day 7/ day 0 |
|---|---|---|---|---|---|
| Q92845 | KIFAP3 | 2 | 1 | 0.01 | 0.97 |
| Q96BM9 | ARL8A | 2 | 1 | 0.01 | 0.76 |
| O15013 | ARHGEF10 | 2 | 2 | 0.58 | 0.83 |
| Q14699 | RFTN1 | 5 | 5 | 0.58 | 0.01 |
| Q6WCQ1 | MPRIP | 11 | 10 | 0.58 | 0.56 |
| Q9HA77 | CARS2 | 6 | 6 | 0.59 | 0.29 |
| Q8IZ07 | ANKRD13A | 3 | 3 | 0.59 | 0.01 |
| O15118 | NPC1 | 8 | 8 | 0.59 | 0.48 |
| P40926 | MDH2 | 15 | 15 | 0.59 | 0.43 |
| O95379 | TNFAIP8 | 3 | 3 | 0.59 | 0.74 |
| O96008 | TOMM40 | 2 | 2 | 0.59 | 0.48 |
| P55809 | OXCT1 | 12 | 12 | 0.59 | 0.43 |
| Q5T8D3 | ACBD5 | 3 | 3 | 0.60 | 0.46 |
| Q9H7C4 | SYNC | 6 | 6 | 0.60 | 0.56 |
| Q9H6X2 | ANTXR1 | 4 | 4 | 0.60 | 0.64 |
| P42126 | ECI1 | 5 | 5 | 0.60 | 0.51 |
| P20962 | PTMS | 4 | 4 | 0.61 | 0.35 |
| P11177 | PDHB | 8 | 8 | 0.61 | 0.52 |
| Q9NS69 | TOMM22 | 3 | 3 | 0.61 | 0.58 |
| Q8N3D4 | EHBP1L1 | 10 | 10 | 0.61 | 0.64 |
| Q5RI15 | COX20 | 2 | 2 | 0.61 | 0.01 |
| Q16832 | DDR2 | 8 | 8 | 0.61 | 0.01 |
| P05141 | SLC25A5 | 13 | 5 | 0.61 | 0.48 |
| Q8IWB1 | ITPRIP | 5 | 5 | 0.61 | 0.01 |
| O75521 | ECI2 | 6 | 6 | 0.61 | 0.35 |
| Q92542 | NCSTN | 2 | 2 | 0.61 | 0.51 |
| O00330 | PDHX | 3 | 3 | 0.61 | 0.44 |
| Q15847 | ADIRF | 6 | 6 | 0.61 | 0.33 |
| Q9Y2D5 | AKAP2 | 15 | 15 | 0.62 | 0.36 |
| Q10471 | GALNT2 | 17 | 15 | 0.62 | 0.90 |
| Q04837 | SSBP1 | 6 | 6 | 0.62 | 0.52 |
| Q8NE86 | MCU | 8 | 8 | 0.62 | 0.44 |
| P05106 | ITGB3 | 2 | 2 | 0.62 | 0.01 |
| P50281 | MMP14 | 7 | 7 | 0.62 | 0.47 |
| P01210 | PENK | 9 | 9 | 0.62 | 0.39 |
| P43897 | TSFM | 2 | 2 | 0.62 | 0.38 |
| Q9ULC3 | RAB23 | 6 | 6 | 0.62 | 0.41 |
| P12236 | SLC25A6 | 14 | 2 | 0.62 | 0.46 |
| Q15746 | MYLK | 15 | 15 | 0.62 | 0.58 |
| O95571 | ETHE1 | 5 | 5 | 0.63 | 0.48 |
| P30084 | ECHS1 | 13 | 13 | 0.63 | 0.51 |
| Q9P0I2 | EMC3 | 4 | 4 | 0.63 | 0.52 |
| Q53H82 | LACTB2 | 3 | 3 | 0.63 | 0.01 |

| Accession | Name | # Peptides | # Unique Peptides | Abundance Ratio: day 3 / day 0 | Abundance Ratio: day 7/ day 0 |
|---|---|---|---|---|---|
| Q92845 | KIFAP3 | 2 | 1 | 0.01 | 0.97 |
| Q96BM9 | ARL8A | 2 | 1 | 0.01 | 0.76 |
| Q7Z3B1 | NEGR1 | 7 | 7 | 0.63 | 0.37 |
| P32456 | GBP2 | 9 | 8 | 0.63 | 0.81 |
| P42224 | STAT1 | 28 | 28 | 0.63 | 0.45 |
| Q9H773 | DCTPP1 | 3 | 3 | 0.63 | 0.50 |
| Q7Z478 | DHX29 | 3 | 3 | 0.63 | 0.63 |
| P38646 | HSPA9 | 40 | 39 | 0.63 | 0.58 |
| Q9Y287 | ITM2B | 4 | 4 | 0.63 | 0.52 |
| P05556 | ITGB1 | 25 | 25 | 0.63 | 0.53 |
| P46199 | MTIF2 | 2 | 2 | 0.63 | 0.44 |
| Q15024 | EXOSC7 | 3 | 3 | 0.63 | 0.63 |
| Q96DB5 | RMDN1 | 6 | 6 | 0.63 | 0.40 |
| P36639 | NUDT1 | 4 | 4 | 0.63 | 0.87 |
| P21796 | VDAC1 | 15 | 15 | 0.63 | 0.51 |
| Q9BT09 | CNPY3 | 2 | 2 | 0.63 | 0.65 |
| O14879 | IFIT3 | 3 | 3 | 0.63 | 0.39 |
| P17301 | ITGA2 | 10 | 10 | 0.63 | 0.26 |
| P15848 | ARSB | 10 | 10 | 0.63 | 0.36 |
| P61604 | HSPE1 | 10 | 10 | 0.63 | 0.54 |
| Q96HS1 | PGAM5 | 4 | 4 | 0.64 | 0.47 |
| Q9UII2 | ATP5IF1 | 5 | 5 | 0.64 | 0.47 |
| Q14192 | FHL2 | 13 | 13 | 0.64 | 0.39 |
| Q5GLZ8 | HERC4 | 17 | 17 | 0.64 | 0.44 |
| P82930 | MRPS34 | 3 | 3 | 0.64 | 0.01 |
| Q14318 | FKBP8 | 10 | 10 | 0.64 | 0.67 |
| P09001 | MRPL3 | 3 | 3 | 0.64 | 0.45 |
| O00483 | NDUFA4 | 3 | 3 | 0.64 | 0.44 |
| Q9BWM7 | SFXN3 | 8 | 8 | 0.64 | 0.47 |
| Q9NP74 | PALMD | 4 | 4 | 0.64 | 0.56 |
| O43760 | SYNGR2 | 2 | 2 | 0.64 | 0.01 |
| Q6Y1H2 | HACD2 | 2 | 2 | 0.64 | 0.80 |
| O14495 | PLPP3 | 2 | 2 | 0.64 | 0.01 |
| Q12929 | EPS8 | 13 | 13 | 0.64 | 0.52 |
| P09622 | DLD | 14 | 14 | 0.64 | 0.52 |
| P08559 | PDHA1 | 15 | 15 | 0.64 | 0.60 |
| O43665 | RGS10 | 2 | 2 | 0.64 | 0.90 |
| Q13586 | STIM1 | 4 | 4 | 0.64 | 0.01 |
| Q92598 | HSPH1 | 22 | 19 | 0.64 | 0.64 |
| P42704 | LRPPRC | 27 | 27 | 0.64 | 0.61 |
| Q99714 | HSD17B10 | 6 | 6 | 0.65 | 0.64 |
| Q00325 | SLC25A3 | 13 | 13 | 0.65 | 0.50 |
| Q8TB61 | SLC35B2 | 3 | 3 | 0.65 | 0.86 |

| Accession | Name | # Peptides | # Unique Peptides | Abundance Ratio: day 3 / day 0 | Abundance Ratio: day 7/ day 0 |
|---|---|---|---|---|---|
| Q92845 | KIFAP3 | 2 | 1 | 0.01 | 0.97 |
| Q96BM9 | ARL8A | 2 | 1 | 0.01 | 0.76 |
| Q9BTY2 | FUCA2 | 6 | 6 | 0.65 | 0.66 |
| P45880 | VDAC2 | 12 | 12 | 0.65 | 0.45 |
| P43304 | GPD2 | 21 | 21 | 0.65 | 0.52 |
| P15151 | PVR | 3 | 3 | 0.65 | 0.49 |
| P07954 | FH | 15 | 15 | 0.65 | 0.62 |
| P35527 | KRT9 | 11 | 10 | 0.65 | 0.65 |
| P19404 | NDUFV2 | 3 | 3 | 0.66 | 0.01 |
| P02786 | TFRC | 17 | 17 | 0.66 | 0.70 |
| P49411 | TUFM | 15 | 15 | 0.66 | 0.53 |
| Q9BPW8 | NIPSNAP1 | 5 | 3 | 0.66 | 0.69 |
| Q13637 | RAB32 | 5 | 5 | 0.66 | 0.63 |
| P00505 | GOT2 | 19 | 19 | 0.66 | 0.49 |
| O00217 | NDUFS8 | 2 | 2 | 0.66 | 0.01 |
| Q9Y305 | ACOT9 | 6 | 6 | 0.66 | 0.62 |
| Q8NBK3 | SUMF1 | 3 | 3 | 0.66 | 1.52 |
| O15269 | SPTLC1 | 11 | 11 | 0.66 | 0.61 |
| P16219 | ACADS | 2 | 2 | 0.66 | 0.45 |
| Q99747 | NAPG | 9 | 9 | 0.66 | 0.57 |
| Q9BQS8 | FYCO1 | 18 | 18 | 0.66 | 0.50 |
| Q9NUI1 | DECR2 | 9 | 9 | 0.67 | 0.01 |
| O43615 | TIMM44 | 12 | 12 | 0.67 | 0.56 |
| Q6PIU2 | NCEH1 | 7 | 7 | 0.67 | 0.32 |
| P08648 | ITGA5 | 15 | 15 | 0.67 | 0.44 |
| Q15651 | HMGN3 | 3 | 3 | 0.67 | 0.01 |
| Q15785 | TOMM34 | 4 | 4 | 0.67 | 0.67 |
| Q86WV6 | STING1 | 8 | 8 | 0.67 | 0.58 |
| O43837 | IDH3B | 4 | 4 | 0.67 | 0.01 |
| Q9UMX0 | UBQLN1 | 6 | 3 | 0.67 | 0.62 |
| P83111 | LACTB | 6 | 6 | 0.67 | 0.01 |
| Q96C19 | EFHD2 | 11 | 11 | 0.67 | 0.43 |
| Q02218 | OGDH | 16 | 10 | 0.67 | 0.60 |
| P07093 | SERPINE2 | 12 | 12 | 0.67 | 0.46 |
| Q8IYB5 | SMAP1 | 3 | 3 | 0.67 | 0.01 |
| P10619 | CTSA | 10 | 10 | 0.67 | 0.43 |
| Q5RKV6 | EXOSC6 | 4 | 4 | 0.68 | 0.01 |
| Q13671 | RIN1 | 6 | 6 | 0.68 | 0.01 |
| Q9UG63 | ABCF2 | 9 | 9 | 0.68 | 0.88 |
| P22307 | SCP2 | 14 | 14 | 0.68 | 0.51 |
| Q5ZPR3 | CD276 | 4 | 4 | 0.68 | 0.54 |
| P48509 | CD151 | 6 | 6 | 0.68 | 0.40 |
| P34897 | SHMT2 | 16 | 16 | 0.68 | 0.65 |

| Accession | Name | # Peptides | # Unique Peptides | Abundance Ratio: day 3 / day 0 | Abundance Ratio: day 7/ day 0 |
|---|---|---|---|---|---|
| Q92845 | KIFAP3 | 2 | 1 | 0.01 | 0.97 |
| Q96BM9 | ARL8A | 2 | 1 | 0.01 | 0.76 |
| Q9HC07 | TMEM165 | 2 | 2 | 0.68 | 0.98 |
| Q6YN16 | HSDL2 | 7 | 7 | 0.68 | 0.01 |
| Q9Y277 | VDAC3 | 11 | 11 | 0.68 | 0.47 |
| P82914 | MRPS15 | 2 | 2 | 0.68 | 0.01 |
| P06454 | PTMA | 5 | 5 | 0.68 | 0.53 |
| O43674 | NDUFB5 | 2 | 2 | 0.68 | 0.73 |
| Q9NXR1 | NDE1 | 3 | 2 | 0.68 | 0.01 |
| Q13526 | PIN1 | 4 | 4 | 0.68 | 0.62 |
| P11310 | ACADM | 7 | 7 | 0.68 | 0.01 |
| P07711 | CTSL | 8 | 7 | 0.68 | 0.32 |
| Q6UB35 | MTHFD1L | 8 | 7 | 0.68 | 0.63 |
| Q5JRX3 | PITRM1 | 27 | 27 | 0.68 | 0.54 |
| Q9UI09 | NDUFA12 | 3 | 3 | 0.68 | 0.48 |
| Q5UCC4 | EMC10 | 3 | 3 | 0.68 | 0.67 |
| Q13336 | SLC14A1 | 6 | 6 | 0.68 | 0.53 |
| Q99623 | PHB2 | 20 | 20 | 0.68 | 0.54 |
| O95299 | NDUFA10 | 5 | 5 | 0.68 | 0.53 |
| P07738 | BPGM | 9 | 9 | 0.68 | 0.44 |
| Q9H0W9 | C11orf54 | 4 | 4 | 0.68 | 0.72 |
| O43678 | NDUFA2 | 6 | 6 | 0.68 | 0.59 |
| O75381 | PEX14 | 6 | 6 | 0.68 | 0.86 |
| O00154 | ACOT7 | 11 | 11 | 0.68 | 0.39 |
| Q9BSJ8 | ESYT1 | 22 | 21 | 0.68 | 0.45 |
| Q9BYD3 | MRPL4 | 2 | 2 | 0.68 | 0.46 |
| Q9H9S4 | CAB39L | 3 | 2 | 0.69 | 0.54 |
| Q8NFQ8 | TOR1AIP2 | 5 | 4 | 0.69 | 0.01 |
| O00148 | DDX39A | 14 | 3 | 0.69 | 0.62 |
| Q9NVI7 | ATAD3A | 20 | 6 | 0.69 | 0.50 |
| Q9BV57 | ADI1 | 4 | 4 | 0.69 | 0.71 |
| P08962 | CD63 | 4 | 4 | 0.69 | 0.57 |
| Q9BTE6 | AARSD1 | 5 | 5 | 0.69 | 0.89 |
| Q9UDT6 | CLIP2 | 12 | 7 | 0.69 | 0.62 |
| P21291 | CSRP1 | 10 | 10 | 0.69 | 0.51 |
| Q02978 | SLC25A11 | 12 | 12 | 0.69 | 0.59 |
| O43314 | PPIP5K2 | 4 | 4 | 0.69 | 0.01 |
| Q8WXF7 | ATL1 | 7 | 6 | 0.69 | 0.78 |
| Q9NWU2 | GID8 | 2 | 2 | 0.69 | 0.78 |
| Q16563 | SYPL1 | 3 | 3 | 0.69 | 0.59 |
| O75306 | NDUFS2 | 8 | 8 | 0.69 | 0.56 |
| Q9Y2Z4 | YARS2 | 2 | 2 | 0.69 | 0.01 |
| Q9BYX2 | TBC1D2 | 4 | 4 | 0.69 | 0.01 |

| Accession | Name | # Peptides | # Unique Peptides | Abundance Ratio: day 3 / day 0 | Abundance Ratio: day 7/ day 0 |
|---|---|---|---|---|---|
| Q92845 | KIFAP3 | 2 | 1 | 0.01 | 0.97 |
| Q96BM9 | ARL8A | 2 | 1 | 0.01 | 0.76 |
| Q9UFN0 | NIPSNAP3A | 4 | 4 | 0.69 | 0.65 |
| P11279 | LAMP1 | 5 | 5 | 0.69 | 0.55 |
| Q15836 | VAMP3 | 7 | 4 | 0.70 | 0.54 |
| Q6P2E9 | EDC4 | 9 | 9 | 0.70 | 0.77 |
| Q99798 | ACO2 | 18 | 18 | 0.70 | 0.62 |
| Q9UIJ7 | AK3 | 12 | 12 | 0.70 | 0.55 |
| Q7L1Q6 | BZW1 | 18 | 18 | 0.70 | 0.63 |
| P22695 | UQCRC2 | 19 | 19 | 0.70 | 0.55 |
| Q6P1M0 | SLC27A4 | 2 | 2 | 0.70 | 0.54 |
| P20645 | M6PR | 6 | 6 | 0.70 | 0.66 |
| Q13620 | CUL4B | 25 | 17 | 0.70 | 0.53 |
| Q8NEY1 | NAV1 | 5 | 5 | 0.70 | 0.01 |
| P19367 | HK1 | 38 | 34 | 0.70 | 0.54 |
| Q9H3U5 | MFSD1 | 2 | 2 | 0.70 | 0.01 |
| Q9NZJ7 | MTCH1 | 5 | 5 | 0.70 | 0.53 |
| P04181 | OAT | 9 | 9 | 0.70 | 1.04 |
| P28331 | NDUFS1 | 13 | 13 | 0.70 | 0.53 |
| P16949 | STMN1 | 15 | 15 | 0.70 | 0.39 |
| Q96EU6 | RRP36 | 2 | 2 | 0.70 | 1.03 |
| P31937 | HIBADH | 7 | 7 | 0.70 | 0.46 |
| Q96I99 | SUCLG2 | 16 | 16 | 0.70 | 0.52 |
| O60684 | KPNA6 | 9 | 7 | 0.70 | 0.79 |
| Q16698 | DECR1 | 9 | 9 | 0.70 | 0.57 |
| O95819 | MAP4K4 | 17 | 14 | 0.70 | 0.53 |
| P05091 | ALDH2 | 16 | 13 | 0.70 | 0.60 |
| Q969N2 | PIGT | 9 | 9 | 0.70 | 0.73 |
| O95573 | ACSL3 | 13 | 12 | 0.70 | 0.58 |
| P21333 | FLNA | 152 | 136 | 0.70 | 0.54 |
| Q9Y2R0 | COA3 | 3 | 3 | 0.71 | 0.52 |
| Q15035 | TRAM2 | 2 | 2 | 0.71 | 0.79 |
| P82675 | MRPS5 | 2 | 2 | 0.71 | 0.01 |
| Q6ZNL6 | FGD5 | 2 | 2 | 0.71 | 0.01 |
| Q9NX08 | COMMD8 | 2 | 2 | 0.71 | 0.01 |
| Q9UDW1 | UQCR10 | 2 | 2 | 0.71 | 0.64 |
| Q5JTH9 | RRP12 | 5 | 5 | 0.71 | 0.65 |
| P09110 | ACAA1 | 7 | 7 | 0.71 | 0.54 |
| Q9NZM1 | MYOF | 85 | 84 | 0.71 | 0.48 |
| O60565 | GREM1 | 3 | 3 | 0.71 | 2.25 |
| O15533 | TAPBP | 4 | 4 | 0.71 | 0.78 |
| O14841 | OPLAH | 4 | 4 | 0.71 | 0.01 |
| P35232 | PHB | 13 | 13 | 0.71 | 0.59 |

| Accession | Name | # Peptides | # Unique Peptides | Abundance Ratio: day 3 / day 0 | Abundance Ratio: day 7/ day 0 |
|---|---|---|---|---|---|
| Q92845 | KIFAP3 | 2 | 1 | 0.01 | 0.97 |
| Q96BM9 | ARL8A | 2 | 1 | 0.01 | 0.76 |
| P45973 | CBX5 | 3 | 3 | 0.71 | 0.56 |
| Q9Y3D6 | FIS1 | 4 | 4 | 0.71 | 0.65 |
| P05121 | SERPINE1 | 7 | 7 | 0.71 | 0.61 |
| Q96K17 | BTF3L4 | 3 | 2 | 0.71 | 0.72 |
| Q96KP1 | EXOC2 | 5 | 5 | 0.71 | 0.70 |
| Q96S52 | PIGS | 5 | 5 | 0.71 | 0.78 |
| Q10472 | GALNT1 | 8 | 8 | 0.71 | 1.25 |
| Q9BUF5 | TUBB6 | 17 | 4 | 0.71 | 0.57 |
| Q13217 | DNAJC3 | 10 | 10 | 0.71 | 0.82 |
| Q92520 | FAM3C | 8 | 8 | 0.71 | 0.80 |
| P06703 | S100A6 | 10 | 10 | 0.71 | 0.57 |
| O43150 | ASAP2 | 2 | 2 | 0.72 | 0.01 |
| P53801 | PTTG1IP | 3 | 3 | 0.72 | 0.41 |
| Q9HB90 | RRAGC | 6 | 6 | 0.72 | 0.01 |
| Q15629 | TRAM1 | 8 | 8 | 0.72 | 0.81 |
| O43795 | MYO1B | 33 | 33 | 0.72 | 0.48 |
| Q01082 | SPTBN1 | 117 | 111 | 0.72 | 0.53 |
| Q16513 | PKN2 | 5 | 5 | 0.72 | 0.01 |
| O14657 | TOR1B | 2 | 2 | 0.72 | 0.82 |
| Q86SX6 | GLRX5 | 3 | 3 | 0.72 | 0.01 |
| Q9BRS8 | LARP6 | 3 | 3 | 0.72 | 0.01 |
| P51688 | SGSH | 6 | 6 | 0.72 | 0.44 |
| Q6NYC8 | PPP1R18 | 17 | 17 | 0.72 | 0.63 |
| Q9BTZ2 | DHRS4 | 3 | 3 | 0.72 | 0.58 |
| P38117 | ETFB | 15 | 15 | 0.72 | 0.55 |
| Q27J81 | INF2 | 25 | 25 | 0.72 | 0.64 |
| O75911 | DHRS3 | 2 | 2 | 0.72 | 0.01 |
| Q9UHB6 | LIMA1 | 24 | 24 | 0.72 | 0.74 |
| Q8N5M4 | TTC9C | 2 | 2 | 0.72 | 0.64 |
| Q9NXC5 | MIOS | 5 | 5 | 0.72 | 0.77 |
| P78362 | SRPK2 | 2 | 1 | 0.72 | 1.00 |
| Q9HAB8 | PPCS | 3 | 3 | 0.72 | 0.87 |
| P13804 | ETFA | 8 | 8 | 0.72 | 0.61 |
| P60981 | DSTN | 14 | 13 | 0.72 | 0.62 |
| Q9H3H3 | C11orf68 | 6 | 6 | 0.72 | 0.56 |
| P50213 | IDH3A | 7 | 7 | 0.72 | 0.63 |
| Q9NSC5 | HOMER3 | 4 | 4 | 0.73 | 0.01 |
| Q8IWB7 | WDFY1 | 6 | 6 | 0.73 | 0.62 |
| Q08945 | SSRP1 | 11 | 11 | 0.73 | 0.65 |
| Q9BPX3 | NCAPG | 2 | 2 | 0.73 | 0.01 |
| O60488 | ACSL4 | 15 | 14 | 0.73 | 0.41 |

| Accession | Name | # Peptides | # Unique Peptides | Abundance Ratio: day 3 / day 0 | Abundance Ratio: day 7/ day 0 |
|---|---|---|---|---|---|
| Q92845 | KIFAP3 | 2 | 1 | 0.01 | 0.97 |
| Q96BM9 | ARL8A | 2 | 1 | 0.01 | 0.76 |
| P13639 | EEF2 | 43 | 42 | 0.73 | 0.91 |
| P36551 | CPOX | 4 | 4 | 0.73 | 0.52 |
| P52630 | STAT2 | 5 | 5 | 0.73 | 0.68 |
| P63172 | DYNLT1 | 2 | 2 | 0.73 | 0.01 |
| Q9NZD2 | GLTP | 3 | 3 | 0.73 | 0.71 |
| Q9Y696 | CLIC4 | 12 | 12 | 0.73 | 0.66 |
| P21980 | TGM2 | 27 | 27 | 0.73 | 0.46 |
| P00403 | MT-CO2 | 3 | 3 | 0.73 | 0.43 |
| P61289 | PSME3 | 5 | 5 | 0.73 | 0.57 |
| Q8TEA8 | DTD1 | 5 | 5 | 0.73 | 0.73 |
| Q9NZU5 | LMCD1 | 9 | 9 | 0.73 | 0.64 |
| P62136 | PPP1CA | 13 | 4 | 0.73 | 0.68 |
| Q9H9C1 | VIPAS39 | 3 | 3 | 0.73 | 0.67 |
| Q6UXV4 | APOOL | 3 | 3 | 0.73 | 0.82 |
| Q13813 | SPTAN1 | 130 | 129 | 0.73 | 0.57 |
| O14646 | CHD1 | 2 | 1 | 0.73 | 0.01 |
| Q6IAA8 | LAMTOR1 | 4 | 4 | 0.73 | 0.52 |
| Q6XE24 | RBMS3 | 5 | 4 | 0.73 | 0.74 |
| Q7Z4W1 | DCXR | 5 | 5 | 0.73 | 0.01 |
| P06132 | UROD | 6 | 6 | 0.73 | 0.64 |
| O75489 | NDUFS3 | 6 | 6 | 0.73 | 0.48 |
| Q96CV9 | OPTN | 17 | 17 | 0.73 | 0.80 |
| Q9UM22 | EPDR1 | 2 | 2 | 0.73 | 0.37 |
| P33316 | DUT | 6 | 6 | 0.73 | 0.80 |
| O75439 | PMPCB | 7 | 7 | 0.73 | 0.57 |
| P36957 | DLST | 9 | 9 | 0.73 | 0.49 |
| Q969V3 | NCLN | 9 | 9 | 0.73 | 0.77 |
| P49756 | RBM25 | 13 | 13 | 0.73 | 0.74 |
| Q8NC51 | SERBP1 | 20 | 20 | 0.73 | 0.87 |
| Q06587 | RING1 | 2 | 2 | 0.74 | 0.01 |
| P30049 | ATP5F1D | 3 | 3 | 0.74 | 0.61 |
| P51148 | RAB5C | 7 | 4 | 0.74 | 0.63 |
| P00387 | CYB5R3 | 14 | 14 | 0.74 | 0.52 |
| Q8NF37 | LPCAT1 | 3 | 3 | 0.74 | 0.92 |
| P47985 | UQCRFS1 | 4 | 4 | 0.74 | 0.60 |
| O43765 | SGTA | 10 | 10 | 0.74 | 0.82 |
| Q13423 | NNT | 27 | 27 | 0.74 | 0.58 |
| P16615 | ATP2A2 | 29 | 29 | 0.74 | 0.64 |
| P53396 | ACLY | 37 | 37 | 0.74 | 0.65 |
| Q9P2X0 | DPM3 | 2 | 2 | 0.74 | 0.64 |
| Q9H0S4 | DDX47 | 2 | 2 | 0.74 | 0.01 |

| Accession | Name | # Peptides | # Unique Peptides | Abundance Ratio: day 3 / day 0 | Abundance Ratio: day 7/ day 0 |
|---|---|---|---|---|---|
| Q92845 | KIFAP3 | 2 | 1 | 0.01 | 0.97 |
| Q96BM9 | ARL8A | 2 | 1 | 0.01 | 0.76 |
| Q96CN9 | GCC1 | 2 | 2 | 0.74 | 0.89 |
| Q13505 | MTX1 | 3 | 3 | 0.74 | 0.65 |
| Q06481 | APLP2 | 4 | 4 | 0.74 | 0.37 |
| Q9HCC0 | MCCC2 | 7 | 7 | 0.74 | 0.01 |
| P53602 | MVD | 2 | 2 | 0.74 | 0.78 |
| Q9Y2Q3 | GSTK1 | 9 | 9 | 0.74 | 0.44 |
| Q9Y6C9 | MTCH2 | 9 | 9 | 0.74 | 0.58 |
| Q15043 | SLC39A14 | 3 | 3 | 0.74 | 0.94 |
| O75251 | NDUFS7 | 5 | 5 | 0.74 | 0.49 |
| O94804 | STK10 | 6 | 6 | 0.74 | 0.01 |
| P11498 | PC | 21 | 21 | 0.74 | 0.46 |
| Q15434 | RBMS2 | 3 | 2 | 0.74 | 0.64 |
| P46734 | MAP2K3 | 7 | 7 | 0.74 | 0.57 |
| Q99447 | PCYT2 | 9 | 9 | 0.74 | 0.75 |
| Q99720 | SIGMAR1 | 2 | 2 | 0.74 | 0.66 |
| Q7L523 | RRAGA | 5 | 5 | 0.74 | 0.50 |
| P24844 | MYL9 | 6 | 3 | 0.74 | 0.77 |
| O75844 | ZMPSTE24 | 7 | 7 | 0.74 | 0.83 |
| Q9H270 | VPS11 | 2 | 2 | 0.74 | 0.01 |
| Q9Y2C4 | EXOG | 5 | 5 | 0.74 | 0.77 |
| Q9NQC3 | RTN4 | 7 | 7 | 0.74 | 0.57 |
| O00629 | KPNA4 | 8 | 4 | 0.74 | 0.91 |
| Q7Z4H8 | POGLUT3 | 9 | 9 | 0.74 | 0.83 |
| P08865 | RPSA | 12 | 12 | 0.74 | 0.82 |
| P04062 | GBA | 13 | 13 | 0.74 | 0.54 |
| P15311 | EZR | 41 | 28 | 0.74 | 0.48 |
| P48506 | GCLC | 9 | 9 | 0.74 | 0.54 |
| P49915 | GMPS | 15 | 15 | 0.74 | 0.72 |
| Q9NSE4 | IARS2 | 15 | 15 | 0.74 | 0.70 |
| Q8WX93 | PALLD | 17 | 17 | 0.74 | 1.10 |
| P35052 | GPC1 | 4 | 4 | 0.75 | 0.85 |
| Q9Y5K8 | ATP6V1D | 8 | 8 | 0.75 | 0.69 |
| Q9H4M9 | EHD1 | 21 | 12 | 0.75 | 0.50 |
| Q9HCN8 | SDF2L1 | 4 | 4 | 0.75 | 0.91 |
| P31930 | UQCRC1 | 12 | 12 | 0.75 | 0.59 |
| P50552 | VASP | 14 | 14 | 0.75 | 0.59 |
| Q15814 | TBCC | 6 | 6 | 0.75 | 0.64 |
| Q86Y82 | STX12 | 7 | 7 | 0.75 | 0.76 |
| Q15907 | RAB11B | 12 | 12 | 0.75 | 0.71 |
| Q9Y520 | PRRC2C | 19 | 17 | 0.75 | 0.79 |
| Q9BQE3 | TUBA1C | 23 | 4 | 0.75 | 0.61 |

| Accession | Name | # Peptides | # Unique Peptides | Abundance Ratio: day 3 / day 0 | Abundance Ratio: day 7/ day 0 |
|---|---|---|---|---|---|
| Q92845 | KIFAP3 | 2 | 1 | 0.01 | 0.97 |
| Q96BM9 | ARL8A | 2 | 1 | 0.01 | 0.76 |
| P17844 | DDX5 | 24 | 17 | 0.75 | 0.99 |
| Q9NUQ6 | SPATS2L | 4 | 4 | 0.75 | 0.93 |
| O14828 | SCAMP3 | 6 | 6 | 0.75 | 1.04 |
| Q9H488 | POFUT1 | 6 | 6 | 0.75 | 0.75 |
| P35914 | HMGCL | 9 | 9 | 0.75 | 0.56 |
| Q15654 | TRIP6 | 12 | 12 | 0.75 | 0.87 |
| Q9NX76 | CMTM6 | 3 | 3 | 0.75 | 0.64 |
| P42765 | ACAA2 | 12 | 12 | 0.75 | 0.54 |
| P04264 | KRT1 | 21 | 19 | 0.75 | 0.71 |
| Q9Y6M9 | NDUFB9 | 3 | 3 | 0.75 | 0.65 |
| P53007 | SLC25A1 | 7 | 7 | 0.75 | 0.62 |
| Q9HDC9 | APMAP | 18 | 18 | 0.75 | 0.68 |
| Q16774 | GUK1 | 7 | 7 | 0.75 | 0.88 |
| O60645 | EXOC3 | 9 | 9 | 0.75 | 0.70 |
| P09497 | CLTB | 9 | 9 | 0.75 | 0.62 |
| P13073 | COX4I1 | 10 | 10 | 0.75 | 0.58 |
| Q02252 | ALDH6A1 | 13 | 13 | 0.75 | 0.63 |
| P49327 | FASN | 53 | 53 | 0.75 | 0.72 |
| Q709F0 | ACAD11 | 7 | 7 | 0.76 | 0.01 |
| P68400 | CSNK2A1 | 7 | 7 | 0.76 | 0.78 |
| Q93050 | ATP6V0A1 | 11 | 11 | 0.76 | 0.01 |
| O75880 | SCO1 | 3 | 3 | 0.76 | 0.62 |
| Q07021 | C1QBP | 3 | 3 | 0.76 | 0.01 |
| P18084 | ITGB5 | 13 | 13 | 0.76 | 0.61 |
| Q9UDY2 | TJP2 | 7 | 7 | 0.76 | 0.01 |
| P10606 | COX5B | 7 | 7 | 0.76 | 0.55 |
| Q9H0U4 | RAB1B | 10 | 5 | 0.76 | 0.74 |
| O75955 | FLOT1 | 10 | 10 | 0.76 | 0.63 |
| Q92879 | CELF1 | 7 | 5 | 0.76 | 0.79 |
| Q6XUX3 | DSTYK | 2 | 2 | 0.76 | 0.63 |
| O43815 | STRN | 3 | 2 | 0.76 | 0.01 |
| Q15061 | WDR43 | 3 | 3 | 0.76 | 0.78 |
| P36776 | LONP1 | 8 | 8 | 0.76 | 0.72 |
| P51149 | RAB7A | 11 | 11 | 0.76 | 0.77 |
| Q96JJ7 | TMX3 | 8 | 8 | 0.76 | 0.68 |
| Q96CT7 | CCDC124 | 10 | 10 | 0.76 | 1.14 |
| O00299 | CLIC1 | 11 | 11 | 0.76 | 0.58 |
| Q5VIR6 | VPS53 | 2 | 2 | 0.76 | 0.78 |
| P62847 | RPS24 | 6 | 6 | 0.76 | 0.91 |
| P55145 | MANF | 10 | 10 | 0.76 | 0.94 |
| P12004 | PCNA | 12 | 12 | 0.76 | 0.66 |

| Accession | Name | # Peptides | # Unique Peptides | Abundance Ratio: day 3 / day 0 | Abundance Ratio: day 7/ day 0 |
|---|---|---|---|---|---|
| Q92845 | KIFAP3 | 2 | 1 | 0.01 | 0.97 |
| Q96BM9 | ARL8A | 2 | 1 | 0.01 | 0.76 |
| Q07866 | KLC1 | 19 | 15 | 0.76 | 0.69 |
| P78344 | EIF4G2 | 25 | 25 | 0.76 | 0.79 |
| O95139 | NDUFB6 | 3 | 3 | 0.76 | 0.71 |
| Q8TCD5 | NT5C | 3 | 3 | 0.76 | 0.01 |
| Q16762 | TST | 5 | 5 | 0.76 | 0.58 |
| P47712 | PLA2G4A | 10 | 10 | 0.76 | 0.71 |
| P35579 | MYH9 | 162 | 133 | 0.76 | 0.67 |
| Q9Y2H6 | FNDC3A | 14 | 14 | 0.76 | 0.65 |
| P50995 | ANXA11 | 21 | 21 | 0.76 | 0.51 |
| Q86W92 | PPFIBP1 | 25 | 24 | 0.76 | 0.69 |
| Q9H3K2 | GHITM | 4 | 4 | 0.76 | 0.84 |
| Q0ZGT2 | NEXN | 7 | 7 | 0.76 | 0.74 |
| Q09666 | AHNAK | 341 | 339 | 0.76 | 0.62 |
| Q9BVG4 | PBDC1 | 3 | 3 | 0.77 | 0.78 |
| Q96RP9 | GFM1 | 3 | 3 | 0.77 | 0.01 |
| Q9UJS0 | SLC25A13 | 6 | 6 | 0.77 | 0.01 |
| P15170 | GSPT1 | 14 | 6 | 0.77 | 0.78 |
| P67809 | YBX1 | 12 | 8 | 0.77 | 0.93 |
| Q9Y266 | NUDC | 12 | 12 | 0.77 | 0.85 |
| P13473 | LAMP2 | 3 | 3 | 0.77 | 0.55 |
| P54920 | NAPA | 11 | 11 | 0.77 | 0.66 |
| Q12931 | TRAP1 | 12 | 12 | 0.77 | 0.81 |
| O14745 | SLC9A3R1 | 19 | 19 | 0.77 | 0.49 |
| P19021 | PAM | 3 | 3 | 0.77 | 0.01 |
| Q5T9L3 | WLS | 4 | 4 | 0.77 | 0.64 |
| Q7LBR1 | CHMP1B | 5 | 5 | 0.77 | 0.41 |
| P18031 | PTPN1 | 10 | 10 | 0.77 | 0.88 |
| Q15046 | KARS1 | 20 | 20 | 0.77 | 0.81 |
| Q4V9L6 | TMEM119 | 3 | 3 | 0.77 | 0.88 |
| P46776 | RPL27A | 5 | 5 | 0.77 | 0.92 |
| P62244 | RPS15A | 8 | 8 | 0.77 | 0.88 |
| Q8TCT9 | HM13 | 8 | 8 | 0.77 | 0.77 |
| P49821 | NDUFV1 | 11 | 11 | 0.77 | 0.50 |
| Q16543 | CDC37 | 12 | 12 | 0.77 | 0.80 |
| Q969T9 | WBP2 | 4 | 4 | 0.77 | 0.46 |
| P21926 | CD9 | 6 | 6 | 0.77 | 0.46 |
| P17812 | CTPS1 | 9 | 8 | 0.77 | 0.81 |
| Q15056 | EIF4H | 10 | 10 | 0.77 | 1.12 |
| P46060 | RANGAP1 | 11 | 11 | 0.77 | 0.82 |
| Q15942 | ZYX | 12 | 12 | 0.77 | 0.68 |
| Q7Z406 | MYH14 | 13 | 1 | 0.77 | 1.06 |

| Accession | Name | # Peptides | # Unique Peptides | Abundance Ratio: day 3 / day 0 | Abundance Ratio: day 7/ day 0 |
|---|---|---|---|---|---|
| Q92845 | KIFAP3 | 2 | 1 | 0.01 | 0.97 |
| Q96BM9 | ARL8A | 2 | 1 | 0.01 | 0.76 |
| Q6DD88 | ATL3 | 14 | 14 | 0.77 | 0.64 |
| Q13509 | TUBB3 | 23 | 5 | 0.77 | 0.70 |
| Q9Y490 | TLN1 | 131 | 111 | 0.77 | 0.59 |
| Q9GZZ9 | UBA5 | 4 | 4 | 0.77 | 0.76 |
| Q9H0A0 | NAT10 | 5 | 5 | 0.77 | 0.01 |
| Q9H7Z7 | PTGES2 | 5 | 5 | 0.77 | 0.75 |
| Q562R1 | ACTBL2 | 9 | 3 | 0.77 | 0.64 |
| Q9HB71 | CACYBP | 13 | 13 | 0.77 | 0.64 |
| Q96S97 | MYADM | 2 | 2 | 0.77 | 0.46 |
| P13987 | CD59 | 4 | 4 | 0.77 | 0.44 |
| P98179 | RBM3 | 6 | 6 | 0.77 | 0.82 |
| Q9HD45 | TM9SF3 | 7 | 7 | 0.77 | 1.05 |
| P62877 | RBX1 | 3 | 3 | 0.77 | 0.71 |
| Q9BYT8 | NLN | 3 | 3 | 0.77 | 0.58 |
| Q99615 | DNAJC7 | 4 | 4 | 0.77 | 1.04 |
| P56537 | EIF6 | 5 | 5 | 0.77 | 0.73 |
| Q9UBV2 | SEL1L | 8 | 8 | 0.77 | 0.82 |
| O95202 | LETM1 | 10 | 10 | 0.77 | 0.65 |
| Q9UL25 | RAB21 | 10 | 10 | 0.77 | 0.76 |
| P36543 | ATP6V1E1 | 10 | 10 | 0.77 | 0.61 |
| O43657 | TSPAN6 | 2 | 2 | 0.77 | 0.01 |
| Q7KZI7 | MARK2 | 3 | 3 | 0.77 | 0.84 |
| Q9Y5P4 | CERT1 | 4 | 4 | 0.77 | 0.01 |
| P14927 | UQCRB | 5 | 5 | 0.77 | 0.63 |
| Q8TBA6 | GOLGA5 | 5 | 5 | 0.77 | 0.01 |
| O60925 | PFDN1 | 7 | 7 | 0.77 | 0.80 |
| P35609 | ACTN2 | 13 | 1 | 0.77 | 0.59 |
| P30520 | ADSS2 | 13 | 12 | 0.77 | 0.62 |
| Q15904 | ATP6AP1 | 3 | 3 | 0.78 | 0.52 |
| O00115 | DNASE2 | 4 | 4 | 0.78 | 0.45 |
| Q9NYL9 | TMOD3 | 10 | 9 | 0.78 | 0.74 |
| P11940 | PABPC1 | 34 | 23 | 0.78 | 0.83 |
| P01033 | TIMP1 | 3 | 3 | 0.78 | 0.74 |
| Q99627 | COPS8 | 3 | 3 | 0.78 | 0.74 |
| P04632 | CAPNS1 | 6 | 6 | 0.78 | 0.58 |
| P48449 | LSS | 8 | 8 | 0.78 | 0.70 |
| Q9Y394 | DHRS7 | 4 | 4 | 0.78 | 0.63 |
| P78357 | CNTNAP1 | 4 | 4 | 0.78 | 0.01 |
| Q8IVM0 | CCDC50 | 5 | 5 | 0.78 | 0.74 |
| P20339 | RAB5A | 7 | 4 | 0.78 | 0.74 |
| Q96EK6 | GNPNAT1 | 8 | 8 | 0.78 | 0.66 |

| Accession | Name | # Peptides | # Unique Peptides | Abundance Ratio: day 3 / day 0 | Abundance Ratio: day 7/ day 0 |
|---|---|---|---|---|---|
| Q92845 | KIFAP3 | 2 | 1 | 0.01 | 0.97 |
| Q96BM9 | ARL8A | 2 | 1 | 0.01 | 0.76 |
| Q9P0K7 | RAI14 | 23 | 23 | 0.78 | 0.61 |
| O75821 | EIF3G | 11 | 11 | 0.78 | 0.80 |
| O60493 | SNX3 | 10 | 8 | 0.78 | 0.77 |
| Q99426 | TBCB | 10 | 10 | 0.78 | 0.79 |
| Q8NBN3 | TMEM87A | 3 | 3 | 0.78 | 1.03 |
| Q15637 | SF1 | 11 | 11 | 0.78 | 0.75 |
| Q9Y4L1 | HYOU1 | 26 | 26 | 0.78 | 1.00 |
| P51659 | HSD17B4 | 35 | 35 | 0.78 | 0.64 |
| Q8TEX9 | IPO4 | 5 | 5 | 0.78 | 0.76 |
| Q92508 | PIEZO1 | 8 | 8 | 0.78 | 0.97 |
| Q8N766 | EMC1 | 10 | 10 | 0.78 | 0.94 |
| Q9Y4F5 | CEP170B | 3 | 3 | 0.78 | 0.01 |
| P22570 | FDXR | 8 | 8 | 0.78 | 0.61 |
| P21283 | ATP6V1C1 | 12 | 12 | 0.78 | 0.56 |
| O15127 | SCAMP2 | 4 | 4 | 0.78 | 0.90 |
| Q9HAV7 | GRPEL1 | 4 | 4 | 0.78 | 0.01 |
| Q9H832 | UBE2Z | 4 | 4 | 0.78 | 0.86 |
| Q9BTU6 | PI4K2A | 4 | 4 | 0.78 | 0.01 |
| O00505 | KPNA3 | 7 | 3 | 0.78 | 0.01 |
| Q9NPQ8 | RIC8A | 8 | 8 | 0.78 | 0.95 |
| Q96A65 | EXOC4 | 11 | 11 | 0.78 | 0.74 |
| Q13409 | DYNC1I2 | 12 | 12 | 0.78 | 0.73 |
| O94905 | ERLIN2 | 13 | 9 | 0.78 | 0.71 |
| O43768 | ENSA | 4 | 3 | 0.78 | 1.08 |
| P43155 | CRAT | 6 | 6 | 0.78 | 0.66 |
| Q7Z7N9 | TMEM179B | 2 | 2 | 0.79 | 0.69 |
| Q96Q11 | TRNT1 | 3 | 3 | 0.79 | 0.79 |
| P36222 | CHI3L1 | 21 | 21 | 0.79 | 0.35 |
| Q86V88 | MDP1 | 2 | 2 | 0.79 | 0.77 |
| P42892 | ECE1 | 2 | 2 | 0.79 | 0.01 |
| O14656 | TOR1A | 2 | 2 | 0.79 | 0.01 |
| Q9BQ67 | GRWD1 | 4 | 4 | 0.79 | 0.01 |
| Q9UP95 | SLC12A4 | 6 | 2 | 0.79 | 0.93 |
| P18077 | RPL35A | 10 | 10 | 0.79 | 0.93 |
| P62070 | RRAS2 | 11 | 9 | 0.79 | 0.43 |
| Q9UI12 | ATP6V1H | 11 | 11 | 0.79 | 0.64 |
| P33176 | KIF5B | 40 | 40 | 0.79 | 0.69 |
| Q9BT40 | INPP5K | 2 | 2 | 0.79 | 0.61 |
| P30536 | TSPO | 2 | 2 | 0.79 | 0.67 |
| P61966 | AP1S1 | 3 | 3 | 0.79 | 0.99 |
| Q9UBI6 | GNG12 | 4 | 4 | 0.79 | 0.51 |

| Accession | Name | # Peptides | # Unique Peptides | Abundance Ratio: day 3 / day 0 | Abundance Ratio: day 7/ day 0 |
|---|---|---|---|---|---|
| Q92845 | KIFAP3 | 2 | 1 | 0.01 | 0.97 |
| Q96BM9 | ARL8A | 2 | 1 | 0.01 | 0.76 |
| Q9BUN8 | DERL1 | 4 | 4 | 0.79 | 1.10 |
| P10515 | DLAT | 4 | 4 | 0.79 | 0.01 |
| Q9H2U2 | PPA2 | 9 | 9 | 0.79 | 0.77 |
| P07437 | TUBB | 24 | 2 | 0.79 | 0.69 |
| P13984 | GTF2F2 | 4 | 4 | 0.79 | 0.60 |
| P50583 | NUDT2 | 4 | 4 | 0.79 | 0.82 |
| O75157 | TSC22D2 | 6 | 4 | 0.79 | 0.90 |
| O75937 | DNAJC8 | 6 | 6 | 0.79 | 0.72 |
| Q00653 | NFKB2 | 10 | 9 | 0.79 | 0.69 |
| P19838 | NFKB1 | 10 | 10 | 0.79 | 1.21 |
| Q8TD55 | PLEKHO2 | 10 | 10 | 0.79 | 0.70 |
| P49748 | ACADVL | 27 | 27 | 0.79 | 0.72 |
| Q9H089 | LSG1 | 2 | 2 | 0.79 | 0.01 |
| Q969E2 | SCAMP4 | 3 | 3 | 0.79 | 0.64 |
| P84085 | ARF5 | 7 | 3 | 0.79 | 0.93 |
| Q9H3U1 | UNC45A | 14 | 14 | 0.79 | 0.62 |
| Q9H3P7 | ACBD3 | 15 | 14 | 0.79 | 0.98 |
| O43852 | CALU | 18 | 18 | 0.79 | 0.81 |
| P99999 | CYCS | 5 | 5 | 0.79 | 0.69 |
| O15400 | STX7 | 8 | 7 | 0.79 | 0.70 |
| Q96CS3 | FAF2 | 8 | 8 | 0.79 | 0.66 |
| Q13011 | ECH1 | 9 | 9 | 0.79 | 0.78 |
| O95881 | TXNDC12 | 2 | 2 | 0.79 | 1.03 |
| P20674 | COX5A | 5 | 5 | 0.79 | 0.58 |
| Q86TX2 | ACOT1 | 9 | 9 | 0.79 | 0.76 |
| Q14444 | CAPRIN1 | 10 | 10 | 0.79 | 0.93 |
| Q13740 | ALCAM | 11 | 11 | 0.79 | 0.62 |
| Q9BZF9 | UACA | 17 | 17 | 0.79 | 1.04 |
| Q9UHD8 | SEPTIN9 | 31 | 31 | 0.79 | 0.62 |
| P11142 | HSPA8 | 41 | 26 | 0.79 | 0.70 |
| P13747 | HLA-E | 3 | 1 | 0.79 | 0.01 |
| Q16890 | TPD52L1 | 5 | 5 | 0.79 | 0.50 |
| Q53GQ0 | HSD17B12 | 5 | 5 | 0.79 | 0.70 |
| P15289 | ARSA | 9 | 8 | 0.79 | 0.67 |
| P24539 | ATP5PB | 10 | 10 | 0.79 | 0.70 |
| Q9UBX3 | SLC25A10 | 2 | 2 | 0.79 | 0.84 |
| Q8IWT0 | ZBTB8OS | 2 | 2 | 0.79 | 0.69 |
| O60831 | PRAF2 | 3 | 3 | 0.79 | 1.02 |
| Q9NXH9 | TRMT1 | 3 | 3 | 0.79 | 0.01 |
| Q9UK41 | VPS28 | 3 | 3 | 0.79 | 0.65 |
| Q9NVS9 | PNPO | 6 | 6 | 0.79 | 0.73 |

| Accession | Name | # Peptides | # Unique Peptides | Abundance Ratio: day 3 / day 0 | Abundance Ratio: day 7/ day 0 |
|---|---|---|---|---|---|
| Q92845 | KIFAP3 | 2 | 1 | 0.01 | 0.97 |
| Q96BM9 | ARL8A | 2 | 1 | 0.01 | 0.76 |
| Q9BWD1 | ACAT2 | 7 | 7 | 0.79 | 0.71 |
| P07737 | PFN1 | 11 | 11 | 0.79 | 0.60 |
| Q9NTK5 | OLA1 | 15 | 15 | 0.79 | 0.67 |
| P11413 | G6PD | 26 | 26 | 0.79 | 0.57 |
| P20933 | AGA | 4 | 4 | 0.79 | 0.01 |
| Q9GZT9 | EGLN1 | 5 | 5 | 0.79 | 0.77 |
| O94903 | PLPBP | 5 | 5 | 0.79 | 0.84 |
| P62942 | FKBP1A | 5 | 5 | 0.79 | 0.78 |
| O00487 | PSMD14 | 9 | 9 | 0.79 | 0.83 |
| Q9Y2Z0 | SUGT1 | 10 | 10 | 0.79 | 0.97 |
| Q8N697 | SLC15A4 | 2 | 2 | 0.80 | 0.91 |
| O96000 | NDUFB10 | 4 | 4 | 0.80 | 0.72 |
| P30837 | ALDH1B1 | 5 | 3 | 0.80 | 0.01 |
| P37235 | HPCAL1 | 8 | 8 | 0.80 | 0.76 |
| Q9UNM6 | PSMD13 | 15 | 15 | 0.80 | 0.77 |
| P51648 | ALDH3A2 | 16 | 15 | 0.80 | 0.56 |
| P60842 | EIF4A1 | 24 | 11 | 0.80 | 0.86 |
| Q9NYZ1 | TVP23B | 3 | 3 | 0.80 | 0.71 |
| Q9NPA0 | EMC7 | 4 | 4 | 0.80 | 0.72 |
| P14324 | FDPS | 5 | 5 | 0.80 | 0.65 |
| P09669 | COX6C | 6 | 6 | 0.80 | 0.56 |
| Q07157 | TJP1 | 13 | 13 | 0.80 | 0.83 |
| O43707 | ACTN4 | 62 | 46 | 0.80 | 0.61 |
| Q9GZM8 | NDEL1 | 4 | 3 | 0.80 | 0.84 |
| P62888 | RPL30 | 7 | 7 | 0.80 | 0.88 |
| O75935 | DCTN3 | 8 | 8 | 0.80 | 0.79 |
| Q9H5N1 | RABEP2 | 9 | 9 | 0.80 | 0.75 |
| Q9NR12 | PDLIM7 | 15 | 15 | 0.80 | 0.94 |
| P55268 | LAMB2 | 28 | 28 | 0.80 | 0.74 |
| Q9Y3E0 | GOLT1B | 2 | 2 | 0.80 | 0.87 |
| Q5VZE5 | NAA35 | 3 | 3 | 0.80 | 0.01 |
| Q6DKJ4 | NXN | 6 | 6 | 0.80 | 0.45 |
| Q99816 | TSG101 | 6 | 6 | 0.80 | 0.68 |
| Q08379 | GOLGA2 | 10 | 10 | 0.80 | 1.10 |
| Q01518 | CAP1 | 30 | 28 | 0.80 | 0.67 |
| Q9Y6I9 | TEX264 | 2 | 2 | 0.80 | 0.01 |
| P35240 | NF2 | 5 | 5 | 0.80 | 0.67 |
| Q9GZZ1 | NAA50 | 5 | 5 | 0.80 | 0.80 |
| Q92974 | ARHGEF2 | 5 | 5 | 0.80 | 0.91 |
| Q969Q0 | RPL36AL | 6 | 2 | 0.80 | 0.73 |
| O00178 | GTPBP1 | 6 | 6 | 0.80 | 0.78 |

| Accession | Name | # Peptides | # Unique Peptides | Abundance Ratio: day 3 / day 0 | Abundance Ratio: day 7/ day 0 |
|---|---|---|---|---|---|
| Q92845 | KIFAP3 | 2 | 1 | 0.01 | 0.97 |
| Q96BM9 | ARL8A | 2 | 1 | 0.01 | 0.76 |
| Q9NYI0 | PSD3 | 9 | 9 | 0.80 | 0.50 |
| Q9UGI8 | TES | 13 | 13 | 0.80 | 0.77 |
| O00429 | DNM1L | 17 | 17 | 0.80 | 0.79 |
| P06576 | ATP5F1B | 18 | 18 | 0.80 | 0.60 |
| Q6NVY1 | HIBCH | 6 | 6 | 0.80 | 0.78 |
| Q9UHD1 | CHORDC1 | 7 | 7 | 0.80 | 0.90 |
| P49023 | PXN | 7 | 7 | 0.80 | 0.67 |
| P36542 | ATP5F1C | 12 | 12 | 0.80 | 0.67 |
| P31040 | SDHA | 14 | 14 | 0.80 | 0.59 |
| P55084 | HADHB | 16 | 16 | 0.80 | 0.66 |
| P06753 | TPM3 | 28 | 10 | 0.80 | 0.69 |
| Q8N5K1 | CISD2 | 3 | 3 | 0.80 | 0.88 |
| P63173 | RPL38 | 4 | 4 | 0.80 | 0.95 |
| P62241 | RPS8 | 10 | 10 | 0.80 | 0.85 |
| P61026 | RAB10 | 11 | 10 | 0.80 | 0.72 |
| P49419 | ALDH7A1 | 19 | 19 | 0.80 | 0.63 |
| O95182 | NDUFA7 | 4 | 4 | 0.80 | 0.01 |
| P51452 | DUSP3 | 4 | 4 | 0.80 | 0.61 |
| Q9BRA2 | TXNDC17 | 5 | 5 | 0.80 | 0.63 |
| P12931 | SRC | 6 | 3 | 0.80 | 0.01 |
| Q9Y4W6 | AFG3L2 | 9 | 9 | 0.80 | 0.71 |
| P40123 | CAP2 | 11 | 9 | 0.80 | 0.84 |
| Q9BXP2 | SLC12A9 | 2 | 2 | 0.80 | 0.62 |
| Q86VR2 | RETREG3 | 3 | 3 | 0.80 | 0.83 |
| Q6L8Q7 | PDE12 | 3 | 3 | 0.80 | 0.74 |
| Q9NR30 | DDX21 | 4 | 3 | 0.80 | 1.08 |
| O95816 | BAG2 | 8 | 8 | 0.80 | 0.51 |
| Q04206 | RELA | 9 | 9 | 0.80 | 0.80 |
| P46777 | RPL5 | 15 | 15 | 0.80 | 0.84 |
| O00410 | IPO5 | 21 | 20 | 0.80 | 0.76 |
| P51808 | DYNLT3 | 3 | 3 | 0.81 | 0.01 |
| P00918 | CA2 | 7 | 7 | 0.81 | 0.49 |
| P54819 | AK2 | 11 | 11 | 0.81 | 0.70 |
| P15559 | NQO1 | 12 | 12 | 0.81 | 0.55 |
| P39023 | RPL3 | 26 | 26 | 0.81 | 0.88 |
| P25705 | ATP5F1A | 29 | 29 | 0.81 | 0.67 |
| Q9UK76 | JPT1 | 2 | 2 | 0.81 | 0.01 |
| Q8WUW1 | BRK1 | 3 | 3 | 0.81 | 1.00 |
| Q96G23 | CERS2 | 4 | 4 | 0.81 | 0.81 |
| P83881 | RPL36A | 6 | 2 | 0.81 | 0.89 |
| P62277 | RPS13 | 10 | 10 | 0.81 | 0.91 |

| Accession | Name | # Peptides | # Unique Peptides | Abundance Ratio: day 3 / day 0 | Abundance Ratio: day 7/ day 0 |
|---|---|---|---|---|---|
| Q92845 | KIFAP3 | 2 | 1 | 0.01 | 0.97 |
| Q96BM9 | ARL8A | 2 | 1 | 0.01 | 0.76 |
| P43686 | PSMC4 | 19 | 18 | 0.81 | 0.71 |
| Q14204 | DYNC1H1 | 169 | 169 | 0.81 | 0.76 |
| P56211 | ARPP19 | 4 | 3 | 0.81 | 1.21 |
| Q9H910 | JPT2 | 5 | 5 | 0.81 | 0.80 |
| P32969 | RPL9 | 9 | 9 | 0.81 | 0.87 |
| P08238 | HSP90AB1 | 44 | 22 | 0.81 | 0.90 |
| P61163 | ACTR1A | 14 | 8 | 0.81 | 0.79 |
| P04083 | ANXA1 | 27 | 27 | 0.81 | 0.53 |
| Q15149 | PLEC | 386 | 378 | 0.81 | 0.57 |
| P19474 | TRIM21 | 2 | 2 | 0.81 | 0.53 |
| O95721 | SNAP29 | 3 | 3 | 0.81 | 0.88 |
| Q9HA65 | TBC1D17 | 5 | 5 | 0.81 | 0.84 |
| Q6UN15 | FIP1L1 | 6 | 6 | 0.81 | 0.70 |
| P61020 | RAB5B | 7 | 4 | 0.81 | 0.80 |
| Q14141 | SEPTIN6 | 9 | 2 | 0.81 | 0.69 |
| Q13596 | SNX1 | 13 | 12 | 0.81 | 0.67 |
| Q9BR76 | CORO1B | 15 | 14 | 0.81 | 0.80 |
| Q06210 | GFPT1 | 31 | 24 | 0.81 | 0.95 |
| Q9C0E2 | XPO4 | 5 | 5 | 0.81 | 0.72 |
| Q9H074 | PAIP1 | 5 | 5 | 0.81 | 0.72 |
| P36915 | GNL1 | 7 | 7 | 0.81 | 0.79 |
| P27635 | RPL10 | 9 | 9 | 0.81 | 0.79 |
| O76003 | GLRX3 | 9 | 9 | 0.81 | 0.74 |
| Q12792 | TWF1 | 15 | 12 | 0.81 | 0.86 |
| Q9UNN5 | FAF1 | 6 | 6 | 0.81 | 0.66 |
| P50453 | SERPINB9 | 8 | 8 | 0.81 | 0.81 |
| P12277 | CKB | 14 | 14 | 0.81 | 0.94 |
| Q00610 | CLTC | 67 | 55 | 0.81 | 0.69 |
| P05204 | HMGN2 | 6 | 5 | 0.81 | 0.82 |
| Q5VT25 | CDC42BPA | 7 | 2 | 0.81 | 0.77 |
| Q9UIQ6 | LNPEP | 10 | 10 | 0.81 | 0.66 |
| Q9ULV4 | CORO1C | 24 | 23 | 0.81 | 0.61 |
| P09132 | SRP19 | 2 | 2 | 0.81 | 0.92 |
| O60443 | GSDME | 2 | 2 | 0.81 | 0.01 |
| Q9UHA4 | LAMTOR3 | 2 | 2 | 0.81 | 0.61 |
| Q9UPT5 | EXOC7 | 3 | 3 | 0.81 | 0.63 |
| O43823 | AKAP8 | 4 | 4 | 0.81 | 0.01 |
| Q13155 | AIMP2 | 7 | 7 | 0.81 | 0.78 |
| O15212 | PFDN6 | 8 | 8 | 0.81 | 0.77 |
| Q99733 | NAP1L4 | 10 | 8 | 0.81 | 0.84 |
| O75822 | EIF3J | 10 | 10 | 0.81 | 1.08 |

| Accession | Name | # Peptides | # Unique Peptides | Abundance Ratio: day 3 / day 0 | Abundance Ratio: day 7/ day 0 |
|---|---|---|---|---|---|
| Q92845 | KIFAP3 | 2 | 1 | 0.01 | 0.97 |
| Q96BM9 | ARL8A | 2 | 1 | 0.01 | 0.76 |
| P11387 | TOP1 | 15 | 15 | 0.81 | 0.75 |
| O60701 | UGDH | 21 | 21 | 0.81 | 0.61 |
| Q06330 | RBPJ | 3 | 3 | 0.81 | 0.56 |
| P49773 | HINT1 | 4 | 4 | 0.81 | 0.68 |
| Q7Z2K6 | ERMP1 | 5 | 5 | 0.81 | 0.80 |
| O75962 | TRIO | 6 | 6 | 0.81 | 0.01 |
| P40429 | RPL13A | 15 | 15 | 0.81 | 0.85 |
| Q9P0J0 | NDUFA13 | 2 | 2 | 0.82 | 0.45 |
| Q8IWR0 | ZC3H7A | 2 | 2 | 0.82 | 0.74 |
| Q9UJY1 | HSPB8 | 4 | 4 | 0.82 | 0.91 |
| P11172 | UMPS | 4 | 4 | 0.82 | 0.78 |
| Q9H9E3 | COG4 | 5 | 5 | 0.82 | 1.16 |
| Q53GS9 | USP39 | 6 | 6 | 0.82 | 0.71 |
| Q96BJ3 | AIDA | 6 | 6 | 0.82 | 0.73 |
| Q92614 | MYO18A | 10 | 10 | 0.82 | 0.91 |
| P17655 | CAPN2 | 20 | 19 | 0.82 | 0.61 |
| Q9NX14 | NDUFB11 | 2 | 2 | 0.82 | 0.69 |
| P16333 | NCK1 | 5 | 5 | 0.82 | 0.81 |
| O60762 | DPM1 | 9 | 9 | 0.82 | 0.64 |
| P54886 | ALDH18A1 | 23 | 23 | 0.82 | 1.13 |
| Q05682 | CALD1 | 43 | 43 | 0.82 | 0.83 |
| Q9H477 | RBKS | 2 | 2 | 0.82 | 0.75 |
| P11234 | RALB | 4 | 3 | 0.82 | 0.59 |
| Q9UM47 | NOTCH3 | 6 | 5 | 0.82 | 0.01 |
| P05067 | APP | 6 | 6 | 0.82 | 0.01 |
| Q12996 | CSTF3 | 8 | 8 | 0.82 | 0.99 |
| Q96PE2 | ARHGEF17 | 5 | 5 | 0.82 | 0.01 |
| Q9P299 | COPZ2 | 7 | 7 | 0.82 | 0.99 |
| P62081 | RPS7 | 11 | 11 | 0.82 | 0.94 |
| Q12904 | AIMP1 | 14 | 14 | 0.82 | 0.84 |
| P15880 | RPS2 | 16 | 16 | 0.82 | 0.84 |
| P47756 | CAPZB | 17 | 17 | 0.82 | 0.72 |
| P05455 | SSB | 19 | 18 | 0.82 | 0.80 |
| O00571 | DDX3X | 22 | 6 | 0.82 | 0.94 |
| Q96QC0 | PPP1R10 | 2 | 2 | 0.82 | 0.01 |
| Q8NEW0 | SLC30A7 | 2 | 2 | 0.82 | 0.80 |
| Q02880 | TOP2B | 6 | 6 | 0.82 | 0.01 |
| Q9NV70 | EXOC1 | 6 | 6 | 0.82 | 0.01 |
| P63241 | EIF5A | 15 | 7 | 0.82 | 0.84 |
| Q96RQ1 | ERGIC2 | 2 | 2 | 0.82 | 0.01 |
| O14558 | HSPB6 | 5 | 5 | 0.82 | 0.73 |

| Accession | Name | # Peptides | # Unique Peptides | Abundance Ratio: day 3 / day 0 | Abundance Ratio: day 7/ day 0 |
|---|---|---|---|---|---|
| Q92845 | KIFAP3 | 2 | 1 | 0.01 | 0.97 |
| Q96BM9 | ARL8A | 2 | 1 | 0.01 | 0.76 |
| Q9UNE7 | STUB1 | 6 | 6 | 0.82 | 0.69 |
| P39748 | FEN1 | 7 | 7 | 0.82 | 0.59 |
| P30626 | SRI | 7 | 7 | 0.82 | 0.83 |
| Q13283 | G3BP1 | 13 | 12 | 0.82 | 0.85 |
| P26038 | MSN | 62 | 47 | 0.82 | 0.71 |
| Q6UW68 | TMEM205 | 2 | 2 | 0.82 | 0.84 |
| P61421 | ATP6V0D1 | 5 | 5 | 0.82 | 0.62 |
| Q92552 | MRPS27 | 5 | 5 | 0.82 | 0.01 |
| Q9UJW0 | DCTN4 | 7 | 7 | 0.82 | 0.80 |
| Q9BQE5 | APOL2 | 8 | 8 | 0.82 | 0.81 |
| Q15005 | SPCS2 | 10 | 10 | 0.82 | 1.01 |
| P84098 | RPL19 | 10 | 10 | 0.82 | 0.87 |
| O95292 | VAPB | 11 | 10 | 0.82 | 0.83 |
| P17612 | PRKACA | 12 | 7 | 0.82 | 0.89 |
| Q9Y5X1 | SNX9 | 12 | 12 | 0.82 | 0.52 |
| P55060 | CSE1L | 19 | 19 | 0.82 | 0.68 |
| P33897 | ABCD1 | 3 | 3 | 0.82 | 0.01 |
| Q8N4C8 | MINK1 | 6 | 3 | 0.82 | 0.62 |
| Q6NUK1 | SLC25A24 | 10 | 10 | 0.82 | 0.64 |
| P62829 | RPL23 | 10 | 10 | 0.82 | 0.87 |
| Q8NEU8 | APPL2 | 14 | 14 | 0.82 | 0.62 |
| P46821 | MAP1B | 62 | 61 | 0.82 | 1.29 |
| Q92609 | TBC1D5 | 2 | 2 | 0.82 | 0.81 |
| Q96BQ5 | CCDC127 | 3 | 3 | 0.82 | 0.01 |
| P04216 | THY1 | 3 | 3 | 0.82 | 0.59 |
| Q9Y2D4 | EXOC6B | 4 | 4 | 0.82 | 0.01 |
| P62987 | UBA52 | 6 | 1 | 0.82 | 1.06 |
| Q9H1H9 | KIF13A | 7 | 7 | 0.82 | 0.01 |
| Q9Y281 | CFL2 | 9 | 7 | 0.82 | 0.61 |
| O15371 | EIF3D | 13 | 13 | 0.82 | 0.80 |
| P49137 | MAPKAPK2 | 5 | 2 | 0.82 | 0.92 |
| Q08J23 | NSUN2 | 7 | 7 | 0.82 | 0.78 |
| Q02543 | RPL18A | 8 | 8 | 0.82 | 0.89 |
| Q8N6H7 | ARFGAP2 | 10 | 10 | 0.82 | 0.85 |
| O94760 | DDAH1 | 11 | 10 | 0.82 | 0.76 |
| P62280 | RPS11 | 11 | 11 | 0.82 | 0.89 |
| O60610 | DIAPH1 | 12 | 12 | 0.82 | 0.73 |
| P54578 | USP14 | 19 | 19 | 0.82 | 0.72 |
| Q9BVC6 | TMEM109 | 6 | 6 | 0.83 | 0.73 |
| P49207 | RPL34 | 8 | 8 | 0.83 | 0.90 |
| P23528 | CFL1 | 16 | 14 | 0.83 | 0.66 |

| Accession | Name | # Peptides | # Unique Peptides | Abundance Ratio: day 3 / day 0 | Abundance Ratio: day 7/ day 0 |
|---|---|---|---|---|---|
| Q92845 | KIFAP3 | 2 | 1 | 0.01 | 0.97 |
| Q96BM9 | ARL8A | 2 | 1 | 0.01 | 0.76 |
| P62333 | PSMC6 | 17 | 17 | 0.83 | 0.78 |
| P11586 | MTHFD1 | 33 | 32 | 0.83 | 0.65 |
| Q9H7D7 | WDR26 | 3 | 3 | 0.83 | 0.01 |
| Q9Y3P9 | RABGAP1 | 4 | 4 | 0.83 | 0.76 |
| P15927 | RPA2 | 5 | 5 | 0.83 | 0.66 |
| P60866 | RPS20 | 5 | 5 | 0.83 | 0.88 |
| Q9H299 | SH3BGRL3 | 7 | 7 | 0.83 | 0.54 |
| P13693 | TPT1 | 10 | 10 | 0.83 | 0.92 |
| O76094 | SRP72 | 19 | 19 | 0.83 | 0.92 |
| Q9NPH2 | ISYNA1 | 4 | 4 | 0.83 | 0.74 |
| P62273 | RPS29 | 5 | 5 | 0.83 | 0.72 |
| P62851 | RPS25 | 5 | 5 | 0.83 | 0.86 |
| Q15738 | NSDHL | 6 | 6 | 0.83 | 0.73 |
| P63167 | DYNLL1 | 7 | 4 | 0.83 | 0.66 |
| P16989 | YBX3 | 12 | 8 | 0.83 | 1.16 |
| O43432 | EIF4G3 | 13 | 9 | 0.83 | 0.63 |
| P62701 | RPS4X | 15 | 6 | 0.83 | 0.86 |
| P23396 | RPS3 | 19 | 19 | 0.83 | 0.83 |
| Q04637 | EIF4G1 | 37 | 33 | 0.83 | 1.00 |
| P07900 | HSP90AA1 | 48 | 28 | 0.83 | 0.82 |
| Q9BQ49 | SMIM7 | 3 | 3 | 0.83 | 1.24 |
| Q9NZZ3 | CHMP5 | 4 | 4 | 0.83 | 0.78 |
| O95372 | LYPLA2 | 5 | 5 | 0.83 | 0.71 |
| Q86XP3 | DDX42 | 14 | 14 | 0.83 | 1.18 |
| P63244 | RACK1 | 22 | 22 | 0.83 | 0.93 |
| O15258 | RER1 | 4 | 4 | 0.83 | 1.03 |
| Q13404 | UBE2V1 | 8 | 4 | 0.83 | 0.81 |
| O60313 | OPA1 | 9 | 9 | 0.83 | 0.71 |
| P30050 | RPL12 | 9 | 9 | 0.83 | 0.87 |
| P48735 | IDH2 | 14 | 14 | 0.83 | 0.90 |
| O43399 | TPD52L2 | 16 | 16 | 0.83 | 0.66 |
| Q9BTV4 | TMEM43 | 16 | 16 | 0.83 | 0.74 |
| P28482 | MAPK1 | 17 | 12 | 0.83 | 0.61 |
| Q14152 | EIF3A | 48 | 48 | 0.83 | 0.81 |
| O75179 | ANKRD17 | 3 | 3 | 0.83 | 0.84 |
| Q9UJZ1 | STOML2 | 9 | 9 | 0.83 | 0.66 |
| P08134 | RHOC | 11 | 4 | 0.83 | 0.69 |
| Q92888 | ARHGEF1 | 12 | 12 | 0.83 | 0.93 |
| P46781 | RPS9 | 15 | 15 | 0.83 | 0.89 |
| P55884 | EIF3B | 19 | 19 | 0.83 | 0.79 |
| Q9Y314 | NOSIP | 2 | 2 | 0.83 | 0.82 |

| Accession | Name | # Peptides | # Unique Peptides | Abundance Ratio: day 3 / day 0 | Abundance Ratio: day 7/ day 0 |
|---|---|---|---|---|---|
| Q92845 | KIFAP3 | 2 | 1 | 0.01 | 0.97 |
| Q96BM9 | ARL8A | 2 | 1 | 0.01 | 0.76 |
| Q9NRW3 | APOBEC3C | 2 | 2 | 0.83 | 0.01 |
| P62266 | RPS23 | 6 | 6 | 0.83 | 0.87 |
| Q09028 | RBBP4 | 10 | 5 | 0.83 | 0.85 |
| O75131 | CPNE3 | 15 | 14 | 0.83 | 0.61 |
| P20073 | ANXA7 | 18 | 18 | 0.83 | 0.70 |
| Q9Y3U8 | RPL36 | 6 | 6 | 0.83 | 0.81 |
| P16070 | CD44 | 12 | 12 | 0.83 | 0.46 |
| P00491 | PNP | 12 | 12 | 0.83 | 0.72 |
| P26373 | RPL13 | 12 | 12 | 0.83 | 0.98 |
| P42226 | STAT6 | 17 | 17 | 0.83 | 0.57 |
| P63261 | ACTG1 | 27 | 1 | 0.83 | 0.72 |
| P23634 | ATP2B4 | 37 | 23 | 0.83 | 0.54 |
| P11021 | HSPA5 | 52 | 50 | 0.83 | 0.89 |
| P82912 | MRPS11 | 2 | 2 | 0.83 | 0.53 |
| Q9NZT2 | OGFR | 5 | 5 | 0.83 | 0.01 |
| O75947 | ATP5PD | 6 | 6 | 0.83 | 0.66 |
| P05388 | RPLP0 | 10 | 10 | 0.83 | 0.94 |
| Q567U6 | CCDC93 | 11 | 11 | 0.83 | 0.69 |
| Q9Y3E5 | PTRH2 | 2 | 2 | 0.84 | 0.01 |
| Q9BWH2 | FUNDC2 | 2 | 2 | 0.84 | 0.01 |
| O75531 | BANF1 | 2 | 2 | 0.84 | 0.51 |
| P62072 | TIMM10 | 3 | 3 | 0.84 | 0.01 |
| P49770 | EIF2B2 | 4 | 4 | 0.84 | 0.77 |
| Q8WTS6 | SETD7 | 4 | 4 | 0.84 | 0.71 |
| P48556 | PSMD8 | 12 | 12 | 0.84 | 0.87 |
| P09382 | LGALS1 | 12 | 12 | 0.84 | 0.56 |
| P52298 | NCBP2 | 4 | 4 | 0.84 | 0.99 |
| Q96FQ6 | S100A16 | 6 | 6 | 0.84 | 0.87 |
| P08708 | RPS17 | 6 | 6 | 0.84 | 0.85 |
| Q6FI81 | CIAPIN1 | 7 | 7 | 0.84 | 0.94 |
| O00303 | EIF3F | 8 | 8 | 0.84 | 0.86 |
| P46778 | RPL21 | 9 | 9 | 0.84 | 0.91 |
| P37802 | TAGLN2 | 13 | 13 | 0.84 | 0.61 |
| Q9Y4P3 | TBL2 | 14 | 14 | 0.84 | 0.95 |
| P17987 | TCP1 | 23 | 23 | 0.84 | 0.81 |
| P68371 | TUBB4B | 24 | 1 | 0.84 | 0.68 |
| O60502 | OGA | 3 | 3 | 0.84 | 1.04 |
| Q96EP5 | DAZAP1 | 5 | 5 | 0.84 | 0.60 |
| P61011 | SRP54 | 14 | 14 | 0.84 | 0.87 |
| P35998 | PSMC2 | 21 | 21 | 0.84 | 0.76 |
| P27797 | CALR | 26 | 26 | 0.84 | 0.92 |

| Accession | Name | # Peptides | # Unique Peptides | Abundance Ratio: day 3 / day 0 | Abundance Ratio: day 7/ day 0 |
|---|---|---|---|---|---|
| Q92845 | KIFAP3 | 2 | 1 | 0.01 | 0.97 |
| Q96BM9 | ARL8A | 2 | 1 | 0.01 | 0.76 |
| P18206 | VCL | 66 | 66 | 0.84 | 0.73 |
| O95989 | NUDT3 | 4 | 3 | 0.84 | 1.10 |
| Q9NV96 | TMEM30A | 4 | 4 | 0.84 | 0.01 |
| Q9UBV8 | PEF1 | 4 | 4 | 0.84 | 0.72 |
| Q3KQU3 | MAP7D1 | 8 | 8 | 0.84 | 0.95 |
| P62263 | RPS14 | 9 | 9 | 0.84 | 0.92 |
| Q14254 | FLOT2 | 11 | 11 | 0.84 | 0.49 |
| P0DP23 | CALM1 | 11 | 11 | 0.84 | 0.60 |
| A0AVT1 | UBA6 | 14 | 14 | 0.84 | 0.79 |
| Q9UKD2 | MRTO4 | 3 | 3 | 0.84 | 0.83 |
| O43684 | BUB3 | 9 | 9 | 0.84 | 0.80 |
| Q8NHP8 | PLBD2 | 10 | 10 | 0.84 | 0.75 |
| Q9NTJ5 | SACM1L | 17 | 17 | 0.84 | 0.85 |
| Q9Y4D7 | PLXND1 | 2 | 2 | 0.84 | 1.26 |
| Q96B36 | AKT1S1 | 3 | 3 | 0.84 | 0.01 |
| O43617 | TRAPPC3 | 4 | 4 | 0.84 | 1.00 |
| Q9UJ41 | RABGEF1 | 4 | 4 | 0.84 | 0.01 |
| P00492 | HPRT1 | 5 | 5 | 0.84 | 0.64 |
| P34059 | GALNS | 7 | 6 | 0.84 | 0.66 |
| Q9NR28 | DIABLO | 8 | 8 | 0.84 | 0.68 |
| Q13247 | SRSF6 | 11 | 7 | 0.84 | 0.74 |
| P17931 | LGALS3 | 11 | 11 | 0.84 | 0.60 |
| P62269 | RPS18 | 13 | 13 | 0.84 | 0.90 |
| P40222 | TXLNA | 15 | 15 | 0.84 | 1.05 |
| Q14157 | UBAP2L | 17 | 17 | 0.84 | 0.91 |
| P30622 | CLIP1 | 19 | 14 | 0.84 | 0.77 |
| Q9NPJ3 | ACOT13 | 4 | 4 | 0.84 | 0.79 |
| Q9Y3B8 | REXO2 | 6 | 6 | 0.84 | 0.87 |
| P21912 | SDHB | 10 | 10 | 0.84 | 0.68 |
| Q99832 | CCT7 | 26 | 26 | 0.84 | 0.79 |
| P19338 | NCL | 48 | 48 | 0.84 | 0.94 |
| O00566 | MPHOSPH10 | 2 | 2 | 0.84 | 1.06 |
| Q96D46 | NMD3 | 3 | 3 | 0.84 | 0.73 |
| P30519 | HMOX2 | 4 | 4 | 0.84 | 1.03 |
| O00743 | PPP6C | 6 | 6 | 0.84 | 0.89 |
| Q9UBF2 | COPG2 | 7 | 6 | 0.84 | 0.80 |
| P51665 | PSMD7 | 7 | 7 | 0.84 | 0.82 |
| O95373 | IPO7 | 13 | 13 | 0.84 | 0.85 |
| Q14019 | COTL1 | 14 | 14 | 0.84 | 0.68 |
| P40227 | CCT6A | 22 | 22 | 0.84 | 0.85 |
| O75083 | WDR1 | 28 | 28 | 0.84 | 0.73 |

| Accession | Name | # Peptides | # Unique Peptides | Abundance Ratio: day 3 / day 0 | Abundance Ratio: day 7/ day 0 |
|---|---|---|---|---|---|
| Q92845 | KIFAP3 | 2 | 1 | 0.01 | 0.97 |
| Q96BM9 | ARL8A | 2 | 1 | 0.01 | 0.76 |
| Q8NBF2 | NHLRC2 | 4 | 4 | 0.84 | 0.92 |
| Q9H078 | CLPB | 4 | 4 | 0.84 | 0.93 |
| P62841 | RPS15 | 5 | 5 | 0.84 | 0.94 |
| Q9HB07 | MYG1 | 6 | 6 | 0.84 | 0.77 |
| Q6YP21 | KYAT3 | 6 | 6 | 0.84 | 0.71 |
| Q03405 | PLAUR | 7 | 7 | 0.84 | 0.42 |
| P12235 | SLC25A4 | 11 | 1 | 0.84 | 0.60 |
| Q9BXJ9 | NAA15 | 14 | 14 | 0.84 | 0.89 |
| Q02878 | RPL6 | 19 | 19 | 0.84 | 0.86 |
| Q2TAA2 | IAH1 | 5 | 5 | 0.84 | 0.83 |
| P52943 | CRIP2 | 7 | 7 | 0.84 | 0.64 |
| P09496 | CLTA | 8 | 8 | 0.84 | 0.80 |
| O00264 | PGRMC1 | 8 | 8 | 0.84 | 0.77 |
| P39019 | RPS19 | 14 | 14 | 0.84 | 0.87 |
| Q9Y3A5 | SBDS | 16 | 16 | 0.84 | 0.74 |
| Q04917 | YWHAH | 18 | 14 | 0.84 | 0.69 |
| P61964 | WDR5 | 2 | 2 | 0.85 | 0.01 |
| Q9UN86 | G3BP2 | 7 | 6 | 0.85 | 1.19 |
| P62906 | RPL10A | 11 | 11 | 0.85 | 0.86 |
| O60522 | TDRD6 | 2 | 2 | 0.85 | 0.85 |
| Q3MHD2 | LSM12 | 2 | 2 | 0.85 | 0.95 |
| P56385 | ATP5ME | 3 | 3 | 0.85 | 0.73 |
| Q9Y3A3 | MOB4 | 3 | 3 | 0.85 | 0.01 |
| Q4V328 | GRIPAP1 | 5 | 5 | 0.85 | 0.83 |
| O15145 | ARPC3 | 6 | 6 | 0.85 | 0.76 |
| P35268 | RPL22 | 6 | 6 | 0.85 | 0.76 |
| Q9NVZ3 | NECAP2 | 7 | 7 | 0.85 | 0.95 |
| O95487 | SEC24B | 8 | 6 | 0.85 | 0.84 |
| P61586 | RHOA | 9 | 3 | 0.85 | 0.72 |
| Q9NR56 | MBNL1 | 10 | 10 | 0.85 | 0.68 |
| P21589 | NT5E | 23 | 23 | 0.85 | 0.43 |
| P20020 | ATP2B1 | 26 | 12 | 0.85 | 0.57 |
| Q96DI7 | SNRNP40 | 5 | 5 | 0.85 | 0.01 |
| P08842 | STS | 6 | 6 | 0.85 | 0.44 |
| Q9H0E2 | TOLLIP | 7 | 7 | 0.85 | 0.69 |
| O94826 | TOMM70 | 20 | 20 | 0.85 | 0.71 |
| P46459 | NSF | 27 | 26 | 0.85 | 0.73 |
| Q14353 | GAMT | 2 | 2 | 0.85 | 0.76 |
| Q13057 | COASY | 4 | 4 | 0.85 | 0.93 |
| P20290 | BTF3 | 6 | 5 | 0.85 | 1.05 |
| Q9NQG5 | RPRD1B | 6 | 6 | 0.85 | 0.61 |

| Accession | Name | # Peptides | # Unique Peptides | Abundance Ratio: day 3 / day 0 | Abundance Ratio: day 7/ day 0 |
|---|---|---|---|---|---|
| Q92845 | KIFAP3 | 2 | 1 | 0.01 | 0.97 |
| Q96BM9 | ARL8A | 2 | 1 | 0.01 | 0.76 |
| Q13439 | GOLGA4 | 9 | 8 | 0.85 | 1.06 |
| P05198 | EIF2S1 | 15 | 15 | 0.85 | 0.83 |
| Q02790 | FKBP4 | 20 | 19 | 0.85 | 0.90 |
| P54136 | RARS1 | 25 | 25 | 0.85 | 0.84 |
| Q9BRT3 | MIEN1 | 3 | 3 | 0.85 | 0.78 |
| P28065 | PSMB9 | 3 | 3 | 0.85 | 0.85 |
| P51397 | DAP | 3 | 3 | 0.85 | 1.02 |
| P25445 | FAS | 4 | 4 | 0.85 | 0.65 |
| Q8NBX0 | SCCPDH | 6 | 6 | 0.85 | 0.62 |
| P55039 | DRG2 | 9 | 8 | 0.85 | 0.85 |
| O15511 | ARPC5 | 12 | 12 | 0.85 | 0.74 |
| Q13177 | PAK2 | 13 | 13 | 0.85 | 0.78 |
| O95861 | BPNT1 | 13 | 13 | 0.85 | 0.75 |
| P62249 | RPS16 | 14 | 14 | 0.85 | 0.84 |
| Q13425 | SNTB2 | 16 | 14 | 0.85 | 0.79 |
| Q9Y262 | EIF3L | 17 | 17 | 0.85 | 0.80 |
| Q9UH65 | SWAP70 | 18 | 18 | 0.85 | 0.56 |
| Q96HC4 | PDLIM5 | 24 | 24 | 0.85 | 0.90 |
| O14880 | MGST3 | 3 | 3 | 0.85 | 0.68 |
| Q9HCD5 | NCOA5 | 4 | 4 | 0.85 | 0.77 |
| P61927 | RPL37 | 5 | 5 | 0.85 | 0.94 |
| P07947 | YES1 | 8 | 4 | 0.85 | 0.67 |
| Q9Y2W2 | WBP11 | 8 | 8 | 0.85 | 1.31 |
| O75165 | DNAJC13 | 11 | 11 | 0.85 | 0.69 |
| P22102 | GART | 18 | 18 | 0.85 | 0.84 |
| Q8NBS9 | TXNDC5 | 19 | 19 | 0.85 | 0.93 |
| P26641 | EEF1G | 23 | 23 | 0.85 | 0.79 |
| P04080 | CSTB | 7 | 7 | 0.85 | 0.78 |
| Q9Y6G9 | DYNC1LI1 | 9 | 9 | 0.85 | 0.70 |
| O14979 | HNRNPDL | 13 | 12 | 0.85 | 0.59 |
| Q7L014 | DDX46 | 15 | 15 | 0.85 | 0.74 |
| Q13561 | DCTN2 | 16 | 16 | 0.85 | 0.79 |
| P53384 | NUBP1 | 3 | 3 | 0.85 | 1.41 |
| P61088 | UBE2N | 9 | 9 | 0.85 | 0.74 |
| O75396 | SEC22B | 9 | 9 | 0.85 | 0.85 |
| P28288 | ABCD3 | 9 | 9 | 0.85 | 0.71 |
| A0FGR8 | ESYT2 | 12 | 11 | 0.85 | 0.94 |
| Q99497 | PARK7 | 15 | 15 | 0.85 | 0.87 |
| Q96AB6 | NTAN1 | 3 | 3 | 0.85 | 0.97 |
| P50897 | PPT1 | 5 | 5 | 0.85 | 0.80 |
| Q9NR31 | SAR1A | 6 | 3 | 0.85 | 1.22 |

| Accession | Name | # Peptides | # Unique Peptides | Abundance Ratio: day 3 / day 0 | Abundance Ratio: day 7/ day 0 |
|---|---|---|---|---|---|
| Q92845 | KIFAP3 | 2 | 1 | 0.01 | 0.97 |
| Q96BM9 | ARL8A | 2 | 1 | 0.01 | 0.76 |
| Q9UHD9 | UBQLN2 | 6 | 3 | 0.85 | 0.78 |
| O43681 | GET3 | 6 | 6 | 0.85 | 0.67 |
| P48739 | PITPNB | 10 | 7 | 0.85 | 0.85 |
| P09651 | HNRNPA1 | 20 | 16 | 0.85 | 0.66 |
| P27824 | CANX | 24 | 24 | 0.85 | 1.00 |
| Q99613 | EIF3C | 30 | 30 | 0.85 | 0.87 |
| O00161 | SNAP23 | 3 | 3 | 0.85 | 0.01 |
| Q01484 | ANK2 | 5 | 5 | 0.85 | 0.65 |
| Q13098 | GPS1 | 9 | 9 | 0.85 | 0.92 |
| Q9UHQ9 | CYB5R1 | 9 | 9 | 0.85 | 0.71 |
| O95793 | STAU1 | 11 | 11 | 0.85 | 0.92 |
| Q01658 | DR1 | 3 | 3 | 0.86 | 0.86 |
| P42677 | RPS27 | 4 | 2 | 0.86 | 1.02 |
| O75964 | ATP5MG | 4 | 4 | 0.86 | 0.66 |
| P61106 | RAB14 | 13 | 13 | 0.86 | 0.87 |
| P98082 | DAB2 | 13 | 13 | 0.86 | 0.85 |
| O95302 | FKBP9 | 15 | 14 | 0.86 | 0.89 |
| Q99536 | VAT1 | 17 | 17 | 0.86 | 0.65 |
| P38606 | ATP6V1A | 19 | 19 | 0.86 | 0.75 |
| Q15155 | NOMO1 | 19 | 19 | 0.86 | 0.98 |
| P23588 | EIF4B | 21 | 21 | 0.86 | 1.07 |
| O14949 | UQCRQ | 2 | 2 | 0.86 | 0.66 |
| Q8NC44 | RETREG2 | 4 | 4 | 0.86 | 0.01 |
| P24593 | IGFBP5 | 7 | 7 | 0.86 | 1.33 |
| Q9UKY7 | CDV3 | 9 | 9 | 0.86 | 0.72 |
| Q9Y639 | NPTN | 11 | 11 | 0.86 | 0.60 |
| P46782 | RPS5 | 11 | 11 | 0.86 | 0.89 |
| Q15691 | MAPRE1 | 12 | 10 | 0.86 | 1.01 |
| Q00688 | FKBP3 | 14 | 14 | 0.86 | 0.84 |
| Q13200 | PSMD2 | 20 | 20 | 0.86 | 0.81 |
| Q9NYU2 | UGGT1 | 33 | 31 | 0.86 | 0.89 |
| Q8NHP6 | MOSPD2 | 2 | 2 | 0.86 | 0.62 |
| Q9BY32 | ITPA | 4 | 4 | 0.86 | 0.76 |
| Q5JTZ9 | AARS2 | 4 | 4 | 0.86 | 0.01 |
| Q9UBM7 | DHCR7 | 5 | 5 | 0.86 | 0.79 |
| Q93034 | CUL5 | 6 | 6 | 0.86 | 0.01 |
| P24534 | EEF1B2 | 8 | 6 | 0.86 | 0.80 |
| Q15008 | PSMD6 | 24 | 24 | 0.86 | 0.84 |
| Q9Y3E7 | CHMP3 | 3 | 3 | 0.86 | 0.99 |
| Q15843 | NEDD8 | 3 | 3 | 0.86 | 0.77 |
| P17900 | GM2A | 4 | 4 | 0.86 | 0.61 |

| Accession | Name | # Peptides | # Unique Peptides | Abundance Ratio: day 3 / day 0 | Abundance Ratio: day 7/ day 0 |
|---|---|---|---|---|---|
| Q92845 | KIFAP3 | 2 | 1 | 0.01 | 0.97 |
| Q96BM9 | ARL8A | 2 | 1 | 0.01 | 0.76 |
| P0DN76 | U2AF1L5 | 6 | 6 | 0.86 | 0.83 |
| P47813 | EIF1AX | 7 | 7 | 0.86 | 0.75 |
| O95486 | SEC24A | 9 | 7 | 0.86 | 1.00 |
| O15372 | EIF3H | 10 | 10 | 0.86 | 0.87 |
| Q06323 | PSME1 | 13 | 13 | 0.86 | 0.64 |
| Q93052 | LPP | 19 | 19 | 0.86 | 0.70 |
| Q96AC1 | FERMT2 | 27 | 27 | 0.86 | 0.72 |
| P49721 | PSMB2 | 5 | 5 | 0.86 | 0.73 |
| O43294 | TGFB1I1 | 5 | 5 | 0.86 | 0.84 |
| Q9NWB6 | ARGLU1 | 7 | 7 | 0.86 | 0.66 |
| Q14315 | FLNC | 47 | 32 | 0.86 | 0.72 |
| Q969X1 | TMBIM1 | 2 | 2 | 0.86 | 0.51 |
| Q92620 | DHX38 | 4 | 4 | 0.86 | 0.74 |
| Q9NW15 | ANO10 | 5 | 5 | 0.86 | 0.90 |
| Q13501 | SQSTM1 | 6 | 6 | 0.86 | 0.61 |
| O94901 | SUN1 | 9 | 9 | 0.86 | 0.66 |
| P62913 | RPL11 | 9 | 9 | 0.86 | 0.92 |
| Q14008 | CKAP5 | 21 | 21 | 0.86 | 0.97 |
| Q15436 | SEC23A | 22 | 20 | 0.86 | 0.90 |
| Q8IZ21 | PHACTR4 | 2 | 2 | 0.86 | 0.01 |
| O14907 | TAX1BP3 | 5 | 5 | 0.86 | 1.01 |
| O60884 | DNAJA2 | 7 | 7 | 0.86 | 0.91 |
| Q9NUQ8 | ABCF3 | 8 | 8 | 0.86 | 1.13 |
| Q9H9B4 | SFXN1 | 9 | 9 | 0.86 | 0.61 |
| Q8TAT6 | NPLOC4 | 11 | 11 | 0.86 | 0.78 |
| Q96TA1 | NIBAN2 | 25 | 25 | 0.86 | 0.57 |
| P78371 | CCT2 | 29 | 29 | 0.86 | 0.89 |
| P40939 | HADHA | 29 | 29 | 0.86 | 0.69 |
| Q9NQT4 | EXOSC5 | 2 | 2 | 0.86 | 0.01 |
| O43310 | CTIF | 2 | 2 | 0.86 | 0.01 |
| Q15006 | EMC2 | 5 | 5 | 0.86 | 0.78 |
| Q6NUM9 | RETSAT | 6 | 6 | 0.86 | 0.61 |
| Q9H269 | VPS16 | 6 | 6 | 0.86 | 0.82 |
| Q6P1J9 | CDC73 | 6 | 6 | 0.86 | 0.71 |
| Q8IY81 | FTSJ3 | 8 | 8 | 0.86 | 0.74 |
| O43290 | SART1 | 10 | 10 | 0.86 | 0.78 |
| P61313 | RPL15 | 13 | 13 | 0.86 | 0.83 |
| O60271 | SPAG9 | 23 | 23 | 0.86 | 0.81 |
| Q13435 | SF3B2 | 23 | 23 | 0.86 | 0.86 |
| Q969P0 | IGSF8 | 2 | 2 | 0.86 | 0.01 |
| Q6DN90 | IQSEC1 | 5 | 5 | 0.86 | 1.17 |

| Accession | Name | # Peptides | # Unique Peptides | Abundance Ratio: day 3 / day 0 | Abundance Ratio: day 7/ day 0 |
|---|---|---|---|---|---|
| Q92845 | KIFAP3 | 2 | 1 | 0.01 | 0.97 |
| Q96BM9 | ARL8A | 2 | 1 | 0.01 | 0.76 |
| Q7Z739 | YTHDF3 | 8 | 5 | 0.86 | 0.82 |
| P22392 | NME2 | 10 | 5 | 0.86 | 0.93 |
| Q9Y5B9 | SUPT16H | 17 | 17 | 0.86 | 0.67 |
| Q9UNF1 | MAGED2 | 21 | 20 | 0.86 | 1.66 |
| Q9BVC4 | MLST8 | 2 | 2 | 0.86 | 0.62 |
| Q8N357 | SLC35F6 | 3 | 3 | 0.86 | 1.00 |
| P00325 | ADH1B | 6 | 6 | 0.86 | 0.53 |
| Q9BT22 | ALG1 | 6 | 6 | 0.86 | 0.84 |
| P84157 | MXRA7 | 6 | 6 | 0.86 | 0.76 |
| P30838 | ALDH3A1 | 9 | 8 | 0.86 | 0.76 |
| Q9H0U3 | MAGT1 | 9 | 9 | 0.86 | 0.84 |
| O43847 | NRDC | 9 | 9 | 0.86 | 1.02 |
| P61254 | RPL26 | 12 | 2 | 0.86 | 0.83 |
| Q9UNF0 | PACSIN2 | 13 | 13 | 0.86 | 0.77 |
| P31943 | HNRNPH1 | 17 | 11 | 0.86 | 0.67 |
| Q9NX63 | CHCHD3 | 17 | 17 | 0.86 | 0.65 |
| O94855 | SEC24D | 21 | 21 | 0.86 | 0.99 |
| Q6SZW1 | SARM1 | 4 | 4 | 0.87 | 0.92 |
| O15067 | PFAS | 6 | 6 | 0.87 | 0.93 |
| Q9UKK9 | NUDT5 | 8 | 8 | 0.87 | 0.97 |
| Q13347 | EIF3I | 10 | 10 | 0.87 | 0.92 |
| P62195 | PSMC5 | 15 | 14 | 0.87 | 0.90 |
| P61247 | RPS3A | 28 | 28 | 0.87 | 0.87 |
| Q8WUM4 | PDCD6IP | 45 | 45 | 0.87 | 0.85 |
| Q96H20 | SNF8 | 3 | 3 | 0.87 | 0.94 |
| Q05048 | CSTF1 | 5 | 5 | 0.87 | 0.72 |
| P63220 | RPS21 | 8 | 8 | 0.87 | 0.84 |
| P52306 | RAP1GDS1 | 9 | 9 | 0.87 | 0.84 |
| Q96IJ6 | GMPPA | 10 | 10 | 0.87 | 0.95 |
| P25786 | PSMA1 | 17 | 17 | 0.87 | 0.80 |
| P63104 | YWHAZ | 23 | 17 | 0.87 | 0.71 |
| P36578 | RPL4 | 28 | 27 | 0.87 | 0.84 |
| P49368 | CCT3 | 30 | 30 | 0.87 | 0.85 |
| Q13641 | TPBG | 2 | 2 | 0.87 | 0.01 |
| P54252 | ATXN3 | 3 | 3 | 0.87 | 0.01 |
| Q14160 | SCRIB | 5 | 5 | 0.87 | 0.01 |
| Q9P035 | HACD3 | 7 | 7 | 0.87 | 0.92 |
| P29218 | IMPA1 | 11 | 11 | 0.87 | 0.82 |
| Q14108 | SCARB2 | 14 | 14 | 0.87 | 0.69 |
| O14974 | PPP1R12A | 18 | 18 | 0.87 | 0.78 |
| Q16891 | IMMT | 36 | 36 | 0.87 | 0.70 |

| Accession | Name | # Peptides | # Unique Peptides | Abundance Ratio: day 3 / day 0 | Abundance Ratio: day 7/ day 0 |
|---|---|---|---|---|---|
| Q92845 | KIFAP3 | 2 | 1 | 0.01 | 0.97 |
| Q96BM9 | ARL8A | 2 | 1 | 0.01 | 0.76 |
| P62310 | LSM3 | 3 | 3 | 0.87 | 0.89 |
| Q16537 | PPP2R5E | 3 | 3 | 0.87 | 0.70 |
| P63165 | SUMO1 | 3 | 3 | 0.87 | 0.01 |
| Q7Z5L9 | IRF2BP2 | 5 | 5 | 0.87 | 1.02 |
| P30566 | ADSL | 7 | 7 | 0.87 | 0.80 |
| Q7Z7H5 | TMED4 | 8 | 7 | 0.87 | 0.97 |
| P04424 | ASL | 8 | 8 | 0.87 | 0.01 |
| P18621 | RPL17 | 11 | 11 | 0.87 | 0.87 |
| P54709 | ATP1B3 | 3 | 3 | 0.87 | 0.59 |
| Q86WR0 | CCDC25 | 3 | 3 | 0.87 | 0.78 |
| Q15286 | RAB35 | 6 | 6 | 0.87 | 0.84 |
| P42766 | RPL35 | 7 | 7 | 0.87 | 0.97 |
| P62899 | RPL31 | 9 | 9 | 0.87 | 0.89 |
| P60660 | MYL6 | 11 | 9 | 0.87 | 0.71 |
| O15143 | ARPC1B | 14 | 14 | 0.87 | 0.75 |
| Q08211 | DHX9 | 38 | 38 | 0.87 | 0.66 |
| O95168 | NDUFB4 | 4 | 4 | 0.87 | 0.57 |
| Q68EM7 | ARHGAP17 | 8 | 8 | 0.87 | 1.05 |
| O14818 | PSMA7 | 15 | 15 | 0.87 | 0.74 |
| P61158 | ACTR3 | 18 | 18 | 0.87 | 0.74 |
| P17275 | JUNB | 5 | 5 | 0.87 | 1.31 |
| O75348 | ATP6V1G1 | 7 | 5 | 0.87 | 1.05 |
| Q96M27 | PRRC1 | 7 | 7 | 0.87 | 1.17 |
| P25398 | RPS12 | 9 | 9 | 0.87 | 0.79 |
| O15144 | ARPC2 | 18 | 18 | 0.87 | 0.81 |
| Q96I24 | FUBP3 | 19 | 16 | 0.87 | 0.83 |
| P56192 | MARS1 | 19 | 19 | 0.87 | 0.86 |
| Q9NQA3 | WASH6P | 4 | 1 | 0.87 | 0.01 |
| Q03519 | TAP2 | 4 | 4 | 0.87 | 0.01 |
| O75477 | ERLIN1 | 12 | 8 | 0.87 | 0.65 |
| P04040 | CAT | 13 | 13 | 0.87 | 0.64 |
| Q16881 | TXNRD1 | 15 | 14 | 0.87 | 0.76 |
| P60228 | EIF3E | 15 | 15 | 0.87 | 0.90 |
| Q9H1E3 | NUCKS1 | 4 | 4 | 0.87 | 1.16 |
| Q6UXH1 | CRELD2 | 4 | 4 | 0.87 | 1.22 |
| P31949 | S100A11 | 5 | 5 | 0.87 | 0.74 |
| P67812 | SEC11A | 5 | 5 | 0.87 | 1.02 |
| P48047 | ATP5PO | 9 | 9 | 0.87 | 0.73 |
| Q96AG4 | LRRC59 | 13 | 13 | 0.87 | 0.89 |
| O00754 | MAN2B1 | 15 | 15 | 0.87 | 0.74 |
| O00232 | PSMD12 | 17 | 17 | 0.87 | 0.87 |

| Accession | Name | # Peptides | # Unique Peptides | Abundance Ratio: day 3 / day 0 | Abundance Ratio: day 7/ day 0 |
|---|---|---|---|---|---|
| Q92845 | KIFAP3 | 2 | 1 | 0.01 | 0.97 |
| Q96BM9 | ARL8A | 2 | 1 | 0.01 | 0.76 |
| P34932 | HSPA4 | 44 | 42 | 0.87 | 0.82 |
| P49902 | NT5C2 | 2 | 2 | 0.87 | 0.73 |
| Q9UL15 | BAG5 | 3 | 3 | 0.87 | 0.81 |
| Q9UBS4 | DNAJB11 | 10 | 10 | 0.87 | 0.99 |
| Q12874 | SF3A3 | 11 | 11 | 0.87 | 0.72 |
| P28074 | PSMB5 | 11 | 11 | 0.87 | 0.86 |
| Q8WWM7 | ATXN2L | 18 | 16 | 0.87 | 0.95 |
| Q9UQ80 | PA2G4 | 18 | 18 | 0.87 | 0.84 |
| O60841 | EIF5B | 18 | 18 | 0.87 | 0.95 |
| Q96AE4 | FUBP1 | 28 | 21 | 0.87 | 0.87 |
| Q9Y3C8 | UFC1 | 3 | 3 | 0.88 | 0.86 |
| Q9Y3D0 | CIAO2B | 3 | 3 | 0.88 | 0.99 |
| P01116 | KRAS | 4 | 1 | 0.88 | 0.75 |
| P23921 | RRM1 | 7 | 7 | 0.88 | 0.84 |
| Q13242 | SRSF9 | 7 | 7 | 0.88 | 0.77 |
| O43854 | EDIL3 | 7 | 7 | 0.88 | 0.71 |
| P23919 | DTYMK | 8 | 8 | 0.88 | 0.69 |
| Q9H8Y8 | GORASP2 | 9 | 9 | 0.88 | 1.05 |
| P43034 | PAFAH1B1 | 16 | 16 | 0.88 | 0.89 |
| Q15019 | SEPTIN2 | 17 | 17 | 0.88 | 0.74 |
| O75534 | CSDE1 | 18 | 18 | 0.88 | 1.12 |
| O43148 | RNMT | 3 | 3 | 0.88 | 0.60 |
| Q13573 | SNW1 | 6 | 6 | 0.88 | 0.85 |
| Q8WVM8 | SCFD1 | 11 | 11 | 0.88 | 0.92 |
| Q9Y285 | FARSA | 14 | 14 | 0.88 | 0.83 |
| P12081 | HARS1 | 17 | 12 | 0.88 | 0.83 |
| P48637 | GSS | 18 | 18 | 0.88 | 0.68 |
| Q16181 | SEPTIN7 | 29 | 29 | 0.88 | 0.72 |
| Q14203 | DCTN1 | 37 | 37 | 0.88 | 0.83 |
| Q9Y2S6 | TMA7 | 3 | 3 | 0.88 | 1.36 |
| Q15165 | PON2 | 3 | 3 | 0.88 | 0.60 |
| P48729 | CSNK1A1 | 5 | 5 | 0.88 | 0.90 |
| Q9NP72 | RAB18 | 7 | 7 | 0.88 | 1.31 |
| P62979 | RPS27A | 10 | 2 | 0.88 | 0.85 |
| Q01085 | TIAL1 | 10 | 7 | 0.88 | 0.81 |
| Q12965 | MYO1E | 17 | 17 | 0.88 | 0.77 |
| P50395 | GDI2 | 27 | 20 | 0.88 | 0.85 |
| Q15233 | NONO | 28 | 24 | 0.88 | 0.78 |
| Q0VDF9 | HSPA14 | 2 | 2 | 0.88 | 0.01 |
| Q96EK5 | KIFBP | 3 | 3 | 0.88 | 1.04 |
| P60983 | GMFB | 5 | 5 | 0.88 | 0.73 |

| Accession | Name | # Peptides | # Unique Peptides | Abundance Ratio: day 3 / day 0 | Abundance Ratio: day 7/ day 0 |
|---|---|---|---|---|---|
| Q92845 | KIFAP3 | 2 | 1 | 0.01 | 0.97 |
| Q96BM9 | ARL8A | 2 | 1 | 0.01 | 0.76 |
| Q7L5N1 | COPS6 | 5 | 5 | 0.88 | 0.89 |
| Q13547 | HDAC1 | 6 | 3 | 0.88 | 0.01 |
| P09661 | SNRPA1 | 7 | 7 | 0.88 | 0.94 |
| Q9Y5M8 | SRPRB | 10 | 10 | 0.88 | 1.29 |
| P62753 | RPS6 | 15 | 15 | 0.88 | 0.89 |
| P31946 | YWHAB | 18 | 9 | 0.88 | 0.87 |
| Q92900 | UPF1 | 27 | 27 | 0.88 | 0.88 |
| O75533 | SF3B1 | 31 | 31 | 0.88 | 0.91 |
| Q16864 | ATP6V1F | 2 | 2 | 0.88 | 0.87 |
| Q15334 | LLGL1 | 5 | 5 | 0.88 | 0.01 |
| O43324 | EEF1E1 | 5 | 5 | 0.88 | 0.85 |
| O95197 | RTN3 | 5 | 5 | 0.88 | 0.65 |
| Q96EM0 | L3HYPDH | 7 | 7 | 0.88 | 1.20 |
| P53999 | SUB1 | 10 | 10 | 0.88 | 0.80 |
| P12955 | PEPD | 11 | 11 | 0.88 | 0.73 |
| Q9NSD9 | FARSB | 19 | 19 | 0.88 | 0.89 |
| P60709 | ACTB | 27 | 1 | 0.88 | 0.73 |
| Q03001 | DST | 34 | 32 | 0.88 | 0.93 |
| Q7L311 | ARMCX2 | 6 | 6 | 0.88 | 0.93 |
| Q8WZA9 | IRGQ | 6 | 6 | 0.88 | 0.01 |
| P41240 | CSK | 8 | 8 | 0.88 | 0.99 |
| Q7Z2W4 | ZC3HAV1 | 10 | 10 | 0.88 | 0.78 |
| Q15435 | PPP1R7 | 16 | 16 | 0.88 | 0.72 |
| Q8N3E9 | PLCD3 | 4 | 4 | 0.88 | 0.01 |
| Q5HYK7 | SH3D19 | 6 | 6 | 0.88 | 0.80 |
| E9PAV3 | NACA | 7 | 7 | 0.88 | 0.95 |
| P62495 | ETF1 | 11 | 11 | 0.88 | 0.85 |
| Q02750 | MAP2K1 | 12 | 8 | 0.88 | 0.80 |
| P26196 | DDX6 | 13 | 13 | 0.88 | 0.79 |
| P61160 | ACTR2 | 17 | 17 | 0.88 | 0.72 |
| P52272 | HNRNPM | 44 | 44 | 0.88 | 0.67 |
| P07814 | EPRS1 | 56 | 56 | 0.88 | 0.94 |
| Q9UK61 | TASOR | 3 | 3 | 0.88 | 0.01 |
| Q92544 | TM9SF4 | 5 | 5 | 0.88 | 1.26 |
| Q9Y3B3 | TMED7 | 8 | 8 | 0.88 | 1.07 |
| P47755 | CAPZA2 | 9 | 7 | 0.88 | 0.75 |
| Q9BRX2 | PELO | 9 | 9 | 0.88 | 0.82 |
| P62910 | RPL32 | 9 | 9 | 0.88 | 0.84 |
| Q9H223 | EHD4 | 10 | 7 | 0.88 | 0.74 |
| P48643 | CCT5 | 26 | 26 | 0.88 | 0.85 |
| Q01813 | PFKP | 31 | 25 | 0.88 | 0.68 |

| Accession | Name | # Peptides | # Unique Peptides | Abundance Ratio: day 3 / day 0 | Abundance Ratio: day 7/ day 0 |
|---|---|---|---|---|---|
| Q92845 | KIFAP3 | 2 | 1 | 0.01 | 0.97 |
| Q96BM9 | ARL8A | 2 | 1 | 0.01 | 0.76 |
| P60510 | PPP4C | 5 | 5 | 0.88 | 0.83 |
| Q99848 | EBNA1BP2 | 7 | 7 | 0.88 | 0.71 |
| Q01105 | SET | 10 | 10 | 0.88 | 0.64 |
| P36507 | MAP2K2 | 12 | 8 | 0.88 | 0.94 |
| Q9BT78 | COPS4 | 15 | 15 | 0.88 | 0.90 |
| Q13642 | FHL1 | 17 | 17 | 0.88 | 0.66 |
| Q14974 | KPNB1 | 20 | 20 | 0.88 | 0.83 |
| P31948 | STIP1 | 41 | 41 | 0.88 | 0.86 |
| Q13232 | NME3 | 3 | 3 | 0.88 | 0.75 |
| P06730 | EIF4E | 4 | 4 | 0.88 | 0.74 |
| Q8TC12 | RDH11 | 7 | 7 | 0.88 | 0.77 |
| Q9BUL8 | PDCD10 | 8 | 8 | 0.88 | 0.75 |
| Q9BTW9 | TBCD | 10 | 10 | 0.88 | 0.74 |
| P62750 | RPL23A | 13 | 13 | 0.88 | 0.99 |
| O76021 | RSL1D1 | 19 | 19 | 0.88 | 0.81 |
| Q16222 | UAP1 | 21 | 19 | 0.88 | 0.94 |
| Q9GZT4 | SRR | 2 | 2 | 0.89 | 0.77 |
| Q10570 | CPSF1 | 3 | 3 | 0.89 | 0.01 |
| Q96JY6 | PDLIM2 | 5 | 5 | 0.89 | 0.71 |
| P28072 | PSMB6 | 5 | 5 | 0.89 | 0.77 |
| Q9Y512 | SAMM50 | 8 | 8 | 0.89 | 0.74 |
| P25787 | PSMA2 | 9 | 9 | 0.89 | 0.82 |
| P50990 | CCT8 | 38 | 38 | 0.89 | 0.82 |
| P01034 | CST3 | 2 | 2 | 0.89 | 0.01 |
| Q8WXA9 | SREK1 | 2 | 2 | 0.89 | 1.71 |
| O95232 | LUC7L3 | 4 | 4 | 0.89 | 0.87 |
| Q15287 | RNPS1 | 4 | 4 | 0.89 | 0.86 |
| Q9NQ55 | PPAN | 4 | 4 | 0.89 | 0.88 |
| O75663 | TIPRL | 5 | 5 | 0.89 | 0.95 |
| P62140 | PPP1CB | 10 | 3 | 0.89 | 0.88 |
| O14964 | HGS | 19 | 19 | 0.89 | 0.77 |
| P28838 | LAP3 | 20 | 20 | 0.89 | 0.80 |
| Q07352 | ZFP36L1 | 4 | 3 | 0.89 | 1.24 |
| O43169 | CYB5B | 5 | 5 | 0.89 | 0.73 |
| Q9HBH5 | RDH14 | 5 | 5 | 0.89 | 0.86 |
| P24666 | ACP1 | 8 | 8 | 0.89 | 0.82 |
| P20618 | PSMB1 | 9 | 9 | 0.89 | 0.85 |
| Q6Y7W6 | GIGYF2 | 14 | 12 | 0.89 | 0.91 |
| P46783 | RPS10 | 14 | 13 | 0.89 | 1.04 |
| P12268 | IMPDH2 | 17 | 14 | 0.89 | 1.19 |
| P18124 | RPL7 | 20 | 20 | 0.89 | 0.95 |

| Accession | Name | # Peptides | # Unique Peptides | Abundance Ratio: day 3 / day 0 | Abundance Ratio: day 7/ day 0 |
|---|---|---|---|---|---|
| Q92845 | KIFAP3 | 2 | 1 | 0.01 | 0.97 |
| Q96BM9 | ARL8A | 2 | 1 | 0.01 | 0.76 |
| P18433 | PTPRA | 2 | 2 | 0.89 | 0.01 |
| Q86TM6 | SYVN1 | 4 | 4 | 0.89 | 0.99 |
| Q9P289 | STK26 | 8 | 3 | 0.89 | 1.25 |
| Q99436 | PSMB7 | 8 | 8 | 0.89 | 0.78 |
| O43633 | CHMP2A | 10 | 10 | 0.89 | 0.91 |
| P28066 | PSMA5 | 10 | 10 | 0.89 | 0.80 |
| P55265 | ADAR | 16 | 16 | 0.89 | 0.83 |
| Q8WYA6 | CTNNBL1 | 3 | 3 | 0.89 | 0.79 |
| Q5VTU8 | ATP5F1EP2 | 4 | 4 | 0.89 | 0.73 |
| P54725 | RAD23A | 5 | 3 | 0.89 | 0.01 |
| P15090 | FABP4 | 5 | 5 | 0.89 | 0.58 |
| Q16134 | ETFDH | 6 | 6 | 0.89 | 0.71 |
| O14744 | PRMT5 | 9 | 9 | 0.89 | 1.07 |
| Q9UMS4 | PRPF19 | 9 | 9 | 0.89 | 0.78 |
| Q15185 | PTGES3 | 6 | 6 | 0.89 | 0.79 |
| P82979 | SARNP | 6 | 6 | 0.89 | 0.81 |
| P14735 | IDE | 7 | 7 | 0.89 | 0.01 |
| P31689 | DNAJA1 | 8 | 8 | 0.89 | 0.99 |
| Q5VYK3 | ECPAS | 15 | 15 | 0.89 | 0.73 |
| P45974 | USP5 | 18 | 18 | 0.89 | 0.74 |
| Q16270 | IGFBP7 | 7 | 7 | 0.89 | 0.79 |
| Q9NP79 | VTA1 | 7 | 7 | 0.89 | 0.83 |
| Q8TC07 | TBC1D15 | 7 | 7 | 0.89 | 0.77 |
| Q9BXS5 | AP1M1 | 11 | 9 | 0.89 | 0.84 |
| P07099 | EPHX1 | 15 | 15 | 0.89 | 0.66 |
| O00203 | AP3B1 | 22 | 22 | 0.89 | 0.83 |
| P35606 | COPB2 | 31 | 31 | 0.89 | 1.02 |
| Q14643 | ITPR1 | 44 | 36 | 0.89 | 0.60 |
| Q14694 | USP10 | 2 | 2 | 0.89 | 1.44 |
| O76024 | WFS1 | 3 | 3 | 0.89 | 0.01 |
| O75915 | ARL6IP5 | 4 | 4 | 0.89 | 0.82 |
| Q92890 | UFD1 | 7 | 7 | 0.89 | 0.94 |
| Q13045 | FLII | 11 | 11 | 0.89 | 0.85 |
| P62937 | PPIA | 16 | 15 | 0.89 | 0.70 |
| Q02818 | NUCB1 | 16 | 16 | 0.89 | 0.96 |
| O00159 | MYO1C | 44 | 43 | 0.89 | 0.65 |
| P55072 | VCP | 46 | 46 | 0.89 | 0.84 |
| Q07065 | CKAP4 | 48 | 47 | 0.89 | 0.90 |
| P29372 | MPG | 3 | 3 | 0.89 | 0.01 |
| Q9UM00 | TMCO1 | 3 | 3 | 0.89 | 1.06 |
| Q9HC35 | EML4 | 4 | 4 | 0.89 | 0.75 |

| Accession | Name | # Peptides | # Unique Peptides | Abundance Ratio: day 3 / day 0 | Abundance Ratio: day 7/ day 0 |
|---|---|---|---|---|---|
| Q92845 | KIFAP3 | 2 | 1 | 0.01 | 0.97 |
| Q96BM9 | ARL8A | 2 | 1 | 0.01 | 0.76 |
| Q16775 | HAGH | 5 | 5 | 0.89 | 0.71 |
| Q9UHV9 | PFDN2 | 7 | 7 | 0.89 | 0.96 |
| P59998 | ARPC4 | 7 | 7 | 0.89 | 0.79 |
| Q14257 | RCN2 | 8 | 8 | 0.89 | 0.99 |
| Q00169 | PITPNA | 9 | 6 | 0.89 | 0.78 |
| P84095 | RHOG | 9 | 8 | 0.89 | 0.73 |
| Q9Y2L1 | DIS3 | 9 | 9 | 0.89 | 0.96 |
| Q9BVK6 | TMED9 | 11 | 10 | 0.89 | 1.10 |
| P52907 | CAPZA1 | 12 | 10 | 0.89 | 0.77 |
| P52597 | HNRNPF | 12 | 11 | 0.89 | 0.71 |
| P60900 | PSMA6 | 15 | 15 | 0.89 | 0.78 |
| P27708 | CAD | 29 | 29 | 0.89 | 1.03 |
| Q9NVV5 | AIG1 | 2 | 2 | 0.89 | 0.94 |
| P51798 | CLCN7 | 3 | 3 | 0.89 | 0.01 |
| Q14498 | RBM39 | 7 | 7 | 0.89 | 0.81 |
| Q13144 | EIF2B5 | 9 | 9 | 0.89 | 0.86 |
| P61619 | SEC61A1 | 10 | 10 | 0.89 | 1.19 |
| P49321 | NASP | 14 | 14 | 0.89 | 0.81 |
| P50991 | CCT4 | 23 | 23 | 0.89 | 0.82 |
| Q0IIM8 | TBC1D8B | 2 | 2 | 0.90 | 0.74 |
| Q9Y6A4 | CFAP20 | 3 | 3 | 0.90 | 0.80 |
| Q9NVH1 | DNAJC11 | 6 | 6 | 0.90 | 0.72 |
| Q13555 | CAMK2G | 7 | 3 | 0.90 | 0.87 |
| P52294 | KPNA1 | 7 | 5 | 0.90 | 1.03 |
| P98175 | RBM10 | 7 | 7 | 0.90 | 0.92 |
| Q96JB5 | CDK5RAP3 | 8 | 8 | 0.90 | 0.93 |
| O00541 | PES1 | 8 | 8 | 0.90 | 0.78 |
| P31942 | HNRNPH3 | 12 | 11 | 0.90 | 0.73 |
| O00584 | RNASET2 | 2 | 2 | 0.90 | 0.01 |
| O96011 | PEX11B | 4 | 4 | 0.90 | 0.01 |
| P61803 | DAD1 | 4 | 4 | 0.90 | 1.13 |
| Q9UBB6 | NCDN | 5 | 5 | 0.90 | 0.76 |
| O76070 | SNCG | 8 | 8 | 0.90 | 1.04 |
| Q12765 | SCRN1 | 9 | 8 | 0.90 | 0.78 |
| Q9UL46 | PSME2 | 9 | 9 | 0.90 | 0.68 |
| P55036 | PSMD4 | 13 | 13 | 0.90 | 0.78 |
| P78417 | GSTO1 | 13 | 13 | 0.90 | 0.67 |
| P83731 | RPL24 | 15 | 15 | 0.90 | 0.96 |
| P25788 | PSMA3 | 15 | 15 | 0.90 | 0.85 |
| O43747 | AP1G1 | 18 | 18 | 0.90 | 0.89 |
| Q8NE71 | ABCF1 | 19 | 18 | 0.90 | 1.14 |

| Accession | Name | # Peptides | # Unique Peptides | Abundance Ratio: day 3 / day 0 | Abundance Ratio: day 7/ day 0 |
|---|---|---|---|---|---|
| Q92845 | KIFAP3 | 2 | 1 | 0.01 | 0.97 |
| Q96BM9 | ARL8A | 2 | 1 | 0.01 | 0.76 |
| P62424 | RPL7A | 20 | 20 | 0.90 | 0.91 |
| O43143 | DHX15 | 23 | 22 | 0.90 | 0.72 |
| Q16555 | DPYSL2 | 25 | 22 | 0.90 | 0.71 |
| O94979 | SEC31A | 30 | 30 | 0.90 | 1.10 |
| P04156 | PRNP | 2 | 2 | 0.90 | 0.01 |
| Q7Z4G1 | COMMD6 | 2 | 2 | 0.90 | 0.01 |
| Q7L2H7 | EIF3M | 7 | 7 | 0.90 | 0.76 |
| Q9H7C9 | AAMDC | 7 | 7 | 0.90 | 0.01 |
| Q9BQG0 | MYBBP1A | 15 | 15 | 0.90 | 0.87 |
| Q8IUD2 | ERC1 | 19 | 19 | 0.90 | 0.93 |
| Q13613 | MTMR1 | 3 | 3 | 0.90 | 0.01 |
| Q07020 | RPL18 | 7 | 7 | 0.90 | 1.00 |
| P61353 | RPL27 | 9 | 9 | 0.90 | 0.90 |
| P62714 | PPP2CB | 12 | 1 | 0.90 | 0.88 |
| Q6PGP7 | TTC37 | 12 | 12 | 0.90 | 0.97 |
| Q9NUU7 | DDX19A | 13 | 13 | 0.90 | 0.87 |
| Q93009 | USP7 | 14 | 14 | 0.90 | 0.76 |
| P51991 | HNRNPA3 | 21 | 18 | 0.90 | 0.76 |
| Q9UHY1 | NRBP1 | 3 | 3 | 0.90 | 1.15 |
| Q96GK7 | FAHD2A | 4 | 4 | 0.90 | 0.92 |
| P61081 | UBE2M | 10 | 10 | 0.90 | 0.74 |
| P29144 | TPP2 | 26 | 26 | 0.90 | 0.80 |
| Q9NQP4 | PFDN4 | 4 | 4 | 0.90 | 0.83 |
| Q9UBB4 | ATXN10 | 4 | 4 | 0.90 | 0.90 |
| Q9H330 | TMEM245 | 4 | 4 | 0.90 | 1.07 |
| O15305 | PMM2 | 6 | 5 | 0.90 | 0.93 |
| Q9NX40 | OCIAD1 | 6 | 6 | 0.90 | 0.81 |
| P53004 | BLVRA | 6 | 6 | 0.90 | 0.90 |
| Q15363 | TMED2 | 6 | 6 | 0.90 | 1.10 |
| O43583 | DENR | 7 | 7 | 0.90 | 1.02 |
| P26368 | U2AF2 | 8 | 8 | 0.90 | 0.74 |
| Q04760 | GLO1 | 10 | 10 | 0.90 | 0.80 |
| A2RRP1 | NBAS | 13 | 13 | 0.90 | 0.81 |
| Q9BS26 | ERP44 | 13 | 13 | 0.90 | 0.92 |
| O00231 | PSMD11 | 18 | 18 | 0.90 | 0.88 |
| Q15459 | SF3A1 | 22 | 22 | 0.90 | 0.89 |
| Q9UM54 | MYO6 | 23 | 23 | 0.90 | 0.75 |
| O95674 | CDS2 | 4 | 4 | 0.90 | 0.79 |
| Q9UH62 | ARMCX3 | 4 | 4 | 0.90 | 0.77 |
| O75886 | STAM2 | 5 | 4 | 0.90 | 0.86 |
| P10599 | TXN | 5 | 5 | 0.90 | 0.66 |

| Accession | Name | # Peptides | # Unique Peptides | Abundance Ratio: day 3 / day 0 | Abundance Ratio: day 7/ day 0 |
|---|---|---|---|---|---|
| Q92845 | KIFAP3 | 2 | 1 | 0.01 | 0.97 |
| Q96BM9 | ARL8A | 2 | 1 | 0.01 | 0.76 |
| Q92820 | GGH | 6 | 6 | 0.90 | 0.56 |
| P25789 | PSMA4 | 9 | 9 | 0.90 | 0.90 |
| Q9BUQ8 | DDX23 | 10 | 10 | 0.90 | 0.65 |
| O75400 | PRPF40A | 10 | 10 | 0.90 | 0.83 |
| O43242 | PSMD3 | 27 | 27 | 0.90 | 0.85 |
| O15347 | HMGB3 | 4 | 2 | 0.90 | 0.01 |
| P78356 | PIP4K2B | 4 | 2 | 0.90 | 1.04 |
| Q9BQ39 | DDX50 | 6 | 5 | 0.90 | 0.01 |
| P28070 | PSMB4 | 6 | 6 | 0.90 | 0.77 |
| Q9UHQ4 | BCAP29 | 6 | 6 | 0.90 | 0.01 |
| Q92973 | TNPO1 | 10 | 7 | 0.90 | 0.92 |
| Q9Y6M1 | IGF2BP2 | 11 | 11 | 0.90 | 0.79 |
| Q15393 | SF3B3 | 32 | 32 | 0.90 | 0.82 |
| O75884 | RBBP9 | 2 | 2 | 0.90 | 0.81 |
| Q9BU89 | DOHH | 3 | 3 | 0.90 | 0.01 |
| O15294 | OGT | 3 | 3 | 0.90 | 0.97 |
| O43670 | ZNF207 | 5 | 5 | 0.90 | 0.81 |
| P54802 | NAGLU | 6 | 6 | 0.90 | 0.92 |
| Q6ZXV5 | TMTC3 | 6 | 6 | 0.90 | 0.78 |
| Q6UWE0 | LRSAM1 | 6 | 6 | 0.90 | 1.12 |
| Q9HA64 | FN3KRP | 7 | 7 | 0.90 | 0.72 |
| Q9Y6B6 | SAR1B | 5 | 2 | 0.90 | 0.93 |
| P42773 | CDKN2C | 6 | 6 | 0.90 | 0.75 |
| Q92930 | RAB8B | 7 | 4 | 0.90 | 0.59 |
| P61764 | STXBP1 | 7 | 7 | 0.90 | 0.62 |
| Q9Y263 | PLAA | 13 | 13 | 0.90 | 0.88 |
| O75347 | TBCA | 13 | 13 | 0.90 | 0.91 |
| P04439 | HLA-A | 15 | 11 | 0.90 | 0.54 |
| P67870 | CSNK2B | 3 | 3 | 0.91 | 0.88 |
| Q96RS6 | NUDCD1 | 7 | 7 | 0.91 | 0.84 |
| Q15366 | PCBP2 | 9 | 3 | 0.91 | 0.87 |
| Q13492 | PICALM | 16 | 16 | 0.91 | 0.82 |
| Q99460 | PSMD1 | 25 | 25 | 0.91 | 0.81 |
| P13667 | PDIA4 | 48 | 48 | 0.91 | 1.14 |
| Q9H6F5 | CCDC86 | 3 | 3 | 0.91 | 0.80 |
| P42566 | EPS15 | 4 | 4 | 0.91 | 0.01 |
| Q9UKS6 | PACSIN3 | 9 | 9 | 0.91 | 0.73 |
| P28062 | PSMB8 | 10 | 10 | 0.91 | 0.85 |
| Q92538 | GBF1 | 14 | 14 | 0.91 | 1.26 |
| Q96PK6 | RBM14 | 18 | 18 | 0.91 | 0.73 |
| P38919 | EIF4A3 | 19 | 15 | 0.91 | 0.76 |

| Accession | Name | # Peptides | # Unique Peptides | Abundance Ratio: day 3 / day 0 | Abundance Ratio: day 7/ day 0 |
|---|---|---|---|---|---|
| Q92845 | KIFAP3 | 2 | 1 | 0.01 | 0.97 |
| Q96BM9 | ARL8A | 2 | 1 | 0.01 | 0.76 |
| P13489 | RNH1 | 19 | 19 | 0.91 | 0.79 |
| Q16706 | MAN2A1 | 3 | 3 | 0.91 | 1.13 |
| P68036 | UBE2L3 | 4 | 4 | 0.91 | 0.69 |
| P42785 | PRCP | 5 | 5 | 0.91 | 0.69 |
| Q9UBR2 | CTSZ | 6 | 6 | 0.91 | 0.77 |
| P62328 | TMSB4X | 7 | 5 | 0.91 | 0.77 |
| Q96QK1 | VPS35 | 19 | 19 | 0.91 | 0.80 |
| P51572 | BCAP31 | 25 | 25 | 0.91 | 0.80 |
| P11717 | IGF2R | 28 | 28 | 0.91 | 1.02 |
| Q9H446 | RWDD1 | 3 | 3 | 0.91 | 0.01 |
| Q9H1E5 | TMX4 | 4 | 4 | 0.91 | 0.61 |
| Q9NSK0 | KLC4 | 5 | 5 | 0.91 | 0.88 |
| Q96N66 | MBOAT7 | 6 | 6 | 0.91 | 0.92 |
| Q05086 | UBE3A | 7 | 7 | 0.91 | 0.98 |
| Q07666 | KHDRBS1 | 8 | 8 | 0.91 | 0.61 |
| P05114 | HMGN1 | 10 | 10 | 0.91 | 1.49 |
| Q9NVA2 | SEPTIN11 | 18 | 11 | 0.91 | 0.79 |
| Q9Y678 | COPG1 | 24 | 23 | 0.91 | 1.14 |
| P23246 | SFPQ | 33 | 31 | 0.91 | 0.75 |
| Q7KZF4 | SND1 | 48 | 48 | 0.91 | 0.99 |
| Q9NQG1 | MANBAL | 2 | 2 | 0.91 | 0.93 |
| Q9NRY4 | ARHGAP35 | 2 | 2 | 0.91 | 1.55 |
| P40818 | USP8 | 3 | 3 | 0.91 | 0.01 |
| Q96IX5 | ATP5MK | 3 | 3 | 0.91 | 0.70 |
| P98170 | XIAP | 4 | 4 | 0.91 | 0.01 |
| Q8WUF5 | PPP1R13L | 5 | 5 | 0.91 | 1.14 |
| Q12791 | KCNMA1 | 8 | 8 | 0.91 | 0.67 |
| Q9P2T1 | GMPR2 | 10 | 10 | 0.91 | 0.83 |
| P80723 | BASP1 | 10 | 10 | 0.91 | 0.90 |
| A5YKK6 | CNOT1 | 12 | 12 | 0.91 | 0.78 |
| Q99873 | PRMT1 | 14 | 14 | 0.91 | 0.87 |
| O15523 | DDX3Y | 18 | 2 | 0.91 | 1.98 |
| Q13310 | PABPC4 | 23 | 12 | 0.91 | 0.98 |
| O75116 | ROCK2 | 24 | 21 | 0.91 | 0.87 |
| P12814 | ACTN1 | 50 | 30 | 0.91 | 0.99 |
| Q9H9A6 | LRRC40 | 3 | 3 | 0.91 | 0.91 |
| Q96F86 | EDC3 | 3 | 3 | 0.91 | 0.01 |
| P00167 | CYB5A | 4 | 4 | 0.91 | 0.97 |
| Q8NCW5 | NAXE | 5 | 5 | 0.91 | 0.81 |
| P30405 | PPIF | 7 | 6 | 0.91 | 0.01 |
| Q15276 | RABEP1 | 7 | 7 | 0.91 | 0.01 |

| Accession | Name | # Peptides | # Unique Peptides | Abundance Ratio: day 3 / day 0 | Abundance Ratio: day 7/ day 0 |
|---|---|---|---|---|---|
| Q92845 | KIFAP3 | 2 | 1 | 0.01 | 0.97 |
| Q96BM9 | ARL8A | 2 | 1 | 0.01 | 0.76 |
| P08195 | SLC3A2 | 9 | 9 | 0.91 | 0.90 |
| P17980 | PSMC3 | 14 | 14 | 0.91 | 0.86 |
| Q9NY33 | DPP3 | 15 | 15 | 0.91 | 0.73 |
| P27348 | YWHAQ | 20 | 14 | 0.91 | 0.79 |
| Q8TDX7 | NEK7 | 6 | 6 | 0.91 | 0.97 |
| Q9UBC2 | EPS15L1 | 6 | 6 | 0.91 | 0.85 |
| P49593 | PPM1F | 7 | 7 | 0.91 | 0.70 |
| Q8IZ83 | ALDH16A1 | 8 | 8 | 0.91 | 0.92 |
| Q13148 | TARDBP | 9 | 9 | 0.91 | 0.81 |
| P26599 | PTBP1 | 11 | 10 | 0.91 | 0.74 |
| P53621 | COPA | 50 | 50 | 0.91 | 1.02 |
| Q5XKP0 | MICOS13 | 2 | 2 | 0.91 | 0.01 |
| Q5TGZ0 | MICOS10 | 2 | 2 | 0.91 | 0.53 |
| Q9Y625 | GPC6 | 7 | 7 | 0.91 | 0.71 |
| P78318 | IGBP1 | 8 | 8 | 0.91 | 0.98 |
| Q9H3S7 | PTPN23 | 13 | 13 | 0.91 | 0.85 |
| P53992 | SEC24C | 14 | 14 | 0.91 | 0.73 |
| Q13557 | CAMK2D | 18 | 14 | 0.91 | 0.72 |
| Q9GZS3 | WDR61 | 5 | 5 | 0.91 | 0.97 |
| Q9Y6I3 | EPN1 | 7 | 7 | 0.91 | 0.73 |
| Q9BRP8 | PYM1 | 7 | 7 | 0.91 | 0.86 |
| O60547 | GMDS | 19 | 19 | 0.91 | 0.74 |
| P22626 | HNRNPA2B1 | 22 | 21 | 0.91 | 0.71 |
| P61769 | B2M | 5 | 5 | 0.91 | 0.58 |
| O00401 | WASL | 6 | 6 | 0.91 | 0.70 |
| Q13363 | CTBP1 | 10 | 5 | 0.91 | 0.83 |
| P18583 | SON | 10 | 10 | 0.91 | 0.01 |
| Q9C035 | TRIM5 | 2 | 2 | 0.92 | 0.01 |
| Q96RK0 | CIC | 2 | 2 | 0.92 | 0.97 |
| Q07960 | ARHGAP1 | 12 | 12 | 0.92 | 0.95 |
| Q9NZN3 | EHD3 | 14 | 4 | 0.92 | 0.90 |
| P23381 | WARS1 | 18 | 18 | 0.92 | 0.82 |
| Q9Y230 | RUVBL2 | 22 | 22 | 0.92 | 0.87 |
| O43491 | EPB41L2 | 39 | 35 | 0.92 | 0.78 |
| Q9UPN3 | MACF1 | 66 | 63 | 0.92 | 0.97 |
| Q9H0L4 | CSTF2T | 5 | 1 | 0.92 | 0.98 |
| Q16186 | ADRM1 | 5 | 5 | 0.92 | 0.85 |
| Q9UBQ5 | EIF3K | 6 | 6 | 0.92 | 0.87 |
| P46779 | RPL28 | 10 | 10 | 0.92 | 0.95 |
| O75475 | PSIP1 | 11 | 10 | 0.92 | 0.66 |
| Q96P70 | IPO9 | 12 | 12 | 0.92 | 0.80 |

| Accession | Name | # Peptides | # Unique Peptides | Abundance Ratio: day 3 / day 0 | Abundance Ratio: day 7/ day 0 |
|---|---|---|---|---|---|
| Q92845 | KIFAP3 | 2 | 1 | 0.01 | 0.97 |
| Q96BM9 | ARL8A | 2 | 1 | 0.01 | 0.76 |
| Q12841 | FSTL1 | 12 | 12 | 0.92 | 1.13 |
| Q16204 | CCDC6 | 14 | 14 | 0.92 | 0.94 |
| P20810 | CAST | 18 | 18 | 0.92 | 0.74 |
| Q07954 | LRP1 | 140 | 140 | 0.92 | 0.87 |
| O15121 | DEGS1 | 2 | 2 | 0.92 | 1.00 |
| P56377 | AP1S2 | 2 | 2 | 0.92 | 0.83 |
| O14531 | DPYSL4 | 2 | 2 | 0.92 | 0.01 |
| P53634 | CTSC | 4 | 4 | 0.92 | 0.01 |
| O00267 | SUPT5H | 7 | 7 | 0.92 | 0.69 |
| O15498 | YKT6 | 8 | 8 | 0.92 | 1.29 |
| P41227 | NAA10 | 9 | 9 | 0.92 | 0.83 |
| Q9UNZ2 | NSFL1C | 16 | 16 | 0.92 | 0.91 |
| P07195 | LDHB | 17 | 14 | 0.92 | 0.87 |
| P67936 | TPM4 | 40 | 20 | 0.92 | 0.88 |
| Q92616 | GCN1 | 63 | 63 | 0.92 | 0.90 |
| Q7KZ85 | SUPT6H | 5 | 5 | 0.92 | 0.73 |
| Q8TD19 | NEK9 | 8 | 8 | 0.92 | 0.93 |
| Q16576 | RBBP7 | 9 | 4 | 0.92 | 0.77 |
| Q13488 | TCIRG1 | 10 | 10 | 0.92 | 0.67 |
| Q8N1G4 | LRRC47 | 13 | 13 | 0.92 | 0.80 |
| P23526 | AHCY | 20 | 20 | 0.92 | 0.96 |
| P62191 | PSMC1 | 22 | 21 | 0.92 | 0.85 |
| A6NDG6 | PGP | 3 | 3 | 0.92 | 1.09 |
| Q9UNW1 | MINPP1 | 4 | 4 | 0.92 | 1.21 |
| P49184 | DNASE1L1 | 4 | 4 | 0.92 | 0.83 |
| Q5T1M5 | FKBP15 | 8 | 8 | 0.92 | 0.90 |
| P18085 | ARF4 | 10 | 6 | 0.92 | 1.16 |
| O14980 | XPO1 | 23 | 23 | 0.92 | 0.81 |
| Q8N2K0 | ABHD12 | 3 | 3 | 0.92 | 0.69 |
| Q969E4 | TCEAL3 | 6 | 6 | 0.92 | 1.84 |
| Q8IYI6 | EXOC8 | 7 | 7 | 0.92 | 0.73 |
| Q9UHB9 | SRP68 | 15 | 15 | 0.92 | 1.05 |
| O00635 | TRIM38 | 3 | 3 | 0.92 | 0.69 |
| P60033 | CD81 | 4 | 4 | 0.92 | 0.78 |
| Q16836 | HADH | 6 | 6 | 0.92 | 0.84 |
| Q9BWF3 | RBM4 | 6 | 6 | 0.92 | 0.97 |
| P15374 | UCHL3 | 7 | 6 | 0.92 | 0.96 |
| O43865 | AHCYL1 | 17 | 7 | 0.92 | 0.78 |
| P61978 | HNRNPK | 22 | 22 | 0.92 | 0.75 |
| P33527 | ABCC1 | 26 | 25 | 0.92 | 0.63 |
| Q9NQ88 | TIGAR | 4 | 4 | 0.92 | 0.67 |

| Accession | Name | # Peptides | # Unique Peptides | Abundance Ratio: day 3 / day 0 | Abundance Ratio: day 7/ day 0 |
|---|---|---|---|---|---|
| Q92845 | KIFAP3 | 2 | 1 | 0.01 | 0.97 |
| Q96BM9 | ARL8A | 2 | 1 | 0.01 | 0.76 |
| P61086 | UBE2K | 7 | 7 | 0.92 | 0.94 |
| Q9BV20 | MRI1 | 8 | 8 | 0.92 | 0.96 |
| P15531 | NME1 | 10 | 5 | 0.92 | 1.03 |
| Q9BY44 | EIF2A | 11 | 11 | 0.92 | 1.12 |
| P09874 | PARP1 | 28 | 28 | 0.92 | 0.74 |
| Q9Y320 | TMX2 | 3 | 3 | 0.92 | 1.10 |
| Q9BVL2 | NUP58 | 4 | 4 | 0.92 | 0.76 |
| Q99805 | TM9SF2 | 7 | 7 | 0.92 | 1.05 |
| O60784 | TOM1 | 7 | 7 | 0.92 | 0.82 |
| Q04323 | UBXN1 | 8 | 8 | 0.92 | 0.89 |
| Q8N3C0 | ASCC3 | 10 | 10 | 0.92 | 0.97 |
| P41252 | IARS1 | 27 | 27 | 0.92 | 1.07 |
| P17050 | NAGA | 2 | 2 | 0.92 | 0.66 |
| Q96A57 | TMEM230 | 3 | 3 | 0.92 | 1.56 |
| O43172 | PRPF4 | 6 | 6 | 0.92 | 0.01 |
| Q969G3 | SMARCE1 | 7 | 6 | 0.92 | 1.00 |
| Q6VY07 | PACS1 | 4 | 4 | 0.93 | 1.14 |
| Q9UI30 | TRMT112 | 4 | 4 | 0.93 | 0.87 |
| P05387 | RPLP2 | 7 | 6 | 0.93 | 0.97 |
| P35270 | SPR | 7 | 7 | 0.93 | 0.89 |
| Q9H444 | CHMP4B | 8 | 8 | 0.93 | 0.81 |
| Q13938 | CAPS | 9 | 9 | 0.93 | 0.70 |
| P54727 | RAD23B | 12 | 10 | 0.93 | 0.88 |
| P41091 | EIF2S3 | 21 | 21 | 0.93 | 0.84 |
| P53582 | METAP1 | 3 | 3 | 0.93 | 0.83 |
| P48507 | GCLM | 5 | 5 | 0.93 | 0.72 |
| Q13951 | CBFB | 5 | 5 | 0.93 | 0.86 |
| P61513 | RPL37A | 6 | 6 | 0.93 | 0.94 |
| Q96A72 | MAGOHB | 7 | 1 | 0.93 | 0.82 |
| Q09161 | NCBP1 | 8 | 8 | 0.93 | 0.73 |
| P23743 | DGKA | 8 | 8 | 0.93 | 0.96 |
| O94776 | MTA2 | 9 | 8 | 0.93 | 1.08 |
| Q9HD20 | ATP13A1 | 11 | 11 | 0.93 | 0.89 |
| P40121 | CAPG | 14 | 14 | 0.93 | 0.63 |
| Q9P0J7 | KCMF1 | 2 | 2 | 0.93 | 0.01 |
| P85037 | FOXK1 | 6 | 6 | 0.93 | 1.14 |
| Q16831 | UPP1 | 6 | 6 | 0.93 | 0.79 |
| Q9Y2T2 | AP3M1 | 9 | 9 | 0.93 | 0.82 |
| Q8NBJ5 | COLGALT1 | 13 | 13 | 0.93 | 1.21 |
| Q9H2G2 | SLK | 21 | 21 | 0.93 | 0.74 |
| O00139 | KIF2A | 3 | 3 | 0.93 | 0.01 |

| Accession | Name | # Peptides | # Unique Peptides | Abundance Ratio: day 3 / day 0 | Abundance Ratio: day 7/ day 0 |
|---|---|---|---|---|---|
| Q92845 | KIFAP3 | 2 | 1 | 0.01 | 0.97 |
| Q96BM9 | ARL8A | 2 | 1 | 0.01 | 0.76 |
| Q7L775 | EPM2AIP1 | 4 | 4 | 0.93 | 0.70 |
| Q5H9R7 | PPP6R3 | 5 | 5 | 0.93 | 0.76 |
| P10301 | RRAS | 10 | 8 | 0.93 | 0.60 |
| O14579 | COPE | 10 | 10 | 0.93 | 1.00 |
| Q07955 | SRSF1 | 13 | 13 | 0.93 | 0.67 |
| P14618 | PKM | 48 | 48 | 0.93 | 0.74 |
| O00748 | CES2 | 2 | 2 | 0.93 | 0.96 |
| Q8TDN6 | BRIX1 | 4 | 4 | 0.93 | 1.11 |
| O60524 | NEMF | 6 | 6 | 0.93 | 0.81 |
| P62316 | SNRPD2 | 7 | 7 | 0.93 | 0.90 |
| O60504 | SORBS3 | 9 | 9 | 0.93 | 0.64 |
| Q13617 | CUL2 | 11 | 11 | 0.93 | 0.65 |
| Q15181 | PPA1 | 14 | 14 | 0.93 | 0.87 |
| P39656 | DDOST | 14 | 14 | 0.93 | 1.03 |
| Q9NTZ6 | RBM12 | 15 | 14 | 0.93 | 0.78 |
| O94906 | PRPF6 | 16 | 16 | 0.93 | 0.74 |
| Q9H2D6 | TRIOBP | 19 | 18 | 0.93 | 0.75 |
| P22314 | UBA1 | 34 | 34 | 0.93 | 0.80 |
| Q9P253 | VPS18 | 4 | 4 | 0.93 | 0.70 |
| Q86V48 | LUZP1 | 7 | 7 | 0.93 | 1.13 |
| Q6ZMZ3 | SYNE3 | 15 | 15 | 0.93 | 0.60 |
| P07686 | HEXB | 15 | 15 | 0.93 | 0.70 |
| Q06830 | PRDX1 | 19 | 15 | 0.93 | 0.84 |
| Q9NZB2 | FAM120A | 19 | 19 | 0.93 | 0.86 |
| P41226 | UBA7 | 5 | 5 | 0.93 | 0.90 |
| O00273 | DFFA | 7 | 7 | 0.93 | 0.73 |
| Q15643 | TRIP11 | 13 | 13 | 0.93 | 1.38 |
| P08133 | ANXA6 | 49 | 49 | 0.93 | 0.83 |
| P43307 | SSR1 | 3 | 3 | 0.93 | 1.20 |
| Q08752 | PPID | 5 | 4 | 0.93 | 1.01 |
| Q13619 | CUL4A | 13 | 5 | 0.93 | 0.83 |
| P84103 | SRSF3 | 13 | 12 | 0.93 | 0.73 |
| P51114 | FXR1 | 14 | 11 | 0.93 | 0.72 |
| P20042 | EIF2S2 | 14 | 14 | 0.93 | 0.93 |
| P10644 | PRKAR1A | 15 | 15 | 0.93 | 0.75 |
| Q9Y2A7 | NCKAP1 | 23 | 23 | 0.93 | 0.83 |
| Q9NVJ2 | ARL8B | 2 | 1 | 0.93 | 0.74 |
| Q15386 | UBE3C | 4 | 4 | 0.93 | 0.01 |
| Q9BQA9 | CYBC1 | 5 | 5 | 0.93 | 0.93 |
| O00764 | PDXK | 10 | 10 | 0.93 | 0.88 |
| O95433 | AHSA1 | 10 | 10 | 0.93 | 0.94 |

| Accession | Name | # Peptides | # Unique Peptides | Abundance Ratio: day 3 / day 0 | Abundance Ratio: day 7/ day 0 |
|---|---|---|---|---|---|
| Q92845 | KIFAP3 | 2 | 1 | 0.01 | 0.97 |
| Q96BM9 | ARL8A | 2 | 1 | 0.01 | 0.76 |
| P31939 | ATIC | 22 | 22 | 0.93 | 0.81 |
| Q9H0R4 | HDHD2 | 2 | 2 | 0.93 | 0.91 |
| Q8N7R7 | CCNYL1 | 3 | 2 | 0.93 | 0.79 |
| Q8N668 | COMMD1 | 3 | 3 | 0.93 | 0.82 |
| Q14155 | ARHGEF7 | 4 | 4 | 0.93 | 0.86 |
| Q53TN4 | CYBRD1 | 4 | 4 | 0.93 | 0.76 |
| P43235 | CTSK | 6 | 5 | 0.93 | 0.81 |
| Q9Y6K9 | IKBKG | 11 | 11 | 0.93 | 0.76 |
| P06748 | NPM1 | 14 | 14 | 0.93 | 0.87 |
| Q14847 | LASP1 | 16 | 16 | 0.93 | 0.92 |
| Q01130 | SRSF2 | 6 | 5 | 0.94 | 0.73 |
| Q969H8 | MYDGF | 6 | 6 | 0.94 | 1.10 |
| Q99459 | CDC5L | 10 | 10 | 0.94 | 0.88 |
| Q9BZE9 | ASPSCR1 | 5 | 5 | 0.94 | 0.01 |
| P36404 | ARL2 | 5 | 5 | 0.94 | 0.82 |
| P08579 | SNRPB2 | 6 | 5 | 0.94 | 0.79 |
| Q9GZT8 | NIF3L1 | 6 | 6 | 0.94 | 0.94 |
| P62995 | TRA2B | 7 | 5 | 0.94 | 0.68 |
| Q9Y376 | CAB39 | 7 | 6 | 0.94 | 0.90 |
| P53618 | COPB1 | 27 | 27 | 0.94 | 1.14 |
| Q9UPT8 | ZC3H4 | 7 | 6 | 0.94 | 1.03 |
| P47914 | RPL29 | 7 | 7 | 0.94 | 0.96 |
| P35637 | FUS | 12 | 10 | 0.94 | 0.80 |
| Q12959 | DLG1 | 13 | 12 | 0.94 | 0.84 |
| Q15084 | PDIA6 | 16 | 16 | 0.94 | 1.02 |
| P62258 | YWHAE | 26 | 23 | 0.94 | 0.82 |
| P57088 | TMEM33 | 5 | 5 | 0.94 | 0.95 |
| Q9NZ08 | ERAP1 | 6 | 6 | 0.94 | 0.80 |
| P61758 | VBP1 | 8 | 8 | 0.94 | 0.83 |
| Q03135 | CAV1 | 9 | 9 | 0.94 | 0.55 |
| Q9H2M9 | RAB3GAP2 | 18 | 18 | 0.94 | 0.97 |
| P26639 | TARS1 | 23 | 22 | 0.94 | 0.89 |
| P46063 | RECQL | 29 | 29 | 0.94 | 0.73 |
| O75643 | SNRNP200 | 33 | 33 | 0.94 | 0.80 |
| P61970 | NUTF2 | 2 | 2 | 0.94 | 0.85 |
| Q9HD42 | CHMP1A | 3 | 3 | 0.94 | 0.97 |
| P33121 | ACSL1 | 3 | 3 | 0.94 | 0.76 |
| Q9H0C8 | ILKAP | 4 | 4 | 0.94 | 0.85 |
| Q86X55 | CARM1 | 6 | 6 | 0.94 | 0.97 |
| P32455 | GBP1 | 13 | 12 | 0.94 | 0.75 |
| P52209 | PGD | 16 | 16 | 0.94 | 0.79 |

| Accession | Name | # Peptides | # Unique Peptides | Abundance Ratio: day 3 / day 0 | Abundance Ratio: day 7/ day 0 |
|---|---|---|---|---|---|
| Q92845 | KIFAP3 | 2 | 1 | 0.01 | 0.97 |
| Q96BM9 | ARL8A | 2 | 1 | 0.01 | 0.76 |
| Q13885 | TUBB2A | 22 | 1 | 0.94 | 1.47 |
| Q4G0F5 | VPS26B | 4 | 4 | 0.94 | 1.21 |
| Q4G0J3 | LARP7 | 7 | 7 | 0.94 | 0.76 |
| P52565 | ARHGDIA | 10 | 10 | 0.94 | 0.80 |
| P06865 | HEXA | 12 | 12 | 0.94 | 0.64 |
| Q9UEY8 | ADD3 | 12 | 12 | 0.94 | 0.90 |
| P16435 | POR | 15 | 15 | 0.94 | 0.74 |
| O15061 | SYNM | 40 | 40 | 0.94 | 0.73 |
| P61956 | SUMO2 | 3 | 2 | 0.94 | 0.78 |
| Q9UEW8 | STK39 | 6 | 4 | 0.94 | 0.88 |
| P35080 | PFN2 | 8 | 8 | 0.94 | 0.79 |
| P11766 | ADH5 | 14 | 14 | 0.94 | 0.76 |
| P49189 | ALDH9A1 | 16 | 16 | 0.94 | 0.86 |
| Q16531 | DDB1 | 33 | 33 | 0.94 | 0.80 |
| Q14697 | GANAB | 40 | 40 | 0.94 | 1.06 |
| Q15428 | SF3A2 | 3 | 3 | 0.94 | 0.98 |
| Q96DZ1 | ERLEC1 | 3 | 3 | 0.94 | 0.94 |
| P61923 | COPZ1 | 3 | 3 | 0.94 | 1.25 |
| Q9Y608 | LRRFIP2 | 7 | 6 | 0.94 | 0.85 |
| Q15369 | ELOC | 7 | 7 | 0.94 | 0.93 |
| Q9UMY4 | SNX12 | 8 | 6 | 0.94 | 0.82 |
| P50479 | PDLIM4 | 13 | 13 | 0.94 | 1.48 |
| Q13418 | ILK | 14 | 14 | 0.94 | 0.85 |
| P07384 | CAPN1 | 21 | 20 | 0.94 | 0.75 |
| P61009 | SPCS3 | 3 | 3 | 0.94 | 0.01 |
| Q99836 | MYD88 | 4 | 4 | 0.94 | 0.93 |
| Q9H173 | SIL1 | 4 | 4 | 0.94 | 1.22 |
| P53680 | AP2S1 | 4 | 4 | 0.94 | 0.79 |
| P61201 | COPS2 | 13 | 13 | 0.94 | 0.88 |
| Q9UBQ0 | VPS29 | 6 | 6 | 0.94 | 0.82 |
| Q15404 | RSU1 | 7 | 7 | 0.94 | 0.74 |
| O43493 | TGOLN2 | 8 | 8 | 0.94 | 0.93 |
| Q9NZW5 | PALS2 | 11 | 11 | 0.94 | 0.86 |
| Q9UKK3 | PARP4 | 16 | 16 | 0.94 | 0.73 |
| P30153 | PPP2R1A | 20 | 15 | 0.94 | 0.93 |
| Q9H2V7 | SPNS1 | 2 | 2 | 0.95 | 0.85 |
| Q14457 | BECN1 | 2 | 2 | 0.95 | 0.01 |
| Q01628 | IFITM3 | 2 | 2 | 0.95 | 0.01 |
| Q9Y3X0 | CCDC9 | 5 | 5 | 0.95 | 0.84 |
| Q9BZF1 | OSBPL8 | 6 | 5 | 0.95 | 0.82 |
| O43598 | DNPH1 | 6 | 6 | 0.95 | 0.94 |

| Accession | Name | # Peptides | # Unique Peptides | Abundance Ratio: day 3 / day 0 | Abundance Ratio: day 7/ day 0 |
|---|---|---|---|---|---|
| Q92845 | KIFAP3 | 2 | 1 | 0.01 | 0.97 |
| Q96BM9 | ARL8A | 2 | 1 | 0.01 | 0.76 |
| P27694 | RPA1 | 12 | 12 | 0.95 | 1.00 |
| Q9UKM9 | RALY | 15 | 15 | 0.95 | 0.81 |
| Q9BRJ6 | C7orf50 | 3 | 3 | 0.95 | 0.96 |
| O95248 | SBF1 | 4 | 4 | 0.95 | 0.01 |
| Q9H2H8 | PPIL3 | 5 | 5 | 0.95 | 0.93 |
| P52788 | SMS | 6 | 6 | 0.95 | 0.74 |
| Q2M389 | WASHC4 | 7 | 7 | 0.95 | 0.85 |
| Q15126 | PMVK | 7 | 7 | 0.95 | 0.83 |
| Q9Y3Z3 | SAMHD1 | 10 | 10 | 0.95 | 0.01 |
| P49755 | TMED10 | 11 | 10 | 0.95 | 1.11 |
| P61221 | ABCE1 | 12 | 12 | 0.95 | 1.02 |
| P62826 | RAN | 13 | 13 | 0.95 | 0.81 |
| P13861 | PRKAR2A | 15 | 13 | 0.95 | 0.83 |
| Q86UU1 | PHLDB1 | 24 | 24 | 0.95 | 1.09 |
| P48444 | ARCN1 | 24 | 24 | 0.95 | 1.15 |
| Q8IVD9 | NUDCD3 | 2 | 2 | 0.95 | 0.01 |
| Q15477 | SKIV2L | 5 | 4 | 0.95 | 0.97 |
| Q7L2J0 | MEPCE | 6 | 6 | 0.95 | 0.81 |
| Q16629 | SRSF7 | 11 | 10 | 0.95 | 0.66 |
| P09543 | CNP | 16 | 15 | 0.95 | 0.77 |
| P35611 | ADD1 | 17 | 17 | 0.95 | 0.89 |
| O60716 | CTNND1 | 18 | 18 | 0.95 | 0.86 |
| O75369 | FLNB | 96 | 81 | 0.95 | 0.80 |
| Q8IZ81 | ELMOD2 | 3 | 3 | 0.95 | 0.81 |
| Q8IXT5 | RBM12B | 4 | 4 | 0.95 | 0.01 |
| O75340 | PDCD6 | 5 | 5 | 0.95 | 0.98 |
| Q9UBQ7 | GRHPR | 9 | 9 | 0.95 | 0.82 |
| P09619 | PDGFRB | 11 | 9 | 0.95 | 0.83 |
| O43396 | TXNL1 | 12 | 12 | 0.95 | 1.00 |
| P63010 | AP2B1 | 28 | 14 | 0.95 | 0.86 |
| P46939 | UTRN | 65 | 65 | 0.95 | 0.68 |
| Q9NYM9 | BET1L | 2 | 2 | 0.95 | 1.56 |
| Q99614 | TTC1 | 2 | 2 | 0.95 | 0.01 |
| P51580 | TPMT | 3 | 3 | 0.95 | 0.64 |
| Q9BZK7 | TBL1XR1 | 4 | 4 | 0.95 | 1.07 |
| O43772 | SLC25A20 | 5 | 5 | 0.95 | 0.01 |
| O15031 | PLXNB2 | 15 | 15 | 0.95 | 0.73 |
| O43776 | NARS1 | 20 | 20 | 0.95 | 0.91 |
| P08758 | ANXA5 | 21 | 21 | 0.95 | 0.88 |
| Q14247 | CTTN | 43 | 43 | 0.95 | 0.91 |
| P20338 | RAB4A | 4 | 4 | 0.95 | 0.98 |

| Accession | Name | # Peptides | # Unique Peptides | Abundance Ratio: day 3 / day 0 | Abundance Ratio: day 7/ day 0 |
|---|---|---|---|---|---|
| Q92845 | KIFAP3 | 2 | 1 | 0.01 | 0.97 |
| Q96BM9 | ARL8A | 2 | 1 | 0.01 | 0.76 |
| Q96HY6 | DDRGK1 | 4 | 4 | 0.95 | 0.90 |
| Q86V81 | ALYREF | 6 | 6 | 0.95 | 0.71 |
| P26447 | S100A4 | 6 | 6 | 0.95 | 0.68 |
| P61916 | NPC2 | 7 | 7 | 0.95 | 1.04 |
| Q8WXF1 | PSPC1 | 8 | 6 | 0.95 | 0.63 |
| Q9UGT4 | SUSD2 | 9 | 9 | 0.95 | 0.61 |
| P35610 | SOAT1 | 2 | 2 | 0.95 | 0.01 |
| Q99470 | SDF2 | 5 | 5 | 0.95 | 0.97 |
| O60264 | SMARCA5 | 6 | 4 | 0.95 | 0.90 |
| Q92783 | STAM | 6 | 5 | 0.95 | 0.84 |
| Q12979 | ABR | 6 | 6 | 0.95 | 0.96 |
| P63000 | RAC1 | 8 | 7 | 0.95 | 0.78 |
| P61019 | RAB2A | 13 | 6 | 0.95 | 1.27 |
| Q969G5 | CAVIN3 | 13 | 13 | 0.95 | 0.74 |
| Q15424 | SAFB | 21 | 13 | 0.95 | 0.83 |
| O43586 | PSTPIP1 | 3 | 2 | 0.95 | 0.01 |
| Q14139 | UBE4A | 4 | 4 | 0.95 | 0.98 |
| Q9Y2G5 | POFUT2 | 8 | 8 | 0.95 | 1.20 |
| Q7L576 | CYFIP1 | 20 | 20 | 0.95 | 0.87 |
| P08237 | PFKM | 25 | 22 | 0.95 | 0.91 |
| P35658 | NUP214 | 6 | 6 | 0.95 | 0.94 |
| Q15050 | RRS1 | 10 | 10 | 0.95 | 0.77 |
| Q96KG9 | SCYL1 | 11 | 11 | 0.95 | 0.87 |
| Q9BS40 | LXN | 4 | 4 | 0.95 | 0.65 |
| Q7RTV0 | PHF5A | 4 | 4 | 0.95 | 0.80 |
| Q86YR5 | GPSM1 | 5 | 5 | 0.95 | 0.74 |
| Q9NX46 | ADPRS | 10 | 10 | 0.95 | 0.80 |
| Q8IXB1 | DNAJC10 | 10 | 10 | 0.95 | 0.85 |
| P19784 | CSNK2A2 | 4 | 4 | 0.96 | 0.01 |
| O96019 | ACTL6A | 5 | 5 | 0.96 | 1.15 |
| P01889 | HLA-B | 8 | 3 | 0.96 | 0.53 |
| P53041 | PPP5C | 8 | 8 | 0.96 | 0.95 |
| P46379 | BAG6 | 9 | 9 | 0.96 | 1.14 |
| O95831 | AIFM1 | 10 | 10 | 0.96 | 0.83 |
| Q96KP4 | CNDP2 | 11 | 11 | 0.96 | 0.83 |
| P43243 | MATR3 | 28 | 28 | 0.96 | 0.83 |
| Q9UP83 | COG5 | 4 | 4 | 0.96 | 1.07 |
| Q00059 | TFAM | 5 | 5 | 0.96 | 0.72 |
| Q96KC8 | DNAJC1 | 6 | 6 | 0.96 | 1.14 |
| Q92572 | AP3S1 | 6 | 6 | 0.96 | 0.85 |
| P19525 | EIF2AK2 | 7 | 7 | 0.96 | 0.84 |

| Accession | Name | # Peptides | # Unique Peptides | Abundance Ratio: day 3 / day 0 | Abundance Ratio: day 7/ day 0 |
|---|---|---|---|---|---|
| Q92845 | KIFAP3 | 2 | 1 | 0.01 | 0.97 |
| Q96BM9 | ARL8A | 2 | 1 | 0.01 | 0.76 |
| P08754 | GNAI3 | 10 | 5 | 0.96 | 0.96 |
| Q92734 | TFG | 10 | 10 | 0.96 | 0.89 |
| Q13442 | PDAP1 | 11 | 11 | 0.96 | 1.08 |
| Q969U7 | PSMG2 | 3 | 3 | 0.96 | 1.15 |
| P30043 | BLVRB | 5 | 5 | 0.96 | 0.73 |
| Q8NBJ7 | SUMF2 | 8 | 8 | 0.96 | 1.12 |
| A1L0T0 | ILVBL | 8 | 8 | 0.96 | 0.80 |
| Q8IX12 | CCAR1 | 9 | 9 | 0.96 | 0.78 |
| P08621 | SNRNP70 | 13 | 13 | 0.96 | 0.78 |
| Q9HB40 | SCPEP1 | 5 | 5 | 0.96 | 0.92 |
| O95834 | EML2 | 6 | 6 | 0.96 | 0.70 |
| P33908 | MAN1A1 | 6 | 6 | 0.96 | 0.01 |
| Q9HCE1 | MOV10 | 7 | 7 | 0.96 | 0.89 |
| P60903 | S100A10 | 7 | 7 | 0.96 | 0.71 |
| Q15365 | PCBP1 | 11 | 7 | 0.96 | 0.89 |
| Q9UNH7 | SNX6 | 15 | 14 | 0.96 | 0.91 |
| P22234 | PAICS | 17 | 17 | 0.96 | 0.93 |
| Q71UM5 | RPS27L | 3 | 1 | 0.96 | 1.30 |
| O60573 | EIF4E2 | 4 | 4 | 0.96 | 0.92 |
| Q99471 | PFDN5 | 4 | 4 | 0.96 | 0.82 |
| Q9H3K6 | BOLA2 | 6 | 6 | 0.96 | 0.90 |
| Q9UIA9 | XPO7 | 8 | 8 | 0.96 | 1.23 |
| P55735 | SEC13 | 9 | 9 | 0.96 | 0.96 |
| P29692 | EEF1D | 11 | 9 | 0.96 | 0.93 |
| Q9UBT2 | UBA2 | 12 | 12 | 0.96 | 0.95 |
| Q15075 | EEA1 | 67 | 67 | 0.96 | 0.90 |
| Q86UP2 | KTN1 | 68 | 68 | 0.96 | 0.86 |
| Q969M3 | YIPF5 | 2 | 2 | 0.96 | 1.13 |
| Q9BRR6 | ADPGK | 4 | 4 | 0.96 | 1.03 |
| Q9P0M6 | MACROH2A2 | 7 | 6 | 0.96 | 0.90 |
| Q13190 | STX5 | 7 | 7 | 0.96 | 1.10 |
| Q9UNS2 | COPS3 | 8 | 8 | 0.96 | 1.06 |
| Q6P996 | PDXDC1 | 10 | 10 | 0.96 | 1.38 |
| P67775 | PPP2CA | 12 | 1 | 0.96 | 0.92 |
| O43252 | PAPSS1 | 17 | 13 | 0.96 | 1.07 |
| Q53EP0 | FNDC3B | 19 | 19 | 0.96 | 1.47 |
| Q8WWM9 | CYGB | 4 | 4 | 0.96 | 0.01 |
| Q9UMX5 | NENF | 5 | 5 | 0.96 | 0.95 |
| P30419 | NMT1 | 11 | 11 | 0.96 | 0.88 |
| O94874 | UFL1 | 21 | 20 | 0.96 | 1.02 |
| P49914 | MTHFS | 2 | 2 | 0.96 | 0.98 |

| Accession | Name | # Peptides | # Unique Peptides | Abundance Ratio: day 3 / day 0 | Abundance Ratio: day 7/ day 0 |
|---|---|---|---|---|---|
| Q92845 | KIFAP3 | 2 | 1 | 0.01 | 0.97 |
| Q96BM9 | ARL8A | 2 | 1 | 0.01 | 0.76 |
| Q7Z434 | MAVS | 5 | 5 | 0.96 | 0.89 |
| Q9BVP2 | GNL3 | 5 | 5 | 0.96 | 0.87 |
| O95870 | ABHD16A | 6 | 6 | 0.96 | 0.74 |
| O75832 | PSMD10 | 7 | 7 | 0.96 | 0.88 |
| Q9BZE4 | GTPBP4 | 8 | 8 | 0.96 | 0.77 |
| O00186 | STXBP3 | 9 | 9 | 0.96 | 0.83 |
| Q13126 | MTAP | 10 | 10 | 0.96 | 0.80 |
| Q8TDZ2 | MICAL1 | 12 | 12 | 0.96 | 0.65 |
| P04899 | GNAI2 | 15 | 10 | 0.96 | 0.86 |
| Q96L92 | SNX27 | 2 | 2 | 0.96 | 0.97 |
| Q7Z4V5 | HDGFL2 | 9 | 8 | 0.96 | 0.75 |
| Q9NVD7 | PARVA | 10 | 10 | 0.96 | 0.88 |
| P42166 | TMPO | 12 | 12 | 0.96 | 0.75 |
| P30740 | SERPINB1 | 15 | 15 | 0.96 | 0.79 |
| Q9NWZ3 | IRAK4 | 2 | 2 | 0.96 | 0.01 |
| Q9NQ48 | LZTFL1 | 6 | 6 | 0.96 | 0.89 |
| Q03154 | ACY1 | 9 | 9 | 0.96 | 0.79 |
| Q9BZQ8 | NIBAN1 | 9 | 9 | 0.96 | 0.93 |
| O14737 | PDCD5 | 11 | 11 | 0.96 | 0.95 |
| Q08257 | CRYZ | 12 | 12 | 0.96 | 0.95 |
| P27361 | MAPK3 | 15 | 10 | 0.96 | 0.83 |
| Q14258 | TRIM25 | 16 | 16 | 0.96 | 0.77 |
| P46940 | IQGAP1 | 83 | 78 | 0.96 | 0.83 |
| Q12846 | STX4 | 4 | 4 | 0.97 | 0.74 |
| Q53FA7 | TP53I3 | 8 | 8 | 0.97 | 0.81 |
| Q9Y5X3 | SNX5 | 12 | 11 | 0.97 | 1.01 |
| P35241 | RDX | 39 | 21 | 0.97 | 0.96 |
| Q16647 | PTGIS | 2 | 2 | 0.97 | 0.82 |
| Q8WUM0 | NUP133 | 3 | 3 | 0.97 | 0.01 |
| Q9Y2V2 | CARHSP1 | 3 | 3 | 0.97 | 0.60 |
| P53367 | ARFIP1 | 6 | 6 | 0.97 | 0.91 |
| Q9Y3C6 | PPIL1 | 6 | 6 | 0.97 | 0.79 |
| Q96IU4 | ABHD14B | 8 | 8 | 0.97 | 0.74 |
| P60891 | PRPS1 | 9 | 5 | 0.97 | 1.09 |
| P46926 | GNPDA1 | 12 | 8 | 0.97 | 0.78 |
| O00534 | VWA5A | 13 | 13 | 0.97 | 0.80 |
| P49585 | PCYT1A | 13 | 13 | 0.97 | 0.81 |
| Q6PKG0 | LARP1 | 17 | 17 | 0.97 | 1.07 |
| P31150 | GDI1 | 21 | 14 | 0.97 | 0.88 |
| Q9H4G0 | EPB41L1 | 22 | 20 | 0.97 | 0.68 |
| Q8NF91 | SYNE1 | 25 | 25 | 0.97 | 0.71 |

| Accession | Name | # Peptides | # Unique Peptides | Abundance Ratio: day 3 / day 0 | Abundance Ratio: day 7/ day 0 |
|---|---|---|---|---|---|
| Q92845 | KIFAP3 | 2 | 1 | 0.01 | 0.97 |
| Q96BM9 | ARL8A | 2 | 1 | 0.01 | 0.76 |
| Q9P258 | RCC2 | 4 | 4 | 0.97 | 1.03 |
| P62312 | LSM6 | 4 | 4 | 0.97 | 0.76 |
| Q12769 | NUP160 | 5 | 5 | 0.97 | 0.87 |
| Q99653 | CHP1 | 6 | 6 | 0.97 | 0.83 |
| Q14344 | GNA13 | 8 | 7 | 0.97 | 0.81 |
| P10321 | HLA-C | 9 | 4 | 0.97 | 0.70 |
| Q32MZ4 | LRRFIP1 | 12 | 11 | 0.97 | 0.95 |
| P47897 | QARS1 | 28 | 28 | 0.97 | 0.94 |
| Q9UKT5 | FBXO4 | 2 | 2 | 0.97 | 1.37 |
| Q15555 | MAPRE2 | 10 | 9 | 0.97 | 0.84 |
| P48059 | LIMS1 | 11 | 7 | 0.97 | 1.00 |
| P14314 | PRKCSH | 23 | 23 | 0.97 | 0.97 |
| Q93008 | USP9X | 29 | 29 | 0.97 | 0.85 |
| P55786 | NPEPPS | 31 | 31 | 0.97 | 0.76 |
| P0DMV8 | HSPA1A | 37 | 30 | 0.97 | 0.86 |
| P50452 | SERPINB8 | 4 | 3 | 0.97 | 0.01 |
| P29353 | SHC1 | 4 | 4 | 0.97 | 0.93 |
| Q9UKG1 | APPL1 | 8 | 8 | 0.97 | 0.83 |
| P30040 | ERP29 | 9 | 9 | 0.97 | 1.04 |
| Q9P0L0 | VAPA | 13 | 12 | 0.97 | 0.85 |
| Q14956 | GPNMB | 2 | 2 | 0.97 | 0.71 |
| Q8TF05 | PPP4R1 | 4 | 4 | 0.97 | 1.06 |
| O00422 | SAP18 | 5 | 5 | 0.97 | 0.74 |
| Q5T6V5 | C9orf64 | 9 | 9 | 0.97 | 0.93 |
| P07858 | CTSB | 11 | 11 | 0.97 | 0.66 |
| Q9Y265 | RUVBL1 | 14 | 14 | 0.97 | 0.84 |
| Q96PE7 | MCEE | 2 | 2 | 0.97 | 0.71 |
| Q96TC7 | RMDN3 | 4 | 4 | 0.97 | 0.67 |
| Q13427 | PPIG | 5 | 5 | 0.97 | 0.01 |
| O95747 | OXSR1 | 8 | 5 | 0.97 | 0.91 |
| Q92905 | COPS5 | 8 | 8 | 0.97 | 0.88 |
| P51608 | MECP2 | 9 | 9 | 0.97 | 1.04 |
| Q8WWY3 | PRPF31 | 9 | 9 | 0.97 | 0.79 |
| Q14165 | MLEC | 12 | 12 | 0.97 | 1.13 |
| Q9UJU6 | DBNL | 15 | 15 | 0.97 | 0.94 |
| Q14764 | MVP | 40 | 40 | 0.97 | 0.71 |
| P62318 | SNRPD3 | 3 | 3 | 0.97 | 0.84 |
| Q9ULF5 | SLC39A10 | 2 | 2 | 0.97 | 0.01 |
| Q99538 | LGMN | 4 | 4 | 0.97 | 0.86 |
| Q9H3N1 | TMX1 | 5 | 5 | 0.97 | 1.07 |
| Q14558 | PRPSAP1 | 5 | 5 | 0.97 | 0.99 |

| Accession | Name | # Peptides | # Unique Peptides | Abundance Ratio: day 3 / day 0 | Abundance Ratio: day 7/ day 0 |
|---|---|---|---|---|---|
| Q92845 | KIFAP3 | 2 | 1 | 0.01 | 0.97 |
| Q96BM9 | ARL8A | 2 | 1 | 0.01 | 0.76 |
| P84090 | ERH | 8 | 8 | 0.97 | 0.78 |
| P55010 | EIF5 | 13 | 13 | 0.97 | 0.82 |
| Q9BXP5 | SRRT | 15 | 15 | 0.97 | 0.81 |
| P12956 | XRCC6 | 26 | 26 | 0.97 | 0.85 |
| Q92945 | KHSRP | 32 | 26 | 0.97 | 0.73 |
| Q05397 | PTK2 | 4 | 4 | 0.97 | 0.01 |
| Q9Y295 | DRG1 | 5 | 4 | 0.97 | 1.33 |
| Q99598 | TSNAX | 7 | 7 | 0.97 | 0.98 |
| Q08170 | SRSF4 | 13 | 9 | 0.97 | 0.82 |
| Q13510 | ASAH1 | 13 | 13 | 0.97 | 0.64 |
| O60506 | SYNCRIP | 26 | 18 | 0.97 | 0.88 |
| P51668 | UBE2D1 | 3 | 2 | 0.98 | 0.69 |
| Q13315 | ATM | 3 | 3 | 0.98 | 0.01 |
| P60953 | CDC42 | 6 | 5 | 0.98 | 0.90 |
| Q9NWV8 | BABAM1 | 6 | 6 | 0.98 | 0.01 |
| Q9H7B2 | RPF2 | 3 | 3 | 0.98 | 0.72 |
| Q01459 | CTBS | 4 | 4 | 0.98 | 0.97 |
| Q9UQN3 | CHMP2B | 5 | 5 | 0.98 | 0.96 |
| Q13616 | CUL1 | 8 | 8 | 0.98 | 0.88 |
| Q92747 | ARPC1A | 9 | 9 | 0.98 | 0.88 |
| P52758 | RIDA | 3 | 3 | 0.98 | 0.85 |
| Q6GMV3 | PTRHD1 | 4 | 4 | 0.98 | 1.02 |
| O75934 | BCAS2 | 5 | 5 | 0.98 | 0.92 |
| P09012 | SNRPA | 10 | 9 | 0.98 | 0.78 |
| Q13228 | SELENBP1 | 20 | 20 | 0.98 | 0.79 |
| Q15029 | EFTUD2 | 21 | 20 | 0.98 | 0.81 |
| Q641Q2 | WASHC2A | 8 | 1 | 0.98 | 0.93 |
| Q9NP61 | ARFGAP3 | 15 | 15 | 0.98 | 1.35 |
| O94832 | MYO1D | 32 | 32 | 0.98 | 0.93 |
| Q9H9Q2 | COPS7B | 3 | 3 | 0.98 | 0.01 |
| P26885 | FKBP2 | 8 | 8 | 0.98 | 1.02 |
| P31749 | AKT1 | 9 | 7 | 0.98 | 0.92 |
| Q9Y570 | PPME1 | 9 | 9 | 0.98 | 1.01 |
| Q14914 | PTGR1 | 10 | 10 | 0.98 | 0.85 |
| O15355 | PPM1G | 11 | 11 | 0.98 | 0.83 |
| P48147 | PREP | 20 | 20 | 0.98 | 0.98 |
| Q5T4S7 | UBR4 | 28 | 28 | 0.98 | 1.03 |
| O95782 | AP2A1 | 34 | 24 | 0.98 | 0.87 |
| Q99961 | SH3GL1 | 13 | 13 | 0.98 | 0.92 |
| O43818 | RRP9 | 4 | 4 | 0.98 | 0.84 |
| Q9H6Z4 | RANBP3 | 6 | 6 | 0.98 | 0.84 |

| Accession | Name | # Peptides | # Unique Peptides | Abundance Ratio: day 3 / day 0 | Abundance Ratio: day 7/ day 0 |
|---|---|---|---|---|---|
| Q92845 | KIFAP3 | 2 | 1 | 0.01 | 0.97 |
| Q96BM9 | ARL8A | 2 | 1 | 0.01 | 0.76 |
| Q9Y5Z4 | HEBP2 | 7 | 7 | 0.98 | 0.86 |
| P43487 | RANBP1 | 8 | 8 | 0.98 | 0.84 |
| Q99729 | HNRNPAB | 11 | 10 | 0.98 | 0.93 |
| Q9NYF8 | BCLAF1 | 19 | 18 | 0.98 | 1.04 |
| Q8NB25 | FAM184A | 2 | 2 | 0.98 | 0.79 |
| O15042 | U2SURP | 5 | 5 | 0.98 | 0.01 |
| P13929 | ENO3 | 7 | 2 | 0.98 | 0.85 |
| Q2M2I8 | AAK1 | 8 | 8 | 0.98 | 1.12 |
| P35659 | DEK | 15 | 15 | 0.98 | 0.83 |
| P14866 | HNRNPL | 20 | 19 | 0.98 | 0.87 |
| O75874 | IDH1 | 21 | 21 | 0.98 | 0.76 |
| Q12906 | ILF3 | 30 | 30 | 0.98 | 0.83 |
| P17676 | CEBPB | 2 | 2 | 0.98 | 0.01 |
| P19532 | TFE3 | 3 | 3 | 0.98 | 1.07 |
| Q9BXK5 | BCL2L13 | 4 | 4 | 0.98 | 0.58 |
| P09601 | HMOX1 | 14 | 14 | 0.98 | 0.45 |
| Q9BZZ5 | API5 | 15 | 15 | 0.98 | 0.84 |
| Q9Y4X5 | ARIH1 | 2 | 2 | 0.98 | 0.92 |
| O15260 | SURF4 | 3 | 3 | 0.98 | 0.99 |
| P35237 | SERPINB6 | 15 | 14 | 0.98 | 0.90 |
| P20700 | LMNB1 | 39 | 37 | 0.98 | 0.78 |
| P19793 | RXRA | 2 | 2 | 0.99 | 0.01 |
| P62314 | SNRPD1 | 2 | 2 | 0.99 | 0.85 |
| Q9Y6K5 | OAS3 | 3 | 3 | 0.99 | 0.01 |
| O95352 | ATG7 | 3 | 3 | 0.99 | 0.01 |
| O00193 | SMAP | 3 | 3 | 0.99 | 0.69 |
| Q9P2K8 | EIF2AK4 | 4 | 4 | 0.99 | 0.81 |
| Q13630 | GFUS | 5 | 5 | 0.99 | 1.03 |
| Q9H307 | PNN | 9 | 9 | 0.99 | 0.82 |
| O00461 | GOLIM4 | 10 | 10 | 0.99 | 1.46 |
| O95425 | SVIL | 12 | 12 | 0.99 | 1.12 |
| Q9BTE3 | MCMBP | 4 | 4 | 0.99 | 1.10 |
| Q15631 | TSN | 12 | 12 | 0.99 | 0.99 |
| Q6NZI2 | CAVIN1 | 24 | 24 | 0.99 | 0.62 |
| P50914 | RPL14 | 10 | 10 | 0.99 | 0.97 |
| Q12907 | LMAN2 | 12 | 12 | 0.99 | 1.25 |
| P09493 | TPM1 | 31 | 13 | 0.99 | 1.59 |
| P14625 | HSP90B1 | 57 | 55 | 0.99 | 1.14 |
| Q01780 | EXOSC10 | 3 | 3 | 0.99 | 0.01 |
| Q9NZ45 | CISD1 | 4 | 4 | 0.99 | 0.87 |
| Q9Y237 | PIN4 | 5 | 5 | 0.99 | 0.99 |

| Accession | Name | # Peptides | # Unique Peptides | Abundance Ratio: day 3 / day 0 | Abundance Ratio: day 7/ day 0 |
|---|---|---|---|---|---|
| Q92845 | KIFAP3 | 2 | 1 | 0.01 | 0.97 |
| Q96BM9 | ARL8A | 2 | 1 | 0.01 | 0.76 |
| Q05209 | PTPN12 | 7 | 7 | 0.99 | 0.75 |
| Q9Y6W5 | WASF2 | 9 | 9 | 0.99 | 1.03 |
| P29992 | GNA11 | 10 | 7 | 0.99 | 0.80 |
| O60568 | PLOD3 | 20 | 20 | 0.99 | 1.05 |
| O60763 | USO1 | 26 | 26 | 0.99 | 1.06 |
| Q6DKI1 | RPL7L1 | 3 | 3 | 0.99 | 0.01 |
| P16234 | PDGFRA | 14 | 12 | 0.99 | 0.68 |
| Q9NR45 | NANS | 16 | 16 | 0.99 | 0.80 |
| O43390 | HNRNPR | 23 | 15 | 0.99 | 0.85 |
| Q6P2Q9 | PRPF8 | 44 | 44 | 0.99 | 0.85 |
| P49720 | PSMB3 | 4 | 4 | 0.99 | 0.88 |
| Q92599 | SEPTIN8 | 9 | 6 | 0.99 | 0.82 |
| Q5SW79 | CEP170 | 9 | 9 | 0.99 | 0.91 |
| O75608 | LYPLA1 | 4 | 4 | 0.99 | 0.90 |
| O75323 | NIPSNAP2 | 5 | 3 | 0.99 | 0.67 |
| Q9UHD2 | TBK1 | 5 | 5 | 0.99 | 1.06 |
| P49757 | NUMB | 6 | 6 | 0.99 | 0.01 |
| Q9Y5X2 | SNX8 | 8 | 8 | 0.99 | 0.76 |
| Q9ULA0 | DNPEP | 8 | 8 | 0.99 | 0.86 |
| P62820 | RAB1A | 11 | 6 | 0.99 | 1.19 |
| Q96CW1 | AP2M1 | 17 | 17 | 0.99 | 0.87 |
| Q92688 | ANP32B | 7 | 5 | 0.99 | 0.79 |
| Q9Y383 | LUC7L2 | 11 | 7 | 0.99 | 0.86 |
| P52888 | THOP1 | 14 | 14 | 0.99 | 0.94 |
| Q8N3Y1 | FBXW8 | 2 | 2 | 0.99 | 0.89 |
| Q13164 | MAPK7 | 5 | 5 | 0.99 | 0.01 |
| Q9P013 | CWC15 | 5 | 5 | 0.99 | 1.08 |
| Q00796 | SORD | 10 | 10 | 0.99 | 0.83 |
| P00966 | ASS1 | 17 | 17 | 0.99 | 1.52 |
| O60664 | PLIN3 | 20 | 20 | 0.99 | 0.87 |
| P17252 | PRKCA | 22 | 21 | 0.99 | 0.92 |
| P04843 | RPN1 | 31 | 31 | 0.99 | 1.14 |
| Q96K37 | SLC35E1 | 2 | 2 | 0.99 | 1.83 |
| P26640 | VARS1 | 34 | 34 | 0.99 | 0.98 |
| P22455 | FGFR4 | 2 | 1 | 1.00 | 0.01 |
| Q9UGP8 | SEC63 | 12 | 12 | 1.00 | 1.20 |
| Q70UQ0 | IKBIP | 19 | 19 | 1.00 | 1.07 |
| Q9NZN4 | EHD2 | 23 | 22 | 1.00 | 0.77 |
| Q712K3 | UBE2R2 | 4 | 3 | 1.00 | 0.99 |
| Q9ULC4 | MCTS1 | 6 | 6 | 1.00 | 0.85 |
| Q92696 | RABGGTA | 7 | 7 | 1.00 | 0.96 |

| Accession | Name | # Peptides | # Unique Peptides | Abundance Ratio: day 3 / day 0 | Abundance Ratio: day 7/ day 0 |
|---|---|---|---|---|---|
| Q92845 | KIFAP3 | 2 | 1 | 0.01 | 0.97 |
| Q96BM9 | ARL8A | 2 | 1 | 0.01 | 0.76 |
| Q15042 | RAB3GAP1 | 8 | 8 | 1.00 | 1.07 |
| O94919 | ENDOD1 | 9 | 9 | 1.00 | 0.83 |
| P30876 | POLR2B | 10 | 10 | 1.00 | 0.88 |
| Q16822 | PCK2 | 10 | 10 | 1.00 | 1.00 |
| Q15717 | ELAVL1 | 12 | 12 | 1.00 | 0.89 |
| Q13464 | ROCK1 | 21 | 18 | 1.00 | 0.65 |
| Q9H4A4 | RNPEP | 21 | 21 | 1.00 | 0.88 |
| Q13263 | TRIM28 | 23 | 23 | 1.00 | 0.90 |
| Q9P2J5 | LARS1 | 31 | 31 | 1.00 | 1.02 |
| P51571 | SSR4 | 5 | 5 | 1.00 | 1.21 |
| P20340 | RAB6A | 9 | 9 | 1.00 | 1.06 |
| O95865 | DDAH2 | 10 | 9 | 1.00 | 0.89 |
| Q15642 | TRIP10 | 10 | 10 | 1.00 | 1.03 |
| Q13838 | DDX39B | 20 | 8 | 1.00 | 0.71 |
| Q14240 | EIF4A2 | 23 | 10 | 1.00 | 1.06 |
| Q9UIC8 | LCMT1 | 2 | 2 | 1.00 | 1.10 |
| Q66PJ3 | ARL6IP4 | 2 | 2 | 1.00 | 1.08 |
| P62330 | ARF6 | 3 | 2 | 1.00 | 0.92 |
| Q86VN1 | VPS36 | 4 | 4 | 1.00 | 1.03 |
| Q9NZ43 | USE1 | 4 | 4 | 1.00 | 1.15 |
| Q5T0F9 | CC2D1B | 5 | 5 | 1.00 | 0.82 |
| P78406 | RAE1 | 7 | 7 | 1.00 | 0.90 |
| O94973 | AP2A2 | 21 | 11 | 1.00 | 1.07 |
| P13010 | XRCC5 | 32 | 32 | 1.00 | 0.90 |
| Q9Y2H0 | DLGAP4 | 6 | 6 | 1.00 | 0.75 |
| P22061 | PCMT1 | 8 | 8 | 1.00 | 0.90 |
| P27144 | AK4 | 8 | 8 | 1.00 | 0.89 |
| O43809 | NUDT21 | 10 | 10 | 1.00 | 0.84 |
| Q96FW1 | OTUB1 | 12 | 12 | 1.00 | 0.89 |
| O14776 | TCERG1 | 13 | 13 | 1.00 | 1.03 |
| P22059 | OSBP | 14 | 14 | 1.00 | 1.01 |
| O00116 | AGPS | 15 | 15 | 1.00 | 0.99 |
| P61981 | YWHAG | 19 | 13 | 1.00 | 0.87 |
| P38159 | RBMX | 28 | 11 | 1.00 | 0.76 |
| Q00839 | HNRNPU | 39 | 39 | 1.00 | 0.88 |
| O75170 | PPP6R2 | 2 | 2 | 1.00 | 0.97 |
| Q9Y6D5 | ARFGEF2 | 5 | 3 | 1.00 | 0.93 |
| P17174 | GOT1 | 11 | 11 | 1.00 | 0.83 |
| O75436 | VPS26A | 11 | 11 | 1.00 | 0.76 |
| Q9Y3F4 | STRAP | 12 | 12 | 1.00 | 1.00 |
| Q6IBS0 | TWF2 | 14 | 11 | 1.00 | 0.95 |

| Accession | Name | # Peptides | # Unique Peptides | Abundance Ratio: day 3 / day 0 | Abundance Ratio: day 7/ day 0 |
|---|---|---|---|---|---|
| Q92845 | KIFAP3 | 2 | 1 | 0.01 | 0.97 |
| Q96BM9 | ARL8A | 2 | 1 | 0.01 | 0.76 |
| O14647 | CHD2 | 2 | 1 | 1.00 | 0.01 |
| Q9H8Y5 | ANKZF1 | 3 | 3 | 1.00 | 1.65 |
| P18615 | NELFE | 3 | 3 | 1.00 | 1.01 |
| P17302 | GJA1 | 9 | 9 | 1.00 | 1.12 |
| Q9Y450 | HBS1L | 11 | 11 | 1.00 | 1.01 |
| Q16658 | FSCN1 | 27 | 27 | 1.00 | 0.90 |
| Q9UHP3 | USP25 | 2 | 2 | 1.00 | 0.90 |
| Q4J6C6 | PREPL | 2 | 2 | 1.00 | 0.01 |
| Q96A26 | FAM162A | 3 | 3 | 1.00 | 0.84 |
| P10620 | MGST1 | 5 | 5 | 1.00 | 0.98 |
| P30044 | PRDX5 | 5 | 5 | 1.00 | 0.98 |
| Q9ULZ3 | PYCARD | 5 | 5 | 1.00 | 0.79 |
| O95336 | PGLS | 9 | 9 | 1.00 | 0.97 |
| Q14789 | GOLGB1 | 29 | 29 | 1.00 | 1.14 |
| Q9Y316 | MEMO1 | 4 | 4 | 1.00 | 0.72 |
| Q92769 | HDAC2 | 7 | 4 | 1.00 | 1.22 |
| Q96EE3 | SEH1L | 7 | 7 | 1.00 | 1.04 |
| O60936 | NOL3 | 7 | 7 | 1.00 | 1.02 |
| Q52LJ0 | FAM98B | 8 | 5 | 1.00 | 0.98 |
| Q16666 | IFI16 | 8 | 8 | 1.00 | 0.70 |
| P46108 | CRK | 10 | 10 | 1.00 | 0.88 |
| P07237 | P4HB | 39 | 39 | 1.00 | 1.20 |
| Q9NPA8 | ENY2 | 2 | 2 | 1.00 | 0.87 |
| O43719 | HTATSF1 | 8 | 8 | 1.00 | 1.08 |
| Q96CX2 | KCTD12 | 10 | 10 | 1.00 | 1.11 |
| Q8WVV9 | HNRNPLL | 2 | 1 | 1.01 | 0.80 |
| Q92633 | LPAR1 | 2 | 2 | 1.01 | 0.72 |
| Q9Y3D9 | MRPS23 | 3 | 3 | 1.01 | 0.01 |
| Q9UEU0 | VTI1B | 4 | 4 | 1.01 | 0.88 |
| Q8WZA0 | LZIC | 6 | 6 | 1.01 | 0.98 |
| Q86UE4 | MTDH | 20 | 20 | 1.01 | 1.07 |
| Q8NFH4 | NUP37 | 2 | 2 | 1.01 | 0.01 |
| P42574 | CASP3 | 4 | 4 | 1.01 | 0.77 |
| Q6UXN9 | WDR82 | 5 | 5 | 1.01 | 0.87 |
| Q14678 | KANK1 | 6 | 5 | 1.01 | 0.01 |
| Q92791 | P3H4 | 6 | 6 | 1.01 | 1.92 |
| Q9NT62 | ATG3 | 6 | 6 | 1.01 | 1.12 |
| Q14677 | CLINT1 | 9 | 9 | 1.01 | 0.90 |
| Q86VS8 | HOOK3 | 14 | 14 | 1.01 | 1.10 |
| P42345 | MTOR | 2 | 2 | 1.01 | 1.02 |
| P52594 | AGFG1 | 4 | 3 | 1.01 | 0.89 |

| Accession | Name | # Peptides | # Unique Peptides | Abundance Ratio: day 3 / day 0 | Abundance Ratio: day 7/ day 0 |
|---|---|---|---|---|---|
| Q92845 | KIFAP3 | 2 | 1 | 0.01 | 0.97 |
| Q96BM9 | ARL8A | 2 | 1 | 0.01 | 0.76 |
| O75494 | SRSF10 | 7 | 7 | 1.01 | 0.86 |
| P05023 | ATP1A1 | 42 | 37 | 1.01 | 0.88 |
| P04921 | GYPC | 2 | 2 | 1.01 | 0.63 |
| P46937 | YAP1 | 5 | 5 | 1.01 | 0.88 |
| Q9Y3T9 | NOC2L | 6 | 6 | 1.01 | 0.77 |
| Q66K14 | TBC1D9B | 6 | 6 | 1.01 | 1.03 |
| Q92878 | RAD50 | 18 | 18 | 1.01 | 1.00 |
| Q9NWM8 | FKBP14 | 3 | 3 | 1.01 | 0.99 |
| Q9NUP9 | LIN7C | 7 | 7 | 1.01 | 0.90 |
| P07355 | ANXA2 | 35 | 35 | 1.01 | 0.77 |
| P47974 | ZFP36L2 | 2 | 1 | 1.01 | 0.71 |
| Q5JTD0 | TJAP1 | 4 | 4 | 1.01 | 1.00 |
| Q9H993 | ARMT1 | 5 | 5 | 1.01 | 0.85 |
| Q9NWV4 | CZIB | 6 | 6 | 1.01 | 1.13 |
| Q9GZP4 | PITHD1 | 6 | 6 | 1.01 | 1.00 |
| Q58FF6 | HSP90AB4P | 7 | 1 | 1.01 | 0.85 |
| Q9UHX1 | PUF60 | 14 | 14 | 1.01 | 0.95 |
| Q13724 | MOGS | 16 | 16 | 1.01 | 1.13 |
| Q5JRA6 | MIA3 | 22 | 22 | 1.01 | 1.07 |
| P23497 | SP100 | 9 | 7 | 1.01 | 0.01 |
| P80303 | NUCB2 | 19 | 19 | 1.01 | 1.14 |
| P55196 | AFDN | 4 | 4 | 1.01 | 0.94 |
| Q8TB22 | SPATA20 | 5 | 5 | 1.01 | 0.01 |
| Q9Y333 | LSM2 | 4 | 4 | 1.01 | 0.89 |
| O95810 | CAVIN2 | 4 | 4 | 1.01 | 0.01 |
| P56182 | RRP1 | 4 | 4 | 1.01 | 0.84 |
| Q8WU90 | ZC3H15 | 9 | 9 | 1.01 | 1.32 |
| O95394 | PGM3 | 17 | 17 | 1.01 | 1.00 |
| Q9BUR5 | APOO | 3 | 3 | 1.01 | 0.01 |
| O95749 | GGPS1 | 2 | 2 | 1.02 | 0.92 |
| P61225 | RAP2B | 4 | 4 | 1.02 | 0.96 |
| Q9NZL4 | HSPBP1 | 3 | 3 | 1.02 | 0.01 |
| Q13243 | SRSF5 | 4 | 3 | 1.02 | 0.88 |
| Q99519 | NEU1 | 4 | 4 | 1.02 | 0.01 |
| Q5J8M3 | EMC4 | 4 | 4 | 1.02 | 0.85 |
| P55769 | SNU13 | 9 | 9 | 1.02 | 0.88 |
| O75351 | VPS4B | 12 | 7 | 1.02 | 1.07 |
| P00441 | SOD1 | 14 | 14 | 1.02 | 0.96 |
| Q9Y646 | CPQ | 7 | 7 | 1.02 | 1.15 |
| O60869 | EDF1 | 8 | 8 | 1.02 | 1.33 |
| P18754 | RCC1 | 8 | 8 | 1.02 | 0.87 |

| Accession | Name | # Peptides | # Unique Peptides | Abundance Ratio: day 3 / day 0 | Abundance Ratio: day 7/ day 0 |
|---|---|---|---|---|---|
| Q92845 | KIFAP3 | 2 | 1 | 0.01 | 0.97 |
| Q96BM9 | ARL8A | 2 | 1 | 0.01 | 0.76 |
| P26583 | HMGB2 | 14 | 11 | 1.02 | 0.69 |
| Q8IWE2 | FAM114A1 | 14 | 14 | 1.02 | 1.09 |
| Q99715 | COL12A1 | 116 | 116 | 1.02 | 1.93 |
| Q9BZH6 | WDR11 | 3 | 3 | 1.02 | 1.29 |
| P35813 | PPM1A | 6 | 4 | 1.02 | 0.96 |
| Q9UBE0 | SAE1 | 15 | 15 | 1.02 | 0.89 |
| Q9Y5S2 | CDC42BPB | 19 | 14 | 1.02 | 0.77 |
| O14562 | UBFD1 | 4 | 4 | 1.02 | 1.01 |
| Q9BY77 | POLDIP3 | 5 | 5 | 1.02 | 1.08 |
| Q96T37 | RBM15 | 5 | 5 | 1.02 | 0.81 |
| P20936 | RASA1 | 7 | 7 | 1.02 | 0.01 |
| P10768 | ESD | 10 | 10 | 1.02 | 0.84 |
| Q92499 | DDX1 | 26 | 26 | 1.02 | 1.10 |
| Q9NVG8 | TBC1D13 | 2 | 2 | 1.02 | 1.13 |
| Q9Y5S9 | RBM8A | 6 | 6 | 1.02 | 0.86 |
| Q15819 | UBE2V2 | 8 | 4 | 1.02 | 1.04 |
| Q12768 | WASHC5 | 9 | 9 | 1.02 | 1.02 |
| P37837 | TALDO1 | 20 | 20 | 1.02 | 0.96 |
| P41250 | GARS1 | 23 | 23 | 1.02 | 1.15 |
| P55854 | SUMO3 | 2 | 1 | 1.02 | 0.01 |
| Q9BPX5 | ARPC5L | 4 | 4 | 1.02 | 0.87 |
| O00170 | AIP | 4 | 4 | 1.02 | 0.75 |
| P46109 | CRKL | 7 | 7 | 1.02 | 0.90 |
| P17813 | ENG | 10 | 10 | 1.02 | 0.56 |
| P13798 | APEH | 14 | 14 | 1.02 | 0.78 |
| P16152 | CBR1 | 15 | 13 | 1.02 | 0.84 |
| P51858 | HDGF | 21 | 20 | 1.02 | 0.87 |
| P17858 | PFKL | 26 | 19 | 1.02 | 1.08 |
| Q9NUQ9 | CYRIB | 4 | 3 | 1.02 | 0.01 |
| Q9NRN7 | AASDHPPT | 4 | 4 | 1.02 | 0.01 |
| P61457 | PCBD1 | 5 | 5 | 1.02 | 0.60 |
| O43488 | AKR7A2 | 10 | 10 | 1.02 | 0.92 |
| Q14683 | SMC1A | 12 | 12 | 1.02 | 0.89 |
| P40925 | MDH1 | 17 | 17 | 1.02 | 0.88 |
| P04406 | GAPDH | 26 | 26 | 1.02 | 0.93 |
| P55263 | ADK | 16 | 16 | 1.02 | 1.06 |
| Q96K76 | USP47 | 9 | 9 | 1.03 | 0.89 |
| Q9UJ70 | NAGK | 14 | 14 | 1.03 | 0.89 |
| O43159 | RRP8 | 2 | 2 | 1.03 | 0.01 |
| Q8WVY7 | UBLCP1 | 5 | 5 | 1.03 | 0.72 |
| P49959 | MRE11 | 6 | 6 | 1.03 | 0.81 |

| Accession | Name | # Peptides | # Unique Peptides | Abundance Ratio: day 3 / day 0 | Abundance Ratio: day 7/ day 0 |
|---|---|---|---|---|---|
| Q92845 | KIFAP3 | 2 | 1 | 0.01 | 0.97 |
| Q96BM9 | ARL8A | 2 | 1 | 0.01 | 0.76 |
| Q15102 | PAFAH1B3 | 7 | 7 | 1.03 | 0.79 |
| Q96T51 | RUFY1 | 12 | 12 | 1.03 | 1.02 |
| P78559 | MAP1A | 53 | 53 | 1.03 | 0.74 |
| P62917 | RPL8 | 15 | 15 | 1.03 | 0.97 |
| Q9Y508 | RNF114 | 5 | 5 | 1.03 | 0.91 |
| P07741 | APRT | 5 | 5 | 1.03 | 1.01 |
| Q9BU23 | LMF2 | 7 | 7 | 1.03 | 1.08 |
| Q14696 | MESD | 8 | 8 | 1.03 | 1.07 |
| O95757 | HSPA4L | 9 | 7 | 1.03 | 0.01 |
| P21964 | COMT | 9 | 9 | 1.03 | 0.80 |
| P68366 | TUBA4A | 18 | 2 | 1.03 | 1.03 |
| Q13459 | MYO9B | 15 | 15 | 1.03 | 0.01 |
| Q16643 | DBN1 | 17 | 17 | 1.03 | 1.14 |
| P07910 | HNRNPC | 18 | 10 | 1.03 | 0.83 |
| Q9Y6Y8 | SEC23IP | 19 | 19 | 1.03 | 1.18 |
| Q01469 | FABP5 | 11 | 11 | 1.03 | 0.90 |
| Q96FJ2 | DYNLL2 | 6 | 3 | 1.03 | 1.00 |
| O43592 | XPOT | 7 | 7 | 1.03 | 0.99 |
| O95210 | STBD1 | 10 | 10 | 1.03 | 0.84 |
| Q9Y2S2 | CRYL1 | 4 | 4 | 1.03 | 0.96 |
| P61006 | RAB8A | 5 | 2 | 1.03 | 1.37 |
| O60749 | SNX2 | 21 | 20 | 1.03 | 0.97 |
| Q1KMD3 | HNRNPUL2 | 27 | 27 | 1.03 | 0.82 |
| P35580 | MYH10 | 75 | 46 | 1.03 | 0.92 |
| O75431 | MTX2 | 3 | 3 | 1.03 | 0.78 |
| Q06278 | AOX1 | 4 | 4 | 1.03 | 0.01 |
| Q01844 | EWSR1 | 8 | 8 | 1.03 | 0.78 |
| Q9NZL9 | MAT2B | 8 | 8 | 1.03 | 0.97 |
| Q10567 | AP1B1 | 23 | 9 | 1.03 | 0.84 |
| Q9Y4Z0 | LSM4 | 2 | 2 | 1.04 | 0.71 |
| O15427 | SLC16A3 | 7 | 7 | 1.04 | 0.73 |
| Q8TED1 | GPX8 | 8 | 8 | 1.04 | 1.42 |
| Q9Y3I0 | RTCB | 21 | 21 | 1.04 | 1.02 |
| Q9NS86 | LANCL2 | 3 | 3 | 1.04 | 0.97 |
| Q9HC38 | GLOD4 | 7 | 7 | 1.04 | 0.89 |
| Q9C0C2 | TNKS1BP1 | 40 | 40 | 1.04 | 1.18 |
| Q9BRF8 | CPPED1 | 9 | 9 | 1.04 | 0.78 |
| P30101 | PDIA3 | 41 | 41 | 1.04 | 1.28 |
| Q9BY43 | CHMP4A | 4 | 4 | 1.04 | 0.95 |
| Q15370 | ELOB | 5 | 5 | 1.04 | 1.07 |
| Q15057 | ACAP2 | 6 | 6 | 1.04 | 0.78 |

| Accession | Name | # Peptides | # Unique Peptides | Abundance Ratio: day 3 / day 0 | Abundance Ratio: day 7/ day 0 |
|---|---|---|---|---|---|
| Q92845 | KIFAP3 | 2 | 1 | 0.01 | 0.97 |
| Q96BM9 | ARL8A | 2 | 1 | 0.01 | 0.76 |
| Q06124 | PTPN11 | 14 | 14 | 1.04 | 0.88 |
| P11216 | PYGB | 46 | 36 | 1.04 | 0.93 |
| Q9NRZ7 | AGPAT3 | 2 | 2 | 1.04 | 0.01 |
| Q8IYB3 | SRRM1 | 3 | 3 | 1.04 | 1.24 |
| Q9NR19 | ACSS2 | 5 | 5 | 1.04 | 1.03 |
| Q5BKZ1 | ZNF326 | 5 | 5 | 1.04 | 0.79 |
| Q8IZL8 | PELP1 | 7 | 7 | 1.04 | 0.81 |
| Q8NCA5 | FAM98A | 11 | 8 | 1.04 | 1.13 |
| P54577 | YARS1 | 25 | 25 | 1.04 | 1.02 |
| P05413 | FABP3 | 8 | 8 | 1.04 | 1.16 |
| Q9NRX4 | PHPT1 | 9 | 9 | 1.04 | 1.04 |
| P14868 | DARS1 | 30 | 30 | 1.04 | 1.07 |
| Q9Y618 | NCOR2 | 4 | 4 | 1.04 | 1.14 |
| P35754 | GLRX | 4 | 4 | 1.04 | 0.70 |
| Q9Y4E8 | USP15 | 6 | 5 | 1.04 | 0.76 |
| Q9UI10 | EIF2B4 | 8 | 8 | 1.04 | 0.91 |
| Q53EL6 | PDCD4 | 11 | 11 | 1.04 | 0.69 |
| P04844 | RPN2 | 14 | 14 | 1.04 | 1.19 |
| Q9Y2W1 | THRAP3 | 24 | 23 | 1.04 | 0.88 |
| Q14657 | LAGE3 | 3 | 3 | 1.04 | 0.90 |
| Q08722 | CD47 | 4 | 4 | 1.04 | 0.84 |
| Q9Y2X3 | NOP58 | 10 | 10 | 1.04 | 0.88 |
| Q9UPU5 | USP24 | 11 | 11 | 1.04 | 0.93 |
| P04075 | ALDOA | 33 | 27 | 1.04 | 0.93 |
| Q8NBI5 | SLC43A3 | 2 | 2 | 1.04 | 0.80 |
| Q96LR5 | UBE2E2 | 4 | 2 | 1.04 | 0.01 |
| Q16401 | PSMD5 | 12 | 12 | 1.04 | 0.96 |
| P30533 | LRPAP1 | 26 | 26 | 1.04 | 0.92 |
| P62306 | SNRPF | 2 | 2 | 1.05 | 0.01 |
| P49790 | NUP153 | 4 | 4 | 1.05 | 1.13 |
| P16930 | FAH | 7 | 7 | 1.05 | 0.99 |
| Q9UHG3 | PCYOX1 | 16 | 16 | 1.05 | 0.90 |
| O95479 | H6PD | 17 | 17 | 1.05 | 0.99 |
| P09525 | ANXA4 | 21 | 21 | 1.05 | 1.04 |
| P27816 | MAP4 | 64 | 64 | 1.05 | 1.20 |
| Q92574 | TSC1 | 2 | 2 | 1.05 | 0.01 |
| Q2TAY7 | SMU1 | 9 | 9 | 1.05 | 1.05 |
| Q13428 | TCOF1 | 14 | 14 | 1.05 | 1.16 |
| Q92841 | DDX17 | 26 | 19 | 1.05 | 0.89 |
| P53990 | IST1 | 7 | 7 | 1.05 | 1.23 |
| Q14166 | TTLL12 | 8 | 8 | 1.05 | 0.97 |

| Accession | Name | # Peptides | # Unique Peptides | Abundance Ratio: day 3 / day 0 | Abundance Ratio: day 7/ day 0 |
|---|---|---|---|---|---|
| Q92845 | KIFAP3 | 2 | 1 | 0.01 | 0.97 |
| Q96BM9 | ARL8A | 2 | 1 | 0.01 | 0.76 |
| P51812 | RPS6KA3 | 9 | 9 | 1.05 | 0.91 |
| Q15293 | RCN1 | 11 | 11 | 1.05 | 1.44 |
| P49588 | AARS1 | 35 | 35 | 1.05 | 1.01 |
| Q9H2P0 | ADNP | 2 | 2 | 1.05 | 0.01 |
| Q9BRK5 | SDF4 | 9 | 9 | 1.05 | 1.26 |
| Q9UQ35 | SRRM2 | 12 | 12 | 1.05 | 1.18 |
| Q8N163 | CCAR2 | 17 | 17 | 1.05 | 0.88 |
| O14617 | AP3D1 | 19 | 19 | 1.05 | 0.94 |
| Q15020 | SART3 | 14 | 14 | 1.05 | 1.14 |
| Q8NBL1 | POGLUT1 | 4 | 4 | 1.05 | 1.06 |
| Q9Y5P6 | GMPPB | 8 | 8 | 1.05 | 1.17 |
| Q14651 | PLS1 | 4 | 1 | 1.05 | 1.77 |
| P46976 | GYG1 | 6 | 6 | 1.05 | 0.92 |
| Q99829 | CPNE1 | 7 | 7 | 1.05 | 1.01 |
| P49750 | YLPM1 | 14 | 13 | 1.05 | 1.03 |
| P21281 | ATP6V1B2 | 15 | 15 | 1.05 | 0.82 |
| O00567 | NOP56 | 15 | 15 | 1.05 | 0.90 |
| Q9P2R3 | ANKFY1 | 16 | 16 | 1.05 | 0.88 |
| Q9NX24 | NHP2 | 4 | 4 | 1.05 | 1.16 |
| O00233 | PSMD9 | 8 | 8 | 1.05 | 0.85 |
| P06756 | ITGAV | 26 | 26 | 1.05 | 0.92 |
| Q9UNL2 | SSR3 | 3 | 3 | 1.05 | 1.47 |
| O14735 | CDIPT | 4 | 4 | 1.05 | 1.11 |
| Q13867 | BLMH | 5 | 5 | 1.05 | 1.13 |
| P55210 | CASP7 | 2 | 2 | 1.05 | 0.96 |
| Q9Y5U2 | TSSC4 | 2 | 2 | 1.05 | 0.95 |
| O95295 | SNAPIN | 4 | 4 | 1.05 | 0.94 |
| O43813 | LANCL1 | 6 | 6 | 1.05 | 0.82 |
| P06733 | ENO1 | 32 | 29 | 1.05 | 0.95 |
| P52434 | POLR2H | 5 | 5 | 1.06 | 1.05 |
| P49589 | CARS1 | 21 | 21 | 1.06 | 1.02 |
| Q15018 | ABRAXAS2 | 2 | 2 | 1.06 | 1.01 |
| Q9Y3Q3 | TMED3 | 3 | 3 | 1.06 | 1.44 |
| P58546 | MTPN | 5 | 5 | 1.06 | 0.88 |
| P50148 | GNAQ | 7 | 4 | 1.06 | 0.01 |
| Q92597 | NDRG1 | 9 | 9 | 1.06 | 0.77 |
| Q15257 | PTPA | 9 | 9 | 1.06 | 0.92 |
| Q92804 | TAF15 | 12 | 10 | 1.06 | 0.89 |
| Q86YP4 | GATAD2A | 2 | 1 | 1.06 | 0.96 |
| P31323 | PRKAR2B | 3 | 1 | 1.06 | 0.01 |
| Q6ZSR9 | | 6 | 6 | 1.06 | 1.05 |

| Accession | Name | # Peptides | # Unique Peptides | Abundance Ratio: day 3 / day 0 | Abundance Ratio: day 7/ day 0 |
|---|---|---|---|---|---|
| Q92845 | KIFAP3 | 2 | 1 | 0.01 | 0.97 |
| Q96BM9 | ARL8A | 2 | 1 | 0.01 | 0.76 |
| P07203 | GPX1 | 7 | 7 | 1.06 | 1.08 |
| O00423 | EML1 | 11 | 11 | 1.06 | 1.09 |
| Q08426 | EHHADH | 3 | 3 | 1.06 | 0.01 |
| O00244 | ATOX1 | 4 | 4 | 1.06 | 1.11 |
| O00471 | EXOC5 | 7 | 6 | 1.06 | 0.01 |
| P51531 | SMARCA2 | 7 | 6 | 1.06 | 1.04 |
| Q9BZL4 | PPP1R12C | 11 | 11 | 1.06 | 0.81 |
| P09104 | ENO2 | 12 | 6 | 1.06 | 0.95 |
| Q92896 | GLG1 | 27 | 27 | 1.06 | 1.05 |
| Q86VP6 | CAND1 | 33 | 33 | 1.06 | 1.03 |
| Q8N3V7 | SYNPO | 6 | 6 | 1.06 | 0.93 |
| Q13185 | CBX3 | 7 | 7 | 1.06 | 0.84 |
| P09211 | GSTP1 | 8 | 8 | 1.06 | 1.13 |
| P35221 | CTNNA1 | 19 | 19 | 1.06 | 1.10 |
| P09960 | LTA4H | 25 | 25 | 1.06 | 0.98 |
| Q96C90 | PPP1R14B | 3 | 3 | 1.06 | 1.11 |
| Q96II8 | LRCH3 | 4 | 4 | 1.06 | 0.01 |
| P49792 | RANBP2 | 28 | 27 | 1.06 | 0.96 |
| P63279 | UBE2I | 5 | 5 | 1.06 | 0.88 |
| Q8N0X7 | SPART | 8 | 8 | 1.06 | 1.30 |
| Q5NDL2 | EOGT | 4 | 4 | 1.06 | 0.01 |
| Q8NBM8 | PCYOX1L | 2 | 2 | 1.06 | 1.01 |
| Q14103 | HNRNPD | 14 | 14 | 1.06 | 0.83 |
| Q96HE7 | ERO1A | 17 | 17 | 1.06 | 0.89 |
| Q63ZY3 | KANK2 | 20 | 19 | 1.06 | 1.00 |
| Q99759 | MAP3K3 | 2 | 1 | 1.06 | 0.73 |
| Q8N684 | CPSF7 | 8 | 8 | 1.06 | 0.92 |
| P32322 | PYCR1 | 10 | 9 | 1.06 | 1.68 |
| Q12905 | ILF2 | 14 | 14 | 1.06 | 0.90 |
| Q7Z3B4 | NUP54 | 4 | 4 | 1.07 | 0.01 |
| P52926 | HMGA2 | 5 | 5 | 1.07 | 1.02 |
| P63151 | PPP2R2A | 8 | 8 | 1.07 | 0.97 |
| Q13131 | PRKAA1 | 11 | 8 | 1.07 | 1.03 |
| P51610 | HCFC1 | 12 | 12 | 1.07 | 0.81 |
| Q02809 | PLOD1 | 26 | 26 | 1.07 | 1.47 |
| Q9BX40 | LSM14B | 3 | 3 | 1.07 | 0.01 |
| Q7Z6Z7 | HUWE1 | 54 | 54 | 1.07 | 1.00 |
| P09417 | QDPR | 3 | 3 | 1.07 | 0.66 |
| Q9NZ01 | TECR | 7 | 7 | 1.07 | 1.29 |
| Q9NWH9 | SLTM | 11 | 11 | 1.07 | 1.04 |
| P00390 | GSR | 12 | 12 | 1.07 | 0.97 |

| Accession | Name | # Peptides | # Unique Peptides | Abundance Ratio: day 3 / day 0 | Abundance Ratio: day 7/ day 0 |
|---|---|---|---|---|---|
| Q92845 | KIFAP3 | 2 | 1 | 0.01 | 0.97 |
| Q96BM9 | ARL8A | 2 | 1 | 0.01 | 0.76 |
| Q06136 | KDSR | 2 | 2 | 1.07 | 1.16 |
| Q8N1B4 | VPS52 | 3 | 3 | 1.07 | 1.15 |
| Q8IZP0 | ABI1 | 5 | 5 | 1.07 | 0.80 |
| Q99567 | NUP88 | 7 | 7 | 1.07 | 0.98 |
| P50570 | DNM2 | 19 | 12 | 1.07 | 1.15 |
| Q9UHY7 | ENOPH1 | 4 | 4 | 1.07 | 0.89 |
| Q96AX1 | VPS33A | 4 | 4 | 1.07 | 0.01 |
| Q14118 | DAG1 | 3 | 3 | 1.07 | 0.01 |
| P62633 | CNBP | 2 | 2 | 1.07 | 1.32 |
| Q9HAT2 | SIAE | 6 | 6 | 1.07 | 1.16 |
| Q00765 | REEP5 | 9 | 9 | 1.07 | 1.00 |
| P17096 | HMGA1 | 9 | 9 | 1.07 | 0.79 |
| Q6GMV2 | SMYD5 | 4 | 4 | 1.07 | 1.32 |
| Q9H4F8 | SMOC1 | 8 | 8 | 1.07 | 2.71 |
| Q9Y224 | RTRAF | 10 | 10 | 1.07 | 1.08 |
| P27695 | APEX1 | 13 | 13 | 1.07 | 0.84 |
| P06744 | GPI | 27 | 26 | 1.07 | 1.06 |
| Q9HCU5 | PREB | 5 | 5 | 1.08 | 1.11 |
| Q8N8S7 | ENAH | 24 | 23 | 1.08 | 0.98 |
| Q9P2E9 | RRBP1 | 117 | 117 | 1.08 | 1.53 |
| P50336 | PPOX | 2 | 2 | 1.08 | 0.01 |
| O95159 | ZFPL1 | 2 | 2 | 1.08 | 0.87 |
| Q8N128 | FAM177A1 | 3 | 3 | 1.08 | 1.14 |
| P62873 | GNB1 | 11 | 6 | 1.08 | 0.84 |
| P29401 | TKT | 29 | 29 | 1.08 | 0.91 |
| Q9NPE3 | NOP10 | 2 | 2 | 1.08 | 0.95 |
| O43149 | ZZEF1 | 2 | 2 | 1.08 | 1.30 |
| Q6P1N0 | CC2D1A | 4 | 3 | 1.08 | 1.03 |
| Q9Y679 | AUP1 | 4 | 4 | 1.08 | 1.17 |
| Q9BV38 | WDR18 | 6 | 6 | 1.08 | 0.87 |
| Q7Z460 | CLASP1 | 8 | 8 | 1.08 | 0.01 |
| Q9Y4I1 | MYO5A | 13 | 13 | 1.08 | 0.65 |
| P50402 | EMD | 13 | 13 | 1.08 | 0.95 |
| P08240 | SRPRA | 13 | 13 | 1.08 | 1.73 |
| Q96A33 | CCDC47 | 14 | 14 | 1.08 | 1.17 |
| P31153 | MAT2A | 15 | 15 | 1.08 | 0.83 |
| O95777 | LSM8 | 3 | 3 | 1.08 | 1.08 |
| Q5JPH6 | EARS2 | 3 | 3 | 1.08 | 0.01 |
| Q7Z2Z2 | EFL1 | 5 | 5 | 1.08 | 0.68 |
| P60468 | SEC61B | 5 | 5 | 1.08 | 1.37 |
| P22694 | PRKACB | 8 | 3 | 1.08 | 1.21 |

| Accession | Name | # Peptides | # Unique Peptides | Abundance Ratio: day 3 / day 0 | Abundance Ratio: day 7/ day 0 |
|---|---|---|---|---|---|
| Q92845 | KIFAP3 | 2 | 1 | 0.01 | 0.97 |
| Q96BM9 | ARL8A | 2 | 1 | 0.01 | 0.76 |
| Q99700 | ATXN2 | 11 | 9 | 1.08 | 0.01 |
| Q5JTV8 | TOR1AIP1 | 20 | 19 | 1.08 | 0.81 |
| P28290 | ITPRID2 | 2 | 2 | 1.08 | 0.01 |
| P21266 | GSTM3 | 9 | 7 | 1.08 | 1.20 |
| Q12972 | PPP1R8 | 2 | 2 | 1.08 | 0.94 |
| Q9BTT0 | ANP32E | 4 | 4 | 1.08 | 0.64 |
| O75223 | GGCT | 5 | 5 | 1.08 | 1.62 |
| Q07812 | BAX | 7 | 7 | 1.08 | 0.89 |
| P18669 | PGAM1 | 21 | 15 | 1.08 | 1.00 |
| Q02318 | CYP27A1 | 4 | 4 | 1.08 | 0.81 |
| O14618 | CCS | 4 | 4 | 1.08 | 0.84 |
| Q9Y4F1 | FARP1 | 12 | 12 | 1.08 | 1.04 |
| P46977 | STT3A | 16 | 14 | 1.08 | 1.39 |
| P84101 | SERF2 | 3 | 3 | 1.08 | 1.28 |
| Q7Z4H3 | HDDC2 | 4 | 4 | 1.08 | 0.01 |
| P62879 | GNB2 | 12 | 3 | 1.08 | 0.95 |
| P41208 | CETN2 | 2 | 2 | 1.08 | 1.57 |
| P09234 | SNRPC | 3 | 3 | 1.08 | 0.84 |
| Q96C86 | DCPS | 7 | 7 | 1.08 | 1.05 |
| Q14151 | SAFB2 | 13 | 5 | 1.08 | 0.91 |
| Q9Y2J2 | EPB41L3 | 14 | 9 | 1.08 | 0.74 |
| Q9NUL5 | SHFL | 7 | 7 | 1.09 | 0.92 |
| Q96AY3 | FKBP10 | 27 | 26 | 1.09 | 1.64 |
| Q12797 | ASPH | 34 | 34 | 1.09 | 1.17 |
| P84243 | H3-3A | 9 | 3 | 1.09 | 0.79 |
| Q14161 | GIT2 | 2 | 2 | 1.09 | 0.01 |
| Q7LG56 | RRM2B | 5 | 5 | 1.09 | 0.93 |
| O14787 | TNPO2 | 7 | 4 | 1.09 | 1.12 |
| P28161 | GSTM2 | 12 | 9 | 1.09 | 0.95 |
| Q14980 | NUMA1 | 74 | 73 | 1.09 | 0.83 |
| Q969T4 | UBE2E3 | 3 | 1 | 1.09 | 0.90 |
| Q6P5R6 | RPL22L1 | 3 | 3 | 1.09 | 1.23 |
| Q14232 | EIF2B1 | 7 | 7 | 1.09 | 1.03 |
| P23284 | PPIB | 18 | 18 | 1.09 | 1.44 |
| Q99996 | AKAP9 | 2 | 2 | 1.09 | 0.01 |
| Q3LXA3 | TKFC | 8 | 8 | 1.09 | 1.03 |
| O75718 | CRTAP | 14 | 14 | 1.09 | 1.72 |
| Q9UKV3 | ACIN1 | 18 | 18 | 1.09 | 1.05 |
| Q03252 | LMNB2 | 43 | 41 | 1.09 | 0.86 |
| Q8IY67 | RAVER1 | 3 | 3 | 1.09 | 0.87 |
| Q99442 | SEC62 | 7 | 7 | 1.09 | 1.15 |

| Accession | Name | # Peptides | # Unique Peptides | Abundance Ratio: day 3 / day 0 | Abundance Ratio: day 7/ day 0 |
|---|---|---|---|---|---|
| Q92845 | KIFAP3 | 2 | 1 | 0.01 | 0.97 |
| Q96BM9 | ARL8A | 2 | 1 | 0.01 | 0.76 |
| Q9HAV0 | GNB4 | 12 | 5 | 1.09 | 1.09 |
| P02545 | LMNA | 70 | 68 | 1.09 | 0.78 |
| P08670 | VIM | 71 | 65 | 1.09 | 0.80 |
| Q14320 | FAM50A | 3 | 3 | 1.09 | 1.43 |
| Q9BZV1 | UBXN6 | 4 | 4 | 1.09 | 1.15 |
| P54289 | CACNA2D1 | 5 | 5 | 1.09 | 0.87 |
| P50135 | HNMT | 2 | 2 | 1.09 | 1.05 |
| P50579 | METAP2 | 8 | 8 | 1.09 | 1.22 |
| Q99575 | POP1 | 3 | 3 | 1.09 | 0.01 |
| P09936 | UCHL1 | 12 | 12 | 1.09 | 1.04 |
| Q8IWE4 | DCUN1D3 | 4 | 4 | 1.10 | 1.16 |
| Q9NX58 | LYAR | 2 | 2 | 1.10 | 0.83 |
| O95671 | ASMTL | 7 | 7 | 1.10 | 1.38 |
| Q96ED9 | HOOK2 | 4 | 4 | 1.10 | 1.81 |
| Q9BUP3 | HTATIP2 | 6 | 6 | 1.10 | 0.85 |
| O75828 | CBR3 | 8 | 6 | 1.10 | 0.86 |
| O43237 | DYNC1LI2 | 10 | 10 | 1.10 | 1.35 |
| P30041 | PRDX6 | 13 | 13 | 1.10 | 1.16 |
| P54105 | CLNS1A | 2 | 2 | 1.10 | 1.23 |
| P62834 | RAP1A | 8 | 1 | 1.10 | 1.28 |
| P27487 | DPP4 | 28 | 28 | 1.10 | 0.74 |
| Q92621 | NUP205 | 10 | 10 | 1.10 | 0.83 |
| Q96G03 | PGM2 | 23 | 21 | 1.10 | 0.94 |
| Q04446 | GBE1 | 26 | 26 | 1.10 | 0.80 |
| Q8TDQ7 | GNPDA2 | 7 | 3 | 1.10 | 1.01 |
| O00469 | PLOD2 | 22 | 22 | 1.10 | 1.56 |
| Q5HYI8 | RABL3 | 2 | 2 | 1.10 | 0.01 |
| P35475 | IDUA | 7 | 7 | 1.10 | 1.04 |
| P19623 | SRM | 10 | 10 | 1.10 | 1.38 |
| P49257 | LMAN1 | 24 | 24 | 1.10 | 1.22 |
| Q8NDH3 | NPEPL1 | 4 | 4 | 1.11 | 0.01 |
| Q71DI3 | H3C13 | 9 | 3 | 1.11 | 0.72 |
| O60832 | DKC1 | 15 | 15 | 1.11 | 1.19 |
| Q9BZL1 | UBL5 | 2 | 2 | 1.11 | 1.12 |
| Q92504 | SLC39A7 | 4 | 4 | 1.11 | 1.27 |
| Q9Y371 | SH3GLB1 | 9 | 8 | 1.11 | 1.20 |
| Q96RF0 | SNX18 | 10 | 10 | 1.11 | 1.36 |
| Q14011 | CIRBP | 5 | 5 | 1.11 | 1.63 |
| Q14728 | MFSD10 | 3 | 3 | 1.11 | 1.04 |
| Q8IWA5 | SLC44A2 | 5 | 5 | 1.11 | 0.88 |
| P14550 | AKR1A1 | 14 | 14 | 1.11 | 1.03 |

| Accession | Name | # Peptides | # Unique Peptides | Abundance Ratio: day 3 / day 0 | Abundance Ratio: day 7/ day 0 |
|---|---|---|---|---|---|
| Q92845 | KIFAP3 | 2 | 1 | 0.01 | 0.97 |
| Q96BM9 | ARL8A | 2 | 1 | 0.01 | 0.76 |
| Q96S44 | TP53RK | 2 | 2 | 1.11 | 0.83 |
| P49354 | FNTA | 4 | 4 | 1.11 | 1.18 |
| P22087 | FBL | 14 | 14 | 1.11 | 0.91 |
| P12270 | TPR | 49 | 49 | 1.11 | 0.96 |
| Q8IWV7 | UBR1 | 2 | 2 | 1.11 | 0.96 |
| Q96AQ6 | PBXIP1 | 9 | 9 | 1.11 | 0.63 |
| Q9C0E8 | LNPK | 9 | 9 | 1.11 | 1.16 |
| P78324 | SIRPA | 12 | 12 | 1.11 | 1.16 |
| Q9NUQ7 | UFSP2 | 2 | 2 | 1.11 | 0.96 |
| P57740 | NUP107 | 4 | 4 | 1.11 | 0.95 |
| Q96GG9 | DCUN1D1 | 2 | 2 | 1.11 | 0.01 |
| Q6PCE3 | PGM2L1 | 14 | 13 | 1.11 | 0.95 |
| Q9Y2U8 | LEMD3 | 6 | 6 | 1.12 | 0.90 |
| Q9NRV9 | HEBP1 | 7 | 7 | 1.12 | 1.11 |
| Q8IVL6 | P3H3 | 11 | 11 | 1.12 | 1.82 |
| Q05519 | SRSF11 | 2 | 2 | 1.12 | 0.01 |
| P10155 | RO60 | 15 | 15 | 1.12 | 1.02 |
| O60437 | PPL | 25 | 25 | 1.12 | 1.05 |
| Q9BUJ2 | HNRNPUL1 | 17 | 17 | 1.12 | 0.85 |
| Q14573 | ITPR3 | 21 | 15 | 1.12 | 0.80 |
| Q15059 | BRD3 | 3 | 3 | 1.12 | 1.01 |
| Q12888 | TP53BP1 | 24 | 24 | 1.12 | 0.94 |
| Q9NP58 | ABCB6 | 4 | 4 | 1.12 | 1.04 |
| Q86U42 | PABPN1 | 5 | 5 | 1.12 | 0.99 |
| O75569 | PRKRA | 8 | 8 | 1.12 | 1.11 |
| P62805 | H4-16 | 9 | 9 | 1.12 | 0.81 |
| P62857 | RPS28 | 4 | 4 | 1.12 | 1.20 |
| Q16630 | CPSF6 | 12 | 12 | 1.12 | 0.84 |
| Q02539 | H1-1 | 10 | 4 | 1.12 | 0.87 |
| P55795 | HNRNPH2 | 14 | 8 | 1.12 | 0.67 |
| Q14919 | DRAP1 | 5 | 5 | 1.12 | 0.98 |
| P13674 | P4HA1 | 33 | 33 | 1.12 | 1.56 |
| Q15650 | TRIP4 | 3 | 3 | 1.13 | 0.01 |
| Q9Y5A7 | NUB1 | 5 | 5 | 1.13 | 0.85 |
| P13797 | PLS3 | 29 | 26 | 1.13 | 1.47 |
| P23193 | TCEA1 | 5 | 5 | 1.13 | 0.97 |
| Q00577 | PURA | 9 | 8 | 1.13 | 1.00 |
| Q8N1F7 | NUP93 | 21 | 21 | 1.13 | 1.09 |
| Q8WVJ2 | NUDCD2 | 2 | 2 | 1.13 | 0.01 |
| Q9UQE7 | SMC3 | 27 | 27 | 1.13 | 0.78 |
| Q9NQR4 | NIT2 | 9 | 9 | 1.13 | 1.05 |

| Accession | Name | # Peptides | # Unique Peptides | Abundance Ratio: day 3 / day 0 | Abundance Ratio: day 7/ day 0 |
|---|---|---|---|---|---|
| Q92845 | KIFAP3 | 2 | 1 | 0.01 | 0.97 |
| Q96BM9 | ARL8A | 2 | 1 | 0.01 | 0.76 |
| P78347 | GTF2I | 15 | 15 | 1.13 | 1.17 |
| P25325 | MPST | 3 | 3 | 1.13 | 1.08 |
| O95140 | MFN2 | 3 | 3 | 1.13 | 0.68 |
| Q9BZG1 | RAB34 | 8 | 8 | 1.13 | 0.95 |
| Q9NVP1 | DDX18 | 9 | 9 | 1.13 | 1.11 |
| Q8NBJ4 | GOLM1 | 12 | 12 | 1.13 | 1.34 |
| O15173 | PGRMC2 | 7 | 7 | 1.13 | 0.89 |
| P15144 | ANPEP | 36 | 36 | 1.13 | 0.71 |
| P16083 | NQO2 | 7 | 7 | 1.13 | 0.99 |
| Q13564 | NAE1 | 9 | 9 | 1.13 | 0.85 |
| Q9NQW7 | XPNPEP1 | 12 | 11 | 1.13 | 1.22 |
| A1X283 | SH3PXD2B | 10 | 9 | 1.13 | 1.15 |
| Q99717 | SMAD5 | 3 | 2 | 1.14 | 1.20 |
| Q99590 | SCAF11 | 3 | 3 | 1.14 | 1.13 |
| Q969X5 | ERGIC1 | 8 | 8 | 1.14 | 1.05 |
| P00568 | AK1 | 19 | 19 | 1.14 | 1.29 |
| Q6PJT7 | ZC3H14 | 6 | 6 | 1.14 | 0.01 |
| Q14376 | GALE | 6 | 6 | 1.14 | 1.03 |
| O95817 | BAG3 | 18 | 18 | 1.14 | 1.23 |
| Q9H0D6 | XRN2 | 5 | 5 | 1.14 | 0.87 |
| Q9ULH1 | ASAP1 | 6 | 6 | 1.14 | 0.01 |
| O75150 | RNF40 | 3 | 2 | 1.14 | 0.01 |
| P36405 | ARL3 | 4 | 4 | 1.14 | 1.22 |
| O75368 | SH3BGRL | 10 | 10 | 1.14 | 1.12 |
| P09429 | HMGB1 | 17 | 14 | 1.14 | 0.89 |
| P29590 | PML | 29 | 29 | 1.14 | 1.06 |
| Q13618 | CUL3 | 12 | 12 | 1.14 | 1.25 |
| P10253 | GAA | 18 | 18 | 1.14 | 1.17 |
| P54840 | GYS2 | 2 | 1 | 1.14 | 0.98 |
| O14684 | PTGES | 5 | 5 | 1.14 | 1.10 |
| P30085 | CMPK1 | 7 | 7 | 1.14 | 1.31 |
| O00625 | PIR | 2 | 2 | 1.14 | 1.39 |
| Q01804 | OTUD4 | 2 | 2 | 1.15 | 0.82 |
| Q9NX62 | BPNT2 | 5 | 5 | 1.15 | 0.91 |
| Q9NY12 | GAR1 | 3 | 3 | 1.15 | 1.05 |
| Q96DG6 | CMBL | 10 | 10 | 1.15 | 1.07 |
| Q96HD1 | CRELD1 | 5 | 5 | 1.15 | 1.14 |
| Q14766 | LTBP1 | 12 | 12 | 1.15 | 1.89 |
| P16401 | H1-5 | 21 | 17 | 1.15 | 0.73 |
| Q9H0W5 | CCDC8 | 6 | 6 | 1.15 | 1.61 |
| P62993 | GRB2 | 6 | 6 | 1.15 | 1.04 |

| Accession | Name | # Peptides | # Unique Peptides | Abundance Ratio: day 3 / day 0 | Abundance Ratio: day 7/ day 0 |
|---|---|---|---|---|---|
| Q92845 | KIFAP3 | 2 | 1 | 0.01 | 0.97 |
| Q96BM9 | ARL8A | 2 | 1 | 0.01 | 0.76 |
| P37108 | SRP14 | 7 | 7 | 1.15 | 1.09 |
| P42330 | AKR1C3 | 17 | 9 | 1.15 | 0.75 |
| Q3SY69 | ALDH1L2 | 17 | 17 | 1.15 | 1.49 |
| Q14195 | DPYSL3 | 24 | 21 | 1.15 | 1.12 |
| P83916 | CBX1 | 2 | 2 | 1.15 | 1.17 |
| Q9UBW8 | COPS7A | 8 | 8 | 1.15 | 0.93 |
| Q16777 | H2AC20 | 9 | 2 | 1.15 | 0.70 |
| Q709C8 | VPS13C | 9 | 9 | 1.15 | 1.15 |
| Q70E73 | RAPH1 | 16 | 16 | 1.15 | 1.53 |
| P63313 | TMSB10 | 4 | 2 | 1.16 | 1.68 |
| Q6EMK4 | VASN | 7 | 7 | 1.16 | 1.34 |
| Q8TBC4 | UBA3 | 7 | 7 | 1.16 | 0.89 |
| O95218 | ZRANB2 | 9 | 9 | 1.16 | 1.14 |
| P13807 | GYS1 | 15 | 14 | 1.16 | 1.03 |
| P09038 | FGF2 | 3 | 3 | 1.16 | 0.01 |
| P11802 | CDK4 | 5 | 5 | 1.16 | 0.97 |
| O00442 | RTCA | 5 | 5 | 1.16 | 0.94 |
| P25686 | DNAJB2 | 4 | 4 | 1.16 | 1.18 |
| P06396 | GSN | 39 | 39 | 1.16 | 1.22 |
| Q92882 | OSTF1 | 7 | 7 | 1.16 | 1.27 |
| Q9UHL4 | DPP7 | 9 | 9 | 1.16 | 1.02 |
| P36871 | PGM1 | 31 | 31 | 1.16 | 1.34 |
| Q9H7N4 | SCAF1 | 3 | 3 | 1.16 | 0.01 |
| Q4KMQ2 | ANO6 | 4 | 4 | 1.16 | 1.19 |
| Q16585 | SGCB | 6 | 6 | 1.16 | 0.89 |
| Q9P016 | THYN1 | 8 | 8 | 1.16 | 0.89 |
| Q8NC56 | LEMD2 | 11 | 11 | 1.16 | 1.14 |
| Q9Y4W2 | LAS1L | 2 | 2 | 1.16 | 0.01 |
| Q9Y3Y2 | CHTOP | 4 | 4 | 1.16 | 0.85 |
| P49458 | SRP9 | 4 | 4 | 1.16 | 1.13 |
| P78527 | PRKDC | 108 | 108 | 1.16 | 0.96 |
| Q14839 | CHD4 | 12 | 8 | 1.17 | 1.11 |
| Q92733 | PRCC | 3 | 3 | 1.17 | 1.56 |
| O75592 | MYCBP2 | 6 | 6 | 1.17 | 0.01 |
| Q9Y2P8 | RCL1 | 2 | 2 | 1.17 | 1.26 |
| Q8N8U9 | BMPER | 3 | 3 | 1.17 | 1.29 |
| Q9UNZ5 | C19orf53 | 3 | 3 | 1.17 | 1.91 |
| Q9NNW7 | TXNRD2 | 6 | 5 | 1.17 | 0.01 |
| Q00341 | HDLBP | 60 | 60 | 1.17 | 1.70 |
| Q01433 | AMPD2 | 13 | 13 | 1.17 | 1.25 |
| Q08209 | PPP3CA | 13 | 7 | 1.17 | 1.19 |

| Accession | Name | # Peptides | # Unique Peptides | Abundance Ratio: day 3 / day 0 | Abundance Ratio: day 7/ day 0 |
|---|---|---|---|---|---|
| Q92845 | KIFAP3 | 2 | 1 | 0.01 | 0.97 |
| Q96BM9 | ARL8A | 2 | 1 | 0.01 | 0.76 |
| Q3KQV9 | UAP1L1 | 9 | 7 | 1.17 | 1.09 |
| P39687 | ANP32A | 9 | 7 | 1.17 | 1.08 |
| P55081 | MFAP1 | 3 | 2 | 1.17 | 0.01 |
| Q8N335 | GPD1L | 4 | 4 | 1.17 | 1.51 |
| P21397 | MAOA | 9 | 7 | 1.17 | 0.99 |
| Q8WUY3 | PRUNE2 | 15 | 15 | 1.17 | 0.97 |
| Q14938 | NFIX | 6 | 4 | 1.17 | 0.97 |
| P16278 | GLB1 | 8 | 8 | 1.17 | 1.17 |
| Q86VM9 | ZC3H18 | 7 | 7 | 1.17 | 1.00 |
| Q5VW32 | BROX | 6 | 6 | 1.18 | 0.83 |
| Q49A26 | GLYR1 | 3 | 3 | 1.18 | 1.31 |
| Q8N9T8 | KRI1 | 4 | 4 | 1.18 | 0.01 |
| Q5SSJ5 | HP1BP3 | 24 | 24 | 1.18 | 0.96 |
| O60885 | BRD4 | 3 | 3 | 1.18 | 0.89 |
| Q15758 | SLC1A5 | 5 | 5 | 1.18 | 1.41 |
| Q9NUJ3 | TCP11L1 | 2 | 2 | 1.18 | 0.01 |
| O75175 | CNOT3 | 3 | 3 | 1.18 | 1.63 |
| Q9BSD7 | NTPCR | 4 | 4 | 1.18 | 0.87 |
| Q8TCJ2 | STT3B | 13 | 11 | 1.18 | 1.01 |
| P15121 | AKR1B1 | 14 | 14 | 1.18 | 1.00 |
| Q9UH99 | SUN2 | 14 | 14 | 1.18 | 0.95 |
| P17480 | UBTF | 9 | 9 | 1.18 | 1.11 |
| Q9UKX5 | ITGA11 | 10 | 10 | 1.18 | 1.18 |
| P63098 | PPP3R1 | 2 | 2 | 1.19 | 1.34 |
| Q9Y3A6 | TMED5 | 3 | 3 | 1.19 | 0.85 |
| Q9P2N5 | RBM27 | 4 | 3 | 1.19 | 0.01 |
| Q8TDY4 | ASAP3 | 2 | 2 | 1.19 | 1.00 |
| P49903 | SEPHS1 | 3 | 3 | 1.19 | 1.02 |
| P46527 | CDKN1B | 2 | 2 | 1.19 | 0.01 |
| P40763 | STAT3 | 16 | 16 | 1.19 | 1.40 |
| Q9UBU9 | NXF1 | 5 | 5 | 1.19 | 0.78 |
| P08243 | ASNS | 9 | 9 | 1.19 | 1.48 |
| P48163 | ME1 | 10 | 10 | 1.19 | 1.56 |
| Q96ST2 | IWS1 | 3 | 3 | 1.20 | 1.26 |
| Q7Z7G0 | ABI3BP | 16 | 16 | 1.20 | 2.47 |
| P51153 | RAB13 | 9 | 7 | 1.20 | 1.27 |
| Q9H4A6 | GOLPH3 | 2 | 2 | 1.20 | 1.02 |
| Q66K74 | MAP1S | 5 | 5 | 1.20 | 1.15 |
| P30086 | PEBP1 | 12 | 12 | 1.20 | 1.17 |
| Q9H6S3 | EPS8L2 | 10 | 10 | 1.20 | 1.90 |
| Q92466 | DDB2 | 3 | 3 | 1.20 | 0.96 |

| Accession | Name | # Peptides | # Unique Peptides | Abundance Ratio: day 3 / day 0 | Abundance Ratio: day 7/ day 0 |
|---|---|---|---|---|---|
| Q92845 | KIFAP3 | 2 | 1 | 0.01 | 0.97 |
| Q96BM9 | ARL8A | 2 | 1 | 0.01 | 0.76 |
| P00558 | PGK1 | 32 | 28 | 1.20 | 1.11 |
| Q9NR99 | MXRA5 | 38 | 38 | 1.20 | 2.89 |
| Q8WTT2 | NOC3L | 2 | 2 | 1.20 | 0.01 |
| Q9BXF6 | RAB11FIP5 | 9 | 9 | 1.20 | 0.88 |
| P42696 | RBM34 | 3 | 3 | 1.21 | 1.12 |
| Q5T8P6 | RBM26 | 4 | 3 | 1.21 | 0.01 |
| Q15121 | PEA15 | 5 | 5 | 1.21 | 0.91 |
| P27105 | STOM | 9 | 9 | 1.21 | 0.89 |
| P09972 | ALDOC | 24 | 18 | 1.21 | 1.17 |
| Q8NB37 | GATD1 | 3 | 3 | 1.21 | 0.80 |
| O43281 | EFS | 4 | 4 | 1.21 | 0.01 |
| P21399 | ACO1 | 25 | 25 | 1.21 | 1.48 |
| Q9Y6D6 | ARFGEF1 | 5 | 3 | 1.21 | 1.04 |
| Q8N6T3 | ARFGAP1 | 7 | 7 | 1.21 | 1.74 |
| P32119 | PRDX2 | 9 | 8 | 1.21 | 1.08 |
| Q9Y613 | FHOD1 | 5 | 5 | 1.21 | 1.24 |
| P52895 | AKR1C2 | 14 | 6 | 1.21 | 0.70 |
| Q9H4A3 | WNK1 | 15 | 15 | 1.21 | 1.41 |
| Q15813 | TBCE | 7 | 7 | 1.21 | 0.82 |
| O94808 | GFPT2 | 17 | 10 | 1.21 | 1.80 |
| P16402 | H1-3 | 18 | 7 | 1.21 | 0.79 |
| P51570 | GALK1 | 13 | 13 | 1.22 | 1.21 |
| Q16678 | CYP1B1 | 9 | 9 | 1.22 | 1.56 |
| Q9ULX6 | AKAP8L | 4 | 4 | 1.22 | 1.26 |
| P52948 | NUP98 | 9 | 9 | 1.22 | 1.03 |
| P05386 | RPLP1 | 2 | 1 | 1.22 | 1.18 |
| Q96BN8 | OTULIN | 2 | 2 | 1.22 | 0.01 |
| Q9H857 | NT5DC2 | 5 | 5 | 1.22 | 1.56 |
| P16104 | H2AX | 10 | 5 | 1.22 | 1.29 |
| Q14684 | RRP1B | 3 | 3 | 1.22 | 0.01 |
| Q96B97 | SH3KBP1 | 10 | 10 | 1.23 | 1.31 |
| Q92575 | UBXN4 | 7 | 7 | 1.23 | 1.10 |
| Q96RY7 | IFT140 | 2 | 2 | 1.23 | 0.01 |
| Q96F85 | CNRIP1 | 4 | 4 | 1.23 | 1.20 |
| Q9HD26 | GOPC | 12 | 12 | 1.23 | 1.55 |
| Q13123 | IK | 3 | 3 | 1.23 | 0.76 |
| Q9H0X9 | OSBPL5 | 7 | 6 | 1.23 | 0.95 |
| Q9Y4G6 | TLN2 | 46 | 26 | 1.23 | 1.16 |
| Q14692 | BMS1 | 4 | 4 | 1.23 | 1.02 |
| P68402 | PAFAH1B2 | 5 | 5 | 1.23 | 1.00 |
| P57772 | EEFSEC | 4 | 4 | 1.24 | 1.25 |

| Accession | Name | # Peptides | # Unique Peptides | Abundance Ratio: day 3 / day 0 | Abundance Ratio: day 7/ day 0 |
|---|---|---|---|---|---|
| Q92845 | KIFAP3 | 2 | 1 | 0.01 | 0.97 |
| Q96BM9 | ARL8A | 2 | 1 | 0.01 | 0.76 |
| Q99584 | S100A13 | 5 | 5 | 1.24 | 1.22 |
| Q13151 | HNRNPA0 | 11 | 11 | 1.24 | 1.36 |
| P23470 | PTPRG | 3 | 3 | 1.24 | 0.01 |
| Q16851 | UGP2 | 20 | 20 | 1.24 | 1.35 |
| Q9Y3E1 | HDGFL3 | 5 | 4 | 1.24 | 1.22 |
| P00338 | LDHA | 22 | 19 | 1.24 | 1.22 |
| P16403 | H1-2 | 22 | 10 | 1.24 | 0.94 |
| Q9P0V9 | SEPTIN10 | 9 | 7 | 1.25 | 0.95 |
| Q96NY7 | CLIC6 | 12 | 12 | 1.25 | 0.01 |
| Q08378 | GOLGA3 | 27 | 27 | 1.25 | 1.38 |
| O95965 | ITGBL1 | 4 | 4 | 1.25 | 0.01 |
| P35613 | BSG | 10 | 10 | 1.25 | 1.17 |
| Q9HBL0 | TNS1 | 42 | 39 | 1.25 | 1.21 |
| Q13541 | EIF4EBP1 | 3 | 3 | 1.25 | 1.24 |
| P35222 | CTNNB1 | 13 | 13 | 1.25 | 1.24 |
| P10412 | H1-4 | 22 | 9 | 1.25 | 0.84 |
| P50454 | SERPINH1 | 25 | 25 | 1.25 | 2.44 |
| O75694 | NUP155 | 10 | 10 | 1.25 | 1.04 |
| P58005 | SESN3 | 3 | 3 | 1.25 | 1.21 |
| P25940 | COL5A3 | 8 | 8 | 1.25 | 1.49 |
| O00418 | EEF2K | 4 | 4 | 1.25 | 1.50 |
| Q9NZ32 | ACTR10 | 2 | 2 | 1.26 | 0.01 |
| Q92629 | SGCD | 7 | 7 | 1.26 | 0.89 |
| Q9BWS9 | CHID1 | 7 | 7 | 1.26 | 1.07 |
| Q8TF66 | LRRC15 | 7 | 7 | 1.26 | 4.89 |
| O75367 | MACROH2A1 | 11 | 10 | 1.26 | 1.07 |
| Q9NXG2 | THUMPD1 | 8 | 8 | 1.26 | 0.94 |
| P19971 | TYMP | 10 | 10 | 1.26 | 1.33 |
| P32321 | DCTD | 4 | 4 | 1.26 | 0.01 |
| Q96SL4 | GPX7 | 4 | 4 | 1.26 | 2.14 |
| Q6NUQ4 | TMEM214 | 16 | 16 | 1.26 | 1.52 |
| Q52LD8 | RFTN2 | 3 | 3 | 1.27 | 1.20 |
| P35908 | KRT2 | 4 | 2 | 1.27 | 1.17 |
| Q9UI14 | RABAC1 | 2 | 2 | 1.27 | 1.96 |
| P07585 | DCN | 14 | 13 | 1.27 | 1.77 |
| P11217 | PYGM | 12 | 1 | 1.27 | 0.95 |
| O94921 | CDK14 | 2 | 2 | 1.27 | 0.01 |
| Q96SB3 | PPP1R9B | 8 | 8 | 1.27 | 1.52 |
| O60216 | RAD21 | 4 | 4 | 1.28 | 1.01 |
| P25685 | DNAJB1 | 9 | 9 | 1.28 | 1.29 |
| P60174 | TPI1 | 17 | 17 | 1.28 | 1.15 |

| Accession | Name | # Peptides | # Unique Peptides | Abundance Ratio: day 3 / day 0 | Abundance Ratio: day 7/ day 0 |
|---|---|---|---|---|---|
| Q92845 | KIFAP3 | 2 | 1 | 0.01 | 0.97 |
| Q96BM9 | ARL8A | 2 | 1 | 0.01 | 0.76 |
| O14936 | CASK | 7 | 7 | 1.28 | 1.25 |
| Q96I15 | SCLY | 2 | 2 | 1.28 | 1.08 |
| Q9UBY9 | HSPB7 | 2 | 2 | 1.28 | 0.66 |
| Q13162 | PRDX4 | 12 | 9 | 1.28 | 1.85 |
| Q9NTX5 | ECHDC1 | 7 | 7 | 1.28 | 1.13 |
| P07477 | PRSS1 | 2 | 2 | 1.29 | 0.98 |
| Q9Y6E2 | BZW2 | 6 | 6 | 1.29 | 1.29 |
| Q14767 | LTBP2 | 23 | 23 | 1.29 | 2.99 |
| Q9ULT8 | HECTD1 | 7 | 7 | 1.29 | 1.27 |
| O95070 | YIF1A | 2 | 2 | 1.29 | 1.43 |
| Q9GZR7 | DDX24 | 11 | 11 | 1.29 | 1.17 |
| Q9NVE7 | PANK4 | 3 | 3 | 1.29 | 1.18 |
| Q9UNN8 | PROCR | 3 | 3 | 1.30 | 0.91 |
| P46439 | GSTM5 | 6 | 2 | 1.30 | 0.01 |
| Q9NX05 | FAM120C | 2 | 2 | 1.30 | 0.01 |
| Q9BRZ2 | TRIM56 | 5 | 5 | 1.30 | 0.73 |
| P51884 | LUM | 9 | 9 | 1.31 | 2.44 |
| P14543 | NID1 | 4 | 3 | 1.31 | 1.47 |
| Q9Y6U3 | SCIN | 16 | 16 | 1.31 | 1.91 |
| P49591 | SARS1 | 19 | 19 | 1.31 | 1.54 |
| P35573 | AGL | 11 | 11 | 1.31 | 1.25 |
| Q9NRX3 | NDUFA4L2 | 6 | 6 | 1.32 | 1.13 |
| Q53F19 | NCBP3 | 2 | 2 | 1.32 | 1.24 |
| P19387 | POLR2C | 4 | 4 | 1.32 | 1.05 |
| Q9UEE9 | CFDP1 | 3 | 3 | 1.32 | 0.01 |
| Q8IY33 | MICALL2 | 3 | 3 | 1.32 | 1.43 |
| Q08AF3 | SLFN5 | 3 | 3 | 1.32 | 0.99 |
| Q96PU8 | QKI | 4 | 4 | 1.32 | 1.04 |
| Q96ME7 | ZNF512 | 3 | 3 | 1.33 | 1.03 |
| Q13643 | FHL3 | 4 | 4 | 1.33 | 1.90 |
| Q08431 | MFGE8 | 11 | 11 | 1.33 | 1.30 |
| O14773 | TPP1 | 4 | 4 | 1.33 | 0.86 |
| Q8WTX7 | CASTOR1 | 2 | 2 | 1.33 | 1.17 |
| P07602 | PSAP | 5 | 5 | 1.33 | 1.57 |
| P08294 | SOD3 | 6 | 6 | 1.34 | 0.84 |
| Q9BUT1 | BDH2 | 9 | 9 | 1.34 | 1.23 |
| P54652 | HSPA2 | 27 | 14 | 1.34 | 1.06 |
| Q5JWF2 | GNAS | 14 | 14 | 1.34 | 1.22 |
| P51178 | PLCD1 | 15 | 15 | 1.34 | 1.58 |
| P22105 | TNXB | 71 | 71 | 1.34 | 0.82 |
| Q01196 | RUNX1 | 2 | 2 | 1.35 | 1.73 |

| Accession | Name | # Peptides | # Unique Peptides | Abundance Ratio: day 3 / day 0 | Abundance Ratio: day 7/ day 0 |
|---|---|---|---|---|---|
| Q92845 | KIFAP3 | 2 | 1 | 0.01 | 0.97 |
| Q96BM9 | ARL8A | 2 | 1 | 0.01 | 0.76 |
| Q32P28 | P3H1 | 20 | 20 | 1.35 | 2.40 |
| O15027 | SEC16A | 13 | 13 | 1.35 | 1.44 |
| O14908 | GIPC1 | 4 | 4 | 1.35 | 0.01 |
| P51151 | RAB9A | 5 | 5 | 1.35 | 1.08 |
| O00291 | HIP1 | 12 | 12 | 1.36 | 1.48 |
| Q9Y2B0 | CNPY2 | 9 | 9 | 1.36 | 1.64 |
| Q96KR1 | ZFR | 9 | 9 | 1.36 | 1.15 |
| P09871 | C1S | 2 | 2 | 1.36 | 0.01 |
| P07339 | CTSD | 17 | 17 | 1.36 | 1.50 |
| O60341 | KDM1A | 4 | 4 | 1.37 | 1.20 |
| Q32P44 | EML3 | 11 | 11 | 1.37 | 1.30 |
| P07951 | TPM2 | 27 | 6 | 1.37 | 2.09 |
| P14174 | MIF | 5 | 5 | 1.37 | 1.47 |
| P55290 | CDH13 | 7 | 7 | 1.37 | 0.94 |
| P48426 | PIP4K2A | 7 | 5 | 1.37 | 1.41 |
| Q06203 | PPAT | 4 | 4 | 1.38 | 1.28 |
| O43556 | SGCE | 3 | 3 | 1.38 | 1.00 |
| Q16539 | MAPK14 | 5 | 5 | 1.38 | 1.02 |
| P05026 | ATP1B1 | 11 | 11 | 1.38 | 1.46 |
| P36269 | GGT5 | 6 | 6 | 1.38 | 1.01 |
| P04792 | HSPB1 | 16 | 16 | 1.38 | 1.73 |
| Q14554 | PDIA5 | 16 | 16 | 1.39 | 1.50 |
| Q16363 | LAMA4 | 23 | 23 | 1.39 | 1.41 |
| O14498 | ISLR | 6 | 6 | 1.40 | 1.65 |
| Q6NW29 | RWDD4 | 3 | 3 | 1.40 | 1.19 |
| P63208 | SKP1 | 3 | 3 | 1.40 | 1.86 |
| Q8TAQ2 | SMARCC2 | 15 | 15 | 1.40 | 1.11 |
| Q8WWX9 | SELENOM | 3 | 3 | 1.41 | 1.84 |
| P07332 | FES | 2 | 2 | 1.41 | 0.01 |
| O00308 | WWP2 | 32 | 29 | 1.41 | 0.75 |
| Q8NFW8 | CMAS | 8 | 8 | 1.41 | 0.94 |
| Q9NTI5 | PDS5B | 11 | 11 | 1.41 | 1.20 |
| P11047 | LAMC1 | 34 | 34 | 1.42 | 1.53 |
| O15484 | CAPN5 | 4 | 4 | 1.42 | 0.01 |
| Q96CG8 | CTHRC1 | 5 | 5 | 1.42 | 1.58 |
| Q8NFH5 | NUP35 | 3 | 3 | 1.42 | 0.01 |
| Q96D15 | RCN3 | 12 | 12 | 1.43 | 1.88 |
| Q96BW5 | PTER | 2 | 2 | 1.43 | 1.40 |
| P00450 | CP | 11 | 11 | 1.43 | 1.05 |
| Q9UKZ9 | PCOLCE2 | 2 | 2 | 1.43 | 1.78 |
| Q9BQA1 | WDR77 | 3 | 3 | 1.43 | 1.38 |

| Accession | Name | # Peptides | # Unique Peptides | Abundance Ratio: day 3 / day 0 | Abundance Ratio: day 7/ day 0 |
|---|---|---|---|---|---|
| Q92845 | KIFAP3 | 2 | 1 | 0.01 | 0.97 |
| Q96BM9 | ARL8A | 2 | 1 | 0.01 | 0.76 |
| P48634 | PRRC2A | 7 | 5 | 1.44 | 2.04 |
| P34741 | SDC2 | 4 | 4 | 1.44 | 1.43 |
| O95400 | CD2BP2 | 2 | 2 | 1.44 | 0.01 |
| Q6UVK1 | CSPG4 | 60 | 60 | 1.45 | 1.66 |
| Q15007 | WTAP | 3 | 3 | 1.45 | 1.47 |
| P23219 | PTGS1 | 3 | 3 | 1.46 | 1.13 |
| O43175 | PHGDH | 21 | 21 | 1.46 | 2.39 |
| Q9BUH6 | PAXX | 2 | 2 | 1.46 | 0.01 |
| Q9H4Z3 | PCIF1 | 2 | 2 | 1.46 | 1.19 |
| P34949 | MPI | 7 | 7 | 1.46 | 1.18 |
| Q9Y617 | PSAT1 | 15 | 15 | 1.46 | 2.09 |
| Q9H4G4 | GLIPR2 | 2 | 2 | 1.47 | 2.54 |
| Q9UHF7 | TRPS1 | 4 | 4 | 1.47 | 2.19 |
| Q96QR8 | PURB | 5 | 4 | 1.48 | 1.34 |
| Q14738 | PPP2R5D | 10 | 10 | 1.48 | 1.32 |
| O00499 | BIN1 | 9 | 7 | 1.49 | 0.01 |
| O95670 | ATP6V1G2 | 3 | 1 | 1.49 | 0.01 |
| P13716 | ALAD | 5 | 5 | 1.49 | 1.30 |
| O15460 | P4HA2 | 19 | 19 | 1.50 | 2.54 |
| Q9P2B2 | PTGFRN | 5 | 5 | 1.50 | 1.83 |
| P38936 | CDKN1A | 2 | 2 | 1.51 | 0.01 |
| P22413 | ENPP1 | 12 | 12 | 1.51 | 1.65 |
| Q08AM6 | VAC14 | 5 | 5 | 1.51 | 0.93 |
| Q9Y680 | FKBP7 | 13 | 13 | 1.51 | 1.99 |
| O60256 | PRPSAP2 | 4 | 4 | 1.52 | 1.49 |
| P43490 | NAMPT | 17 | 17 | 1.53 | 1.24 |
| P49746 | THBS3 | 9 | 9 | 1.53 | 2.45 |
| Q05193 | DNM1 | 18 | 11 | 1.53 | 2.71 |
| Q96LD4 | TRIM47 | 9 | 9 | 1.53 | 1.36 |
| Q8NEZ5 | FBXO22 | 2 | 2 | 1.54 | 1.79 |
| P53985 | SLC16A1 | 9 | 9 | 1.55 | 1.81 |
| Q12884 | FAP | 22 | 22 | 1.55 | 1.99 |
| Q8IVF7 | FMNL3 | 5 | 2 | 1.56 | 1.64 |
| P07902 | GALT | 4 | 4 | 1.56 | 1.02 |
| Q96C23 | GALM | 8 | 8 | 1.57 | 1.67 |
| P82909 | MRPS36 | 4 | 4 | 1.57 | 1.06 |
| Q9NZJ9 | NUDT4 | 5 | 3 | 1.57 | 1.33 |
| P36969 | GPX4 | 3 | 3 | 1.58 | 1.53 |
| Q9UBG0 | MRC2 | 35 | 35 | 1.58 | 1.78 |
| Q76M96 | CCDC80 | 13 | 13 | 1.59 | 3.04 |
| Q6YHK3 | CD109 | 29 | 29 | 1.59 | 1.25 |

| Accession | Name | # Peptides | # Unique Peptides | Abundance Ratio: day 3 / day 0 | Abundance Ratio: day 7/ day 0 |
|---|---|---|---|---|---|
| Q92845 | KIFAP3 | 2 | 1 | 0.01 | 0.97 |
| Q96BM9 | ARL8A | 2 | 1 | 0.01 | 0.76 |
| Q8N2G8 | GHDC | 5 | 5 | 1.59 | 1.03 |
| P07108 | DBI | 6 | 6 | 1.59 | 2.06 |
| P40261 | NNMT | 8 | 8 | 1.59 | 2.16 |
| P55287 | CDH11 | 8 | 8 | 1.59 | 1.66 |
| Q08397 | LOXL1 | 10 | 10 | 1.59 | 2.76 |
| O95340 | PAPSS2 | 29 | 25 | 1.61 | 2.87 |
| Q8IV08 | PLD3 | 10 | 10 | 1.62 | 1.48 |
| Q92522 | H1-10 | 15 | 15 | 1.63 | 0.99 |
| P08651 | NFIC | 5 | 4 | 1.63 | 1.22 |
| Q9UPA5 | BSN | 2 | 2 | 1.63 | 0.01 |
| P10909 | CLU | 13 | 13 | 1.64 | 1.69 |
| P13645 | KRT10 | 7 | 6 | 1.64 | 2.24 |
| P06737 | PYGL | 38 | 31 | 1.65 | 1.77 |
| P23142 | FBLN1 | 20 | 20 | 1.66 | 1.75 |
| O00479 | HMGN4 | 5 | 4 | 1.67 | 1.70 |
| Q92954 | PRG4 | 57 | 57 | 1.67 | 3.71 |
| P28799 | GRN | 6 | 6 | 1.70 | 1.32 |
| Q9BQB6 | VKORC1 | 4 | 4 | 1.70 | 2.49 |
| O76054 | SEC14L2 | 4 | 4 | 1.70 | 1.57 |
| P08572 | COL4A2 | 13 | 13 | 1.77 | 2.69 |
| Q99808 | SLC29A1 | 6 | 6 | 1.77 | 2.68 |
| P00736 | C1R | 6 | 6 | 1.78 | 2.09 |
| Q99541 | PLIN2 | 19 | 19 | 1.80 | 1.42 |
| P07942 | LAMB1 | 40 | 40 | 1.80 | 1.79 |
| P14209 | CD99 | 3 | 3 | 1.81 | 1.67 |
| Q6PCB0 | VWA1 | 4 | 4 | 1.83 | 2.00 |
| P02768 | ALB | 4 | 4 | 1.84 | 1.70 |
| P08582 | MELTF | 25 | 25 | 1.89 | 1.88 |
| P14555 | PLA2G2A | 4 | 4 | 1.90 | 5.79 |
| P04179 | SOD2 | 11 | 11 | 1.90 | 1.94 |
| P02795 | MT2A | 4 | 1 | 1.90 | 3.49 |
| Q68CZ2 | TNS3 | 12 | 9 | 1.92 | 2.56 |
| P49006 | MARCKSL1 | 5 | 5 | 1.92 | 1.84 |
| P98160 | HSPG2 | 133 | 133 | 1.92 | 2.27 |
| P09486 | SPARC | 12 | 12 | 1.93 | 4.06 |
| P08493 | MGP | 3 | 3 | 1.97 | 2.59 |
| P07305 | H1-0 | 11 | 11 | 1.97 | 1.90 |
| P98095 | FBLN2 | 13 | 13 | 2.00 | 2.32 |
| Q6UY14 | ADAMTSL4 | 6 | 6 | 2.03 | 1.62 |
| O14494 | PLPP1 | 2 | 2 | 2.03 | 1.78 |
| Q05707 | COL14A1 | 47 | 47 | 2.05 | 2.78 |

| Accession | Name | # Peptides | # Unique Peptides | Abundance Ratio: day 3 / day 0 | Abundance Ratio: day 7/ day 0 |
|---|---|---|---|---|---|
| Q92845 | KIFAP3 | 2 | 1 | 0.01 | 0.97 |
| Q96BM9 | ARL8A | 2 | 1 | 0.01 | 0.76 |
| Q9NYL4 | FKBP11 | 2 | 2 | 2.09 | 2.69 |
| O94875 | SORBS2 | 12 | 12 | 2.11 | 1.94 |
| P02751 | FN1 | 98 | 98 | 2.12 | 3.51 |
| Q8TAE6 | PPP1R14C | 3 | 3 | 2.21 | 2.52 |
| P21741 | MDK | 3 | 3 | 2.22 | 2.18 |
| P12109 | COL6A1 | 47 | 47 | 2.22 | 2.73 |
| O95967 | EFEMP2 | 9 | 9 | 2.27 | 2.88 |
| Q8TER0 | SNED1 | 5 | 5 | 2.29 | 1.83 |
| Q658P3 | STEAP3 | 7 | 7 | 2.36 | 3.38 |
| P12110 | COL6A2 | 45 | 45 | 2.36 | 2.99 |
| Q8N142 | ADSS1 | 8 | 7 | 2.37 | 2.18 |
| P02794 | FTH1 | 12 | 12 | 2.44 | 2.15 |
| P02511 | CRYAB | 14 | 14 | 2.51 | 2.92 |
| P54687 | BCAT1 | 7 | 7 | 2.53 | 2.89 |
| P12111 | COL6A3 | 165 | 165 | 2.53 | 3.10 |
| Q15113 | PCOLCE | 16 | 16 | 2.55 | 3.69 |
| Q6UX71 | PLXDC2 | 7 | 7 | 2.61 | 4.16 |
| P15104 | GLUL | 8 | 8 | 2.62 | 1.27 |
| P01011 | SERPINA3 | 6 | 6 | 2.64 | 2.73 |
| P02458 | COL2A1 | 67 | 61 | 2.68 | 32.94 |
| Q92743 | HTRA1 | 10 | 10 | 2.69 | 3.53 |
| Q07092 | COL16A1 | 17 | 17 | 2.71 | 5.78 |
| Q13451 | FKBP5 | 21 | 20 | 2.77 | 3.02 |
| P35555 | FBN1 | 101 | 95 | 2.79 | 4.38 |
| P13611 | VCAN | 27 | 27 | 2.80 | 7.65 |
| P10915 | HAPLN1 | 15 | 15 | 2.86 | 4.39 |
| Q14112 | NID2 | 33 | 32 | 2.94 | 3.32 |
| P35556 | FBN2 | 17 | 11 | 3.00 | 3.34 |
| Q9Y6C2 | EMILIN1 | 38 | 38 | 3.09 | 3.88 |
| P12107 | COL11A1 | 35 | 28 | 3.11 | 10.84 |
| P07996 | THBS1 | 34 | 33 | 3.19 | 6.06 |
| P20908 | COL5A1 | 37 | 30 | 3.23 | 6.27 |
| Q15582 | TGFBI | 33 | 33 | 3.30 | 4.17 |
| P02462 | COL4A1 | 9 | 9 | 3.42 | 4.29 |
| P02792 | FTL | 10 | 10 | 3.43 | 3.31 |
| P24821 | TNC | 106 | 106 | 3.45 | 5.86 |
| P05997 | COL5A2 | 50 | 48 | 3.61 | 7.87 |
| P21810 | BGN | 12 | 11 | 3.73 | 9.34 |
| Q8IUX7 | AEBP1 | 38 | 37 | 3.90 | 8.02 |
| P51888 | PRELP | 12 | 12 | 4.07 | 9.67 |
| Q15063 | POSTN | 39 | 39 | 4.31 | 12.43 |

| Accession | Name | # Peptides | # Unique Peptides | Abundance Ratio: day 3 / day 0 | Abundance Ratio: day 7/ day 0 |
|---|---|---|---|---|---|
| Q92845 | KIFAP3 | 2 | 1 | 0.01 | 0.97 |
| Q96BM9 | ARL8A | 2 | 1 | 0.01 | 0.76 |
| Q16790 | CA9 | 6 | 6 | 4.47 | 5.54 |
| Q5JTB6 | PLAC9 | 4 | 4 | 4.54 | 5.75 |
| P08123 | COL1A2 | 94 | 93 | 4.99 | 14.41 |
| O00339 | MATN2 | 22 | 22 | 5.10 | 11.95 |
| P16112 | ACAN | 43 | 43 | 5.28 | 13.75 |
| P55001 | MFAP2 | 4 | 4 | 5.28 | 9.44 |
| P49747 | COMP | 21 | 21 | 5.31 | 20.25 |
| Q96GY0 | ZC2HC1A | 2 | 2 | 5.37 | 2.30 |
| P05090 | APOD | 7 | 7 | 5.83 | 9.43 |
| Q02388 | COL7A1 | 7 | 7 | 6.74 | 23.52 |
| Q07507 | DPT | 5 | 5 | 6.86 | 16.10 |
| P02452 | COL1A1 | 109 | 105 | 7.03 | 22.97 |
| Q06828 | FMOD | 8 | 8 | 7.68 | 25.00 |
| Q15782 | CHI3L2 | 9 | 9 | 9.03 | 10.04 |
| P02461 | COL3A1 | 109 | 108 | 11.89 | 27.14 |
| P22914 | CRYGS | 9 | 9 | 12.54 | 31.15 |
| P27658 | COL8A1 | 6 | 6 | 13.27 | 40.19 |

**Table S5** - Differential protein expression between chondrocyte pellet cultures with and without upregulation of WWP2

| Accession | Name | # Peptides | # Unique Peptides | Abundance Ratio: WWP2 / control |
|---|---|---|---|---|
| O00308 | WWP2 | 31 | 28 | 10.10 |
| P09914 | IFIT1 | 2 | 1 | 1.89 |
| P26022 | PTX3 | 2 | 2 | 1.78 |
| Q16832 | DDR2 | 2 | 2 | 1.53 |
| Q96LR5 | UBE2E2 | 4 | 2 | 1.52 |
| P33897 | ABCD1 | 2 | 2 | 1.52 |
| P51965 | UBE2E1 | 3 | 1 | 1.44 |
| P28300 | LOX | 3 | 3 | 1.43 |
| Q96B36 | AKT1S1 | 2 | 2 | 1.38 |
| Q9P2K8 | EIF2AK4 | 3 | 3 | 1.37 |
| Q969T4 | UBE2E3 | 3 | 1 | 1.36 |
| Q9UBX5 | FBLN5 | 5 | 5 | 1.33 |
| P54652 | HSPA2 | 27 | 14 | 1.32 |
| Q01780 | EXOSC10 | 2 | 2 | 1.32 |
| P21741 | MDK | 3 | 3 | 1.31 |
| Q96GY0 | ZC2HC1A | 2 | 2 | 1.31 |

| Accession | Name | # Peptides | # Unique Peptides | Abundance Ratio: WWP2 / control |
|---|---|---|---|---|
| Q9Y4K4 | MAP4K5 | 2 | 2 | 1.31 |
| Q92541 | RTF1 | 3 | 3 | 1.30 |
| P05161 | ISG15 | 3 | 3 | 1.30 |
| P47974 | ZFP36L2 | 2 | 1 | 1.30 |
| P25205 | MCM3 | 2 | 2 | 1.30 |
| P61077 | UBE2D3 | 3 | 2 | 1.30 |
| P62837 | UBE2D2 | 3 | 2 | 1.30 |
| Q9Y4F5 | CEP170B | 2 | 2 | 1.30 |
| Q13613 | MTMR1 | 3 | 3 | 1.30 |
| Q9H7D7 | WDR26 | 2 | 2 | 1.29 |
| O00148 | DDX39A | 14 | 3 | 1.29 |
| Q5RI15 | COX20 | 2 | 2 | 1.28 |
| Q13586 | STIM1 | 2 | 2 | 1.28 |
| Q9Y625 | GPC6 | 6 | 6 | 1.28 |
| P55290 | CDH13 | 6 | 6 | 1.27 |
| Q9HA77 | CARS2 | 4 | 4 | 1.26 |
| Q9H6T3 | RPAP3 | 2 | 2 | 1.26 |
| Q9H2K8 | TAOK3 | 3 | 2 | 1.26 |
| Q5T9L3 | WLS | 2 | 2 | 1.26 |
| Q5RKV6 | EXOSC6 | 3 | 3 | 1.26 |
| Q8IVF2 | AHNAK2 | 15 | 13 | 1.26 |
| Q8IWT0 | ZBTB8OS | 2 | 2 | 1.25 |
| Q05655 | PRKCD | 3 | 2 | 1.25 |
| O94804 | STK10 | 7 | 6 | 1.25 |
| Q96RK0 | CIC | 2 | 2 | 1.25 |
| P83111 | LACTB | 4 | 4 | 1.25 |
| Q9UM22 | EPDR1 | 2 | 2 | 1.25 |
| Q9BQE4 | SELENOS | 2 | 2 | 1.25 |
| Q9HAV7 | GRPEL1 | 2 | 2 | 1.25 |
| P28845 | HSD11B1 | 3 | 3 | 1.25 |
| P29373 | CRABP2 | 3 | 3 | 1.25 |
| Q93062 | RBPMS | 2 | 2 | 1.24 |
| Q5VWZ2 | LYPLAL1 | 2 | 2 | 1.24 |
| O43172 | PRPF4 | 4 | 4 | 1.24 |
| O75179 | ANKRD17 | 3 | 3 | 1.24 |
| Q96II8 | LRCH3 | 2 | 2 | 1.24 |
| Q15276 | RABEP1 | 6 | 6 | 1.24 |
| Q15063 | POSTN | 39 | 39 | 1.24 |
| Q8IY33 | MICALL2 | 3 | 3 | 1.23 |
| O43150 | ASAP2 | 3 | 3 | 1.23 |
| Q9BQS8 | FYCO1 | 10 | 10 | 1.23 |
| O96011 | PEX11B | 4 | 4 | 1.23 |
| Q96GQ7 | DDX27 | 2 | 2 | 1.23 |
| P51668 | UBE2D1 | 2 | 1 | 1.23 |

| Accession | Name | # Peptides | # Unique Peptides | Abundance Ratio: WWP2 / control |
|---|---|---|---|---|
| Q9Y2X8 | UBE2D4 | 2 | 1 | 1.23 |
| O95182 | NDUFA7 | 3 | 3 | 1.23 |
| Q06278 | AOX1 | 3 | 3 | 1.23 |
| P30838 | ALDH3A1 | 4 | 4 | 1.22 |
| O95155 | UBE4B | 3 | 3 | 1.22 |
| Q96A57 | TMEM230 | 3 | 3 | 1.22 |
| P23743 | DGKA | 7 | 7 | 1.22 |
| P20020 | ATP2B1 | 23 | 11 | 1.22 |
| P25067 | COL8A2 | 2 | 2 | 1.22 |
| P35052 | GPC1 | 5 | 5 | 1.22 |
| Q13451 | FKBP5 | 21 | 20 | 1.22 |
| Q9BXR0 | QTRT1 | 2 | 2 | 1.21 |
| O00629 | KPNA4 | 6 | 3 | 1.21 |
| O15523 | DDX3Y | 17 | 2 | 1.21 |
| Q9H0B6 | KLC2 | 5 | 1 | 1.21 |
| Q5HYK7 | SH3D19 | 4 | 4 | 1.21 |
| P19823 | ITIH2 | 2 | 2 | 1.21 |
| O75150 | RNF40 | 2 | 2 | 1.21 |
| O00160 | MYO1F | 3 | 3 | 1.20 |
| Q15007 | WTAP | 4 | 4 | 1.20 |
| O60637 | TSPAN3 | 2 | 2 | 1.20 |
| Q13740 | ALCAM | 9 | 9 | 1.20 |
| P05121 | SERPINE1 | 4 | 4 | 1.20 |
| Q53H96 | PYCR3 | 2 | 2 | 1.20 |
| O60437 | PPL | 17 | 17 | 1.19 |
| O75521 | ECI2 | 4 | 4 | 1.19 |
| O75175 | CNOT3 | 3 | 3 | 1.19 |
| Q14C86 | GAPVD1 | 5 | 5 | 1.19 |
| Q9BWH6 | RPAP1 | 3 | 3 | 1.19 |
| P15104 | GLUL | 8 | 8 | 1.19 |
| Q96B96 | TMEM159 | 3 | 3 | 1.19 |
| Q86U38 | NOP9 | 2 | 2 | 1.18 |
| Q8NEY1 | NAV1 | 3 | 3 | 1.18 |
| O15118 | NPC1 | 5 | 5 | 1.18 |
| Q99700 | ATXN2 | 6 | 4 | 1.18 |
| Q01814 | ATP2B2 | 11 | 1 | 1.18 |
| Q9UDT6 | CLIP2 | 7 | 3 | 1.18 |
| P08294 | SOD3 | 6 | 6 | 1.18 |
| Q14728 | MFSD10 | 3 | 3 | 1.18 |
| P15848 | ARSB | 9 | 9 | 1.18 |
| Q5JPH6 | EARS2 | 3 | 3 | 1.18 |
| P04062 | GBA | 12 | 12 | 1.18 |
| Q9H9C1 | VIPAS39 | 2 | 2 | 1.18 |
| Q5T8D3 | ACBD5 | 3 | 3 | 1.18 |

| Accession | Name | # Peptides | # Unique Peptides | Abundance Ratio: WWP2 / control |
|---|---|---|---|---|
| Q9NQZ5 | STARD7 | 2 | 2 | 1.18 |
| Q96NY7 | CLIC6 | 10 | 10 | 1.17 |
| Q9BZF1 | OSBPL8 | 6 | 5 | 1.17 |
| Q15847 | ADIRF | 6 | 6 | 1.17 |
| Q9Y6N5 | SQOR | 15 | 15 | 1.17 |
| P20810 | CAST | 19 | 19 | 1.17 |
| Q49A26 | GLYR1 | 2 | 2 | 1.17 |
| P13747 | HLA-E | 3 | 1 | 1.17 |
| Q7L7L0 | H2AW | 9 | 4 | 1.17 |
| P04908 | H2AC4 | 9 | 4 | 1.17 |
| P51688 | SGSH | 6 | 6 | 1.17 |
| Q93045 | STMN2 | 3 | 3 | 1.17 |
| Q13336 | SLC14A1 | 3 | 3 | 1.17 |
| Q9H270 | VPS11 | 2 | 2 | 1.17 |
| Q0ZGT2 | NEXN | 6 | 6 | 1.16 |
| Q9BTU6 | PI4K2A | 2 | 2 | 1.16 |
| Q16134 | ETFDH | 4 | 4 | 1.16 |
| P04179 | SOD2 | 10 | 10 | 1.16 |
| Q7L775 | EPM2AIP1 | 3 | 3 | 1.16 |
| O75348 | ATP6V1G1 | 6 | 4 | 1.16 |
| P19532 | TFE3 | 2 | 2 | 1.16 |
| P49356 | FNTB | 2 | 2 | 1.16 |
| P16949 | STMN1 | 15 | 15 | 1.16 |
| Q96HP0 | DOCK6 | 2 | 2 | 1.16 |
| P45877 | PPIC | 3 | 3 | 1.16 |
| Q9NYI0 | PSD3 | 9 | 9 | 1.16 |
| Q9GZT9 | EGLN1 | 5 | 5 | 1.16 |
| P53367 | ARFIP1 | 3 | 3 | 1.16 |
| Q12802 | AKAP13 | 2 | 2 | 1.16 |
| P0DP23 | CALM1 | 11 | 11 | 1.16 |
| Q02952 | AKAP12 | 34 | 34 | 1.16 |
| Q7Z417 | NUFIP2 | 2 | 2 | 1.16 |
| P04792 | HSPB1 | 17 | 17 | 1.16 |
| P22830 | FECH | 3 | 3 | 1.16 |
| Q9Y6D6 | ARFGEF1 | 5 | 3 | 1.16 |
| P31323 | PRKAR2B | 3 | 1 | 1.16 |
| P04350 | TUBB4A | 21 | 1 | 1.16 |
| O75381 | PEX14 | 4 | 4 | 1.16 |
| P10301 | RRAS | 9 | 7 | 1.16 |
| Q03405 | PLAUR | 6 | 6 | 1.16 |
| Q15149 | PLEC | 374 | 366 | 1.16 |
| P34741 | SDC2 | 4 | 4 | 1.15 |
| Q15334 | LLGL1 | 4 | 4 | 1.15 |
| P09936 | UCHL1 | 14 | 14 | 1.15 |

| Accession | Name | # Peptides | # Unique Peptides | Abundance Ratio: WWP2 / control |
|---|---|---|---|---|
| Q13439 | GOLGA4 | 8 | 7 | 1.15 |
| P42356 | PI4KA | 3 | 3 | 1.15 |
| Q9UBN7 | HDAC6 | 2 | 2 | 1.15 |
| P63172 | DYNLT1 | 2 | 2 | 1.15 |
| Q12965 | MYO1E | 11 | 11 | 1.15 |
| Q9BWD1 | ACAT2 | 5 | 5 | 1.15 |
| P52306 | RAP1GDS1 | 9 | 9 | 1.15 |
| Q93008 | USP9X | 24 | 24 | 1.15 |
| P15311 | EZR | 40 | 26 | 1.15 |
| O14933 | UBE2L6 | 2 | 2 | 1.15 |
| Q96C19 | EFHD2 | 10 | 10 | 1.15 |
| Q92598 | HSPH1 | 14 | 11 | 1.15 |
| P04424 | ASL | 6 | 6 | 1.15 |
| Q6ZMZ3 | SYNE3 | 13 | 13 | 1.15 |
| Q9P013 | CWC15 | 5 | 5 | 1.15 |
| Q9Y3A5 | SBDS | 14 | 14 | 1.15 |
| Q9NVH1 | DNAJC11 | 5 | 5 | 1.15 |
| P53801 | PTTG1IP | 2 | 2 | 1.15 |
| Q9Y5P4 | CERT1 | 3 | 3 | 1.15 |
| O00139 | KIF2A | 2 | 2 | 1.15 |
| Q92974 | ARHGEF2 | 4 | 4 | 1.15 |
| Q96GQ5 | RUSF1 | 3 | 3 | 1.15 |
| Q7Z5L9 | IRF2BP2 | 5 | 5 | 1.15 |
| Q99615 | DNAJC7 | 4 | 4 | 1.15 |
| Q6WCQ1 | MPRIP | 5 | 5 | 1.15 |
| Q06481 | APLP2 | 3 | 3 | 1.15 |
| O15347 | HMGB3 | 2 | 1 | 1.15 |
| O96005 | CLPTM1 | 3 | 3 | 1.15 |
| Q14108 | SCARB2 | 14 | 14 | 1.15 |
| P34059 | GALNS | 4 | 3 | 1.15 |
| O00330 | PDHX | 2 | 2 | 1.15 |
| Q6NUK4 | REEP3 | 2 | 2 | 1.15 |
| Q15555 | MAPRE2 | 8 | 7 | 1.15 |
| Q8N3D4 | EHBP1L1 | 4 | 4 | 1.15 |
| Q9UN86 | G3BP2 | 5 | 4 | 1.15 |
| Q6SZW1 | SARM1 | 3 | 3 | 1.15 |
| P35754 | GLRX | 4 | 4 | 1.15 |
| O15484 | CAPN5 | 2 | 2 | 1.14 |
| O14841 | OPLAH | 4 | 4 | 1.14 |
| Q9UPT8 | ZC3H4 | 5 | 4 | 1.14 |
| Q8IY17 | PNPLA6 | 5 | 5 | 1.14 |
| P22570 | FDXR | 6 | 6 | 1.14 |
| Q9Y2L1 | DIS3 | 7 | 7 | 1.14 |
| O43657 | TSPAN6 | 2 | 2 | 1.14 |

| Accession | Name | # Peptides | # Unique Peptides | Abundance Ratio: WWP2 / control |
|---|---|---|---|---|
| P49757 | NUMB | 3 | 2 | 1.14 |
| P17936 | IGFBP3 | 7 | 7 | 1.14 |
| P19793 | RXRA | 2 | 2 | 1.14 |
| Q7L2J0 | MEPCE | 6 | 6 | 1.14 |
| P55196 | AFDN | 4 | 4 | 1.14 |
| Q7Z2W4 | ZC3HAV1 | 8 | 8 | 1.14 |
| Q9Y2D5 | AKAP2 | 11 | 11 | 1.14 |
| P29372 | MPG | 2 | 2 | 1.14 |
| O94805 | ACTL6B | 2 | 2 | 1.14 |
| P02649 | APOE | 6 | 6 | 1.14 |
| Q13938 | CAPS | 10 | 10 | 1.14 |
| Q15650 | TRIP4 | 2 | 2 | 1.14 |
| O60610 | DIAPH1 | 10 | 10 | 1.14 |
| Q12800 | TFCP2 | 2 | 1 | 1.14 |
| P36543 | ATP6V1E1 | 10 | 10 | 1.14 |
| Q8IVM0 | CCDC50 | 3 | 3 | 1.14 |
| P33121 | ACSL1 | 3 | 3 | 1.14 |
| Q92466 | DDB2 | 3 | 3 | 1.14 |
| Q8WXF0 | SRSF12 | 2 | 2 | 1.14 |
| Q6PJT7 | ZC3H14 | 5 | 5 | 1.14 |
| Q9BSD7 | NTPCR | 4 | 4 | 1.13 |
| Q86X76 | NIT1 | 3 | 3 | 1.13 |
| Q14315 | FLNC | 42 | 29 | 1.13 |
| P15144 | ANPEP | 35 | 35 | 1.13 |
| P30711 | GSTT1 | 3 | 2 | 1.13 |
| Q12929 | EPS8 | 9 | 9 | 1.13 |
| P36873 | PPP1CC | 9 | 1 | 1.13 |
| P21266 | GSTM3 | 7 | 5 | 1.13 |
| Q9Y5B9 | SUPT16H | 14 | 14 | 1.13 |
| Q7Z4G1 | COMMD6 | 2 | 2 | 1.13 |
| O15020 | SPTBN2 | 8 | 4 | 1.13 |
| Q6NUM9 | RETSAT | 5 | 5 | 1.13 |
| Q9UHJ6 | SHPK | 2 | 2 | 1.13 |
| Q15654 | TRIP6 | 8 | 8 | 1.13 |
| Q9HAB8 | PPCS | 2 | 2 | 1.13 |
| Q14694 | USP10 | 2 | 2 | 1.13 |
| P55268 | LAMB2 | 21 | 21 | 1.13 |
| P51580 | TPMT | 2 | 2 | 1.13 |
| Q9H078 | CLPB | 2 | 2 | 1.13 |
| P19474 | TRIM21 | 2 | 2 | 1.13 |
| Q96AQ6 | PBXIP1 | 7 | 7 | 1.13 |
| Q99614 | TTC1 | 2 | 2 | 1.13 |
| Q9GZM8 | NDEL1 | 3 | 2 | 1.13 |
| Q8IVD9 | NUDCD3 | 2 | 2 | 1.13 |

| Accession | Name | # Peptides | # Unique Peptides | Abundance Ratio: WWP2 / control |
|---|---|---|---|---|
| P17050 | NAGA | 3 | 3 | 1.13 |
| O95248 | SBF1 | 3 | 3 | 1.13 |
| Q9UKE5 | TNIK | 3 | 1 | 1.13 |
| P26447 | S100A4 | 6 | 6 | 1.13 |
| Q8TEA8 | DTD1 | 4 | 4 | 1.13 |
| Q96K37 | SLC35E1 | 2 | 2 | 1.13 |
| Q9H1E5 | TMX4 | 4 | 4 | 1.13 |
| Q9Y2S2 | CRYL1 | 3 | 3 | 1.13 |
| Q14671 | PUM1 | 2 | 2 | 1.13 |
| Q9NXR1 | NDE1 | 2 | 1 | 1.13 |
| P62070 | RRAS2 | 6 | 4 | 1.13 |
| Q8TB72 | PUM2 | 2 | 2 | 1.13 |
| P08648 | ITGA5 | 13 | 13 | 1.13 |
| P42226 | STAT6 | 15 | 15 | 1.13 |
| P11277 | SPTB | 6 | 4 | 1.13 |
| Q9C0B1 | FTO | 4 | 4 | 1.13 |
| Q86UU1 | PHLDB1 | 22 | 22 | 1.13 |
| Q9UN37 | VPS4A | 8 | 3 | 1.12 |
| P62072 | TIMM10 | 2 | 2 | 1.12 |
| O75688 | PPM1B | 3 | 1 | 1.12 |
| Q7Z4I7 | LIMS2 | 5 | 2 | 1.12 |
| Q8WTS6 | SETD7 | 2 | 2 | 1.12 |
| Q08431 | MFGE8 | 11 | 11 | 1.12 |
| Q9H3H3 | C11orf68 | 4 | 4 | 1.12 |
| Q86V88 | MDP1 | 2 | 2 | 1.12 |
| A6QL63 | BTBD11 | 2 | 2 | 1.12 |
| O14647 | CHD2 | 2 | 1 | 1.12 |
| Q09666 | AHNAK | 328 | 326 | 1.12 |
| Q2TAL8 | QRICH1 | 2 | 2 | 1.12 |
| Q9H173 | SIL1 | 4 | 4 | 1.12 |
| Q15651 | HMGN3 | 2 | 2 | 1.12 |
| O00507 | USP9Y | 7 | 7 | 1.12 |
| P50416 | CPT1A | 5 | 5 | 1.12 |
| Q06787 | FMR1 | 5 | 2 | 1.12 |
| P49736 | MCM2 | 2 | 2 | 1.12 |
| Q9NZ08 | ERAP1 | 3 | 3 | 1.12 |
| Q9BUR5 | APOO | 2 | 2 | 1.12 |
| Q93034 | CUL5 | 6 | 6 | 1.12 |
| P06493 | CDK1 | 2 | 2 | 1.12 |
| Q96ED9 | HOOK2 | 3 | 3 | 1.12 |
| O15533 | TAPBP | 4 | 4 | 1.12 |
| P46937 | YAP1 | 5 | 5 | 1.12 |
| P29353 | SHC1 | 4 | 4 | 1.12 |
| Q9NZT2 | OGFR | 3 | 3 | 1.12 |

| Accession | Name | # Peptides | # Unique Peptides | Abundance Ratio: WWP2 / control |
|---|---|---|---|---|
| Q15645 | TRIP13 | 2 | 2 | 1.12 |
| Q05707 | COL14A1 | 46 | 46 | 1.12 |
| P68036 | UBE2L3 | 4 | 4 | 1.12 |
| Q03001 | DST | 21 | 20 | 1.12 |
| Q96CV9 | OPTN | 13 | 13 | 1.12 |
| Q9H1E3 | NUCKS1 | 3 | 3 | 1.12 |
| Q13671 | RIN1 | 2 | 2 | 1.12 |
| Q96HE7 | ERO1A | 17 | 17 | 1.12 |
| P0DMV8 | HSPA1A | 35 | 28 | 1.12 |
| Q9BXP2 | SLC12A9 | 2 | 2 | 1.12 |
| P07738 | BPGM | 5 | 5 | 1.12 |
| Q9BV20 | MRI1 | 6 | 6 | 1.11 |
| O15294 | OGT | 3 | 3 | 1.11 |
| Q6PIU2 | NCEH1 | 6 | 6 | 1.11 |
| Q8TBX8 | PIP4K2C | 3 | 3 | 1.11 |
| Q96PC5 | MIA2 | 3 | 2 | 1.11 |
| P11172 | UMPS | 3 | 3 | 1.11 |
| P63218 | GNG5 | 2 | 2 | 1.11 |
| A0A1B0GUS4 | UBE2L5 | 3 | 3 | 1.11 |
| Q9UEW8 | STK39 | 6 | 4 | 1.11 |
| Q13619 | CUL4A | 12 | 5 | 1.11 |
| Q66PJ3 | ARL6IP4 | 3 | 3 | 1.11 |
| Q27J81 | INF2 | 18 | 18 | 1.11 |
| P52701 | MSH6 | 2 | 2 | 1.11 |
| Q9Y6H1 | CHCHD2 | 2 | 2 | 1.11 |
| Q96JY6 | PDLIM2 | 3 | 3 | 1.11 |
| P78559 | MAP1A | 48 | 48 | 1.11 |
| Q7L1Q6 | BZW1 | 17 | 17 | 1.11 |
| P18084 | ITGB5 | 7 | 7 | 1.11 |
| Q8WWI1 | LMO7 | 26 | 26 | 1.11 |
| Q5T1J5 | CHCHD2P9 | 2 | 2 | 1.11 |
| Q9H0A0 | NAT10 | 5 | 5 | 1.11 |
| Q96MW5 | COG8 | 3 | 3 | 1.11 |
| Q14160 | SCRIB | 2 | 2 | 1.11 |
| Q5SW79 | CEP170 | 8 | 8 | 1.11 |
| Q96ET8 | TVP23C | 2 | 2 | 1.11 |
| P17900 | GM2A | 2 | 2 | 1.11 |
| Q9GZR7 | DDX24 | 10 | 10 | 1.11 |
| P26583 | HMGB2 | 12 | 11 | 1.11 |
| Q9HD67 | MYO10 | 2 | 2 | 1.11 |
| Q9NS15 | LTBP3 | 2 | 2 | 1.11 |
| O75131 | CPNE3 | 14 | 13 | 1.11 |
| P60510 | PPP4C | 4 | 4 | 1.11 |
| Q8IYB5 | SMAP1 | 3 | 3 | 1.11 |

| Accession | Name | # Peptides | # Unique Peptides | Abundance Ratio: WWP2 / control |
|---|---|---|---|---|
| P14543 | NID1 | 4 | 3 | 1.11 |
| Q63HN8 | RNF213 | 6 | 6 | 1.11 |
| Q9NNW7 | TXNRD2 | 3 | 2 | 1.11 |
| P17096 | HMGA1 | 8 | 8 | 1.11 |
| P59768 | GNG2 | 2 | 2 | 1.11 |
| O60443 | GSDME | 2 | 2 | 1.11 |
| Q9NZL4 | HSPBP1 | 3 | 3 | 1.11 |
| O75439 | PMPCB | 5 | 4 | 1.11 |
| Q9BX68 | HINT2 | 3 | 3 | 1.11 |
| P01889 | HLA-B | 8 | 3 | 1.11 |
| O94885 | SASH1 | 6 | 6 | 1.11 |
| Q6P1N0 | CC2D1A | 3 | 2 | 1.11 |
| P80723 | BASP1 | 8 | 8 | 1.11 |
| Q96A05 | ATP6V1E2 | 6 | 6 | 1.11 |
| P15090 | FABP4 | 5 | 5 | 1.11 |
| P36404 | ARL2 | 5 | 5 | 1.11 |
| O00116 | AGPS | 11 | 11 | 1.11 |
| O43760 | SYNGR2 | 2 | 2 | 1.11 |
| Q8N697 | SLC15A4 | 2 | 2 | 1.11 |
| Q9UI12 | ATP6V1H | 10 | 10 | 1.11 |
| Q15942 | ZYX | 11 | 11 | 1.11 |
| P19784 | CSNK2A2 | 4 | 3 | 1.11 |
| Q03154 | ACY1 | 8 | 8 | 1.11 |
| Q8N128 | FAM177A1 | 3 | 3 | 1.10 |
| Q6PKG0 | LARP1 | 16 | 16 | 1.10 |
| Q13501 | SQSTM1 | 5 | 5 | 1.10 |
| P20962 | PTMS | 4 | 4 | 1.10 |
| O43768 | ENSA | 5 | 4 | 1.10 |
| O43617 | TRAPPC3 | 3 | 3 | 1.10 |
| P54709 | ATP1B3 | 3 | 3 | 1.10 |
| Q13526 | PIN1 | 3 | 3 | 1.10 |
| P41208 | CETN2 | 3 | 3 | 1.10 |
| Q7LBR1 | CHMP1B | 5 | 5 | 1.10 |
| P29279 | CCN2 | 7 | 7 | 1.10 |
| P15502 | ELN | 4 | 4 | 1.10 |
| Q9H1H9 | KIF13A | 3 | 2 | 1.10 |
| O14782 | KIF3C | 2 | 2 | 1.10 |
| Q9NYZ1 | TVP23B | 3 | 3 | 1.10 |
| Q15056 | EIF4H | 10 | 10 | 1.10 |
| Q96KP1 | EXOC2 | 5 | 5 | 1.10 |
| P82094 | TMF1 | 2 | 2 | 1.10 |
| P23497 | SP100 | 7 | 5 | 1.10 |
| Q8NBK3 | SUMF1 | 2 | 2 | 1.10 |
| O60264 | SMARCA5 | 5 | 4 | 1.10 |

| Accession | Name | # Peptides | # Unique Peptides | Abundance Ratio: WWP2 / control |
|---|---|---|---|---|
| Q8IXT5 | RBM12B | 4 | 4 | 1.10 |
| Q9P0I2 | EMC3 | 3 | 3 | 1.10 |
| O94776 | MTA2 | 7 | 6 | 1.10 |
| O00442 | RTCA | 3 | 3 | 1.10 |
| P32456 | GBP2 | 9 | 8 | 1.10 |
| Q9Y4K0 | LOXL2 | 8 | 8 | 1.10 |
| Q9NXC5 | MIOS | 2 | 2 | 1.10 |
| O00154 | ACOT7 | 7 | 7 | 1.10 |
| O95670 | ATP6V1G2 | 3 | 1 | 1.10 |
| Q9UJW0 | DCTN4 | 9 | 9 | 1.10 |
| O95479 | H6PD | 16 | 16 | 1.10 |
| O60684 | KPNA6 | 10 | 7 | 1.10 |
| Q9Y223 | GNE | 3 | 3 | 1.10 |
| Q9BWH2 | FUNDC2 | 2 | 2 | 1.10 |
| Q8N3Y1 | FBXW8 | 2 | 2 | 1.10 |
| P21283 | ATP6V1C1 | 9 | 9 | 1.10 |
| Q9Y6K9 | IKBKG | 8 | 8 | 1.10 |
| P20936 | RASA1 | 4 | 4 | 1.10 |
| Q16831 | UPP1 | 6 | 6 | 1.10 |
| Q96DE0 | NUDT16 | 2 | 2 | 1.10 |
| Q92597 | NDRG1 | 9 | 9 | 1.10 |
| Q709F0 | ACAD11 | 2 | 2 | 1.10 |
| Q15814 | TBCC | 4 | 4 | 1.10 |
| P53004 | BLVRA | 6 | 6 | 1.10 |
| Q8IUL8 | CILP2 | 5 | 5 | 1.10 |
| P08779 | KRT16 | 2 | 1 | 1.10 |
| Q02388 | COL7A1 | 8 | 8 | 1.10 |
| P35914 | HMGCL | 6 | 6 | 1.10 |
| P62979 | RPS27A | 10 | 2 | 1.10 |
| O14657 | TOR1B | 2 | 2 | 1.10 |
| P42785 | PRCP | 5 | 5 | 1.10 |
| P01034 | CST3 | 2 | 2 | 1.10 |
| O75694 | NUP155 | 10 | 10 | 1.10 |
| P61601 | NCALD | 3 | 3 | 1.10 |
| O60925 | PFDN1 | 5 | 5 | 1.10 |
| Q9Y2U8 | LEMD3 | 5 | 5 | 1.10 |
| O43314 | PPIP5K2 | 3 | 3 | 1.10 |
| Q969G5 | CAVIN3 | 13 | 13 | 1.10 |
| Q3KQU3 | MAP7D1 | 5 | 5 | 1.10 |
| P68366 | TUBA4A | 17 | 2 | 1.10 |
| P61289 | PSME3 | 3 | 3 | 1.10 |
| P53602 | MVD | 2 | 2 | 1.10 |
| Q14157 | UBAP2L | 15 | 15 | 1.10 |
| Q96DI7 | SNRNP40 | 4 | 4 | 1.10 |

| Accession | Name | # Peptides | # Unique Peptides | Abundance Ratio: WWP2 / control |
|---|---|---|---|---|
| Q16610 | ECM1 | 2 | 2 | 1.10 |
| P52788 | SMS | 6 | 6 | 1.10 |
| Q8IYB3 | SRRM1 | 3 | 3 | 1.10 |
| Q9Y394 | DHRS7 | 5 | 5 | 1.10 |
| P43304 | GPD2 | 17 | 17 | 1.10 |
| Q9Y5A7 | NUB1 | 4 | 4 | 1.09 |
| Q9Y4I1 | MYO5A | 7 | 7 | 1.09 |
| Q9C0E2 | XPO4 | 3 | 3 | 1.09 |
| Q9NPH2 | ISYNA1 | 3 | 3 | 1.09 |
| Q9H7C9 | AAMDC | 6 | 6 | 1.09 |
| Q9BRX2 | PELO | 8 | 8 | 1.09 |
| P56545 | CTBP2 | 6 | 2 | 1.09 |
| O43290 | SART1 | 8 | 8 | 1.09 |
| Q13617 | CUL2 | 11 | 11 | 1.09 |
| Q9H0L4 | CSTF2T | 3 | 3 | 1.09 |
| P16435 | POR | 13 | 13 | 1.09 |
| Q16864 | ATP6V1F | 2 | 2 | 1.09 |
| P62310 | LSM3 | 3 | 3 | 1.09 |
| Q9UDY4 | DNAJB4 | 2 | 2 | 1.09 |
| Q8IYK4 | COLGALT2 | 2 | 2 | 1.09 |
| Q9Y2H6 | FNDC3A | 11 | 11 | 1.09 |
| O95070 | YIF1A | 2 | 2 | 1.09 |
| Q9UI10 | EIF2B4 | 7 | 7 | 1.09 |
| Q96TA1 | NIBAN2 | 23 | 23 | 1.09 |
| Q9NZD2 | GLTP | 2 | 2 | 1.09 |
| Q8WUM0 | NUP133 | 3 | 3 | 1.09 |
| Q9BUF5 | TUBB6 | 16 | 4 | 1.09 |
| Q8WVJ2 | NUDCD2 | 3 | 3 | 1.09 |
| Q99447 | PCYT2 | 5 | 5 | 1.09 |
| Q96EK5 | KIFBP | 3 | 3 | 1.09 |
| P17812 | CTPS1 | 7 | 6 | 1.09 |
| O75880 | SCO1 | 3 | 3 | 1.09 |
| Q16539 | MAPK14 | 4 | 4 | 1.09 |
| P49770 | EIF2B2 | 4 | 4 | 1.09 |
| Q7LG56 | RRM2B | 5 | 5 | 1.09 |
| P12110 | COL6A2 | 46 | 46 | 1.09 |
| Q53H82 | LACTB2 | 2 | 2 | 1.09 |
| P62993 | GRB2 | 6 | 6 | 1.09 |
| Q9UBB4 | ATXN10 | 4 | 4 | 1.09 |
| Q9H8H3 | METTL7A | 3 | 3 | 1.09 |
| Q14573 | ITPR3 | 17 | 13 | 1.09 |
| Q9BUT1 | BDH2 | 9 | 9 | 1.09 |
| Q9Y450 | HBS1L | 12 | 12 | 1.09 |
| P68363 | TUBA1B | 21 | 2 | 1.09 |

| Accession | Name | # Peptides | # Unique Peptides | Abundance Ratio: WWP2 / control |
|---|---|---|---|---|
| Q7Z3B1 | NEGR1 | 6 | 6 | 1.09 |
| P36915 | GNL1 | 5 | 5 | 1.09 |
| Q99720 | SIGMAR1 | 2 | 2 | 1.09 |
| Q9UNH6 | SNX7 | 2 | 2 | 1.09 |
| Q71U36 | TUBA1A | 22 | 2 | 1.09 |
| Q9UP83 | COG5 | 2 | 2 | 1.09 |
| Q14643 | ITPR1 | 34 | 28 | 1.09 |
| O60504 | SORBS3 | 7 | 7 | 1.09 |
| P15313 | ATP6V1B1 | 3 | 3 | 1.09 |
| P02511 | CRYAB | 14 | 14 | 1.09 |
| O43148 | RNMT | 2 | 2 | 1.09 |
| Q92522 | H1-10 | 14 | 14 | 1.09 |
| Q9BRZ2 | TRIM56 | 4 | 4 | 1.09 |
| O75976 | CPD | 2 | 2 | 1.09 |
| Q92973 | TNPO1 | 8 | 5 | 1.09 |
| Q92643 | PIGK | 2 | 2 | 1.09 |
| P37235 | HPCAL1 | 7 | 7 | 1.09 |
| P84074 | HPCA | 6 | 6 | 1.09 |
| P56539 | CAV3 | 2 | 2 | 1.09 |
| O95817 | BAG3 | 17 | 17 | 1.09 |
| Q13464 | ROCK1 | 18 | 15 | 1.09 |
| Q5T4B2 | CERCAM | 2 | 2 | 1.09 |
| Q12979 | ABR | 4 | 4 | 1.09 |
| P60033 | CD81 | 4 | 4 | 1.09 |
| P49419 | ALDH7A1 | 16 | 16 | 1.09 |
| Q9BYT8 | NLN | 3 | 3 | 1.09 |
| Q9P1U1 | ACTR3B | 2 | 2 | 1.09 |
| Q8IZ21 | PHACTR4 | 2 | 2 | 1.09 |
| P28065 | PSMB9 | 3 | 3 | 1.09 |
| Q9H330 | TMEM245 | 2 | 2 | 1.09 |
| P35568 | IRS1 | 4 | 4 | 1.09 |
| Q86YR5 | GPSM1 | 4 | 4 | 1.09 |
| A8K0Z3 | WASHC1 | 4 | 1 | 1.09 |
| A8MWX3 | WASH4P | 3 | 1 | 1.09 |
| Q15061 | WDR43 | 3 | 3 | 1.09 |
| Q13813 | SPTAN1 | 115 | 115 | 1.09 |
| P11142 | HSPA8 | 39 | 24 | 1.09 |
| P11234 | RALB | 4 | 3 | 1.09 |
| Q08378 | GOLGA3 | 22 | 22 | 1.08 |
| Q712K3 | UBE2R2 | 4 | 3 | 1.08 |
| Q9BQA1 | WDR77 | 3 | 3 | 1.08 |
| Q99590 | SCAF11 | 2 | 2 | 1.08 |
| Q01484 | ANK2 | 5 | 4 | 1.08 |
| Q14789 | GOLGB1 | 26 | 26 | 1.08 |

| Accession | Name | # Peptides | # Unique Peptides | Abundance Ratio: WWP2 / control |
|---|---|---|---|---|
| P23921 | RRM1 | 5 | 5 | 1.08 |
| P49821 | NDUFV1 | 10 | 10 | 1.08 |
| O14602 | EIF1AY | 6 | 6 | 1.08 |
| Q9UHB6 | LIMA1 | 22 | 22 | 1.08 |
| Q9Y3D9 | MRPS23 | 2 | 2 | 1.08 |
| Q9BRS8 | LARP6 | 2 | 2 | 1.08 |
| P82912 | MRPS11 | 2 | 2 | 1.08 |
| P11586 | MTHFD1 | 29 | 28 | 1.08 |
| O43432 | EIF4G3 | 12 | 8 | 1.08 |
| P47813 | EIF1AX | 6 | 6 | 1.08 |
| P52926 | HMGA2 | 5 | 5 | 1.08 |
| Q15386 | UBE3C | 3 | 3 | 1.08 |
| P16278 | GLB1 | 8 | 8 | 1.08 |
| P23919 | DTYMK | 7 | 7 | 1.08 |
| P42330 | AKR1C3 | 17 | 9 | 1.08 |
| Q6QNY0 | BLOC1S3 | 2 | 2 | 1.08 |
| Q9UIQ6 | LNPEP | 7 | 7 | 1.08 |
| Q5T0F9 | CC2D1B | 2 | 2 | 1.08 |
| Q9Y6D9 | MAD1L1 | 4 | 4 | 1.08 |
| Q99426 | TBCB | 10 | 10 | 1.08 |
| Q8TD55 | PLEKHO2 | 7 | 7 | 1.08 |
| Q2TB10 | ZNF800 | 2 | 2 | 1.08 |
| Q6GMV2 | SMYD5 | 4 | 4 | 1.08 |
| Q12846 | STX4 | 4 | 4 | 1.08 |
| Q06203 | PPAT | 4 | 4 | 1.08 |
| Q9H0C8 | ILKAP | 3 | 3 | 1.08 |
| Q13951 | CBFB | 4 | 4 | 1.08 |
| Q08AM6 | VAC14 | 4 | 4 | 1.08 |
| Q9Y608 | LRRFIP2 | 6 | 5 | 1.08 |
| O60271 | SPAG9 | 21 | 21 | 1.08 |
| O15321 | TM9SF1 | 5 | 5 | 1.08 |
| Q96DB5 | RMDN1 | 4 | 4 | 1.08 |
| Q8NFH3 | NUP43 | 2 | 2 | 1.08 |
| P35611 | ADD1 | 15 | 15 | 1.08 |
| O95819 | MAP4K4 | 13 | 11 | 1.08 |
| P26196 | DDX6 | 12 | 12 | 1.08 |
| Q12931 | TRAP1 | 7 | 7 | 1.08 |
| Q9UGT4 | SUSD2 | 7 | 7 | 1.08 |
| P78406 | RAE1 | 7 | 7 | 1.08 |
| P08962 | CD63 | 4 | 4 | 1.08 |
| Q9UM47 | NOTCH3 | 4 | 3 | 1.08 |
| P78318 | IGBP1 | 7 | 7 | 1.08 |
| Q9UGI8 | TES | 11 | 11 | 1.08 |
| Q6P996 | PDXDC1 | 9 | 9 | 1.08 |

| Accession | Name | # Peptides | # Unique Peptides | Abundance Ratio: WWP2 / control |
|---|---|---|---|---|
| Q7Z460 | CLASP1 | 7 | 7 | 1.08 |
| P17931 | LGALS3 | 12 | 12 | 1.08 |
| Q14320 | FAM50A | 3 | 3 | 1.08 |
| P68371 | TUBB4B | 24 | 1 | 1.08 |
| P56537 | EIF6 | 5 | 5 | 1.08 |
| Q5VT25 | CDC42BPA | 5 | 1 | 1.08 |
| Q9NXH9 | TRMT1 | 3 | 3 | 1.08 |
| Q6UW68 | TMEM205 | 2 | 2 | 1.08 |
| Q99714 | HSD17B10 | 6 | 6 | 1.08 |
| P04083 | ANXA1 | 26 | 26 | 1.08 |
| O95674 | CDS2 | 3 | 3 | 1.08 |
| Q96DU9 | PABPC5 | 2 | 2 | 1.08 |
| P12109 | COL6A1 | 46 | 46 | 1.08 |
| Q86WR0 | CCDC25 | 2 | 2 | 1.08 |
| Q5TFE4 | NT5DC1 | 2 | 2 | 1.08 |
| P09661 | SNRPA1 | 7 | 7 | 1.08 |
| P10321 | HLA-C | 9 | 4 | 1.08 |
| Q70CQ2 | USP34 | 3 | 3 | 1.08 |
| O75116 | ROCK2 | 22 | 19 | 1.08 |
| P30085 | CMPK1 | 8 | 8 | 1.08 |
| Q92888 | ARHGEF1 | 11 | 11 | 1.08 |
| Q5JSZ5 | PRRC2B | 2 | 2 | 1.08 |
| Q14966 | ZNF638 | 5 | 5 | 1.08 |
| P40818 | USP8 | 3 | 3 | 1.08 |
| Q9Y5X3 | SNX5 | 12 | 11 | 1.08 |
| Q8NB37 | GATD1 | 3 | 3 | 1.08 |
| Q15102 | PAFAH1B3 | 7 | 7 | 1.08 |
| Q9GZP4 | PITHD1 | 5 | 5 | 1.08 |
| P05204 | HMGN2 | 6 | 5 | 1.08 |
| Q8WVY7 | UBLCP1 | 5 | 5 | 1.08 |
| Q92930 | RAB8B | 7 | 4 | 1.08 |
| P78344 | EIF4G2 | 20 | 20 | 1.08 |
| Q6P2E9 | EDC4 | 6 | 6 | 1.08 |
| Q13509 | TUBB3 | 22 | 5 | 1.08 |
| P48741 | HSPA7 | 5 | 4 | 1.08 |
| P05106 | ITGB3 | 2 | 2 | 1.08 |
| P41227 | NAA10 | 5 | 5 | 1.08 |
| P20648 | ATP4A | 5 | 5 | 1.08 |
| Q9ULF5 | SLC39A10 | 2 | 2 | 1.07 |
| Q15628 | TRADD | 3 | 3 | 1.07 |
| Q9H993 | ARMT1 | 5 | 5 | 1.07 |
| Q13554 | CAMK2B | 5 | 1 | 1.07 |
| Q16647 | PTGIS | 2 | 2 | 1.07 |
| Q99816 | TSG101 | 4 | 4 | 1.07 |

| Accession | Name | # Peptides | # Unique Peptides | Abundance Ratio: WWP2 / control |
|---|---|---|---|---|
| Q13555 | CAMK2G | 7 | 3 | 1.07 |
| Q9BRJ6 | C7orf50 | 2 | 2 | 1.07 |
| P51648 | ALDH3A2 | 14 | 14 | 1.07 |
| P51114 | FXR1 | 11 | 8 | 1.07 |
| O95816 | BAG2 | 8 | 8 | 1.07 |
| Q9H6F5 | CCDC86 | 3 | 3 | 1.07 |
| P98082 | DAB2 | 13 | 13 | 1.07 |
| Q9UBR2 | CTSZ | 6 | 6 | 1.07 |
| Q9UI30 | TRMT112 | 4 | 4 | 1.07 |
| P49184 | DNASE1L1 | 4 | 4 | 1.07 |
| Q9H2G2 | SLK | 16 | 15 | 1.07 |
| Q9BQ49 | SMIM7 | 3 | 3 | 1.07 |
| Q9Y277 | VDAC3 | 10 | 10 | 1.07 |
| Q53EL6 | PDCD4 | 10 | 10 | 1.07 |
| Q9H9E3 | COG4 | 4 | 4 | 1.07 |
| Q96B97 | SH3KBP1 | 10 | 10 | 1.07 |
| P30519 | HMOX2 | 4 | 4 | 1.07 |
| P36776 | LONP1 | 6 | 6 | 1.07 |
| Q15057 | ACAP2 | 6 | 6 | 1.07 |
| Q01433 | AMPD2 | 14 | 14 | 1.07 |
| O75914 | PAK3 | 3 | 3 | 1.07 |
| Q16563 | SYPL1 | 3 | 3 | 1.07 |
| Q14657 | LAGE3 | 3 | 3 | 1.07 |
| Q96GK7 | FAHD2A | 3 | 3 | 1.07 |
| Q68CQ7 | GLT8D1 | 2 | 2 | 1.07 |
| Q96G61 | NUDT11 | 4 | 2 | 1.07 |
| Q8NFP7 | NUDT10 | 4 | 2 | 1.07 |
| Q01518 | CAP1 | 27 | 26 | 1.07 |
| Q9UPT6 | MAPK8IP3 | 2 | 2 | 1.07 |
| Q12768 | WASHC5 | 9 | 9 | 1.07 |
| Q9HA65 | TBC1D17 | 4 | 4 | 1.07 |
| P46821 | MAP1B | 55 | 54 | 1.07 |
| P50995 | ANXA11 | 21 | 21 | 1.07 |
| P42126 | ECI1 | 2 | 2 | 1.07 |
| Q02252 | ALDH6A1 | 13 | 13 | 1.07 |
| Q04446 | GBE1 | 25 | 25 | 1.07 |
| Q96QC0 | PPP1R10 | 2 | 2 | 1.07 |
| C4AMC7 | WASH3P | 5 | 2 | 1.07 |
| Q6VEQ5 | WASH2P | 5 | 2 | 1.07 |
| Q96S97 | MYADM | 2 | 2 | 1.07 |
| Q9Y3B8 | REXO2 | 6 | 6 | 1.07 |
| Q14204 | DYNC1H1 | 160 | 160 | 1.07 |
| Q96A72 | MAGOHB | 7 | 1 | 1.07 |
| P19021 | PAM | 2 | 2 | 1.07 |

| Accession | Name | # Peptides | # Unique Peptides | Abundance Ratio: WWP2 / control |
|---|---|---|---|---|
| Q99439 | CNN2 | 11 | 9 | 1.07 |
| Q9UBM7 | DHCR7 | 4 | 4 | 1.07 |
| P56199 | ITGA1 | 3 | 3 | 1.07 |
| Q01082 | SPTBN1 | 112 | 106 | 1.07 |
| Q9UMX0 | UBQLN1 | 6 | 3 | 1.07 |
| Q03518 | TAP1 | 2 | 2 | 1.07 |
| Q13315 | ATM | 3 | 3 | 1.07 |
| P26038 | MSN | 63 | 47 | 1.07 |
| Q15042 | RAB3GAP1 | 5 | 5 | 1.07 |
| O15127 | SCAMP2 | 4 | 4 | 1.07 |
| Q12972 | PPP1R8 | 2 | 2 | 1.07 |
| Q9Y6Q5 | AP1M2 | 3 | 1 | 1.07 |
| O60841 | EIF5B | 15 | 15 | 1.07 |
| P57740 | NUP107 | 3 | 3 | 1.07 |
| P14324 | FDPS | 6 | 6 | 1.07 |
| Q05209 | PTPN12 | 7 | 7 | 1.07 |
| Q9NVI7 | ATAD3A | 16 | 15 | 1.07 |
| Q9UK41 | VPS28 | 2 | 2 | 1.07 |
| Q13637 | RAB32 | 4 | 4 | 1.07 |
| Q9BXJ9 | NAA15 | 12 | 12 | 1.07 |
| Q12888 | TP53BP1 | 20 | 20 | 1.07 |
| P42765 | ACAA2 | 12 | 12 | 1.07 |
| O95757 | HSPA4L | 7 | 5 | 1.07 |
| P12111 | COL6A3 | 166 | 166 | 1.07 |
| Q14498 | RBM39 | 7 | 7 | 1.07 |
| O94906 | PRPF6 | 15 | 15 | 1.07 |
| O43847 | NRDC | 9 | 9 | 1.07 |
| Q9UEY8 | ADD3 | 9 | 9 | 1.07 |
| P06737 | PYGL | 40 | 31 | 1.07 |
| Q5T2N8 | ATAD3C | 4 | 4 | 1.07 |
| Q86YZ3 | HRNR | 6 | 6 | 1.07 |
| O00506 | STK25 | 2 | 1 | 1.07 |
| P18031 | PTPN1 | 7 | 7 | 1.07 |
| Q6UB35 | MTHFD1L | 4 | 3 | 1.07 |
| Q16678 | CYP1B1 | 9 | 9 | 1.07 |
| O75369 | FLNB | 94 | 80 | 1.07 |
| Q8NDH3 | NPEPL1 | 4 | 4 | 1.07 |
| Q9H0X9 | OSBPL5 | 7 | 6 | 1.07 |
| Q9H4M9 | EHD1 | 19 | 11 | 1.07 |
| Q96P70 | IPO9 | 12 | 12 | 1.07 |
| P16333 | NCK1 | 4 | 4 | 1.07 |
| Q9Y2U5 | MAP3K2 | 3 | 2 | 1.07 |
| Q16881 | TXNRD1 | 14 | 13 | 1.07 |
| Q8TDZ2 | MICAL1 | 9 | 9 | 1.07 |

| Accession | Name | # Peptides | # Unique Peptides | Abundance Ratio: WWP2 / control |
|---|---|---|---|---|
| Q9BRP8 | PYM1 | 4 | 4 | 1.07 |
| P46060 | RANGAP1 | 9 | 9 | 1.07 |
| Q9Y2G5 | POFUT2 | 7 | 7 | 1.07 |
| Q9Y5S9 | RBM8A | 5 | 5 | 1.07 |
| P27487 | DPP4 | 28 | 28 | 1.07 |
| P24752 | ACAT1 | 7 | 7 | 1.07 |
| P15559 | NQO1 | 12 | 12 | 1.07 |
| P27694 | RPA1 | 12 | 12 | 1.07 |
| Q7Z6Z7 | HUWE1 | 44 | 44 | 1.07 |
| P04040 | CAT | 11 | 11 | 1.07 |
| P15927 | RPA2 | 5 | 5 | 1.07 |
| Q14008 | CKAP5 | 16 | 16 | 1.07 |
| Q9Y6D5 | ARFGEF2 | 4 | 2 | 1.07 |
| Q9BXK5 | BCL2L13 | 3 | 3 | 1.07 |
| Q8TDY4 | ASAP3 | 2 | 2 | 1.07 |
| Q13057 | COASY | 4 | 4 | 1.07 |
| P54707 | ATP12A | 5 | 5 | 1.07 |
| P05155 | SERPING1 | 2 | 2 | 1.07 |
| Q2TB90 | HKDC1 | 3 | 2 | 1.07 |
| Q9BTY2 | FUCA2 | 4 | 4 | 1.07 |
| Q6YN16 | HSDL2 | 6 | 6 | 1.07 |
| Q8NBX0 | SCCPDH | 5 | 5 | 1.07 |
| Q7Z7N9 | TMEM179B | 2 | 2 | 1.07 |
| Q15836 | VAMP3 | 7 | 7 | 1.07 |
| P27449 | ATP6V0C | 2 | 2 | 1.07 |
| Q9NRX5 | SERINC1 | 2 | 2 | 1.07 |
| O75475 | PSIP1 | 8 | 7 | 1.07 |
| O95232 | LUC7L3 | 2 | 2 | 1.06 |
| Q9NZJ9 | NUDT4 | 5 | 3 | 1.06 |
| Q8IX12 | CCAR1 | 8 | 7 | 1.06 |
| O95372 | LYPLA2 | 4 | 4 | 1.06 |
| Q96PE2 | ARHGEF17 | 3 | 3 | 1.06 |
| Q5T8P6 | RBM26 | 3 | 2 | 1.06 |
| Q06330 | RBPJ | 3 | 3 | 1.06 |
| Q9BUQ8 | DDX23 | 8 | 8 | 1.06 |
| P49023 | PXN | 7 | 7 | 1.06 |
| Q13045 | FLII | 13 | 13 | 1.06 |
| O00299 | CLIC1 | 11 | 11 | 1.06 |
| P48634 | PRRC2A | 4 | 3 | 1.06 |
| P13807 | GYS1 | 14 | 13 | 1.06 |
| Q9UHD9 | UBQLN2 | 6 | 3 | 1.06 |
| Q9NYM9 | BET1L | 2 | 2 | 1.06 |
| P43490 | NAMPT | 17 | 17 | 1.06 |
| Q9Y4E8 | USP15 | 4 | 4 | 1.06 |

| Accession | Name | # Peptides | # Unique Peptides | Abundance Ratio: WWP2 / control |
|---|---|---|---|---|
| O43447 | PPIH | 2 | 1 | 1.06 |
| Q9UJ41 | RABGEF1 | 3 | 3 | 1.06 |
| Q14974 | KPNB1 | 20 | 20 | 1.06 |
| P31327 | CPS1 | 2 | 2 | 1.06 |
| Q96SU4 | OSBPL9 | 3 | 3 | 1.06 |
| A5YKK6 | CNOT1 | 10 | 10 | 1.06 |
| Q5T9A4 | ATAD3B | 10 | 9 | 1.06 |
| Q8NFH5 | NUP35 | 3 | 3 | 1.06 |
| O60885 | BRD4 | 3 | 3 | 1.06 |
| P36639 | NUDT1 | 4 | 4 | 1.06 |
| Q9NQC3 | RTN4 | 7 | 7 | 1.06 |
| O00186 | STXBP3 | 7 | 7 | 1.06 |
| P17516 | AKR1C4 | 8 | 2 | 1.06 |
| Q8NEU8 | APPL2 | 15 | 15 | 1.06 |
| Q9ULA0 | DNPEP | 7 | 7 | 1.06 |
| P54105 | CLNS1A | 2 | 2 | 1.06 |
| Q92820 | GGH | 6 | 6 | 1.06 |
| Q9NW64 | RBM22 | 2 | 2 | 1.06 |
| Q8WU90 | ZC3H15 | 8 | 8 | 1.06 |
| Q9NR19 | ACSS2 | 3 | 3 | 1.06 |
| P46734 | MAP2K3 | 7 | 7 | 1.06 |
| Q9Y2P8 | RCL1 | 2 | 2 | 1.06 |
| O94901 | SUN1 | 7 | 7 | 1.06 |
| Q5T4S7 | UBR4 | 22 | 22 | 1.06 |
| O95429 | BAG4 | 2 | 2 | 1.06 |
| Q53FA7 | TP53I3 | 7 | 7 | 1.06 |
| Q99962 | SH3GL2 | 5 | 5 | 1.06 |
| Q13733 | ATP1A4 | 8 | 8 | 1.06 |
| Q7L9L4 | MOB1B | 2 | 2 | 1.06 |
| Q9H8S9 | MOB1A | 2 | 2 | 1.06 |
| P06241 | FYN | 4 | 2 | 1.06 |
| Q8WWY3 | PRPF31 | 9 | 9 | 1.06 |
| Q8N142 | ADSS1 | 8 | 7 | 1.06 |
| P30566 | ADSL | 7 | 7 | 1.06 |
| Q9H9G7 | AGO3 | 3 | 1 | 1.06 |
| P09601 | HMOX1 | 13 | 13 | 1.06 |
| P07947 | YES1 | 7 | 3 | 1.06 |
| Q9UL18 | AGO1 | 2 | 1 | 1.06 |
| Q04637 | EIF4G1 | 36 | 32 | 1.06 |
| P20933 | AGA | 4 | 4 | 1.06 |
| Q9UHL4 | DPP7 | 9 | 9 | 1.06 |
| O00625 | PIR | 2 | 2 | 1.06 |
| Q96ST3 | SIN3A | 2 | 2 | 1.06 |
| O60828 | PQBP1 | 4 | 4 | 1.06 |

| Accession | Name | # Peptides | # Unique Peptides | Abundance Ratio: WWP2 / control |
|---|---|---|---|---|
| Q9NVZ3 | NECAP2 | 6 | 6 | 1.06 |
| O60716 | CTNND1 | 16 | 16 | 1.06 |
| Q9NX46 | ADPRS | 10 | 10 | 1.06 |
| Q14139 | UBE4A | 4 | 4 | 1.06 |
| A0A024RBG1 | NUDT4B | 4 | 2 | 1.06 |
| Q9BY77 | POLDIP3 | 3 | 3 | 1.06 |
| Q9HB07 | MYG1 | 5 | 5 | 1.06 |
| Q9H5N1 | RABEP2 | 7 | 7 | 1.06 |
| Q96C86 | DCPS | 7 | 7 | 1.06 |
| Q96HY6 | DDRGK1 | 4 | 4 | 1.06 |
| Q9NWB6 | ARGLU1 | 7 | 7 | 1.06 |
| Q9BY43 | CHMP4A | 4 | 4 | 1.06 |
| Q9UKX5 | ITGA11 | 9 | 9 | 1.06 |
| P17655 | CAPN2 | 19 | 18 | 1.06 |
| P61966 | AP1S1 | 2 | 2 | 1.06 |
| O95487 | SEC24B | 8 | 6 | 1.06 |
| O15083 | ERC2 | 4 | 4 | 1.06 |
| Q93050 | ATP6V0A1 | 5 | 5 | 1.06 |
| P52758 | RIDA | 3 | 3 | 1.06 |
| P30043 | BLVRB | 5 | 5 | 1.06 |
| Q99567 | NUP88 | 5 | 5 | 1.06 |
| P38606 | ATP6V1A | 16 | 16 | 1.06 |
| Q9H2M9 | RAB3GAP2 | 16 | 16 | 1.06 |
| P53999 | SUB1 | 8 | 8 | 1.06 |
| O14745 | SLC9A3R1 | 17 | 17 | 1.06 |
| Q9Y295 | DRG1 | 5 | 4 | 1.06 |
| P49959 | MRE11 | 5 | 5 | 1.06 |
| Q13618 | CUL3 | 10 | 10 | 1.06 |
| O60568 | PLOD3 | 21 | 21 | 1.06 |
| O94925 | GLS | 8 | 8 | 1.06 |
| Q96DG6 | CMBL | 8 | 8 | 1.06 |
| Q96HC4 | PDLIM5 | 24 | 24 | 1.06 |
| P13473 | LAMP2 | 3 | 3 | 1.06 |
| O75477 | ERLIN1 | 12 | 8 | 1.06 |
| Q92882 | OSTF1 | 6 | 6 | 1.06 |
| P10620 | MGST1 | 5 | 5 | 1.06 |
| P09525 | ANXA4 | 21 | 21 | 1.06 |
| P30536 | TSPO | 2 | 2 | 1.06 |
| Q9H910 | JPT2 | 5 | 5 | 1.06 |
| Q9BQE5 | APOL2 | 8 | 8 | 1.06 |
| Q6P2Q9 | PRPF8 | 43 | 43 | 1.06 |
| Q9BTE3 | MCMBP | 4 | 4 | 1.06 |
| P22694 | PRKACB | 8 | 3 | 1.06 |
| P52895 | AKR1C2 | 14 | 6 | 1.06 |

| Accession | Name | # Peptides | # Unique Peptides | Abundance Ratio: WWP2 / control |
|---|---|---|---|---|
| Q16513 | PKN2 | 4 | 4 | 1.06 |
| P33908 | MAN1A1 | 4 | 4 | 1.06 |
| Q86WV6 | STING1 | 5 | 5 | 1.06 |
| P42696 | RBM34 | 2 | 2 | 1.06 |
| P30049 | ATP5F1D | 3 | 3 | 1.06 |
| Q13459 | MYO9B | 10 | 10 | 1.06 |
| Q9HD42 | CHMP1A | 3 | 3 | 1.06 |
| Q92552 | MRPS27 | 4 | 4 | 1.06 |
| P09972 | ALDOC | 23 | 17 | 1.06 |
| Q96RF0 | SNX18 | 9 | 9 | 1.06 |
| P15289 | ARSA | 7 | 6 | 1.06 |
| P13798 | APEH | 13 | 13 | 1.06 |
| P63165 | SUMO1 | 4 | 4 | 1.06 |
| Q6DN90 | IQSEC1 | 2 | 2 | 1.06 |
| P78352 | DLG4 | 2 | 1 | 1.06 |
| A0AVT1 | UBA6 | 13 | 13 | 1.05 |
| Q01804 | OTUD4 | 2 | 2 | 1.05 |
| Q00059 | TFAM | 3 | 3 | 1.05 |
| P29803 | PDHA2 | 3 | 3 | 1.05 |
| P50552 | VASP | 13 | 13 | 1.05 |
| Q13418 | ILK | 13 | 13 | 1.05 |
| Q8TCT9 | HM13 | 8 | 8 | 1.05 |
| Q9UK61 | TASOR | 3 | 3 | 1.05 |
| Q9UH65 | SWAP70 | 12 | 12 | 1.05 |
| Q9H444 | CHMP4B | 7 | 7 | 1.05 |
| Q92575 | UBXN4 | 5 | 5 | 1.05 |
| Q7Z478 | DHX29 | 3 | 3 | 1.05 |
| Q15020 | SART3 | 14 | 14 | 1.05 |
| Q15599 | SLC9A3R2 | 2 | 2 | 1.05 |
| Q5VZM2 | RRAGB | 4 | 4 | 1.05 |
| Q7L523 | RRAGA | 4 | 4 | 1.05 |
| P54840 | GYS2 | 2 | 1 | 1.05 |
| Q16790 | CA9 | 6 | 6 | 1.05 |
| Q9UQE7 | SMC3 | 24 | 24 | 1.05 |
| P40123 | CAP2 | 6 | 5 | 1.05 |
| Q99747 | NAPG | 7 | 7 | 1.05 |
| P30040 | ERP29 | 9 | 9 | 1.05 |
| Q9NR30 | DDX21 | 4 | 4 | 1.05 |
| Q6ZSR9 | | 4 | 4 | 1.05 |
| Q9UHY1 | NRBP1 | 3 | 3 | 1.05 |
| Q9H1I8 | ASCC2 | 3 | 3 | 1.05 |
| Q9NQ55 | PPAN | 3 | 3 | 1.05 |
| Q96CS3 | FAF2 | 6 | 6 | 1.05 |
| Q9Y2V7 | COG6 | 2 | 2 | 1.05 |

| Accession | Name | # Peptides | # Unique Peptides | Abundance Ratio: WWP2 / control |
|---|---|---|---|---|
| Q86TI2 | DPP9 | 6 | 6 | 1.05 |
| P51798 | CLCN7 | 2 | 2 | 1.05 |
| Q6UXN9 | WDR82 | 5 | 5 | 1.05 |
| Q9UNF0 | PACSIN2 | 11 | 11 | 1.05 |
| O95379 | TNFAIP8 | 2 | 2 | 1.05 |
| Q86W92 | PPFIBP1 | 21 | 19 | 1.05 |
| P55899 | FCGRT | 4 | 4 | 1.05 |
| P49746 | THBS3 | 9 | 9 | 1.05 |
| O14498 | ISLR | 6 | 6 | 1.05 |
| O14880 | MGST3 | 3 | 3 | 1.05 |
| Q9BUL8 | PDCD10 | 7 | 7 | 1.05 |
| P07437 | TUBB | 24 | 2 | 1.05 |
| O75157 | TSC22D2 | 6 | 4 | 1.05 |
| Q9Y490 | TLN1 | 122 | 102 | 1.05 |
| P54252 | ATXN3 | 3 | 3 | 1.05 |
| Q96EY1 | DNAJA3 | 3 | 3 | 1.05 |
| P80297 | MT1X | 4 | 1 | 1.05 |
| P21926 | CD9 | 6 | 6 | 1.05 |
| Q01813 | PFKP | 30 | 24 | 1.05 |
| Q15404 | RSU1 | 7 | 7 | 1.05 |
| Q96SZ5 | ADO | 2 | 2 | 1.05 |
| Q9BXF6 | RAB11FIP5 | 5 | 5 | 1.05 |
| Q9Y5X1 | SNX9 | 11 | 11 | 1.05 |
| Q9UPU5 | USP24 | 8 | 8 | 1.05 |
| Q9UGV2 | NDRG3 | 2 | 2 | 1.05 |
| P27708 | CAD | 24 | 24 | 1.05 |
| Q9UDY2 | TJP2 | 2 | 2 | 1.05 |
| Q7KZ85 | SUPT6H | 3 | 3 | 1.05 |
| Q9Y3X0 | CCDC9 | 2 | 2 | 1.05 |
| P10768 | ESD | 10 | 10 | 1.05 |
| Q9UKT5 | FBXO4 | 2 | 2 | 1.05 |
| Q13177 | PAK2 | 11 | 11 | 1.05 |
| Q9UBS8 | RNF14 | 2 | 2 | 1.05 |
| Q9H0R4 | HDHD2 | 2 | 2 | 1.05 |
| Q96CT7 | CCDC124 | 8 | 8 | 1.05 |
| B7ZW38 | HNRNPCL3 | 7 | 1 | 1.05 |
| P36969 | GPX4 | 3 | 3 | 1.05 |
| O94966 | USP19 | 2 | 2 | 1.05 |
| Q8WX93 | PALLD | 14 | 14 | 1.05 |
| Q9UBV8 | PEF1 | 4 | 4 | 1.05 |
| Q9H9S4 | CAB39L | 2 | 1 | 1.05 |
| Q13885 | TUBB2A | 22 | 1 | 1.05 |
| Q9NYL9 | TMOD3 | 10 | 9 | 1.05 |
| O60812 | HNRNPCL1 | 8 | 1 | 1.05 |

| Accession | Name | # Peptides | # Unique Peptides | Abundance Ratio: WWP2 / control |
|---|---|---|---|---|
| P43155 | CRAT | 4 | 4 | 1.05 |
| Q9HB40 | SCPEP1 | 5 | 5 | 1.05 |
| Q00653 | NFKB2 | 6 | 5 | 1.05 |
| A0A0G2JPF8 | HNRNPCL4 | 7 | 1 | 1.05 |
| Q9Y2W2 | WBP11 | 6 | 6 | 1.05 |
| Q9H267 | VPS33B | 3 | 3 | 1.05 |
| Q9H3P7 | ACBD3 | 14 | 14 | 1.05 |
| Q9NX05 | FAM120C | 2 | 2 | 1.05 |
| Q03135 | CAV1 | 9 | 9 | 1.05 |
| P51116 | FXR2 | 7 | 3 | 1.05 |
| P09496 | CLTA | 8 | 8 | 1.05 |
| P61586 | RHOA | 9 | 3 | 1.05 |
| P51151 | RAB9A | 5 | 5 | 1.05 |
| Q9HC35 | EML4 | 3 | 3 | 1.05 |
| P28072 | PSMB6 | 5 | 5 | 1.05 |
| P40121 | CAPG | 14 | 14 | 1.05 |
| P49585 | PCYT1A | 12 | 12 | 1.05 |
| O76003 | GLRX3 | 8 | 8 | 1.05 |
| Q01459 | CTBS | 3 | 3 | 1.05 |
| Q9HA64 | FN3KRP | 7 | 7 | 1.05 |
| P61803 | DAD1 | 4 | 4 | 1.05 |
| Q9Y5V3 | MAGED1 | 4 | 3 | 1.05 |
| Q13573 | SNW1 | 5 | 5 | 1.05 |
| Q9H074 | PAIP1 | 5 | 5 | 1.05 |
| Q9HCD5 | NCOA5 | 4 | 4 | 1.05 |
| O00423 | EML1 | 9 | 9 | 1.05 |
| Q9NUU7 | DDX19A | 9 | 9 | 1.05 |
| P09497 | CLTB | 9 | 9 | 1.05 |
| Q6NW29 | RWDD4 | 2 | 2 | 1.05 |
| P40222 | TXLNA | 14 | 14 | 1.05 |
| P30876 | POLR2B | 10 | 10 | 1.05 |
| Q99848 | EBNA1BP2 | 6 | 6 | 1.05 |
| Q9UMS4 | PRPF19 | 9 | 9 | 1.05 |
| Q96A65 | EXOC4 | 11 | 11 | 1.05 |
| Q8N7H5 | PAF1 | 2 | 2 | 1.05 |
| Q14376 | GALE | 6 | 6 | 1.05 |
| Q9UMR2 | DDX19B | 9 | 9 | 1.05 |
| Q02218 | OGDH | 15 | 8 | 1.05 |
| Q08722 | CD47 | 4 | 4 | 1.05 |
| Q9NV70 | EXOC1 | 4 | 4 | 1.05 |
| P32455 | GBP1 | 8 | 7 | 1.05 |
| Q6NZI2 | CAVIN1 | 22 | 22 | 1.05 |
| Q13564 | NAE1 | 8 | 8 | 1.05 |
| Q9H6X2 | ANTXR1 | 2 | 2 | 1.05 |

| Accession | Name | # Peptides | # Unique Peptides | Abundance Ratio: WWP2 / control |
|---|---|---|---|---|
| P39748 | FEN1 | 5 | 5 | 1.05 |
| O60518 | RANBP6 | 2 | 1 | 1.05 |
| Q32MK0 | MYLK3 | 2 | 2 | 1.05 |
| Q5TCZ1 | SH3PXD2A | 3 | 2 | 1.05 |
| P51610 | HCFC1 | 10 | 10 | 1.05 |
| P04632 | CAPNS1 | 6 | 6 | 1.05 |
| Q96T51 | RUFY1 | 9 | 9 | 1.05 |
| P52564 | MAP2K6 | 3 | 3 | 1.05 |
| Q562R1 | ACTBL2 | 8 | 3 | 1.05 |
| P54725 | RAD23A | 5 | 3 | 1.05 |
| Q99805 | TM9SF2 | 6 | 6 | 1.05 |
| Q99733 | NAP1L4 | 9 | 7 | 1.05 |
| Q9Y5L4 | TIMM13 | 2 | 2 | 1.05 |
| Q66K74 | MAP1S | 5 | 5 | 1.05 |
| Q9BXW6 | OSBPL1A | 2 | 2 | 1.05 |
| Q9H269 | VPS16 | 5 | 5 | 1.05 |
| P08559 | PDHA1 | 12 | 12 | 1.05 |
| P21281 | ATP6V1B2 | 14 | 14 | 1.05 |
| Q12765 | SCRN1 | 8 | 8 | 1.05 |
| Q6UVY6 | MOXD1 | 2 | 2 | 1.05 |
| Q9H3U1 | UNC45A | 11 | 11 | 1.05 |
| Q9BQ67 | GRWD1 | 3 | 3 | 1.05 |
| Q9ULH1 | ASAP1 | 3 | 3 | 1.05 |
| Q6IPX3 | TCEAL6 | 3 | 3 | 1.05 |
| P53582 | METAP1 | 4 | 4 | 1.05 |
| P21291 | CSRP1 | 7 | 7 | 1.05 |
| Q5H9L2 | TCEAL5 | 4 | 4 | 1.05 |
| Q6PCB0 | VWA1 | 4 | 4 | 1.05 |
| P04080 | CSTB | 7 | 7 | 1.04 |
| Q09161 | NCBP1 | 8 | 8 | 1.04 |
| O43586 | PSTPIP1 | 2 | 1 | 1.04 |
| Q9Y3D0 | CIAO2B | 3 | 3 | 1.04 |
| Q8TB61 | SLC35B2 | 3 | 3 | 1.04 |
| Q15477 | SKIV2L | 6 | 5 | 1.04 |
| Q96L46 | CAPNS2 | 2 | 2 | 1.04 |
| Q14839 | CHD4 | 11 | 8 | 1.04 |
| P09110 | ACAA1 | 6 | 6 | 1.04 |
| Q13442 | PDAP1 | 11 | 11 | 1.04 |
| Q63HR2 | TNS2 | 3 | 1 | 1.04 |
| P52789 | HK2 | 12 | 8 | 1.04 |
| Q15746 | MYLK | 6 | 6 | 1.04 |
| P05026 | ATP1B1 | 11 | 11 | 1.04 |
| Q8WUF5 | PPP1R13L | 2 | 2 | 1.04 |
| P43487 | RANBP1 | 9 | 9 | 1.04 |

| Accession | Name | # Peptides | # Unique Peptides | Abundance Ratio: WWP2 / control |
|---|---|---|---|---|
| Q14232 | EIF2B1 | 6 | 6 | 1.04 |
| Q6GMV3 | PTRHD1 | 3 | 3 | 1.04 |
| Q9HAU6 | TPT1P8 | 2 | 2 | 1.04 |
| Q9UNN5 | FAF1 | 6 | 6 | 1.04 |
| Q8N6H7 | ARFGAP2 | 6 | 6 | 1.04 |
| Q96H20 | SNF8 | 3 | 3 | 1.04 |
| Q6UXV4 | APOOL | 3 | 3 | 1.04 |
| Q9UHD1 | CHORDC1 | 6 | 6 | 1.04 |
| O75828 | CBR3 | 6 | 4 | 1.04 |
| Q9BRA2 | TXNDC17 | 4 | 4 | 1.04 |
| P98175 | RBM10 | 7 | 7 | 1.04 |
| P43686 | PSMC4 | 17 | 16 | 1.04 |
| P06132 | UROD | 5 | 5 | 1.04 |
| Q99808 | SLC29A1 | 5 | 5 | 1.04 |
| Q9HBH5 | RDH14 | 4 | 4 | 1.04 |
| P35270 | SPR | 6 | 6 | 1.04 |
| Q14767 | LTBP2 | 22 | 22 | 1.04 |
| P51659 | HSD17B4 | 31 | 31 | 1.04 |
| Q04206 | RELA | 7 | 7 | 1.04 |
| P05771 | PRKCB | 6 | 5 | 1.04 |
| Q9NR56 | MBNL1 | 9 | 9 | 1.04 |
| Q13164 | MAPK7 | 4 | 4 | 1.04 |
| Q9Y6A4 | CFAP20 | 3 | 3 | 1.04 |
| P53992 | SEC24C | 11 | 11 | 1.04 |
| Q9BV38 | WDR18 | 5 | 5 | 1.04 |
| Q29RF7 | PDS5A | 3 | 3 | 1.04 |
| Q9HC38 | GLOD4 | 9 | 9 | 1.04 |
| Q6FI81 | CIAPIN1 | 5 | 5 | 1.04 |
| P49327 | FASN | 49 | 49 | 1.04 |
| Q92542 | NCSTN | 2 | 2 | 1.04 |
| Q86V48 | LUZP1 | 6 | 6 | 1.04 |
| P08842 | STS | 6 | 6 | 1.04 |
| O00178 | GTPBP1 | 2 | 2 | 1.04 |
| Q8WW12 | PCNP | 2 | 2 | 1.04 |
| Q9BV57 | ADI1 | 4 | 4 | 1.04 |
| Q96ME7 | ZNF512 | 3 | 3 | 1.04 |
| Q9UFN0 | NIPSNAP3A | 3 | 3 | 1.04 |
| Q99653 | CHP1 | 6 | 6 | 1.04 |
| P34913 | EPHX2 | 2 | 2 | 1.04 |
| Q709C8 | VPS13C | 6 | 6 | 1.04 |
| P13716 | ALAD | 6 | 6 | 1.04 |
| P54619 | PRKAG1 | 4 | 4 | 1.04 |
| Q8WXE0 | CASKIN2 | 3 | 2 | 1.04 |
| P39059 | COL15A1 | 2 | 2 | 1.04 |

| Accession | Name | # Peptides | # Unique Peptides | Abundance Ratio: WWP2 / control |
|---|---|---|---|---|
| Q92783 | STAM | 4 | 3 | 1.04 |
| Q8TEX9 | IPO4 | 5 | 5 | 1.04 |
| Q9H7Z7 | PTGES2 | 4 | 4 | 1.04 |
| O95571 | ETHE1 | 3 | 3 | 1.04 |
| Q8WZA9 | IRGQ | 6 | 6 | 1.04 |
| Q99961 | SH3GL1 | 12 | 12 | 1.04 |
| Q05086 | UBE3A | 4 | 4 | 1.04 |
| Q13131 | PRKAA1 | 9 | 9 | 1.04 |
| Q2M389 | WASHC4 | 5 | 5 | 1.04 |
| O95865 | DDAH2 | 9 | 8 | 1.04 |
| P00492 | HPRT1 | 3 | 3 | 1.04 |
| P26440 | IVD | 5 | 5 | 1.04 |
| Q9NYF8 | BCLAF1 | 17 | 16 | 1.04 |
| Q8WUM4 | PDCD6IP | 41 | 41 | 1.04 |
| Q9H7B2 | RPF2 | 2 | 2 | 1.04 |
| P68400 | CSNK2A1 | 6 | 5 | 1.04 |
| P50148 | GNAQ | 7 | 4 | 1.04 |
| P21589 | NT5E | 25 | 25 | 1.04 |
| Q15435 | PPP1R7 | 16 | 16 | 1.04 |
| O95433 | AHSA1 | 8 | 8 | 1.04 |
| O14672 | ADAM10 | 3 | 3 | 1.04 |
| Q6UY14 | ADAMTSL4 | 6 | 6 | 1.04 |
| Q9UKM9 | RALY | 13 | 13 | 1.04 |
| P07858 | CTSB | 11 | 11 | 1.04 |
| P07942 | LAMB1 | 38 | 38 | 1.04 |
| Q8NE86 | MCU | 6 | 6 | 1.04 |
| Q9UPQ0 | LIMCH1 | 2 | 2 | 1.04 |
| Q9NW15 | ANO10 | 5 | 5 | 1.04 |
| Q13011 | ECH1 | 8 | 8 | 1.04 |
| Q5JTV8 | TOR1AIP1 | 18 | 17 | 1.04 |
| P22314 | UBA1 | 35 | 35 | 1.04 |
| Q86VS8 | HOOK3 | 10 | 10 | 1.04 |
| Q96LD4 | TRIM47 | 8 | 8 | 1.04 |
| P02795 | MT2A | 4 | 1 | 1.04 |
| P53675 | CLTCL1 | 12 | 12 | 1.04 |
| Q8WWM9 | CYGB | 2 | 2 | 1.04 |
| Q00005 | PPP2R2B | 2 | 2 | 1.04 |
| Q13325 | IFIT5 | 3 | 2 | 1.04 |
| Q96AT9 | RPE | 2 | 2 | 1.04 |
| O76070 | SNCG | 7 | 7 | 1.04 |
| Q13363 | CTBP1 | 8 | 4 | 1.04 |
| O00429 | DNM1L | 14 | 14 | 1.04 |
| O43294 | TGFB1I1 | 5 | 5 | 1.04 |
| Q86YP4 | GATAD2A | 2 | 1 | 1.04 |

| Accession | Name | # Peptides | # Unique Peptides | Abundance Ratio: WWP2 / control |
|---|---|---|---|---|
| Q13423 | NNT | 23 | 23 | 1.04 |
| P12236 | SLC25A6 | 14 | 2 | 1.04 |
| Q2M2I8 | AAK1 | 8 | 8 | 1.04 |
| Q10472 | GALNT1 | 5 | 5 | 1.04 |
| Q96EE3 | SEH1L | 7 | 7 | 1.04 |
| Q9Y2H0 | DLGAP4 | 6 | 6 | 1.04 |
| Q8NCW5 | NAXE | 3 | 3 | 1.04 |
| P09104 | ENO2 | 11 | 6 | 1.04 |
| Q96C90 | PPP1R14B | 3 | 3 | 1.04 |
| Q13424 | SNTA1 | 2 | 1 | 1.04 |
| Q9NS69 | TOMM22 | 3 | 3 | 1.04 |
| Q96IU4 | ABHD14B | 8 | 8 | 1.04 |
| Q9NQL2 | RRAGD | 3 | 3 | 1.04 |
| O95630 | STAMBP | 6 | 6 | 1.04 |
| Q9GZT8 | NIF3L1 | 4 | 4 | 1.04 |
| O43747 | AP1G1 | 16 | 16 | 1.04 |
| Q16222 | UAP1 | 21 | 19 | 1.04 |
| Q6NYC8 | PPP1R18 | 11 | 11 | 1.04 |
| Q14258 | TRIM25 | 15 | 15 | 1.04 |
| P37802 | TAGLN2 | 13 | 13 | 1.04 |
| Q99541 | PLIN2 | 18 | 18 | 1.04 |
| P52294 | KPNA1 | 7 | 4 | 1.04 |
| Q9HCN8 | SDF2L1 | 2 | 2 | 1.04 |
| Q9NQ48 | LZTFL1 | 5 | 5 | 1.04 |
| P15586 | GNS | 9 | 9 | 1.04 |
| Q8IVF7 | FMNL3 | 6 | 2 | 1.04 |
| Q14914 | PTGR1 | 9 | 9 | 1.04 |
| P24539 | ATP5PB | 9 | 9 | 1.04 |
| P49792 | RANBP2 | 27 | 26 | 1.04 |
| P00390 | GSR | 12 | 12 | 1.04 |
| P08670 | VIM | 68 | 62 | 1.04 |
| Q8NE71 | ABCF1 | 15 | 14 | 1.04 |
| Q9HB90 | RRAGC | 4 | 4 | 1.04 |
| Q16204 | CCDC6 | 12 | 12 | 1.04 |
| Q9UQ35 | SRRM2 | 12 | 12 | 1.04 |
| P47712 | PLA2G4A | 7 | 7 | 1.04 |
| Q04917 | YWHAH | 17 | 13 | 1.04 |
| O75608 | LYPLA1 | 4 | 4 | 1.04 |
| Q6DKI1 | RPL7L1 | 3 | 3 | 1.04 |
| Q9H9A5 | CNOT10 | 2 | 2 | 1.04 |
| Q9H0D6 | XRN2 | 4 | 4 | 1.04 |
| O15031 | PLXNB2 | 13 | 13 | 1.04 |
| P21397 | MAOA | 9 | 7 | 1.04 |
| P63167 | DYNLL1 | 7 | 4 | 1.04 |

| Accession | Name | # Peptides | # Unique Peptides | Abundance Ratio: WWP2 / control |
|---|---|---|---|---|
| Q9Y5Z4 | HEBP2 | 7 | 7 | 1.04 |
| Q8NBF2 | NHLRC2 | 4 | 4 | 1.04 |
| Q9Y3D6 | FIS1 | 4 | 4 | 1.04 |
| Q9UNZ5 | C19orf53 | 2 | 2 | 1.04 |
| Q9BQE3 | TUBA1C | 22 | 4 | 1.04 |
| P33176 | KIF5B | 36 | 36 | 1.04 |
| P41219 | PRPH | 5 | 3 | 1.04 |
| Q15785 | TOMM34 | 3 | 3 | 1.04 |
| O00267 | SUPT5H | 5 | 5 | 1.04 |
| Q8NC44 | RETREG2 | 2 | 2 | 1.04 |
| O00483 | NDUFA4 | 2 | 2 | 1.04 |
| Q9BVL2 | NUP58 | 4 | 4 | 1.04 |
| Q9UJV8 | PURG | 2 | 1 | 1.04 |
| P16157 | ANK1 | 2 | 1 | 1.04 |
| O00541 | PES1 | 6 | 6 | 1.04 |
| O60869 | EDF1 | 8 | 8 | 1.04 |
| P48637 | GSS | 18 | 18 | 1.04 |
| P54289 | CACNA2D1 | 4 | 4 | 1.04 |
| Q9UPN3 | MACF1 | 48 | 46 | 1.04 |
| Q5J8M3 | EMC4 | 3 | 3 | 1.04 |
| Q96I99 | SUCLG2 | 12 | 12 | 1.04 |
| Q9HB71 | CACYBP | 12 | 12 | 1.04 |
| P18846 | ATF1 | 2 | 2 | 1.04 |
| Q3KQV9 | UAP1L1 | 6 | 4 | 1.04 |
| Q9BU23 | LMF2 | 7 | 7 | 1.04 |
| Q8WTX7 | CASTOR1 | 2 | 2 | 1.04 |
| P42224 | STAT1 | 25 | 25 | 1.04 |
| Q14192 | FHL2 | 10 | 10 | 1.04 |
| P63104 | YWHAZ | 23 | 17 | 1.04 |
| Q7L5N1 | COPS6 | 3 | 3 | 1.04 |
| Q9H223 | EHD4 | 9 | 5 | 1.04 |
| Q9Y680 | FKBP7 | 12 | 12 | 1.04 |
| P16220 | CREB1 | 2 | 2 | 1.04 |
| Q99704 | DOK1 | 2 | 2 | 1.04 |
| P22059 | OSBP | 10 | 10 | 1.04 |
| Q92917 | GPKOW | 2 | 2 | 1.04 |
| Q9UJS0 | SLC25A13 | 4 | 4 | 1.04 |
| Q9Y6C2 | EMILIN1 | 38 | 38 | 1.04 |
| P00491 | PNP | 10 | 10 | 1.04 |
| P84095 | RHOG | 7 | 6 | 1.04 |
| Q96IP4 | TENT5A | 2 | 2 | 1.04 |
| O94826 | TOMM70 | 17 | 17 | 1.04 |
| P52434 | POLR2H | 4 | 4 | 1.04 |
| P62857 | RPS28 | 4 | 4 | 1.04 |

| Accession | Name | # Peptides | # Unique Peptides | Abundance Ratio: WWP2 / control |
|---|---|---|---|---|
| P04181 | OAT | 9 | 9 | 1.04 |
| P07954 | FH | 12 | 12 | 1.04 |
| Q92544 | TM9SF4 | 5 | 5 | 1.04 |
| Q92520 | FAM3C | 7 | 7 | 1.04 |
| Q96N66 | MBOAT7 | 5 | 5 | 1.04 |
| P08133 | ANXA6 | 50 | 50 | 1.04 |
| O60645 | EXOC3 | 7 | 7 | 1.04 |
| Q9BZE4 | GTPBP4 | 6 | 6 | 1.04 |
| Q13347 | EIF3I | 9 | 9 | 1.04 |
| Q15035 | TRAM2 | 2 | 2 | 1.04 |
| Q9P253 | VPS18 | 4 | 4 | 1.04 |
| P22102 | GART | 15 | 15 | 1.04 |
| P49593 | PPM1F | 7 | 7 | 1.03 |
| Q8IVL6 | P3H3 | 10 | 10 | 1.03 |
| Q02318 | CYP27A1 | 4 | 4 | 1.03 |
| P30622 | CLIP1 | 16 | 12 | 1.03 |
| Q12981 | BNIP1 | 3 | 3 | 1.03 |
| Q06323 | PSME1 | 13 | 13 | 1.03 |
| Q9Y2J2 | EPB41L3 | 10 | 7 | 1.03 |
| Q99715 | COL12A1 | 115 | 115 | 1.03 |
| Q15417 | CNN3 | 12 | 9 | 1.03 |
| Q969N2 | PIGT | 6 | 6 | 1.03 |
| Q05048 | CSTF1 | 3 | 3 | 1.03 |
| Q9H115 | NAPB | 2 | 2 | 1.03 |
| P48059 | LIMS1 | 11 | 8 | 1.03 |
| Q13126 | MTAP | 10 | 10 | 1.03 |
| Q9ULZ3 | PYCARD | 4 | 4 | 1.03 |
| Q8TB92 | HMGCLL1 | 2 | 2 | 1.03 |
| P07205 | PGK2 | 6 | 1 | 1.03 |
| O43772 | SLC25A20 | 2 | 2 | 1.03 |
| P27105 | STOM | 8 | 8 | 1.03 |
| Q9UHG3 | PCYOX1 | 17 | 17 | 1.03 |
| Q9BUK6 | MSTO1 | 2 | 2 | 1.03 |
| P00918 | CA2 | 6 | 6 | 1.03 |
| Q08397 | LOXL1 | 10 | 10 | 1.03 |
| Q969U7 | PSMG2 | 2 | 2 | 1.03 |
| O14776 | TCERG1 | 13 | 13 | 1.03 |
| O60739 | EIF1B | 2 | 2 | 1.03 |
| Q09028 | RBBP4 | 9 | 4 | 1.03 |
| Q13098 | GPS1 | 8 | 8 | 1.03 |
| Q8TC07 | TBC1D15 | 7 | 7 | 1.03 |
| P41567 | EIF1 | 2 | 2 | 1.03 |
| O43615 | TIMM44 | 9 | 9 | 1.03 |
| Q8WZA0 | LZIC | 6 | 6 | 1.03 |

| Accession | Name | # Peptides | # Unique Peptides | Abundance Ratio: WWP2 / control |
|---|---|---|---|---|
| Q8N2G8 | GHDC | 4 | 4 | 1.03 |
| Q9Y4G6 | TLN2 | 44 | 24 | 1.03 |
| Q15631 | TSN | 10 | 10 | 1.03 |
| P33240 | CSTF2 | 5 | 5 | 1.03 |
| P13489 | RNH1 | 19 | 19 | 1.03 |
| Q9NUQ6 | SPATS2L | 3 | 3 | 1.03 |
| Q9BTT0 | ANP32E | 3 | 3 | 1.03 |
| Q63ZY3 | KANK2 | 20 | 19 | 1.03 |
| Q567U6 | CCDC93 | 8 | 8 | 1.03 |
| Q969V3 | NCLN | 10 | 10 | 1.03 |
| Q9NUM4 | TMEM106B | 2 | 2 | 1.03 |
| P42025 | ACTR1B | 5 | 2 | 1.03 |
| Q9BT78 | COPS4 | 14 | 14 | 1.03 |
| P13637 | ATP1A3 | 18 | 18 | 1.03 |
| Q96EK7 | FAM120B | 3 | 3 | 1.03 |
| Q9BZK7 | TBL1XR1 | 4 | 4 | 1.03 |
| P35659 | DEK | 13 | 13 | 1.03 |
| Q9H832 | UBE2Z | 4 | 4 | 1.03 |
| Q96AB6 | NTAN1 | 2 | 2 | 1.03 |
| P54578 | USP14 | 18 | 18 | 1.03 |
| P10644 | PRKAR1A | 14 | 14 | 1.03 |
| O94875 | SORBS2 | 12 | 12 | 1.03 |
| Q15714 | TSC22D1 | 4 | 2 | 1.03 |
| P54802 | NAGLU | 7 | 7 | 1.03 |
| Q92626 | PXDN | 16 | 16 | 1.03 |
| Q13620 | CUL4B | 22 | 15 | 1.03 |
| Q9UIA9 | XPO7 | 6 | 6 | 1.03 |
| Q07666 | KHDRBS1 | 8 | 8 | 1.03 |
| P58215 | LOXL3 | 4 | 4 | 1.03 |
| Q9Y6G9 | DYNC1LI1 | 7 | 7 | 1.03 |
| P14618 | PKM | 44 | 44 | 1.03 |
| Q9NP72 | RAB18 | 6 | 6 | 1.03 |
| Q9BSJ8 | ESYT1 | 20 | 19 | 1.03 |
| Q14203 | DCTN1 | 34 | 34 | 1.03 |
| P00558 | PGK1 | 33 | 28 | 1.03 |
| Q13546 | RIPK1 | 2 | 2 | 1.03 |
| P24821 | TNC | 106 | 106 | 1.03 |
| P11413 | G6PD | 26 | 26 | 1.03 |
| P52888 | THOP1 | 13 | 13 | 1.03 |
| Q8TAT6 | NPLOC4 | 10 | 10 | 1.03 |
| Q7L576 | CYFIP1 | 17 | 17 | 1.03 |
| P62854 | RPS26 | 3 | 3 | 1.03 |
| Q96S52 | PIGS | 4 | 4 | 1.03 |
| P04406 | GAPDH | 26 | 26 | 1.03 |

| Accession | Name | # Peptides | # Unique Peptides | Abundance Ratio: WWP2 / control |
|---|---|---|---|---|
| A6NEC2 | NPEPPSL1 | 13 | 13 | 1.03 |
| P49756 | RBM25 | 9 | 9 | 1.03 |
| Q5JNZ5 | RPS26P11 | 3 | 3 | 1.03 |
| Q14155 | ARHGEF7 | 3 | 3 | 1.03 |
| Q14738 | PPP2R5D | 10 | 10 | 1.03 |
| O00748 | CES2 | 2 | 2 | 1.03 |
| Q5VIR6 | VPS53 | 2 | 2 | 1.03 |
| P21333 | FLNA | 144 | 128 | 1.03 |
| P78357 | CNTNAP1 | 2 | 2 | 1.03 |
| Q9UJU6 | DBNL | 14 | 14 | 1.03 |
| Q92614 | MYO18A | 7 | 7 | 1.03 |
| P06396 | GSN | 38 | 38 | 1.03 |
| Q14677 | CLINT1 | 9 | 9 | 1.03 |
| Q8WWM7 | ATXN2L | 15 | 13 | 1.03 |
| P06703 | S100A6 | 10 | 10 | 1.03 |
| P55786 | NPEPPS | 29 | 29 | 1.03 |
| Q9Y3E7 | CHMP3 | 3 | 3 | 1.03 |
| Q9UJZ1 | STOML2 | 8 | 8 | 1.03 |
| Q9NVD7 | PARVA | 9 | 9 | 1.03 |
| P22105 | TNXB | 63 | 63 | 1.03 |
| P36405 | ARL3 | 4 | 4 | 1.03 |
| Q9UL15 | BAG5 | 3 | 3 | 1.03 |
| P18583 | SON | 6 | 6 | 1.03 |
| Q02880 | TOP2B | 7 | 7 | 1.03 |
| P20073 | ANXA7 | 18 | 18 | 1.03 |
| Q15059 | BRD3 | 3 | 3 | 1.03 |
| O43493 | TGOLN2 | 6 | 6 | 1.03 |
| O15067 | PFAS | 6 | 6 | 1.03 |
| Q9BTW9 | TBCD | 10 | 10 | 1.03 |
| Q15126 | PMVK | 6 | 6 | 1.03 |
| O95777 | LSM8 | 3 | 3 | 1.03 |
| P19525 | EIF2AK2 | 7 | 7 | 1.03 |
| Q16630 | CPSF6 | 12 | 12 | 1.03 |
| Q5JRX3 | PITRM1 | 22 | 22 | 1.03 |
| Q9NVE7 | PANK4 | 3 | 3 | 1.03 |
| P18615 | NELFE | 2 | 2 | 1.03 |
| Q6DKJ4 | NXN | 6 | 6 | 1.03 |
| P49720 | PSMB3 | 4 | 4 | 1.03 |
| P31321 | PRKAR1B | 4 | 4 | 1.03 |
| P20618 | PSMB1 | 9 | 9 | 1.03 |
| Q08J23 | NSUN2 | 5 | 5 | 1.03 |
| Q8NFW8 | CMAS | 7 | 7 | 1.03 |
| P55072 | VCP | 45 | 45 | 1.03 |
| Q9NX08 | COMMD8 | 2 | 2 | 1.03 |

| Accession | Name | # Peptides | # Unique Peptides | Abundance Ratio: WWP2 / control |
|---|---|---|---|---|
| Q9HD26 | GOPC | 12 | 12 | 1.03 |
| Q9UJ70 | NAGK | 13 | 13 | 1.03 |
| Q10570 | CPSF1 | 3 | 3 | 1.03 |
| Q9Y266 | NUDC | 12 | 12 | 1.03 |
| Q13523 | PRPF4B | 2 | 2 | 1.03 |
| O15357 | INPPL1 | 3 | 3 | 1.03 |
| P54687 | BCAT1 | 7 | 7 | 1.03 |
| P19634 | SLC9A1 | 2 | 2 | 1.03 |
| P63208 | SKP1 | 3 | 3 | 1.03 |
| P56211 | ARPP19 | 4 | 3 | 1.03 |
| P55039 | DRG2 | 9 | 8 | 1.03 |
| Q9NRH3 | TUBG2 | 3 | 3 | 1.03 |
| Q9UL46 | PSME2 | 8 | 8 | 1.03 |
| Q8NEV1 | CSNK2A3 | 5 | 4 | 1.03 |
| Q9NR31 | SAR1A | 6 | 3 | 1.03 |
| P11498 | PC | 10 | 10 | 1.03 |
| Q15437 | SEC23B | 3 | 1 | 1.03 |
| O14617 | AP3D1 | 15 | 15 | 1.03 |
| Q9UBT2 | UBA2 | 11 | 11 | 1.03 |
| Q9BPX5 | ARPC5L | 4 | 4 | 1.03 |
| Q8IUD2 | ERC1 | 15 | 15 | 1.03 |
| Q6PGP7 | TTC37 | 9 | 9 | 1.03 |
| Q92538 | GBF1 | 13 | 13 | 1.03 |
| O60664 | PLIN3 | 20 | 20 | 1.03 |
| Q92616 | GCN1 | 51 | 51 | 1.03 |
| Q9H2D6 | TRIOBP | 13 | 13 | 1.03 |
| P13987 | CD59 | 4 | 4 | 1.03 |
| Q9Y2D4 | EXOC6B | 4 | 4 | 1.03 |
| O14980 | XPO1 | 21 | 21 | 1.03 |
| O00231 | PSMD11 | 16 | 16 | 1.03 |
| P15170 | GSPT1 | 13 | 6 | 1.03 |
| P23258 | TUBG1 | 3 | 3 | 1.03 |
| P61981 | YWHAG | 18 | 12 | 1.03 |
| P31040 | SDHA | 12 | 12 | 1.03 |
| P42229 | STAT5A | 3 | 3 | 1.03 |
| Q9P0K7 | RAI14 | 13 | 13 | 1.03 |
| Q6NUK1 | SLC25A24 | 9 | 9 | 1.03 |
| Q9Y5K8 | ATP6V1D | 4 | 4 | 1.03 |
| Q16891 | IMMT | 33 | 33 | 1.03 |
| O00170 | AIP | 4 | 4 | 1.03 |
| Q92609 | TBC1D5 | 4 | 4 | 1.03 |
| Q6ZVM7 | TOM1L2 | 2 | 2 | 1.03 |
| Q8TF68 | ZNF384 | 2 | 2 | 1.03 |
| O14531 | DPYSL4 | 2 | 2 | 1.03 |

| Accession | Name | # Peptides | # Unique Peptides | Abundance Ratio: WWP2 / control |
|---|---|---|---|---|
| P51692 | STAT5B | 3 | 3 | 1.03 |
| Q9BRR6 | ADPGK | 3 | 3 | 1.03 |
| Q9HCE1 | MOV10 | 6 | 6 | 1.03 |
| P49448 | GLUD2 | 10 | 10 | 1.03 |
| O60826 | CCDC22 | 5 | 4 | 1.03 |
| Q9NSC5 | HOMER3 | 3 | 3 | 1.03 |
| Q01658 | DR1 | 3 | 3 | 1.03 |
| Q9Y284 | WDR83OS | 2 | 2 | 1.03 |
| Q8N4C8 | MINK1 | 3 | 1 | 1.03 |
| O43396 | TXNL1 | 11 | 11 | 1.03 |
| P48047 | ATP5PO | 8 | 8 | 1.03 |
| P05062 | ALDOB | 2 | 1 | 1.03 |
| O95721 | SNAP29 | 2 | 2 | 1.03 |
| Q14019 | COTL1 | 12 | 12 | 1.03 |
| P61081 | UBE2M | 10 | 10 | 1.03 |
| Q96K17 | BTF3L4 | 2 | 1 | 1.03 |
| Q9NWH9 | SLTM | 9 | 9 | 1.03 |
| O00161 | SNAP23 | 3 | 3 | 1.03 |
| P07305 | H1-0 | 11 | 11 | 1.03 |
| O75165 | DNAJC13 | 7 | 7 | 1.03 |
| P11908 | PRPS2 | 6 | 1 | 1.03 |
| O95747 | OXSR1 | 8 | 5 | 1.03 |
| P05023 | ATP1A1 | 41 | 41 | 1.03 |
| Q53TN4 | CYBRD1 | 3 | 3 | 1.03 |
| P30520 | ADSS2 | 11 | 10 | 1.03 |
| P50993 | ATP1A2 | 15 | 15 | 1.03 |
| P04439 | HLA-A | 14 | 10 | 1.02 |
| Q06830 | PRDX1 | 18 | 14 | 1.02 |
| P11021 | HSPA5 | 49 | 47 | 1.02 |
| Q15436 | SEC23A | 21 | 19 | 1.02 |
| Q13427 | PPIG | 3 | 3 | 1.02 |
| P62877 | RBX1 | 3 | 3 | 1.02 |
| P27361 | MAPK3 | 14 | 9 | 1.02 |
| Q9H0R5 | GBP3 | 3 | 3 | 1.02 |
| Q9NVS9 | PNPO | 5 | 5 | 1.02 |
| P46940 | IQGAP1 | 77 | 72 | 1.02 |
| Q13243 | SRSF5 | 2 | 1 | 1.02 |
| Q9NZB2 | FAM120A | 12 | 12 | 1.02 |
| Q9Y5A9 | YTHDF2 | 5 | 2 | 1.02 |
| Q92879 | CELF1 | 5 | 5 | 1.02 |
| P36269 | GGT5 | 4 | 4 | 1.02 |
| Q9NSD9 | FARSB | 15 | 15 | 1.02 |
| O14618 | CCS | 4 | 4 | 1.02 |
| Q8N684 | CPSF7 | 7 | 7 | 1.02 |

| Accession | Name | # Peptides | # Unique Peptides | Abundance Ratio: WWP2 / control |
|---|---|---|---|---|
| Q9H6R4 | NOL6 | 2 | 2 | 1.02 |
| O43684 | BUB3 | 8 | 8 | 1.02 |
| O95292 | VAPB | 11 | 10 | 1.02 |
| O76094 | SRP72 | 16 | 16 | 1.02 |
| Q12805 | EFEMP1 | 3 | 3 | 1.02 |
| Q9UHB9 | SRP68 | 13 | 13 | 1.02 |
| P63027 | VAMP2 | 3 | 3 | 1.02 |
| P46459 | NSF | 27 | 26 | 1.02 |
| Q9Y4D7 | PLXND1 | 2 | 2 | 1.02 |
| P84101 | SERF2 | 4 | 4 | 1.02 |
| P82909 | MRPS36 | 3 | 3 | 1.02 |
| Q16762 | TST | 5 | 5 | 1.02 |
| O00193 | SMAP | 3 | 3 | 1.02 |
| Q99880 | H2BC13 | 10 | 1 | 1.02 |
| P0DPI2 | GATD3A | 5 | 5 | 1.02 |
| O14818 | PSMA7 | 15 | 15 | 1.02 |
| P25325 | MPST | 3 | 3 | 1.02 |
| Q9Y4Z0 | LSM4 | 2 | 2 | 1.02 |
| P62316 | SNRPD2 | 8 | 8 | 1.02 |
| Q96K76 | USP47 | 8 | 8 | 1.02 |
| O00217 | NDUFS8 | 2 | 2 | 1.02 |
| A0A0B4J2D5 | GATD3B | 5 | 5 | 1.02 |
| Q8IUE6 | H2AC21 | 6 | 1 | 1.02 |
| Q96A08 | H2BC1 | 5 | 1 | 1.02 |
| Q07866 | KLC1 | 17 | 13 | 1.02 |
| P10619 | CTSA | 8 | 8 | 1.02 |
| P30533 | LRPAP1 | 21 | 21 | 1.02 |
| Q9Y2W1 | THRAP3 | 20 | 19 | 1.02 |
| P52948 | NUP98 | 8 | 8 | 1.02 |
| Q15024 | EXOSC7 | 3 | 3 | 1.02 |
| Q00796 | SORD | 9 | 9 | 1.02 |
| P63313 | TMSB10 | 4 | 2 | 1.02 |
| O15269 | SPTLC1 | 10 | 10 | 1.02 |
| P07902 | GALT | 3 | 3 | 1.02 |
| Q9UQ80 | PA2G4 | 18 | 18 | 1.02 |
| Q15907 | RAB11B | 11 | 11 | 1.02 |
| P12277 | CKB | 12 | 12 | 1.02 |
| O15258 | RER1 | 4 | 4 | 1.02 |
| Q969Q0 | RPL36AL | 6 | 2 | 1.02 |
| Q9P2R7 | SUCLA2 | 8 | 8 | 1.02 |
| Q9P289 | STK26 | 7 | 3 | 1.02 |
| P62491 | RAB11A | 10 | 10 | 1.02 |
| O95831 | AIFM1 | 9 | 9 | 1.02 |
| Q9H7C4 | SYNC | 4 | 4 | 1.02 |

| Accession | Name | # Peptides | # Unique Peptides | Abundance Ratio: WWP2 / control |
|---|---|---|---|---|
| Q99832 | CCT7 | 25 | 25 | 1.02 |
| Q969X1 | TMBIM1 | 2 | 2 | 1.02 |
| Q9NTI5 | PDS5B | 9 | 9 | 1.02 |
| Q8TEQ6 | GEMIN5 | 6 | 6 | 1.02 |
| P78371 | CCT2 | 27 | 27 | 1.02 |
| Q92747 | ARPC1A | 6 | 6 | 1.02 |
| Q7Z2Z2 | EFL1 | 2 | 2 | 1.02 |
| Q9BR76 | CORO1B | 13 | 13 | 1.02 |
| O60701 | UGDH | 18 | 18 | 1.02 |
| P62195 | PSMC5 | 15 | 14 | 1.02 |
| O95163 | ELP1 | 4 | 4 | 1.02 |
| P07093 | SERPINE2 | 9 | 9 | 1.02 |
| P36507 | MAP2K2 | 10 | 6 | 1.02 |
| Q9NVP1 | DDX18 | 9 | 9 | 1.02 |
| P21912 | SDHB | 8 | 8 | 1.02 |
| Q2TAA2 | IAH1 | 5 | 5 | 1.02 |
| P21964 | COMT | 9 | 9 | 1.02 |
| Q04837 | SSBP1 | 6 | 6 | 1.02 |
| Q5T6V5 | C9orf64 | 8 | 8 | 1.02 |
| O15511 | ARPC5 | 11 | 11 | 1.02 |
| Q92997 | DVL3 | 3 | 2 | 1.02 |
| Q969E4 | TCEAL3 | 6 | 6 | 1.02 |
| Q9UP95 | SLC12A4 | 6 | 6 | 1.02 |
| Q96D46 | NMD3 | 2 | 2 | 1.02 |
| P45880 | VDAC2 | 11 | 11 | 1.02 |
| Q9BUP3 | HTATIP2 | 5 | 5 | 1.02 |
| O60831 | PRAF2 | 3 | 3 | 1.02 |
| O00559 | EBAG9 | 2 | 2 | 1.02 |
| Q8N766 | EMC1 | 8 | 8 | 1.02 |
| Q96EK6 | GNPNAT1 | 8 | 8 | 1.02 |
| P31689 | DNAJA1 | 6 | 6 | 1.02 |
| P00352 | ALDH1A1 | 3 | 1 | 1.02 |
| P78356 | PIP4K2B | 3 | 2 | 1.02 |
| O43633 | CHMP2A | 9 | 9 | 1.02 |
| Q99759 | MAP3K3 | 2 | 1 | 1.02 |
| P25705 | ATP5F1A | 27 | 27 | 1.02 |
| Q13724 | MOGS | 16 | 16 | 1.02 |
| P12270 | TPR | 41 | 41 | 1.02 |
| P11387 | TOP1 | 14 | 14 | 1.02 |
| Q9NZL9 | MAT2B | 6 | 6 | 1.02 |
| Q9BU89 | DOHH | 2 | 2 | 1.02 |
| P61020 | RAB5B | 6 | 3 | 1.02 |
| P52630 | STAT2 | 5 | 5 | 1.02 |
| P48506 | GCLC | 7 | 7 | 1.02 |

| Accession | Name | # Peptides | # Unique Peptides | Abundance Ratio: WWP2 / control |
|---|---|---|---|---|
| Q96IJ6 | GMPPA | 9 | 9 | 1.02 |
| Q9Y2H1 | STK38L | 4 | 4 | 1.02 |
| Q9Y6M9 | NDUFB9 | 2 | 2 | 1.02 |
| Q8WYA6 | CTNNBL1 | 2 | 2 | 1.02 |
| Q9NY12 | GAR1 | 3 | 3 | 1.02 |
| Q99459 | CDC5L | 9 | 9 | 1.02 |
| Q92890 | UFD1 | 6 | 6 | 1.02 |
| Q9NQG5 | RPRD1B | 5 | 5 | 1.02 |
| Q9NVJ2 | ARL8B | 2 | 1 | 1.02 |
| Q32MZ4 | LRRFIP1 | 10 | 9 | 1.02 |
| Q9Y2Q3 | GSTK1 | 9 | 9 | 1.02 |
| Q04323 | UBXN1 | 8 | 8 | 1.02 |
| Q8TD47 | RPS4Y2 | 9 | 2 | 1.02 |
| P53007 | SLC25A1 | 6 | 6 | 1.02 |
| Q58FF6 | HSP90AB4P | 7 | 1 | 1.02 |
| P13693 | TPT1 | 9 | 9 | 1.02 |
| P50213 | IDH3A | 6 | 6 | 1.02 |
| Q9BZZ5 | API5 | 13 | 13 | 1.02 |
| P25786 | PSMA1 | 15 | 15 | 1.02 |
| Q8NF91 | SYNE1 | 18 | 18 | 1.02 |
| P45974 | USP5 | 17 | 17 | 1.02 |
| Q9Y646 | CPQ | 7 | 7 | 1.02 |
| O60749 | SNX2 | 21 | 20 | 1.02 |
| P34932 | HSPA4 | 43 | 41 | 1.02 |
| O75937 | DNAJC8 | 5 | 5 | 1.02 |
| P51452 | DUSP3 | 4 | 4 | 1.02 |
| Q4G0J3 | LARP7 | 6 | 6 | 1.02 |
| P16403 | H1-2 | 22 | 10 | 1.02 |
| O75821 | EIF3G | 9 | 9 | 1.02 |
| Q9NVA2 | SEPTIN11 | 16 | 9 | 1.02 |
| P41240 | CSK | 5 | 5 | 1.02 |
| Q6NVY1 | HIBCH | 6 | 6 | 1.02 |
| P50452 | SERPINB8 | 3 | 2 | 1.02 |
| Q8TDQ7 | GNPDA2 | 7 | 3 | 1.02 |
| O43681 | GET3 | 6 | 6 | 1.02 |
| Q9NR12 | PDLIM7 | 14 | 14 | 1.02 |
| Q13641 | TPBG | 2 | 2 | 1.02 |
| Q9Y618 | NCOR2 | 4 | 4 | 1.02 |
| Q9Y4X5 | ARIH1 | 2 | 2 | 1.02 |
| Q96C23 | GALM | 7 | 7 | 1.02 |
| Q15050 | RRS1 | 9 | 9 | 1.02 |
| O14936 | CASK | 6 | 6 | 1.02 |
| P09874 | PARP1 | 24 | 24 | 1.02 |
| Q8IZL8 | PELP1 | 6 | 6 | 1.02 |

| Accession | Name | # Peptides | # Unique Peptides | Abundance Ratio: WWP2 / control |
|---|---|---|---|---|
| Q9Y639 | NPTN | 10 | 10 | 1.02 |
| Q5JXB2 | UBE2NL | 6 | 6 | 1.02 |
| P35232 | PHB | 13 | 13 | 1.02 |
| Q99598 | TSNAX | 7 | 7 | 1.02 |
| P13995 | MTHFD2 | 3 | 3 | 1.02 |
| P00387 | CYB5R3 | 12 | 12 | 1.02 |
| P35221 | CTNNA1 | 17 | 17 | 1.02 |
| P27695 | APEX1 | 13 | 13 | 1.02 |
| Q4ZHG4 | FNDC1 | 4 | 4 | 1.02 |
| Q8N8U9 | BMPER | 2 | 2 | 1.02 |
| Q9UM00 | TMCO1 | 3 | 3 | 1.02 |
| O95202 | LETM1 | 8 | 8 | 1.02 |
| O14737 | PDCD5 | 11 | 11 | 1.02 |
| Q9H9B4 | SFXN1 | 9 | 9 | 1.02 |
| O14744 | PRMT5 | 8 | 8 | 1.02 |
| Q9UI14 | RABAC1 | 2 | 2 | 1.02 |
| Q9NP97 | DYNLRB1 | 3 | 3 | 1.02 |
| Q6DD88 | ATL3 | 15 | 15 | 1.02 |
| Q13867 | BLMH | 4 | 4 | 1.02 |
| Q15084 | PDIA6 | 15 | 15 | 1.02 |
| Q14152 | EIF3A | 45 | 45 | 1.02 |
| Q00169 | PITPNA | 7 | 5 | 1.02 |
| Q06124 | PTPN11 | 11 | 11 | 1.02 |
| Q15642 | TRIP10 | 7 | 7 | 1.02 |
| Q16531 | DDB1 | 33 | 33 | 1.02 |
| Q8TF09 | DYNLRB2 | 3 | 3 | 1.02 |
| Q9ULC4 | MCTS1 | 5 | 5 | 1.02 |
| P11310 | ACADM | 4 | 4 | 1.02 |
| Q9C0C2 | TNKS1BP1 | 33 | 33 | 1.02 |
| Q9UHQ9 | CYB5R1 | 10 | 10 | 1.02 |
| Q9NZ43 | USE1 | 3 | 3 | 1.02 |
| Q9UM54 | MYO6 | 20 | 20 | 1.02 |
| O60341 | KDM1A | 2 | 2 | 1.02 |
| Q5JTH9 | RRP12 | 4 | 4 | 1.02 |
| P08237 | PFKM | 22 | 19 | 1.02 |
| P19338 | NCL | 46 | 46 | 1.02 |
| P63173 | RPL38 | 4 | 4 | 1.02 |
| P33316 | DUT | 6 | 6 | 1.02 |
| A1L0T0 | ILVBL | 7 | 7 | 1.02 |
| A0A1B0GUL7 |  | 7 | 4 | 1.02 |
| P35527 | KRT9 | 11 | 10 | 1.02 |
| Q96HN2 | AHCYL2 | 8 | 8 | 1.02 |
| Q01130 | SRSF2 | 5 | 4 | 1.02 |
| P60983 | GMFB | 4 | 4 | 1.02 |

| Accession | Name | # Peptides | # Unique Peptides | Abundance Ratio: WWP2 / control |
|---|---|---|---|---|
| Q9ULT8 | HECTD1 | 7 | 7 | 1.02 |
| Q9UNH7 | SNX6 | 14 | 13 | 1.02 |
| P46109 | CRKL | 8 | 8 | 1.02 |
| Q9Y4P3 | TBL2 | 13 | 13 | 1.02 |
| Q92620 | DHX38 | 2 | 2 | 1.02 |
| P27816 | MAP4 | 62 | 62 | 1.02 |
| Q6ZMI0 | PPP1R21 | 3 | 3 | 1.02 |
| Q9UKK3 | PARP4 | 13 | 13 | 1.02 |
| P61923 | COPZ1 | 3 | 3 | 1.02 |
| Q8TD19 | NEK9 | 7 | 7 | 1.02 |
| P31939 | ATIC | 22 | 22 | 1.01 |
| P35080 | PFN2 | 7 | 7 | 1.01 |
| P45973 | CBX5 | 3 | 3 | 1.01 |
| Q86U42 | PABPN1 | 5 | 5 | 1.01 |
| Q15287 | RNPS1 | 4 | 4 | 1.01 |
| O15173 | PGRMC2 | 7 | 7 | 1.01 |
| P52272 | HNRNPM | 44 | 44 | 1.01 |
| P78417 | GSTO1 | 12 | 12 | 1.01 |
| P13984 | GTF2F2 | 4 | 4 | 1.01 |
| P62633 | CNBP | 3 | 3 | 1.01 |
| P28838 | LAP3 | 19 | 19 | 1.01 |
| P19387 | POLR2C | 4 | 4 | 1.01 |
| Q96AX1 | VPS33A | 2 | 2 | 1.01 |
| Q6Y1H2 | HACD2 | 2 | 2 | 1.01 |
| Q9Y520 | PRRC2C | 15 | 14 | 1.01 |
| Q9NUQ8 | ABCF3 | 5 | 5 | 1.01 |
| Q8IY81 | FTSJ3 | 5 | 5 | 1.01 |
| Q9Y4W6 | AFG3L2 | 6 | 6 | 1.01 |
| Q9GZS3 | WDR61 | 4 | 4 | 1.01 |
| P54920 | NAPA | 9 | 9 | 1.01 |
| O43242 | PSMD3 | 26 | 26 | 1.01 |
| Q96L92 | SNX27 | 2 | 2 | 1.01 |
| Q86V81 | ALYREF | 6 | 6 | 1.01 |
| O43491 | EPB41L2 | 37 | 33 | 1.01 |
| P48449 | LSS | 7 | 7 | 1.01 |
| P85037 | FOXK1 | 4 | 4 | 1.01 |
| Q9NRY4 | ARHGAP35 | 2 | 2 | 1.01 |
| Q9H3S7 | PTPN23 | 10 | 10 | 1.01 |
| Q9P2T1 | GMPR2 | 9 | 9 | 1.01 |
| Q9H4G0 | EPB41L1 | 20 | 18 | 1.01 |
| Q13616 | CUL1 | 8 | 8 | 1.01 |
| P16930 | FAH | 7 | 7 | 1.01 |
| P28288 | ABCD3 | 9 | 9 | 1.01 |
| P06865 | HEXA | 12 | 12 | 1.01 |

| Accession | Name | # Peptides | # Unique Peptides | Abundance Ratio: WWP2 / control |
|---|---|---|---|---|
| Q12959 | DLG1 | 12 | 11 | 1.01 |
| P61201 | COPS2 | 12 | 12 | 1.01 |
| P28482 | MAPK1 | 17 | 12 | 1.01 |
| Q9Y2A7 | NCKAP1 | 21 | 21 | 1.01 |
| P11047 | LAMC1 | 35 | 35 | 1.01 |
| A6NHG4 | DDTL | 7 | 7 | 1.01 |
| P16401 | H1-5 | 20 | 16 | 1.01 |
| O95989 | NUDT3 | 4 | 3 | 1.01 |
| P30046 | DDT | 7 | 7 | 1.01 |
| P07711 | CTSL | 8 | 7 | 1.01 |
| O60488 | ACSL4 | 9 | 9 | 1.01 |
| Q9NX40 | OCIAD1 | 6 | 6 | 1.01 |
| P55265 | ADAR | 14 | 14 | 1.01 |
| Q0IIM8 | TBC1D8B | 2 | 2 | 1.01 |
| O75431 | MTX2 | 2 | 2 | 1.01 |
| P52594 | AGFG1 | 3 | 2 | 1.01 |
| P78347 | GTF2I | 13 | 13 | 1.01 |
| Q9Y262 | EIF3L | 18 | 18 | 1.01 |
| P49914 | MTHFS | 2 | 2 | 1.01 |
| A6NDG6 | PGP | 3 | 3 | 1.01 |
| Q15019 | SEPTIN2 | 15 | 15 | 1.01 |
| P06730 | EIF4E | 5 | 5 | 1.01 |
| O75436 | VPS26A | 10 | 10 | 1.01 |
| Q9BWM7 | SFXN3 | 8 | 8 | 1.01 |
| P35998 | PSMC2 | 20 | 20 | 1.01 |
| P13611 | VCAN | 28 | 28 | 1.01 |
| A6NKT7 | RGPD3 | 7 | 7 | 1.01 |
| Q9Y3T9 | NOC2L | 5 | 5 | 1.01 |
| Q13188 | STK3 | 2 | 1 | 1.01 |
| P11717 | IGF2R | 20 | 20 | 1.01 |
| Q5TGZ0 | MICOS10 | 2 | 2 | 1.01 |
| Q16401 | PSMD5 | 12 | 12 | 1.01 |
| P35610 | SOAT1 | 2 | 2 | 1.01 |
| Q7Z3J3 | RGPD4 | 6 | 6 | 1.01 |
| Q8N1G4 | LRRC47 | 13 | 13 | 1.01 |
| O43765 | SGTA | 7 | 7 | 1.01 |
| P62314 | SNRPD1 | 2 | 2 | 1.01 |
| Q9P0M6 | MACROH2A2 | 7 | 6 | 1.01 |
| P84243 | H3-3A | 10 | 3 | 1.01 |
| Q8IY67 | RAVER1 | 2 | 2 | 1.01 |
| Q16777 | H2AC20 | 9 | 2 | 1.01 |
| Q7Z739 | YTHDF3 | 7 | 4 | 1.01 |
| P00367 | GLUD1 | 19 | 19 | 1.01 |
| Q9NQ88 | TIGAR | 3 | 3 | 1.01 |

| Accession | Name | # Peptides | # Unique Peptides | Abundance Ratio: WWP2 / control |
|---|---|---|---|---|
| Q86VR2 | RETREG3 | 3 | 3 | 1.01 |
| P49902 | NT5C2 | 2 | 2 | 1.01 |
| Q96CX2 | KCTD12 | 9 | 9 | 1.01 |
| Q8N3C0 | ASCC3 | 8 | 8 | 1.01 |
| Q13409 | DYNC1I2 | 12 | 12 | 1.01 |
| P07099 | EPHX1 | 15 | 15 | 1.01 |
| Q5VYK3 | ECPAS | 11 | 11 | 1.01 |
| P50579 | METAP2 | 7 | 7 | 1.01 |
| O00303 | EIF3F | 7 | 7 | 1.01 |
| A0A3B3IRV3 | MCTS2P | 4 | 4 | 1.01 |
| Q9GZZ1 | NAA50 | 2 | 2 | 1.01 |
| O15144 | ARPC2 | 18 | 18 | 1.01 |
| Q9BPW8 | NIPSNAP1 | 5 | 3 | 1.01 |
| Q14764 | MVP | 38 | 38 | 1.01 |
| Q96PK6 | RBM14 | 14 | 14 | 1.01 |
| O95394 | PGM3 | 15 | 15 | 1.01 |
| P46527 | CDKN1B | 2 | 2 | 1.01 |
| P09382 | LGALS1 | 12 | 12 | 1.01 |
| Q13232 | NME3 | 3 | 3 | 1.01 |
| O14974 | PPP1R12A | 12 | 12 | 1.01 |
| Q96JC1 | VPS39 | 2 | 2 | 1.01 |
| Q9BXS5 | AP1M1 | 7 | 5 | 1.01 |
| P40926 | MDH2 | 14 | 14 | 1.01 |
| P25445 | FAS | 2 | 2 | 1.01 |
| O95197 | RTN3 | 5 | 5 | 1.01 |
| Q13247 | SRSF6 | 8 | 5 | 1.01 |
| Q13488 | TCIRG1 | 7 | 7 | 1.01 |
| Q9P258 | RCC2 | 3 | 3 | 1.01 |
| P54136 | RARS1 | 23 | 23 | 1.01 |
| Q9UHD8 | SEPTIN9 | 29 | 29 | 1.01 |
| Q9BZF9 | UACA | 10 | 10 | 1.01 |
| P48509 | CD151 | 6 | 6 | 1.01 |
| Q14847 | LASP1 | 15 | 15 | 1.01 |
| P10809 | HSPD1 | 24 | 24 | 1.01 |
| Q92508 | PIEZO1 | 5 | 5 | 1.01 |
| O60502 | OGA | 4 | 4 | 1.01 |
| Q9NY33 | DPP3 | 11 | 11 | 1.01 |
| P98160 | HSPG2 | 130 | 130 | 1.01 |
| Q15046 | KARS1 | 18 | 18 | 1.01 |
| Q16836 | HADH | 5 | 5 | 1.01 |
| P19367 | HK1 | 35 | 31 | 1.01 |
| Q16473 | TNXA | 7 | 7 | 1.01 |
| Q15369 | ELOC | 6 | 6 | 1.01 |
| Q9UI08 | EVL | 5 | 4 | 1.01 |

| Accession | Name | # Peptides | # Unique Peptides | Abundance Ratio: WWP2 / control |
|---|---|---|---|---|
| P17987 | TCP1 | 22 | 22 | 1.01 |
| Q96FQ6 | S100A16 | 6 | 6 | 1.01 |
| Q9UBI6 | GNG12 | 4 | 4 | 1.01 |
| Q9Y5X2 | SNX8 | 5 | 5 | 1.01 |
| Q96I24 | FUBP3 | 16 | 13 | 1.01 |
| O14715 | RGPD8 | 4 | 4 | 1.01 |
| P38117 | ETFB | 12 | 12 | 1.01 |
| A6NHR9 | SMCHD1 | 6 | 6 | 1.01 |
| Q9NYL2 | MAP3K20 | 2 | 2 | 1.01 |
| P36542 | ATP5F1C | 12 | 12 | 1.01 |
| Q9P0J0 | NDUFA13 | 2 | 2 | 1.01 |
| P29966 | MARCKS | 8 | 8 | 1.01 |
| Q9BZL4 | PPP1R12C | 9 | 9 | 1.01 |
| O00754 | MAN2B1 | 14 | 14 | 1.01 |
| P23193 | TCEA1 | 4 | 4 | 1.01 |
| Q8N0X7 | SPART | 6 | 6 | 1.01 |
| P30405 | PPIF | 6 | 5 | 1.01 |
| Q9Y263 | PLAA | 12 | 12 | 1.01 |
| Q99666 | RGPD5 | 4 | 4 | 1.01 |
| Q8TBC4 | UBA3 | 6 | 6 | 1.01 |
| Q6P158 | DHX57 | 3 | 3 | 1.01 |
| Q9BVC4 | MLST8 | 2 | 2 | 1.01 |
| P09012 | SNRPA | 10 | 9 | 1.01 |
| P52298 | NCBP2 | 3 | 3 | 1.01 |
| Q8WUW1 | BRK1 | 3 | 3 | 1.01 |
| Q56UQ5 | | 2 | 2 | 1.01 |
| Q9NZJ7 | MTCH1 | 3 | 3 | 1.01 |
| O95425 | SVIL | 12 | 12 | 1.01 |
| Q07352 | ZFP36L1 | 2 | 1 | 1.01 |
| Q9UNM6 | PSMD13 | 12 | 12 | 1.01 |
| Q96S59 | RANBP9 | 2 | 2 | 1.01 |
| P09429 | HMGB1 | 16 | 14 | 1.01 |
| O43169 | CYB5B | 6 | 6 | 1.01 |
| P23588 | EIF4B | 19 | 19 | 1.01 |
| Q08945 | SSRP1 | 7 | 7 | 1.01 |
| Q9GZZ9 | UBA5 | 4 | 4 | 1.01 |
| P60900 | PSMA6 | 14 | 14 | 1.01 |
| Q15637 | SF1 | 9 | 9 | 1.01 |
| O15143 | ARPC1B | 14 | 14 | 1.01 |
| P84103 | SRSF3 | 14 | 13 | 1.01 |
| P36551 | CPOX | 3 | 3 | 1.01 |
| P08621 | SNRNP70 | 13 | 13 | 1.01 |
| Q13561 | DCTN2 | 15 | 15 | 1.01 |
| Q96S44 | TP53RK | 2 | 2 | 1.01 |

| Accession | Name | # Peptides | # Unique Peptides | Abundance Ratio: WWP2 / control |
|---|---|---|---|---|
| P62714 | PPP2CB | 12 | 1 | 1.01 |
| Q00839 | HNRNPU | 37 | 37 | 1.01 |
| O00232 | PSMD12 | 17 | 17 | 1.01 |
| O00401 | WASL | 4 | 4 | 1.01 |
| Q13153 | PAK1 | 3 | 3 | 1.01 |
| Q96F07 | CYFIP2 | 7 | 7 | 1.01 |
| P35579 | MYH9 | 157 | 134 | 1.01 |
| P11177 | PDHB | 8 | 8 | 1.01 |
| Q15843 | NEDD8 | 4 | 4 | 1.01 |
| P41091 | EIF2S3 | 20 | 20 | 1.01 |
| P07355 | ANXA2 | 35 | 35 | 1.01 |
| P07741 | APRT | 5 | 5 | 1.01 |
| O43813 | LANCL1 | 6 | 6 | 1.01 |
| P55795 | HNRNPH2 | 15 | 9 | 1.01 |
| P49368 | CCT3 | 30 | 30 | 1.01 |
| P20290 | BTF3 | 5 | 4 | 1.01 |
| Q6L8Q7 | PDE12 | 3 | 3 | 1.01 |
| Q96KP4 | CNDP2 | 10 | 10 | 1.01 |
| O60936 | NOL3 | 6 | 6 | 1.01 |
| Q9BW83 | IFT27 | 2 | 2 | 1.01 |
| P57772 | EEFSEC | 4 | 4 | 1.01 |
| P59998 | ARPC4 | 6 | 6 | 1.01 |
| P47755 | CAPZA2 | 10 | 6 | 1.01 |
| P61026 | RAB10 | 10 | 9 | 1.01 |
| Q9UKV3 | ACIN1 | 16 | 16 | 1.01 |
| O43678 | NDUFA2 | 6 | 6 | 1.01 |
| P53618 | COPB1 | 25 | 25 | 1.01 |
| Q14699 | RFTN1 | 3 | 3 | 1.01 |
| P48163 | ME1 | 10 | 10 | 1.01 |
| Q9ULD0 | OGDHL | 8 | 1 | 1.01 |
| Q9UKS6 | PACSIN3 | 8 | 8 | 1.01 |
| Q70UQ0 | IKBIP | 18 | 18 | 1.01 |
| Q8TDN6 | BRIX1 | 4 | 4 | 1.01 |
| Q9Y281 | CFL2 | 8 | 6 | 1.01 |
| O43827 | ANGPTL7 | 5 | 5 | 1.01 |
| P31948 | STIP1 | 37 | 37 | 1.01 |
| P08236 | GUSB | 4 | 4 | 1.01 |
| O00244 | ATOX1 | 4 | 4 | 1.01 |
| Q9H488 | POFUT1 | 5 | 5 | 1.01 |
| O95249 | GOSR1 | 2 | 2 | 1.01 |
| Q9P016 | THYN1 | 6 | 6 | 1.01 |
| P67775 | PPP2CA | 12 | 1 | 1.01 |
| Q9NZW5 | PALS2 | 10 | 10 | 1.01 |
| Q6PCE3 | PGM2L1 | 12 | 11 | 1.01 |

| Accession | Name | # Peptides | # Unique Peptides | Abundance Ratio: WWP2 / control |
|---|---|---|---|---|
| Q96BW5 | PTER | 2 | 2 | 1.01 |
| P60709 | ACTB | 26 | 1 | 1.01 |
| P61006 | RAB8A | 5 | 2 | 1.01 |
| Q969G3 | SMARCE1 | 7 | 6 | 1.01 |
| Q66LE6 | PPP2R2D | 3 | 3 | 1.01 |
| Q07812 | BAX | 7 | 7 | 1.01 |
| Q9BYJ9 | YTHDF1 | 4 | 2 | 1.00 |
| Q13505 | MTX1 | 3 | 3 | 1.00 |
| Q6ZNL6 | FGD5 | 2 | 2 | 1.00 |
| O00273 | DFFA | 7 | 7 | 1.00 |
| Q15691 | MAPRE1 | 12 | 10 | 1.00 |
| P12235 | SLC25A4 | 11 | 1 | 1.00 |
| Q5JTD0 | TJAP1 | 4 | 4 | 1.00 |
| Q13596 | SNX1 | 12 | 11 | 1.00 |
| P49589 | CARS1 | 20 | 20 | 1.00 |
| Q13155 | AIMP2 | 7 | 7 | 1.00 |
| P29218 | IMPA1 | 11 | 11 | 1.00 |
| P22307 | SCP2 | 12 | 12 | 1.00 |
| Q9NTZ6 | RBM12 | 14 | 14 | 1.00 |
| Q00610 | CLTC | 64 | 64 | 1.00 |
| O60573 | EIF4E2 | 4 | 4 | 1.00 |
| P46108 | CRK | 9 | 9 | 1.00 |
| Q5GLZ8 | HERC4 | 12 | 12 | 1.00 |
| P51888 | PRELP | 12 | 12 | 1.00 |
| P53597 | SUCLG1 | 6 | 6 | 1.00 |
| P52943 | CRIP2 | 6 | 6 | 1.00 |
| Q8IZP0 | ABI1 | 5 | 5 | 1.00 |
| Q13630 | GFUS | 6 | 6 | 1.00 |
| Q9Y230 | RUVBL2 | 21 | 21 | 1.00 |
| P56182 | RRP1 | 2 | 2 | 1.00 |
| Q7Z2K6 | ERMP1 | 2 | 2 | 1.00 |
| Q8N5J2 | MINDY1 | 2 | 2 | 1.00 |
| P26368 | U2AF2 | 9 | 9 | 1.00 |
| Q16181 | SEPTIN7 | 27 | 27 | 1.00 |
| Q9BUN8 | DERL1 | 3 | 3 | 1.00 |
| Q08379 | GOLGA2 | 8 | 8 | 1.00 |
| Q9HBL0 | TNS1 | 41 | 38 | 1.00 |
| P11117 | ACP2 | 2 | 2 | 1.00 |
| P61011 | SRP54 | 14 | 14 | 1.00 |
| Q6DRA6 | H2BC19P | 5 | 2 | 1.00 |
| Q9Y679 | AUP1 | 2 | 2 | 1.00 |
| P35573 | AGL | 8 | 8 | 1.00 |
| P06576 | ATP5F1B | 18 | 18 | 1.00 |
| O60762 | DPM1 | 7 | 7 | 1.00 |

| Accession | Name | # Peptides | # Unique Peptides | Abundance Ratio: WWP2 / control |
|---|---|---|---|---|
| Q9Y2Z0 | SUGT1 | 9 | 9 | 1.00 |
| O95210 | STBD1 | 10 | 10 | 1.00 |
| P53814 | SMTN | 4 | 4 | 1.00 |
| P46063 | RECQL | 27 | 27 | 1.00 |
| P52907 | CAPZA1 | 14 | 10 | 1.00 |
| Q8N1F7 | NUP93 | 18 | 18 | 1.00 |
| Q13428 | TCOF1 | 9 | 9 | 1.00 |
| O75935 | DCTN3 | 8 | 8 | 1.00 |
| P78527 | PRKDC | 104 | 104 | 1.00 |
| Q6DN03 | H2BC20P | 5 | 2 | 1.00 |
| Q86VN1 | VPS36 | 4 | 4 | 1.00 |
| Q99436 | PSMB7 | 8 | 8 | 1.00 |
| O75534 | CSDE1 | 15 | 15 | 1.00 |
| Q8N0Y7 | PGAM4 | 7 | 4 | 1.00 |
| O00159 | MYO1C | 43 | 42 | 1.00 |
| O95749 | GGPS1 | 2 | 2 | 1.00 |
| O14562 | UBFD1 | 4 | 4 | 1.00 |
| Q16576 | RBBP7 | 9 | 4 | 1.00 |
| P08238 | HSP90AB1 | 43 | 21 | 1.00 |
| Q9Y2V2 | CARHSP1 | 3 | 3 | 1.00 |
| Q53GQ0 | HSD17B12 | 5 | 5 | 1.00 |
| P46439 | GSTM5 | 5 | 2 | 1.00 |
| Q14696 | MESD | 8 | 8 | 1.00 |
| Q12874 | SF3A3 | 11 | 11 | 1.00 |
| P09622 | DLD | 11 | 11 | 1.00 |
| P63151 | PPP2R2A | 8 | 8 | 1.00 |
| P49354 | FNTA | 4 | 4 | 1.00 |
| Q99497 | PARK7 | 15 | 15 | 1.00 |
| P08579 | SNRPB2 | 6 | 5 | 1.00 |
| O96000 | NDUFB10 | 4 | 4 | 1.00 |
| P98095 | FBLN2 | 13 | 13 | 1.00 |
| Q96RS6 | NUDCD1 | 7 | 7 | 1.00 |
| P61158 | ACTR3 | 17 | 17 | 1.00 |
| Q08170 | SRSF4 | 12 | 9 | 1.00 |
| O43390 | HNRNPR | 22 | 13 | 1.00 |
| Q04760 | GLO1 | 8 | 8 | 1.00 |
| Q7Z434 | MAVS | 6 | 6 | 1.00 |
| Q9H361 | PABPC3 | 17 | 14 | 1.00 |
| P28074 | PSMB5 | 10 | 10 | 1.00 |
| Q9UIJ7 | AK3 | 10 | 10 | 1.00 |
| Q92733 | PRCC | 3 | 3 | 1.00 |
| Q96TC7 | RMDN3 | 4 | 4 | 1.00 |
| P56192 | MARS1 | 17 | 17 | 1.00 |
| Q16774 | GUK1 | 4 | 4 | 1.00 |

| Accession | Name | # Peptides | # Unique Peptides | Abundance Ratio: WWP2 / control |
|---|---|---|---|---|
| Q13042 | CDC16 | 2 | 2 | 1.00 |
| Q93009 | USP7 | 12 | 12 | 1.00 |
| Q6P9B6 | MEAK7 | 4 | 4 | 1.00 |
| Q12904 | AIMP1 | 14 | 14 | 1.00 |
| Q96N67 | DOCK7 | 5 | 5 | 1.00 |
| P0CAP2 | POLR2M | 2 | 2 | 1.00 |
| P62736 | ACTA2 | 20 | 7 | 1.00 |
| Q99627 | COPS8 | 3 | 3 | 1.00 |
| P61225 | RAP2B | 4 | 4 | 1.00 |
| P55809 | OXCT1 | 7 | 7 | 1.00 |
| P15374 | UCHL3 | 6 | 5 | 1.00 |
| P40939 | HADHA | 29 | 29 | 1.00 |
| P07384 | CAPN1 | 20 | 19 | 1.00 |
| P62191 | PSMC1 | 20 | 19 | 1.00 |
| Q96JB5 | CDK5RAP3 | 8 | 8 | 1.00 |
| Q4VXU2 | PABPC1L | 8 | 6 | 1.00 |
| Q9NRX3 | NDUFA4L2 | 6 | 6 | 1.00 |
| Q86UP2 | KTN1 | 62 | 62 | 1.00 |
| O00410 | IPO5 | 14 | 13 | 1.00 |
| O75223 | GGCT | 5 | 5 | 1.00 |
| Q9Y6Y8 | SEC23IP | 18 | 18 | 1.00 |
| P06899 | H2BC11 | 10 | 2 | 1.00 |
| P50135 | HNMT | 2 | 2 | 1.00 |
| Q15008 | PSMD6 | 21 | 21 | 1.00 |
| P23527 | H2BC17 | 10 | 2 | 1.00 |
| P68032 | ACTC1 | 21 | 7 | 1.00 |
| Q96KR1 | ZFR | 7 | 7 | 1.00 |
| Q9UK45 | LSM7 | 2 | 1 | 1.00 |
| O43865 | AHCYL1 | 15 | 15 | 1.00 |
| Q96SB3 | PPP1R9B | 8 | 8 | 1.00 |
| O75155 | CAND2 | 4 | 4 | 1.00 |
| Q99536 | VAT1 | 16 | 16 | 1.00 |
| Q4G0F5 | VPS26B | 4 | 4 | 1.00 |
| Q9BZD3 | | 2 | 2 | 1.00 |
| Q8TAE6 | PPP1R14C | 2 | 2 | 1.00 |
| Q8IWR0 | ZC3H7A | 2 | 2 | 1.00 |
| Q16778 | H2BC21 | 10 | 2 | 1.00 |
| P38405 | GNAL | 3 | 3 | 1.00 |
| Q9H0U3 | MAGT1 | 9 | 9 | 1.00 |
| Q96D15 | RCN3 | 12 | 12 | 1.00 |
| O00571 | DDX3X | 21 | 6 | 1.00 |
| Q9UBW8 | COPS7A | 7 | 7 | 1.00 |
| P26599 | PTBP1 | 12 | 12 | 1.00 |
| P53990 | IST1 | 6 | 6 | 1.00 |

| Accession | Name | # Peptides | # Unique Peptides | Abundance Ratio: WWP2 / control |
|---|---|---|---|---|
| P20338 | RAB4A | 4 | 4 | 1.00 |
| Q9Y3E0 | GOLT1B | 2 | 2 | 1.00 |
| O43674 | NDUFB5 | 2 | 2 | 1.00 |
| Q9BYX2 | TBC1D2 | 2 | 2 | 1.00 |
| P33778 | H2BC3 | 10 | 2 | 1.00 |
| Q9Y2T4 | PPP2R2C | 2 | 2 | 1.00 |
| Q96QR8 | PURB | 6 | 5 | 1.00 |
| Q9HCU5 | PREB | 5 | 5 | 1.00 |
| Q9HDC9 | APMAP | 16 | 16 | 1.00 |
| Q00688 | FKBP3 | 14 | 14 | 1.00 |
| P28070 | PSMB4 | 6 | 6 | 1.00 |
| Q07021 | C1QBP | 3 | 3 | 1.00 |
| P12956 | XRCC6 | 26 | 26 | 1.00 |
| Q9NQR4 | NIT2 | 7 | 7 | 1.00 |
| Q9Y5S2 | CDC42BPB | 11 | 7 | 1.00 |
| P02794 | FTH1 | 12 | 12 | 1.00 |
| Q02978 | SLC25A11 | 10 | 10 | 1.00 |
| P33241 | LSP1 | 3 | 3 | 1.00 |
| O43598 | DNPH1 | 4 | 4 | 1.00 |
| P15531 | NME1 | 9 | 4 | 1.00 |
| P0DJD0 | RGPD1 | 6 | 6 | 1.00 |
| O14929 | HAT1 | 3 | 3 | 1.00 |
| Q86X83 | COMMD2 | 2 | 2 | 1.00 |
| O00471 | EXOC5 | 5 | 5 | 1.00 |
| Q9UMY4 | SNX12 | 7 | 6 | 1.00 |
| P0DJD1 | RGPD2 | 6 | 6 | 1.00 |
| Q5JWF2 | GNAS | 14 | 14 | 1.00 |
| Q92878 | RAD50 | 15 | 15 | 1.00 |
| P26006 | ITGA3 | 3 | 3 | 1.00 |
| Q9NP58 | ABCB6 | 3 | 3 | 1.00 |
| O75351 | VPS4B | 12 | 7 | 1.00 |
| Q9Y696 | CLIC4 | 12 | 12 | 1.00 |
| Q9NQP4 | PFDN4 | 3 | 3 | 1.00 |
| O96008 | TOMM40 | 2 | 2 | 1.00 |
| O75340 | PDCD6 | 5 | 5 | 1.00 |
| P09211 | GSTP1 | 8 | 8 | 1.00 |
| P07203 | GPX1 | 7 | 7 | 1.00 |
| P20042 | EIF2S2 | 13 | 13 | 1.00 |
| Q14980 | NUMA1 | 68 | 68 | 1.00 |
| Q7Z7H5 | TMED4 | 6 | 4 | 1.00 |
| P36957 | DLST | 9 | 9 | 1.00 |
| Q9NVV5 | AIG1 | 2 | 2 | 1.00 |
| P35658 | NUP214 | 5 | 5 | 1.00 |
| P50991 | CCT4 | 23 | 23 | 1.00 |

| Accession | Name | # Peptides | # Unique Peptides | Abundance Ratio: WWP2 / control |
|---|---|---|---|---|
| O14964 | HGS | 18 | 18 | 1.00 |
| Q9Y6W5 | WASF2 | 10 | 10 | 1.00 |
| P46781 | RPS9 | 14 | 14 | 1.00 |
| Q15365 | PCBP1 | 11 | 7 | 1.00 |
| Q15643 | TRIP11 | 7 | 7 | 1.00 |
| O15027 | SEC16A | 9 | 9 | 1.00 |
| P06753 | TPM3 | 27 | 10 | 1.00 |
| P62258 | YWHAE | 23 | 21 | 1.00 |
| P20645 | M6PR | 5 | 5 | 1.00 |
| Q14692 | BMS1 | 4 | 4 | 1.00 |
| P55010 | EIF5 | 10 | 10 | 1.00 |
| Q8N1B4 | VPS52 | 2 | 2 | 1.00 |
| O75874 | IDH1 | 20 | 20 | 1.00 |
| O43583 | DENR | 7 | 7 | 1.00 |
| P48507 | GCLM | 5 | 5 | 1.00 |
| Q86VM9 | ZC3H18 | 4 | 4 | 1.00 |
| P25788 | PSMA3 | 13 | 13 | 1.00 |
| P42704 | LRPPRC | 21 | 21 | 1.00 |
| P04921 | GYPC | 2 | 2 | 1.00 |
| P62241 | RPS8 | 10 | 10 | 1.00 |
| P28066 | PSMA5 | 10 | 10 | 1.00 |
| P09234 | SNRPC | 3 | 3 | 1.00 |
| P16152 | CBR1 | 13 | 11 | 1.00 |
| P19105 | MYL12A | 6 | 3 | 1.00 |
| P51858 | HDGF | 19 | 18 | 1.00 |
| Q6Y7W6 | GIGYF2 | 13 | 12 | 1.00 |
| P17676 | CEBPB | 2 | 2 | 1.00 |
| P31942 | HNRNPH3 | 10 | 9 | 1.00 |
| B2RPK0 | HMGB1P1 | 7 | 5 | 1.00 |
| O75400 | PRPF40A | 9 | 9 | 1.00 |
| O43795 | MYO1B | 32 | 32 | 1.00 |
| P24666 | ACP1 | 7 | 7 | 1.00 |
| Q6YHK3 | CD109 | 29 | 29 | 1.00 |
| P01033 | TIMP1 | 3 | 3 | 1.00 |
| P63267 | ACTG2 | 19 | 6 | 1.00 |
| Q9UBC2 | EPS15L1 | 5 | 5 | 1.00 |
| P46926 | GNPDA1 | 8 | 4 | 1.00 |
| Q86X55 | CARM1 | 5 | 5 | 1.00 |
| Q9NWV4 | CZIB | 6 | 6 | 1.00 |
| P07996 | THBS1 | 34 | 33 | 1.00 |
| P00403 | MT-CO2 | 3 | 3 | 1.00 |
| O75746 | SLC25A12 | 2 | 2 | 1.00 |
| P31153 | MAT2A | 15 | 15 | 1.00 |
| P49189 | ALDH9A1 | 15 | 15 | 1.00 |

| Accession | Name | # Peptides | # Unique Peptides | Abundance Ratio: WWP2 / control |
|---|---|---|---|---|
| P62269 | RPS18 | 13 | 13 | 1.00 |
| O95834 | EML2 | 3 | 3 | 1.00 |
| Q5JRA6 | MIA3 | 21 | 21 | 1.00 |
| O14950 | MYL12B | 6 | 3 | 1.00 |
| Q6VY07 | PACS1 | 3 | 3 | 1.00 |
| P68133 | ACTA1 | 20 | 6 | 1.00 |
| Q9P0P0 | RNF181 | 2 | 2 | 1.00 |
| O15061 | SYNM | 33 | 33 | 1.00 |
| P12081 | HARS1 | 12 | 8 | 1.00 |
| Q9NUP9 | LIN7C | 6 | 6 | 1.00 |
| Q06210 | GFPT1 | 29 | 23 | 1.00 |
| P17612 | PRKACA | 11 | 6 | 1.00 |
| O00487 | PSMD14 | 8 | 8 | 1.00 |
| O95870 | ABHD16A | 4 | 4 | 1.00 |
| P62937 | PPIA | 17 | 16 | 1.00 |
| P48643 | CCT5 | 26 | 26 | 1.00 |
| Q641Q2 | WASHC2A | 7 | 1 | 1.00 |
| Q00325 | SLC25A3 | 12 | 12 | 1.00 |
| P42166 | TMPO | 9 | 9 | 1.00 |
| Q6UXH1 | CRELD2 | 3 | 3 | 1.00 |
| Q9H4A4 | RNPEP | 20 | 20 | 1.00 |
| Q8TF05 | PPP4R1 | 2 | 2 | 1.00 |
| P30419 | NMT1 | 8 | 8 | 1.00 |
| P29992 | GNA11 | 10 | 7 | 1.00 |
| P17480 | UBTF | 9 | 9 | 1.00 |
| O75643 | SNRNP200 | 31 | 31 | 1.00 |
| Q9UNF1 | MAGED2 | 18 | 17 | 1.00 |
| P46783 | RPS10 | 15 | 14 | 1.00 |
| Q9NUJ3 | TCP11L1 | 2 | 2 | 1.00 |
| Q8IZ83 | ALDH16A1 | 6 | 6 | 1.00 |
| P61604 | HSPE1 | 9 | 9 | 1.00 |
| Q8WVM8 | SCFD1 | 11 | 11 | 1.00 |
| Q9H2H8 | PPIL3 | 4 | 4 | 1.00 |
| Q9NX14 | NDUFB11 | 2 | 2 | 1.00 |
| P17174 | GOT1 | 11 | 11 | 1.00 |
| Q99584 | S100A13 | 5 | 5 | 1.00 |
| P09543 | CNP | 14 | 13 | 1.00 |
| Q9NR22 | PRMT8 | 4 | 4 | 1.00 |
| P61009 | SPCS3 | 2 | 2 | 1.00 |
| Q08257 | CRYZ | 9 | 9 | 1.00 |
| Q9Y315 | DERA | 4 | 4 | 1.00 |
| P23634 | ATP2B4 | 30 | 18 | 1.00 |
| Q9HC07 | TMEM165 | 2 | 2 | 1.00 |
| O60282 | KIF5C | 7 | 7 | 1.00 |

| Accession | Name | # Peptides | # Unique Peptides | Abundance Ratio: WWP2 / control |
|---|---|---|---|---|
| P09038 | FGF2 | 2 | 2 | 1.00 |
| Q9BT40 | INPP5K | 2 | 2 | 1.00 |
| Q9UNL2 | SSR3 | 3 | 3 | 1.00 |
| P53621 | COPA | 50 | 50 | 1.00 |
| P36959 | GMPR | 2 | 2 | 1.00 |
| Q9Y3A3 | MOB4 | 3 | 3 | 1.00 |
| P61764 | STXBP1 | 6 | 6 | 1.00 |
| O75964 | ATP5MG | 4 | 4 | 1.00 |
| Q9NR28 | DIABLO | 7 | 7 | 1.00 |
| P28331 | NDUFS1 | 14 | 14 | 1.00 |
| Q06547 | GABPB1 | 3 | 3 | 1.00 |
| Q15811 | ITSN1 | 3 | 3 | 1.00 |
| P55060 | CSE1L | 21 | 21 | 1.00 |
| Q92572 | AP3S1 | 6 | 6 | 1.00 |
| Q8TAA3 | PSMA8 | 5 | 5 | 1.00 |
| Q9BZJ0 | CRNKL1 | 2 | 2 | 1.00 |
| Q7Z3B4 | NUP54 | 2 | 2 | 1.00 |
| Q6P1J9 | CDC73 | 5 | 5 | 1.00 |
| Q01995 | TAGLN | 17 | 17 | 1.00 |
| O14773 | TPP1 | 4 | 4 | 1.00 |
| P40227 | CCT6A | 18 | 18 | 0.99 |
| Q92504 | SLC39A7 | 4 | 4 | 0.99 |
| Q9NP61 | ARFGAP3 | 13 | 13 | 0.99 |
| P27348 | YWHAQ | 19 | 14 | 0.99 |
| Q8N7R7 | CCNYL1 | 2 | 1 | 0.99 |
| Q5NE16 | CTSL3P | 2 | 1 | 0.99 |
| O76054 | SEC14L2 | 4 | 4 | 0.99 |
| P14868 | DARS1 | 29 | 29 | 0.99 |
| Q5H9R7 | PPP6R3 | 4 | 4 | 0.99 |
| O00635 | TRIM38 | 2 | 2 | 0.99 |
| Q99798 | ACO2 | 18 | 18 | 0.99 |
| Q9UH99 | SUN2 | 13 | 13 | 0.99 |
| P10155 | RO60 | 13 | 13 | 0.99 |
| Q9Y383 | LUC7L2 | 10 | 6 | 0.99 |
| Q16629 | SRSF7 | 11 | 10 | 0.99 |
| Q92845 | KIFAP3 | 2 | 1 | 0.99 |
| P51911 | CNN1 | 3 | 1 | 0.99 |
| O95793 | STAU1 | 8 | 8 | 0.99 |
| O76021 | RSL1D1 | 17 | 17 | 0.99 |
| A2RRP1 | NBAS | 9 | 9 | 0.99 |
| Q99471 | PFDN5 | 4 | 4 | 0.99 |
| O00422 | SAP18 | 4 | 4 | 0.99 |
| Q9BTV4 | TMEM43 | 14 | 14 | 0.99 |
| Q9H7N4 | SCAF1 | 2 | 2 | 0.99 |

| Accession | Name | # Peptides | # Unique Peptides | Abundance Ratio: WWP2 / control |
|---|---|---|---|---|
| Q9UQN3 | CHMP2B | 4 | 4 | 0.99 |
| Q9UHF7 | TRPS1 | 4 | 4 | 0.99 |
| Q13144 | EIF2B5 | 6 | 6 | 0.99 |
| Q96RQ1 | ERGIC2 | 2 | 2 | 0.99 |
| P02545 | LMNA | 68 | 66 | 0.99 |
| P35268 | RPL22 | 5 | 5 | 0.99 |
| P16615 | ATP2A2 | 26 | 26 | 0.99 |
| P62495 | ETF1 | 11 | 11 | 0.99 |
| P05455 | SSB | 16 | 15 | 0.99 |
| Q86TM6 | SYVN1 | 4 | 4 | 0.99 |
| P09132 | SRP19 | 2 | 2 | 0.99 |
| P67812 | SEC11A | 5 | 5 | 0.99 |
| Q7Z4V5 | HDGFL2 | 6 | 5 | 0.99 |
| P30740 | SERPINB1 | 15 | 15 | 0.99 |
| Q58FG1 | HSP90AA4P | 3 | 2 | 0.99 |
| Q14558 | PRPSAP1 | 4 | 4 | 0.99 |
| P13010 | XRCC5 | 31 | 31 | 0.99 |
| Q9NSK0 | KLC4 | 3 | 3 | 0.99 |
| P83916 | CBX1 | 2 | 2 | 0.99 |
| Q9UKK9 | NUDT5 | 8 | 8 | 0.99 |
| P53384 | NUBP1 | 3 | 3 | 0.99 |
| Q5NDL2 | EOGT | 3 | 3 | 0.99 |
| Q9P2B2 | PTGFRN | 4 | 4 | 0.99 |
| Q9NZN3 | EHD3 | 11 | 2 | 0.99 |
| Q9P2E9 | RRBP1 | 117 | 117 | 0.99 |
| P63279 | UBE2I | 5 | 5 | 0.99 |
| Q4J6C6 | PREPL | 2 | 2 | 0.99 |
| Q9BRF8 | CPPED1 | 8 | 8 | 0.99 |
| Q92945 | KHSRP | 29 | 24 | 0.99 |
| Q92841 | DDX17 | 27 | 19 | 0.99 |
| Q9UHW9 | SLC12A6 | 4 | 4 | 0.99 |
| A0FGR8 | ESYT2 | 8 | 7 | 0.99 |
| Q9NPJ3 | ACOT13 | 4 | 4 | 0.99 |
| Q15717 | ELAVL1 | 12 | 12 | 0.99 |
| O60884 | DNAJA2 | 5 | 5 | 0.99 |
| Q9NYU2 | UGGT1 | 32 | 30 | 0.99 |
| P62826 | RAN | 13 | 13 | 0.99 |
| P35606 | COPB2 | 29 | 29 | 0.99 |
| Q9NZN4 | EHD2 | 24 | 22 | 0.99 |
| O60313 | OPA1 | 6 | 6 | 0.99 |
| Q9BZE9 | ASPSCR1 | 4 | 4 | 0.99 |
| Q9H3N1 | TMX1 | 5 | 5 | 0.99 |
| Q16775 | HAGH | 5 | 5 | 0.99 |
| P17980 | PSMC3 | 14 | 14 | 0.99 |

| Accession | Name | # Peptides | # Unique Peptides | Abundance Ratio: WWP2 / control |
|---|---|---|---|---|
| P62333 | PSMC6 | 16 | 16 | 0.99 |
| Q9Y5L0 | TNPO3 | 2 | 2 | 0.99 |
| O60763 | USO1 | 26 | 26 | 0.99 |
| P62841 | RPS15 | 6 | 5 | 0.99 |
| Q6ZXV5 | TMTC3 | 5 | 5 | 0.99 |
| P05362 | ICAM1 | 11 | 11 | 0.99 |
| Q9UJY1 | HSPB8 | 2 | 2 | 0.99 |
| O00400 | SLC33A1 | 3 | 3 | 0.99 |
| P07814 | EPRS1 | 51 | 51 | 0.99 |
| P51153 | RAB13 | 9 | 7 | 0.99 |
| P06454 | PTMA | 5 | 5 | 0.99 |
| Q6P2I3 | FAHD2B | 2 | 2 | 0.99 |
| Q15233 | NONO | 26 | 22 | 0.99 |
| P08758 | ANXA5 | 22 | 22 | 0.99 |
| Q9H553 | ALG2 | 5 | 5 | 0.99 |
| Q8TED1 | GPX8 | 8 | 8 | 0.99 |
| P49790 | NUP153 | 3 | 3 | 0.99 |
| Q00535 | CDK5 | 2 | 2 | 0.99 |
| Q9Y371 | SH3GLB1 | 9 | 9 | 0.99 |
| P35580 | MYH10 | 67 | 44 | 0.99 |
| P29692 | EEF1D | 11 | 9 | 0.99 |
| Q92734 | TFG | 10 | 10 | 0.99 |
| Q13043 | STK4 | 2 | 1 | 0.99 |
| P50990 | CCT8 | 38 | 38 | 0.99 |
| P30048 | PRDX3 | 7 | 7 | 0.99 |
| O43488 | AKR7A2 | 9 | 9 | 0.99 |
| P61163 | ACTR1A | 10 | 7 | 0.99 |
| Q9NTJ5 | SACM1L | 15 | 15 | 0.99 |
| Q9Y282 | ERGIC3 | 5 | 5 | 0.99 |
| P49748 | ACADVL | 26 | 26 | 0.99 |
| P62805 | H4-16 | 9 | 9 | 0.99 |
| O75886 | STAM2 | 5 | 4 | 0.99 |
| Q1KMD3 | HNRNPUL2 | 24 | 24 | 0.99 |
| Q9UKZ9 | PCOLCE2 | 2 | 2 | 0.99 |
| Q92696 | RABGGTA | 5 | 5 | 0.99 |
| P55287 | CDH11 | 8 | 8 | 0.99 |
| P12268 | IMPDH2 | 16 | 14 | 0.99 |
| Q9NV96 | TMEM30A | 4 | 4 | 0.99 |
| P62318 | SNRPD3 | 3 | 3 | 0.99 |
| Q16555 | DPYSL2 | 25 | 22 | 0.99 |
| Q12906 | ILF3 | 27 | 27 | 0.99 |
| Q9Y6M1 | IGF2BP2 | 9 | 9 | 0.99 |
| P22234 | PAICS | 16 | 16 | 0.99 |
| Q96QK1 | VPS35 | 18 | 18 | 0.99 |

| Accession | Name | # Peptides | # Unique Peptides | Abundance Ratio: WWP2 / control |
|---|---|---|---|---|
| Q8TF66 | LRRC15 | 7 | 7 | 0.99 |
| P14735 | IDE | 5 | 5 | 0.99 |
| P62847 | RPS24 | 6 | 6 | 0.99 |
| P20700 | LMNB1 | 38 | 35 | 0.99 |
| P17252 | PRKCA | 21 | 20 | 0.99 |
| O43707 | ACTN4 | 59 | 44 | 0.99 |
| P60660 | MYL6 | 11 | 9 | 0.99 |
| P22695 | UQCRC2 | 16 | 16 | 0.99 |
| Q6UN15 | FIP1L1 | 6 | 6 | 0.99 |
| O43823 | AKAP8 | 4 | 4 | 0.99 |
| P13073 | COX4I1 | 10 | 10 | 0.99 |
| Q9Y3C8 | UFC1 | 3 | 3 | 0.99 |
| Q58FF7 | HSP90AB3P | 21 | 6 | 0.99 |
| Q86XP3 | DDX42 | 11 | 11 | 0.99 |
| P21399 | ACO1 | 26 | 26 | 0.99 |
| Q75LS8 | FKBP9P1 | 2 | 2 | 0.99 |
| Q9NTK5 | OLA1 | 14 | 14 | 0.99 |
| O00233 | PSMD9 | 7 | 7 | 0.99 |
| Q9UEU0 | VTI1B | 4 | 4 | 0.99 |
| Q16698 | DECR1 | 9 | 9 | 0.99 |
| Q8WVV9 | HNRNPLL | 2 | 1 | 0.99 |
| P39023 | RPL3 | 25 | 25 | 0.99 |
| P35241 | RDX | 39 | 20 | 0.99 |
| Q9UK59 | DBR1 | 2 | 2 | 0.99 |
| P62136 | PPP1CA | 12 | 4 | 0.99 |
| Q02809 | PLOD1 | 25 | 25 | 0.99 |
| P48729 | CSNK1A1 | 5 | 5 | 0.99 |
| Q9NPA0 | EMC7 | 4 | 4 | 0.99 |
| O43592 | XPOT | 6 | 6 | 0.99 |
| Q99613 | EIF3C | 28 | 28 | 0.99 |
| Q7L7X3 | TAOK1 | 4 | 3 | 0.99 |
| Q9Y2X3 | NOP58 | 10 | 10 | 0.99 |
| P11802 | CDK4 | 4 | 4 | 0.99 |
| O43818 | RRP9 | 3 | 3 | 0.99 |
| Q14651 | PLS1 | 4 | 1 | 0.99 |
| P29558 | RBMS1 | 3 | 3 | 0.99 |
| Q9H201 | EPN3 | 2 | 2 | 0.99 |
| O94760 | DDAH1 | 11 | 10 | 0.99 |
| Q9HAT2 | SIAE | 5 | 5 | 0.99 |
| Q9UBQ5 | EIF3K | 6 | 6 | 0.99 |
| O60361 | NME2P1 | 6 | 4 | 0.99 |
| Q9UHV9 | PFDN2 | 6 | 6 | 0.99 |
| O95837 | GNA14 | 3 | 1 | 0.99 |
| P43243 | MATR3 | 25 | 25 | 0.99 |

| Accession | Name | # Peptides | # Unique Peptides | Abundance Ratio: WWP2 / control |
|---|---|---|---|---|
| O00264 | PGRMC1 | 8 | 8 | 0.99 |
| Q12841 | FSTL1 | 12 | 12 | 0.99 |
| Q8IWB7 | WDFY1 | 5 | 5 | 0.99 |
| P49750 | YLPM1 | 11 | 10 | 0.99 |
| P14314 | PRKCSH | 23 | 23 | 0.99 |
| P05787 | KRT8 | 6 | 3 | 0.99 |
| P11940 | PABPC1 | 33 | 22 | 0.99 |
| Q9Y285 | FARSA | 13 | 13 | 0.99 |
| Q9BVC6 | TMEM109 | 5 | 5 | 0.99 |
| Q8NC56 | LEMD2 | 9 | 9 | 0.99 |
| Q9Y6K5 | OAS3 | 2 | 2 | 0.99 |
| Q7L014 | DDX46 | 13 | 13 | 0.99 |
| Q9UBE0 | SAE1 | 14 | 14 | 0.99 |
| Q9UHY7 | ENOPH1 | 3 | 3 | 0.99 |
| Q13217 | DNAJC3 | 8 | 8 | 0.99 |
| P34897 | SHMT2 | 15 | 15 | 0.99 |
| P62277 | RPS13 | 10 | 10 | 0.99 |
| Q9BVP2 | GNL3 | 6 | 6 | 0.99 |
| O43815 | STRN | 2 | 2 | 0.99 |
| P31946 | YWHAB | 17 | 8 | 0.99 |
| Q92900 | UPF1 | 23 | 23 | 0.99 |
| P41226 | UBA7 | 5 | 5 | 0.99 |
| P12955 | PEPD | 11 | 11 | 0.99 |
| P00441 | SOD1 | 15 | 15 | 0.99 |
| O96019 | ACTL6A | 4 | 4 | 0.99 |
| P23142 | FBLN1 | 20 | 20 | 0.99 |
| P31930 | UQCRC1 | 11 | 10 | 0.99 |
| Q96FJ2 | DYNLL2 | 6 | 3 | 0.99 |
| O00203 | AP3B1 | 21 | 21 | 0.99 |
| Q14141 | SEPTIN6 | 9 | 2 | 0.99 |
| Q70E73 | RAPH1 | 15 | 15 | 0.99 |
| Q7Z4H3 | HDDC2 | 4 | 4 | 0.99 |
| Q96FW1 | OTUB1 | 11 | 11 | 0.99 |
| P38646 | HSPA9 | 37 | 36 | 0.99 |
| Q7Z406 | MYH14 | 12 | 1 | 0.99 |
| O43399 | TPD52L2 | 15 | 15 | 0.99 |
| P60866 | RPS20 | 5 | 5 | 0.99 |
| Q13185 | CBX3 | 7 | 7 | 0.99 |
| Q12840 | KIF5A | 6 | 6 | 0.99 |
| Q9NZ45 | CISD1 | 3 | 3 | 0.99 |
| O75663 | TIPRL | 5 | 5 | 0.99 |
| Q9Y570 | PPME1 | 8 | 8 | 0.99 |
| Q96A33 | CCDC47 | 11 | 11 | 0.99 |
| P61160 | ACTR2 | 14 | 14 | 0.99 |

| Accession | Name | # Peptides | # Unique Peptides | Abundance Ratio: WWP2 / control |
|---|---|---|---|---|
| Q12797 | ASPH | 33 | 33 | 0.99 |
| P49755 | TMED10 | 11 | 10 | 0.99 |
| P62312 | LSM6 | 4 | 4 | 0.99 |
| Q8NC51 | SERBP1 | 15 | 15 | 0.99 |
| Q02750 | MAP2K1 | 12 | 8 | 0.99 |
| P62753 | RPS6 | 16 | 16 | 0.99 |
| Q9NX24 | NHP2 | 4 | 4 | 0.99 |
| Q9Y265 | RUVBL1 | 13 | 13 | 0.99 |
| Q93052 | LPP | 13 | 13 | 0.99 |
| Q2VIR3 | EIF2S3B | 18 | 18 | 0.99 |
| Q92526 | CCT6B | 5 | 5 | 0.99 |
| O75884 | RBBP9 | 2 | 2 | 0.99 |
| O75934 | BCAS2 | 4 | 4 | 0.99 |
| P02792 | FTL | 11 | 11 | 0.99 |
| Q9NRV9 | HEBP1 | 8 | 8 | 0.99 |
| Q8N5K1 | CISD2 | 3 | 3 | 0.99 |
| Q52LJ0 | FAM98B | 8 | 5 | 0.99 |
| Q9H4G4 | GLIPR2 | 2 | 2 | 0.99 |
| Q9Y508 | RNF114 | 4 | 4 | 0.99 |
| P55036 | PSMD4 | 10 | 10 | 0.99 |
| P49915 | GMPS | 11 | 11 | 0.99 |
| Q8N573 | OXR1 | 3 | 3 | 0.99 |
| Q99873 | PRMT1 | 14 | 14 | 0.99 |
| Q9HCC0 | MCCC2 | 6 | 6 | 0.99 |
| Q96AY3 | FKBP10 | 27 | 26 | 0.99 |
| P31943 | HNRNPH1 | 14 | 8 | 0.99 |
| P00450 | CP | 11 | 11 | 0.99 |
| Q86UE4 | MTDH | 19 | 19 | 0.99 |
| B5ME19 | EIF3CL | 27 | 27 | 0.99 |
| Q66K14 | TBC1D9B | 5 | 5 | 0.98 |
| P82979 | SARNP | 6 | 6 | 0.98 |
| Q9Y237 | PIN4 | 4 | 4 | 0.98 |
| P25789 | PSMA4 | 9 | 9 | 0.98 |
| P22087 | FBL | 13 | 13 | 0.98 |
| Q15459 | SF3A1 | 17 | 17 | 0.98 |
| O15230 | LAMA5 | 3 | 3 | 0.98 |
| Q8N163 | CCAR2 | 17 | 17 | 0.98 |
| Q9Y2S6 | TMA7 | 3 | 3 | 0.98 |
| Q14678 | KANK1 | 3 | 2 | 0.98 |
| P51991 | HNRNPA3 | 19 | 16 | 0.98 |
| Q9BYM8 | RBCK1 | 2 | 2 | 0.98 |
| P55769 | SNU13 | 8 | 8 | 0.98 |
| P21796 | VDAC1 | 15 | 15 | 0.98 |
| P51148 | RAB5C | 7 | 4 | 0.98 |

| Accession | Name | # Peptides | # Unique Peptides | Abundance Ratio: WWP2 / control |
|---|---|---|---|---|
| Q8TDX7 | NEK7 | 5 | 5 | 0.98 |
| Q9NSE4 | IARS2 | 9 | 9 | 0.98 |
| O75533 | SF3B1 | 28 | 28 | 0.98 |
| P00338 | LDHA | 23 | 20 | 0.98 |
| P08123 | COL1A2 | 94 | 93 | 0.98 |
| O14979 | HNRNPDL | 13 | 12 | 0.98 |
| P31749 | AKT1 | 8 | 8 | 0.98 |
| Q5VZF2 | MBNL2 | 5 | 5 | 0.98 |
| P84157 | MXRA7 | 5 | 5 | 0.98 |
| Q9P299 | COPZ2 | 7 | 7 | 0.98 |
| O43149 | ZZEF1 | 2 | 2 | 0.98 |
| Q9UNW1 | MINPP1 | 3 | 3 | 0.98 |
| P09651 | HNRNPA1 | 18 | 14 | 0.98 |
| Q9BQG0 | MYBBP1A | 12 | 12 | 0.98 |
| Q9BXP5 | SRRT | 13 | 13 | 0.98 |
| Q9UK32 | RPS6KA6 | 2 | 2 | 0.98 |
| P25787 | PSMA2 | 9 | 9 | 0.98 |
| P22061 | PCMT1 | 8 | 8 | 0.98 |
| P61086 | UBE2K | 7 | 7 | 0.98 |
| P04075 | ALDOA | 33 | 27 | 0.98 |
| P18754 | RCC1 | 4 | 4 | 0.98 |
| O60784 | TOM1 | 7 | 7 | 0.98 |
| Q9BS40 | LXN | 3 | 3 | 0.98 |
| Q15782 | CHI3L2 | 9 | 9 | 0.98 |
| Q9ULV4 | CORO1C | 22 | 22 | 0.98 |
| O75306 | NDUFS2 | 7 | 7 | 0.98 |
| Q15257 | PTPA | 8 | 8 | 0.98 |
| Q92922 | SMARCC1 | 3 | 3 | 0.98 |
| P14550 | AKR1A1 | 14 | 14 | 0.98 |
| Q9UL25 | RAB21 | 9 | 9 | 0.98 |
| Q15029 | EFTUD2 | 19 | 18 | 0.98 |
| P37837 | TALDO1 | 18 | 18 | 0.98 |
| Q9UHP3 | USP25 | 2 | 2 | 0.98 |
| Q14444 | CAPRIN1 | 9 | 9 | 0.98 |
| Q969E2 | SCAMP4 | 2 | 2 | 0.98 |
| P61421 | ATP6V0D1 | 4 | 4 | 0.98 |
| Q15424 | SAFB | 19 | 11 | 0.98 |
| Q9BRT3 | MIEN1 | 2 | 2 | 0.98 |
| Q16658 | FSCN1 | 27 | 27 | 0.98 |
| Q96I15 | SCLY | 2 | 2 | 0.98 |
| E9PAV3 | NACA | 6 | 6 | 0.98 |
| Q9BUJ2 | HNRNPUL1 | 16 | 16 | 0.98 |
| P51665 | PSMD7 | 7 | 7 | 0.98 |
| P37198 | NUP62 | 3 | 3 | 0.98 |

| Accession | Name | # Peptides | # Unique Peptides | Abundance Ratio: WWP2 / control |
|---|---|---|---|---|
| P23246 | SFPQ | 31 | 29 | 0.98 |
| P08134 | RHOC | 10 | 4 | 0.98 |
| Q9NQG1 | MANBAL | 2 | 2 | 0.98 |
| O43809 | NUDT21 | 10 | 10 | 0.98 |
| O60524 | NEMF | 3 | 3 | 0.98 |
| Q9Y3U8 | RPL36 | 6 | 6 | 0.98 |
| O00115 | DNASE2 | 4 | 4 | 0.98 |
| Q9UBU9 | NXF1 | 4 | 4 | 0.98 |
| P30154 | PPP2R1B | 5 | 5 | 0.98 |
| Q8TER0 | SNED1 | 4 | 4 | 0.98 |
| P18206 | VCL | 66 | 66 | 0.98 |
| Q8WXF7 | ATL1 | 6 | 6 | 0.98 |
| Q96A26 | FAM162A | 3 | 3 | 0.98 |
| P49411 | TUFM | 13 | 13 | 0.98 |
| P11766 | ADH5 | 14 | 14 | 0.98 |
| Q96EP5 | DAZAP1 | 4 | 4 | 0.98 |
| P29144 | TPP2 | 22 | 22 | 0.98 |
| Q8NB25 | FAM184A | 2 | 2 | 0.98 |
| Q9Y3F4 | STRAP | 12 | 12 | 0.98 |
| P0DME0 | SETSIP | 7 | 7 | 0.98 |
| O75503 | CLN5 | 2 | 2 | 0.98 |
| P62942 | FKBP1A | 5 | 5 | 0.98 |
| P30084 | ECHS1 | 11 | 11 | 0.98 |
| O75832 | PSMD10 | 7 | 7 | 0.98 |
| Q9UMX5 | NENF | 5 | 5 | 0.98 |
| P13797 | PLS3 | 29 | 26 | 0.98 |
| Q14938 | NFIX | 6 | 4 | 0.98 |
| Q9UHX1 | PUF60 | 13 | 13 | 0.98 |
| Q8WTW3 | COG1 | 2 | 2 | 0.98 |
| P40925 | MDH1 | 18 | 18 | 0.98 |
| Q99836 | MYD88 | 4 | 4 | 0.98 |
| Q14247 | CTTN | 41 | 41 | 0.98 |
| P49903 | SEPHS1 | 2 | 2 | 0.98 |
| P01116 | KRAS | 5 | 2 | 0.98 |
| P84098 | RPL19 | 10 | 10 | 0.98 |
| P16402 | H1-3 | 17 | 6 | 0.98 |
| Q12905 | ILF2 | 14 | 14 | 0.98 |
| P37108 | SRP14 | 7 | 7 | 0.98 |
| Q9Y2T2 | AP3M1 | 8 | 8 | 0.98 |
| P27701 | CD82 | 2 | 2 | 0.98 |
| P30837 | ALDH1B1 | 4 | 1 | 0.98 |
| Q9H299 | SH3BGRL3 | 7 | 7 | 0.98 |
| Q7Z304 | MAMDC2 | 16 | 16 | 0.98 |
| Q9UEE9 | CFDP1 | 2 | 2 | 0.98 |

| Accession | Name | # Peptides | # Unique Peptides | Abundance Ratio: WWP2 / control |
|---|---|---|---|---|
| P61956 | SUMO2 | 2 | 1 | 0.98 |
| P38159 | RBMX | 27 | 10 | 0.98 |
| Q01105 | SET | 10 | 10 | 0.98 |
| Q16363 | LAMA4 | 18 | 18 | 0.98 |
| P48739 | PITPNB | 7 | 5 | 0.98 |
| Q3LXA3 | TKFC | 8 | 8 | 0.98 |
| P07477 | PRSS1 | 2 | 2 | 0.98 |
| Q9Y5P6 | GMPPB | 9 | 9 | 0.98 |
| O14579 | COPE | 10 | 10 | 0.98 |
| P10606 | COX5B | 6 | 6 | 0.98 |
| Q6UW63 | POGLUT2 | 2 | 2 | 0.98 |
| O95218 | ZRANB2 | 9 | 9 | 0.98 |
| O94905 | ERLIN2 | 13 | 9 | 0.98 |
| P52209 | PGD | 16 | 16 | 0.98 |
| Q14344 | GNA13 | 6 | 5 | 0.98 |
| Q71DI3 | H3C13 | 10 | 3 | 0.98 |
| Q8N3V7 | SYNPO | 5 | 5 | 0.98 |
| P47756 | CAPZB | 16 | 16 | 0.98 |
| P0C0S5 | H2AZ1 | 6 | 4 | 0.98 |
| Q08209 | PPP3CA | 14 | 8 | 0.98 |
| O43301 | HSPA12A | 2 | 2 | 0.98 |
| P61106 | RAB14 | 12 | 12 | 0.98 |
| Q71UI9 | H2AZ2 | 6 | 4 | 0.98 |
| P02768 | ALB | 4 | 4 | 0.98 |
| O60832 | DKC1 | 13 | 13 | 0.98 |
| O14656 | TOR1A | 2 | 2 | 0.98 |
| Q07020 | RPL18 | 6 | 6 | 0.98 |
| P19838 | NFKB1 | 7 | 7 | 0.98 |
| Q9Y3C6 | PPIL1 | 6 | 6 | 0.98 |
| P55084 | HADHB | 15 | 15 | 0.98 |
| P12004 | PCNA | 11 | 11 | 0.98 |
| P23526 | AHCY | 18 | 18 | 0.98 |
| Q8IXM2 | BAP18 | 3 | 3 | 0.98 |
| O15372 | EIF3H | 10 | 10 | 0.98 |
| P22392 | NME2 | 10 | 5 | 0.98 |
| O14983 | ATP2A1 | 11 | 11 | 0.98 |
| Q9C0E8 | LNPK | 8 | 8 | 0.98 |
| P61758 | VBP1 | 6 | 6 | 0.98 |
| Q96GX9 | APIP | 3 | 3 | 0.98 |
| Q15700 | DLG2 | 2 | 1 | 0.98 |
| Q9Y5K3 | PCYT1B | 3 | 3 | 0.98 |
| Q9NQW7 | XPNPEP1 | 12 | 11 | 0.98 |
| Q6P474 | PDXDC2P | 3 | 3 | 0.98 |
| Q5UCC4 | EMC10 | 3 | 3 | 0.98 |

| Accession | Name | # Peptides | # Unique Peptides | Abundance Ratio: WWP2 / control |
|---|---|---|---|---|
| Q03113 | GNA12 | 2 | 1 | 0.98 |
| P55884 | EIF3B | 18 | 18 | 0.98 |
| Q08752 | PPID | 5 | 4 | 0.98 |
| Q13404 | UBE2V1 | 8 | 4 | 0.98 |
| Q10713 | PMPCA | 3 | 3 | 0.98 |
| Q9NR99 | MXRA5 | 40 | 40 | 0.98 |
| Q9Y376 | CAB39 | 7 | 6 | 0.98 |
| Q6UVK1 | CSPG4 | 58 | 58 | 0.98 |
| P28161 | GSTM2 | 10 | 8 | 0.98 |
| P78330 | PSPH | 3 | 3 | 0.98 |
| O43719 | HTATSF1 | 7 | 7 | 0.98 |
| O95861 | BPNT1 | 13 | 13 | 0.98 |
| P48426 | PIP4K2A | 6 | 5 | 0.98 |
| P40763 | STAT3 | 16 | 16 | 0.98 |
| Q9UKG1 | APPL1 | 5 | 5 | 0.98 |
| Q13367 | AP3B2 | 3 | 3 | 0.98 |
| P12814 | ACTN1 | 49 | 30 | 0.98 |
| Q9NRW1 | RAB6B | 2 | 2 | 0.98 |
| O43670 | ZNF207 | 5 | 5 | 0.98 |
| Q14318 | FKBP8 | 8 | 8 | 0.98 |
| P25686 | DNAJB2 | 2 | 2 | 0.98 |
| P62701 | RPS4X | 16 | 7 | 0.98 |
| Q6XE24 | RBMS3 | 2 | 2 | 0.98 |
| P62266 | RPS23 | 6 | 6 | 0.98 |
| P26639 | TARS1 | 19 | 18 | 0.98 |
| Q9NYU1 | UGGT2 | 5 | 3 | 0.98 |
| P05067 | APP | 4 | 4 | 0.98 |
| Q13435 | SF3B2 | 19 | 19 | 0.98 |
| Q04721 | NOTCH2 | 4 | 3 | 0.98 |
| P99999 | CYCS | 5 | 5 | 0.98 |
| O00764 | PDXK | 10 | 10 | 0.98 |
| A2A3N6 | PIPSL | 3 | 3 | 0.98 |
| P38919 | EIF4A3 | 17 | 13 | 0.98 |
| Q8IXB1 | DNAJC10 | 8 | 8 | 0.98 |
| P55145 | MANF | 9 | 9 | 0.98 |
| O94874 | UFL1 | 20 | 19 | 0.98 |
| O95486 | SEC24A | 7 | 5 | 0.98 |
| P22413 | ENPP1 | 13 | 13 | 0.98 |
| P07900 | HSP90AA1 | 47 | 27 | 0.98 |
| Q96Q11 | TRNT1 | 3 | 3 | 0.98 |
| O95140 | MFN2 | 3 | 3 | 0.98 |
| P47985 | UQCRFS1 | 4 | 4 | 0.98 |
| P62263 | RPS14 | 9 | 9 | 0.97 |
| Q96MU7 | YTHDC1 | 3 | 3 | 0.97 |

| Accession | Name | # Peptides | # Unique Peptides | Abundance Ratio: WWP2 / control |
|---|---|---|---|---|
| Q8NHP1 | AKR7L | 3 | 3 | 0.97 |
| P14866 | HNRNPL | 18 | 17 | 0.97 |
| P49137 | MAPKAPK2 | 5 | 5 | 0.97 |
| P61254 | RPL26 | 11 | 2 | 0.97 |
| Q5VWX1 | KHDRBS2 | 3 | 3 | 0.97 |
| O75915 | ARL6IP5 | 4 | 4 | 0.97 |
| Q9UNE7 | STUB1 | 6 | 6 | 0.97 |
| P62995 | TRA2B | 6 | 4 | 0.97 |
| Q16644 | MAPKAPK3 | 3 | 3 | 0.97 |
| P20340 | RAB6A | 9 | 9 | 0.97 |
| P17844 | DDX5 | 24 | 16 | 0.97 |
| Q9C035 | TRIM5 | 2 | 2 | 0.97 |
| P06733 | ENO1 | 32 | 29 | 0.97 |
| P38435 | GGCX | 2 | 2 | 0.97 |
| P10253 | GAA | 18 | 18 | 0.97 |
| Q9Y333 | LSM2 | 4 | 4 | 0.97 |
| O75947 | ATP5PD | 6 | 6 | 0.97 |
| P55263 | ADK | 15 | 15 | 0.97 |
| Q9Y673 | ALG5 | 2 | 2 | 0.97 |
| Q9Y512 | SAMM50 | 6 | 6 | 0.97 |
| P50402 | EMD | 13 | 13 | 0.97 |
| Q14697 | GANAB | 39 | 39 | 0.97 |
| O75438 | NDUFB1 | 2 | 2 | 0.97 |
| Q14103 | HNRNPD | 14 | 14 | 0.97 |
| Q9UH62 | ARMCX3 | 4 | 4 | 0.97 |
| P35637 | FUS | 12 | 10 | 0.97 |
| Q9ULX6 | AKAP8L | 3 | 3 | 0.97 |
| Q15853 | USF2 | 2 | 2 | 0.97 |
| Q15155 | NOMO1 | 16 | 16 | 0.97 |
| Q9NWV8 | BABAM1 | 2 | 2 | 0.97 |
| P69849 | NOMO3 | 14 | 14 | 0.97 |
| Q96IX5 | ATP5MK | 3 | 3 | 0.97 |
| Q9UNZ2 | NSFL1C | 16 | 16 | 0.97 |
| Q2TAY7 | SMU1 | 9 | 9 | 0.97 |
| Q5JPE7 | NOMO2 | 15 | 15 | 0.97 |
| P31150 | GDI1 | 19 | 12 | 0.97 |
| Q86VP6 | CAND1 | 33 | 33 | 0.97 |
| P22415 | USF1 | 2 | 2 | 0.97 |
| Q92621 | NUP205 | 9 | 9 | 0.97 |
| P49257 | LMAN1 | 24 | 24 | 0.97 |
| P47897 | QARS1 | 24 | 24 | 0.97 |
| P49458 | SRP9 | 4 | 4 | 0.97 |
| P34949 | MPI | 8 | 8 | 0.97 |
| O15371 | EIF3D | 11 | 11 | 0.97 |

| Accession | Name | # Peptides | # Unique Peptides | Abundance Ratio: WWP2 / control |
|---|---|---|---|---|
| P42773 | CDKN2C | 5 | 5 | 0.97 |
| Q9UKU9 | ANGPTL2 | 2 | 2 | 0.97 |
| Q9UPT5 | EXOC7 | 2 | 2 | 0.97 |
| Q9Y3P9 | RABGAP1 | 3 | 3 | 0.97 |
| Q9H6Z4 | RANBP3 | 4 | 4 | 0.97 |
| Q9NPQ8 | RIC8A | 7 | 7 | 0.97 |
| Q14254 | FLOT2 | 8 | 8 | 0.97 |
| P48444 | ARCN1 | 23 | 23 | 0.97 |
| P17858 | PFKL | 25 | 18 | 0.97 |
| P84090 | ERH | 7 | 7 | 0.97 |
| Q96KC8 | DNAJC1 | 4 | 4 | 0.97 |
| P08708 | RPS17 | 6 | 6 | 0.97 |
| Q15738 | NSDHL | 3 | 3 | 0.97 |
| P31949 | S100A11 | 5 | 5 | 0.97 |
| Q15006 | EMC2 | 4 | 4 | 0.97 |
| Q9NT62 | ATG3 | 6 | 6 | 0.97 |
| P00966 | ASS1 | 18 | 18 | 0.97 |
| Q9BY44 | EIF2A | 11 | 11 | 0.97 |
| Q96L21 | RPL10L | 4 | 4 | 0.97 |
| Q9UNS2 | COPS3 | 6 | 6 | 0.97 |
| Q9BZL1 | UBL5 | 2 | 2 | 0.97 |
| Q8NF37 | LPCAT1 | 3 | 3 | 0.97 |
| P50570 | DNM2 | 17 | 10 | 0.97 |
| O75489 | NDUFS3 | 4 | 4 | 0.97 |
| P05198 | EIF2S1 | 13 | 13 | 0.97 |
| Q5T1M5 | FKBP15 | 7 | 7 | 0.97 |
| P60981 | DSTN | 12 | 11 | 0.97 |
| Q9UI09 | NDUFA12 | 3 | 3 | 0.97 |
| O75368 | SH3BGRL | 11 | 11 | 0.97 |
| P84085 | ARF5 | 7 | 3 | 0.97 |
| Q9Y3B3 | TMED7 | 8 | 8 | 0.97 |
| Q96AC1 | FERMT2 | 24 | 24 | 0.97 |
| Q8TC12 | RDH11 | 5 | 5 | 0.97 |
| Q96AE4 | FUBP1 | 25 | 19 | 0.97 |
| Q12769 | NUP160 | 4 | 4 | 0.97 |
| P52565 | ARHGDIA | 10 | 10 | 0.97 |
| Q16851 | UGP2 | 20 | 20 | 0.97 |
| O00566 | MPHOSPH10 | 2 | 2 | 0.97 |
| Q96EM0 | L3HYPDH | 6 | 6 | 0.97 |
| Q8N9T8 | KRI1 | 2 | 2 | 0.97 |
| Q9UBV2 | SEL1L | 6 | 6 | 0.97 |
| Q9UDW1 | UQCR10 | 2 | 2 | 0.97 |
| P62913 | RPL11 | 8 | 8 | 0.97 |
| Q92896 | GLG1 | 23 | 23 | 0.97 |

| Accession | Name | # Peptides | # Unique Peptides | Abundance Ratio: WWP2 / control |
|---|---|---|---|---|
| P52597 | HNRNPF | 12 | 11 | 0.97 |
| O15212 | PFDN6 | 8 | 8 | 0.97 |
| Q10567 | AP1B1 | 23 | 9 | 0.97 |
| Q9NWB1 | RBFOX1 | 2 | 2 | 0.97 |
| Q96G03 | PGM2 | 21 | 19 | 0.97 |
| O15145 | ARPC3 | 6 | 6 | 0.97 |
| Q99460 | PSMD1 | 24 | 24 | 0.97 |
| O43251 | RBFOX2 | 2 | 2 | 0.97 |
| O75390 | CS | 11 | 11 | 0.97 |
| P54819 | AK2 | 10 | 10 | 0.97 |
| Q96KG9 | SCYL1 | 11 | 11 | 0.97 |
| A6NFN3 | RBFOX3 | 2 | 2 | 0.97 |
| Q9NR45 | NANS | 16 | 16 | 0.97 |
| Q14011 | CIRBP | 5 | 5 | 0.97 |
| Q92599 | SEPTIN8 | 9 | 6 | 0.97 |
| Q8NEZ5 | FBXO22 | 2 | 2 | 0.97 |
| P05091 | ALDH2 | 14 | 11 | 0.97 |
| Q9Y320 | TMX2 | 3 | 3 | 0.97 |
| Q9UBQ0 | VPS29 | 6 | 6 | 0.97 |
| Q92905 | COPS5 | 6 | 6 | 0.97 |
| Q96G23 | CERS2 | 3 | 3 | 0.97 |
| Q7L2H7 | EIF3M | 4 | 4 | 0.97 |
| P48556 | PSMD8 | 11 | 11 | 0.97 |
| Q9HAV0 | GNB4 | 12 | 5 | 0.97 |
| Q8N2K0 | ABHD12 | 2 | 2 | 0.97 |
| Q96HD1 | CRELD1 | 5 | 5 | 0.97 |
| Q9H2U2 | PPA2 | 6 | 6 | 0.97 |
| P35475 | IDUA | 7 | 7 | 0.97 |
| P0CB38 | PABPC4L | 2 | 2 | 0.97 |
| Q15813 | TBCE | 4 | 4 | 0.97 |
| O75170 | PPP6R2 | 2 | 2 | 0.97 |
| Q9UBI1 | COMMD3 | 2 | 2 | 0.97 |
| P78345 | RPP38 | 2 | 2 | 0.97 |
| P63010 | AP2B1 | 27 | 13 | 0.97 |
| Q9Y3L5 | RAP2C | 2 | 2 | 0.97 |
| Q9UGJ0 | PRKAG2 | 2 | 2 | 0.97 |
| Q9H8Y8 | GORASP2 | 8 | 8 | 0.97 |
| Q03252 | LMNB2 | 43 | 40 | 0.97 |
| P07737 | PFN1 | 9 | 9 | 0.97 |
| P05556 | ITGB1 | 21 | 21 | 0.97 |
| P61088 | UBE2N | 9 | 9 | 0.97 |
| P11217 | PYGM | 14 | 1 | 0.97 |
| Q13148 | TARDBP | 9 | 9 | 0.97 |
| Q14919 | DRAP1 | 4 | 4 | 0.97 |

| Accession | Name | # Peptides | # Unique Peptides | Abundance Ratio: WWP2 / control |
|---|---|---|---|---|
| P28062 | PSMB8 | 8 | 8 | 0.97 |
| Q13151 | HNRNPA0 | 10 | 10 | 0.97 |
| Q9Y6I3 | EPN1 | 6 | 6 | 0.97 |
| Q9P0V9 | SEPTIN10 | 8 | 6 | 0.97 |
| Q53GS9 | USP39 | 4 | 4 | 0.97 |
| P20674 | COX5A | 5 | 5 | 0.97 |
| P40429 | RPL13A | 15 | 15 | 0.97 |
| P10114 | RAP2A | 2 | 2 | 0.97 |
| P14927 | UQCRB | 4 | 4 | 0.97 |
| Q96CW1 | AP2M1 | 16 | 16 | 0.97 |
| Q9Y678 | COPG1 | 24 | 23 | 0.97 |
| P13861 | PRKAR2A | 13 | 11 | 0.97 |
| P07910 | HNRNPC | 17 | 10 | 0.97 |
| Q9H9A6 | LRRC40 | 3 | 3 | 0.97 |
| Q9NZZ3 | CHMP5 | 4 | 4 | 0.97 |
| Q01469 | FABP5 | 10 | 10 | 0.97 |
| O75844 | ZMPSTE24 | 7 | 7 | 0.97 |
| P18124 | RPL7 | 20 | 20 | 0.97 |
| O75251 | NDUFS7 | 3 | 3 | 0.97 |
| O00567 | NOP56 | 14 | 14 | 0.97 |
| Q13838 | DDX39B | 20 | 9 | 0.97 |
| Q5SSJ5 | HP1BP3 | 23 | 23 | 0.97 |
| Q969H8 | MYDGF | 6 | 6 | 0.97 |
| Q14739 | LBR | 2 | 2 | 0.97 |
| Q13228 | SELENBP1 | 19 | 19 | 0.97 |
| O15498 | YKT6 | 10 | 10 | 0.97 |
| Q9NUI1 | DECR2 | 5 | 5 | 0.97 |
| Q99442 | SEC62 | 6 | 6 | 0.97 |
| P30153 | PPP2R1A | 19 | 19 | 0.97 |
| P06744 | GPI | 26 | 25 | 0.97 |
| P62330 | ARF6 | 3 | 2 | 0.97 |
| O43237 | DYNC1LI2 | 12 | 12 | 0.97 |
| P35609 | ACTN2 | 12 | 1 | 0.97 |
| P01210 | PENK | 4 | 4 | 0.97 |
| P68431 | H3C1 | 9 | 2 | 0.97 |
| Q13242 | SRSF9 | 6 | 6 | 0.97 |
| P55854 | SUMO3 | 2 | 1 | 0.97 |
| Q15075 | EEA1 | 63 | 63 | 0.97 |
| O95373 | IPO7 | 13 | 13 | 0.97 |
| Q16543 | CDC37 | 12 | 12 | 0.97 |
| P00505 | GOT2 | 18 | 18 | 0.97 |
| P19971 | TYMP | 10 | 10 | 0.97 |
| A6NMY6 | ANXA2P2 | 28 | 28 | 0.97 |
| Q5TEC6 | H3-2 | 7 | 2 | 0.97 |

| Accession | Name | # Peptides | # Unique Peptides | Abundance Ratio: WWP2 / control |
|---|---|---|---|---|
| P46977 | STT3A | 15 | 13 | 0.97 |
| O60506 | SYNCRIP | 24 | 16 | 0.97 |
| Q16695 | H3-4 | 9 | 2 | 0.97 |
| P06280 | GLA | 3 | 3 | 0.97 |
| Q9UKY7 | CDV3 | 8 | 8 | 0.97 |
| P09669 | COX6C | 4 | 4 | 0.97 |
| P50479 | PDLIM4 | 13 | 13 | 0.97 |
| P46782 | RPS5 | 10 | 10 | 0.97 |
| Q68EM7 | ARHGAP17 | 8 | 8 | 0.97 |
| P17275 | JUNB | 5 | 5 | 0.97 |
| P82930 | MRPS34 | 2 | 2 | 0.97 |
| P50395 | GDI2 | 25 | 18 | 0.97 |
| O94832 | MYO1D | 29 | 29 | 0.97 |
| Q9H3K6 | BOLA2 | 6 | 6 | 0.97 |
| Q10471 | GALNT2 | 17 | 16 | 0.97 |
| P39019 | RPS19 | 14 | 14 | 0.97 |
| P13674 | P4HA1 | 32 | 32 | 0.97 |
| Q8N129 | CNPY4 | 3 | 3 | 0.97 |
| P51570 | GALK1 | 13 | 13 | 0.96 |
| Q9UG63 | ABCF2 | 7 | 7 | 0.96 |
| P61247 | RPS3A | 28 | 28 | 0.96 |
| Q8NBI5 | SLC43A3 | 2 | 2 | 0.96 |
| O75396 | SEC22B | 9 | 9 | 0.96 |
| P30101 | PDIA3 | 42 | 42 | 0.96 |
| Q8IWA5 | SLC44A2 | 4 | 4 | 0.96 |
| Q9NUK0 | MBNL3 | 3 | 3 | 0.96 |
| P62140 | PPP1CB | 11 | 4 | 0.96 |
| Q9NYL4 | FKBP11 | 2 | 2 | 0.96 |
| P62750 | RPL23A | 13 | 13 | 0.96 |
| P39687 | ANP32A | 9 | 7 | 0.96 |
| P49721 | PSMB2 | 5 | 5 | 0.96 |
| Q96AG4 | LRRC59 | 13 | 13 | 0.96 |
| P42566 | EPS15 | 4 | 4 | 0.96 |
| Q99623 | PHB2 | 18 | 18 | 0.96 |
| P47989 | XDH | 2 | 2 | 0.96 |
| Q7L311 | ARMCX2 | 5 | 5 | 0.96 |
| Q9Y2R0 | COA3 | 3 | 3 | 0.96 |
| P46379 | BAG6 | 6 | 6 | 0.96 |
| P61019 | RAB2A | 13 | 6 | 0.96 |
| O14646 | CHD1 | 2 | 1 | 0.96 |
| P06748 | NPM1 | 13 | 13 | 0.96 |
| P61313 | RPL15 | 12 | 12 | 0.96 |
| P26640 | VARS1 | 30 | 30 | 0.96 |
| P62820 | RAB1A | 11 | 6 | 0.96 |

| Accession | Name | # Peptides | # Unique Peptides | Abundance Ratio: WWP2 / control |
|---|---|---|---|---|
| Q12996 | CSTF3 | 6 | 6 | 0.96 |
| Q01844 | EWSR1 | 5 | 5 | 0.96 |
| O00241 | SIRPB1 | 3 | 3 | 0.96 |
| P49321 | NASP | 12 | 12 | 0.96 |
| Q15005 | SPCS2 | 10 | 10 | 0.96 |
| P98066 | TNFAIP6 | 2 | 2 | 0.96 |
| Q14166 | TTLL12 | 7 | 7 | 0.96 |
| P46778 | RPL21 | 9 | 9 | 0.96 |
| Q9H477 | RBKS | 2 | 2 | 0.96 |
| P63244 | RACK1 | 21 | 21 | 0.96 |
| P46776 | RPL27A | 5 | 5 | 0.96 |
| P53680 | AP2S1 | 4 | 4 | 0.96 |
| P27797 | CALR | 26 | 26 | 0.96 |
| P0DN76 | U2AF1L5 | 6 | 6 | 0.96 |
| O00461 | GOLIM4 | 10 | 10 | 0.96 |
| Q8NHW5 | RPLP0P6 | 8 | 8 | 0.96 |
| P39656 | DDOST | 14 | 14 | 0.96 |
| Q13200 | PSMD2 | 19 | 19 | 0.96 |
| O75822 | EIF3J | 9 | 9 | 0.96 |
| Q86Y82 | STX12 | 5 | 5 | 0.96 |
| Q14457 | BECN1 | 2 | 2 | 0.96 |
| Q9P1W8 | SIRPG | 2 | 2 | 0.96 |
| O75347 | TBCA | 12 | 12 | 0.96 |
| Q15434 | RBMS2 | 2 | 2 | 0.96 |
| P27144 | AK4 | 7 | 7 | 0.96 |
| P18669 | PGAM1 | 21 | 15 | 0.96 |
| P05997 | COL5A2 | 48 | 46 | 0.96 |
| P35237 | SERPINB6 | 14 | 13 | 0.96 |
| P35813 | PPM1A | 6 | 4 | 0.96 |
| P62081 | RPS7 | 9 | 9 | 0.96 |
| O95782 | AP2A1 | 34 | 23 | 0.96 |
| Q9BY32 | ITPA | 4 | 4 | 0.96 |
| Q15370 | ELOB | 5 | 5 | 0.96 |
| P02452 | COL1A1 | 108 | 103 | 0.96 |
| Q9BS26 | ERP44 | 12 | 12 | 0.96 |
| P56381 | ATP5F1E | 3 | 3 | 0.96 |
| P61353 | RPL27 | 9 | 9 | 0.96 |
| Q00765 | REEP5 | 9 | 9 | 0.96 |
| P09960 | LTA4H | 24 | 24 | 0.96 |
| Q9UKM7 | MAN1B1 | 4 | 4 | 0.96 |
| P28290 | ITPRID2 | 2 | 2 | 0.96 |
| Q99829 | CPNE1 | 8 | 8 | 0.96 |
| Q96BQ5 | CCDC127 | 2 | 2 | 0.96 |
| Q13263 | TRIM28 | 21 | 21 | 0.96 |

| Accession | Name | # Peptides | # Unique Peptides | Abundance Ratio: WWP2 / control |
|---|---|---|---|---|
| P26232 | CTNNA2 | 4 | 4 | 0.96 |
| P62829 | RPL23 | 10 | 10 | 0.96 |
| Q9H4A6 | GOLPH3 | 2 | 2 | 0.96 |
| Q9NUQ9 | CYRIB | 3 | 2 | 0.96 |
| P04843 | RPN1 | 31 | 31 | 0.96 |
| Q92769 | HDAC2 | 6 | 3 | 0.96 |
| P56385 | ATP5ME | 3 | 3 | 0.96 |
| Q9BWF3 | RBM4 | 6 | 6 | 0.96 |
| Q9BZG1 | RAB34 | 7 | 7 | 0.96 |
| P17301 | ITGA2 | 6 | 6 | 0.96 |
| Q9Y3E1 | HDGFL3 | 5 | 4 | 0.96 |
| P43034 | PAFAH1B1 | 13 | 13 | 0.96 |
| P13804 | ETFA | 8 | 8 | 0.96 |
| Q9NRX4 | PHPT1 | 9 | 9 | 0.96 |
| Q9NX63 | CHCHD3 | 10 | 10 | 0.96 |
| O15355 | PPM1G | 9 | 9 | 0.96 |
| P47914 | RPL29 | 7 | 7 | 0.96 |
| Q14240 | EIF4A2 | 23 | 10 | 0.96 |
| Q9BVG4 | PBDC1 | 2 | 2 | 0.96 |
| Q9Y6B6 | SAR1B | 5 | 2 | 0.96 |
| P25940 | COL5A3 | 8 | 8 | 0.96 |
| P26373 | RPL13 | 10 | 10 | 0.96 |
| P42677 | RPS27 | 4 | 2 | 0.96 |
| Q96BM9 | ARL8A | 2 | 1 | 0.96 |
| Q9Y3I0 | RTCB | 21 | 21 | 0.96 |
| Q07954 | LRP1 | 135 | 135 | 0.96 |
| Q9NQ39 | RPS10P5 | 9 | 8 | 0.96 |
| Q9UN70 | PCDHGC3 | 5 | 3 | 0.96 |
| O60522 | TDRD6 | 2 | 2 | 0.96 |
| P30050 | RPL12 | 9 | 9 | 0.96 |
| P26641 | EEF1G | 23 | 23 | 0.96 |
| Q99729 | HNRNPAB | 9 | 8 | 0.96 |
| P20908 | COL5A1 | 37 | 30 | 0.96 |
| Q9NXG2 | THUMPD1 | 7 | 7 | 0.96 |
| Q6IBS0 | TWF2 | 14 | 11 | 0.96 |
| P62851 | RPS25 | 6 | 6 | 0.96 |
| P12931 | SRC | 4 | 1 | 0.96 |
| P10412 | H1-4 | 22 | 9 | 0.96 |
| Q14766 | LTBP1 | 10 | 10 | 0.96 |
| Q9BVK6 | TMED9 | 12 | 10 | 0.96 |
| O14735 | CDIPT | 4 | 4 | 0.96 |
| O43776 | NARS1 | 18 | 18 | 0.96 |
| Q08211 | DHX9 | 37 | 37 | 0.96 |
| P27658 | COL8A1 | 6 | 6 | 0.96 |

| Accession | Name | # Peptides | # Unique Peptides | Abundance Ratio: WWP2 / control |
|---|---|---|---|---|
| Q07157 | TJP1 | 11 | 11 | 0.96 |
| O15042 | U2SURP | 6 | 6 | 0.96 |
| Q9H0W5 | CCDC8 | 5 | 5 | 0.96 |
| P21108 | PRPS1L1 | 4 | 2 | 0.96 |
| P21246 | PTN | 2 | 2 | 0.96 |
| Q9NZU5 | LMCD1 | 7 | 7 | 0.96 |
| Q9UIC8 | LCMT1 | 2 | 2 | 0.96 |
| O95810 | CAVIN2 | 4 | 4 | 0.96 |
| Q53EP0 | FNDC3B | 19 | 19 | 0.96 |
| P46976 | GYG1 | 6 | 6 | 0.96 |
| P51149 | RAB7A | 11 | 11 | 0.96 |
| P04844 | RPN2 | 13 | 13 | 0.96 |
| P51608 | MECP2 | 8 | 8 | 0.96 |
| P33527 | ABCC1 | 21 | 21 | 0.96 |
| P06756 | ITGAV | 23 | 23 | 0.96 |
| Q8IW45 | NAXD | 3 | 3 | 0.96 |
| Q9H089 | LSG1 | 2 | 2 | 0.96 |
| Q58FF3 | HSP90B2P | 10 | 10 | 0.96 |
| Q08AF3 | SLFN5 | 2 | 2 | 0.96 |
| Q9Y5M8 | SRPRB | 9 | 9 | 0.96 |
| P57088 | TMEM33 | 5 | 5 | 0.96 |
| P14625 | HSP90B1 | 57 | 55 | 0.96 |
| O00534 | VWA5A | 11 | 11 | 0.96 |
| Q8NBJ4 | GOLM1 | 12 | 12 | 0.96 |
| Q00587 | CDC42EP1 | 3 | 3 | 0.96 |
| Q16537 | PPP2R5E | 3 | 3 | 0.96 |
| O94903 | PLPBP | 5 | 5 | 0.96 |
| P07237 | P4HB | 40 | 40 | 0.96 |
| O43324 | EEF1E1 | 6 | 6 | 0.96 |
| O14910 | LIN7A | 2 | 2 | 0.96 |
| P00167 | CYB5A | 4 | 4 | 0.96 |
| P15880 | RPS2 | 16 | 16 | 0.96 |
| P16104 | H2AX | 10 | 5 | 0.96 |
| P40261 | NNMT | 8 | 8 | 0.96 |
| Q9Y305 | ACOT9 | 5 | 5 | 0.96 |
| Q4V9L6 | TMEM119 | 3 | 3 | 0.96 |
| O43143 | DHX15 | 21 | 20 | 0.96 |
| Q14568 | HSP90AA2P | 8 | 2 | 0.96 |
| Q9H845 | ACAD9 | 2 | 2 | 0.96 |
| O14558 | HSPB6 | 4 | 4 | 0.96 |
| O15427 | SLC16A3 | 7 | 7 | 0.96 |
| P11233 | RALA | 4 | 3 | 0.96 |
| Q12882 | DPYD | 6 | 6 | 0.96 |
| Q96F85 | CNRIP1 | 4 | 4 | 0.96 |

| Accession | Name | # Peptides | # Unique Peptides | Abundance Ratio: WWP2 / control |
|---|---|---|---|---|
| P60228 | EIF3E | 14 | 14 | 0.96 |
| O95299 | NDUFA10 | 4 | 4 | 0.96 |
| Q58FF8 | HSP90AB2P | 9 | 3 | 0.96 |
| Q92499 | DDX1 | 26 | 26 | 0.96 |
| Q05682 | CALD1 | 42 | 41 | 0.96 |
| Q15366 | PCBP2 | 9 | 3 | 0.96 |
| Q13190 | STX5 | 6 | 6 | 0.96 |
| Q8IZ81 | ELMOD2 | 2 | 2 | 0.96 |
| Q92688 | ANP32B | 7 | 5 | 0.96 |
| P46777 | RPL5 | 15 | 15 | 0.95 |
| Q13557 | CAMK2D | 16 | 12 | 0.95 |
| P62910 | RPL32 | 9 | 9 | 0.95 |
| Q6UWE0 | LRSAM1 | 4 | 4 | 0.95 |
| Q14195 | DPYSL3 | 23 | 20 | 0.95 |
| P61916 | NPC2 | 7 | 7 | 0.95 |
| Q07092 | COL16A1 | 17 | 17 | 0.95 |
| Q8N9N2 | ASCC1 | 3 | 3 | 0.95 |
| O76071 | CIAO1 | 2 | 2 | 0.95 |
| P62280 | RPS11 | 11 | 11 | 0.95 |
| P30086 | PEBP1 | 12 | 12 | 0.95 |
| Q8NBQ5 | HSD17B11 | 2 | 2 | 0.95 |
| Q92804 | TAF15 | 12 | 10 | 0.95 |
| P29590 | PML | 28 | 28 | 0.95 |
| Q15185 | PTGES3 | 6 | 6 | 0.95 |
| Q9Y316 | MEMO1 | 3 | 3 | 0.95 |
| P09493 | TPM1 | 29 | 13 | 0.95 |
| Q9BWS9 | CHID1 | 6 | 6 | 0.95 |
| O95336 | PGLS | 8 | 8 | 0.95 |
| Q8NHP8 | PLBD2 | 10 | 10 | 0.95 |
| O15400 | STX7 | 8 | 7 | 0.95 |
| P23284 | PPIB | 17 | 17 | 0.95 |
| Q8IWE2 | FAM114A1 | 14 | 14 | 0.95 |
| P05388 | RPLP0 | 11 | 11 | 0.95 |
| Q4KMQ2 | ANO6 | 4 | 4 | 0.95 |
| O43252 | PAPSS1 | 16 | 12 | 0.95 |
| Q9BVJ6 | UTP14A | 2 | 2 | 0.95 |
| P51812 | RPS6KA3 | 7 | 7 | 0.95 |
| P63241 | EIF5A | 14 | 14 | 0.95 |
| Q9UGP8 | SEC63 | 13 | 13 | 0.95 |
| O95168 | NDUFB4 | 4 | 4 | 0.95 |
| Q9H307 | PNN | 8 | 8 | 0.95 |
| P62906 | RPL10A | 11 | 11 | 0.95 |
| P13796 | LCP1 | 3 | 2 | 0.95 |
| Q9H4Z3 | PCIF1 | 2 | 2 | 0.95 |

| Accession | Name | # Peptides | # Unique Peptides | Abundance Ratio: WWP2 / control |
|---|---|---|---|---|
| Q9UBS4 | DNAJB11 | 10 | 10 | 0.95 |
| P13667 | PDIA4 | 46 | 46 | 0.95 |
| P60842 | EIF4A1 | 23 | 10 | 0.95 |
| P05114 | HMGN1 | 10 | 10 | 0.95 |
| Q9P2J5 | LARS1 | 27 | 27 | 0.95 |
| Q9Y6C9 | MTCH2 | 8 | 8 | 0.95 |
| O00743 | PPP6C | 4 | 4 | 0.95 |
| O94973 | AP2A2 | 20 | 9 | 0.95 |
| O94979 | SEC31A | 28 | 28 | 0.95 |
| P62879 | GNB2 | 12 | 3 | 0.95 |
| P63220 | RPS21 | 8 | 8 | 0.95 |
| P08865 | RPSA | 12 | 12 | 0.95 |
| Q02543 | RPL18A | 8 | 8 | 0.95 |
| Q13547 | HDAC1 | 5 | 2 | 0.95 |
| Q8N6T3 | ARFGAP1 | 8 | 8 | 0.95 |
| Q9Y3Q3 | TMED3 | 4 | 4 | 0.95 |
| P67936 | TPM4 | 39 | 20 | 0.95 |
| P61978 | HNRNPK | 22 | 22 | 0.95 |
| Q92791 | P3H4 | 6 | 6 | 0.95 |
| Q9BQ04 | RBM4B | 4 | 4 | 0.95 |
| Q32P44 | EML3 | 9 | 9 | 0.95 |
| P50583 | NUDT2 | 4 | 4 | 0.95 |
| Q00266 | MAT1A | 2 | 2 | 0.95 |
| P20339 | RAB5A | 7 | 4 | 0.95 |
| Q8WWX9 | SELENOM | 3 | 3 | 0.95 |
| O43819 | SCO2 | 2 | 2 | 0.95 |
| O15305 | PMM2 | 5 | 5 | 0.95 |
| P28370 | SMARCA1 | 3 | 2 | 0.95 |
| Q969X5 | ERGIC1 | 8 | 8 | 0.95 |
| Q9NPA8 | ENY2 | 2 | 2 | 0.95 |
| Q13492 | PICALM | 15 | 15 | 0.95 |
| P13639 | EEF2 | 42 | 41 | 0.95 |
| Q9HD45 | TM9SF3 | 7 | 7 | 0.95 |
| P18621 | RPL17 | 10 | 10 | 0.95 |
| Q9Y3Y2 | CHTOP | 3 | 3 | 0.95 |
| Q9Y314 | NOSIP | 2 | 2 | 0.95 |
| Q9NQ29 | LUC7L | 5 | 1 | 0.95 |
| Q8NBM8 | PCYOX1L | 2 | 2 | 0.95 |
| Q8IWE4 | DCUN1D3 | 3 | 3 | 0.95 |
| Q96DZ1 | ERLEC1 | 3 | 3 | 0.95 |
| P23528 | CFL1 | 16 | 14 | 0.95 |
| Q9BRK3 | MXRA8 | 4 | 4 | 0.95 |
| P51531 | SMARCA2 | 6 | 5 | 0.95 |
| Q9P291 | ARMCX1 | 2 | 2 | 0.95 |

| Accession | Name | # Peptides | # Unique Peptides | Abundance Ratio: WWP2 / control |
|---|---|---|---|---|
| P23396 | RPS3 | 16 | 16 | 0.95 |
| Q9UBF2 | COPG2 | 5 | 4 | 0.95 |
| Q9BZH6 | WDR11 | 2 | 2 | 0.95 |
| Q9BT09 | CNPY3 | 2 | 2 | 0.95 |
| Q93084 | ATP2A3 | 4 | 4 | 0.95 |
| Q9UKV8 | AGO2 | 3 | 1 | 0.95 |
| O95302 | FKBP9 | 14 | 13 | 0.95 |
| Q92901 | RPL3L | 2 | 2 | 0.95 |
| Q6P5R6 | RPL22L1 | 3 | 3 | 0.95 |
| Q9H0Q0 | CYRIA | 2 | 1 | 0.95 |
| Q14151 | SAFB2 | 13 | 5 | 0.95 |
| Q7KZF4 | SND1 | 45 | 45 | 0.95 |
| Q15293 | RCN1 | 11 | 11 | 0.95 |
| O94855 | SEC24D | 20 | 20 | 0.95 |
| Q8NBL1 | POGLUT1 | 4 | 4 | 0.95 |
| Q16270 | IGFBP7 | 5 | 5 | 0.95 |
| P42766 | RPL35 | 5 | 5 | 0.95 |
| Q9UNN8 | PROCR | 4 | 4 | 0.95 |
| P61204 | ARF3 | 9 | 5 | 0.95 |
| Q6NVV1 | RPL13AP3 | 3 | 3 | 0.95 |
| P84077 | ARF1 | 9 | 5 | 0.95 |
| P32969 | RPL9 | 8 | 8 | 0.95 |
| Q6NUQ4 | TMEM214 | 16 | 16 | 0.95 |
| Q6YP21 | KYAT3 | 5 | 5 | 0.95 |
| P55210 | CASP7 | 2 | 2 | 0.95 |
| P62888 | RPL30 | 7 | 7 | 0.95 |
| P41252 | IARS1 | 26 | 26 | 0.95 |
| Q02790 | FKBP4 | 18 | 17 | 0.95 |
| Q9UHD2 | TBK1 | 4 | 4 | 0.95 |
| Q99717 | SMAD5 | 3 | 3 | 0.95 |
| Q7Z4H8 | POGLUT3 | 7 | 7 | 0.95 |
| Q9UL26 | RAB22A | 2 | 2 | 0.95 |
| P14136 | GFAP | 2 | 1 | 0.95 |
| O75339 | CILP | 29 | 29 | 0.95 |
| O43423 | ANP32C | 2 | 2 | 0.95 |
| P22626 | HNRNPA2B1 | 22 | 21 | 0.95 |
| Q12792 | TWF1 | 14 | 11 | 0.95 |
| O60565 | GREM1 | 3 | 3 | 0.95 |
| Q9UHA4 | LAMTOR3 | 2 | 2 | 0.95 |
| Q9Y4L1 | HYOU1 | 24 | 24 | 0.95 |
| Q9BQA9 | CYBC1 | 4 | 4 | 0.94 |
| Q16186 | ADRM1 | 4 | 4 | 0.94 |
| Q8NBS9 | TXNDC5 | 19 | 19 | 0.94 |
| Q5VTU8 | ATP5F1EP2 | 4 | 4 | 0.94 |

| Accession | Name | # Peptides | # Unique Peptides | Abundance Ratio: WWP2 / control |
|---|---|---|---|---|
| Q9H0U4 | RAB1B | 10 | 5 | 0.94 |
| Q9BQB6 | VKORC1 | 3 | 3 | 0.94 |
| Q02878 | RPL6 | 17 | 17 | 0.94 |
| Q00341 | HDLBP | 59 | 59 | 0.94 |
| Q8TAQ2 | SMARCC2 | 13 | 13 | 0.94 |
| P61457 | PCBD1 | 3 | 3 | 0.94 |
| P43235 | CTSK | 4 | 3 | 0.94 |
| P21980 | TGM2 | 24 | 24 | 0.94 |
| Q16890 | TPD52L1 | 3 | 3 | 0.94 |
| P30626 | SRI | 9 | 9 | 0.94 |
| Q16643 | DBN1 | 16 | 16 | 0.94 |
| P25685 | DNAJB1 | 8 | 8 | 0.94 |
| Q9UBB6 | NCDN | 4 | 4 | 0.94 |
| P42574 | CASP3 | 4 | 4 | 0.94 |
| Q15629 | TRAM1 | 6 | 6 | 0.94 |
| P67870 | CSNK2B | 3 | 3 | 0.94 |
| P49207 | RPL34 | 8 | 8 | 0.94 |
| Q9P035 | HACD3 | 7 | 7 | 0.94 |
| Q01085 | TIAL1 | 9 | 6 | 0.94 |
| O75955 | FLOT1 | 8 | 8 | 0.94 |
| Q5BKZ1 | ZNF326 | 3 | 3 | 0.94 |
| P27635 | RPL10 | 9 | 9 | 0.94 |
| P02751 | FN1 | 97 | 97 | 0.94 |
| P49588 | AARS1 | 34 | 34 | 0.94 |
| P49773 | HINT1 | 4 | 4 | 0.94 |
| Q8IYI6 | EXOC8 | 3 | 3 | 0.94 |
| P62249 | RPS16 | 14 | 14 | 0.94 |
| P48147 | PREP | 17 | 17 | 0.94 |
| P11216 | PYGB | 46 | 34 | 0.94 |
| Q15393 | SF3B3 | 27 | 27 | 0.94 |
| P49753 | ACOT2 | 5 | 5 | 0.94 |
| Q9Y536 | PPIAL4A | 5 | 4 | 0.94 |
| Q13310 | PABPC4 | 24 | 13 | 0.94 |
| Q06546 | GABPA | 2 | 2 | 0.94 |
| P32322 | PYCR1 | 9 | 9 | 0.94 |
| O95671 | ASMTL | 5 | 5 | 0.94 |
| O75592 | MYCBP2 | 4 | 4 | 0.94 |
| Q96JJ7 | TMX3 | 7 | 7 | 0.94 |
| Q8NCA5 | FAM98A | 10 | 7 | 0.94 |
| Q9P2N5 | RBM27 | 3 | 2 | 0.94 |
| Q9NX58 | LYAR | 2 | 2 | 0.94 |
| Q9NX76 | CMTM6 | 3 | 3 | 0.94 |
| Q9Y224 | RTRAF | 10 | 10 | 0.94 |
| Q96CG8 | CTHRC1 | 4 | 4 | 0.94 |

| Accession | Name | # Peptides | # Unique Peptides | Abundance Ratio: WWP2 / control |
|---|---|---|---|---|
| Q8WXF1 | PSPC1 | 7 | 5 | 0.94 |
| P53041 | PPP5C | 6 | 6 | 0.94 |
| P07951 | TPM2 | 25 | 6 | 0.94 |
| O60551 | NMT2 | 2 | 2 | 0.94 |
| Q7RTV0 | PHF5A | 3 | 3 | 0.94 |
| Q58FG0 | HSP90AA5P | 4 | 1 | 0.94 |
| P14174 | MIF | 5 | 5 | 0.94 |
| Q14257 | RCN2 | 7 | 7 | 0.94 |
| Q8IUX7 | AEBP1 | 39 | 39 | 0.94 |
| Q53S08 | RAB6D | 2 | 2 | 0.94 |
| P60763 | RAC3 | 4 | 3 | 0.94 |
| Q13643 | FHL3 | 4 | 4 | 0.94 |
| Q9Y4W2 | LAS1L | 2 | 2 | 0.94 |
| O60256 | PRPSAP2 | 4 | 4 | 0.94 |
| Q14165 | MLEC | 11 | 11 | 0.94 |
| Q15181 | PPA1 | 13 | 13 | 0.94 |
| P05141 | SLC25A5 | 13 | 5 | 0.94 |
| Q96RP9 | GFM1 | 2 | 2 | 0.94 |
| P24928 | POLR2A | 2 | 2 | 0.94 |
| Q32P51 | HNRNPA1L2 | 13 | 9 | 0.94 |
| Q8TDD1 | DDX54 | 3 | 3 | 0.94 |
| P60891 | PRPS1 | 10 | 5 | 0.94 |
| Q07960 | ARHGAP1 | 11 | 11 | 0.94 |
| Q96SI9 | STRBP | 3 | 3 | 0.94 |
| Q9Y4F1 | FARP1 | 10 | 10 | 0.94 |
| O75083 | WDR1 | 28 | 28 | 0.94 |
| P07585 | DCN | 14 | 13 | 0.94 |
| Q14554 | PDIA5 | 15 | 15 | 0.94 |
| Q969M3 | YIPF5 | 2 | 2 | 0.94 |
| P36578 | RPL4 | 28 | 27 | 0.94 |
| Q15819 | UBE2V2 | 8 | 4 | 0.94 |
| P68104 | EEF1A1 | 25 | 25 | 0.94 |
| Q9H6B4 | CLMP | 4 | 4 | 0.94 |
| A0A075B767 | PPIAL4H | 4 | 3 | 0.94 |
| P0DN37 | PPIAL4G | 4 | 3 | 0.94 |
| P35222 | CTNNB1 | 13 | 13 | 0.94 |
| P24534 | EEF1B2 | 7 | 5 | 0.94 |
| Q9UII2 | ATP5IF1 | 5 | 5 | 0.94 |
| P54727 | RAD23B | 10 | 8 | 0.94 |
| Q07065 | CKAP4 | 47 | 46 | 0.94 |
| P63000 | RAC1 | 8 | 7 | 0.94 |
| O94919 | ENDOD1 | 8 | 8 | 0.94 |
| P80303 | NUCB2 | 18 | 18 | 0.94 |
| Q14683 | SMC1A | 11 | 11 | 0.94 |

| Accession | Name | # Peptides | # Unique Peptides | Abundance Ratio: WWP2 / control |
|---|---|---|---|---|
| F5H284 | PPIAL4D | 4 | 3 | 0.94 |
| A0A0B4J2A2 | PPIAL4C | 4 | 3 | 0.94 |
| A0A075B759 | PPIAL4E | 4 | 3 | 0.94 |
| Q86TX2 | ACOT1 | 7 | 7 | 0.94 |
| Q5VTE0 | EEF1A1P5 | 24 | 24 | 0.94 |
| P18077 | RPL35A | 10 | 10 | 0.94 |
| P07686 | HEXB | 14 | 14 | 0.94 |
| Q07955 | SRSF1 | 13 | 13 | 0.94 |
| P43307 | SSR1 | 3 | 3 | 0.94 |
| P51178 | PLCD1 | 14 | 14 | 0.94 |
| Q96GC5 | MRPL48 | 2 | 2 | 0.94 |
| Q9P0L0 | VAPA | 14 | 13 | 0.94 |
| P39060 | COL18A1 | 7 | 7 | 0.94 |
| Q9NTX5 | ECHDC1 | 5 | 5 | 0.94 |
| Q96BJ3 | AIDA | 4 | 4 | 0.94 |
| O00291 | HIP1 | 11 | 11 | 0.94 |
| O75367 | MACROH2A1 | 11 | 10 | 0.94 |
| Q9H4A3 | WNK1 | 13 | 13 | 0.94 |
| Q15286 | RAB35 | 5 | 5 | 0.94 |
| Q16822 | PCK2 | 9 | 9 | 0.94 |
| Q8IZP2 | ST13P4 | 6 | 6 | 0.94 |
| P48723 | HSPA13 | 2 | 2 | 0.94 |
| Q9NRN7 | AASDHPPT | 3 | 3 | 0.94 |
| Q7Z4N8 | P4HA3 | 4 | 4 | 0.94 |
| Q92633 | LPAR1 | 2 | 2 | 0.93 |
| Q9BTC0 | DIDO1 | 2 | 2 | 0.93 |
| Q52LD8 | RFTN2 | 3 | 3 | 0.93 |
| A6NIZ1 | | 9 | 4 | 0.93 |
| Q8NBJ9 | SIDT2 | 2 | 2 | 0.93 |
| Q5VW32 | BROX | 5 | 5 | 0.93 |
| P15121 | AKR1B1 | 13 | 13 | 0.93 |
| P61221 | ABCE1 | 11 | 11 | 0.93 |
| P62424 | RPL7A | 20 | 20 | 0.93 |
| P07332 | FES | 2 | 2 | 0.93 |
| O95139 | NDUFB6 | 2 | 2 | 0.93 |
| P19623 | SRM | 10 | 10 | 0.93 |
| P07195 | LDHB | 16 | 13 | 0.93 |
| P62899 | RPL31 | 9 | 9 | 0.93 |
| O00469 | PLOD2 | 21 | 21 | 0.93 |
| Q9BV86 | NTMT1 | 2 | 2 | 0.93 |
| Q06828 | FMOD | 8 | 8 | 0.93 |
| P83881 | RPL36A | 6 | 2 | 0.93 |
| Q08426 | EHHADH | 3 | 3 | 0.93 |
| P54646 | PRKAA2 | 2 | 2 | 0.93 |

| Accession | Name | # Peptides | # Unique Peptides | Abundance Ratio: WWP2 / control |
|---|---|---|---|---|
| Q00577 | PURA | 8 | 7 | 0.93 |
| O43281 | EFS | 3 | 3 | 0.93 |
| Q8NBN3 | TMEM87A | 3 | 3 | 0.93 |
| Q15121 | PEA15 | 5 | 5 | 0.93 |
| Q9BQ39 | DDX50 | 3 | 3 | 0.93 |
| Q9H920 | RNF121 | 2 | 2 | 0.93 |
| A0A2R8Y4L2 | HNRNPA1P48 | 12 | 9 | 0.93 |
| Q14353 | GAMT | 2 | 2 | 0.93 |
| O75494 | SRSF10 | 6 | 6 | 0.93 |
| P25398 | RPS12 | 9 | 9 | 0.93 |
| P05413 | FABP3 | 8 | 8 | 0.93 |
| Q13884 | SNTB1 | 2 | 1 | 0.93 |
| Q7Z2X4 | PID1 | 2 | 2 | 0.93 |
| Q99519 | NEU1 | 4 | 4 | 0.93 |
| P61927 | RPL37 | 7 | 7 | 0.93 |
| P83731 | RPL24 | 15 | 15 | 0.93 |
| P53396 | ACLY | 33 | 33 | 0.93 |
| Q9H0E2 | TOLLIP | 5 | 5 | 0.93 |
| Q53RD9 | FBLN7 | 3 | 3 | 0.93 |
| Q9NP79 | VTA1 | 7 | 7 | 0.93 |
| Q99674 | CGREF1 | 4 | 4 | 0.93 |
| Q7Z7G0 | ABI3BP | 17 | 17 | 0.93 |
| P61224 | RAP1B | 11 | 4 | 0.93 |
| P46779 | RPL28 | 9 | 9 | 0.93 |
| Q9UBQ7 | GRHPR | 9 | 9 | 0.93 |
| Q8TBA6 | GOLGA5 | 2 | 2 | 0.93 |
| P35556 | FBN2 | 17 | 11 | 0.93 |
| Q12791 | KCNMA1 | 6 | 6 | 0.93 |
| Q12907 | LMAN2 | 12 | 12 | 0.93 |
| P10398 | ARAF | 5 | 3 | 0.93 |
| Q9BXY0 | MAK16 | 2 | 2 | 0.93 |
| Q96D71 | REPS1 | 4 | 4 | 0.93 |
| P29401 | TKT | 29 | 29 | 0.93 |
| Q8NEW0 | SLC30A7 | 2 | 2 | 0.93 |
| P04264 | KRT1 | 20 | 18 | 0.93 |
| Q8N5M4 | TTC9C | 2 | 2 | 0.93 |
| P50453 | SERPINB9 | 8 | 8 | 0.93 |
| Q92954 | PRG4 | 58 | 58 | 0.93 |
| Q9H2J4 | PDCL3 | 2 | 2 | 0.93 |
| Q15428 | SF3A2 | 3 | 3 | 0.93 |
| Q9BRK5 | SDF4 | 10 | 10 | 0.93 |
| P63261 | ACTG1 | 26 | 1 | 0.93 |
| Q3SY69 | ALDH1L2 | 15 | 15 | 0.93 |
| O95965 | ITGBL1 | 4 | 4 | 0.93 |

| Accession | Name | # Peptides | # Unique Peptides | Abundance Ratio: WWP2 / control |
|---|---|---|---|---|
| P10599 | TXN | 5 | 5 | 0.93 |
| P16989 | YBX3 | 10 | 6 | 0.93 |
| Q13283 | G3BP1 | 13 | 12 | 0.93 |
| Q9BT22 | ALG1 | 6 | 6 | 0.93 |
| Q71UM5 | RPS27L | 3 | 1 | 0.93 |
| P16070 | CD44 | 10 | 10 | 0.93 |
| Q13510 | ASAH1 | 11 | 11 | 0.93 |
| Q9NUL5 | SHFL | 3 | 3 | 0.93 |
| O95967 | EFEMP2 | 9 | 9 | 0.93 |
| Q8TCJ2 | STT3B | 12 | 10 | 0.93 |
| Q03519 | TAP2 | 2 | 2 | 0.93 |
| P62917 | RPL8 | 13 | 13 | 0.93 |
| Q9H9S3 | SEC61A2 | 6 | 6 | 0.93 |
| Q96M27 | PRRC1 | 6 | 6 | 0.93 |
| Q9H3U7 | SMOC2 | 4 | 4 | 0.92 |
| P61619 | SEC61A1 | 10 | 10 | 0.92 |
| P18085 | ARF4 | 10 | 6 | 0.92 |
| P58546 | MTPN | 5 | 5 | 0.92 |
| P50502 | ST13 | 12 | 12 | 0.92 |
| Q05193 | DNM1 | 19 | 12 | 0.92 |
| O60547 | GMDS | 19 | 19 | 0.92 |
| P50454 | SERPINH1 | 25 | 25 | 0.92 |
| Q6IS14 | EIF5AL1 | 11 | 11 | 0.92 |
| Q05639 | EEF1A2 | 13 | 13 | 0.92 |
| Q9BW91 | NUDT9 | 4 | 4 | 0.92 |
| P26885 | FKBP2 | 8 | 8 | 0.92 |
| P32321 | DCTD | 4 | 4 | 0.92 |
| Q9Y3A6 | TMED5 | 3 | 3 | 0.92 |
| Q9Y6E2 | BZW2 | 6 | 6 | 0.92 |
| Q8N5V2 | NGEF | 2 | 2 | 0.92 |
| P27482 | CALML3 | 2 | 2 | 0.92 |
| Q9H0S4 | DDX47 | 2 | 2 | 0.92 |
| Q9HD20 | ATP13A1 | 8 | 8 | 0.92 |
| P10915 | HAPLN1 | 15 | 15 | 0.92 |
| O15260 | SURF4 | 3 | 3 | 0.92 |
| O75962 | TRIO | 4 | 4 | 0.92 |
| Q9Y613 | FHOD1 | 6 | 6 | 0.92 |
| Q96PU8 | QKI | 4 | 3 | 0.92 |
| Q8IWB1 | ITPRIP | 3 | 3 | 0.92 |
| Q9BZV1 | UBXN6 | 3 | 3 | 0.92 |
| Q13425 | SNTB2 | 14 | 13 | 0.92 |
| P54577 | YARS1 | 25 | 25 | 0.92 |
| Q9H3K2 | GHITM | 3 | 3 | 0.92 |
| P51572 | BCAP31 | 23 | 23 | 0.92 |

| Accession | Name | # Peptides | # Unique Peptides | Abundance Ratio: WWP2 / control |
|---|---|---|---|---|
| Q9BTE6 | AARSD1 | 4 | 4 | 0.92 |
| Q9Y2B0 | CNPY2 | 9 | 9 | 0.92 |
| P62273 | RPS29 | 5 | 5 | 0.92 |
| A1X283 | SH3PXD2B | 10 | 9 | 0.92 |
| Q96PE7 | MCEE | 2 | 2 | 0.92 |
| Q16706 | MAN2A1 | 2 | 2 | 0.92 |
| P24593 | IGFBP5 | 6 | 6 | 0.92 |
| Q92928 | RAB1C | 3 | 2 | 0.92 |
| Q9UK22 | FBXO2 | 2 | 2 | 0.92 |
| Q9H446 | RWDD1 | 2 | 2 | 0.92 |
| O75578 | ITGA10 | 2 | 2 | 0.92 |
| Q8N335 | GPD1L | 4 | 4 | 0.92 |
| Q15363 | TMED2 | 6 | 6 | 0.92 |
| O75569 | PRKRA | 6 | 6 | 0.92 |
| Q8WXA9 | SREK1 | 2 | 2 | 0.92 |
| P32119 | PRDX2 | 9 | 8 | 0.92 |
| Q4V328 | GRIPAP1 | 5 | 5 | 0.92 |
| Q15797 | SMAD1 | 2 | 2 | 0.92 |
| O00468 | AGRN | 3 | 3 | 0.92 |
| Q9NZM1 | MYOF | 74 | 74 | 0.92 |
| P28799 | GRN | 4 | 4 | 0.92 |
| P04731 | MT1A | 2 | 1 | 0.92 |
| Q8TEW0 | PARD3 | 2 | 2 | 0.92 |
| P49591 | SARS1 | 19 | 19 | 0.92 |
| P35240 | NF2 | 2 | 2 | 0.92 |
| O14949 | UQCRQ | 2 | 2 | 0.92 |
| P12107 | COL11A1 | 35 | 28 | 0.92 |
| Q8NBJ5 | COLGALT1 | 12 | 12 | 0.92 |
| O95295 | SNAPIN | 4 | 4 | 0.92 |
| P16520 | GNB3 | 3 | 1 | 0.92 |
| P08240 | SRPRA | 12 | 12 | 0.92 |
| Q8ND76 | CCNY | 2 | 1 | 0.92 |
| Q9Y6E0 | STK24 | 6 | 2 | 0.92 |
| P15153 | RAC2 | 4 | 3 | 0.91 |
| Q9NX62 | BPNT2 | 4 | 4 | 0.91 |
| Q68CZ2 | TNS3 | 11 | 8 | 0.91 |
| Q9Y2C4 | EXOG | 3 | 3 | 0.91 |
| P16083 | NQO2 | 5 | 5 | 0.91 |
| Q8IWV7 | UBR1 | 2 | 2 | 0.91 |
| P55735 | SEC13 | 9 | 9 | 0.91 |
| O43852 | CALU | 15 | 15 | 0.91 |
| Q8N5I4 | DHRSX | 2 | 2 | 0.91 |
| O00505 | KPNA3 | 6 | 3 | 0.91 |
| P68402 | PAFAH1B2 | 5 | 5 | 0.91 |

| Accession | Name | # Peptides | # Unique Peptides | Abundance Ratio: WWP2 / control |
|---|---|---|---|---|
| Q13162 | PRDX4 | 12 | 9 | 0.91 |
| O00499 | BIN1 | 8 | 7 | 0.91 |
| P35613 | BSG | 10 | 10 | 0.91 |
| P00568 | AK1 | 18 | 18 | 0.91 |
| P31937 | HIBADH | 6 | 6 | 0.91 |
| P08572 | COL4A2 | 14 | 14 | 0.91 |
| Q14978 | NOLC1 | 8 | 8 | 0.91 |
| Q9BWJ5 | SF3B5 | 2 | 2 | 0.91 |
| Q9H0W9 | C11orf54 | 4 | 4 | 0.91 |
| Q8NBJ7 | SUMF2 | 8 | 8 | 0.91 |
| O60493 | SNX3 | 9 | 8 | 0.91 |
| Q99571 | P2RX4 | 2 | 2 | 0.91 |
| P42345 | MTOR | 2 | 2 | 0.91 |
| Q9GZV4 | EIF5A2 | 8 | 8 | 0.91 |
| Q8NFI4 | ST13P5 | 10 | 10 | 0.91 |
| O14907 | TAX1BP3 | 5 | 5 | 0.91 |
| P46939 | UTRN | 54 | 54 | 0.91 |
| Q13642 | FHL1 | 13 | 13 | 0.91 |
| P51884 | LUM | 9 | 9 | 0.91 |
| P51397 | DAP | 3 | 3 | 0.91 |
| P51571 | SSR4 | 4 | 4 | 0.91 |
| P17405 | SMPD1 | 5 | 5 | 0.91 |
| O95785 | WIZ | 2 | 2 | 0.91 |
| P60903 | S100A10 | 7 | 7 | 0.91 |
| Q9UHQ4 | BCAP29 | 5 | 5 | 0.91 |
| P55081 | MFAP1 | 3 | 2 | 0.91 |
| Q02539 | H1-1 | 10 | 4 | 0.91 |
| P04899 | GNAI2 | 16 | 10 | 0.91 |
| Q9NS86 | LANCL2 | 3 | 3 | 0.91 |
| P60953 | CDC42 | 6 | 5 | 0.91 |
| Q9H254 | SPTBN4 | 7 | 2 | 0.91 |
| P13942 | COL11A2 | 4 | 2 | 0.91 |
| P11279 | LAMP1 | 5 | 5 | 0.91 |
| P16112 | ACAN | 43 | 43 | 0.91 |
| Q8WU68 | U2AF1L4 | 2 | 2 | 0.91 |
| P62987 | UBA52 | 6 | 1 | 0.91 |
| P30041 | PRDX6 | 13 | 13 | 0.91 |
| Q99470 | SDF2 | 3 | 3 | 0.90 |
| Q9BZQ8 | NIBAN1 | 8 | 8 | 0.90 |
| P07602 | PSAP | 5 | 5 | 0.90 |
| P04216 | THY1 | 3 | 3 | 0.90 |
| P50281 | MMP14 | 7 | 7 | 0.90 |
| P62834 | RAP1A | 8 | 1 | 0.90 |
| O14787 | TNPO2 | 7 | 4 | 0.90 |

| Accession | Name | # Peptides | # Unique Peptides | Abundance Ratio: WWP2 / control |
|---|---|---|---|---|
| P60468 | SEC61B | 5 | 5 | 0.90 |
| Q5M775 | SPECC1 | 3 | 3 | 0.90 |
| Q9H8Y5 | ANKZF1 | 3 | 3 | 0.90 |
| Q8IYD1 | GSPT2 | 8 | 1 | 0.90 |
| Q8NDT2 | RBM15B | 2 | 2 | 0.90 |
| O00418 | EEF2K | 3 | 3 | 0.90 |
| P27824 | CANX | 21 | 21 | 0.90 |
| P58005 | SESN3 | 3 | 3 | 0.90 |
| P04156 | PRNP | 2 | 2 | 0.90 |
| Q13136 | PPFIA1 | 4 | 4 | 0.90 |
| P41250 | GARS1 | 25 | 25 | 0.90 |
| Q9H936 | SLC25A22 | 2 | 2 | 0.90 |
| P63098 | PPP3R1 | 2 | 2 | 0.90 |
| P24844 | MYL9 | 6 | 3 | 0.90 |
| Q01196 | RUNX1 | 2 | 2 | 0.90 |
| Q96JD6 | AKR1E2 | 2 | 2 | 0.90 |
| P24043 | LAMA2 | 3 | 3 | 0.90 |
| Q8NFQ8 | TOR1AIP2 | 3 | 2 | 0.90 |
| P61970 | NUTF2 | 2 | 2 | 0.90 |
| Q96PY5 | FMNL2 | 6 | 3 | 0.90 |
| P62244 | RPS15A | 8 | 8 | 0.90 |
| P62328 | TMSB4X | 6 | 4 | 0.90 |
| Q9Y6K8 | AK5 | 2 | 2 | 0.90 |
| Q92743 | HTRA1 | 12 | 12 | 0.90 |
| Q13362 | PPP2R5C | 2 | 2 | 0.90 |
| Q9HCJ1 | ANKH | 2 | 2 | 0.90 |
| O95573 | ACSL3 | 11 | 11 | 0.90 |
| Q9NWM8 | FKBP14 | 3 | 3 | 0.90 |
| Q53F19 | NCBP3 | 2 | 2 | 0.90 |
| Q8N8S7 | ENAH | 22 | 21 | 0.90 |
| O15460 | P4HA2 | 19 | 19 | 0.90 |
| P17813 | ENG | 9 | 9 | 0.90 |
| P07108 | DBI | 7 | 7 | 0.90 |
| P49006 | MARCKSL1 | 5 | 5 | 0.90 |
| Q14684 | RRP1B | 2 | 2 | 0.90 |
| P36871 | PGM1 | 33 | 33 | 0.90 |
| Q32P28 | P3H1 | 20 | 20 | 0.90 |
| Q9NZ01 | TECR | 8 | 8 | 0.90 |
| Q8WUY3 | PRUNE2 | 15 | 15 | 0.90 |
| P23219 | PTGS1 | 3 | 3 | 0.90 |
| Q9UKD2 | MRTO4 | 3 | 3 | 0.90 |
| O00479 | HMGN4 | 5 | 4 | 0.89 |
| Q9ULC3 | RAB23 | 4 | 4 | 0.89 |
| Q96SL4 | GPX7 | 5 | 5 | 0.89 |

| Accession | Name | # Peptides | # Unique Peptides | Abundance Ratio: WWP2 / control |
|---|---|---|---|---|
| Q9BX40 | LSM14B | 2 | 2 | 0.89 |
| P54886 | ALDH18A1 | 21 | 21 | 0.89 |
| P60174 | TPI1 | 17 | 17 | 0.89 |
| P04732 | MT1E | 3 | 1 | 0.89 |
| Q05397 | PTK2 | 3 | 3 | 0.89 |
| Q9H4F8 | SMOC1 | 8 | 8 | 0.89 |
| P05387 | RPLP2 | 7 | 6 | 0.89 |
| P08574 | CYC1 | 3 | 3 | 0.89 |
| P01023 | A2M | 6 | 6 | 0.89 |
| Q8N3E9 | PLCD3 | 2 | 2 | 0.89 |
| O75718 | CRTAP | 14 | 14 | 0.89 |
| Q9P2R3 | ANKFY1 | 15 | 15 | 0.89 |
| Q8IVL5 | P3H2 | 4 | 4 | 0.89 |
| P01011 | SERPINA3 | 6 | 6 | 0.89 |
| Q9NPE3 | NOP10 | 2 | 2 | 0.89 |
| A8MWD9 | SNRPGP15 | 3 | 3 | 0.89 |
| Q07507 | DPT | 5 | 5 | 0.89 |
| Q96T37 | RBM15 | 3 | 3 | 0.89 |
| P62308 | SNRPG | 3 | 3 | 0.89 |
| P67809 | YBX1 | 10 | 6 | 0.89 |
| Q99538 | LGMN | 3 | 3 | 0.89 |
| O15121 | DEGS1 | 2 | 2 | 0.89 |
| Q9NP74 | PALMD | 2 | 2 | 0.89 |
| Q9BST9 | RTKN | 3 | 3 | 0.89 |
| Q9NWU2 | GID8 | 2 | 2 | 0.89 |
| P61513 | RPL37A | 6 | 6 | 0.88 |
| O43854 | EDIL3 | 7 | 7 | 0.88 |
| Q6UX71 | PLXDC2 | 7 | 7 | 0.88 |
| P62873 | GNB1 | 10 | 5 | 0.88 |
| Q6K0P9 | PYHIN1 | 2 | 2 | 0.88 |
| P23470 | PTPRG | 3 | 3 | 0.88 |
| P08754 | GNAI3 | 10 | 4 | 0.88 |
| P61769 | B2M | 5 | 5 | 0.88 |
| P08651 | NFIC | 4 | 3 | 0.88 |
| Q12884 | FAP | 23 | 23 | 0.88 |
| Q99575 | POP1 | 2 | 2 | 0.88 |
| Q14112 | NID2 | 33 | 32 | 0.88 |
| P49840 | GSK3A | 2 | 2 | 0.88 |
| Q14690 | PDCD11 | 7 | 7 | 0.88 |
| Q9H6S3 | EPS8L2 | 9 | 9 | 0.87 |
| Q16666 | IFI16 | 8 | 8 | 0.87 |
| P15151 | PVR | 3 | 3 | 0.87 |
| P14678 | SNRPB | 5 | 5 | 0.87 |
| P10909 | CLU | 12 | 12 | 0.87 |

| Accession | Name | # Peptides | # Unique Peptides | Abundance Ratio: WWP2 / control |
|---|---|---|---|---|
| P63162 | SNRPN | 5 | 5 | 0.87 |
| P00736 | C1R | 6 | 6 | 0.87 |
| P02461 | COL3A1 | 109 | 108 | 0.87 |
| P30044 | PRDX5 | 5 | 5 | 0.87 |
| Q9H583 | HEATR1 | 2 | 2 | 0.87 |
| P08582 | MELTF | 25 | 25 | 0.87 |
| P14209 | CD99 | 3 | 3 | 0.87 |
| P13929 | ENO3 | 7 | 2 | 0.87 |
| Q5ZPR3 | CD276 | 2 | 2 | 0.87 |
| P50914 | RPL14 | 9 | 9 | 0.87 |
| P16234 | PDGFRA | 9 | 7 | 0.87 |
| Q7Z4W1 | DCXR | 2 | 2 | 0.87 |
| Q96A19 | CCDC102A | 5 | 4 | 0.87 |
| P02462 | COL4A1 | 9 | 9 | 0.87 |
| O76041 | NEBL | 2 | 2 | 0.87 |
| P05090 | APOD | 7 | 7 | 0.87 |
| Q6IAA8 | LAMTOR1 | 3 | 3 | 0.86 |
| O60279 | SUSD5 | 2 | 2 | 0.86 |
| Q14055 | COL9A2 | 2 | 2 | 0.86 |
| Q8IV08 | PLD3 | 11 | 11 | 0.86 |
| Q9UBG0 | MRC2 | 36 | 36 | 0.86 |
| P23381 | WARS1 | 17 | 17 | 0.86 |
| P98179 | RBM3 | 6 | 6 | 0.86 |
| O14494 | PLPP1 | 2 | 2 | 0.86 |
| P53985 | SLC16A1 | 9 | 9 | 0.86 |
| P31751 | AKT2 | 2 | 2 | 0.86 |
| Q02818 | NUCB1 | 16 | 16 | 0.86 |
| P35908 | KRT2 | 4 | 2 | 0.86 |
| Q9Y6U3 | SCIN | 16 | 16 | 0.86 |
| Q5KU26 | COLEC12 | 4 | 4 | 0.85 |
| Q53GT1 | KLHL22 | 2 | 2 | 0.85 |
| Q01628 | IFITM3 | 2 | 2 | 0.85 |
| P20774 | OGN | 5 | 5 | 0.85 |
| O75531 | BANF1 | 2 | 2 | 0.85 |
| Q9Y3Q8 | TSC22D4 | 3 | 1 | 0.85 |
| Q16585 | SGCB | 6 | 6 | 0.85 |
| O60216 | RAD21 | 4 | 4 | 0.85 |
| Q15582 | TGFBI | 32 | 32 | 0.85 |
| P0CG29 | GSTT2 | 3 | 2 | 0.85 |
| P0CG30 | GSTT2B | 3 | 2 | 0.85 |
| P22455 | FGFR4 | 2 | 1 | 0.85 |
| Q99719 | SEPTIN5 | 2 | 2 | 0.85 |
| Q5JTB6 | PLAC9 | 4 | 4 | 0.85 |
| P09486 | SPARC | 12 | 12 | 0.85 |

| Accession | Name | # Peptides | # Unique Peptides | Abundance Ratio: WWP2 / control |
|---|---|---|---|---|
| P28289 | TMOD1 | 2 | 1 | 0.84 |
| Q92629 | SGCD | 6 | 6 | 0.84 |
| O14684 | PTGES | 4 | 4 | 0.84 |
| Q06136 | KDSR | 2 | 2 | 0.84 |
| Q15758 | SLC1A5 | 4 | 4 | 0.84 |
| Q9H857 | NT5DC2 | 5 | 5 | 0.84 |
| P00325 | ADH1B | 5 | 5 | 0.84 |
| P05386 | RPLP1 | 2 | 1 | 0.84 |
| Q13123 | IK | 2 | 2 | 0.83 |
| P08243 | ASNS | 8 | 8 | 0.83 |
| P35555 | FBN1 | 99 | 93 | 0.83 |
| Q9BX66 | SORBS1 | 2 | 2 | 0.83 |
| P08253 | MMP2 | 4 | 4 | 0.83 |
| Q13541 | EIF4EBP1 | 3 | 3 | 0.83 |
| O75420 | GIGYF1 | 2 | 1 | 0.83 |
| P78324 | SIRPA | 10 | 10 | 0.82 |
| Q15113 | PCOLCE | 16 | 16 | 0.82 |
| O43175 | PHGDH | 20 | 20 | 0.82 |
| P02458 | COL2A1 | 66 | 60 | 0.82 |
| Q76M96 | CCDC80 | 13 | 13 | 0.82 |
| P48735 | IDH2 | 11 | 11 | 0.82 |
| P08493 | MGP | 3 | 3 | 0.82 |
| Q9H2U1 | DHX36 | 3 | 3 | 0.82 |
| P07339 | CTSD | 17 | 17 | 0.82 |
| O00339 | MATN2 | 22 | 22 | 0.81 |
| P08195 | SLC3A2 | 8 | 8 | 0.81 |
| P00326 | ADH1C | 4 | 4 | 0.81 |
| Q13308 | PTK7 | 7 | 7 | 0.81 |
| P09619 | PDGFRB | 9 | 7 | 0.81 |
| O94808 | GFPT2 | 16 | 10 | 0.81 |
| P21810 | BGN | 12 | 11 | 0.80 |
| O14828 | SCAMP3 | 4 | 4 | 0.80 |
| Q99983 | OMD | 3 | 3 | 0.80 |
| P55001 | MFAP2 | 4 | 4 | 0.79 |
| P50897 | PPT1 | 5 | 5 | 0.79 |
| O43556 | SGCE | 2 | 2 | 0.79 |
| P49747 | COMP | 21 | 21 | 0.79 |
| Q9BVA1 | TUBB2B | 22 | 1 | 0.79 |
| P07327 | ADH1A | 2 | 2 | 0.79 |
| Q6AZY7 | SCARA3 | 3 | 3 | 0.78 |
| Q9Y3Z3 | SAMHD1 | 4 | 4 | 0.78 |
| Q9NQA3 | WASH6P | 4 | 1 | 0.78 |
| Q6EMK4 | VASN | 7 | 7 | 0.77 |
| P35442 | THBS2 | 7 | 6 | 0.77 |

| Accession | Name | # Peptides | # Unique Peptides | Abundance Ratio: WWP2 / control |
|---|---|---|---|---|
| Q6IQ23 | PLEKHA7 | 2 | 2 | 0.76 |
| Q9Y617 | PSAT1 | 14 | 14 | 0.76 |
| O95340 | PAPSS2 | 29 | 25 | 0.75 |
| P02786 | TFRC | 15 | 15 | 0.75 |
| P18827 | SDC1 | 2 | 2 | 0.75 |
| Q01650 | SLC7A5 | 2 | 1 | 0.75 |
| P36222 | CHI3L1 | 19 | 19 | 0.75 |
| P01009 | SERPINA1 | 7 | 7 | 0.75 |
| Q658P3 | STEAP3 | 7 | 7 | 0.75 |
| P62306 | SNRPF | 2 | 2 | 0.74 |
| Q15036 | SNX17 | 2 | 2 | 0.74 |
| P15259 | PGAM2 | 9 | 3 | 0.73 |
| Q15043 | SLC39A14 | 3 | 3 | 0.73 |
| P13645 | KRT10 | 7 | 6 | 0.73 |
| Q14118 | DAG1 | 3 | 3 | 0.68 |
| P51808 | DYNLT3 | 2 | 2 | 0.66 |
| P22914 | CRYGS | 11 | 11 | 0.62 |
| P14555 | PLA2G2A | 4 | 4 | 0.62 |
| P17302 | GJA1 | 8 | 8 | 0.54 |
| O95460 | MATN4 | 8 | 8 | 0.51 |
